# A TMT-based quantitative proteomics approach toward α-syn PFF associated Lewy Body Dementia (LBD) using α-syn PFF-injected mouse brain tissues

**DOI:** 10.1101/2022.10.10.511662

**Authors:** Fatih Akkentli, In kyu Jang, Yoonseop Choi, Dongyoon Yoo, Eric Joon-Hun Oh, Chan Hyun Na, Shinwon Ha, Sung-Ung Kang

## Abstract

The aggregation of α-synuclein in the nervous system leads to a class of neurodegenerative disorders termed α-synucleinopathies. A form of primary degenerative dementia called Lewy body dementia (LBD) often develops in the case these aggregations develop into intracellular inclusions called Lewy bodies (LB) and Lewy neurites (LN). Despite the high frequency of LBD, being the leading cause of dementia following Alzheimer’s disease (AD), there is relatively little information discovered about its pathological pathway or diagnostic criteria. In this report, we attempt to address such shortcomings via utilizing a proteomic approach to identify the proteomic changes following intrastriatal injection of α-synuclein preformed fibril (α-syn PFF). Through mass spectrometry, we have identified a total of 179 proteins that were either up- or down-regulated at different time points, with the four proteins – TPP3, RAB10, CAMK2A, and DYNLL1 – displaying the most significant changes throughout the timeframe. Further examining the modulated proteins with network-based enrichment analyses, we have found that 1) the most significantly associated neurodegenerative pathways were Parkinson’s (*p*V = 3.0e-16) and Huntington’s (*p*V = 1.9e-15) disease, and 2) the majority of molecular functions specific to the pathology only appeared at later time points. While these results do not expose a conclusive biomarker for LBD, they suggest a potential framework that may be utilized to diagnose and differentiate LBD pathology from other forms of dementia by focusing on the cortical proteomic changes which occur in a later time span.

## Introduction

⍺-synucleinopathies are a class of neurodegenerative disorders characterized by insoluble fibrillary aggregations of ⍺-synuclein inside the cytoplasm of neurons and glial cells (Fujiwara et al., 2002; Jellinger, 2003; Marti et al., 2003; Lee et al., 2008). In the case of neuronal inclusions, known as Lewy bodies (LB) and Lewy neurites (LN), they lead to a subclass of disorders called Lewy body diseases which include Dementia with Lewy bodies (DLB), Parkinson’s Disease (PD), and Parkinson’s Disease with Dementia (PDD) (Mueller et al., 2017; Surendranathan et al., 2020). Although the Lewy body dementias (LBD) - comprised of DLB and PDD are presumed to be the second most frequent case of primary degenerative dementia following Alzheimer’s Disease (AD), their exact clinical diagnosis and prevalence, especially for DLB, remain relatively unclear due to its concurrence or similarities with other types of dementia (Mueller et al., 2017; Kane et al., 2018; Matar et al., 2020). Thus, establishing clear clinical features and reliable diagnostic biomarkers for LBD have long been of interest in the field of neurology, with the international consensus criteria for DLB being revised four times since its foundation in 1996 (Yamada et al., 2020). Unlike the pathology of AD which involves the deposit of amyloid-β plaques in the cholinergic system and hippocampus, such areas are relatively spared in the case of DLB, in which Lewy bodies ascend to the cortical/subcortical pathways (Gibb et al., 1987; Gibb et al., 1989; Armstrong, 2020). Yet, due to its history of misdiagnoses and ambiguity, there is a relative deficit in data dedicated to identifying biomarkers or pathological conditions for the disease. Discovering progressive proteomic changes indicative of LBD will be critical in developing early diagnostic kits, a key component in ensuring the effectiveness of most treatments and medication. In this study, we utilize α-Synuclein performed fibrils (α-syn PFF) - injected mice models to analyze proteomic changes at different time points following injection, attempting to elucidate the pathway and effects of α-Synuclein aggregation pathology.

## Materials and methods

### Intrastriatal injection of α-syn PFF

Mice were anesthetized with isoflurane (1~2% inhalation), and small incision was made to expose the cranium. α-syn PFF is injected into the striatum (2 μL per hemisphere at 0.4 μL/min). The stereotaxic coordinates, relative to the bregma were anteroposterior (AP) = +0.2 mm, mediolateral (ML) = +2.0 mm, dorsoventral (DV) = +2.8 mm. After each injection was completed, the needle was maintained in position for 5 mins. Following needle removal, the wound was closed with silk suture and antibiotic ointment was applied to the surgical site. Animal was gently warmed on a 37°C heating pad for 3-5 minutes and returned to their original nesting material.

### Sample preparation and TMT labeling

The cerebral cortices for each time point were dissected and collected. The tissues were homogenized using a Dounce homogenizer with CHAPS lysis buffer (150 mM KCl, 50 mM HEPES pH = 7.4, 0.1% CHAPS, and 1 protease inhibitor cocktail tablet per 50 mL of buffer). Subsequently, samples were sonicated (30 sec on/off for 10 times at 4 ° C water bath) and centrifuged (16,000 x *g* on 4 ° C for 10 min). Protein concentrations of the supernatant were measured using the Bicinchoninic acid assay (Pierce, Waltham, MA, USA). Reducing of cysteine chains from equal amounts of proteins (0.5 mg per condition) were performed using 5 mM DTT at 55°C for 60 min and alkylated using 5 mM iodoacetamide for 45 min at room temperature (RT) in the dark. The samples were then trypsinized for 3 hours before Tandem mass tag (TMT) labeling. TMT labeling was carried out following as the manufacturer instructions (Dayon et al., 2008). Briefly, trypsinized peptides from four conditions were reconstituted in 50 mM TEABC buffer (2% SDS, 50 mM triethyl ammonium bicarbonate, 5mM sodium fluoride, 1mM sodium orthovanadate, and 1 mM β-glycerophosphate) and mixed with the TMT reagent and incubated at RT for 1 h. After the labeling, all samples were pooled and desalted using Sep-Pak C18 cartridges.

### Mass Spectrometry

LTQ-Orbitrap Elite mass spectrometer (Thermo Electron, Bremen, Germany) coupled with Easy-nLC II nanoflow LC system (Thermo Scientific, Odense, Denmark) was applied to identify sequence of peptides and quantification. Briefly, the pooled TMT-labeled peptides were reconstituted in 0.1% formic acid and loaded on an analytical column (75 µm x 50 cm) at a flow rate of 300 nL/min using a linear gradient of 10-35% solvent B (0.1% formic acid in 95% acetonitrile) over 120 min. Data-dependent acquisition with full scans in 350-1700 m/z range was carried out using an Orbitrap mass analyzer at a mass resolution of 120,000 at 400 m/z. The fifteen most intense precursor ions from a survey scan were selected for MS/MS fragmentation using higher-energy collisional dissociation (HCD) fragmentation with 32% normalized collision energy and detected at a mass resolution of 30,000 at 400 m/z. Automatic gain control for full MS was set to 1 × 10^6^ for MS and 5 × 10^4^ ions for MS/MS with a maximum ion injection time of 100 ms. Dynamic exclusion was set to 30 sec. and singly charged ions were rejected. Internal calibration was carried out using the lock mass option (m/z 445.1200025) in ambient air.

### Immunoblot Assay

Samples were separated by 8-16% SDS-PAGE and transferred to nitrocellulose membrane (0.45 μm). The membrane were incubated with PBS-T (0.05 % Tween 20, v/v) containing bovine serum albumin (5%, w/v) for blocking, and the membranes were applied with antibodies; polyclonal rabbit anti-TPPP3 (15057-1-AP, ProteinTech), polyclonal rabbit anti-RAB10 (11808-1-AP, ProteinTech), polyclonal rabbit anti-CaMKII-α (3357, Cell Signaling Technology), polyclonal rabbit anti-DYNLL1 (A53885, EpigenTek), and rabbit anti-beta-Actin HRP conjugate (13E5, Cell Signaling Technology). Horseradish peroxidase-conjugated secondary anti-rabbit or anti-mouse IgG (Amersham Bioscience) was used to detecte in X-ray film (AGFA) by an ECL method (Thermo Scientific). Densitometry was performed by Image J, and values were averaged from three-independent experiments.

### Data analysis

Cytoscape v3.9.1 software with the GlueGo v2.5.7 and BiNGO v3.0.3 plugins, coupled with reference networks such as KEGG, Gene Ontology (GO): Biological Process (BP) and Molecular Function (MF) resources were applied by according to the instruction manual (Maere et al., 2005; Bindea et al., 2009).

### Statistical analysis

Quantitative data were presented as mean ± SEM using GraphPad prism software with unpaired student’s t-test. Statistic approaches for functional analysis: ClueGO and BiNGO were assessed by the Two-sided hypergeometric test coupled with the Bonferroni step-down for correction and hypergeometric test coupled with Benjamini&Hochberg False Discovery Rate (FDR) correction respectively.

## Results

### Quantitative proteomic screening for α-syn PFF-injected mouse brain tissues

A high-throughput proteomic profiling was performed to identify large-scale proteome changes following intrastriatal injection of α-synuclein pre-formed fibril (α-syn PFF) in C57BL6 mice (Fig. 1A). After injecting α-syn PFF on 8 male mice per time point, 6 mice were sacrificed and grouped into Set_1 and Set_2 (n = 3 mice per set with biological replication) for proteomic profiling, and the remaining mice were used for immunostaining to check the propagation of α-syn PFF. α-syn PFF was stably injected, distributed to cerebral cortex, and aggregated in cerebral cortex at 2-months as evidenced by immunostaining using anti-phospho-S129 α-synuclein antibody (Fig. 1B). Two sets of quantitative proteomic screening using Tandem mass tag (TMT) technique were compared for reliability, whose results are illustrated in Fig. 1C. A regression analysis of sample ratios revealed a strong positive correlation between Set_1 and Set_2 (Fig. 1C). For Set_1 and Set_2, the number of quantified peptides were 4,523 and 4,387, with 3,457 shared (76.4% and 78.8%, respectively) between the two sets; the number of quantified protein groups were 796 and 797, with 660 shared (82.9% and 82.8%, respectively) (Fig. 1D, Table S1). Among the 660 shared protein groups, 37 were classified as up-regulated (AVR > 5%) and 142 as down-regulated (AVR > 20%) (Table 1 and S2).

**Table 1.**
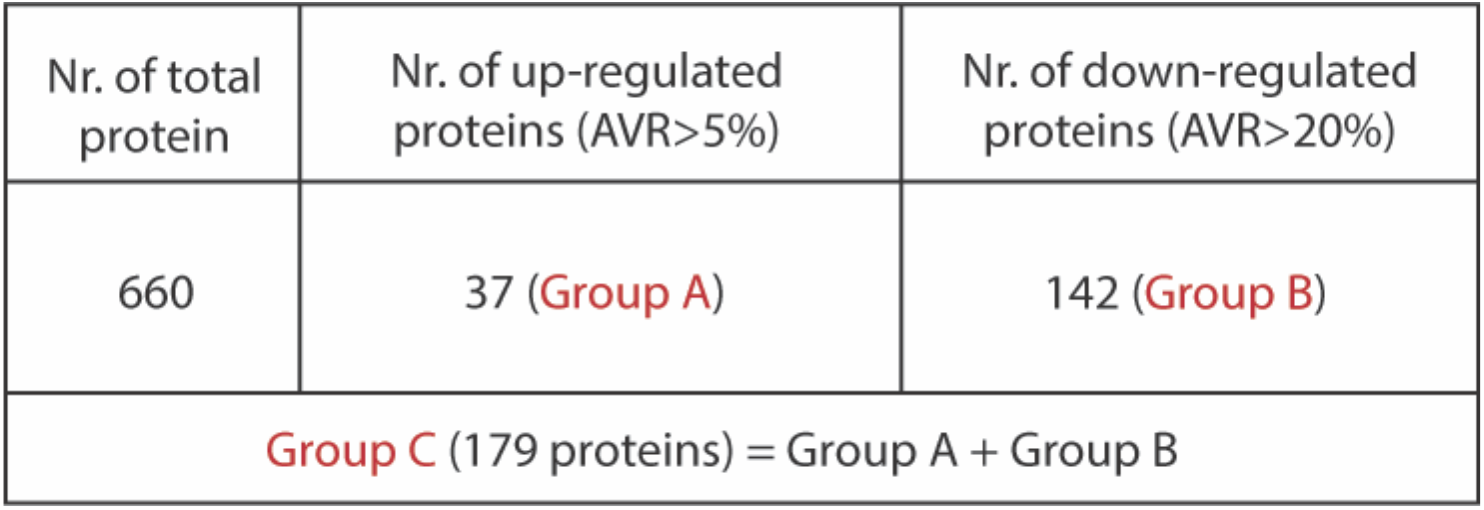
37 proteins (AVR>5% up-regulated; Group A) had significance among 186 up-regulated proteins, and 142 proteins (AVR>20% down-regulated; Group B) had significance among 473 down-regulated proteins. 179 proteins (Group C = Group A+B) were used for functional network analysis.

| Nr. of total protein | Nr. of up-regulated proteins (AVR>5%) | Nr. of down-regulated proteins (AVR>20%) |
| --- | --- | --- |
| 660 | 37 (Group A) | 142 (Group B) |
| Group C (179 proteins) = Group A + Group B |  |  |

**Figure 1.**
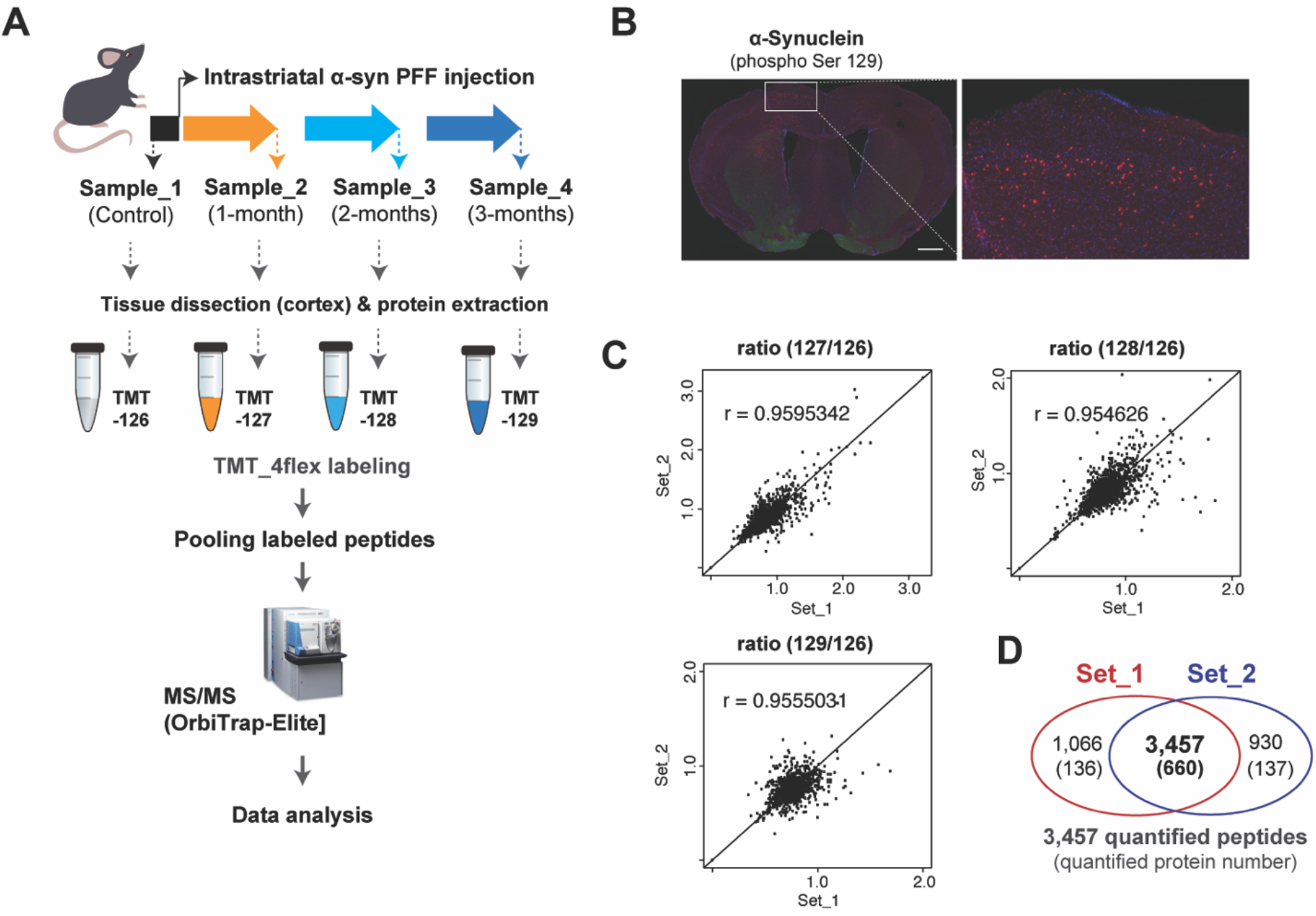
TMT-based protein quantification using α-syn PFF-injected mouse brain tissues. (A) Experimental illustration for the procedures of quantitative proteomics approaches including intrastriatal injection of α-syn PFF, sampling, TMT-labeling, and mass spectrometry is shown. (B) Representative immuno-stained images of coronal sections after 2 months of α-syn PFF injection using phospho-S129 α-synuclein antibody (red), indicating biodistribution and aggregation of α-syn PFF. Scale bars: 1 mm. (C) Scatter plots showing the reproducibility of transcriptomic data between biological replications of Set_1 and Set_2. (D) The Venn diagram indicates the overlapped number of peptides and proteins between two data of our biological replications obtained by TMT-based protein quantification.

### Functional alteration following intrastriatal α-syn PFF injection

A comparative analysis of average fold changes relative to the control revealed changes in gene expression over time by Proteome Discoverer v1.4, with a range of up- and down-regulated proteins after 1-, 2-, and 3-month following α-syn PFF injection (Fig. 2A and Table S2). The four most prominent alterations presented in all three time points included the upregulation of Tubulin polymerization promoting protein family member 3 (TPPP3) and Ras-related protein Rab-10 (RAB10), and the downregulation of Calcium/calmodulin dependent protein kinase II alpha (CAMK2A) and Dynein light chain 1 (DYNLL1). All changes were statistically significant with the exception of TPPP3 at 2 and 3 months (Fig. 2B and C).

**Figure 2.**
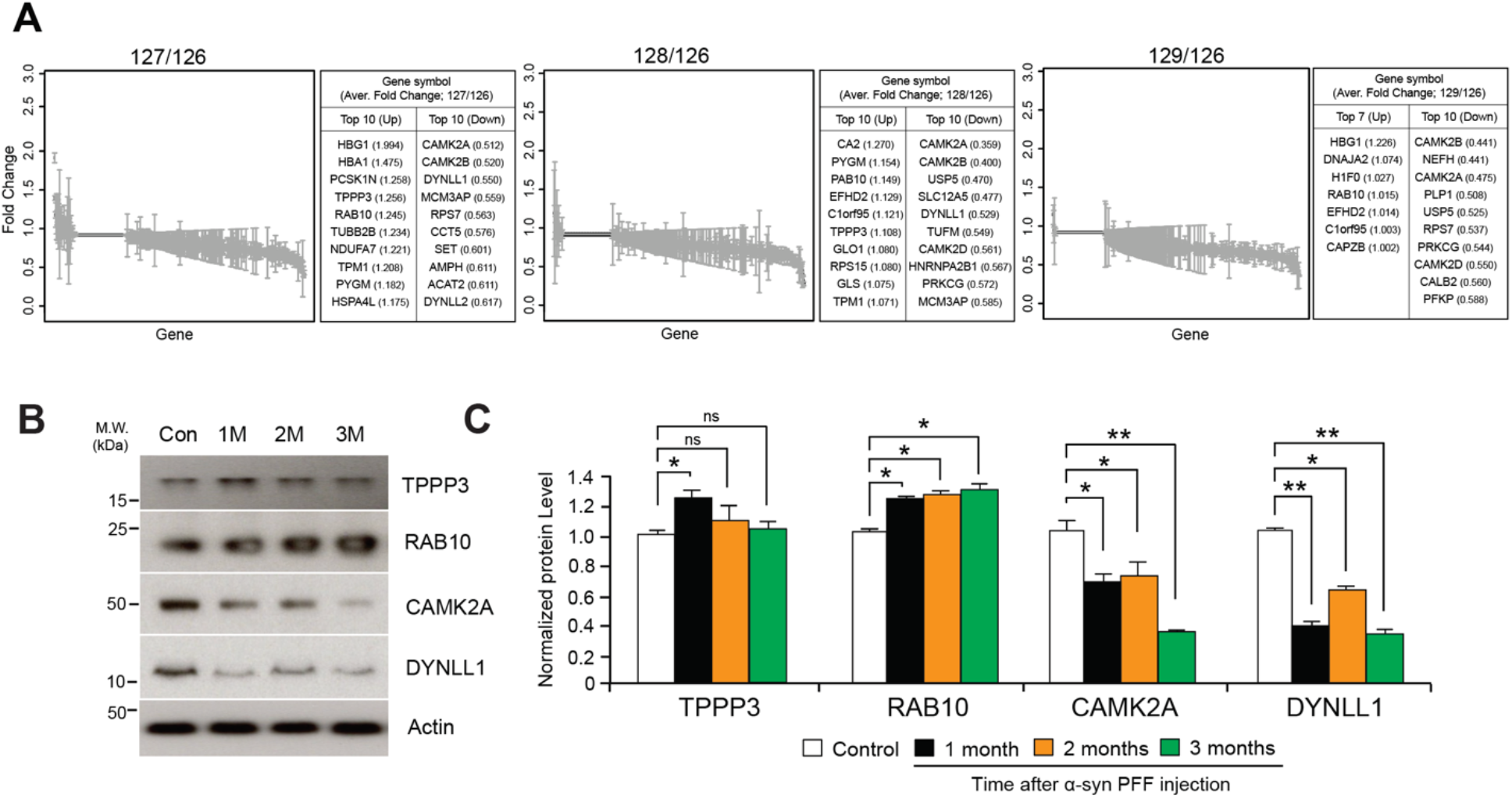
Protein quantification using TMT-labeled peptides. (A) Fold changes of 660 commonly identified proteins after normalization using peptides with expression of mass reporter (126 Da). (B) Western blot validation of significantly up- and down-regulated proteins including Tubulin polymerization-promoting protein family member 3 (TPPP3), Ras-related protein Rab-10 (RAB10), Calcium/calmodulin-dependent protein kinase type II subunit alpha (CAMK2A), Dynein light chain 1 (DYNLL1). (C) Quantification graphs of expression ratio obtained from triplicate samples and normalized using expression of β-Actin; two-tailed unpaired Student’s t-test.

### Network-based KEGG and GO: Biological Process (BP) Analysis

The network analysis of significantly up- and down-regulated proteins identified several enriched Kyoto Encyclopedia of Genes and Genomes (KEGG) pathways and GO biological processes (BP), designated by a minimum association of 6 or 8% of genes and *p*-value < 1.0e-5 (Fig. 3A and B). Total 17 representative pathways spanning over a wide range of networks for metabolism, environmental information processing, cellular processes, organismal systems, and human diseases were discovered in KEGG pathways (Fig. 1A, Table S3). The pathways with the greatest significance were neurodegenerative disorders including Parkinson’s (*p*V = 3.0e-16) and Huntington’s (*p*V = 1.9e-15) disease, endocrine and other factor-regulated calcium reabsorption (*p*V = 8.8e-14), and synaptic vesicle cycle (*p*V = 5.6e-13) (Table S3). GO analysis elucidated five networks displaying high significance, including vesicle-mediated transport (*p*V = 1.1e-13) and substantia nigra development (*p*V = 1.1e-10). The relative *p*-values and percentage per number of associated genes for all KEGG pathways and GO: BP are depicted in Fig. 3B.

**Figure 3.**
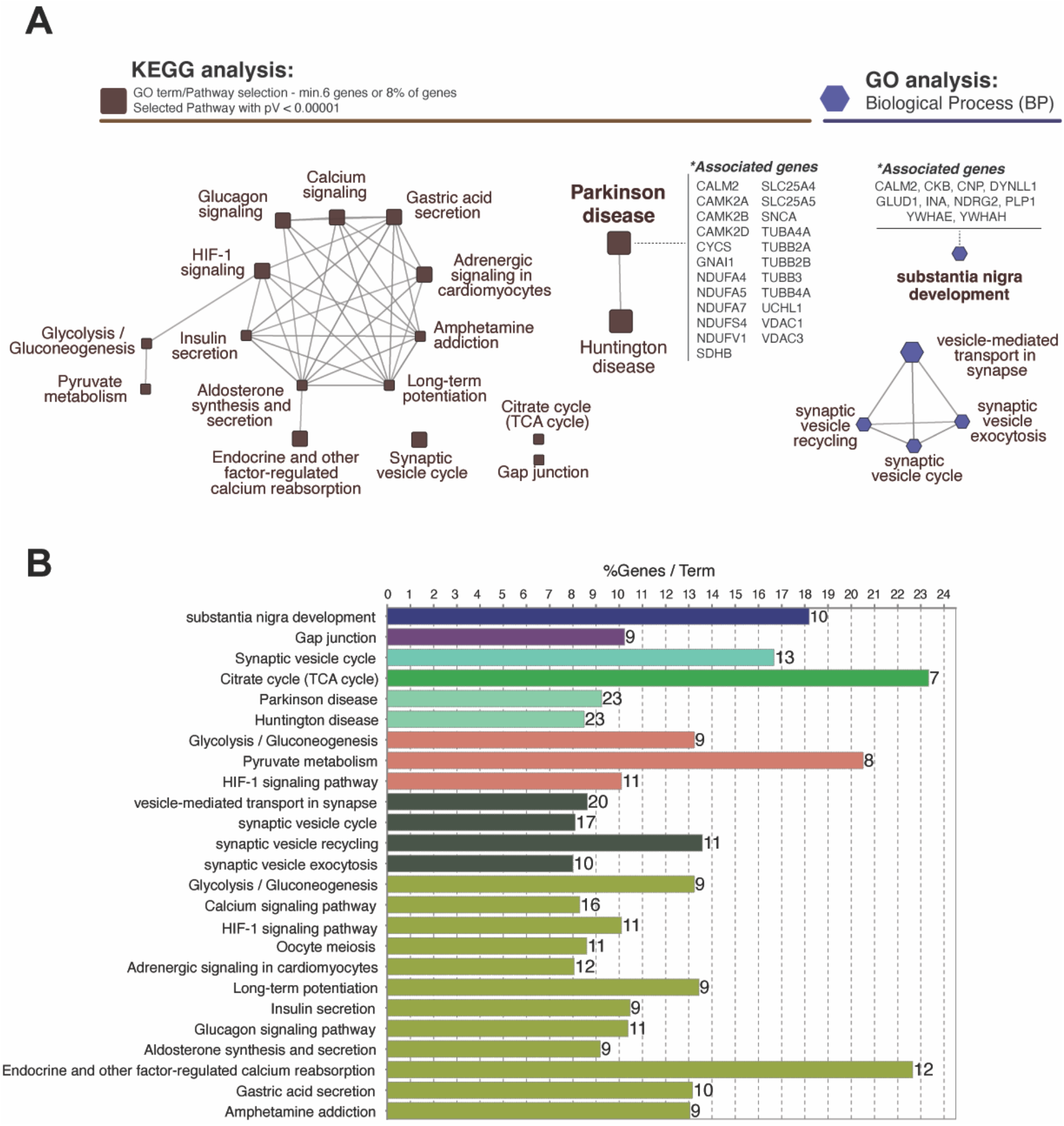
Analysis of functional networks using ClueGo Cytoscape plugin. (A) Network-based KEGG and GO analysis represent proteins classified into specific pathways which are labeled beside the nodes. Total 17 representative pathways with a high significance (*p*V<0.00001) including Parkinson’s (23 genes; 9.24%) and Huntington diseases (23 genes; 8.49%) were identified in KEGG pathways. GO:Biological Process analysis identified 5 enriched networks including substantia nigra development (10 genes; 18.18%). (B) Bar graph represents the number of genes, terms, and %geneset in each pathway shown in (A).

### Time-dependent Network-based GO: Molecular Function (MF) Analysis

Additionally, a network-based GO: molecular function (MF) analysis was performed to investigate the molecular progression of α-Synuclein aggregates in significantly up- and down-regulated proteins for the three time points of interest. We found that the number of significantly (*p*V < 1.0e-5) enriched GO: MF clusters increased over time, with 37 unique clusters including ATPase (*p*V = 8.9e-11) and Catalytic (*p*V = 8.1e-9) activity only appearing at 3 months after α-syn PFF injection. Nevertheless, the most significant and frequent clusters appeared to be those common across multiple time points, such as nucleotide (pV = 1.2e-20 at 3-month) and protein (*p*V = 1.3e-17) binding present at 1-, 2-, and 3-month, and GTPase activity (*p*V = 4.3e-15) present at 2- and 3-month. Information regarding *p*-values, cluster frequency, and associated genes for all significantly enriched functions are provided in Fig. 4 and Table S4.

**Figure 4.**
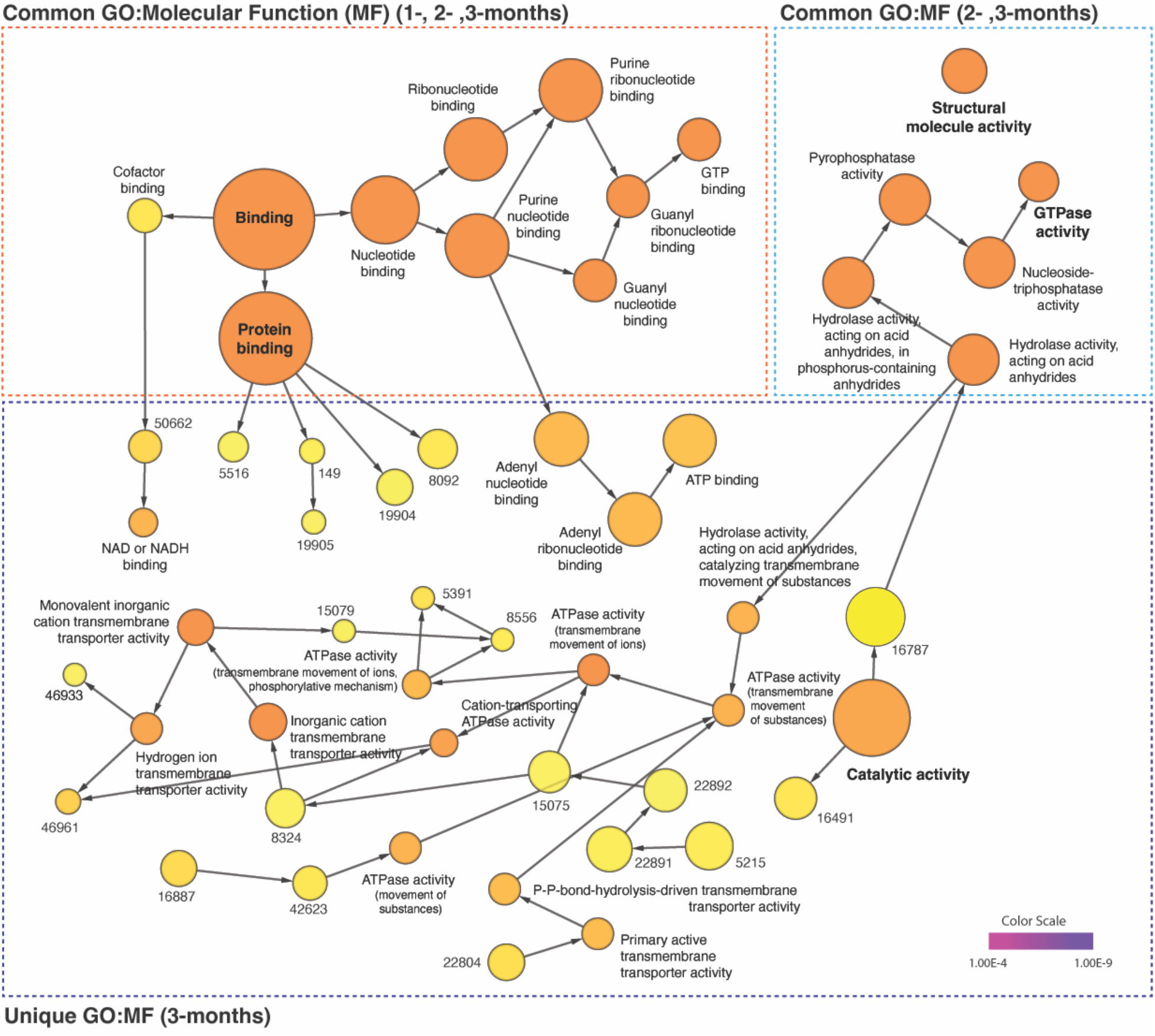
Distinct time-dependent GO profiles and network dynamics in post-α-syn PFF-injection. The gene ontology (GO) enrichment analysis of up-(AVR>5%) and down-regulated (AVR>20%) proteins in each time points were performed on the network using BiNGO plugin of Cytoscape (v.3.9.1). The common and unique clusters of molecular function (MF) were shown. The color gradient of the cluster distribution stands for the *p*-value of each cluster associated with the term. Darker (orange) color indicates a lower *p*-value. Numbers on yellow color (relevantly higher *p*-value) indicate GO-ID number, which provided in Table S4.

## Discussion

In this study, we have attempted to elucidate the pathological pathways and mechanisms associated with α-synuclein propagation, often leading to Lewy body dementias (LBD) in human patients, using a proteomic approach in the mice brain. Previous literature has discovered that while Lewy bodies originate in the substantia nigra, their ascension to the cortex is the pathway lying behind Parkinson’s disease and Dementia with Lewy bodies (Obeso et al., 2010). Consistent with this theory, we have observed α-synuclein aggregations in the mice cortex through coronal sections, supporting the clinical relevance of the study for human models. Investigating the range of varying gene expression levels through mass spectrometry analysis, we have identified several genes which were commonly altered at all time points after α-syn PFF injection, including TPPP3 and RAB10, which were upregulated, and CAMK2A and DYNLL1, which were downregulated. These results agree with prior reports on neurodegenerative diseases, where all four genes displaying association with not only Parkinson’s (Simunovic et al., 2009; Zhang et al., 2014; Olah et al., 2020; Fellgett et al., 2021), but also with Alzheimer’s (Kong et al., 2009; Ghosh and Giese, 2015; Ridge et al., 2017; Singh et al., 2022) disease. Thus, while we may conclude that these genes may serve as potential biomarkers for early detection of general neurodegeneration, it appears difficult to assign them to a particular diagnosis.

Yet, our network-based analysis results suggest a method that may help differentiate specific pathologies from a range of similar disorders, in accordance with the difference in associated genes found and more detailed pathways/molecular functions (MF) being enriched at different time points. For instance, the same number of genes with modified expression were associated with the enrichment of both Parkinson’s and Huntington’s disease pathways via KEGG analysis, with the majority of genes shared between the two. However, the deciding factor between these two similar motor disorders was that the unshared genes for Parkinson’s disease were related to calcium-dependent signaling (CAMK2-α, β, δ), while those for Huntington’s disease were related to adaptor protein complexes (AP2-α1, β1, Mu1). Additionally, our GO:MF analysis revealed that calmodulin binding was only enriched 2 and 3 months after α-syn PFF injection. We believe this implies that while the two diseases may share common signatures early on during their progression, the detailed pathways appearing over time, in this case the calcium/calmodulin-dependent pathways, may serve as a diagnosis criterion in distinguishing a specific condition. This idea is further strengthened by the fact that while a relatively small number (11 modules) of general, significant functions emerged at 1-month, a large number (53 modules) of uncommon, specific functions emerged at 3-month.

Being one of the first of its kind, our study illustrates the possibility of utilizing a proteomic approach in identifying Lewy-body related neurodegeneration. However, it is also not without its limitations. First, while the human neurodegenerative dementias develop gradually over a prolonged timeframe of years, our analysis studied differences in gene expression over 3 months. Thus, the definitive signatures in human models may differ significantly from those in our report. Additionally, due to the lack of similar studies, the verification of the potential biomarkers we identified remains a challenge.

## Acknowledgements

This work was supported by the National Institutes of Health (NIH)/National Institute of Neurological Disorders and Stroke (NINDS) R01 NS123456.

## Conflicts of interest

The authors declare that they have no conflict of interest.

## Author contributions

F.A., I.J., Y.C., C.H.N., S.H., and S.U.K. designed the study and wrote the paper. F.A. performed intrastriatal injection of α-syn PFF and sample collection. C.H.N., S.H., and S.U.K. performed enzyme digestion, TMT-labeling, peptide preparation and mass spectrometry. F.A., I.J., Y.C., D.Y., E.J.H.O, C.H.N., S.H., and S.U.K. analyzed the data.

## Supporting information

**Table S1**. List of identified genes with TMT quantification

**Table S2**. Comparative analysis of overlapped 660 genes over three different time points

**Table S3**. Network-based KEGG and GO analysis

**Table S4**. List of distinct time-dependent GO profiles in post-a-syn PFF-injection

## Notes

### Competing Interest Statement

The authors have declared no competing interest.

