## Supplemental table 1 for "A TMT-based quantitative proteomics approach toward α-syn PFF associated Lewy Body Dementia (LBD) using α-syn PFF-injected mouse brain tissues"

| Accession | Gene symbol | Protein description | Refseq protein accession | Exp. q-value | Sum PEP Score |
|---|---|---|---|---|---|
| 4501867 | ACO2 | aconitate hydratase, r | NP_001089.1 | 0 | 58.6020271 |
| 4501881 | ACTA1 | actin, alpha skeletal m | NP_001091.1 | 0 | 101.071356 |
| 4501913 | ADAM23 | disintegrin and metallo | NP_003803.1 | 0 | 3.33292079 |
| 4502027 | ALB | serum albumin preprop | NP_000468.1 | 0.00269139 | 1.93554201 |
| 4502049 | AKR1B1 | aldose reductase [Hor | NP_001619.1 | 0.01631637 | 1.28743447 |
| 4502107 | ANXA5 | annexin A5 [Homo sap | NP_001145.1 | 0 | 5.33703492 |
| 4502201 | ARF1 | ADP-ribosylation facto | NP_001649.1 | 0 | 8.25740886 |
| 4502209 | ARF5 | ADP-ribosylation facto | NP_001653.1 | 0.00030826 | 2.35981681 |
| 4502227 | ARL1 | ADP-ribosylation facto | NP_001168.1 | 0 | 6.86518563 |
| 4502271 | ATP1A2 | sodium/potassium-trar | NP_000693.1 | 0 | 138.630545 |
| 4502277 | ATP1B1 | sodium/potassium-trar | NP_001668.1 | 0 | 35.2885103 |
| 4502303 | ATP5O | ATP synthase subunit | NP_001688.1 | 0 | 13.7897103 |
| 4502315 | ATP6V1C1 | V-type proton ATPase | NP_001686.1 | 0 | 13.8015924 |
| 4502317 | ATP6V1E1 | V-type proton ATPase | NP_001687.1 | 0 | 55.5790016 |
| 4502491 | C1QBP | complement componer | NP_001203.1 | 0 | 4.74569367 |
| 4502673 | CD47 | leukocyte surface anti | NP_001768.1 | 0 | 9.58142164 |
| 4502993 | COX7C | cytochrome c oxidase | NP_001858.1 | 0.0065658 | 1.50584541 |
| 4503009 | CPE | carboxypeptidase E pi | NP_001864.1 | 0 | 7.40540726 |
| 4503065 | CRYM | ketimine reductase mu | NP_001879.1 | 0.01226652 | 1.36291073 |
| 4503095 | CSNK2A1 | casein kinase II subur | NP_001886.1 | 0 | 15.5556482 |
| 4503379 | DPYSL3 | dihydropyrimidinase-re | NP_001378.1 | 0 | 53.0674973 |
| 4503471 | EEF1A1 | elongation factor 1-alp | NP_001393.1 | 0 | 32.3872905 |
| 4503475 | EEF1A2 | elongation factor 1-alp | NP_001949.1 | 0 | 27.7375983 |
| 4503481 | EEF1G | elongation factor 1-gai | NP_001395.1 | 0 | 11.5351208 |
| 4503483 | EEF2 | elongation factor 2 [H | NP_001952.1 | 0 | 12.9573597 |
| 4503571 | ENO1 | alpha-enolase isoform | NP_001419.1 | 0 | 60.2652538 |
| 4503607 | ETFA | electron transfer flavo | NP_000117.1 | 0 | 10.821088 |
| 4503971 | GDI1 | rab GDP dissociation i | NP_001484.1 | 0 | 70.587378 |
| 4504041 | GNAI2 | guanine nucleotide-bin | NP_002061.1 | 0 | 20.4911774 |
| 4504067 | GOT1 | aspartate aminotransf | NP_002070.1 | 0 | 24.2939918 |
| 4504251 | HIST2H2AA3 | histone H2A type 2-A | NP_003507.1 | 0 | 22.4873918 |
| 4504317 | HIST1H4L | histone H4 [Homo sap | NP_003537.1 | 0 | 25.1126626 |
| 4504347 | HBA1 | hemoglobin subunit al | NP_000549.1 | 0 | 14.2517872 |
| 4504511 | DNAJA1 | dnaJ homolog subfami | NP_001530.1 | 0 | 7.08622818 |
| 4504523 | HSPE1 | 10 kDa heat shock pro | NP_002148.1 | 0 | 15.7892183 |
| 4505029 | LTA4H | leukotriene A-4 hydrol | NP_000886.1 | 0.00406268 | 1.73969005 |
| 4505185 | MIF | macrophage migration | NP_002406.1 | 0 | 3.52549236 |
| 4505357 | NDUFA4 | cytochrome c oxidase | NP_002480.1 | 0 | 4.02553436 |
| 4505369 | NDUFS4 | NADH dehydrogenase | NP_002486.1 | 0 | 4.00638744 |
| 4505451 | NRAS | GTPase NRas [Homo s | NP_002515.1 | 0 | 23.0609189 |
| 4505505 | OPCML | opioid-binding protein/ | NP_002536.1 | 0 | 5.62857803 |
| 4505531 | OSBP | oxysterol-binding prot | NP_002547.1 | 0 | 4.39707229 |
| 4505585 | PAFAH1B2 | platelet-activating fact | NP_002563.1 | 0 | 2.65896137 |
| 4505621 | PEBP1 | phosphatidylethanolan | NP_002558.1 | 0 | 7.48886862 |
| 4505657 | PDE2A | cGMP-dependent 3',5' | NP_002590.1 | 0 | 7.51723615 |
| 4505735 | PF4V1 | platelet factor 4 variar | NP_002611.1 | 0.0188216 | 1.19873353 |
| 4505753 | PGAM1 | phosphoglycerate mut | NP_002620.1 | 0 | 79.0178887 |
| 4505763 | PGK1 | phosphoglycerate kina | NP_000282.1 | 0 | 44.3832681 |
| 4506013 | PPP1R7 | protein phosphatase 1 | NP_002703.1 | 0 | 10.6546263 |
| 4506017 | PPP2CA | serine/threonine-protei | NP_002706.1 | 0 | 4.37861597 |
| 4506025 | PPP3R1 | calcineurin subunit B t | NP_000936.1 | 0 | 6.83283381 |
| 4506055 | PRKACA | cAMP-dependent prote | NP_002721.1 | 0 | 8.22049266 |
| 4506181 | PSMA2 | proteasome subunit al | NP_002778.1 | 0 | 3.16966801 |
| 4506183 | PSMA3 | proteasome subunit al | NP_002779.1 | 0.00030826 | 2.12726117 |
| 4506193 | PSMB1 | proteasome subunit be | NP_002784.1 | 0.00181214 | 2.07052998 |
| 4506209 | PSMC2 | 26S protease regulato | NP_002794.1 | 0 | 15.6604939 |
| 4506365 | RAB2A | ras-related protein Rab | NP_002856.1 | 0 | 25.5046932 |

| | | | | | |
|---|---|---|---|---|---|
| 4506367 | RAB3A | ras-related protein Ral | NP_002857.1 | 0 | 75.5265923 |
| 4506413 | RAP1A | ras-related protein Ra | NP_002875.1 | 0 | 17.4617889 |
| 4506597 | RPL12 | 60S ribosomal protein | NP_000967.1 | 0.01712234 | 1.27237742 |
| 4506605 | RPL23 | 60S ribosomal protein | NP_000969.1 | 0.00030826 | 2.31264943 |
| 4506625 | RPL27A | 60S ribosomal protein | NP_000981.1 | 0.00030826 | 2.16411927 |
| 4506631 | RPL30 | 60S ribosomal protein | NP_000980.1 | 0 | 3.35912122 |
| 4506645 | RPL38 | 60S ribosomal protein | NP_000990.1 | 0 | 6.39008239 |
| 4506649 | RPL3 | 60S ribosomal protein | NP_000958.1 | 0.00030826 | 2.19907682 |
| 4506661 | RPL7A | 60S ribosomal protein | NP_000963.1 | 0 | 8.7609197 |
| 4506663 | RPL8 | 60S ribosomal protein | NP_000964.1 | 0 | 4.37582107 |
| 4506671 | RPLP2 | 60S acidic ribosomal p | NP_000995.1 | 0 | 9.17615502 |
| 4506675 | RPN1 | dolichyl-diphosphoolig | NP_002941.1 | 0.00487945 | 1.59074335 |
| 4506681 | RPS11 | 40S ribosomal protein | NP_001006.1 | 0.00030826 | 2.19791074 |
| 4506685 | RPS13 | 40S ribosomal protein | NP_001008.1 | 0 | 8.49703002 |
| 4506691 | RPS16 | 40S ribosomal protein | NP_001011.1 | 0 | 5.33171268 |
| 4506695 | RPS19 | 40S ribosomal protein | NP_001013.1 | 0 | 7.17186353 |
| 4506715 | RPS28 | 40S ribosomal protein | NP_001022.1 | 0 | 5.46584693 |
| 4506723 | RPS3A | 40S ribosomal protein | NP_000997.1 | 0 | 3.94493646 |
| 4506725 | RPS4X | 40S ribosomal protein | NP_000998.1 | 0 | 4.57643057 |
| 4506741 | RPS7 | 40S ribosomal protein | NP_001002.1 | 0 | 11.8825929 |
| 4506743 | RPS8 | 40S ribosomal protein | NP_001003.1 | 0 | 10.2223663 |
| 4507115 | FSCN1 | fascin [Homo sapiens] | NP_003079.1 | 0 | 8.31043711 |
| 4507297 | STXBP1 | syntaxin-binding prote | NP_003156.1 | 0 | 83.3268248 |
| 4507677 | HSP90B1 | endoplasmin precursoı | NP_003290.1 | 0 | 23.6185484 |
| 4507729 | TUBB2A | tubulin beta-2A chain i | NP_001060.1 | 0 | 253.475156 |
| 4507791 | UBE2M | NEDD8-conjugating en | NP_003960.1 | 0 | 3.79615154 |
| 4507793 | UBE2N | ubiquitin-conjugating e | NP_003339.1 | 0 | 18.2968873 |
| 4507879 | VDAC1 | voltage-dependent ani | NP_003365.1 | 0 | 50.1360755 |
| 4507947 | YARS | tyrosine--tRNA ligase, | NP_003671.1 | 0 | 3.45308736 |
| 4507949 | YWHAB | 14-3-3 protein beta/alp | NP_003395.1 | 0 | 39.3802212 |
| 4507951 | YWHAH | 14-3-3 protein eta [Hoı | NP_003396.1 | 0 | 39.4913538 |
| 4557237 | ACAT1 | acetyl-CoA acetyltrans | NP_000010.1 | 0 | 6.27711528 |
| 4557367 | BLMH | bleomycin hydrolase [I | NP_000377.1 | 0 | 4.43332686 |
| 4557395 | CA2 | carbonic anhydrase 2 | NP_000058.1 | 0 | 15.0092573 |
| 4557581 | FABP5 | fatty acid-binding prot | NP_001435.1 | 0.00350365 | 1.82506841 |
| 4557707 | L1CAM | neural cell adhesion m | NP_000416.1 | 0 | 8.15776356 |
| 4757714 | ACP1 | low molecular weight p | NP_004291.1 | 0 | 4.2651134 |
| 4757732 | AIFM1 | apoptosis-inducing fac | NP_004199.1 | 0 | 3.72815839 |
| 4757774 | ARL3 | ADP-ribosylation facto | NP_004302.1 | 0 | 2.84832377 |
| 4757818 | ATP6V1G1 | V-type proton ATPase | NP_004879.1 | 0 | 3.74617756 |
| 4757900 | CALR | calreticulin precursor [ | NP_004334.1 | 0 | 4.15181088 |
| 4758086 | CSRP1 | cysteine and glycine-r | NP_004069.1 | 0 | 9.41199265 |
| 4758152 | TIMM8A | mitochondrial import in | NP_004076.1 | 0.00406268 | 1.68465952 |
| 4758328 | FABP3 | fatty acid-binding prot | NP_004093.1 | 0 | 2.72193267 |
| 4758442 | GMFB | glia maturation factor I | NP_004115.1 | 0.00708617 | 1.43687504 |
| 4758516 | HDGF | hepatoma-derived grov | NP_004485.1 | 0 | 4.16858087 |
| 4758582 | IDH3G | isocitrate dehydrogena | NP_004126.1 | 0.00434279 | 1.62397082 |
| 4758638 | PRDX6 | peroxiredoxin-6 [Homo | NP_004896.1 | 0 | 10.584857 |
| 4758650 | KIF5C | kinesin heavy chain is | NP_004513.1 | 0 | 24.0195323 |
| 4758786 | NDUFS2 | NADH dehydrogenase | NP_004541.1 | 0 | 3.6567504 |
| 4758788 | NDUFS3 | NADH dehydrogenase | NP_004542.1 | 0 | 16.0169999 |
| 4758950 | PPIB | peptidyl-prolyl cis-tran | NP_000933.1 | 0 | 8.22796963 |
| 4758988 | RAB1A | ras-related protein Ral | NP_004152.1 | 0 | 21.451008 |
| 4759160 | SNRPD3 | small nuclear ribonucl | NP_004166.1 | 0 | 2.92299567 |
| 4759182 | STX1A | syntaxin-1A isoform 1 | NP_004594.1 | 0 | 38.1854218 |
| 4759198 | HOMER1 | homer protein homolog | NP_004263.1 | 0 | 5.92995445 |
| 4759302 | VAPB | vesicle-associated me | NP_004729.1 | 0 | 5.91257354 |
| 4759306 | LIN7A | protein lin-7 homolog A | NP_004655.1 | 0.00030826 | 2.2974833 |
| 4826655 | CALB1 | calbindin [Homo sapieı | NP_004920.1 | 0 | 17.3748446 |
| 4826675 | CDK5 | cyclin-dependent-like I | NP_004926.1 | 0.00461494 | 1.6064248 |
| 4826734 | FUS | RNA-binding protein Fl | NP_004951.1 | 0 | 3.38002299 |
| 4826816 | LGI1 | leucine-rich glioma-ina | NP_005088.1 | 0 | 8.33682034 |

| | | | | | |
|---|---|---|---|---|---|
| 4826854 | NDUFB8 | NADH dehydrogenase | NP_004995.1 | 0 | 3.50473326 |
| 4826898 | PFN1 | profilin-1 [Homo sapier | NP_005013.1 | 0 | 6.15610849 |
| 4826932 | PPID | peptidyl-prolyl cis-tran | NP_005029.1 | 0 | 8.40144147 |
| 4885281 | GLUD1 | glutamate dehydrogen | NP_005262.1 | 0 | 52.7506578 |
| 4885371 | H1F0 | histone H1.0 [Homo sa | NP_005309.1 | 0 | 2.66548079 |
| 4885377 | HIST1H1D | histone H1.3 [Homo sa | NP_005311.1 | 0 | 24.004603 |
| 4885563 | PRKCE | protein kinase C epsilo | NP_005391.1 | 0 | 3.1875546 |
| 5031569 | ACTR1A | alpha-centractin [Hom | NP_005727.1 | 0 | 25.8719163 |
| 5031571 | ACTR2 | actin-related protein 2 | NP_005713.1 | 0 | 15.041713 |
| 5031573 | ACTR3 | actin-related protein 3 | NP_005712.1 | 0 | 19.7162014 |
| 5031599 | ARPC2 | actin-related protein 2/ | NP_005722.1 | 0 | 5.9800107 |
| 5031635 | CFL1 | cofilin-1 [Homo sapien | NP_005498.1 | 0 | 18.9369437 |
| 5031711 | EIF1B | eukaryotic translation | NP_005866.1 | 0.00708617 | 1.43297363 |
| 5031741 | DNAJA2 | dnaJ homolog subfami | NP_005871.1 | 0 | 6.37222462 |
| 5031777 | IDH3A | isocitrate dehydrogena | NP_005521.1 | 0 | 15.2855231 |
| 5031985 | NUTF2 | nuclear transport facto | NP_005787.1 | 0 | 3.44285386 |
| 5032007 | PURA | transcriptional activato | NP_005850.1 | 0 | 23.2922209 |
| 5032009 | PYGM | glycogen phosphoryla: | NP_005600.1 | 0 | 18.8683685 |
| 5032139 | SYT1 | synaptotagmin-1 isofo | NP_005630.1 | 0 | 80.5897595 |
| 5174391 | AKR1A1 | alcohol dehydrogenas | NP_006057.1 | 0 | 5.50028955 |
| 5174447 | GNB2L1 | guanine nucleotide-bin | NP_006089.1 | 0 | 7.61522257 |
| 5174529 | MAT2A | S-adenosylmethionine | NP_005902.1 | 0 | 3.33059013 |
| 5174735 | TUBB4B | tubulin beta-4B chain | | NP_006079.1 | 0 | 245.210527 |
| 5453549 | PRDX4 | peroxiredoxin-4 precur | NP_006397.1 | 0 | 9.99772396 |
| 5453555 | RAN | GTP-binding nuclear p | NP_006316.1 | 0 | 18.5592539 |
| 5453559 | ATP5H | ATP synthase subunit | NP_006347.1 | 0.00350365 | 1.81559251 |
| 5453593 | CAP2 | adenylyl cyclase-asso | NP_006357.1 | 0 | 11.4892751 |
| 5453599 | CAPZA2 | F-actin-capping proteir | NP_006127.1 | 0 | 8.13410459 |
| 5453603 | CCT2 | T-complex protein 1 su | NP_006422.1 | 0 | 20.5123188 |
| 5453607 | CCT7 | T-complex protein 1 su | NP_006420.1 | 0 | 9.83550815 |
| 5453629 | DCTN2 | dynactin subunit 2 iso | NP_006391.1 | 0 | 6.75444559 |
| 5453710 | LASP1 | LIM and SH3 domain p | NP_006139.1 | 0 | 10.9424806 |
| 5453958 | PPP5C | serine/threonine-protei | NP_006238.1 | 0.00708617 | 1.46205504 |
| 5579478 | MAP2K1 | dual specificity mitoge | NP_002746.1 | 0 | 12.821004 |
| 5729781 | CPLX1 | complexin-1 [Homo sa | NP_006642.1 | 0 | 6.88498161 |
| 5729875 | PGRMC1 | membrane-associated | NP_006658.1 | 0 | 4.68047755 |
| 5729937 | MTX2 | metaxin-2 [Homo sapie | NP_006545.1 | 0 | 4.0051991 |
| 5802966 | DSTN | destrin isoform a [Hom | NP_006861.1 | 0 | 7.16441418 |
| 5803011 | ENO2 | gamma-enolase [Homo | NP_001966.1 | 0 | 85.4092193 |
| 5803013 | ERP29 | endoplasmic reticulum | NP_006808.1 | 0 | 6.92497169 |
| 5803187 | TALDO1 | transaldolase [Homo s | NP_006746.1 | 0 | 7.33573368 |
| 5803225 | YWHAE | 14-3-3 protein epsilon | NP_006752.1 | 0 | 40.9469585 |
| 5803227 | YWHAQ | 14-3-3 protein theta [H | NP_006817.1 | 0 | 37.3202929 |
| 5902018 | TPPP | tubulin polymerization- | NP_008961.1 | 0 | 29.5091198 |
| 5902076 | SRSF1 | serine/arginine-rich sp | NP_008855.1 | 0 | 4.51930061 |
| 5902102 | SNRPD1 | small nuclear ribonucle | NP_008869.1 | 0 | 5.26265479 |
| 6005717 | ATP5I | ATP synthase subunit | NP_009031.1 | 0 | 3.05754547 |
| 6005942 | VCP | transitional endoplasm | NP_009057.1 | 0 | 77.6904428 |
| 6005993 | CLTA | clathrin light chain A is | NP_009027.1 | 0 | 14.3508478 |
| 6005995 | CLTB | clathrin light chain B is | NP_009028.1 | 0 | 4.18973952 |
| 6031192 | SLC25A3 | phosphate carrier prot | NP_005879.1 | 0 | 10.1663654 |
| 6598323 | GDI2 | rab GDP dissociation i | NP_001485.2 | 0 | 47.5376958 |
| 6715568 | PPP3CA | serine/threonine-protei | NP_000935.1 | 0 | 53.5076276 |
| 6912238 | PRDX5 | peroxiredoxin-5, mitoc | NP_036226.1 | 0 | 6.44170577 |
| 6912280 | AHSA1 | activator of 90 kDa he | NP_036243.1 | 0 | 3.95624487 |
| 6912328 | DDAH1 | N(G),N(G)-dimethylarg | NP_036269.1 | 0 | 10.963064 |
| 6912482 | LETM1 | LETM1 and EF-hand d | NP_036450.1 | 0 | 17.3287223 |
| 6912634 | RPL13A | 60S ribosomal protein | NP_036555.1 | 0 | 4.86742015 |
| 6912646 | NPTN | neuroplastin isoform b | NP_036560.1 | 0 | 5.34625797 |
| 7019485 | PDCD6 | programmed cell death | NP_037364.1 | 0.00487945 | 1.58037464 |
| 7019519 | PCSK1N | proSAAS preproproteir | NP_037403.1 | 0 | 4.46715444 |
| 7549809 | PLS3 | plastin-3 isoform 1 [Hc | NP_005023.2 | 0 | 6.363743 |

| | | | | | |
|---|---|---|---|---|---|
| 7657015 | RTCB | tRNA-splicing ligase R | NP_055121.1 | 0 | 10.6062974 |
| 7657056 | EHD3 | EH domain-containing | NP_055415.1 | 0 | 13.8865019 |
| 7657381 | PRPF19 | pre-mRNA-processing | NP_055317.1 | 0 | 2.70158362 |
| 7705419 | HPCAL4 | hippocalcin-like protei | NP_057341.1 | 0 | 17.0172055 |
| 7706497 | CMPK1 | UMP-CMP kinase isof | NP_057392.1 | 0 | 11.4619032 |
| 7706567 | GNG13 | guanine nucleotide-bin | NP_057625.1 | 0 | 5.80910828 |
| 7706757 | ATP6V1D | V-type proton ATPase | NP_057078.1 | 0 | 2.66776358 |
| 8922720 | TMEM30A | cell cycle control prot | NP_060717.1 | 0.0188216 | 1.1890286 |
| 8923911 | LANCL2 | lanC-like protein 2 [Ho | NP_061167.1 | 0 | 2.78701382 |
| 8923930 | CISD1 | CDGSH iron-sulfur dor | NP_060934.1 | 0 | 3.07727454 |
| 9257257 | WDR1 | WD repeat-containing | NP_059830.1 | 0 | 15.2833451 |
| 9845509 | RAC1 | ras-related C3 botulin | NP_061485.1 | 0 | 12.0230043 |
| 9945322 | SLC17A7 | vesicular glutamate tr | NP_064705.1 | 0 | 9.98058027 |
| 9945439 | 5-Sep | septin-5 isoform 1 [Ho | NP_002679.2 | 0 | 20.5366622 |
| 9951915 | AHCY | adenosylhomocystein | NP_000678.1 | 0 | 3.99131987 |
| 9966913 | ACTR3B | actin-related protein 3 | NP_065178.1 | 0 | 13.7070972 |
| 10092657 | NDUFA12 | NADH dehydrogenase | NP_061326.1 | 0.00030826 | 2.36211018 |
| 10567816 | GNAO1 | guanine nucleotide-bin | NP_066268.1 | 0 | 50.0583461 |
| 10800140 | HIST1H2BB | histone H2B type 1-B | NP_066406.1 | 0 | 15.2883359 |
| 10800412 | MAPRE3 | microtubule-associate | NP_036458.2 | 0 | 7.6436753 |
| 10835063 | NPM1 | nucleophosmin isofor | NP_002511.1 | 0.00813008 | 1.40616034 |
| 10863895 | TMSB10 | thymosin beta-10 [Hor | NP_066926.1 | 0 | 2.57202729 |
| 10863927 | PPIA | peptidyl-prolyl cis-tran | NP_066953.1 | 0 | 36.5297465 |
| 10880989 | RAB18 | ras-related protein Ra | NP_067075.1 | 0 | 4.6452393 |
| 11024700 | TIMM13 | mitochondrial import in | NP_036590.1 | 0 | 6.37726803 |
| 11056044 | PPA1 | inorganic pyrophospha | NP_066952.1 | 0 | 9.81936753 |
| 11056046 | CADM3 | cell adhesion molecul | NP_067012.1 | 0 | 4.27769521 |
| 11056061 | TMSB4X | thymosin beta-4 [Hom | NP_066932.1 | 0.00813008 | 1.40860145 |
| 11128019 | CYCS | cytochrome c [Homo s | NP_061820.1 | 0 | 13.2745194 |
| 11225258 | MAG | myelin-associated gly | NP_002352.1 | 0 | 2.88945631 |
| 11321583 | SUCLA2 | succinyl-CoA ligase [ | NP_003841.1 | 0 | 9.76613181 |
| 11545841 | OXCT2 | succinyl-CoA:3-ketoac | NP_071403.1 | 0 | 8.61771742 |
| 11612659 | FXYD7 | FXYD domain-containi | NP_071289.1 | 0.01658833 | 1.27901426 |
| 12025678 | ACTN4 | alpha-actinin-4 [Homo | NP_004915.2 | 0 | 25.0498258 |
| 12083581 | PLCB1 | 1-phosphatidylinositol | NP_056007.1 | 0 | 25.3316274 |
| 12548785 | GABRB2 | gamma-aminobutyric a | NP_068711.1 | 0 | 5.65975424 |
| 13259510 | DCTN1 | dynactin subunit 1 iso | NP_004073.2 | 0 | 5.7052323 |
| 13569885 | ITM2C | integral membrane pro | NP_112188.1 | 0 | 4.70071067 |
| 13775198 | SH3BGRL3 | SH3 domain-binding g | NP_112576.1 | 0 | 4.47729501 |
| 13775600 | SIRT2 | NAD-dependent protei | NP_036369.2 | 0 | 6.20465884 |
| 14043072 | HNRNPA2B1 | heterogeneous nuclea | NP_112533.1 | 0 | 8.78601063 |
| 14110420 | HNRNPD | heterogeneous nuclea | NP_112738.1 | 0 | 6.35457468 |
| 14141152 | HNRNPM | heterogeneous nuclea | NP_005959.2 | 0 | 3.43238556 |
| 14141168 | PCBP2 | poly(rC)-binding protei | NP_005007.2 | 0 | 7.1121281 |
| 14165437 | HNRNPK | heterogeneous nuclea | NP_112553.1 | 0 | 12.6884931 |
| 14210536 | TUBB6 | tubulin beta-6 chain is | NP_115914.1 | 0 | 110.750182 |
| 14249342 | INA | alpha-internexin [Hom | NP_116116.1 | 0 | 48.4322927 |
| 14277700 | RPS12 | 40S ribosomal protein | NP_001007.2 | 0 | 2.53536144 |
| 14591909 | RPL5 | 60S ribosomal protein | NP_000960.2 | 0.00269139 | 1.93367407 |
| 15011936 | RPS26 | 40S ribosomal protein | NP_001020.2 | 0.00322486 | 1.83743559 |
| 15055539 | RPS2 | 40S ribosomal protein | NP_002943.2 | 0 | 3.12499665 |
| 15082258 | CBX3 | chromobox protein ho | NP_009207.2 | 0.00030826 | 2.30434324 |
| 15147219 | PURB | transcriptional activat | NP_150093.1 | 0 | 3.17660416 |
| 15431288 | RPL10A | 60S ribosomal protein | NP_009035.3 | 0 | 6.36517563 |
| 15431290 | RPL11 | 60S ribosomal protein | NP_000966.2 | 0 | 9.77146465 |
| 15431293 | RPL15 | 60S ribosomal protein | NP_002939.2 | 0 | 7.65070449 |
| 15431295 | RPL13 | 60S ribosomal protein | NP_150254.1 | 0 | 6.11907213 |
| 15431301 | RPL7 | 60S ribosomal protein | NP_000962.2 | 0 | 10.1350486 |
| 15617199 | HIST3H2A | histone H2A type 3 [H | NP_254280.1 | 0 | 11.1330311 |
| 15812198 | FBXO2 | F-box only protein 2 [ | NP_036300.2 | 0.00030826 | 2.26816958 |
| 16357472 | CDC42 | cell division control pr | NP_426359.1 | 0 | 6.41032792 |
| 16418379 | STX1B | syntaxin-1B [Homo sa | NP_443106.1 | 0 | 59.1416329 |

| | | | | | |
|---|---|---|---|---|---|
| 16418397 | MAL2 | protein MAL2 [Homo s | NP_443118.1 | 0.00406268 | 1.72978715 |
| 16507237 | HSPA5 | 78 kDa glucose-regula | NP_005338.1 | 0 | 75.4617904 |
| 16579885 | RPL4 | 60S ribosomal protein | NP_000959.2 | 0 | 5.5798643 |
| 16753215 | PFN2 | profilin-2 isoform a [Hc | NP_444252.1 | 0 | 5.32292538 |
| 16933546 | RPLP0 | 60S acidic ribosomal p | NP_444505.1 | 0 | 6.41217683 |
| 17105394 | RPL23A | 60S ribosomal protein | NP_000975.2 | 0 | 9.51956613 |
| 17149836 | FKBP1A | peptidyl-prolyl cis-tran | NP_463460.1 | 0 | 12.3873178 |
| 17149842 | FKBP2 | peptidyl-prolyl cis-tran | NP_004461.2 | 0 | 4.22428632 |
| 17158044 | RPS6 | 40S ribosomal protein | NP_001001.2 | 0.01226652 | 1.36311109 |
| 17921989 | TUBA4A | tubulin alpha-4A chain | NP_005991.1 | 0 | 130.365842 |
| 17986258 | MYL6 | myosin light polypeptic | NP_066299.2 | 0 | 13.8372238 |
| 17986283 | TUBA1A | tubulin alpha-1A chain | NP_006000.2 | 0 | 152.626265 |
| 17999528 | COX6A1 | cytochrome c oxidase | NP_004364.2 | 0 | 4.09701274 |
| 17999541 | VPS35 | vacuolar protein sortin | NP_060676.2 | 0 | 10.3751622 |
| 18079216 | CASKIN1 | caskin-1 [Homo sapier | NP_065815.1 | 0 | 5.17829432 |
| 18087855 | DYNLL2 | dynein light chain 2, c | NP_542408.1 | 0 | 10.2195686 |
| 18104948 | RPL21 | 60S ribosomal protein | NP_000973.2 | 0.00708617 | 1.48306819 |
| 18379349 | VAT1 | synaptic vesicle meml | NP_006364.2 | 0 | 6.78225293 |
| 18497300 | ATP6V1G2 | V-type proton ATPase | NP_569730.1 | 0 | 8.06248751 |
| 18765733 | SNAP25 | synaptosomal-associa | NP_003072.2 | 0 | 30.5154543 |
| 19718741 | OSBPL1A | oxysterol-binding prote | NP_542164.2 | 0 | 2.6466609 |
| 19743875 | FH | fumarate hydratase, m | NP_000134.2 | 0 | 22.3194031 |
| 19913414 | AP2A1 | AP-2 complex subunit | NP_055018.2 | 0 | 35.5426543 |
| 19913424 | ATP6V1A | V-type proton ATPase | NP_001681.2 | 0 | 48.4589654 |
| 19913428 | ATP6V1B2 | V-type proton ATPase | NP_001684.2 | 0 | 64.4065658 |
| 19913432 | ATP6V0D1 | V-type proton ATPase | NP_004682.2 | 0 | 19.7456728 |
| 19923142 | KPNB1 | importin subunit beta-1 | NP_002256.2 | 0 | 10.0749568 |
| 19923191 | MCM3AP | germinal-center assoc | NP_003897.2 | 0.00296384 | 1.90170246 |
| 19923193 | ST13 | hsc70-interacting prot | NP_003923.2 | 0 | 11.8857926 |
| 19923233 | SCP2 | non-specific lipid-trans | NP_002970.2 | 0 | 2.62616885 |
| 19923445 | ATL1 | atlastin-1 isoform a [H | NP_056999.2 | 0 | 6.58469271 |
| 19923483 | RAB14 | ras-related protein Ral | NP_057406.2 | 0 | 13.8188818 |
| 19923653 | PSIP1 | PC4 and SFRS1-intera | NP_150091.2 | 0 | 4.45855276 |
| 19923748 | DLST | dihydrolipoyllysine-res | NP_001924.2 | 0.00030826 | 2.44081181 |
| 19923821 | DPYSL5 | dihydropyrimidinase-re | NP_064519.2 | 0 | 2.51984927 |
| 19924099 | SYN1 | synapsin-1 isoform Ia | NP_008881.2 | 0 | 80.9437985 |
| 19924103 | SYN2 | synapsin-2 isoform IIa | NP_598328.1 | 0 | 22.2040152 |
| 20070125 | P4HB | protein disulfide-isome | NP_000909.2 | 0 | 6.11954268 |
| 20127450 | PRKCB | protein kinase C beta | NP_002729.2 | 0 | 13.3751441 |
| 20149568 | NDUFV1 | NADH dehydrogenase | NP_009034.2 | 0 | 8.5466478 |
| 20149594 | HSP90AB1 | heat shock protein HS | NP_031381.2 | 0 | 100.660044 |
| 20149675 | EFHD2 | EF-hand domain-conta | NP_077305.2 | 0 | 7.22729767 |
| 20270343 | ARL8A | ADP-ribosylation facto | NP_620150.1 | 0 | 4.39329626 |
| 20336761 | HEBP1 | heme-binding protein 1 | NP_057071.2 | 0.00708617 | 1.47146894 |
| 21361091 | UCHL1 | ubiquitin carboxyl-term | NP_004172.2 | 0 | 20.7623333 |
| 21361103 | SLC25A12 | calcium-binding mitoch | NP_003696.2 | 0 | 39.9366569 |
| 21361114 | SLC25A11 | mitochondrial 2-oxoglu | NP_003553.2 | 0 | 10.6047914 |
| 21361176 | ALDH1A1 | retinal dehydrogenase | NP_000680.2 | 0 | 9.75029598 |
| 21361181 | ATP1A1 | sodium/potassium-trar | NP_000692.2 | 0 | 151.770038 |
| 21361337 | EIF5 | eukaryotic translation | NP_001960.2 | 0 | 2.90448196 |
| 21361370 | PYGB | glycogen phosphoryla: | NP_002853.2 | 0 | 28.6239437 |
| 21361399 | PPP2R1A | serine/threonine-protei | NP_055040.2 | 0 | 23.4426884 |
| 21361559 | VSNL1 | visinin-like protein 1 [I | NP_003376.2 | 0 | 17.5395765 |
| 21361619 | TOLLIP | toll-interacting protein | NP_061882.2 | 0 | 4.78674795 |
| 21361647 | AHCYL1 | adenosylhomocysteina | NP_006612.2 | 0 | 8.83558432 |
| 21361657 | PDIA3 | protein disulfide-isome | NP_005304.3 | 0 | 15.1596698 |
| 21361794 | CAND1 | cullin-associated NEDI | NP_060918.2 | 0 | 12.624118 |
| 21396489 | LONP1 | lon protease homolog, | NP_004784.2 | 0 | 4.84713559 |
| 21464101 | YWHAG | 14-3-3 protein gamma | NP_036611.2 | 0 | 51.2508857 |
| 21536286 | CKB | creatine kinase B-type | NP_001814.2 | 0 | 90.4893819 |
| 21624607 | COTL1 | coactosin-like protein | NP_066972.1 | 0 | 3.11599808 |
| 21686977 | CADM4 | cell adhesion molecule | NP_660339.1 | 0.00030826 | 2.2349278 |

| | | | | | |
|---|---|---|---|---|---|
| 21703367 | DIRAS2 | GTP-binding protein Di | NP_060064.2 | 0 | 3.9788107 |
| 21735492 | PPP1R1B | protein phosphatase 1 | NP_115568.2 | 0 | 3.70663745 |
| 21735621 | MDH2 | malate dehydrogenase | NP_005909.2 | 0 | 81.9749634 |
| 22035696 | SYNGR1 | synaptogyrin-1 isoform | NP_004702.2 | 0 | 3.72653573 |
| 22907052 | ARPC1A | actin-related protein 2/ | NP_006400.2 | 0 | 6.82601798 |
| 23065544 | GSTM1 | glutathione S-transfera | NP_000552.2 | 0 | 2.55548679 |
| 23110935 | PSMA1 | proteasome subunit al | NP_683877.1 | 0 | 4.85076918 |
| 23110942 | PSMA5 | proteasome subunit al | NP_002781.2 | 0 | 15.022123 |
| 23110944 | PSMA6 | proteasome subunit al | NP_002782.1 | 0 | 3.46571999 |
| 23503295 | CSNK2B | casein kinase II subur | NP_001311.3 | 0 | 2.55720677 |
| 24234688 | HSPA9 | stress-70 protein, mitc | NP_004125.3 | 0 | 44.2894732 |
| 24234747 | ILF2 | interleukin enhancer-b | NP_004506.2 | 0 | 4.23987921 |
| 24307939 | CCT5 | T-complex protein 1 su | NP_036205.1 | 0 | 17.0327625 |
| 24307949 | NCDN | neurochondrin isoform | NP_055099.1 | 0 | 13.4526445 |
| 24307999 | GPD1L | glycerol-3-phosphate d | NP_055956.1 | 0 | 4.38407897 |
| 24308071 | CNRIP1 | CB1 cannabinoid recep | NP_056278.1 | 0 | 7.58648266 |
| 24308257 | VAT1L | synaptic vesicle memb | NP_065978.1 | 0 | 12.0800904 |
| 24430151 | PSMC1 | 26S protease regulato | NP_002793.2 | 0 | 3.40252421 |
| 24497435 | PSMC5 | 26S protease regulato | NP_002796.4 | 0 | 7.06313454 |
| 24638454 | ATP2A2 | sarcoplasmic/endoplas | NP_733765.1 | 0 | 26.0402209 |
| 25777600 | PSMD1 | 26S proteasome non-A | NP_002798.2 | 0 | 3.36091213 |
| 25777713 | SKP1 | S-phase kinase-assoc | NP_733779.1 | 0.00030826 | 2.47495519 |
| 25777732 | ALDH2 | aldehyde dehydrogena | NP_000681.2 | 0 | 6.61121102 |
| 25777736 | ALDH4A1 | delta-1-pyrroline-5-carl | NP_733844.1 | 0 | 3.70885324 |
| 25952118 | CAMK2A | calcium/calmodulin-dep | NP_741960.1 | 0 | 78.2587673 |
| 27436946 | LMNA | lamin isoform A [Homo | NP_733821.1 | 0 | 6.27754872 |
| 27764867 | SYP | synaptophysin [Homo | NP_003170.1 | 0 | 3.3290124 |
| 28178832 | IDH2 | isocitrate dehydrogena | NP_002159.2 | 0 | 11.6503292 |
| 28302131 | HBG1 | hemoglobin subunit ga | NP_000550.2 | 0 | 4.3230077 |
| 29171702 | PPA2 | inorganic pyrophospha | NP_789845.1 | 0 | 3.57105571 |
| 29788768 | TUBB2B | tubulin beta-2B chain | | NP_821080.1 | 0 | 247.775042 |
| 29788785 | TUBB | tubulin beta chain isof | NP_821133.1 | 0 | 222.220383 |
| 29826321 | ADD1 | alpha-adducin isoform | NP_054908.2 | 0 | 13.7268889 |
| 30089916 | PACS1 | phosphofurin acidic cl | NP_060496.2 | 0.00210021 | 2.00621119 |
| 30089948 | PPM1E | protein phosphatase 1 | NP_055721.3 | 0 | 5.73306309 |
| 31541941 | HSPA4L | heat shock 70 kDa pro | NP_055093.2 | 0 | 16.8881323 |
| 31542528 | NECAP1 | adaptin ear-binding co | NP_056324.2 | 0 | 5.14175981 |
| 31542947 | HSPD1 | 60 kDa heat shock pro | NP_002147.2 | 0 | 103.256696 |
| 31621303 | SFXN3 | sideroflexin-3 [Homo s | NP_112233.2 | 0.00269139 | 1.96297212 |
| 31711992 | DLAT | dihydrolipoyllysine-res | NP_001922.2 | 0 | 14.7726784 |
| 32189362 | PPFIA3 | liprin-alpha-3 [Homo sa | NP_003651.1 | 0 | 2.95624487 |
| 32189394 | ATP5B | ATP synthase subunit | NP_001677.2 | 0 | 208.922047 |
| 32307148 | OGT | UDP-N-acetylglucosan | NP_858058.1 | 0.00030826 | 2.10974669 |
| 32483399 | PAK2 | serine/threonine-protei | NP_002568.2 | 0 | 4.52038762 |
| 32483416 | NEFH | neurofilament heavy p | NP_066554.2 | 0 | 22.1133592 |
| 32528282 | ACOT7 | cytosolic acyl coenzyr | NP_863654.1 | 0 | 15.4296382 |
| 32967601 | ANK3 | ankyrin-3 isoform 1 [H | NP_066267.2 | 0 | 5.50295166 |
| 33350932 | DYNC1H1 | cytoplasmic dynein 1 | | NP_001367.2 | 0 | 97.6521964 |
| 33457311 | MDP1 | magnesium-dependent | NP_612485.2 | 0 | 2.74982405 |
| 33946324 | GNAI1 | guanine nucleotide-bin | NP_002060.4 | 0 | 10.6974055 |
| 33946329 | RALA | ras-related protein Ral | NP_005393.2 | 0 | 11.1697673 |
| 34147513 | RAB7A | ras-related protein Rat | NP_004628.4 | 0 | 20.3646134 |
| 34147630 | TUFM | elongation factor Tu, r | NP_003312.3 | 0 | 14.2992095 |
| 34419635 | HSPA6 | heat shock 70 kDa pro | NP_002146.2 | 0 | 16.0279858 |
| 34482047 | AP3B2 | AP-3 complex subunit | NP_004635.2 | 0 | 4.60005772 |
| 38201690 | RAP2B | ras-related protein Rap | NP_002877.2 | 0 | 3.73306309 |
| 38327039 | HSPA4 | heat shock 70 kDa pro | NP_002145.3 | 0 | 51.8593199 |
| 38327625 | CS | citrate synthase, mito | NP_004068.2 | 0 | 19.2821378 |
| 38455427 | CCT4 | T-complex protein 1 su | NP_006421.2 | 0 | 9.55163363 |
| 40254462 | GNAQ | guanine nucleotide-bin | NP_002063.2 | 0 | 18.4025088 |
| 40254947 | TMX4 | thioredoxin-related tra | NP_066979.2 | 0 | 2.90274269 |
| 40538799 | UBQLN4 | ubiquilin-4 isoform 1 [H | NP_064516.2 | 0.00813008 | 1.38774609 |

| | | | | | |
|---|---|---|---|---|---|
| 40806190 | TMEM189-UBE2V1 | TMEM189-UBE2V1 fus | NP_954673.1 | 0 | 2.49962629 |
| 41281564 | WDR37 | WD repeat-containing | NP_054742.2 | 0 | 3.21759905 |
| 41327712 | CRK | adapter molecule crk i | NP_058431.2 | 0 | 2.84893675 |
| 41393561 | LAP3 | cytosol aminopeptidas | NP_056991.2 | 0 | 8.86975739 |
| 42794769 | PAK1 | serine/threonine-protei | NP_002567.3 | 0 | 22.3611045 |
| 42794771 | TXNDC5 | thioredoxin domain-coı | NP_110437.2 | 0 | 3.55752023 |
| 44680154 | MTMR2 | myotubularin-related p | NP_057240.3 | 0.0065658 | 1.53224395 |
| 45827706 | QKI | protein quaking isoforı | NP_006766.1 | 0.00487945 | 1.58485965 |
| 45827776 | RTN1 | reticulon-1 isoform C [ | NP_996734.1 | 0.01631637 | 1.28416372 |
| 46249393 | RHOG | rho-related GTP-bindin | NP_001656.2 | 0 | 8.97958841 |
| 46389550 | ENSA | alpha-endosulfine isof | NP_996925.1 | 0 | 3.31080395 |
| 46593007 | UQCRC1 | cytochrome b-c1 comp | NP_003356.2 | 0 | 15.052629 |
| 47132585 | PRKAR2B | cAMP-dependent prote | NP_002727.2 | 0 | 29.1464586 |
| 47132622 | MTAP | S-methyl-5'-thioadenos | NP_002442.2 | 0 | 3.2052332 |
| 47717102 | ATP6V1H | V-type proton ATPase | NP_998785.1 | 0 | 25.7041387 |
| 48255951 | ATP2B2 | plasma membrane calc | NP_001001331.1 | 0 | 37.0650151 |
| 48255966 | UGP2 | UTP--glucose-1-phosp | NP_006750.3 | 0 | 15.2776989 |
| 48375173 | MAP6 | microtubule-associate | NP_149052.1 | 0.00708617 | 1.43711262 |
| 48762918 | PCP4 | Purkinje cell protein 4 | NP_006189.2 | 0 | 12.2885422 |
| 48762932 | CCT8 | T-complex protein 1 sı | NP_006576.2 | 0 | 10.2460869 |
| 49574491 | ATP1B2 | sodium/potassium-trar | NP_001669.3 | 0 | 4.02642127 |
| 49574532 | GSK3A | glycogen synthase kir | NP_063937.2 | 0 | 8.1758157 |
| 50263048 | GPM6B | neuronal membrane gl | NP_001001995.1 | 0 | 7.25273163 |
| 50345984 | ATP5A1 | ATP synthase subunit | NP_001001937.1 | 0 | 177.686269 |
| 50345988 | ATP5C1 | ATP synthase subunit | NP_001001973.1 | 0 | 3.75597041 |
| 50345991 | ATP5D | ATP synthase subunit | NP_001001975.1 | 0 | 4.31776464 |
| 50428938 | ASNA1 | ATPase ASNA1 [Homc | NP_004308.2 | 0 | 8.78153537 |
| 50592988 | UQCRC2 | cytochrome b-c1 comp | NP_003357.2 | 0.00708617 | 1.48691564 |
| 50592994 | TXN | thioredoxin isoform 1 [ | NP_003320.2 | 0.00708617 | 1.4661009 |
| 50592996 | TUBB3 | tubulin beta-3 chain is | NP_006077.2 | 0 | 198.628142 |
| 51092270 | C1orf95 | protein stum homolog | NP_001003665.1 | 0 | 3.38436553 |
| 51317370 | NDUFA6 | NADH dehydrogenase | NP_002481.2 | 0 | 2.49200928 |
| 52426787 | OMG | oligodendrocyte-myelir | NP_002535.3 | 0 | 2.93367407 |
| 52630440 | FKBP8 | peptidyl-prolyl cis-tran | NP_036313.3 | 0 | 5.74208155 |
| 52632383 | HNRNPL | heterogeneous nuclea | NP_001524.2 | 0 | 11.4575995 |
| 54607031 | CADM2 | cell adhesion molecule | NP_694854.2 | 0 | 4.27761291 |
| 55749577 | SLC25A4 | ADP/ATP translocase | NP_001142.2 | 0 | 37.9715863 |
| 55770878 | NPTX1 | neuronal pentraxin-1 p | NP_002513.2 | 0 | 6.60449888 |
| 55956919 | HNRNPAB | heterogeneous nuclea | NP_112556.2 | 0.00030826 | 2.38278991 |
| 56549135 | TAGLN3 | transgelin-3 [Homo sa | NP_037391.2 | 0 | 10.3644532 |
| 56676375 | TPPP3 | tubulin polymerization- | NP_057224.2 | 0 | 9.4600439 |
| 56699409 | RBMX | RNA-binding motif prot | NP_002130.2 | 0 | 4.2588484 |
| 56788381 | MOG | myelin-oligodendrocyte | NP_996532.2 | 0 | 5.25696085 |
| 57163987 | SLC8A2 | sodium/calcium excha | NP_055878.1 | 0 | 24.3266082 |
| 57165410 | ACSL6 | long-chain-fatty-acid-- | NP_056071.2 | 0 | 5.38341947 |
| 57863257 | TCP1 | T-complex protein 1 sı | NP_110379.2 | 0 | 8.46750041 |
| 58331274 | 5-Sep | septin-5 isoform 2 [Ho | NP_001009939.1 | 0 | 15.0743185 |
| 58761500 | OLA1 | obg-like ATPase 1 isoi | NP_037473.3 | 0 | 9.21469793 |
| 61743942 | ABI1 | abl interactor 1 isoforr | NP_005461.2 | 0 | 3.12569217 |
| 61744477 | SLC3A2 | 4F2 cell-surface antig | NP_001012680.1 | 0 | 4.70245833 |
| 62420877 | ETFB | electron transfer flavo | NP_001014763.1 | 0 | 4.14029623 |
| 62422571 | CRMP1 | dihydropyrimidinase-re | NP_001014809.1 | 0 | 56.0680345 |
| 63162572 | CCT3 | T-complex protein 1 sı | NP_005989.3 | 0 | 24.933217 |
| 63253298 | SRM | spermidine synthase [ | NP_003123.2 | 0 | 3.34775366 |
| 66392205 | NME2 | nucleoside diphosphat | NP_001018148.1 | 0 | 15.2219914 |
| 66932916 | MAPK1 | mitogen-activated prot | NP_002736.3 | 0 | 13.5229797 |
| 67782307 | SOD2 | superoxide dismutase | NP_001019636.1 | 0 | 7.93752885 |
| 68160915 | RPS14 | 40S ribosomal protein | NP_001020241.1 | 0 | 5.8068754 |
| 68163411 | ALCAM | CD166 antigen isoforr | NP_001618.2 | 0.00030826 | 2.29115436 |
| 68303561 | PSMA8 | proteasome subunit al | NP_653263.2 | 0 | 4.49349497 |
| 71565154 | ADH5 | alcohol dehydrogenas | NP_000662.3 | 0 | 5.94271436 |
| 71773010 | AP1G1 | AP-1 complex subunit | NP_001025178.1 | 0.00708617 | 1.45692576 |

| | | | | | | |
|---|---|---|---|---|---|---|
| 71773329 | ANXA6 | | annexin A6 isoform 1 [ | NP_001146.2 | 0 | 7.34769299 |
| 72534660 | SRSF7 | | serine/arginine-rich sp | NP_001026854.1 | 0 | 4.5457651 |
| 73486658 | GOT2 | | aspartate aminotransf | NP_002071.2 | 0 | 29.0450316 |
| 74136883 | HNRNPU | | heterogeneous nuclea | NP_114032.2 | 0 | 5.20954749 |
| 74271837 | GLUL | | glutamine synthetase | NP_001028216.1 | 0 | 5.51426438 |
| 75750480 | RHOT1 | | mitochondrial Rho GTF | NP_001028740.1 | 0 | 4.58536085 |
| 76159293 | THEM4 | | acyl-coenzyme A thioe | NP_444283.2 | 0.00406268 | 1.68634365 |
| 78000181 | RPL14 | | 60S ribosomal protein | NP_003964.3 | 0 | 4.14002156 |
| 82546843 | BCR | | breakpoint cluster regi | NP_004318.3 | 0 | 2.86518563 |
| 83267868 | DYNLL1 | | dynein light chain 1, c | NP_001032584.1 | 0 | 4.52782885 |
| 83700235 | EIF4A2 | | eukaryotic initiation fa | NP_001958.2 | 0 | 14.9202713 |
| 88758565 | KCNA3 | | potassium voltage-gat | NP_002223.3 | 0 | 5.0075718 |
| 91199540 | DLD | | dihydrolipoyl dehydrog | NP_000099.2 | 0 | 19.2440979 |
| 91208420 | BSN | | protein bassoon [Hom | NP_003449.2 | 0 | 3.57511836 |
| 91208428 | PTPRZ1 | | receptor-type tyrosine | NP_002842.2 | 0 | 12.5836593 |
| 94538322 | HAGH | | hydroxyacylglutathion | NP_005317.2 | 0 | 4.14068153 |
| 94721250 | VAPA | | vesicle-associated me | NP_003565.4 | 0 | 5.25586942 |
| 94721261 | CNP | | 2',3'-cyclic-nucleotide | NP_149124.3 | 0 | 58.6448518 |
| 95147555 | MAP1A | | microtubule-associate | NP_002364.5 | 0 | 33.1384377 |
| 96975097 | RAB6B | | ras-related protein Ral | NP_057661.3 | 0 | 13.3798163 |
| 98986464 | TMED10 | | transmembrane emp24 | NP_006818.3 | 0 | 5.95921979 |
| 103472001 | NDUFA7 | | NADH dehydrogenase | NP_004992.2 | 0.00030826 | 2.40516564 |
| 104487006 | PTPRS | | receptor-type tyrosine | NP_002841.3 | 0 | 4.5818645 |
| 105990539 | NEFL | | neurofilament light pol | NP_006149.2 | 0 | 76.5442852 |
| 109148508 | OTUB1 | | ubiquitin thioesterase | NP_060140.2 | 0 | 7.21572557 |
| 109148542 | AARS | | alanine--tRNA ligase, c | NP_001596.2 | 0.00322486 | 1.85917782 |
| 109240550 | PSPC1 | | paraspeckle compone | NP_001035879.1 | 0 | 5.23754652 |
| 109452591 | SUCLG1 | | succinyl-CoA ligase [A | NP_003840.2 | 0 | 4.52157781 |
| 112382250 | SPTBN1 | | spectrin beta chain, n | NP_003119.2 | 0 | 214.804981 |
| 114155144 | TPM3 | | tropomyosin alpha-3 c | NP_001036816.1 | 0 | 30.4784428 |
| 115334682 | SRCIN1 | | SRC kinase signaling i | NP_079524.2 | 0 | 9.21393864 |
| 115387094 | SDHB | | succinate dehydrogen | NP_002991.2 | 0 | 5.32393391 |
| 115511049 | GNA11 | | guanine nucleotide-bin | NP_002058.2 | 0 | 10.8302236 |
| 116256358 | SCAMP1 | | secretory carrier-asso | NP_004857.4 | 0 | 6.5595604 |
| 117938759 | GNAS | | protein GNAS isoform | NP_536350.2 | 0 | 14.000116 |
| 118402586 | GLO1 | | lactoylglutathione lyas | NP_006699.2 | 0 | 2.66494348 |
| 118600983 | NCAN | | neurocan core protein | NP_004377.2 | 0 | 8.52480913 |
| 122939155 | NNT | | NAD(P) transhydrogen | NP_036475.3 | 0.00296384 | 1.86614187 |
| 145386562 | MRAS | | ras-related protein M-F | NP_036351.3 | 0 | 4.88372441 |
| 148233338 | CTNNB1 | | catenin beta-1 [Homo | NP_001091679.1 | 0 | 5.95467702 |
| 148277037 | AAK1 | | AP2-associated protei | NP_055726.3 | 0 | 10.7253804 |
| 148612849 | KIF2A | | kinesin-like protein KII | NP_001091981.1 | 0 | 3.23642228 |
| 148727331 | USP5 | | ubiquitin carboxyl-term | NP_001092006.1 | 0 | 29.2229467 |
| 148727341 | STRAP | | serine-threonine kinas | NP_009109.3 | 0.00322486 | 1.83564714 |
| 150170704 | PHYHIP | | phytanoyl-CoA hydrox | NP_055574.3 | 0 | 6.08634072 |
| 153070260 | MARCKS | | myristoylated alanine-r | NP_002347.5 | 0.01226652 | 1.35477429 |
| 153791492 | EIF3CL | | eukaryotic translation | NP_001093131.1 | 0.00461494 | 1.60924147 |
| 153792590 | HSP90AA1 | | heat shock protein HS | NP_001017963.2 | 0 | 84.5833689 |
| 153945728 | MAP1B | | microtubule-associate | NP_005900.2 | 0 | 16.9487218 |
| 153946409 | CALB2 | | calretinin isoform 1 [H | NP_001731.2 | 0 | 16.6110891 |
| 154354964 | IMMT | | MICOS complex subur | NP_006830.2 | 0 | 19.8575093 |
| 155030192 | RAP1GDS1 | | rap1 GTPase-GDP dis | NP_001093896.1 | 0 | 5.32661042 |
| 156071459 | SLC25A5 | | ADP/ATP translocase | NP_001143.2 | 0 | 48.2901535 |
| 156104878 | GLS | | glutaminase kidney isc | NP_055720.3 | 0 | 23.3095948 |
| 156104880 | GMPR | | GMP reductase 1 [Hor | NP_006868.3 | 0 | 6.20950372 |
| 156151366 | LGALSL | | galectin-related protei | NP_054900.2 | 0 | 5.89414933 |
| 156416000 | UFC1 | | ubiquitin-fold modifier- | NP_057490.2 | 0.00406268 | 1.7212464 |
| 156416003 | SDHA | | succinate dehydrogen | NP_004159.2 | 0 | 4.69742303 |
| 156564403 | PDHB | | pyruvate dehydrogena | NP_000916.2 | 0 | 17.818221 |
| 156602655 | | 3-Sep | neuronal-specific sept | NP_663786.2 | 0 | 8.6494213 |
| 157384973 | TNR | | tenascin-R precursor [ | NP_003276.3 | 0 | 5.90130307 |
| 157412328 | CORO1C | | coronin-1C isoform a [ | NP_001098707.1 | 0.00210021 | 2.01605814 |

| | | | | | |
|---|---|---|---|---|---|
| 157649073 | CAP1 | adenylyl cyclase-asso | NP_001099000.1 | 0 | 3.97930906 |
| 157738645 | PLXNA4 | plexin-A4 isoform 1 pr | NP_065962.1 | 0 | 4.63357704 |
| 157738649 | NEFM | neurofilament medium | NP_005373.2 | 0 | 66.0923758 |
| 158937236 | NPEPPS | puromycin-sensitive a | NP_006301.3 | 0 | 25.4032815 |
| 163644321 | UQCRFS1 | cytochrome b-c1 comp | NP_005994.2 | 0 | 3.72223935 |
| 163659856 | GRIA3 | glutamate receptor 3 i | NP_015564.4 | 0 | 6.78879955 |
| 166795297 | DYNC1LI1 | cytoplasmic dynein 1 l | NP_057225.2 | 0 | 6.46136997 |
| 189083724 | GABRA1 | gamma-aminobutyric a | NP_001121115.1 | 0.00030826 | 2.34151162 |
| 189491755 | PAK3 | serine/threonine-protei | NP_001121640.1 | 0 | 11.056038 |
| 190194363 | DPYSL4 | dihydropyrimidinase-re | NP_006417.2 | 0.00030826 | 2.47198366 |
| 190341060 | PHYHIPL | phytanoyl-CoA hydrox | NP_115815.2 | 0.00030826 | 2.13229687 |
| 190358517 | RAB11B | ras-related protein Ral | NP_004209.2 | 0 | 9.10684279 |
| 190885499 | COX5A | cytochrome c oxidase | NP_004246.2 | 0 | 21.839264 |
| 192449447 | PLP1 | myelin proteolipid prot | NP_001122306.1 | 0 | 28.6572196 |
| 193211616 | NIPSNAP1 | protein NipSnap homol | NP_003625.2 | 0 | 7.15009239 |
| 194018537 | PRPSAP1 | phosphoribosyl pyroph | NP_002757.2 | 0 | 4.19581809 |
| 194097323 | ECHS1 | enoyl-CoA hydratase, | NP_004083.3 | 0 | 4.9437947 |
| 194578909 | EIF4E | eukaryotic translation | NP_001124151.1 | 0.01444444 | 1.30812318 |
| 197927160 | SLC4A4 | electrogenic sodium bi | NP_001128214.1 | 0 | 13.8055514 |
| 198041678 | SLC12A5 | solute carrier family 12 | NP_001128243.1 | 0 | 11.5821977 |
| 203098753 | PDHX | pyruvate dehydrogena | NP_003468.2 | 0 | 5.29311706 |
| 208431771 | C11orf68 | UPF0696 protein C11o | NP_001129107.1 | 0 | 4.5636783 |
| 208879465 | VDAC3 | voltage-dependent ani | NP_001129166.1 | 0 | 39.9034898 |
| 208973246 | QDPR | dihydropteridine reduc | NP_000311.2 | 0 | 9.29972906 |
| 212549553 | ILF3 | interleukin enhancer-b | NP_060090.2 | 0 | 3.5965363 |
| 215490011 | AIMP1 | aminoacyl tRNA synth | NP_001135888.1 | 0 | 6.61834352 |
| 216548223 | SV2A | synaptic vesicle glyco | NP_055664.3 | 0 | 6.90362801 |
| 217330598 | GLOD4 | glyoxalase domain-cor | NP_057164.3 | 0 | 2.65345144 |
| 219555707 | EIF5A | eukaryotic translation | NP_001137232.1 | 0 | 6.18087826 |
| 219804203 | RPH3A | rabphilin-3A isoform 1 | NP_001137326.1 | 0 | 9.74268254 |
| 221307584 | PHB2 | prohibitin-2 isoform 1 [ | NP_001138303.1 | 0 | 17.6083249 |
| 221625487 | IMPA1 | inositol monophosphat | NP_001138350.1 | 0 | 8.7307407 |
| 222352151 | PCBP1 | poly(rC)-binding protei | NP_006187.2 | 0 | 4.70092874 |
| 224028244 | NONO | non-POU domain-conta | NP_001138880.1 | 0 | 2.89997427 |
| 224451142 | STMN1 | stathmin isoform b [Ho | NP_001138926.1 | 0 | 18.9723239 |
| 224465190 | SCRN1 | secernin-1 isoform b [ | NP_001138986.1 | 0 | 10.9911694 |
| 224831253 | OPA1 | dynamin-like 120 kDa | NP_570850.2 | 0 | 14.0824593 |
| 226529917 | TPI1 | triosephosphate isome | NP_001152759.1 | 0 | 40.2224639 |
| 237757310 | SYNJ1 | synaptojanin-1 isoform | NP_003886.3 | 0 | 36.1394032 |
| 241982780 | PDCD6IP | programmed cell death | NP_001155901.1 | 0 | 4.97290044 |
| 256222019 | RAB10 | ras-related protein Ral | NP_057215.3 | 0 | 21.9467801 |
| 260099723 | LDHA | L-lactate dehydrogena | NP_001158886.1 | 0 | 33.5806529 |
| 260436862 | AP1B1 | AP-1 complex subunit | NP_001118.3 | 0 | 25.0961895 |
| 260763955 | NDUFA13 | NADH dehydrogenase | NP_057049.5 | 0.00210021 | 2.01542768 |
| 285002233 | GPD2 | glycerol-3-phosphate c | NP_000399.3 | 0 | 30.0176024 |
| 289666746 | FBXL16 | F-box/LRR-repeat prot | NP_699181.2 | 0 | 5.91039582 |
| 290463102 | PGM1 | phosphoglucomutase-1 | NP_001166289.1 | 0 | 9.23436057 |
| 291575128 | LDHB | L-lactate dehydrogena | NP_001167568.1 | 0 | 55.9053559 |
| 294832006 | PPP2R2A | serine/threonine-protei | NP_001171062.1 | 0 | 11.730831 |
| 294862256 | SUCLG2 | succinyl-CoA ligase [G | NP_001171070.1 | 0 | 2.62269375 |
| 295821216 | PCCB | propionyl-CoA carboxy | NP_001171485.1 | 0 | 3.82798119 |
| 296080746 | CTTN | src substrate cortactin | NP_001171669.1 | 0 | 5.40403336 |
| 296317337 | VDAC2 | voltage-dependent ani | NP_001171712.1 | 0 | 15.8622021 |
| 297747286 | PFDN6 | prefoldin subunit 6 [Ho | NP_001172110.1 | 0.01985854 | 1.16222223 |
| 302058252 | RASAL1 | rasGAP-activating-like | NP_001180449.1 | 0.01906318 | 1.16551571 |
| 308818195 | DPYSL2 | dihydropyrimidinase-re | NP_001184222.1 | 0 | 129.30119 |
| 311771647 | ARPC4 | actin-related protein 2/ | NP_001185709.1 | 0 | 5.60748304 |
| 313569822 | RPL17-C18orf32 | RPL17-C18orf32 prote | NP_001186284.1 | 0 | 2.55533077 |
| 315259111 | NEDD8-MDP1 | NEDD8-MDP1 protein | | NP_001186752.1 | 0.00030826 | 2.24078557 |
| 316659409 | ACTG1 | actin, cytoplasmic 2 [ | NP_001186883.1 | 0 | 144.759462 |
| 316983160 | NDUFS1 | NADH-ubiquinone oxid | NP_001186913.1 | 0 | 28.5222256 |
| 333944015 | CDH13 | cadherin-13 isoform 2 | NP_001207417.1 | 0 | 3.37530547 |

| | | | | | |
|---|---|---|---|---|---|
| 334688844 | GFAP | glial fibrillary acidic pro | NP_001229305.1 | 0 | 17.4418599 |
| 336285440 | NCAM1 | neural cell adhesion m | NP_001229536.1 | 0 | 8.10575405 |
| 338827685 | AP2A2 | AP-2 complex subunit | NP_001229766.1 | 0 | 41.6599545 |
| 342187211 | ALDOA | fructose-bisphosphate | NP_001230106.1 | 0 | 52.0258736 |
| 348041302 | PI4KA | phosphatidylinositol 4- | NP_477352.3 | 0 | 7.16279047 |
| 354983493 | PCMT1 | protein-L-isoaspartate | NP_001238978.1 | 0.00030826 | 2.30777016 |
| 363498929 | UQCRB | cytochrome b-c1 comp | NP_001241681.1 | 0 | 4.97387548 |
| 371940940 | ATP1A3 | sodium/potassium-tran | NP_001243143.1 | 0 | 195.837585 |
| 379030615 | DNAJC6 | putative tyrosine-prote | NP_001243793.1 | 0 | 6.59879871 |
| 380837121 | SEC22B | vesicle-trafficking prot | NP_004883.3 | 0 | 3.37396775 |
| 384381488 | GRIA1 | glutamate receptor 1 is | NP_001244950.1 | 0.00322486 | 1.84193921 |
| 384475521 | TKT | transketolase isoform | NP_001244957.1 | 0 | 14.7696392 |
| 386869503 | RPS3 | 40S ribosomal protein | NP_001247435.1 | 0 | 2.67427914 |
| 388240768 | RTN3 | reticulon-3 isoform e [I | NP_001252518.1 | 0 | 2.64608377 |
| 507834101 | MPP2 | MAGUK p55 subfamily | NP_001265299.1 | 0.00181214 | 2.10757145 |
| 510937025 | DNM1L | dynamin-1-like protein | NP_001265393.1 | 0 | 33.9804659 |
| 514052670 | ARPC3 | actin-related protein 2/ | NP_001265485.1 | 0.00813008 | 1.41623463 |
| 525507379 | CPNE6 | copine-6 isoform 1 [Ho | NP_001267487.1 | 0 | 3.99584412 |
| 527498279 | PHB | prohibitin isoform 1 [Ho | NP_001268425.1 | 0 | 29.5409357 |
| 530360487 | PARK7 | protein DJ-1 isoform X | XP_005263481.1 | 0 | 12.1337723 |
| 530363144 | SFPQ | splicing factor, proline | XP_005271170.1 | 0 | 6.15329185 |
| 530364544 | DNM3 | dynamin-3 isoform X1 | XP_005245136.1 | 0 | 62.7601445 |
| 530365106 | PSMD4 | 26S proteasome non-A | XP_005245411.1 | 0 | 6.4867824 |
| 530365929 | BPNT1 | 3'(2'),5'-bisphosphate | XP_005273056.1 | 0.01859956 | 1.21795758 |
| 530366978 | ROCK2 | rho-associated protein | XP_005246247.1 | 0.01836623 | 1.22738495 |
| 530367503 | NRBP1 | nuclear receptor-bindir | XP_005264331.1 | 0 | 4.1406215 |
| 530369146 | BIN1 | myc box-dependent-in | XP_005263699.1 | 0 | 12.9285757 |
| 530369643 | ABI2 | abl interactor 2 isoforr | XP_005246274.1 | 0 | 6.81407262 |
| 530369990 | HNRNPA3 | heterogeneous nuclea | XP_005246437.1 | 0 | 12.3819857 |
| 530372834 | PRKAR2A | cAMP-dependent prote | XP_005265370.1 | 0 | 10.3772598 |
| 530374617 | LSAMP | limbic system-associa | XP_005247511.1 | 0 | 3.48958905 |
| 530378226 | CAMK2D | calcium/calmodulin-de | XP_005263308.1 | 0 | 37.5692231 |
| 530380562 | PCDH1 | protocadherin-1 isofor | XP_005268509.1 | 0.0065658 | 1.52302353 |
| 530380819 | CPLX2 | complexin-2 isoform X | XP_005265856.1 | 0 | 6.58055472 |
| 530381854 | HIST1H2BD | histone H2B type 1-D | XP_005249096.1 | 0 | 21.9121424 |
| 530382286 | CPNE5 | copine-5 isoform X4 [H | XP_005249304.1 | 0 | 6.14738486 |
| 530387660 | PTK2B | protein-tyrosine kinase | XP_005273504.1 | 0 | 7.79951306 |
| 530388906 | PABPC1 | polyadenylate-binding | XP_005250918.1 | 0 | 13.0811118 |
| 530389143 | PLEC | plectin isoform X3 [Ho | XP_005251033.1 | 0.00181214 | 2.09996076 |
| 530389317 | YWHAZ | 14-3-3 protein zeta/de | XP_005251118.1 | 0 | 83.2981917 |
| 530390646 | DNM1 | dynamin-1 isoform X4 | XP_005251820.1 | 0 | 134.423294 |
| 530390694 | AK1 | adenylate kinase isoer | XP_005251843.1 | 0 | 3.01623844 |
| 530391273 | AUH | methylglutaconyl-CoA | XP_005252123.1 | 0 | 3.92081875 |
| 530392189 | PFKP | ATP-dependent 6-phos | XP_005252522.1 | 0 | 10.1098954 |
| 530393366 | HSPA12A | heat shock 70 kDa pro | XP_005269729.1 | 0 | 42.5772356 |
| 530393496 | HK1 | hexokinase-1 isoform | XP_005269792.1 | 0 | 45.7227677 |
| 530396203 | ME3 | NADP-dependent malic | XP_005273774.1 | 0 | 4.8645493 |
| 530396818 | NDUFS8 | NADH dehydrogenase | XP_005274070.1 | 0 | 2.64187471 |
| 530397184 | SPTBN2 | spectrin beta chain, no | XP_005274250.1 | 0 | 57.6533625 |
| 530398179 | NTM | neurotrimin isoform X2 | XP_005271637.1 | 0 | 5.35506929 |
| 530399089 | GABARAPL1 | gamma-aminobutyric a | XP_005253401.1 | 0.00487945 | 1.58720357 |
| 530400348 | ATP2B1 | plasma membrane calc | XP_005268976.1 | 0 | 25.9988711 |
| 530400466 | PFKM | ATP-dependent 6-phos | XP_005269033.1 | 0 | 13.9982686 |
| 530402316 | HMGB1 | high mobility group pro | XP_005266422.1 | 0 | 4.00421349 |
| 530402792 | DCLK1 | serine/threonine-protei | XP_005266649.1 | 0 | 20.2375809 |
| 530405580 | DMXL2 | dmX-like protein 2 isof | XP_005254312.1 | 0.00030826 | 2.28962874 |
| 530405783 | IVD | isovaleryl-CoA dehydro | XP_005254407.1 | 0 | 6.16024085 |
| 530406406 | TPM1 | tropomyosin alpha-1 c | XP_005254704.1 | 0 | 17.6682596 |
| 530406443 | AAGAB | alpha- and gamma-ada | XP_005254721.1 | 0.01859956 | 1.21666824 |
| 530406532 | TLN2 | talin-2 isoform X2 [Hor | XP_005254765.1 | 0 | 4.54959691 |
| 530409932 | TOM1L2 | TOM1-like protein 2 iso | XP_005256518.1 | 0 | 9.12694459 |
| 530410596 | VAMP2 | vesicle-associated me | XP_005256832.1 | 0 | 24.6771677 |

| | | | | | | |
|---|---|---|---|---|---|---|
| 530410945 | AP2B1 | | AP-2 complex subunit | XP_005257994.1 | 0 | 45.5800488 |
| 530410969 | ALDOC | | fructose-bisphosphate | XP_005258006.1 | 0 | 65.8446392 |
| 530411491 | CLTC | | clathrin heavy chain 1 | XP_005257069.1 | 0 | 241.677585 |
| 530412219 | MAPT | | microtubule-associate | XP_005257419.1 | 0 | 18.237553 |
| 530412393 | LUC7L3 | | luc7-like protein 3 isof | XP_005257505.1 | 0.01631637 | 1.29490639 |
| 530412417 | ATP6V0A1 | | V-type proton ATPase | XP_005257516.1 | 0 | 24.7745682 |
| 530415146 | UBA52 | | ubiquitin-60S ribosoma | XP_005260108.1 | 0 | 14.4287937 |
| 530417532 | PDCD5 | | programmed cell death | XP_005259449.1 | 0 | 3.1534155 |
| 530418431 | DYNLRB1 | | dynein light chain roac | XP_005260625.1 | 0 | 9.12096149 |
| 530420843 | SBF1 | | myotubularin-related p | XP_005261988.1 | 0 | 3.15608236 |
| 530423397 | TPP2 | | tripeptidyl-peptidase 2 | XP_005254127.1 | 0.00461494 | 1.61207653 |
| 530425723 | IDH3B | | isocitrate dehydrogena | XP_005260773.1 | 0 | 22.1019851 |
| 531990816 | CAPZB | | F-actin-capping protein | NP_001269091.1 | 0 | 8.23646645 |
| 538917681 | IDH1 | | isocitrate dehydrogena | NP_001269316.1 | 0.01985854 | 1.15839029 |
| 540344518 | EHD1 | | EH domain-containing | NP_001269374.1 | 0 | 5.09615066 |
| 543173169 | GNB1 | | guanine nucleotide-bin | NP_001269468.1 | 0 | 16.0333664 |
| 543871463 | OLFM1 | | noelin isoform 4 precu | NP_001269540.1 | 0 | 4.69572495 |
| 544063423 | STIP1 | | stress-induced-phosph | NP_001269581.1 | 0 | 13.7617498 |
| 544186032 | PDIA6 | | protein disulfide-isome | NP_001269633.1 | 0 | 11.32002 |
| 545688329 | NAPB | | beta-soluble NSF attac | NP_001269947.1 | 0 | 25.1774727 |
| 546232203 | ADAP1 | | arf-GAP with dual PH | NP_001271237.1 | 0 | 3.06153031 |
| 555943884 | SLC2A14 | | solute carrier family 2, | NP_001273166.1 | 0.00406268 | 1.73992861 |
| 555943903 | TPT1 | | translationally-controll | NP_001273201.1 | 0 | 4.49689056 |
| 557357733 | HSPH1 | | heat shock protein 10 | NP_001273433.1 | 0 | 12.6131526 |
| 574584816 | TUBB4A | | tubulin beta-4A chain i | NP_001276058.1 | 0 | 192.244411 |
| 576583524 | GAPDH | | glyceraldehyde-3-phos | NP_001276675.1 | 0 | 98.9063719 |
| 578798717 | ADGRL2 | | latrophilin-2 isoform X | XP_006710551.1 | 0.00030826 | 2.27286558 |
| 578799563 | SARS | | serine--tRNA ligase, c | XP_006710876.1 | 0 | 5.50513361 |
| 578799914 | SGIP1 | | SH3-containing GRB2- | XP_006711024.1 | 0 | 3.73423908 |
| 578803186 | FAM49A | | protein FAM49A isofor | XP_006712171.1 | 0.00210021 | 1.9863203 |
| 578803214 | DPY30 | | protein dpy-30 homolo | XP_006712181.1 | 0 | 2.58854866 |
| 578804380 | | 2-Sep | septin-2 isoform X2 [H | XP_006712610.1 | 0 | 5.8224635 |
| 578804626 | SLC4A10 | | sodium-driven chloride | XP_006712706.1 | 0 | 11.8743213 |
| 578806987 | TFG | | protein TFG isoform X | XP_006713536.1 | 0 | 2.81276138 |
| 578808960 | TRIM2 | | tripartite motif-containi | XP_006714220.1 | 0 | 2.96137984 |
| 578811231 | SNCB | | beta-synuclein isoform | XP_006714977.1 | 0 | 31.1465182 |
| 578813410 | AMPH | | amphiphysin isoform X | XP_006715752.1 | 0 | 20.7102186 |
| 578813710 | | 7-Sep | septin-7 isoform X6 [H | XP_006715869.1 | 0 | 38.8072023 |
| 578814598 | CACNA2D1 | | voltage-dependent cal | XP_006716181.1 | 0 | 11.2130507 |
| 578816062 | OXR1 | | oxidation resistance p | XP_006716657.1 | 0 | 8.62132526 |
| 578817145 | DNM1 | | dynamin-1 isoform X3 | XP_006717055.1 | 0 | 132.775213 |
| 578817242 | SLC44A1 | | choline transporter-like | XP_006717090.1 | 0.00030826 | 2.36431524 |
| 578817664 | SH3GLB2 | | endophilin-B2 isoform | XP_006717251.1 | 0 | 4.4571788 |
| 578817797 | SPTAN1 | | spectrin alpha chain, r | XP_006717308.1 | 0 | 325.476207 |
| 578818230 | CELF2 | | CUGBP Elav-like famil | XP_006717432.1 | 0 | 7.46117501 |
| 578818565 | VIM | | vimentin isoform X1 [H | XP_006717563.1 | 0 | 19.7155039 |
| 578818924 | ADD3 | | gamma-adducin isofor | XP_006717690.1 | 0.00030826 | 2.38902062 |
| 578819834 | CAMK2G | | calcium/calmodulin-de | XP_006718056.1 | 0 | 29.4989448 |
| 578822042 | HEPACAM | | hepatocyte cell adhes | XP_006718849.1 | 0.0188216 | 1.19935164 |
| 578826688 | SCAMP5 | | secretory carrier-asso | XP_006720483.1 | 0 | 6.05065879 |
| 578827087 | PKM | | pyruvate kinase PKM | XP_006720633.1 | 0 | 85.3349673 |
| 578828264 | ROGDI | | protein rogdi homolog | XP_006721010.1 | 0 | 5.44381815 |
| 578832288 | MAPRE2 | | microtubule-associate | XP_006722438.1 | 0 | 11.6782121 |
| 578832421 | WDR7 | | WD repeat-containing | XP_006722494.1 | 0 | 3.50473326 |
| 578836565 | PFKL | | ATP-dependent 6-phos | XP_006724074.1 | 0 | 4.17923308 |
| 604723356 | NDUFA5 | | NADH dehydrogenase | NP_001278233.1 | 0 | 3.71737789 |
| 662033922 | PEA15 | | astrocytic phosphopro | NP_001284505.1 | 0 | 3.35575841 |
| 664805999 | EIF4B | | eukaryotic translation | NP_001287750.1 | 0 | 3.24902902 |
| 665821284 | PTBP2 | | polypyrimidine tract-bi | NP_001287916.1 | 0 | 3.92885471 |
| 735997436 | ACAT2 | | acetyl-CoA acetyltrans | NP_001290182.1 | 0 | 4.14399655 |
| 740086846 | ACLY | | ATP-citrate synthase i | NP_001290203.1 | 0 | 5.14739803 |
| 740087210 | MYL12A | | myosin regulatory ligh | NP_001289978.1 | 0.00296384 | 1.88239731 |

| | | | | | |
|---|---|---|---|---|---|
| 746817428 | RPL10 | 60S ribosomal protein | NP_001290553.1 | 0 | 2.80604102 |
| 747165389 | RPSA | 40S ribosomal protein | NP_001291217.1 | 0 | 10.8201769 |
| 767900757 | LOC102724023 | ES1 protein homolog, | XP_011507206.1 | 0 | 9.13639976 |
| 767903710 | PHGDH | D-3-phosphoglycerate | XP_011539528.1 | 0 | 19.0440298 |
| 767905392 | RAP1GAP | rap1 GTPase-activatin | XP_006710867.2 | 0 | 5.33068312 |
| 767908737 | NFASC | neurofascin isoform X | XP_011507620.1 | 0 | 21.3208755 |
| 767910140 | BCAN | brevican core protein i | XP_011508168.1 | 0 | 2.53357728 |
| 767910855 | ARHGEF2 | rho guanine nucleotide | XP_011508441.1 | 0.00181214 | 2.04253638 |
| 767913715 | ADD2 | beta-adducin isoform ) | XP_011530804.1 | 0 | 11.732705 |
| 767914416 | HADHB | trifunctional enzyme s | XP_011531105.1 | 0 | 2.90204893 |
| 767915105 | SLC8A1 | sodium/calcium excha | XP_011531358.1 | 0.00350365 | 1.7490923 |
| 767917469 | NGEF | ephexin-1 isoform X1 | | XP_011509225.1 | 0 | 6.64435695 |
| 767918167 | MAP2 | microtubule-associate | XP_011509492.1 | 0 | 28.7238031 |
| 767918388 | FAHD2A | fumarylacetoacetate h | XP_011509586.1 | 0.00813008 | 1.38795826 |
| 767918809 | KANSL3 | KAT8 regulatory NSL ( | XP_011509757.1 | 0.01836623 | 1.22635981 |
| 767922705 | CLASP2 | CLIP-associating prote | XP_011531807.1 | 0 | 5.72630441 |
| 767924436 | CADPS | calcium-dependent se | XP_011532473.1 | 0 | 18.4664681 |
| 767924440 | CADPS | calcium-dependent se | XP_011532475.1 | 0 | 20.4807756 |
| 767924749 | SRGAP3 | SLIT-ROBO Rho GTPa | XP_011532597.1 | 0 | 5.69314625 |
| 767924768 | IQSEC1 | IQ motif and SEC7 do | XP_011532606.1 | 0 | 3.82131076 |
| 767925902 | DLG1 | disks large homolog 1 | XP_011510797.1 | 0 | 11.9420067 |
| 767926887 | PEX5L | PEX5-related protein i | XP_011511184.1 | 0 | 3.62782471 |
| 767927354 | BDH1 | D-beta-hydroxybutyrat | XP_011511369.1 | 0 | 3.42608456 |
| 767930319 | KCTD8 | BTB/POZ domain-cont | XP_011511991.1 | 0 | 3.46192921 |
| 767931419 | RUFY3 | protein RUFY3 isoform | XP_011530052.1 | 0 | 4.48838398 |
| 767931770 | ANK2 | ankyrin-2 isoform X5 [ | XP_011530193.1 | 0 | 50.3341993 |
| 767932537 | SNCA | alpha-synuclein isofor | XP_011530509.1 | 0 | 40.0884475 |
| 767933143 | G3BP2 | ras GTPase-activating | XP_011530742.1 | 0 | 4.82594019 |
| 767935355 | HAPLN1 | hyaluronan and proteo | XP_011541470.1 | 0.00322486 | 1.83713701 |
| 767937841 | UBE2D2 | ubiquitin-conjugating e | XP_011535981.1 | 0 | 2.61924623 |
| 767938112 | DBN1 | drebrin isoform X1 [Hc | XP_011532748.1 | 0 | 16.3014828 |
| 767938356 | HNRNPH1 | heterogeneous nuclea | XP_011532847.1 | 0 | 20.6407371 |
| 767938662 | CANX | calnexin isoform X2 [H | XP_011532966.1 | 0 | 5.06383826 |
| 767939866 | PACSIN1 | protein kinase C and c | XP_011512843.1 | 0 | 10.8585922 |
| 767941917 | STXBP5 | syntaxin-binding prote | XP_011533751.1 | 0 | 3.22300842 |
| 767943839 | WASF1 | wiskott-Aldrich syndro | XP_011534535.1 | 0 | 9.41715325 |
| 767943934 | SNAP91 | clathrin coat assembly | XP_011534567.1 | 0 | 28.0061661 |
| 767945521 | OGDH | 2-oxoglutarate dehydr | XP_011513710.1 | 0 | 18.1268153 |
| 767945559 | MPP6 | MAGUK p55 subfamily | XP_011513727.1 | 0 | 4.03475125 |
| 767945863 | CAMK2B | calcium/calmodulin-de | XP_011513849.1 | 0 | 47.4392154 |
| 767947915 | NRCAM | neuronal cell adhesion | XP_011514557.1 | 0 | 7.77873173 |
| 767948171 | CHCHD3 | MICOS complex subur | XP_011514665.1 | 0 | 5.74933608 |
| 767948221 | PPP1R9A | neurabin-1 isoform X3 | XP_011514685.1 | 0.00296384 | 1.92009553 |
| 767948685 | EIF4H | eukaryotic translation | XP_011514858.1 | 0 | 13.2514559 |
| 767950720 | PSD3 | PH and SEC7 domain- | XP_011542764.1 | 0 | 7.66064557 |
| 767951827 | GDAP1 | ganglioside-induced di | XP_011515853.1 | 0 | 14.4003762 |
| 767953037 | ASAP1 | arf-GAP with SH3 dom | XP_011515354.1 | 0.00181214 | 2.05645602 |
| 767953162 | FAM49B | protein FAM49B isofor | XP_011515409.1 | 0 | 6.4711347 |
| 767953706 | NCALD | neurocalcin-delta isofc | XP_011515637.1 | 0 | 7.1482472 |
| 767954696 | ALDH1B1 | aldehyde dehydrogena | XP_011516104.1 | 0 | 4.70333141 |
| 767954754 | KIAA1045 | protein KIAA1045 isof | XP_011516129.1 | 0 | 2.71264623 |
| 767955207 | SH3GL2 | endophilin-A1 isoform | XP_011516307.1 | 0 | 27.0998059 |
| 767958030 | SET | protein SET isoform X | XP_011517213.1 | 0 | 7.15428198 |
| 767958792 | GDA | guanine deaminase is | XP_011517524.1 | 0.00181214 | 2.06844043 |
| 767963194 | PPP3CB | serine/threonine-protei | XP_011538223.1 | 0 | 31.4508843 |
| 767963256 | OGDHL | 2-oxoglutarate dehydr | XP_011538249.1 | 0 | 24.6673782 |
| 767966201 | PSMC3 | 26S protease regulato | XP_011518535.1 | 0 | 7.97767062 |
| 767966323 | SLC1A2 | excitatory amino acid | XP_011518586.1 | 0 | 28.6705176 |
| 767967976 | PGM2L1 | glucose 1,6-bisphospl | XP_011543255.1 | 0.00813008 | 1.38817052 |
| 767969491 | HYOU1 | hypoxia up-regulated | XP_011540859.1 | 0 | 8.56720898 |
| 767970091 | HSPA8 | heat shock cognate 7 | XP_011541100.1 | 0 | 183.052455 |
| 767971264 | ERC1 | ELKS/Rab6-interacting | XP_011519238.1 | 0 | 3.82413029 |

| | | | | | | |
|---|---|---|---|---|---|---|
| 767971402 | PTMS | | parathymosin isoform | XP_011519289.1 | 0 | 6.49948909 |
| 767973233 | CNTN1 | | contactin-1 isoform X1 | XP_011536228.1 | 0 | 38.4298339 |
| 767974361 | NAP1L1 | | nucleosome assembly | XP_011536692.1 | 0 | 7.07788269 |
| 767977196 | WASF3 | | wiskott-Aldrich syndro | XP_011533192.1 | 0.00269139 | 1.94309515 |
| 767980422 | HNRNPC | | heterogeneous nuclea | XP_011535010.1 | 0 | 7.71018816 |
| 767981118 | NDRG2 | | protein NDRG2 isoforn | XP_011535300.1 | 0 | 4.28726614 |
| 767981762 | ACTN1 | | alpha-actinin-1 isoforn | XP_011535567.1 | 0 | 46.3225143 |
| 767982072 | RGS6 | | regulator of G-protein | XP_011535691.1 | 0 | 3.95000714 |
| 767983063 | CYFIP1 | | cytoplasmic FMR1-inte | XP_011542175.1 | 0 | 16.7632946 |
| 767983219 | CKMT1B | | creatine kinase U-type | XP_011519499.1 | 0 | 26.0172843 |
| 767983614 | FES | | tyrosine-protein kinase | XP_011519661.1 | 0.01444444 | 1.31704307 |
| 767984251 | MYO5A | | unconventional myosir | XP_011519908.1 | 0 | 20.0515898 |
| 767985907 | SLC12A6 | | solute carrier family 12 | XP_011520569.1 | 0 | 4.66552453 |
| 767986980 | ABAT | | 4-aminobutyrate aminc | XP_011520702.1 | 0 | 8.98909944 |
| 767987102 | CARHSP1 | | calcium-regulated heat | XP_011520746.1 | 0 | 6.76649624 |
| 767987604 | UBE2I | | SUMO-conjugating enz | XP_011520947.1 | 0.00461494 | 1.59894427 |
| 767988216 | CORO1A | | coronin-1A isoform X1 | XP_011544016.1 | 0 | 16.1873588 |
| 767991742 | DLG4 | | disks large homolog 4 | XP_011522000.1 | 0 | 2.70487291 |
| 767992184 | MYH10 | | myosin-10 isoform X2 | XP_011522177.1 | 0 | 2.50111706 |
| 767992241 | PAFAH1B1 | | platelet-activating fact | XP_011522203.1 | 0.00322486 | 1.85761053 |
| 767993234 | BAIAP2 | | brain-specific angiogei | XP_011522496.1 | 0 | 5.67592342 |
| 767993263 | | 9-Sep | septin-9 isoform X3 [H | XP_011522506.1 | 0 | 4.28362432 |
| 767994460 | GIT1 | | ARF GTPase-activatin | XP_011522986.1 | 0 | 7.56177419 |
| 767994910 | NMT1 | | glycylpeptide N-tetrad | XP_011523160.1 | 0 | 5.58070528 |
| 767994920 | NSF | | vesicle-fusing ATPase | XP_011523165.1 | 0 | 89.8132042 |
| 767995054 | | 4-Sep | septin-4 isoform X2 [H | XP_011523212.1 | 0 | 5.83914639 |
| 767995236 | PRKAR1A | | cAMP-dependent prote | XP_011523286.1 | 0 | 5.83416238 |
| 767995537 | UBE2O | | E2/E3 hybrid ubiquitin- | XP_011523409.1 | 0 | 7.21282301 |
| 767996099 | PIP4K2B | | phosphatidylinositol 5- | XP_011523628.1 | 0 | 6.17548784 |
| 767997328 | FASN | | fatty acid synthase is | XP_011521840.1 | 0 | 2.54867419 |
| 767997413 | ARHGDIA | | rho GDP-dissociation i | XP_011521876.1 | 0 | 10.8506697 |
| 767997588 | EPB41L3 | | band 4.1-like protein 3 | XP_011523918.1 | 0 | 14.3892533 |
| 767997921 | NAPG | | gamma-soluble NSF at | XP_011524057.1 | 0 | 39.8802826 |
| 767998780 | MBP | | Golli-MBP isoform X1 | | XP_011524311.1 | 0 | 31.4421631 |
| 767999390 | HDHD2 | | haloacid dehalogenase | XP_011524529.1 | 0 | 4.1834273 |
| 768001322 | ADGRL1 | | latrophilin-1 isoform X | XP_011526099.1 | 0 | 3.46724562 |
| 768003717 | ICAM5 | | intercellular adhesion | XP_011526531.1 | 0.00350365 | 1.8173001 |
| 768006376 | AP2S1 | | AP-2 complex subunit | XP_011524725.1 | 0.01790634 | 1.23987921 |
| 768008086 | GPI | | glucose-6-phosphate i | XP_011525056.1 | 0 | 4.1715185 |
| 768014078 | NSFL1C | | NSFL1 cofactor p47 is | XP_011527602.1 | 0 | 4.57218943 |
| 768016461 | EPB41L1 | | band 4.1-like protein 1 | XP_011526969.1 | 0 | 10.1351871 |
| 768017580 | ARFGAP1 | | ADP-ribosylation facto | XP_011527203.1 | 0 | 2.78225293 |
| 768022887 | SLC25A18 | | mitochondrial glutamat | XP_011544451.1 | 0 | 7.64351285 |
| 768023906 | EWSR1 | | RNA-binding protein E | XP_011528297.1 | 0.00322486 | 1.83833259 |
| 768032495 | PDHA1 | | pyruvate dehydrogena | XP_011543833.1 | 0 | 6.88101335 |
| 768033352 | UBA1 | | ubiquitin-like modifier-a | XP_011542255.1 | 0 | 31.8342532 |
| 768037765 | MSN | | moesin isoform X1 [Hc | XP_011529261.1 | 0.01006993 | 1.38404995 |
| 768039038 | | 6-Sep | septin-6 isoform X2 [H | XP_011529619.1 | 0 | 20.9050447 |
| 768039088 | HPRT1 | | hypoxanthine-guanine | XP_011529630.1 | 0 | 8.92336261 |
| 768054000 | RPS9 | | 40S ribosomal protein | XP_011546661.1 | 0 | 5.46673648 |
| 779176942 | CALM2 | | calmodulin isoform 1 [ | NP_001292553.1 | 0 | 19.9508093 |
| 807066321 | | 11-Sep | septin-11 isoform 1 [H | NP_001293076.1 | 0 | 23.2406876 |
| 815891093 | RPS15 | | 40S ribosomal protein | NP_001295155.1 | 0 | 6.49811951 |
| 909618061 | AP2M1 | | AP-2 complex subunit | NP_001298127.1 | 0 | 27.766074 |
| 927669104 | RHOA | | transforming protein R | NP_001300870.1 | 0 | 2.98653329 |
| 938403405 | PRKCG | | protein kinase C gamn | NP_001303258.1 | 0 | 11.6585472 |
| 939646432 | MDH1 | | malate dehydrogenase | NP_001303303.1 | 0 | 20.6737961 |

| Coverage | # Peptides | # PSMs | # Unique Peptides | # Protein Groups | # AAs | MW [kDa] | calc. pI | Abundance Ratio: (F2, 127) / (F2, 126) |
|---|---|---|---|---|---|---|---|---|
| 16.5384615 | 12 | 38 | 12 | 1 | 780 | 85.372 | 7.61 | 0.85 |
| 32.0954907 | 17 | 88 | 3 | 1 | 377 | 42.024 | 5.39 | 0.671 |
| 2.04326923 | 1 | 2 | 1 | 1 | 832 | 91.867 | 7.64 | |
| 1.47783251 | 1 | 2 | 1 | 1 | 609 | 69.321 | 6.28 | 2.491 |
| 9.49367089 | 1 | 2 | 1 | 1 | 316 | 35.83 | 6.98 | 0.853 |
| 10.3125 | 3 | 6 | 3 | 1 | 320 | 35.914 | 5.05 | 0.794 |
| 28.7292818 | 3 | 8 | 3 | 1 | 181 | 20.684 | 6.8 | 0.612 |
| 5.55555556 | 1 | 2 | 1 | 1 | 180 | 20.517 | 6.79 | 0.813 |
| 8.83977901 | 1 | 2 | 1 | 1 | 181 | 20.404 | 5.72 | |
| 30.6862745 | 30 | 77 | 14 | 1 | 1020 | 112.193 | 5.66 | 0.866 |
| 22.7722772 | 10 | 33 | 10 | 1 | 303 | 35.039 | 8.53 | 0.797 |
| 11.7370892 | 2 | 7 | 2 | 1 | 213 | 23.263 | 9.96 | 0.864 |
| 9.68586387 | 4 | 8 | 4 | 1 | 382 | 43.914 | 7.46 | 0.78 |
| 42.0353982 | 13 | 32 | 13 | 1 | 226 | 26.129 | 8 | 0.754 |
| 12.0567376 | 1 | 2 | 1 | 1 | 282 | 31.343 | 4.84 | 1.635 |
| 4.33436533 | 2 | 4 | 2 | 1 | 323 | 35.191 | 7.21 | 0.86 |
| 14.2857143 | 1 | 2 | 1 | 1 | 63 | 7.241 | 10.27 | |
| 2.94117647 | 2 | 4 | 2 | 1 | 476 | 53.117 | 5.14 | 0.832 |
| 3.18471338 | 1 | 1 | 1 | 1 | 314 | 33.754 | 5.14 | |
| 14.8337596 | 3 | 11 | 3 | 1 | 391 | 45.115 | 7.74 | 0.812 |
| 26.1403509 | 10 | 36 | 7 | 1 | 570 | 61.924 | 6.49 | 0.888 |
| 22.2943723 | 8 | 20 | 3 | 1 | 462 | 50.109 | 9.01 | 0.796 |
| 19.0064795 | 7 | 18 | 2 | 1 | 463 | 50.438 | 9.03 | 0.931 |
| 7.55148741 | 4 | 9 | 4 | 1 | 437 | 50.087 | 6.67 | 0.957 |
| 8.62470862 | 4 | 10 | 4 | 1 | 858 | 95.277 | 6.83 | 0.85 |
| 30.6451613 | 12 | 44 | 11 | 1 | 434 | 47.139 | 7.39 | 0.714 |
| 16.2162162 | 3 | 6 | 3 | 1 | 333 | 35.058 | 8.38 | 1.286 |
| 29.082774 | 10 | 35 | 4 | 1 | 447 | 50.55 | 5.14 | 0.826 |
| 12.6760563 | 3 | 12 | 2 | 1 | 355 | 40.425 | 5.54 | 0.832 |
| 18.1598063 | 6 | 12 | 6 | 1 | 413 | 46.219 | 7.01 | 0.714 |
| 50.7692308 | 4 | 10 | 2 | 1 | 130 | 14.087 | 10.9 | |
| 52.4271845 | 7 | 18 | 7 | 1 | 103 | 11.36 | 11.36 | 0.853 |
| 25.3521127 | 4 | 12 | 4 | 1 | 142 | 15.248 | 8.68 | 1.495 |
| 5.28967254 | 2 | 4 | 2 | 1 | 397 | 44.839 | 7.08 | |
| 49.0196078 | 6 | 12 | 6 | 1 | 102 | 10.925 | 8.92 | 0.867 |
| 1.96399345 | 1 | 1 | 1 | 1 | 611 | 69.241 | 6.18 | |
| 9.56521739 | 1 | 1 | 1 | 1 | 115 | 12.468 | 7.88 | |
| 12.345679 | 1 | 4 | 1 | 1 | 81 | 9.364 | 9.38 | 0.724 |
| 5.71428571 | 1 | 2 | 1 | 1 | 175 | 20.095 | 10.3 | 0.763 |
| 41.2698413 | 5 | 10 | 1 | 1 | 189 | 21.216 | 5.17 | |
| 7.82608696 | 2 | 6 | 2 | 1 | 345 | 37.983 | 6.87 | 0.837 |
| 1.11524164 | 1 | 2 | 1 | 1 | 807 | 89.365 | 7.3 | |
| 3.930131 | 1 | 2 | 1 | 1 | 229 | 25.553 | 5.92 | 0.815 |
| 13.9037433 | 2 | 4 | 2 | 1 | 187 | 21.044 | 7.53 | 0.788 |
| 4.1445271 | 2 | 4 | 2 | 1 | 941 | 105.649 | 5.41 | |
| 10.5769231 | 1 | 1 | 1 | 1 | 104 | 11.545 | 9.1 | |
| 50.3937008 | 10 | 38 | 10 | 1 | 254 | 28.786 | 7.18 | 0.849 |
| 27.8177458 | 11 | 30 | 11 | 1 | 417 | 44.586 | 8.1 | 0.716 |
| 5 | 1 | 2 | 1 | 1 | 360 | 41.539 | 4.91 | |
| 3.55987055 | 1 | 2 | 1 | 1 | 309 | 35.571 | 5.54 | |
| 11.7647059 | 1 | 6 | 1 | 1 | 170 | 19.288 | 4.81 | 0.734 |
| 9.4017094 | 3 | 6 | 3 | 1 | 351 | 40.564 | 8.79 | 0.693 |
| 8.11965812 | 1 | 2 | 1 | 1 | 234 | 25.882 | 7.43 | |
| 3.52941176 | 1 | 2 | 1 | 1 | 255 | 28.415 | 5.33 | |
| 8.71369295 | 1 | 2 | 1 | 1 | 241 | 26.472 | 8.13 | 0.919 |
| 11.3163972 | 3 | 8 | 3 | 1 | 433 | 48.603 | 5.95 | 0.987 |
| 27.3584906 | 5 | 12 | 5 | 1 | 212 | 23.531 | 6.54 | 0.709 |

| | | | | | | | | |
|---|---|---|---|---|---|---|---|---|
| 55.4545455 | 13 | 46 | 11 | 1 | 220 | 24.968 | 5.03 | 0.668 |
| 35.8695652 | 5 | 10 | 5 | 1 | 184 | 20.974 | 6.67 | 0.794 |
| 5.45454545 | 1 | 2 | 1 | 1 | 165 | 17.808 | 9.42 | |
| 5.71428571 | 1 | 2 | 1 | 1 | 140 | 14.856 | 10.51 | |
| 8.78378378 | 1 | 2 | 1 | 1 | 148 | 16.551 | 11 | |
| 10.4347826 | 1 | 2 | 1 | 1 | 115 | 12.776 | 9.63 | |
| 34.2857143 | 2 | 6 | 2 | 1 | 70 | 8.213 | 10.1 | 0.866 |
| 2.72952854 | 1 | 2 | 1 | 1 | 403 | 46.08 | 10.18 | |
| 9.39849624 | 2 | 4 | 2 | 1 | 266 | 29.977 | 10.61 | |
| 6.22568093 | 1 | 2 | 1 | 1 | 257 | 28.007 | 11.03 | 0.94 |
| 14.7826087 | 2 | 4 | 2 | 1 | 115 | 11.658 | 4.54 | |
| 1.31795717 | 1 | 2 | 1 | 1 | 607 | 68.527 | 6.38 | |
| 6.96202532 | 1 | 2 | 1 | 1 | 158 | 18.419 | 10.3 | 0.863 |
| 27.1523179 | 4 | 10 | 4 | 1 | 151 | 17.212 | 10.54 | 0.818 |
| 15.7534247 | 3 | 6 | 3 | 1 | 146 | 16.435 | 10.21 | 0.781 |
| 11.7241379 | 3 | 6 | 3 | 1 | 145 | 16.051 | 10.32 | 1.074 |
| 23.1884058 | 1 | 2 | 1 | 1 | 69 | 7.836 | 10.7 | |
| 10.6060606 | 2 | 4 | 2 | 1 | 264 | 29.926 | 9.73 | 0.862 |
| 7.2243346 | 2 | 4 | 2 | 1 | 263 | 29.579 | 10.15 | |
| 18.556701 | 4 | 9 | 4 | 1 | 194 | 22.113 | 10.1 | 0.306 |
| 6.25 | 1 | 4 | 1 | 1 | 208 | 24.19 | 10.32 | 0.764 |
| 8.51926978 | 4 | 8 | 4 | 1 | 493 | 54.496 | 7.24 | 0.858 |
| 39.800995 | 21 | 95 | 21 | 1 | 603 | 68.692 | 6.77 | 0.814 |
| 10.4607721 | 7 | 18 | 5 | 1 | 803 | 92.411 | 4.84 | 0.723 |
| 57.9775281 | 28 | 261 | 2 | 1 | 445 | 49.875 | 4.89 | 0.643 |
| 4.91803279 | 1 | 2 | 1 | 1 | 183 | 20.887 | 7.69 | 0.835 |
| 35.5263158 | 3 | 6 | 3 | 1 | 152 | 17.127 | 6.57 | 0.772 |
| 39.9293286 | 11 | 36 | 10 | 1 | 283 | 30.754 | 8.54 | 0.794 |
| 3.40909091 | 1 | 2 | 1 | 1 | 528 | 59.106 | 7.05 | |
| 39.8373984 | 9 | 28 | 4 | 1 | 246 | 28.065 | 4.83 | 0.686 |
| 37.398374 | 10 | 25 | 7 | 1 | 246 | 28.201 | 4.84 | 0.81 |
| 3.7470726 | 2 | 4 | 2 | 1 | 427 | 45.171 | 8.85 | |
| 2.85714286 | 1 | 2 | 1 | 1 | 455 | 52.528 | 6.27 | 0.812 |
| 11.1538462 | 2 | 6 | 2 | 1 | 260 | 29.228 | 7.4 | 0.641 |
| 7.40740741 | 1 | 2 | 1 | 1 | 135 | 15.155 | 7.01 | |
| 2.70485282 | 3 | 6 | 3 | 1 | 1257 | 139.915 | 6.24 | 0.943 |
| 12.6582278 | 2 | 4 | 2 | 1 | 158 | 18.031 | 6.74 | |
| 3.0995106 | 1 | 2 | 1 | 1 | 613 | 66.859 | 8.95 | 0.901 |
| 6.04395604 | 1 | 2 | 1 | 1 | 182 | 20.443 | 7.24 | |
| 9.3220339 | 1 | 2 | 1 | 1 | 118 | 13.749 | 8.79 | 0.737 |
| 5.27577938 | 1 | 4 | 1 | 1 | 417 | 48.112 | 4.44 | |
| 19.1709845 | 2 | 4 | 2 | 1 | 193 | 20.554 | 8.57 | 0.8 |
| 11.3402062 | 1 | 1 | 1 | 1 | 97 | 10.991 | 5.16 | 1.102 |
| 6.76691729 | 1 | 2 | 1 | 1 | 133 | 14.849 | 6.8 | |
| 4.92957746 | 1 | 2 | 1 | 1 | 142 | 16.702 | 5.29 | 0.739 |
| 8.75 | 2 | 4 | 2 | 1 | 240 | 26.772 | 4.73 | 0.804 |
| 1.78117048 | 1 | 2 | 1 | 1 | 393 | 42.767 | 8.5 | 0.877 |
| 24.1071429 | 4 | 8 | 4 | 1 | 224 | 25.019 | 6.38 | 0.697 |
| 7.62800418 | 4 | 8 | 4 | 1 | 957 | 109.427 | 6.19 | 0.86 |
| 5.18358531 | 2 | 4 | 2 | 1 | 463 | 52.512 | 7.55 | 0.925 |
| 10.9848485 | 3 | 6 | 3 | 1 | 264 | 30.223 | 7.5 | 1.166 |
| 12.037037 | 2 | 4 | 2 | 1 | 216 | 23.728 | 9.41 | |
| 31.7073171 | 5 | 12 | 3 | 1 | 205 | 22.663 | 6.21 | 0.902 |
| 7.14285714 | 1 | 2 | 1 | 1 | 126 | 13.907 | 10.32 | |
| 21.5277778 | 8 | 16 | 8 | 1 | 288 | 33.003 | 5.24 | 0.681 |
| 5.93220339 | 2 | 3 | 2 | 1 | 354 | 40.252 | 5.44 | |
| 5.76131687 | 1 | 2 | 1 | 1 | 243 | 27.211 | 7.3 | 0.974 |
| 7.72532189 | 1 | 2 | 1 | 1 | 233 | 25.981 | 8.72 | |
| 18.3908046 | 3 | 9 | 2 | 1 | 261 | 30.006 | 4.83 | 0.861 |
| 3.76712329 | 1 | 2 | 1 | 1 | 292 | 33.283 | 7.66 | |
| 7.79467681 | 2 | 4 | 2 | 1 | 526 | 53.394 | 9.36 | 0.847 |
| 6.10412926 | 3 | 6 | 3 | 1 | 557 | 63.777 | 8.24 | 0.853 |

| | | | | | | | | |
|---|---|---|---|---|---|---|---|---|
| 5.91397849 | 1 | 2 | 1 | 1 | 186 | 21.751 | 6.8 | 0.913 |
| 14.2857143 | 2 | 6 | 2 | 1 | 140 | 15.045 | 8.27 | 0.64 |
| 7.83783784 | 2 | 6 | 2 | 1 | 370 | 40.738 | 7.21 | |
| 24.3727599 | 13 | 30 | 2 | 1 | 558 | 61.359 | 7.8 | 0.688 |
| 11.3402062 | 2 | 4 | 2 | 1 | 194 | 20.85 | 10.84 | 1.022 |
| 17.1945701 | 6 | 12 | 6 | 1 | 221 | 22.336 | 11.02 | 1.114 |
| 2.44233379 | 1 | 4 | 1 | 1 | 737 | 83.62 | 7.12 | |
| 14.0957447 | 4 | 10 | 4 | 1 | 376 | 42.587 | 6.64 | 0.844 |
| 18.2741117 | 3 | 8 | 3 | 1 | 394 | 44.732 | 6.74 | 0.818 |
| 13.6363636 | 5 | 12 | 4 | 1 | 418 | 47.341 | 5.88 | 0.779 |
| 8 | 3 | 4 | 3 | 1 | 300 | 34.311 | 7.36 | |
| 31.9277108 | 4 | 13 | 3 | 1 | 166 | 18.491 | 8.09 | 0.76 |
| 14.159292 | 1 | 2 | 1 | 1 | 113 | 12.816 | 7.37 | |
| 7.76699029 | 1 | 2 | 1 | 1 | 412 | 45.717 | 6.48 | 1.195 |
| 12.295082 | 5 | 12 | 5 | 1 | 366 | 39.566 | 6.92 | 0.918 |
| 6.2992126 | 1 | 2 | 1 | 1 | 127 | 14.469 | 5.38 | |
| 28.2608696 | 6 | 12 | 6 | 1 | 322 | 34.889 | 6.44 | 0.756 |
| 8.43230404 | 5 | 11 | 1 | 1 | 842 | 97.031 | 7.03 | 1.363 |
| 36.492891 | 16 | 48 | 16 | 1 | 422 | 47.543 | 8.12 | 0.714 |
| 7.38461538 | 2 | 4 | 2 | 1 | 325 | 36.55 | 6.79 | |
| 10.0946372 | 3 | 6 | 3 | 1 | 317 | 35.055 | 7.69 | 0.89 |
| 3.79746835 | 1 | 2 | 1 | 1 | 395 | 43.633 | 6.48 | 0.787 |
| 63.1460674 | 28 | 254 | 1 | 1 | 445 | 49.799 | 4.89 | 0.694 |
| 8.48708487 | 3 | 7 | 1 | 1 | 271 | 30.521 | 6.29 | 0.783 |
| 24.0740741 | 5 | 10 | 5 | 1 | 216 | 24.408 | 7.49 | 0.619 |
| 8.07453416 | 1 | 2 | 1 | 1 | 161 | 18.479 | 5.3 | |
| 4.82180294 | 2 | 4 | 1 | 1 | 477 | 52.791 | 6.37 | 0.951 |
| 9.79020979 | 2 | 4 | 2 | 1 | 286 | 32.929 | 5.85 | 0.888 |
| 11.7757009 | 4 | 8 | 4 | 1 | 535 | 57.452 | 6.46 | |
| 6.81399632 | 3 | 6 | 3 | 1 | 543 | 59.329 | 7.65 | |
| 8.12807882 | 3 | 5 | 3 | 1 | 406 | 44.792 | 5.17 | 0.884 |
| 18.7739464 | 4 | 8 | 4 | 1 | 261 | 29.698 | 7.05 | 0.99 |
| 3.20641283 | 1 | 1 | 1 | 1 | 499 | 56.842 | 6.28 | |
| 18.3206107 | 6 | 11 | 6 | 1 | 393 | 43.411 | 6.62 | 0.859 |
| 28.358209 | 2 | 6 | 2 | 1 | 134 | 15.02 | 4.97 | 0.824 |
| 7.69230769 | 1 | 2 | 1 | 1 | 195 | 21.658 | 4.7 | |
| 8.36501901 | 1 | 2 | 1 | 1 | 263 | 29.744 | 6.29 | |
| 13.3333333 | 2 | 3 | 1 | 1 | 165 | 18.493 | 7.85 | 0.82 |
| 35.2534562 | 12 | 61 | 11 | 1 | 434 | 47.239 | 5.03 | 0.821 |
| 10.7279693 | 2 | 3 | 2 | 1 | 261 | 28.975 | 7.31 | 1.032 |
| 9.49554896 | 3 | 6 | 3 | 1 | 337 | 37.516 | 6.81 | 0.744 |
| 43.1372549 | 12 | 27 | 10 | 1 | 255 | 29.155 | 4.74 | 0.654 |
| 24.0816327 | 6 | 23 | 3 | 1 | 245 | 27.747 | 4.78 | 0.823 |
| 13.6986301 | 3 | 12 | 3 | 1 | 219 | 23.679 | 9.44 | 1.011 |
| 8.06451613 | 2 | 4 | 2 | 1 | 248 | 27.728 | 10.36 | 1.04 |
| 16.8067227 | 1 | 3 | 1 | 1 | 119 | 13.273 | 11.56 | |
| 15.942029 | 1 | 1 | 1 | 1 | 69 | 7.928 | 9.35 | 0.742 |
| 33.7468983 | 20 | 54 | 20 | 1 | 806 | 89.266 | 5.26 | 0.803 |
| 10.483871 | 3 | 10 | 3 | 1 | 248 | 27.06 | 4.51 | 0.739 |
| 4.36681223 | 2 | 4 | 2 | 1 | 229 | 25.175 | 4.64 | 0.893 |
| 11.878453 | 5 | 9 | 5 | 1 | 362 | 40.069 | 9.38 | 0.7 |
| 17.3033708 | 7 | 27 | 1 | 1 | 445 | 50.631 | 6.47 | |
| 28.9827255 | 14 | 40 | 8 | 1 | 521 | 58.65 | 5.86 | 0.756 |
| 8.87850467 | 2 | 4 | 2 | 1 | 214 | 22.012 | 8.57 | 0.757 |
| 4.73372781 | 1 | 2 | 1 | 1 | 338 | 38.25 | 5.53 | |
| 15.4385965 | 4 | 8 | 4 | 1 | 285 | 31.102 | 5.81 | 0.749 |
| 5.1420839 | 3 | 6 | 3 | 1 | 739 | 83.302 | 6.7 | 0.78 |
| 5.41871921 | 1 | 2 | 1 | 1 | 203 | 23.562 | 10.93 | 0.76 |
| 5.27638191 | 2 | 4 | 2 | 1 | 398 | 44.36 | 7.99 | 0.719 |
| 5.7591623 | 1 | 2 | 1 | 1 | 191 | 21.855 | 5.4 | |
| 5 | 1 | 4 | 1 | 1 | 260 | 27.356 | 6.62 | 1.286 |
| 5.71428571 | 3 | 6 | 3 | 1 | 630 | 70.766 | 5.6 | |

| | | | | | | | | |
|---|---|---|---|---|---|---|---|---|
| 10.6930693 | 3 | 6 | 3 | 1 | 505 | 55.175 | 7.23 | 0.851 |
| 7.28971963 | 2 | 6 | 2 | 1 | 535 | 60.849 | 6.57 | 0.943 |
| 2.77777778 | 1 | 2 | 1 | 1 | 504 | 55.146 | 6.61 | |
| 27.7486911 | 3 | 7 | 2 | 1 | 191 | 22.188 | 4.89 | 0.608 |
| 11.4035088 | 3 | 8 | 3 | 1 | 228 | 25.838 | 7.97 | 1.109 |
| 26.8656716 | 1 | 2 | 1 | 1 | 67 | 7.944 | 5.11 | |
| 7.69230769 | 1 | 2 | 1 | 1 | 247 | 28.245 | 9.36 | |
| 3.04709141 | 1 | 1 | 1 | 1 | 361 | 40.658 | 8.59 | |
| 4 | 1 | 2 | 1 | 1 | 450 | 50.821 | 7.43 | 0.64 |
| 13.8888889 | 1 | 2 | 1 | 1 | 108 | 12.191 | 9.09 | |
| 11.7161716 | 6 | 12 | 6 | 1 | 606 | 66.152 | 6.65 | 0.936 |
| 10.9004739 | 3 | 8 | 3 | 1 | 211 | 23.452 | 8.63 | 0.752 |
| 2.14285714 | 2 | 4 | 2 | 1 | 560 | 61.573 | 7.47 | 0.809 |
| 15.1761518 | 5 | 12 | 2 | 1 | 369 | 42.75 | 6.67 | |
| 4.62962963 | 2 | 4 | 1 | 1 | 432 | 47.685 | 6.34 | |
| 7.89473684 | 2 | 6 | 1 | 1 | 418 | 47.577 | 5.88 | |
| 12.4137931 | 1 | 2 | 1 | 1 | 145 | 17.104 | 9.63 | |
| 31.6384181 | 11 | 36 | 10 | 1 | 354 | 40.025 | 5.53 | 0.798 |
| 26.1904762 | 3 | 12 | 1 | 1 | 126 | 13.942 | 10.32 | 0.778 |
| 16.7259786 | 3 | 9 | 3 | 1 | 281 | 31.962 | 5.54 | 0.712 |
| 3.7414966 | 1 | 2 | 1 | 1 | 294 | 32.555 | 4.78 | |
| 29.5454545 | 1 | 2 | 1 | 1 | 44 | 5.023 | 5.36 | 0.767 |
| 39.3939394 | 9 | 42 | 9 | 1 | 165 | 18.001 | 7.81 | 0.931 |
| 10.6796117 | 2 | 4 | 2 | 1 | 206 | 22.963 | 5.24 | |
| 14.7368421 | 1 | 2 | 1 | 1 | 95 | 10.493 | 8.18 | |
| 17.6470588 | 4 | 8 | 4 | 1 | 289 | 32.639 | 5.86 | 1.015 |
| 4.16666667 | 1 | 2 | 1 | 1 | 432 | 46.991 | 5.38 | 0.859 |
| 43.1818182 | 1 | 2 | 1 | 1 | 44 | 5.05 | 5.06 | 0.854 |
| 20 | 4 | 12 | 2 | 1 | 105 | 11.741 | 9.57 | |
| 5.11182109 | 2 | 4 | 2 | 1 | 626 | 69.025 | 5.03 | 0.82 |
| 5.83153348 | 3 | 6 | 3 | 1 | 463 | 50.285 | 7.42 | 0.698 |
| 5.80270793 | 2 | 4 | 2 | 1 | 517 | 56.104 | 7.14 | 0.763 |
| 16.25 | 1 | 2 | 1 | 1 | 80 | 8.518 | 8.29 | 1.043 |
| 7.35455543 | 6 | 19 | 3 | 1 | 911 | 104.788 | 5.44 | 0.955 |
| 6.82565789 | 6 | 12 | 6 | 1 | 1216 | 138.48 | 6.23 | 0.888 |
| 3.3203125 | 1 | 2 | 1 | 1 | 512 | 59.112 | 9.36 | |
| 2.73865415 | 2 | 4 | 2 | 1 | 1278 | 141.607 | 5.81 | |
| 8.23970037 | 1 | 2 | 1 | 1 | 267 | 30.204 | 8 | |
| 20.4301075 | 1 | 2 | 1 | 1 | 93 | 10.431 | 4.93 | |
| 6.42673522 | 2 | 4 | 2 | 1 | 389 | 43.154 | 5.36 | 0.62 |
| 10.7648725 | 3 | 5 | 3 | 1 | 353 | 37.407 | 8.95 | 0.672 |
| 6.1971831 | 2 | 4 | 2 | 1 | 355 | 38.41 | 7.81 | 0.676 |
| 1.50684932 | 1 | 2 | 1 | 1 | 730 | 77.464 | 8.7 | |
| 12.0218579 | 2 | 4 | 2 | 1 | 366 | 38.627 | 6.79 | 0.875 |
| 11.4224138 | 3 | 6 | 3 | 1 | 464 | 50.996 | 5.33 | 0.859 |
| 28.4753363 | 16 | 102 | 3 | 1 | 446 | 49.825 | 4.88 | 0.992 |
| 21.4428858 | 12 | 32 | 10 | 1 | 499 | 55.357 | 5.4 | 0.732 |
| 6.06060606 | 1 | 2 | 1 | 1 | 132 | 14.505 | 7.21 | |
| 2.35690236 | 1 | 2 | 1 | 1 | 297 | 34.341 | 9.72 | 0.818 |
| 13.0434783 | 1 | 2 | 1 | 1 | 115 | 13.007 | 11 | |
| 4.09556314 | 1 | 2 | 1 | 1 | 293 | 31.305 | 10.24 | 0.895 |
| 7.10382514 | 1 | 2 | 1 | 1 | 183 | 20.798 | 5.33 | |
| 6.73076923 | 2 | 4 | 2 | 1 | 312 | 33.22 | 5.43 | |
| 12.4423963 | 2 | 4 | 2 | 1 | 217 | 24.816 | 9.94 | |
| 12.9213483 | 2 | 4 | 2 | 1 | 178 | 20.24 | 9.6 | 0.805 |
| 20.5882353 | 3 | 5 | 3 | 1 | 204 | 24.131 | 11.62 | 0.637 |
| 5.21327014 | 1 | 2 | 1 | 1 | 211 | 24.247 | 11.65 | 0.679 |
| 13.3064516 | 2 | 6 | 2 | 1 | 248 | 29.207 | 10.65 | 0.794 |
| 28.4615385 | 3 | 5 | 1 | 1 | 130 | 14.113 | 11.05 | |
| 4.72972973 | 1 | 2 | 1 | 1 | 296 | 33.307 | 4.37 | |
| 16.7539267 | 2 | 8 | 2 | 1 | 191 | 21.297 | 6.04 | 0.908 |
| 31.9444444 | 11 | 21 | 11 | 1 | 288 | 33.224 | 5.38 | 0.792 |

| | | | | | | | | |
|---|---|---|---|---|---|---|---|---|
| 6.25 | 1 | 2 | 1 | 1 | 176 | 19.113 | 6.24 | 0.641 |
| 29.3577982 | 17 | 38 | 15 | 1 | 654 | 72.288 | 5.16 | 1.077 |
| 6.79156909 | 2 | 4 | 2 | 1 | 427 | 47.667 | 11.06 | 0.785 |
| 17.1428571 | 2 | 6 | 2 | 1 | 140 | 15.036 | 6.99 | 1.123 |
| 3.78548896 | 1 | 2 | 1 | 1 | 317 | 34.252 | 5.97 | |
| 14.1025641 | 3 | 6 | 3 | 1 | 156 | 17.684 | 10.45 | 0.813 |
| 25 | 2 | 3 | 2 | 1 | 108 | 11.943 | 8.16 | 0.824 |
| 22.5352113 | 2 | 4 | 2 | 1 | 142 | 15.639 | 9.13 | |
| 4.01606426 | 1 | 2 | 1 | 1 | 249 | 28.663 | 10.84 | 1.058 |
| 41.5178571 | 20 | 111 | 8 | 1 | 448 | 49.892 | 5.06 | 0.818 |
| 29.1390728 | 3 | 6 | 3 | 1 | 151 | 16.919 | 4.65 | 0.743 |
| 47.6718404 | 23 | 159 | 11 | 1 | 451 | 50.104 | 5.06 | 0.98 |
| 26.6055046 | 1 | 4 | 1 | 1 | 109 | 12.147 | 9.32 | 0.987 |
| 3.64321608 | 2 | 4 | 2 | 1 | 796 | 91.649 | 5.49 | |
| 1.25786164 | 2 | 4 | 2 | 1 | 1431 | 149.722 | 9.14 | |
| 13.4831461 | 2 | 6 | 2 | 1 | 89 | 10.343 | 7.37 | 0.55 |
| 4.375 | 1 | 2 | 1 | 1 | 160 | 18.553 | 10.49 | |
| 4.07124682 | 1 | 2 | 1 | 1 | 393 | 41.893 | 6.29 | 0.832 |
| 30.5084746 | 3 | 6 | 3 | 1 | 118 | 13.596 | 10.26 | 1.089 |
| 37.3786408 | 8 | 17 | 8 | 1 | 206 | 23.321 | 4.86 | 0.939 |
| 2.42105263 | 1 | 2 | 1 | 1 | 950 | 108.401 | 6.38 | |
| 10.3921569 | 3 | 8 | 3 | 1 | 510 | 54.602 | 8.76 | |
| 16.2743091 | 10 | 25 | 7 | 1 | 977 | 107.478 | 7.03 | 0.795 |
| 25.2836305 | 10 | 26 | 10 | 1 | 617 | 68.26 | 5.52 | 0.901 |
| 30.5283757 | 12 | 47 | 12 | 1 | 511 | 56.465 | 5.81 | 0.92 |
| 12.8205128 | 6 | 12 | 6 | 1 | 351 | 40.303 | 5 | 0.826 |
| 2.62557078 | 2 | 8 | 2 | 1 | 876 | 97.108 | 4.78 | |
| 0.55555556 | 1 | 2 | 1 | 1 | 1980 | 218.267 | 6.39 | 0.491 |
| 8.67208672 | 4 | 8 | 4 | 1 | 369 | 41.305 | 5.27 | 1.003 |
| 1.46252285 | 1 | 2 | 1 | 1 | 547 | 58.956 | 6.89 | 0.877 |
| 3.40501792 | 1 | 2 | 1 | 1 | 558 | 63.503 | 6.18 | |
| 9.76744186 | 3 | 12 | 1 | 1 | 215 | 23.882 | 6.21 | |
| 4.71698113 | 2 | 3 | 2 | 1 | 530 | 60.067 | 9.13 | |
| 1.76600442 | 1 | 2 | 1 | 1 | 453 | 48.724 | 8.95 | 0.679 |
| 1.77304965 | 1 | 2 | 1 | 1 | 564 | 61.382 | 7.2 | |
| 34.3262411 | 16 | 42 | 14 | 1 | 705 | 74.066 | 9.83 | 0.882 |
| 7.04467354 | 4 | 10 | 2 | 1 | 582 | 62.957 | 8.41 | 0.702 |
| 4.33070866 | 2 | 4 | 2 | 1 | 508 | 57.081 | 4.87 | |
| 4.90341753 | 3 | 6 | 3 | 1 | 673 | 76.962 | 7.02 | 0.855 |
| 6.25 | 3 | 6 | 3 | 1 | 464 | 50.785 | 8.21 | 0.75 |
| 31.4917127 | 24 | 62 | 13 | 1 | 724 | 83.212 | 5.03 | 0.792 |
| 14.1666667 | 3 | 6 | 3 | 1 | 240 | 26.68 | 5.2 | 0.883 |
| 9.13978495 | 1 | 2 | 1 | 1 | 186 | 21.402 | 7.77 | |
| 11.1111111 | 1 | 2 | 1 | 1 | 189 | 21.084 | 5.8 | |
| 36.32287 | 7 | 19 | 7 | 1 | 223 | 24.808 | 5.48 | 0.693 |
| 18.4365782 | 9 | 18 | 9 | 1 | 678 | 74.715 | 8.38 | 0.826 |
| 5.0955414 | 1 | 4 | 1 | 1 | 314 | 34.04 | 9.91 | |
| 4.59081836 | 2 | 4 | 1 | 1 | 501 | 54.827 | 6.73 | 0.974 |
| 27.2727273 | 27 | 79 | 10 | 1 | 1023 | 112.824 | 5.49 | 0.726 |
| 2.32018561 | 1 | 2 | 1 | 1 | 431 | 49.192 | 5.58 | |
| 8.54092527 | 7 | 16 | 3 | 1 | 843 | 96.635 | 6.86 | 0.883 |
| 14.770798 | 5 | 12 | 5 | 1 | 589 | 65.267 | 5.11 | 0.779 |
| 24.0837696 | 4 | 8 | 3 | 1 | 191 | 22.128 | 5.15 | 0.783 |
| 5.10948905 | 1 | 2 | 1 | 1 | 274 | 30.262 | 5.97 | |
| 6.22641509 | 3 | 6 | 2 | 1 | 530 | 58.913 | 6.89 | 0.995 |
| 10.4950495 | 5 | 10 | 5 | 1 | 505 | 56.747 | 6.35 | 0.878 |
| 4.95934959 | 3 | 6 | 3 | 1 | 1230 | 136.289 | 5.78 | 0.797 |
| 5.00521376 | 2 | 3 | 2 | 1 | 959 | 106.422 | 6.39 | 0.843 |
| 38.4615385 | 11 | 32 | 6 | 1 | 247 | 28.285 | 4.89 | 0.799 |
| 27.5590551 | 10 | 72 | 10 | 1 | 381 | 42.617 | 5.59 | 0.722 |
| 5.63380282 | 1 | 2 | 1 | 1 | 142 | 15.935 | 5.67 | |
| 2.31958763 | 1 | 2 | 1 | 1 | 388 | 42.759 | 6.3 | |

| | | | | | | | | |
|---|---|---|---|---|---|---|---|---|
| 4.52261307 | 1 | 2 | 1 | 1 | 199 | 22.471 | 8.76 | 0.824 |
| 7.35294118 | 1 | 2 | 1 | 1 | 204 | 22.949 | 4.53 | |
| 31.6568047 | 13 | 44 | 13 | 1 | 338 | 35.481 | 8.68 | 0.804 |
| 5.15021459 | 1 | 2 | 1 | 1 | 233 | 25.439 | 4.68 | |
| 5.67567568 | 2 | 4 | 2 | 1 | 370 | 41.543 | 8.18 | |
| 3.66972477 | 1 | 2 | 1 | 1 | 218 | 25.695 | 6.7 | |
| 7.06319703 | 2 | 4 | 2 | 1 | 269 | 30.22 | 6.99 | 0.87 |
| 12.8630705 | 2 | 4 | 2 | 1 | 241 | 26.394 | 4.79 | 0.877 |
| 3.65853659 | 1 | 2 | 1 | 1 | 246 | 27.382 | 6.76 | |
| 3.72093023 | 1 | 2 | 1 | 1 | 215 | 24.926 | 5.55 | 0.79 |
| 15.7584683 | 9 | 22 | 9 | 1 | 679 | 73.635 | 6.16 | 0.852 |
| 4.1025641 | 1 | 2 | 1 | 1 | 390 | 43.035 | 5.26 | 0.671 |
| 12.3844732 | 4 | 8 | 4 | 1 | 541 | 59.633 | 5.66 | 0.527 |
| 5.62414266 | 3 | 8 | 3 | 1 | 729 | 78.814 | 5.48 | 0.792 |
| 6.26780627 | 2 | 4 | 2 | 1 | 351 | 38.394 | 7.02 | 0.798 |
| 22.5609756 | 2 | 4 | 2 | 1 | 164 | 18.637 | 7.94 | 0.851 |
| 7.6372315 | 2 | 4 | 2 | 1 | 419 | 45.87 | 5.06 | 0.941 |
| 2.04545455 | 1 | 2 | 1 | 1 | 440 | 49.154 | 6.21 | |
| 5.17241379 | 1 | 2 | 1 | 1 | 406 | 45.597 | 7.55 | 0.849 |
| 7.10172745 | 5 | 10 | 5 | 1 | 1042 | 114.683 | 5.34 | 0.878 |
| 1.57397692 | 1 | 2 | 1 | 1 | 953 | 105.769 | 5.39 | |
| 8.58895706 | 1 | 2 | 1 | 1 | 163 | 18.646 | 4.54 | |
| 5.99613153 | 3 | 6 | 2 | 1 | 517 | 56.346 | 7.05 | 0.709 |
| 3.73001776 | 1 | 4 | 1 | 1 | 563 | 61.681 | 8.07 | |
| 34.1004184 | 19 | 56 | 13 | 1 | 478 | 54.054 | 7.08 | 0.532 |
| 3.76506024 | 2 | 4 | 2 | 1 | 664 | 74.095 | 7.02 | |
| 3.19488818 | 1 | 2 | 1 | 1 | 313 | 33.823 | 4.81 | |
| 11.9469027 | 4 | 8 | 4 | 1 | 452 | 50.877 | 8.69 | 0.869 |
| 6.80272109 | 1 | 7 | 1 | 1 | 147 | 16.118 | 7.2 | 1.949 |
| 4.19161677 | 1 | 2 | 1 | 1 | 334 | 37.896 | 7.39 | 0.817 |
| 57.9775281 | 28 | 259 | 2 | 1 | 445 | 49.921 | 4.89 | 1.025 |
| 64.1891892 | 27 | 251 | 5 | 1 | 444 | 49.639 | 4.89 | 0.921 |
| 5.72916667 | 4 | 8 | 3 | 1 | 768 | 84.25 | 5.96 | 0.845 |
| 0.93457944 | 1 | 2 | 1 | 1 | 963 | 104.833 | 7.74 | |
| 1.32450331 | 1 | 2 | 1 | 1 | 755 | 83.899 | 5.01 | |
| 9.65435042 | 6 | 12 | 6 | 1 | 839 | 94.453 | 5.88 | 0.976 |
| 9.45454545 | 2 | 4 | 2 | 1 | 275 | 29.719 | 6.8 | |
| 39.0924956 | 21 | 61 | 21 | 1 | 573 | 61.016 | 5.87 | 0.884 |
| 2.76923077 | 1 | 2 | 1 | 1 | 325 | 35.956 | 9.09 | 0.897 |
| 9.89180835 | 5 | 10 | 5 | 1 | 647 | 68.953 | 7.84 | 0.786 |
| 1.3400335 | 1 | 4 | 1 | 1 | 1194 | 133.414 | 5.68 | |
| 69.9432892 | 25 | 141 | 25 | 1 | 529 | 56.525 | 5.4 | 0.855 |
| 0.95602294 | 1 | 2 | 1 | 1 | 1046 | 116.85 | 6.7 | |
| 3.6259542 | 2 | 4 | 1 | 1 | 524 | 58.006 | 5.96 | |
| 7.64705882 | 7 | 16 | 6 | 1 | 1020 | 111.771 | 6.11 | 0.922 |
| 15.5263158 | 4 | 12 | 4 | 1 | 380 | 41.769 | 8.54 | 0.798 |
| 0.70824766 | 3 | 5 | 1 | 1 | 4377 | 480.113 | 6.49 | |
| 9.3844167 | 27 | 54 | 27 | 1 | 4646 | 532.072 | 6.4 | 0.828 |
| 14.2045455 | 1 | 2 | 1 | 1 | 176 | 20.096 | 6.39 | |
| 12.7118644 | 3 | 10 | 2 | 1 | 354 | 40.335 | 5.97 | 0.782 |
| 12.6213592 | 2 | 4 | 2 | 1 | 206 | 23.552 | 7.11 | 0.749 |
| 36.2318841 | 5 | 10 | 5 | 1 | 207 | 23.475 | 6.7 | |
| 9.67032967 | 4 | 8 | 4 | 1 | 455 | 49.843 | 7.61 | 0.703 |
| 8.24261275 | 4 | 8 | 1 | 1 | 643 | 70.984 | 6.14 | 1.102 |
| 2.03327172 | 2 | 4 | 2 | 1 | 1082 | 118.985 | 5.59 | 0.746 |
| 7.65027322 | 1 | 2 | 1 | 1 | 183 | 20.491 | 4.81 | |
| 17.8571429 | 11 | 22 | 11 | 1 | 840 | 94.271 | 5.19 | 0.77 |
| 10.0858369 | 5 | 10 | 5 | 1 | 466 | 51.68 | 8.32 | 0.851 |
| 10.5751391 | 4 | 7 | 4 | 1 | 539 | 57.888 | 7.83 | 1.073 |
| 12.5348189 | 3 | 6 | 2 | 1 | 359 | 42.115 | 5.68 | 0.673 |
| 3.72492837 | 1 | 2 | 1 | 1 | 349 | 38.928 | 4.37 | 0.812 |
| 2.82861897 | 1 | 2 | 1 | 1 | 601 | 63.812 | 5.22 | 0.904 |

| | | | | | | | | |
|---|---|---|---|---|---|---|---|---|
| 2.7027027 | 1 | 2 | 1 | 1 | 370 | 42.182 | 6.61 | |
| 5.26315789 | 1 | 2 | 1 | 1 | 494 | 54.632 | 7.23 | 0.973 |
| 4.60526316 | 1 | 2 | 1 | 1 | 304 | 33.81 | 5.55 | |
| 5.20231214 | 3 | 8 | 3 | 1 | 519 | 56.131 | 7.93 | 0.863 |
| 18.1651376 | 7 | 15 | 4 | 1 | 545 | 60.609 | 5.76 | |
| 4.62962963 | 1 | 2 | 1 | 1 | 432 | 47.599 | 5.97 | |
| 2.48833593 | 1 | 2 | 1 | 1 | 643 | 73.335 | 7.4 | |
| 4.69208211 | 1 | 2 | 1 | 1 | 341 | 37.647 | 8.56 | |
| 4.80769231 | 1 | 2 | 1 | 1 | 208 | 23.561 | 8.91 | |
| 19.895288 | 2 | 4 | 2 | 1 | 191 | 21.295 | 8.12 | 0.838 |
| 19.7080292 | 2 | 4 | 2 | 1 | 137 | 15.272 | 6.04 | 0.902 |
| 6.25 | 2 | 8 | 2 | 1 | 480 | 52.612 | 6.37 | 0.762 |
| 17.7033493 | 6 | 18 | 6 | 1 | 418 | 46.273 | 4.92 | 0.763 |
| 6.00706714 | 1 | 2 | 1 | 1 | 283 | 31.216 | 7.18 | |
| 18.2194617 | 6 | 14 | 6 | 1 | 483 | 55.847 | 6.48 | 0.811 |
| 8.04505229 | 8 | 16 | 6 | 1 | 1243 | 136.789 | 5.91 | 0.8 |
| 9.84251969 | 4 | 8 | 4 | 1 | 508 | 56.905 | 8.15 | 1.066 |
| 1.47601476 | 1 | 2 | 1 | 1 | 813 | 86.452 | 9.16 | 0.972 |
| 45.1612903 | 3 | 9 | 3 | 1 | 62 | 6.787 | 6.71 | 0.95 |
| 12.9562044 | 4 | 8 | 4 | 1 | 548 | 59.583 | 5.6 | 0.809 |
| 10 | 2 | 4 | 2 | 1 | 290 | 33.345 | 8.31 | 1.026 |
| 6.6252588 | 2 | 6 | 2 | 1 | 483 | 50.949 | 8.75 | 0.957 |
| 6.40243902 | 2 | 6 | 2 | 1 | 328 | 36.196 | 6.09 | |
| 49.3670886 | 32 | 130 | 32 | 1 | 553 | 59.714 | 9.13 | 0.863 |
| 3.3557047 | 1 | 2 | 1 | 1 | 298 | 32.975 | 9.22 | |
| 5.35714286 | 1 | 2 | 1 | 1 | 168 | 17.479 | 5.49 | 0.825 |
| 6.6091954 | 2 | 4 | 2 | 1 | 348 | 38.767 | 4.91 | |
| 2.20750552 | 1 | 2 | 1 | 1 | 453 | 48.413 | 8.63 | 0.774 |
| 10.4761905 | 1 | 2 | 1 | 1 | 105 | 11.73 | 4.92 | |
| 59.7777778 | 27 | 194 | 10 | 1 | 450 | 50.4 | 4.93 | 0.766 |
| 9.21985816 | 1 | 2 | 1 | 1 | 141 | 14.997 | 7.21 | |
| 5.19480519 | 1 | 2 | 1 | 1 | 154 | 17.859 | 10.14 | |
| 3.18181818 | 1 | 2 | 1 | 1 | 440 | 49.576 | 7.94 | |
| 3.87409201 | 1 | 2 | 1 | 1 | 413 | 44.621 | 4.84 | 0.789 |
| 10.3565365 | 3 | 6 | 3 | 1 | 589 | 64.092 | 8.22 | 0.77 |
| 2.74599542 | 1 | 2 | 1 | 1 | 437 | 47.534 | 5.33 | 0.761 |
| 20.8053691 | 8 | 20 | 3 | 1 | 298 | 33.043 | 9.76 | 0.634 |
| 4.16666667 | 1 | 2 | 1 | 1 | 432 | 47.093 | 6.55 | 1.026 |
| 2.40963855 | 1 | 2 | 1 | 1 | 332 | 35.945 | 6.95 | 0.855 |
| 30.1507538 | 4 | 10 | 4 | 1 | 199 | 22.458 | 7.33 | 0.919 |
| 15.3409091 | 2 | 4 | 2 | 1 | 176 | 18.974 | 9.13 | 1.367 |
| 4.60358056 | 1 | 2 | 1 | 1 | 391 | 42.306 | 10.05 | |
| 5.26315789 | 1 | 4 | 1 | 1 | 247 | 28.175 | 8.56 | |
| 10.3148751 | 5 | 10 | 5 | 1 | 921 | 100.304 | 5.15 | 0.685 |
| 2.49307479 | 1 | 2 | 1 | 1 | 722 | 80.477 | 7.18 | 0.781 |
| 2.69784173 | 1 | 2 | 1 | 1 | 556 | 60.306 | 6.11 | 0.788 |
| 12.4277457 | 4 | 10 | 1 | 1 | 346 | 39.307 | 7.2 | |
| 6.06060606 | 2 | 6 | 2 | 1 | 396 | 44.715 | 7.81 | 0.896 |
| 3.1496063 | 1 | 2 | 1 | 1 | 508 | 55.047 | 7.06 | |
| 1.42630745 | 1 | 2 | 1 | 1 | 631 | 68.059 | 5.05 | 0.544 |
| 6.3583815 | 2 | 4 | 2 | 1 | 346 | 37.411 | 7.17 | 0.955 |
| 24.9271137 | 9 | 26 | 7 | 1 | 686 | 74.216 | 6.86 | 0.923 |
| 18.5321101 | 7 | 14 | 7 | 1 | 545 | 60.495 | 6.49 | 0.836 |
| 4.63576159 | 1 | 2 | 1 | 1 | 302 | 33.803 | 5.49 | |
| 31.5789474 | 5 | 9 | 5 | 1 | 152 | 17.287 | 8.41 | 0.841 |
| 10.2777778 | 3 | 10 | 3 | 1 | 360 | 41.363 | 6.98 | 0.967 |
| 10.3603604 | 2 | 7 | 2 | 1 | 222 | 24.735 | 8.25 | 0.661 |
| 13.9072848 | 1 | 2 | 1 | 1 | 151 | 16.263 | 10.05 | |
| 2.22984563 | 1 | 2 | 1 | 1 | 583 | 65.061 | 6.25 | |
| 4.296875 | 1 | 2 | 1 | 1 | 256 | 28.512 | 8.98 | |
| 5.88235294 | 1 | 2 | 1 | 1 | 374 | 39.698 | 7.49 | 0.757 |
| 1.09090909 | 1 | 2 | 1 | 1 | 825 | 91.664 | 6.8 | |

| | | | | | | | | |
|---|---|---|---|---|---|---|---|---|
| 4.01188707 | 3 | 6 | 3 | 1 | 673 | 75.826 | 5.6 | |
| 8.82352941 | 1 | 4 | 1 | 1 | 238 | 27.35 | 11.82 | |
| 14.4186047 | 8 | 22 | 8 | 1 | 430 | 47.487 | 9.01 | 0.696 |
| 2.42424242 | 2 | 4 | 2 | 1 | 825 | 90.528 | 6 | |
| 3.21715818 | 1 | 4 | 1 | 1 | 373 | 42.037 | 6.89 | 0.906 |
| 1.5918958 | 1 | 2 | 1 | 1 | 691 | 79.496 | 6.9 | 0.889 |
| 3.33333333 | 1 | 2 | 1 | 1 | 240 | 27.112 | 8.28 | 0.95 |
| 5.58139535 | 1 | 2 | 1 | 1 | 215 | 23.417 | 10.93 | |
| 0.78678206 | 1 | 2 | 1 | 1 | 1271 | 142.73 | 7.03 | 0.8 |
| 13.4831461 | 1 | 2 | 1 | 1 | 89 | 10.359 | 7.4 | 0.517 |
| 10.8108108 | 3 | 6 | 3 | 1 | 407 | 46.373 | 5.48 | 0.856 |
| 4.52173913 | 2 | 4 | 2 | 1 | 575 | 63.801 | 5.87 | |
| 5.89390963 | 2 | 8 | 2 | 1 | 509 | 54.143 | 7.85 | 0.88 |
| 0.22924096 | 1 | 2 | 1 | 1 | 3926 | 416.214 | 7.55 | 0.83 |
| 1.12311015 | 3 | 6 | 3 | 1 | 2315 | 254.429 | 4.88 | 0.805 |
| 5.19480519 | 1 | 2 | 1 | 1 | 308 | 33.784 | 8.12 | 1.133 |
| 8.50340136 | 2 | 4 | 2 | 1 | 294 | 32.593 | 8.91 | 0.898 |
| 15.6769596 | 11 | 32 | 11 | 1 | 421 | 47.549 | 9.07 | 0.84 |
| 3.17516946 | 6 | 16 | 5 | 1 | 2803 | 305.298 | 4.92 | 0.831 |
| 19.2307692 | 4 | 10 | 3 | 1 | 208 | 23.447 | 5.53 | 0.688 |
| 9.13242009 | 2 | 4 | 2 | 1 | 219 | 24.96 | 7.44 | 0.895 |
| 13.2743363 | 1 | 2 | 1 | 1 | 113 | 12.544 | 10.18 | 1.288 |
| 0.71868583 | 1 | 2 | 1 | 1 | 1948 | 216.905 | 6.46 | |
| 35.174954 | 18 | 45 | 17 | 1 | 543 | 61.479 | 4.65 | 0.742 |
| 16.2361624 | 3 | 6 | 3 | 1 | 271 | 31.264 | 4.94 | 0.832 |
| 0.82644628 | 1 | 2 | 1 | 1 | 968 | 106.743 | 5.53 | |
| 3.25047801 | 1 | 2 | 1 | 1 | 523 | 58.706 | 6.67 | 0.993 |
| 4.33526012 | 1 | 2 | 1 | 1 | 346 | 36.227 | 8.79 | |
| 27.8341794 | 55 | 128 | 52 | 1 | 2364 | 274.439 | 5.57 | 0.891 |
| 41.1290323 | 11 | 21 | 7 | 1 | 248 | 28.853 | 4.75 | 0.876 |
| 3.04311074 | 3 | 5 | 3 | 1 | 1183 | 127.027 | 9.38 | 0.757 |
| 11.4285714 | 3 | 6 | 3 | 1 | 280 | 31.609 | 8.76 | 1.27 |
| 8.91364903 | 2 | 4 | 1 | 1 | 359 | 42.097 | 5.69 | 0.976 |
| 7.98816568 | 2 | 4 | 2 | 1 | 338 | 37.896 | 7.42 | 0.833 |
| 4.05014465 | 4 | 12 | 3 | 1 | 1037 | 110.956 | 5.03 | 0.898 |
| 10.8695652 | 1 | 2 | 1 | 1 | 184 | 20.764 | 5.31 | 1.208 |
| 1.665405 | 2 | 4 | 2 | 1 | 1321 | 143.003 | 5.38 | 0.994 |
| 0.73664825 | 1 | 2 | 1 | 1 | 1086 | 113.823 | 8.09 | |
| 4.80769231 | 1 | 2 | 1 | 1 | 208 | 23.831 | 8.79 | |
| 1.53649168 | 1 | 2 | 1 | 1 | 781 | 85.442 | 5.86 | 0.921 |
| 1.66493236 | 1 | 2 | 1 | 1 | 961 | 103.821 | 6.51 | 0.774 |
| 2.28494624 | 1 | 2 | 1 | 1 | 744 | 84.037 | 6.37 | |
| 10.7226107 | 6 | 12 | 6 | 1 | 858 | 95.725 | 5.03 | 0.707 |
| 4.28571429 | 1 | 2 | 1 | 1 | 350 | 38.414 | 5.12 | |
| 6.66666667 | 2 | 4 | 2 | 1 | 330 | 37.548 | 7.01 | |
| 2.10843373 | 1 | 2 | 1 | 1 | 332 | 31.536 | 4.45 | 0.804 |
| 1.31291028 | 1 | 2 | 1 | 1 | 914 | 105.407 | 5.64 | |
| 20.4918033 | 19 | 51 | 10 | 1 | 854 | 98.099 | 5.16 | 1.041 |
| 2.8363047 | 6 | 14 | 5 | 1 | 2468 | 270.468 | 4.81 | 0.893 |
| 18.8191882 | 5 | 11 | 4 | 1 | 271 | 31.52 | 5.15 | 0.788 |
| 7.51978892 | 6 | 11 | 6 | 1 | 758 | 83.626 | 6.48 | 0.689 |
| 1.97368421 | 1 | 2 | 1 | 1 | 608 | 66.346 | 5.31 | 0.91 |
| 26.1744966 | 10 | 36 | 5 | 1 | 298 | 32.831 | 9.69 | 0.661 |
| 11.3602392 | 5 | 10 | 5 | 1 | 669 | 73.414 | 7.77 | 0.937 |
| 6.37681159 | 1 | 4 | 1 | 1 | 345 | 37.395 | 7.06 | |
| 6.39534884 | 1 | 2 | 1 | 1 | 172 | 18.974 | 5.35 | 0.741 |
| 5.98802395 | 1 | 2 | 1 | 1 | 167 | 19.446 | 7.4 | |
| 2.86144578 | 2 | 4 | 2 | 1 | 664 | 72.645 | 7.39 | |
| 22.005571 | 5 | 12 | 5 | 1 | 359 | 39.208 | 6.65 | 0.764 |
| 8.93854749 | 3 | 6 | 3 | 1 | 358 | 40.678 | 7.2 | 0.799 |
| 2.65095729 | 3 | 8 | 3 | 1 | 1358 | 149.467 | 4.82 | |
| 1.51802657 | 1 | 2 | 1 | 1 | 527 | 58.91 | 7.75 | |

| | | | | | | | | |
|---|---|---|---|---|---|---|---|---|
| 6.94736842 | 2 | 6 | 1 | 1 | 475 | 51.641 | 8.02 | 0.94 |
| 0.73917635 | 1 | 2 | 1 | 1 | 1894 | 212.318 | 6.86 | |
| 17.7947598 | 16 | 41 | 14 | 1 | 916 | 102.411 | 4.91 | 0.853 |
| 9.57562568 | 8 | 16 | 8 | 1 | 919 | 103.211 | 5.72 | 0.908 |
| 7.66423358 | 2 | 4 | 2 | 1 | 274 | 29.649 | 8.32 | 0.806 |
| 2.79642058 | 2 | 4 | 2 | 1 | 894 | 101.064 | 8.54 | 0.841 |
| 4.78011472 | 2 | 4 | 2 | 1 | 523 | 56.544 | 6.42 | 0.916 |
| 2.19298246 | 1 | 2 | 1 | 1 | 456 | 51.769 | 9.22 | |
| 7.06896552 | 4 | 10 | 1 | 1 | 580 | 64.49 | 5.53 | |
| 2.27272727 | 1 | 2 | 1 | 1 | 572 | 61.838 | 7.09 | 0.858 |
| 2.65957447 | 1 | 2 | 1 | 1 | 376 | 42.459 | 6.42 | |
| 11.0091743 | 2 | 4 | 2 | 1 | 218 | 24.473 | 5.94 | 0.894 |
| 44.6666667 | 6 | 20 | 6 | 1 | 150 | 16.752 | 6.79 | 0.955 |
| 16.2454874 | 4 | 20 | 4 | 1 | 277 | 30.057 | 8.35 | 0.663 |
| 7.74647887 | 2 | 3 | 2 | 1 | 284 | 33.289 | 9.31 | 0.87 |
| 10.1298701 | 2 | 4 | 2 | 1 | 385 | 42.44 | 8.47 | 0.907 |
| 9.65517241 | 2 | 4 | 2 | 1 | 290 | 31.367 | 8.07 | |
| 2.82258065 | 1 | 2 | 1 | 1 | 248 | 28.76 | 6.42 | |
| 4.2047532 | 3 | 6 | 3 | 1 | 1094 | 123.102 | 6.87 | |
| 4.12642669 | 4 | 8 | 3 | 1 | 1139 | 126.102 | 6.73 | 0.609 |
| 3.59281437 | 2 | 4 | 2 | 1 | 501 | 54.089 | 8.66 | |
| 4.43686007 | 1 | 2 | 1 | 1 | 293 | 31.498 | 6.32 | |
| 30.6338028 | 8 | 24 | 7 | 1 | 284 | 30.77 | 8.66 | 0.76 |
| 6.55737705 | 1 | 2 | 1 | 1 | 244 | 25.773 | 7.37 | 0.815 |
| 1.00222717 | 1 | 2 | 1 | 1 | 898 | 95.748 | 8.81 | |
| 3.57142857 | 1 | 2 | 1 | 1 | 336 | 37.016 | 8.65 | |
| 5.66037736 | 3 | 6 | 3 | 1 | 742 | 82.642 | 5.57 | 0.903 |
| 3.3557047 | 1 | 2 | 1 | 1 | 298 | 33.212 | 5.6 | |
| 10.8695652 | 2 | 4 | 2 | 1 | 184 | 20.157 | 7.01 | 0.83 |
| 4.32276657 | 2 | 3 | 2 | 1 | 694 | 76.824 | 8.34 | 0.868 |
| 21.7391304 | 5 | 12 | 5 | 1 | 299 | 33.276 | 9.83 | 1.015 |
| 8.03571429 | 2 | 4 | 2 | 1 | 336 | 36.671 | 7.91 | 0.983 |
| 3.65168539 | 1 | 2 | 1 | 1 | 356 | 37.474 | 7.09 | |
| 3.3970276 | 1 | 2 | 1 | 1 | 471 | 54.197 | 8.95 | |
| 19.5402299 | 4 | 10 | 4 | 1 | 174 | 19.811 | 7.02 | 1.081 |
| 6.22119816 | 3 | 8 | 3 | 1 | 434 | 48.683 | 4.78 | 1.188 |
| 6.99507389 | 5 | 11 | 5 | 1 | 1015 | 117.67 | 7.77 | 0.833 |
| 37.0629371 | 8 | 24 | 8 | 1 | 286 | 30.772 | 5.92 | 0.879 |
| 5.89330025 | 7 | 18 | 7 | 1 | 1612 | 177.438 | 8.22 | 0.773 |
| 2.63459336 | 2 | 4 | 2 | 1 | 873 | 96.711 | 6.52 | |
| 23.5 | 5 | 16 | 2 | 1 | 200 | 22.527 | 8.38 | 1.409 |
| 19.9445983 | 6 | 14 | 5 | 1 | 361 | 39.812 | 8.43 | 0.774 |
| 8.64067439 | 6 | 12 | 1 | 1 | 949 | 104.54 | 5.06 | 0.643 |
| 12.5 | 1 | 2 | 1 | 1 | 144 | 16.688 | 8.43 | 0.924 |
| 13.480055 | 7 | 16 | 7 | 1 | 727 | 80.802 | 7.69 | 0.883 |
| 7.09812109 | 2 | 6 | 2 | 1 | 479 | 51.624 | 6.52 | 0.746 |
| 3.96551724 | 2 | 4 | 2 | 1 | 580 | 63.75 | 5.83 | 0.794 |
| 31.7365269 | 12 | 28 | 11 | 1 | 334 | 36.615 | 6.05 | 0.97 |
| 9.19037199 | 3 | 10 | 3 | 1 | 457 | 52.966 | 6.49 | 0.705 |
| 1.59090909 | 1 | 4 | 1 | 1 | 440 | 47.702 | 6.43 | |
| 2.3255814 | 1 | 2 | 1 | 1 | 559 | 60.483 | 7.24 | |
| 4.7318612 | 2 | 4 | 2 | 1 | 634 | 70.915 | 6.64 | 0.77 |
| 15.5339806 | 5 | 10 | 5 | 1 | 309 | 33.351 | 7.59 | 0.882 |
| 7.75193798 | 1 | 2 | 1 | 1 | 129 | 14.574 | 8.88 | 0.797 |
| 0.86848635 | 1 | 2 | 1 | 1 | 806 | 90.216 | 6.43 | |
| 40.4726736 | 18 | 64 | 14 | 1 | 677 | 73.457 | 6.35 | 0.844 |
| 11.7647059 | 3 | 6 | 3 | 1 | 187 | 21.574 | 8.59 | 0.641 |
| 4.38596491 | 1 | 2 | 1 | 1 | 228 | 26.356 | 10.1 | |
| 7.25388601 | 1 | 2 | 1 | 1 | 193 | 22.014 | 6.37 | |
| 45.8666667 | 20 | 109 | 6 | 1 | 375 | 41.766 | 5.48 | 0.793 |
| 10.9311741 | 7 | 16 | 7 | 1 | 741 | 80.945 | 6.64 | 0.837 |
| 1.84210526 | 1 | 2 | 1 | 1 | 760 | 83.345 | 5.12 | |

| | | | | | | | | |
|---|---|---|---|---|---|---|---|---|
| 10.0456621 | 4 | 12 | 4 | 1 | 438 | 50.258 | 5.67 | 0.857 |
| 5.42986425 | 4 | 8 | 4 | 1 | 884 | 97.284 | 4.93 | 0.866 |
| 14.5744681 | 8 | 22 | 5 | 1 | 940 | 104.024 | 6.86 | 0.785 |
| 30.1435407 | 12 | 32 | 10 | 1 | 418 | 45.232 | 8.25 | 1.165 |
| 0.95147479 | 1 | 2 | 1 | 1 | 2102 | 236.678 | 7.06 | 0.818 |
| 5.94405594 | 1 | 2 | 1 | 1 | 286 | 30.339 | 6.73 | |
| 7.14285714 | 1 | 2 | 1 | 1 | 140 | 16.256 | 8.97 | 0.889 |
| 36.4522417 | 37 | 116 | 17 | 1 | 1026 | 112.987 | 5.38 | 0.837 |
| 2.88659794 | 2 | 4 | 2 | 1 | 970 | 105.608 | 7.47 | 0.978 |
| 9.30232558 | 1 | 2 | 1 | 1 | 215 | 24.725 | 8.51 | 0.732 |
| 1.20087336 | 1 | 2 | 1 | 1 | 916 | 102.822 | 7.55 | |
| 9.50871632 | 5 | 10 | 5 | 1 | 631 | 68.77 | 7.52 | 0.869 |
| 5.40540541 | 1 | 2 | 1 | 1 | 259 | 28.468 | 9.73 | |
| 1.25968992 | 1 | 2 | 1 | 1 | 1032 | 112.541 | 4.96 | 0.806 |
| 2.34505863 | 1 | 1 | 1 | 1 | 597 | 66.765 | 6.61 | 0.917 |
| 14.6862483 | 7 | 18 | 7 | 1 | 749 | 83.347 | 7.08 | 0.849 |
| 6.17977528 | 1 | 2 | 1 | 1 | 178 | 20.533 | 8.59 | |
| 1.96078431 | 2 | 4 | 1 | 1 | 612 | 67.598 | 5.91 | 0.721 |
| 35.6617647 | 8 | 18 | 8 | 1 | 272 | 29.786 | 5.76 | 0.965 |
| 12.1693122 | 3 | 6 | 3 | 1 | 189 | 19.878 | 6.79 | 0.879 |
| 2.40452617 | 1 | 2 | 1 | 1 | 707 | 76.102 | 9.44 | |
| 10.1947308 | 10 | 34 | 1 | 1 | 873 | 98.098 | 8.46 | |
| 4.73684211 | 1 | 2 | 1 | 1 | 380 | 41.053 | 4.81 | |
| 2.47678019 | 1 | 2 | 1 | 1 | 323 | 35.246 | 6.05 | |
| 0.98800282 | 1 | 2 | 1 | 1 | 1417 | 164.203 | 5.69 | 0.903 |
| 2.80373832 | 1 | 2 | 1 | 1 | 535 | 59.807 | 5.08 | |
| 9.86842105 | 4 | 7 | 4 | 1 | 608 | 66.586 | 5.45 | 0.861 |
| 5.16605166 | 2 | 4 | 2 | 1 | 542 | 58.741 | 6.02 | |
| 12.4338624 | 3 | 8 | 3 | 1 | 378 | 39.571 | 9.01 | |
| 6.93069307 | 3 | 5 | 3 | 1 | 404 | 45.49 | 5.07 | 0.713 |
| 3.04709141 | 1 | 2 | 1 | 1 | 361 | 39.92 | 7.64 | 1.125 |
| 16.8855535 | 9 | 23 | 3 | 1 | 533 | 59.965 | 7.27 | 0.684 |
| 0.70257611 | 1 | 2 | 1 | 1 | 1281 | 138.748 | 5.17 | |
| 19.4029851 | 3 | 7 | 3 | 1 | 134 | 15.385 | 5.08 | 0.93 |
| 24 | 5 | 17 | 3 | 1 | 150 | 16.347 | 10.32 | 0.925 |
| 4.59016393 | 2 | 6 | 1 | 1 | 610 | 67.468 | 5.88 | |
| 2.18037661 | 2 | 6 | 2 | 1 | 1009 | 115.8 | 6.25 | 0.774 |
| 7.2327044 | 4 | 8 | 4 | 1 | 636 | 70.626 | 9.5 | 1.003 |
| 0.2345916 | 1 | 2 | 1 | 1 | 4689 | 532.145 | 5.97 | |
| 48.1632653 | 15 | 42 | 13 | 1 | 245 | 27.728 | 4.79 | 0.734 |
| 33.8709677 | 30 | 87 | 1 | 1 | 868 | 97.647 | 7.34 | |
| 5.71428571 | 1 | 2 | 1 | 1 | 210 | 23.396 | 8.6 | |
| 6.01719198 | 1 | 2 | 1 | 1 | 349 | 36.75 | 9.38 | |
| 4.19161677 | 3 | 7 | 1 | 1 | 835 | 90.822 | 7.11 | 1.532 |
| 16.0404624 | 10 | 26 | 10 | 1 | 692 | 76.97 | 7.17 | 0.814 |
| 12.6041667 | 11 | 28 | 11 | 1 | 960 | 106.871 | 6.98 | 0.904 |
| 3.80794702 | 1 | 2 | 1 | 1 | 604 | 67.026 | 7.97 | |
| 5.23809524 | 1 | 2 | 1 | 1 | 210 | 23.69 | 6.34 | 0.821 |
| 8.45188285 | 14 | 28 | 11 | 1 | 2390 | 271.157 | 6.11 | 0.802 |
| 3.54223433 | 1 | 2 | 1 | 1 | 367 | 40.498 | 7.64 | 0.838 |
| 8.21917808 | 1 | 2 | 1 | 1 | 146 | 16.997 | 9.5 | |
| 7.28582866 | 7 | 14 | 5 | 1 | 1249 | 137.717 | 6 | 0.888 |
| 6.56851642 | 5 | 11 | 3 | 1 | 883 | 96.846 | 8.12 | 0.842 |
| 11.1627907 | 2 | 4 | 2 | 1 | 215 | 24.878 | 5.74 | 1.059 |
| 11.4864865 | 7 | 14 | 7 | 1 | 740 | 82.173 | 8.66 | 0.743 |
| 0.32701112 | 1 | 2 | 1 | 1 | 3058 | 341.933 | 6.34 | |
| 6.98151951 | 2 | 4 | 2 | 1 | 487 | 53.437 | 8.31 | |
| 25.8964143 | 6 | 11 | 2 | 1 | 251 | 28.806 | 4.83 | 1.24 |
| 10.1941748 | 1 | 1 | 1 | 1 | 206 | 23.012 | 4.82 | |
| 0.43019163 | 1 | 2 | 1 | 1 | 2557 | 273.039 | 5.58 | |
| 8.20895522 | 2 | 6 | 2 | 1 | 536 | 58.682 | 4.79 | |
| 42.3728814 | 4 | 12 | 4 | 1 | 118 | 12.926 | 8.13 | 0.868 |

| | | | | | | | | |
|---|---|---|---|---|---|---|---|---|
| 14.0904311 | 11 | 22 | 6 | 1 | 951 | 105.625 | 5.34 | 0.674 |
| 35.1648352 | 13 | 31 | 11 | 1 | 364 | 39.431 | 6.87 | 0.787 |
| 33.6504162 | 46 | 133 | 46 | 1 | 1682 | 192.276 | 5.69 | 0.721 |
| 6.26450116 | 6 | 12 | 6 | 1 | 862 | 89.739 | 6.65 | 0.946 |
| 5.11247444 | 1 | 2 | 1 | 1 | 489 | 58.185 | 9.92 | |
| 8.6492891 | 6 | 18 | 6 | 1 | 844 | 97.194 | 6.43 | 0.815 |
| 22.8571429 | 4 | 8 | 4 | 1 | 175 | 19.446 | 9.22 | 0.742 |
| 10.0775194 | 1 | 2 | 1 | 1 | 129 | 14.988 | 6.93 | |
| 13.2231405 | 1 | 2 | 1 | 1 | 121 | 13.354 | 10.08 | |
| 0.79197466 | 1 | 2 | 1 | 1 | 1894 | 211.044 | 7.11 | |
| 0.76687117 | 1 | 2 | 1 | 1 | 1304 | 144.36 | 6.67 | |
| 9.04392765 | 5 | 12 | 5 | 1 | 387 | 42.384 | 8.46 | 0.77 |
| 9.30232558 | 3 | 6 | 3 | 1 | 301 | 33.76 | 6.43 | 0.963 |
| 1.69082126 | 1 | 2 | 1 | 1 | 414 | 46.63 | 7.01 | |
| 4.19708029 | 1 | 2 | 1 | 1 | 548 | 61.888 | 6.71 | |
| 14.4117647 | 4 | 12 | 4 | 1 | 340 | 37.353 | 6 | 0.623 |
| 2.26804124 | 1 | 2 | 1 | 1 | 485 | 55.307 | 6.95 | 0.753 |
| 10.1694915 | 5 | 10 | 5 | 1 | 590 | 68.037 | 7.74 | 0.826 |
| 7.5203252 | 3 | 8 | 3 | 1 | 492 | 53.867 | 5.33 | 0.855 |
| 24.1721854 | 7 | 15 | 7 | 1 | 302 | 33.896 | 5.47 | 0.802 |
| 5.19480519 | 1 | 2 | 1 | 1 | 385 | 44.938 | 8.95 | |
| 1.68224299 | 1 | 2 | 1 | 1 | 535 | 58.211 | 7.23 | |
| 6.59898477 | 1 | 2 | 1 | 1 | 197 | 22.559 | 5.24 | |
| 5.69767442 | 3 | 6 | 3 | 1 | 860 | 97.398 | 5.72 | 1.026 |
| 60.5855856 | 27 | 212 | 5 | 1 | 444 | 49.554 | 4.88 | 0.772 |
| 21.1940299 | 7 | 64 | 7 | 1 | 335 | 36.03 | 8.46 | 0.91 |
| 0.54644809 | 1 | 2 | 1 | 1 | 1464 | 163.843 | 6.48 | 0.684 |
| 5.03731343 | 2 | 4 | 2 | 1 | 536 | 61.274 | 7.06 | 0.921 |
| 1.91846523 | 1 | 2 | 1 | 1 | 834 | 89.776 | 8.51 | |
| 5.57275542 | 1 | 2 | 1 | 1 | 323 | 37.289 | 6.01 | |
| 16.6666667 | 1 | 2 | 1 | 1 | 120 | 13.857 | 7.5 | 1.276 |
| 4.49172577 | 1 | 2 | 1 | 1 | 423 | 48.768 | 7.17 | 0.849 |
| 4.00696864 | 2 | 4 | 2 | 1 | 1148 | 129.473 | 6.8 | |
| 2.5 | 1 | 2 | 1 | 1 | 400 | 43.421 | 5.1 | |
| 2.96774194 | 1 | 2 | 1 | 1 | 775 | 84.878 | 7.37 | 0.576 |
| 38.1578947 | 6 | 12 | 3 | 1 | 152 | 15.964 | 4.7 | 0.859 |
| 5.14285714 | 5 | 11 | 5 | 1 | 1225 | 134.798 | 4.55 | 0.585 |
| 24.3303571 | 9 | 25 | 8 | 1 | 448 | 52.011 | 8.73 | 0.764 |
| 4.41441441 | 3 | 8 | 3 | 1 | 1110 | 125.229 | 5.36 | |
| 2.56696429 | 2 | 4 | 2 | 1 | 896 | 100.527 | 5.4 | 0.969 |
| 33.7557604 | 30 | 85 | 1 | 1 | 868 | 97.502 | 7.01 | |
| 1.89959294 | 1 | 2 | 1 | 1 | 737 | 82.883 | 8.21 | 1.155 |
| 8.17757009 | 3 | 6 | 3 | 1 | 428 | 47.239 | 5.68 | 0.945 |
| 30.9163347 | 67 | 186 | 67 | 1 | 2510 | 288.945 | 5.41 | 0.925 |
| 3.61904762 | 1 | 2 | 1 | 1 | 525 | 56.095 | 8.6 | |
| 7.72532189 | 4 | 11 | 2 | 1 | 466 | 53.619 | 5.12 | 0.905 |
| 1.41643059 | 1 | 2 | 1 | 1 | 706 | 79.105 | 6.32 | |
| 12.755102 | 7 | 17 | 1 | 1 | 588 | 65.201 | 7.94 | |
| 5.43735225 | 1 | 1 | 1 | 1 | 423 | 46.705 | 9 | |
| 4.34782609 | 1 | 2 | 1 | 1 | 253 | 28.005 | 8.81 | 0.722 |
| 29.0909091 | 17 | 63 | 17 | 1 | 605 | 65.764 | 7.99 | 0.749 |
| 6.46258503 | 1 | 2 | 1 | 1 | 294 | 32.921 | 7.88 | |
| 19.6374622 | 4 | 8 | 4 | 1 | 331 | 37.383 | 5.91 | 0.659 |
| 0.60402685 | 1 | 2 | 1 | 1 | 1490 | 163.705 | 6.92 | |
| 3.04806565 | 2 | 4 | 1 | 1 | 853 | 93.176 | 8.16 | |
| 4.94505495 | 1 | 2 | 1 | 1 | 182 | 21.203 | 9.01 | 0.754 |
| 11.2582781 | 1 | 4 | 1 | 1 | 151 | 17.296 | 4.89 | |
| 2.5974026 | 1 | 2 | 1 | 1 | 616 | 69.657 | 5.67 | |
| 3.64963504 | 1 | 2 | 1 | 1 | 548 | 59.583 | 9.23 | |
| 2.58215962 | 1 | 2 | 1 | 1 | 426 | 44.614 | 8.46 | 0.65 |
| 1.2987013 | 1 | 2 | 1 | 1 | 1155 | 126.138 | 8.18 | 0.968 |
| 5.64971751 | 1 | 2 | 1 | 1 | 177 | 20.444 | 4.75 | |

| | | | | | | | | |
|---|---|---|---|---|---|---|---|---|
| 6.54205607 | 1 | 2 | 1 | 1 | 214 | 24.561 | 10.08 | |
| 21.3333333 | 3 | 6 | 3 | 1 | 300 | 33.293 | 4.87 | 0.888 |
| 13.4328358 | 2 | 4 | 2 | 1 | 268 | 28.125 | 8.27 | 0.736 |
| 7.08401977 | 4 | 8 | 4 | 1 | 607 | 64.653 | 6.87 | 0.704 |
| 1.60427807 | 1 | 2 | 1 | 1 | 748 | 81.681 | 6.14 | |
| 4.5890411 | 5 | 10 | 5 | 1 | 1460 | 162.147 | 7.15 | 0.749 |
| 1.09769484 | 1 | 2 | 1 | 1 | 911 | 99.056 | 4.64 | |
| 0.87796313 | 1 | 2 | 1 | 1 | 1139 | 127.704 | 7.42 | |
| 5.09641873 | 3 | 6 | 2 | 1 | 726 | 80.803 | 5.92 | 0.935 |
| 1.89873418 | 1 | 2 | 1 | 1 | 474 | 51.262 | 9.41 | |
| 1.13052415 | 1 | 2 | 1 | 1 | 973 | 108.478 | 5 | 0.701 |
| 3.09859155 | 1 | 2 | 1 | 1 | 710 | 82.445 | 5.57 | |
| 5.95116989 | 8 | 16 | 8 | 1 | 1966 | 213.957 | 4.92 | 0.917 |
| 2.06489676 | 1 | 1 | 1 | 1 | 339 | 37.656 | 8.5 | |
| 3.16248637 | 1 | 2 | 1 | 1 | 917 | 97.54 | 9.38 | |
| 1.20942075 | 1 | 2 | 1 | 1 | 1571 | 171.721 | 8.47 | 1.333 |
| 5.55170021 | 4 | 10 | 1 | 1 | 1441 | 162.579 | 5.97 | 0.82 |
| 5.2887961 | 4 | 10 | 1 | 1 | 1437 | 162.241 | 5.97 | 0.909 |
| 1.08695652 | 1 | 4 | 1 | 1 | 1104 | 124.998 | 6.73 | |
| 1.39116203 | 1 | 2 | 1 | 1 | 1222 | 136.028 | 8.47 | |
| 5.54371002 | 4 | 8 | 4 | 1 | 938 | 104.494 | 5.82 | |
| 3.08370044 | 1 | 2 | 1 | 1 | 681 | 75.415 | 5.36 | |
| 2.88184438 | 1 | 2 | 1 | 1 | 347 | 38.8 | 8.19 | |
| 2.1484375 | 1 | 2 | 1 | 1 | 512 | 56.854 | 8.66 | 0.837 |
| 1.61764706 | 1 | 2 | 1 | 1 | 680 | 75.968 | 6.15 | |
| 3.94230769 | 12 | 26 | 10 | 1 | 4160 | 456.149 | 5.12 | 1.015 |
| 25.8823529 | 6 | 16 | 3 | 1 | 170 | 18.367 | 10.01 | 1.052 |
| 2.0746888 | 1 | 2 | 1 | 1 | 482 | 54.088 | 5.55 | |
| 2.25988701 | 1 | 2 | 1 | 1 | 354 | 40.14 | 7.42 | |
| 6.3583815 | 1 | 2 | 1 | 1 | 173 | 19.618 | 5.52 | |
| 7.31707317 | 3 | 6 | 3 | 1 | 697 | 76.422 | 4.58 | 0.877 |
| 7.20338983 | 2 | 6 | 2 | 1 | 472 | 51.197 | 6.8 | 0.773 |
| 3.09597523 | 1 | 2 | 1 | 1 | 646 | 73.364 | 4.65 | 0.929 |
| 13.963964 | 7 | 13 | 7 | 1 | 444 | 50.934 | 5.24 | 0.992 |
| 1.95080577 | 1 | 3 | 1 | 1 | 1179 | 130.542 | 7.44 | |
| 7.33452594 | 3 | 6 | 3 | 1 | 559 | 61.614 | 6.46 | 0.731 |
| 13.1201764 | 8 | 20 | 8 | 1 | 907 | 92.444 | 4.86 | 0.91 |
| 6.26204239 | 4 | 8 | 3 | 1 | 1038 | 117.59 | 6.92 | 0.825 |
| 2.22222222 | 1 | 2 | 1 | 1 | 540 | 61.079 | 6.18 | |
| 15.796897 | 11 | 30 | 4 | 1 | 709 | 77.208 | 7.2 | 0.47 |
| 2.21205187 | 2 | 6 | 2 | 1 | 1311 | 144.601 | 5.71 | 0.861 |
| 4.74137931 | 1 | 2 | 1 | 1 | 232 | 26.737 | 7.75 | |
| 1.57706093 | 1 | 2 | 1 | 1 | 1395 | 156.359 | 5.62 | |
| 30.8 | 5 | 10 | 5 | 1 | 250 | 27.604 | 9.07 | 0.692 |
| 2.55615802 | 2 | 4 | 2 | 1 | 1291 | 142.185 | 6.67 | 1.009 |
| 7.6754386 | 3 | 7 | 3 | 1 | 456 | 52.158 | 9.69 | 0.719 |
| 1.59010601 | 1 | 1 | 1 | 1 | 1132 | 125.82 | 7.52 | |
| 9.30232558 | 2 | 4 | 2 | 1 | 344 | 38.866 | 6.38 | 0.751 |
| 17.0984456 | 2 | 6 | 2 | 1 | 193 | 22.231 | 5.35 | 0.644 |
| 4.44874275 | 2 | 4 | 1 | 1 | 517 | 57.213 | 6.99 | |
| 2.70935961 | 1 | 2 | 1 | 1 | 406 | 45.81 | 5.67 | |
| 14.7668394 | 5 | 18 | 5 | 1 | 386 | 44.295 | 5.48 | 0.736 |
| 11.0344828 | 1 | 2 | 1 | 1 | 290 | 33.469 | 4.32 | 0.548 |
| 1.57170923 | 1 | 2 | 1 | 1 | 509 | 57.082 | 6.25 | |
| 17.9775281 | 8 | 24 | 2 | 1 | 534 | 60.187 | 6.1 | |
| 5.74257426 | 3 | 8 | 2 | 1 | 1010 | 114.409 | 6.65 | 0.778 |
| 9.71922246 | 2 | 5 | 2 | 1 | 463 | 51.535 | 5.43 | |
| 3.72881356 | 3 | 20 | 3 | 1 | 590 | 64.036 | 6.87 | 0.938 |
| 1.39968896 | 1 | 2 | 1 | 1 | 643 | 72.849 | 7.33 | |
| 3.44827586 | 3 | 6 | 3 | 1 | 1044 | 116.288 | 5.38 | |
| 43.6532508 | 30 | 116 | 26 | 1 | 646 | 70.854 | 5.52 | 0.907 |
| 1.73761946 | 2 | 4 | 2 | 1 | 1151 | 131.787 | 6.25 | |

| | | | | | | | | |
|---|---|---|---|---|---|---|---|---|
| 10.5769231 | 1 | 4 | 1 | 1 | 104 | 12.067 | 11 | 1.029 |
| 16.8958743 | 14 | 31 | 14 | 1 | 1018 | 113.249 | 5.9 | 0.833 |
| 5.45905707 | 3 | 6 | 3 | 1 | 403 | 46.803 | 4.46 | 0.829 |
| 2.98804781 | 1 | 2 | 1 | 1 | 502 | 55.259 | 6.43 | |
| 3.92156863 | 1 | 2 | 1 | 1 | 306 | 33.65 | 5.08 | |
| 2.69541779 | 1 | 2 | 1 | 1 | 371 | 40.772 | 5.21 | 0.712 |
| 12.2712594 | 10 | 24 | 7 | 1 | 929 | 106.714 | 6.05 | 0.772 |
| 1.75438596 | 1 | 2 | 1 | 1 | 684 | 77.007 | 8.25 | 0.787 |
| 4.76190476 | 5 | 10 | 5 | 1 | 1386 | 159.629 | 6.98 | 0.755 |
| 21.3429257 | 7 | 21 | 7 | 1 | 417 | 47.007 | 8.34 | 0.949 |
| 2.26190476 | 1 | 1 | 1 | 1 | 840 | 95.272 | 6.73 | |
| 3.40063762 | 6 | 16 | 6 | 1 | 1882 | 218.693 | 8.47 | 0.725 |
| 1.65217391 | 2 | 4 | 1 | 1 | 1150 | 127.534 | 7.08 | 0.749 |
| 5.2 | 2 | 6 | 2 | 1 | 500 | 56.403 | 7.96 | 0.747 |
| 10.8843537 | 1 | 2 | 1 | 1 | 147 | 15.882 | 8.21 | |
| 5.43478261 | 1 | 3 | 1 | 1 | 184 | 20.444 | 8.46 | |
| 14.3167028 | 5 | 10 | 5 | 1 | 461 | 50.994 | 6.68 | 0.747 |
| 1.88205772 | 1 | 2 | 1 | 1 | 797 | 88.51 | 6.1 | |
| 1.17820324 | 2 | 4 | 2 | 1 | 2037 | 235.705 | 5.6 | |
| 1.63551402 | 1 | 2 | 1 | 1 | 428 | 48.739 | 7.06 | |
| 4.75352113 | 1 | 2 | 1 | 1 | 568 | 62.982 | 9.13 | |
| 3.24149109 | 2 | 4 | 2 | 1 | 617 | 68.635 | 8.97 | |
| 1.8018018 | 1 | 2 | 1 | 1 | 777 | 86.156 | 7.03 | |
| 2.52427184 | 1 | 2 | 1 | 1 | 515 | 58.757 | 8.84 | |
| 32.89987 | 22 | 59 | 22 | 1 | 769 | 85.081 | 7.62 | 0.728 |
| 3.63636364 | 2 | 6 | 1 | 1 | 495 | 56.778 | 6.23 | |
| 4.72440945 | 1 | 2 | 1 | 1 | 381 | 42.955 | 5.35 | |
| 1.35440181 | 1 | 2 | 1 | 1 | 1329 | 145.107 | 5.11 | |
| 6.09480813 | 2 | 3 | 2 | 1 | 443 | 50.177 | 7.81 | |
| 0.35842294 | 1 | 2 | 1 | 1 | 2511 | 273.254 | 6.44 | |
| 15.2610442 | 3 | 5 | 3 | 1 | 249 | 27.501 | 5.01 | 0.919 |
| 4.89795918 | 4 | 8 | 4 | 1 | 1225 | 136.405 | 5.3 | 1.025 |
| 27.8287462 | 10 | 22 | 10 | 1 | 327 | 36.483 | 5.67 | 0.864 |
| 9.375 | 4 | 28 | 4 | 1 | 384 | 41.735 | 9.82 | 0.972 |
| 5.01930502 | 1 | 2 | 1 | 1 | 259 | 28.518 | 6.24 | |
| 1.39534884 | 1 | 2 | 1 | 1 | 1505 | 166.006 | 6.4 | |
| 1.2987013 | 1 | 2 | 1 | 1 | 924 | 97.056 | 5.95 | 0.923 |
| 13.9534884 | 1 | 1 | 1 | 1 | 172 | 20.313 | 7.36 | |
| 1.3400335 | 1 | 4 | 1 | 1 | 597 | 67.243 | 8.97 | 0.699 |
| 3.2 | 1 | 2 | 1 | 1 | 375 | 41.205 | 5.21 | |
| 2.58675079 | 3 | 6 | 3 | 1 | 1585 | 173.984 | 5.53 | 1.009 |
| 2.16346154 | 1 | 2 | 1 | 1 | 416 | 45.875 | 5.78 | |
| 5.12195122 | 2 | 4 | 2 | 1 | 410 | 43.958 | 9.22 | 0.777 |
| 2.23880597 | 1 | 2 | 1 | 1 | 670 | 69.974 | 9.16 | |
| 5.28735632 | 2 | 4 | 2 | 1 | 435 | 48.361 | 8.54 | 0.775 |
| 9.66850829 | 8 | 18 | 8 | 1 | 1086 | 120.85 | 5.71 | 0.838 |
| 1.14754098 | 1 | 2 | 1 | 1 | 610 | 71.32 | 6.06 | 0.687 |
| 16.539924 | 7 | 18 | 1 | 1 | 526 | 59.924 | 7.37 | 0.781 |
| 8.92857143 | 2 | 4 | 2 | 1 | 224 | 25.438 | 7.83 | |
| 6.18556701 | 1 | 2 | 1 | 1 | 194 | 22.578 | 10.65 | |
| 43.1472081 | 5 | 17 | 5 | 1 | 197 | 22.173 | 4.63 | 0.733 |
| 19.5899772 | 7 | 18 | 1 | 1 | 439 | 50.791 | 7.01 | 0.744 |
| 14.4736842 | 1 | 2 | 1 | 1 | 152 | 17.712 | 10.39 | 0.756 |
| 18.9130435 | 7 | 20 | 7 | 1 | 460 | 52.27 | 9.5 | 0.789 |
| 13.9896373 | 2 | 3 | 2 | 1 | 193 | 21.754 | 6.1 | |
| 5.49295775 | 3 | 6 | 3 | 1 | 710 | 79.787 | 7.55 | 0.834 |
| 21.529745 | 6 | 14 | 6 | 1 | 353 | 38.593 | 7.3 | 0.848 |

| Abundance Ratio: (F2, 128) / (F2, 126) | Abundance Ratio: (F2, 129) / (F2, 126) | Abundance: F2: 126, Control | Abundance: F2: 127, Sample | Abundance: F2: 128, Sample | Abundance: F2: 129, Sample | Abundances Count: F2: 126, Control | Abundances Count: F2: 127, Sample | Abundances Count: F2: 128, Sample |
|---|---|---|---|---|---|---|---|---|
| 0.876 | 0.758 | 654.6 | 556.7 | 573.6 | 496.1 | 7 | 7 | 7 |
| 0.711 | 0.768 | 204.4 | 137.1 | 145.3 | 157 | 3 | 3 | 3 |
| 2.743 | 1.278 | 43.6 | 108.6 | 119.6 | 55.7 | 1 | 1 | 1 |
| 0.821 | 0.79 | 34.7 | 29.6 | 28.5 | 27.4 | 1 | 1 | 1 |
| 0.723 | 0.608 | 133.8 | 106.3 | 96.7 | 81.4 | 1 | 1 | 1 |
| 0.678 | 0.725 | 25.5 | 15.6 | 17.3 | 18.5 | 1 | 1 | 1 |
| 0.699 | 0.819 | 86.6 | 70.4 | 60.5 | 70.9 | 1 | 1 | 1 |
| 0.861 | 0.712 | 697.1 | 603.5 | 599.9 | 496.1 | 8 | 8 | 8 |
| 0.798 | 0.694 | 875 | 697.8 | 698.2 | 606.9 | 9 | 9 | 9 |
| 0.79 | 0.758 | 183.4 | 158.5 | 144.8 | 139.1 | 2 | 2 | 2 |
| 0.805 | 0.794 | 230.8 | 180 | 185.9 | 183.3 | 2 | 2 | 2 |
| 0.836 | 0.723 | 282.7 | 213.2 | 236.2 | 204.4 | 7 | 7 | 7 |
| 1.865 | 1.176 | 7.4 | 12.1 | 13.8 | 8.7 | 1 | 1 | 1 |
| 0.82 | 0.834 | 129 | 111 | 105.8 | 107.6 | 1 | 1 | 1 |
| 0.807 | 0.699 | 126.2 | 105 | 101.9 | 88.2 | 1 | 1 | 1 |
| 0.803 | 0.727 | 47.3 | 38.4 | 38 | 34.4 | 1 | 1 | 1 |
| 0.858 | 0.763 | 1028.9 | 913.9 | 883.1 | 784.6 | 5 | 5 | 5 |
| 0.618 | 0.624 | 128.2 | 102.1 | 79.2 | 80 | 3 | 3 | 3 |
| 0.902 | 0.825 | 91.5 | 85.2 | 82.5 | 75.5 | 2 | 2 | 2 |
| 0.853 | 0.776 | 154.1 | 147.5 | 131.5 | 119.6 | 3 | 3 | 3 |
| 0.803 | 0.718 | 80.8 | 68.7 | 64.9 | 58 | 2 | 2 | 2 |
| 0.725 | 0.71 | 588.7 | 420.2 | 427.1 | 417.9 | 7 | 7 | 7 |
| 1.274 | 0.619 | 8.4 | 10.8 | 10.7 | 5.2 | 1 | 1 | 1 |
| 0.867 | 0.766 | 295 | 243.6 | 255.9 | 226 | 4 | 4 | 4 |
| 0.678 | 0.675 | 63.1 | 52.5 | 42.8 | 42.6 | 1 | 1 | 1 |
| 0.742 | 0.684 | 437.4 | 312.4 | 324.5 | 299.1 | 5 | 5 | 5 |
| 0.785 | 0.813 | 215.8 | 184.1 | 169.4 | 175.5 | 4 | 4 | 4 |
| 0.827 | 0.855 | 226.9 | 339.3 | 187.6 | 194 | 3 | 3 | 3 |
| 0.837 | 0.787 | 232.7 | 201.7 | 194.7 | 183.2 | 3 | 3 | 3 |
| 0.787 | 0.754 | 140.4 | 101.6 | 110.5 | 105.8 | 1 | 1 | 1 |
| 0.88 | 0.805 | 103.6 | 79 | 91.2 | 83.4 | 1 | 1 | 1 |
| 0.863 | 0.826 | 282.8 | 236.8 | 244.1 | 233.5 | 2 | 2 | 2 |
| 0.923 | 0.743 | 60.7 | 49.5 | 56 | 45.1 | 1 | 1 | 1 |
| 0.534 | 0.579 | 37.8 | 29.8 | 20.2 | 21.9 | 1 | 1 | 1 |
| 0.812 | 0.728 | 748.7 | 636 | 607.9 | 545.4 | 10 | 10 | 10 |
| 0.727 | 0.695 | 435.1 | 311.6 | 316.2 | 302.5 | 5 | 5 | 5 |
| 0.694 | 0.655 | 72.8 | 53.4 | 50.5 | 47.7 | 1 | 1 | 1 |
| 0.628 | 0.766 | 73.4 | 50.9 | 46.1 | 56.2 | 1 | 1 | 1 |
| 0.818 | 0.912 | 30.8 | 28.3 | 25.2 | 28.1 | 1 | 1 | 1 |
| 0.826 | 0.708 | 95.9 | 94.7 | 79.2 | 67.9 | 1 | 1 | 1 |
| 0.708 | 0.72 | 191.3 | 135.7 | 135.4 | 137.7 | 2 | 2 | 2 |

| | | | | | | | | |
|---|---|---|---|---|---|---|---|---|
| 0.76 | 0.757 | 1042.6 | 696.4 | 792.3 | 789.5 | 11 | 11 | 11 |
| 0.801 | 0.666 | 126.4 | 100.3 | 101.2 | 84.2 | 2 | 2 | 2 |
| 0.709 | 0.687 | 81.1 | 70.2 | 57.5 | 55.7 | 1 | 1 | 1 |
| 0.949 | 0.843 | 43.4 | 40.8 | 41.2 | 36.6 | 1 | 1 | 1 |
| 0.873 | 0.688 | 72.2 | 62.3 | 63 | 49.7 | 1 | 1 | 1 |
| 0.81 | 0.683 | 111.2 | 91 | 90.1 | 75.9 | 1 | 1 | 1 |
| 0.852 | 0.735 | 19.6 | 15.3 | 16.7 | 14.4 | 1 | 1 | 1 |
| 0.823 | 0.863 | 35.1 | 37.7 | 28.9 | 30.3 | 1 | 1 | 1 |
| 0.821 | 0.76 | 123 | 106 | 101 | 93.5 | 2 | 2 | 2 |
| 0.531 | 0.347 | 4.9 | 1.5 | 2.6 | 1.7 | 1 | 1 | 1 |
| 0.739 | 0.552 | 16.5 | 12.6 | 12.2 | 9.1 | 1 | 1 | 1 |
| 0.808 | 0.774 | 147.6 | 126.6 | 119.2 | 114.2 | 2 | 2 | 2 |
| 0.822 | 0.77 | 1104.1 | 898.3 | 907.9 | 850.7 | 15 | 15 | 15 |
| 0.805 | 0.778 | 118.9 | 86 | 95.7 | 92.5 | 2 | 2 | 2 |
| 0.626 | 0.635 | 372 | 239.3 | 232.9 | 236.3 | 2 | 2 | 2 |
| 0.787 | 0.8 | 80.7 | 67.4 | 63.5 | 64.6 | 1 | 1 | 1 |
| 0.825 | 0.729 | 155.1 | 119.7 | 127.9 | 113.1 | 2 | 2 | 2 |
| 0.875 | 0.769 | 812.8 | 645.7 | 711.5 | 625.4 | 7 | 7 | 7 |
| 0.684 | 0.738 | 130.7 | 89.7 | 89.4 | 96.5 | 3 | 3 | 3 |
| 0.773 | 0.756 | 707.9 | 573.3 | 547.2 | 535 | 7 | 7 | 7 |
| 0.739 | 0.7 | 80.4 | 65.3 | 59.4 | 56.3 | 1 | 1 | 1 |
| 0.875 | 0.703 | 6.4 | 4.1 | 5.6 | 4.5 | 1 | 1 | 1 |
| 0.859 | 0.872 | 63.2 | 59.6 | 54.3 | 55.1 | 1 | 1 | 1 |
| 0.909 | 0.825 | 58.4 | 52.6 | 53.1 | 48.2 | 1 | 1 | 1 |
| 0.817 | 0.67 | 74.9 | 55.2 | 61.2 | 50.2 | 1 | 1 | 1 |
| 0.912 | 0.791 | 106.3 | 85 | 96.9 | 84.1 | 2 | 2 | 2 |
| 0.778 | 0.842 | 78.5 | 86.5 | 61.1 | 66.1 | 1 | 1 | 1 |
| 0.678 | 0.697 | 78.3 | 57.9 | 53.1 | 54.6 | 1 | 1 | 1 |
| 0.771 | 0.693 | 131.1 | 105.4 | 101.1 | 90.9 | 1 | 1 | 1 |
| 1.071 | 0.785 | 109.1 | 95.7 | 116.9 | 85.6 | 1 | 1 | 1 |
| 0.663 | 0.702 | 101.9 | 71 | 67.6 | 71.5 | 2 | 2 | 2 |
| 0.754 | 0.835 | 72.7 | 62.5 | 54.8 | 60.7 | 1 | 1 | 1 |
| 0.936 | 0.744 | 138.2 | 127.9 | 129.4 | 102.8 | 2 | 2 | 2 |
| 0.947 | 0.795 | 76.1 | 88.7 | 72.1 | 60.5 | 1 | 1 | 1 |
| 0.8 | 0.685 | 48.9 | 44.1 | 39.1 | 33.5 | 2 | 2 | 2 |
| 0.765 | 0.784 | 300.7 | 204.9 | 230.1 | 235.6 | 4 | 4 | 4 |
| 0.848 | 0.819 | 42 | 40.9 | 35.6 | 34.4 | 1 | 1 | 1 |
| 0.84 | 0.671 | 74.2 | 63.9 | 62.3 | 49.8 | 1 | 1 | 1 |
| 0.831 | 0.949 | 5.9 | 5 | 4.9 | 5.6 | 1 | 1 | 1 |
| 0.814 | 0.75 | 213.3 | 182 | 173.6 | 160 | 2 | 2 | 2 |

| | | | | | | | | |
|---|---|---|---|---|---|---|---|---|
| 0.956 | 0.748 | 91.7 | 83.7 | 87.7 | 68.6 | 1 | 1 | 1 |
| 0.656 | 0.64 | 172 | 110 | 112.9 | 110.1 | 2 | 2 | 2 |
| 0.632 | 0.644 | 41 | 28.2 | 25.9 | 26.4 | 1 | 1 | 1 |
| 0.93 | 1.054 | 18.6 | 19 | 17.3 | 19.6 | 1 | 1 | 1 |
| 0.704 | 0.806 | 170.6 | 190.1 | 120.1 | 137.5 | 3 | 3 | 3 |
| 0.768 | 0.739 | 157.6 | 133 | 121 | 116.4 | 2 | 2 | 2 |
| 0.825 | 0.776 | 137.1 | 112.2 | 113.1 | 106.4 | 2 | 2 | 2 |
| 0.792 | 0.87 | 33.1 | 25.8 | 26.2 | 28.8 | 1 | 1 | 1 |
| 0.714 | 0.712 | 338 | 256.8 | 241.5 | 240.7 | 2 | 2 | 2 |
| 0.87 | 1.221 | 7.7 | 9.2 | 6.7 | 9.4 | 1 | 1 | 1 |
| 0.883 | 0.814 | 102.6 | 94.2 | 90.6 | 83.5 | 2 | 2 | 2 |
| 0.713 | 0.73 | 165.4 | 125.1 | 117.9 | 120.7 | 3 | 3 | 3 |
| 1.308 | 0.835 | 9.1 | 12.4 | 11.9 | 7.6 | 1 | 1 | 1 |
| 0.781 | 0.767 | 931.9 | 665.1 | 727.4 | 714.4 | 10 | 10 | 10 |
| 0.88 | 0.782 | 32.6 | 29 | 28.7 | 25.5 | 1 | 1 | 1 |
| 0.807 | 0.717 | 96.5 | 75.9 | 77.9 | 69.2 | 1 | 1 | 1 |
| 0.67 | 0.637 | 153.4 | 106.5 | 102.8 | 97.7 | 1 | 1 | 1 |
| 0.859 | 0.704 | 70 | 54.8 | 60.1 | 49.3 | 1 | 1 | 1 |
| 0.702 | 0.713 | 131.9 | 81.6 | 92.6 | 94.1 | 1 | 1 | 1 |
| 0.781 | 0.906 | 119.8 | 113.9 | 93.6 | 108.5 | 1 | 1 | 1 |
| 0.837 | 0.98 | 9.8 | 8.7 | 8.2 | 9.6 | 1 | 1 | 1 |
| 1.092 | 0.98 | 25 | 22.1 | 27.3 | 24.5 | 1 | 1 | 1 |
| 0.876 | 0.783 | 148.1 | 146.6 | 129.7 | 115.9 | 3 | 3 | 3 |
| 0.848 | 0.729 | 150.2 | 129 | 127.3 | 109.5 | 4 | 4 | 4 |
| 0.687 | 0.678 | 105.9 | 87.3 | 72.8 | 71.8 | 2 | 2 | 2 |
| 0.788 | 0.727 | 169.6 | 139 | 133.6 | 123.3 | 1 | 1 | 1 |
| 0.838 | 0.715 | 1027.3 | 843.2 | 861.1 | 735 | 8 | 8 | 8 |
| 0.774 | 0.874 | 34.1 | 35.2 | 26.4 | 29.8 | 1 | 1 | 1 |
| 0.731 | 0.669 | 61 | 45.4 | 44.6 | 40.8 | 1 | 1 | 1 |
| 0.75 | 0.706 | 385.3 | 251.9 | 288.8 | 272 | 6 | 6 | 6 |
| 0.778 | 0.733 | 161.6 | 133 | 125.8 | 118.4 | 3 | 3 | 3 |
| 0.944 | 0.841 | 212.5 | 214.8 | 200.5 | 178.7 | 3 | 3 | 3 |
| 0.685 | 0.75 | 161.1 | 167.6 | 110.3 | 120.8 | 2 | 2 | 2 |
| 0.695 | 0.604 | 76.8 | 57 | 53.4 | 46.4 | 1 | 1 | 1 |
| 0.803 | 0.739 | 388.1 | 311.7 | 311.5 | 286.9 | 9 | 9 | 9 |
| 0.784 | 0.753 | 377.6 | 278.9 | 296 | 284.2 | 2 | 2 | 2 |
| 0.788 | 0.768 | 34.5 | 30.8 | 27.2 | 26.5 | 1 | 1 | 1 |
| 0.816 | 0.735 | 336.2 | 235.2 | 274.5 | 247 | 4 | 4 | 4 |
| 0.756 | 0.748 | 533.5 | 403.3 | 403.3 | 398.8 | 7 | 7 | 7 |
| 0.821 | 0.812 | 123.7 | 93.6 | 101.6 | 100.5 | 1 | 1 | 1 |
| 0.853 | 0.804 | 135.9 | 101.8 | 115.9 | 109.2 | 2 | 2 | 2 |
| 0.855 | 0.745 | 137 | 106.9 | 117.2 | 102 | 1 | 1 | 1 |
| 0.831 | 0.632 | 132.7 | 100.9 | 110.3 | 83.9 | 1 | 1 | 1 |
| 0.756 | 0.755 | 111.2 | 80 | 84.1 | 84 | 1 | 1 | 1 |
| 0.891 | 0.798 | 11.9 | 15.3 | 10.6 | 9.5 | 1 | 1 | 1 |

| | | | | | | | | |
|---|---|---|---|---|---|---|---|---|
| 0.651 | 0.687 | 49.5 | 42.1 | 32.2 | 34 | 1 | 1 | 1 |
| 0.815 | 0.649 | 48.7 | 45.9 | 39.7 | 31.6 | 2 | 2 | 2 |
| 0.72 | 0.61 | 61 | 37.1 | 43.9 | 37.2 | 1 | 1 | 1 |
| 0.916 | 0.831 | 168.6 | 186.9 | 154.5 | 140.1 | 2 | 2 | 2 |
| 0.711 | 0.639 | 82 | 52.5 | 58.3 | 52.4 | 1 | 1 | 1 |
| 0.853 | 0.8 | 201.5 | 188.6 | 171.8 | 161.2 | 2 | 2 | 2 |
| 0.811 | 0.726 | 194.9 | 146.6 | 158 | 141.5 | 2 | 2 | 2 |
| 0.781 | 0.837 | 158.4 | 128.1 | 123.7 | 132.6 | 2 | 2 | 2 |
| 0.709 | 0.772 | 829.2 | 662 | 588.2 | 640 | 10 | 10 | 10 |
| 0.7 | 0.703 | 127.6 | 99.3 | 89.3 | 89.7 | 1 | 1 | 1 |
| 0.695 | 0.753 | 43 | 30.6 | 29.9 | 32.4 | 1 | 1 | 1 |
| 0.794 | 0.644 | 73.8 | 56.6 | 58.6 | 47.5 | 1 | 1 | 1 |
| 0.853 | 0.784 | 1124.8 | 1047.1 | 960 | 881.4 | 6 | 6 | 6 |
| 0.888 | 0.882 | 47.4 | 48.1 | 42.1 | 41.8 | 1 | 1 | 1 |
| 0.871 | 0.704 | 61.9 | 53.2 | 53.9 | 43.6 | 1 | 1 | 1 |
| 0.637 | 0.657 | 64.4 | 55 | 41 | 42.3 | 1 | 1 | 1 |
| 0.755 | 0.592 | 105 | 86.1 | 79.3 | 62.2 | 1 | 1 | 1 |
| 0.59 | 0.673 | 106.7 | 74.5 | 63 | 71.8 | 1 | 1 | 1 |
| 0.811 | 0.745 | 59.9 | 45.7 | 48.6 | 44.6 | 1 | 1 | 1 |
| 0.889 | 0.868 | 28 | 29.2 | 24.9 | 24.3 | 1 | 1 | 1 |
| 0.921 | 0.85 | 59.4 | 56.7 | 54.7 | 50.5 | 1 | 1 | 1 |
| 1.003 | 0.952 | 93.7 | 83.2 | 94 | 89.2 | 2 | 2 | 2 |
| 0.786 | 0.722 | 18.7 | 11.6 | 14.7 | 13.5 | 1 | 1 | 1 |
| 0.564 | 0.684 | 201 | 135 | 113.4 | 137.5 | 2 | 2 | 2 |
| 0.686 | 0.63 | 117 | 79.1 | 80.3 | 73.7 | 1 | 1 | 1 |
| 0.648 | 0.795 | 17.6 | 15.4 | 11.4 | 14 | 1 | 1 | 1 |
| 0.832 | 0.828 | 70.8 | 60.8 | 58.9 | 58.6 | 2 | 2 | 2 |
| 0.763 | 0.792 | 106.9 | 106 | 81.6 | 84.7 | 3 | 3 | 3 |
| 0.802 | 0.656 | 776.8 | 568.8 | 622.9 | 509.3 | 8 | 8 | 8 |
| 0.8 | 0.716 | 159 | 130.1 | 127.2 | 113.8 | 1 | 1 | 1 |
| 0.826 | 0.853 | 50.5 | 45.2 | 41.7 | 43.1 | 1 | 1 | 1 |
| 0.79 | 0.669 | 179.7 | 144.7 | 141.9 | 120.2 | 2 | 2 | 2 |
| 0.704 | 0.644 | 28.4 | 18.1 | 20 | 18.3 | 2 | 2 | 2 |
| 0.595 | 0.612 | 58.2 | 39.5 | 34.6 | 35.6 | 1 | 1 | 1 |
| 0.861 | 0.782 | 374.3 | 297.2 | 322.2 | 292.6 | 1 | 1 | 1 |
| 0.887 | 0.9 | 91.3 | 82.9 | 81 | 82.2 | 1 | 1 | 1 |
| 0.859 | 0.771 | 626.5 | 496.2 | 538.4 | 482.9 | 9 | 9 | 9 |

| | | | | | | | | |
|---|---|---|---|---|---|---|---|---|
| 0.665 | 0.575 | 83.1 | 53.3 | 55.3 | 47.8 | 1 | 1 | 1 |
| 0.993 | 0.851 | 222.9 | 240 | 221.4 | 189.7 | 5 | 5 | 5 |
| 0.824 | 0.729 | 34 | 26.7 | 28 | 24.8 | 1 | 1 | 1 |
| 1.015 | 0.877 | 32.5 | 36.5 | 33 | 28.5 | 1 | 1 | 1 |
| 0.777 | 0.747 | 113.1 | 92 | 87.9 | 84.5 | 1 | 1 | 1 |
| 0.737 | 0.796 | 94.3 | 77.7 | 69.5 | 75.1 | 1 | 1 | 1 |
| 0.907 | 0.92 | 31.2 | 33 | 28.3 | 28.7 | 1 | 1 | 1 |
| 0.741 | 0.725 | 628.3 | 514.2 | 465.7 | 455.4 | 6 | 6 | 6 |
| 0.812 | 0.818 | 141.8 | 105.4 | 115.2 | 116 | 2 | 2 | 2 |
| 0.824 | 0.737 | 1108.4 | 1086 | 913.4 | 816.4 | 9 | 9 | 9 |
| 0.888 | 0.978 | 22.4 | 22.1 | 19.9 | 21.9 | 1 | 1 | 1 |
| 0.659 | 0.672 | 264.9 | 145.8 | 174.5 | 177.9 | 1 | 1 | 1 |
| 0.839 | 0.651 | 96.3 | 80.1 | 80.8 | 62.7 | 1 | 1 | 1 |
| 0.843 | 0.881 | 72.2 | 78.6 | 60.9 | 63.6 | 3 | 3 | 3 |
| 0.798 | 0.649 | 88.1 | 82.7 | 70.3 | 57.2 | 3 | 3 | 3 |
| 0.767 | 0.741 | 189.8 | 150.8 | 145.6 | 140.7 | 4 | 4 | 4 |
| 0.899 | 0.803 | 252.3 | 227.4 | 226.8 | 202.7 | 5 | 5 | 5 |
| 0.866 | 0.784 | 866.8 | 797.1 | 750.8 | 679.2 | 11 | 11 | 11 |
| 0.844 | 0.789 | 426 | 351.7 | 359.6 | 336.1 | 3 | 3 | 3 |
| 0.431 | 0.69 | 11.6 | 5.7 | 5 | 8 | 1 | 1 | 1 |
| 0.982 | 0.765 | 90.4 | 90.7 | 88.8 | 69.2 | 2 | 2 | 2 |
| 0.718 | 0.737 | 104 | 91.2 | 74.7 | 76.6 | 1 | 1 | 1 |
| 0.778 | 0.64 | 70.2 | 47.7 | 54.6 | 44.9 | 1 | 1 | 1 |
| 0.861 | 0.775 | 939.3 | 828 | 809.2 | 728.2 | 12 | 12 | 12 |
| 0.807 | 0.68 | 46.6 | 32.7 | 37.6 | 31.7 | 1 | 1 | 1 |
| 0.77 | 0.702 | 58.7 | 50.2 | 45.2 | 41.2 | 1 | 1 | 1 |
| 0.752 | 0.683 | 242 | 181.4 | 182.1 | 165.2 | 2 | 2 | 2 |
| 0.72 | 0.747 | 549.6 | 435.1 | 395.5 | 410.8 | 9 | 9 | 9 |
| 1.056 | 0.973 | 44.4 | 39.2 | 46.9 | 43.2 | 1 | 1 | 1 |
| 0.749 | 0.627 | 339.6 | 235.4 | 254.5 | 213 | 5 | 5 | 5 |
| 0.778 | 0.752 | 511.8 | 422.5 | 398.4 | 384.9 | 8 | 8 | 8 |
| 0.853 | 0.714 | 46.9 | 45.7 | 40 | 33.5 | 1 | 1 | 1 |
| 0.818 | 0.786 | 521.2 | 378.5 | 426.6 | 409.8 | 8 | 8 | 8 |
| 0.784 | 0.784 | 69.5 | 61.4 | 54.5 | 54.5 | 1 | 1 | 1 |
| 0.732 | 0.789 | 167 | 130.1 | 122.3 | 131.8 | 2 | 2 | 2 |
| 0.828 | 0.683 | 113.6 | 89 | 94.1 | 77.6 | 2 | 2 | 2 |
| 0.949 | 0.842 | 39.2 | 39 | 37.2 | 33 | 1 | 1 | 1 |
| 0.763 | 0.789 | 156.8 | 137.7 | 119.7 | 123.7 | 3 | 3 | 3 |
| 0.883 | 0.74 | 78.7 | 62.7 | 69.5 | 58.2 | 1 | 1 | 1 |
| 0.663 | 0.646 | 17.8 | 15 | 11.8 | 11.5 | 1 | 1 | 1 |
| 0.824 | 0.783 | 501 | 400.4 | 412.6 | 392.2 | 7 | 7 | 7 |
| 0.635 | 0.648 | 1145.8 | 827.3 | 727.5 | 743 | 12 | 12 | 12 |

| | | | | | | | | |
|---|---|---|---|---|---|---|---|---|
| 0.814 | 0.712 | 113.7 | 93.7 | 92.6 | 80.9 | 1 | 1 | 1 |
| 0.783 | 0.711 | 1022.6 | 822.1 | 800.5 | 726.7 | 10 | 10 | 10 |
| 0.881 | 0.828 | 43 | 37.4 | 37.9 | 35.6 | 1 | 1 | 1 |
| 1.095 | 0.801 | 50.3 | 44.1 | 55.1 | 40.3 | 1 | 1 | 1 |
| 0.783 | 0.673 | 136.1 | 107.5 | 106.5 | 91.6 | 1 | 1 | 1 |
| 0.718 | 0.673 | 449.7 | 383 | 322.9 | 302.7 | 5 | 5 | 5 |
| 0.694 | 0.754 | 51.7 | 34.7 | 35.9 | 39 | 1 | 1 | 1 |
| 0.571 | 0.637 | 9.1 | 4.8 | 5.2 | 5.8 | 1 | 1 | 1 |
| 0.615 | 0.594 | 100.5 | 79.6 | 61.8 | 59.7 | 2 | 2 | 2 |
| 0.887 | 0.791 | 80.7 | 64.4 | 71.6 | 63.8 | 1 | 1 | 1 |
| 0.72 | 0.829 | 32.2 | 27.4 | 23.2 | 26.7 | 1 | 1 | 1 |
| 0.88 | 0.675 | 110.7 | 104.2 | 97.4 | 74.7 | 2 | 2 | 2 |
| 0.885 | 0.656 | 38.4 | 32.6 | 34 | 25.2 | 1 | 1 | 1 |
| 0.879 | 0.842 | 194.5 | 170.7 | 170.9 | 163.8 | 3 | 3 | 3 |
| 0.756 | 0.77 | 81.2 | 57.6 | 61.4 | 62.5 | 1 | 1 | 1 |
| 0.378 | 0.483 | 1821.4 | 968.1 | 687.7 | 880.2 | 12 | 12 | 12 |
| 0.798 | 0.764 | 91.6 | 79.6 | 73.1 | 70 | 3 | 3 | 3 |
| 0.947 | 1.137 | 97.9 | 190.8 | 92.7 | 111.3 | 1 | 1 | 1 |
| 0.881 | 0.701 | 46.9 | 38.3 | 41.3 | 32.9 | 1 | 1 | 1 |
| 0.838 | 0.738 | 164.8 | 168.9 | 138.1 | 121.6 | 2 | 2 | 2 |
| 0.799 | 0.723 | 569.5 | 524.7 | 454.8 | 412 | 5 | 5 | 5 |
| 0.758 | 0.737 | 219 | 185.1 | 166 | 161.3 | 3 | 3 | 3 |
| 0.868 | 0.753 | 156.9 | 153.2 | 136.2 | 118.1 | 2 | 2 | 2 |
| 0.82 | 0.745 | 990.8 | 875.6 | 812.2 | 738.6 | 14 | 14 | 14 |
| 0.882 | 0.805 | 77.4 | 69.4 | 68.3 | 62.3 | 1 | 1 | 1 |
| 0.834 | 0.834 | 150.8 | 118.5 | 125.7 | 125.8 | 2 | 2 | 2 |
| 0.845 | 0.737 | 1769.7 | 1512.5 | 1494.6 | 1304.4 | 26 | 26 | 26 |
| 0.878 | 0.322 | 11.5 | 10.6 | 10.1 | 3.7 | 1 | 1 | 1 |
| 0.858 | 0.72 | 257.6 | 205.6 | 221 | 185.4 | 2 | 2 | 2 |
| 0.791 | 0.736 | 829.2 | 686.8 | 655.6 | 610.6 | 12 | 12 | 12 |
| 0.874 | 0.767 | 147.3 | 115.2 | 128.7 | 113 | 2 | 2 | 2 |
| 0.795 | 0.685 | 153.8 | 115.2 | 122.3 | 105.3 | 2 | 2 | 2 |
| 0.52 | 0.644 | 224 | 157.4 | 116.4 | 144.3 | 3 | 3 | 3 |
| 1.12 | 1.253 | 38.4 | 42.3 | 43 | 48.1 | 1 | 1 | 1 |
| 0.672 | 0.762 | 103.2 | 77 | 69.3 | 78.6 | 1 | 1 | 1 |
| 0.773 | 0.748 | 480.2 | 369.8 | 371 | 359.3 | 5 | 5 | 5 |
| 0.902 | 0.786 | 214.7 | 182.7 | 193.7 | 168.7 | 4 | 4 | 4 |
| 0.97 | 0.816 | 100.4 | 107.7 | 97.4 | 81.9 | 3 | 3 | 3 |
| 0.737 | 0.74 | 129.5 | 87.1 | 95.4 | 95.8 | 2 | 2 | 2 |
| 0.798 | 0.671 | 37.7 | 30.6 | 30.1 | 25.3 | 1 | 1 | 1 |
| 0.872 | 0.78 | 34.5 | 31.2 | 30.1 | 26.9 | 1 | 1 | 1 |

| | | | | | | | | |
|---|---|---|---|---|---|---|---|---|
| 0.678 | 0.493 | 14.6 | 14.2 | 9.9 | 7.2 | 1 | 1 | 1 |
| 0.782 | 0.715 | 79.7 | 68.8 | 62.3 | 57 | 1 | 1 | 1 |
| 0.793 | 0.689 | 69.7 | 58.4 | 55.3 | 48 | 2 | 2 | 2 |
| 0.727 | 0.807 | 72.2 | 65.1 | 52.5 | 58.3 | 1 | 1 | 1 |
| 0.713 | 0.732 | 93.3 | 71.1 | 66.5 | 68.3 | 2 | 2 | 2 |
| 0.672 | 0.61 | 183.8 | 140.2 | 123.6 | 112.2 | 3 | 3 | 3 |
| 0.758 | 0.747 | 260.4 | 211.1 | 197.5 | 194.4 | 2 | 2 | 2 |
| 0.838 | 0.845 | 213.3 | 170.7 | 178.7 | 180.3 | 4 | 4 | 4 |
| 1.102 | 0.889 | 24.4 | 26 | 26.9 | 21.7 | 1 | 1 | 1 |
| 0.937 | 0.868 | 31.9 | 31 | 29.9 | 27.7 | 1 | 1 | 1 |
| 0.736 | 0.702 | 512.5 | 486.8 | 377 | 360 | 3 | 3 | 3 |
| 0.592 | 0.796 | 15.7 | 12.7 | 9.3 | 12.5 | 1 | 1 | 1 |
| 0.895 | 0.658 | 26.6 | 27.3 | 23.8 | 17.5 | 1 | 1 | 1 |
| 0.856 | 0.759 | 51.5 | 49.3 | 44.1 | 39.1 | 1 | 1 | 1 |
| 0.88 | 0.751 | 2812 | 2427.1 | 2474.1 | 2112.2 | 31 | 31 | 31 |
| 0.754 | 0.65 | 136.7 | 112.8 | 103.1 | 88.8 | 1 | 1 | 1 |
| 0.776 | 0.779 | 129.3 | 100.1 | 100.3 | 100.7 | 1 | 1 | 1 |
| 0.769 | 0.73 | 685.6 | 525.4 | 527.3 | 500.2 | 7 | 7 | 7 |
| 0.683 | 0.604 | 142.4 | 112.3 | 97.2 | 86 | 1 | 1 | 1 |
| 0.732 | 0.694 | 77.1 | 59.4 | 56.4 | 53.5 | 2 | 2 | 2 |
| 0.857 | 0.839 | 44.8 | 34.1 | 38.4 | 37.6 | 1 | 1 | 1 |
| 0.767 | 0.738 | 198.9 | 126.1 | 152.5 | 146.7 | 2 | 2 | 2 |
| 1.035 | 0.907 | 31.2 | 32 | 32.3 | 28.3 | 1 | 1 | 1 |
| 0.797 | 0.862 | 60.7 | 51.9 | 48.4 | 52.3 | 1 | 1 | 1 |
| 0.962 | 0.844 | 229.5 | 210.9 | 220.7 | 193.6 | 3 | 3 | 3 |
| 1.2 | 0.763 | 76.9 | 105.1 | 92.3 | 58.7 | 1 | 1 | 1 |
| 0.655 | 0.696 | 82.9 | 56.8 | 54.3 | 57.7 | 2 | 2 | 2 |
| 0.684 | 0.63 | 80 | 62.5 | 54.7 | 50.4 | 1 | 1 | 1 |
| 0.825 | 0.703 | 109.6 | 86.4 | 90.4 | 77.1 | 1 | 1 | 1 |
| 0.884 | 0.764 | 95 | 85.1 | 84 | 72.6 | 1 | 1 | 1 |
| 0.64 | 0.633 | 50.9 | 27.7 | 32.6 | 32.2 | 1 | 1 | 1 |
| 0.988 | 0.653 | 24.5 | 23.4 | 24.2 | 16 | 1 | 1 | 1 |
| 0.899 | 0.838 | 335.9 | 310 | 302.1 | 281.4 | 5 | 5 | 5 |
| 0.872 | 0.794 | 107.5 | 89.9 | 93.7 | 85.4 | 3 | 3 | 3 |
| 0.857 | 0.786 | 207.4 | 174.4 | 177.7 | 163 | 3 | 3 | 3 |
| 0.892 | 0.749 | 218.6 | 211.4 | 195 | 163.8 | 2 | 2 | 2 |
| 0.742 | 0.679 | 214.5 | 141.7 | 159.2 | 145.7 | 2 | 2 | 2 |
| 0.863 | 0.952 | 48.1 | 36.4 | 41.5 | 45.8 | 1 | 1 | 1 |

| | | | | | | | | |
|---|---|---|---|---|---|---|---|---|
| 0.758 | 0.669 | 742.6 | 517 | 562.6 | 496.9 | 7 | 7 | 7 |
| 1 | 0.906 | 6.4 | 5.8 | 6.4 | 5.8 | 1 | 1 | 1 |
| 0.724 | 0.677 | 107.9 | 95.9 | 78.1 | 73.1 | 1 | 1 | 1 |
| 0.763 | 0.636 | 33.8 | 32.1 | 25.8 | 21.5 | 1 | 1 | 1 |
| 0.782 | 0.615 | 54.6 | 43.7 | 42.7 | 33.6 | 1 | 1 | 1 |
| 0.591 | 0.739 | 46 | 23.8 | 27.2 | 34 | 1 | 1 | 1 |
| 0.747 | 0.729 | 150.3 | 128.6 | 112.3 | 109.5 | 2 | 2 | 2 |
| 0.848 | 0.63 | 88.1 | 77.5 | 74.7 | 55.5 | 2 | 2 | 2 |
| 0.697 | 0.702 | 127.5 | 105.8 | 88.9 | 89.5 | 1 | 1 | 1 |
| 0.885 | 0.783 | 120.8 | 97.3 | 106.9 | 94.6 | 2 | 2 | 2 |
| 1.343 | 1.01 | 10.5 | 11.9 | 14.1 | 10.6 | 1 | 1 | 1 |
| 0.97 | 0.869 | 49.8 | 44.7 | 48.3 | 43.3 | 1 | 1 | 1 |
| 0.768 | 0.675 | 433.8 | 364.4 | 333.3 | 292.6 | 6 | 6 | 6 |
| 0.795 | 0.872 | 67.9 | 56.4 | 54 | 59.2 | 1 | 1 | 1 |
| 0.697 | 0.667 | 44.5 | 30.6 | 31 | 29.7 | 1 | 1 | 1 |
| 0.807 | 0.76 | 52.4 | 46.9 | 42.3 | 39.8 | 1 | 1 | 1 |
| 0.712 | 0.808 | 7.3 | 9.4 | 5.2 | 5.9 | 1 | 1 | 1 |
| 0.778 | 0.61 | 1134.2 | 841.5 | 882 | 692.3 | 15 | 15 | 15 |
| 0.882 | 0.844 | 33.9 | 28.2 | 29.9 | 28.6 | 1 | 1 | 1 |
| 0.901 | 0.737 | 44.5 | 44.2 | 40.1 | 32.8 | 1 | 1 | 1 |
| 0.911 | 0.832 | 1681.6 | 1497.8 | 1531.3 | 1398.7 | 27 | 27 | 27 |
| 0.884 | 0.821 | 243.6 | 213.4 | 215.3 | 200.1 | 5 | 5 | 5 |
| 0.794 | 0.8 | 166.9 | 126.3 | 132.6 | 133.6 | 2 | 2 | 2 |
| 1.066 | 0.869 | 49.7 | 63.1 | 53 | 43.2 | 2 | 2 | 2 |
| 0.794 | 0.683 | 12.6 | 12.3 | 10 | 8.6 | 1 | 1 | 1 |
| 0.872 | 0.803 | 50.8 | 42.3 | 44.3 | 40.8 | 1 | 1 | 1 |
| 0.817 | 0.663 | 98.4 | 88.4 | 80.4 | 65.2 | 1 | 1 | 1 |
| 1.16 | 0.752 | 12.5 | 15.1 | 14.5 | 9.4 | 1 | 1 | 1 |
| 0.986 | 0.891 | 84.5 | 84 | 83.3 | 75.3 | 1 | 1 | 1 |
| 0.751 | 0.767 | 93.1 | 85.7 | 69.9 | 71.4 | 1 | 1 | 1 |
| 0.634 | 0.575 | 100.3 | 77.6 | 63.6 | 57.7 | 1 | 1 | 1 |
| 0.472 | 0.543 | 52.3 | 37 | 24.7 | 28.4 | 1 | 1 | 1 |
| 0.749 | 0.68 | 164.9 | 132.5 | 123.5 | 112.2 | 1 | 1 | 1 |
| 0.888 | 0.773 | 298 | 310.1 | 264.5 | 230.5 | 8 | 8 | 8 |
| 0.772 | 0.678 | 258 | 230.5 | 199.3 | 174.9 | 2 | 2 | 2 |
| 0.844 | 0.542 | 67.5 | 53.2 | 57 | 36.6 | 1 | 1 | 1 |
| 0.841 | 0.753 | 29.6 | 20.4 | 24.9 | 22.3 | 1 | 1 | 1 |
| 0.932 | 0.86 | 44.2 | 40.2 | 41.2 | 38 | 1 | 1 | 1 |
| 0.703 | 0.671 | 707.7 | 467.7 | 497.3 | 474.7 | 3 | 3 | 3 |
| 1.008 | 0.929 | 25.5 | 23.9 | 25.7 | 23.7 | 1 | 1 | 1 |
| 0.743 | 0.785 | 142.8 | 105.8 | 106.1 | 112.1 | 1 | 1 | 1 |
| 0.775 | 0.728 | 174.1 | 133 | 135 | 126.7 | 3 | 3 | 3 |
| 0.724 | 0.617 | 109.7 | 87.6 | 79.4 | 67.7 | 1 | 1 | 1 |

| | | | | | | | | |
|---|---|---|---|---|---|---|---|---|
| 1.015 | 0.814 | 19.9 | 18.7 | 20.2 | 16.2 | 1 | 1 | 1 |
| 0.806 | 0.662 | 547.7 | 467.1 | 441.6 | 362.8 | 10 | 10 | 10 |
| 0.969 | 0.835 | 106.3 | 96.5 | 103 | 88.8 | 1 | 1 | 1 |
| 0.813 | 0.743 | 132.5 | 106.8 | 107.7 | 98.5 | 1 | 1 | 1 |
| 0.849 | 0.803 | 68.1 | 57.3 | 57.8 | 54.7 | 1 | 1 | 1 |
| 0.948 | 0.705 | 46.5 | 42.6 | 44.1 | 32.8 | 1 | 1 | 1 |
| 0.897 | 0.866 | 90 | 77.2 | 80.7 | 77.9 | 1 | 1 | 1 |
| 0.828 | 0.754 | 88.9 | 79.5 | 73.6 | 67 | 1 | 1 | 1 |
| 0.926 | 0.79 | 358.4 | 342.1 | 331.8 | 283.2 | 4 | 4 | 4 |
| 0.663 | 0.51 | 843 | 559 | 559.3 | 430.1 | 5 | 5 | 5 |
| 0.929 | 0.703 | 71.8 | 62.5 | 66.7 | 50.5 | 1 | 1 | 1 |
| 0.813 | 0.813 | 10.7 | 9.7 | 8.7 | 8.7 | 1 | 1 | 1 |
| 0.359 | 0.511 | 9.2 | 5.6 | 3.3 | 4.7 | 1 | 1 | 1 |
| 0.858 | 0.747 | 379.6 | 288.5 | 325.8 | 283.6 | 3 | 3 | 3 |
| 0.819 | 0.622 | 220.5 | 179.7 | 180.6 | 137.1 | 1 | 1 | 1 |
| 0.876 | 0.733 | 93.7 | 84.6 | 82.1 | 68.7 | 2 | 2 | 2 |
| 0.749 | 0.705 | 119.1 | 98.9 | 89.2 | 84 | 1 | 1 | 1 |
| 0.807 | 0.744 | 188 | 163.2 | 151.7 | 139.8 | 2 | 2 | 2 |
| 0.966 | 0.844 | 26.3 | 26.7 | 25.4 | 22.2 | 1 | 1 | 1 |
| 0.844 | 0.776 | 96.5 | 94.9 | 81.4 | 74.9 | 1 | 1 | 1 |
| 0.874 | 0.785 | 226.9 | 245.2 | 198.3 | 178.1 | 2 | 2 | 2 |
| 1.014 | 0.919 | 99 | 117.6 | 100.4 | 91 | 2 | 2 | 2 |
| 0.798 | 0.72 | 125.5 | 104.6 | 100.1 | 90.4 | 2 | 2 | 2 |
| 0.878 | 0.815 | 314.3 | 276.2 | 276 | 256.3 | 4 | 4 | 4 |
| 0.77 | 0.742 | 452.6 | 349.7 | 348.6 | 336 | 5 | 5 | 5 |
| 1.705 | 1.318 | 4.4 | 6.2 | 7.5 | 5.8 | 1 | 1 | 1 |
| 0.737 | 0.801 | 315.8 | 244.5 | 232.7 | 252.8 | 3 | 3 | 3 |
| 0.692 | 0.696 | 145.5 | 93.5 | 100.7 | 101.2 | 1 | 1 | 1 |
| 0.83 | 0.818 | 39.5 | 36.5 | 32.8 | 32.3 | 1 | 1 | 1 |
| 0.883 | 0.866 | 181.6 | 160.3 | 160.3 | 157.3 | 3 | 3 | 3 |
| 0.74 | 0.74 | 38.9 | 29 | 28.8 | 28.8 | 1 | 1 | 1 |
| 0.887 | 0.83 | 42.3 | 33.6 | 37.5 | 35.1 | 1 | 1 | 1 |
| 0.864 | 0.748 | 378.8 | 367.3 | 327.1 | 283.5 | 5 | 5 | 5 |
| 0.763 | 0.738 | 172.3 | 121.5 | 131.4 | 127.2 | 2 | 2 | 2 |
| 0.641 | 0.722 | 127 | 97.8 | 81.4 | 91.7 | 1 | 1 | 1 |
| 0.93 | 0.778 | 482.9 | 425.8 | 449 | 375.6 | 5 | 5 | 5 |
| 0.731 | 0.714 | 54.2 | 43.2 | 39.6 | 38.7 | 1 | 1 | 1 |
| 0.855 | 0.755 | 1007.7 | 850.5 | 861.8 | 760.9 | 17 | 17 | 17 |
| 0.821 | 0.833 | 81.6 | 52.3 | 67 | 68 | 1 | 1 | 1 |
| 0.95 | 0.94 | 383 | 303.7 | 364 | 360.1 | 7 | 7 | 7 |
| 0.847 | 0.764 | 503.3 | 421.2 | 426.1 | 384.5 | 5 | 5 | 5 |

| | | | | | | | | |
|---|---|---|---|---|---|---|---|---|
| 0.865 | 0.745 | 213.8 | 183.3 | 185 | 159.2 | 3 | 3 | 3 |
| 0.795 | 0.764 | 102.4 | 88.7 | 81.4 | 78.2 | 2 | 2 | 2 |
| 0.808 | 0.751 | 334.7 | 262.9 | 270.4 | 251.2 | 5 | 5 | 5 |
| 0.994 | 0.805 | 368.3 | 429.1 | 366.1 | 296.6 | 8 | 8 | 8 |
| 0.647 | 0.676 | 52.1 | 42.6 | 33.7 | 35.2 | 1 | 1 | 1 |
| 0.825 | 0.71 | 209 | 185.7 | 172.4 | 148.3 | 1 | 1 | 1 |
| 0.833 | 0.749 | 1199.9 | 1003.9 | 999.8 | 898.7 | 16 | 16 | 16 |
| 0.936 | 0.76 | 26.7 | 26.1 | 25 | 20.3 | 1 | 1 | 1 |
| 0.989 | 0.684 | 19 | 13.9 | 18.8 | 13 | 1 | 1 | 1 |
| 0.839 | 0.723 | 228.5 | 198.5 | 191.7 | 165.1 | 4 | 4 | 4 |
| 0.837 | 0.837 | 28.8 | 23.2 | 24.1 | 24.1 | 1 | 1 | 1 |
| 0.788 | 0.715 | 50.5 | 46.3 | 39.8 | 36.1 | 1 | 1 | 1 |
| 0.839 | 0.688 | 175.4 | 149 | 147.1 | 120.6 | 3 | 3 | 3 |
| 0.597 | 0.693 | 54.1 | 39 | 32.3 | 37.5 | 1 | 1 | 1 |
| 0.907 | 0.771 | 161.8 | 156.2 | 146.8 | 124.8 | 3 | 3 | 3 |
| 0.673 | 0.781 | 38.8 | 34.1 | 26.1 | 30.3 | 1 | 1 | 1 |
| 0.852 | 0.932 | 35.2 | 31.8 | 30 | 32.8 | 1 | 1 | 1 |
| 0.883 | 0.711 | 128.5 | 110.7 | 113.5 | 91.4 | 3 | 3 | 3 |
| 0.896 | 0.782 | 91.4 | 65.2 | 81.9 | 71.5 | 1 | 1 | 1 |
| 1.037 | 1.015 | 46.4 | 52.2 | 48.1 | 47.1 | 1 | 1 | 1 |
| 0.523 | 0.551 | 136.5 | 93.3 | 71.4 | 75.2 | 1 | 1 | 1 |
| 0.809 | 0.734 | 274.8 | 255.6 | 222.3 | 201.6 | 3 | 3 | 3 |
| 0.753 | 0.859 | 218.8 | 202.3 | 164.7 | 187.9 | 3 | 3 | 3 |
| 0.769 | 0.689 | 111.9 | 86.6 | 86 | 77.1 | 1 | 1 | 1 |
| 0.898 | 0.866 | 34.4 | 34.5 | 30.9 | 29.8 | 1 | 1 | 1 |
| 0.743 | 0.723 | 683 | 501.2 | 507.5 | 493.8 | 10 | 10 | 10 |
| 1.277 | 0.66 | 4.7 | 7.2 | 6 | 3.1 | 1 | 1 | 1 |
| 0.726 | 0.706 | 355 | 288.9 | 257.8 | 250.6 | 5 | 5 | 5 |
| 1.016 | 0.891 | 433.2 | 391.6 | 440 | 386.1 | 6 | 6 | 6 |
| 0.917 | 0.924 | 110.2 | 90.5 | 101 | 101.8 | 1 | 1 | 1 |
| 0.814 | 0.806 | 174.6 | 140.1 | 142.1 | 140.7 | 4 | 4 | 4 |
| 0.765 | 0.803 | 149.1 | 125 | 114 | 119.7 | 1 | 1 | 1 |
| 0.792 | 0.742 | 105.6 | 93.8 | 83.6 | 78.4 | 2 | 2 | 2 |
| 0.902 | 0.8 | 61.4 | 51.7 | 55.4 | 49.1 | 2 | 2 | 2 |
| 0.957 | 0.893 | 42.2 | 44.7 | 40.4 | 37.7 | 1 | 1 | 1 |
| 0.728 | 0.744 | 365.7 | 271.6 | 266.3 | 272 | 4 | 4 | 4 |
| 1.031 | 0.906 | 28.8 | 35.7 | 29.7 | 26.1 | 2 | 2 | 2 |
| 0.869 | 0.725 | 308.7 | 268.1 | 268.4 | 223.8 | 3 | 3 | 3 |

| | | | | | | | | |
|---|---|---|---|---|---|---|---|---|
| 0.723 | 0.758 | 430.3 | 290.1 | 311.2 | 326.1 | 4 | 4 | 4 |
| 0.801 | 0.699 | 609.3 | 479.7 | 487.8 | 426.1 | 8 | 8 | 8 |
| 0.769 | 0.769 | 2406.7 | 1734.1 | 1850.6 | 1851.7 | 33 | 33 | 33 |
| 0.823 | 0.766 | 312.7 | 295.7 | 257.2 | 239.5 | 4 | 4 | 4 |
| 0.861 | 0.743 | 202.2 | 164.7 | 174.1 | 150.3 | 3 | 3 | 3 |
| 0.705 | 0.772 | 393.3 | 291.9 | 277.1 | 303.7 | 4 | 4 | 4 |
| 0.7 | 0.642 | 326.4 | 251.3 | 228.6 | 209.7 | 4 | 4 | 4 |
| 0.94 | 0.969 | 91.2 | 87.8 | 85.7 | 88.4 | 1 | 1 | 1 |
| 0.637 | 0.69 | 150.5 | 93.8 | 95.8 | 103.9 | 1 | 1 | 1 |
| 0.861 | 0.789 | 130.1 | 98 | 112 | 102.7 | 1 | 1 | 1 |
| 0.801 | 0.755 | 233.3 | 192.6 | 186.9 | 176.1 | 2 | 2 | 2 |
| 0.845 | 0.831 | 233.5 | 199.6 | 197.3 | 194.1 | 1 | 1 | 1 |
| 0.858 | 0.739 | 325.5 | 261 | 279.2 | 240.4 | 5 | 5 | 5 |
| 1.035 | 0.894 | 65.3 | 67 | 67.6 | 58.4 | 1 | 1 | 1 |
| 0.715 | 0.698 | 573.8 | 443.2 | 410 | 400.5 | 4 | 4 | 4 |
| 0.869 | 0.789 | 528.6 | 480.8 | 459.5 | 417 | 10 | 10 | 10 |
| 0.656 | 0.677 | 89.2 | 61 | 58.5 | 60.4 | 1 | 1 | 1 |
| 0.913 | 0.739 | 36.8 | 33.9 | 33.6 | 27.2 | 1 | 1 | 1 |
| 0.943 | 1.172 | 8.7 | 11.1 | 8.2 | 10.2 | 1 | 1 | 1 |
| 0.792 | 0.707 | 43.7 | 37.1 | 34.6 | 30.9 | 1 | 1 | 1 |
| 0.819 | 0.799 | 14.4 | 8.3 | 11.8 | 11.5 | 1 | 1 | 1 |
| 0.882 | 0.756 | 39.7 | 34.1 | 35 | 30 | 1 | 1 | 1 |
| 0.685 | 0.693 | 183.4 | 107.2 | 125.7 | 127.1 | 2 | 2 | 2 |
| 0.813 | 0.758 | 297.2 | 227.2 | 241.7 | 225.3 | 6 | 6 | 6 |
| 0.952 | 0.872 | 70.5 | 68.3 | 67.1 | 61.5 | 1 | 1 | 1 |
| 1.31 | 1.207 | 5.8 | 6.7 | 7.6 | 7 | 1 | 1 | 1 |
| 0.989 | 0.663 | 18.1 | 17.1 | 17.9 | 12 | 1 | 1 | 1 |
| 0.91 | 0.837 | 2478 | 2292.5 | 2254.5 | 2073.3 | 39 | 39 | 39 |
| 0.804 | 0.681 | 159.7 | 144.6 | 128.4 | 108.8 | 2 | 2 | 2 |
| 0.792 | 0.745 | 145.7 | 105.2 | 115.4 | 108.6 | 1 | 1 | 1 |
| 0.817 | 0.756 | 1044.1 | 782.1 | 853.4 | 789.3 | 11 | 11 | 11 |
| 0.596 | 0.567 | 41.6 | 27.4 | 24.8 | 23.6 | 1 | 1 | 1 |
| 0.779 | 0.74 | 101.8 | 76.8 | 79.3 | 75.3 | 1 | 1 | 1 |
| 0.848 | 0.924 | 22.3 | 14.5 | 18.9 | 20.6 | 1 | 1 | 1 |
| 1.078 | 0.747 | 43.8 | 42.4 | 47.2 | 32.7 | 1 | 1 | 1 |

| | | | | | | | | |
|---|---|---|---|---|---|---|---|---|
| 0.81 | 0.731 | 112.1 | 99.6 | 90.8 | 82 | 2 | 2 | 2 |
| 0.753 | 0.661 | 36 | 26.5 | 27.1 | 23.8 | 1 | 1 | 1 |
| 0.763 | 0.722 | 317.4 | 223.3 | 242.1 | 229.1 | 3 | 3 | 3 |
| 0.783 | 0.664 | 422.3 | 316.5 | 330.7 | 280.6 | 5 | 5 | 5 |
| 0.865 | 0.838 | 81.7 | 76.4 | 70.7 | 68.5 | 1 | 1 | 1 |
| 0.845 | 0.79 | 52.8 | 37 | 44.6 | 41.7 | 1 | 1 | 1 |
| 0.738 | 0.7 | 351.9 | 322.8 | 259.6 | 246.4 | 4 | 4 | 4 |
| 0.908 | 0.867 | 12 | 16 | 10.9 | 10.4 | 1 | 1 | 1 |
| 0.896 | 0.671 | 58.9 | 48.3 | 52.8 | 39.5 | 1 | 1 | 1 |
| 0.84 | 0.746 | 81 | 73.6 | 68 | 60.4 | 1 | 1 | 1 |
| 0.869 | 0.837 | 24.5 | 20.5 | 21.3 | 20.5 | 1 | 1 | 1 |
| 0.888 | 0.81 | 116.6 | 118.3 | 103.5 | 94.5 | 1 | 1 | 1 |
| 0.908 | 0.933 | 104.6 | 110 | 95 | 97.6 | 2 | 2 | 2 |
| 0.768 | 0.848 | 73.8 | 64.7 | 56.7 | 62.6 | 2 | 2 | 2 |
| 0.609 | 0.584 | 177.6 | 137.3 | 108.1 | 103.8 | 1 | 1 | 1 |
| 0.971 | 0.794 | 51.9 | 48.2 | 50.4 | 41.2 | 1 | 1 | 1 |
| 0.943 | 0.81 | 159.7 | 158.5 | 150.6 | 129.4 | 5 | 5 | 5 |
| 0.805 | 0.826 | 78.9 | 57.7 | 63.5 | 65.2 | 1 | 1 | 1 |
| 0.831 | 0.752 | 187.7 | 170.8 | 156 | 141.2 | 3 | 3 | 3 |
| 0.839 | 0.814 | 43.5 | 35.9 | 36.5 | 35.4 | 1 | 1 | 1 |
| 0.363 | 0.435 | 218.5 | 102.7 | 79.3 | 95.1 | 1 | 1 | 1 |
| 0.909 | 0.883 | 214.3 | 184.6 | 194.9 | 189.3 | 1 | 1 | 1 |
| 0.842 | 0.692 | 36 | 24.9 | 30.3 | 24.9 | 2 | 2 | 2 |
| 0.852 | 0.889 | 69.5 | 70.1 | 59.2 | 61.8 | 1 | 1 | 1 |
| 0.699 | 0.613 | 55.1 | 39.6 | 38.5 | 33.8 | 1 | 1 | 1 |
| 0.851 | 0.714 | 67.8 | 50.9 | 57.7 | 48.4 | 2 | 2 | 2 |
| 0.627 | 0.653 | 88.4 | 56.9 | 55.4 | 57.7 | 1 | 1 | 1 |
| 0.816 | 0.763 | 411.8 | 303 | 336.1 | 314.3 | 2 | 2 | 2 |
| 1.151 | 0.753 | 7.3 | 4 | 8.4 | 5.5 | 1 | 1 | 1 |
| 0.829 | 0.781 | 65.4 | 50.9 | 54.2 | 51.1 | 2 | 2 | 2 |
| 0.911 | 0.785 | 248.2 | 232.7 | 226.1 | 194.8 | 3 | 3 | 3 |
| 0.847 | 0.759 | 2315 | 2099.5 | 1960.9 | 1758.1 | 25 | 25 | 25 |

| | | | | | | | | |
|---|---|---|---|---|---|---|---|---|
| 0.901 | 0.836 | 255.2 | 262.6 | 229.9 | 213.3 | 1 | 1 | 1 |
| 0.898 | 0.79 | 294.1 | 245.1 | 264.2 | 232.4 | 5 | 5 | 5 |
| 0.641 | 0.765 | 35.7 | 29.6 | 22.9 | 27.3 | 1 | 1 | 1 |
| 0.742 | 0.627 | 172.2 | 122.6 | 127.8 | 108 | 1 | 1 | 1 |
| 0.848 | 0.827 | 260.6 | 201.2 | 220.9 | 215.4 | 5 | 5 | 5 |
| 0.705 | 0.698 | 82 | 64.5 | 57.8 | 57.2 | 1 | 1 | 1 |
| 0.768 | 0.756 | 138.2 | 104.3 | 106.1 | 104.5 | 2 | 2 | 2 |
| 1.008 | 0.818 | 191 | 181.3 | 192.5 | 156.2 | 4 | 4 | 4 |
| 0.713 | 0.765 | 173 | 125.5 | 123.4 | 132.3 | 2 | 2 | 2 |
| 0.648 | 0.704 | 96.9 | 72.6 | 62.8 | 68.2 | 1 | 1 | 1 |
| 0.768 | 0.676 | 97.5 | 72.8 | 74.9 | 65.9 | 1 | 1 | 1 |
| 0.575 | 0.759 | 8.7 | 6.5 | 5 | 6.6 | 1 | 1 | 1 |
| 0.637 | 0.645 | 1263 | 919.7 | 803.9 | 814.9 | 13 | 13 | 13 |
| 0.885 | 0.807 | 192.5 | 177 | 170.4 | 155.3 | 3 | 3 | 3 |
| 0.891 | 0.766 | 118.2 | 121.1 | 105.3 | 90.5 | 3 | 3 | 3 |
| 0.879 | 0.759 | 345.1 | 298.3 | 303.4 | 261.8 | 5 | 5 | 5 |
| 0.899 | 0.66 | 324.6 | 315.6 | 291.8 | 214.2 | 2 | 2 | 2 |
| 0.769 | 0.753 | 82.3 | 76 | 63.3 | 62 | 1 | 1 | 1 |
| 0.693 | 0.656 | 16.3 | 11.4 | 11.3 | 10.7 | 1 | 1 | 1 |
| 0.82 | 0.709 | 54.3 | 54.8 | 44.5 | 38.5 | 1 | 1 | 1 |
| 0.827 | 0.795 | 259.1 | 201.2 | 214.4 | 206 | 2 | 2 | 2 |
| 0.689 | 0.685 | 232.3 | 180.1 | 160 | 159.1 | 2 | 2 | 2 |
| 0.686 | 0.715 | 351.6 | 294.8 | 241.3 | 251.4 | 4 | 4 | 4 |
| 0.706 | 0.554 | 54 | 37.1 | 38.1 | 29.9 | 1 | 1 | 1 |
| 0.696 | 0.646 | 26 | 20.3 | 18.1 | 16.8 | 1 | 1 | 1 |
| 0.822 | 0.751 | 196.4 | 143.9 | 161.5 | 147.4 | 2 | 2 | 2 |
| 0.654 | 0.82 | 28.9 | 21.5 | 18.9 | 23.7 | 1 | 1 | 1 |
| 0.733 | 0.674 | 8.6 | 6.5 | 6.3 | 5.8 | 1 | 1 | 1 |
| 0.783 | 0.742 | 548.7 | 432.7 | 429.8 | 407 | 5 | 5 | 5 |
| 0.561 | 0.55 | 89.3 | 74.5 | 50.1 | 49.1 | 2 | 2 | 2 |
| 0.907 | 0.789 | 429.7 | 364.4 | 389.7 | 338.9 | 4 | 4 | 4 |

| Abundances Count: F2: 129, Sample | emPAI | Score Sequest HT | # Peptides Sequest HT | Score Mascot | # Peptides Mascot |
|---|---|---|---|---|---|
| 7 | 1.336 | 67.2704573 | 12 | 435.369954 | 12 |
| 3 | 55.234 | 116.948996 | 17 | 443.92572 | 17 |
| | 0.058 | 2.6680634 | 1 | 0 | 1 |
| 1 | 0.064 | 0 | 1 | 0 | 1 |
| 1 | 0.136 | 2.52889824 | 1 | 0 | 1 |
| 1 | 0.369 | 7.95092702 | 3 | 51.2953576 | 3 |
| 1 | 0.995 | 4.88091946 | 3 | 27.22 | 3 |
| 1 | 0.233 | 2.95353675 | 1 | 31.29 | 1 |
| | 0.292 | 5.16537809 | 1 | 62.3 | 1 |
| 8 | 4.179 | 136.963808 | 30 | 811.768702 | 30 |
| 9 | 5.494 | 43.3620131 | 10 | 127.447594 | 9 |
| 2 | 1.154 | 8.8761344 | 2 | 81.6270762 | 2 |
| 2 | 0.389 | 11.8906677 | 4 | 80.0990659 | 4 |
| 7 | 13.251 | 48.1256802 | 13 | 325.950418 | 13 |
| 1 | 0.179 | 5.47677231 | 1 | 33.67 | 1 |
| 1 | 0.468 | 6.97250652 | 2 | 0 | 2 |
| | 0.585 | 2.99119473 | 1 | 0 | 1 |
| 1 | 0.186 | 4.85131884 | 2 | 67.35 | 2 |
| | 0.166 | 1.66663337 | 1 | | |
| 1 | 0.468 | 10.7276418 | 3 | 71.2964165 | 3 |
| 5 | 1.438 | 49.4954681 | 10 | 302.00703 | 10 |
| 3 | 1.683 | 32.5879722 | 8 | 156.10536 | 8 |
| 2 | 1.683 | 25.6133728 | 7 | 134.14 | 7 |
| 3 | 0.557 | 8.6690793 | 3 | 58.1739597 | 4 |
| 2 | 0.216 | 10.6795108 | 4 | 46.8043805 | 4 |
| 7 | 3.806 | 84.1911628 | 12 | 422.096404 | 12 |
| 1 | 0.413 | 10.5738263 | 3 | 63.2999303 | 3 |
| 4 | 2.631 | 74.6511772 | 10 | 507.002804 | 10 |
| 1 | 0.668 | 22.6568878 | 3 | 207.74979 | 3 |
| 5 | 0.823 | 20.8007877 | 6 | 151.31 | 6 |
| | 9 | 21.3149526 | 4 | 160.60531 | 4 |
| 4 | 20.544 | 18.8681724 | 7 | 87.74 | 7 |
| 3 | 3.217 | 19.0098178 | 4 | 104.332307 | 4 |
| | 0.274 | 5.44620109 | 2 | 0 | 2 |
| 3 | 4.623 | 16.9399357 | 6 | 66.8 | 6 |
| | 0.066 | 2.01075053 | 1 | | |
| | 0.778 | | | 41.63 | 1 |
| 1 | 0.468 | 7.60506511 | 1 | 40.48 | 1 |
| 1 | 0.292 | 3.998878 | 1 | 40.9 | 1 |
| | 2.162 | 18.771455 | 5 | 139.217747 | 5 |
| 2 | 0.274 | 7.57760763 | 2 | 37.6668761 | 2 |
| | 0.056 | 2.99518228 | 1 | 56.64 | 1 |
| 1 | 0.259 | 2.58011079 | 1 | 50.21 | 1 |
| 1 | 0.52 | 7.87093234 | 2 | 33.58 | 2 |
| | 0.093 | 7.62195325 | 2 | 0 | 2 |
| | 0.468 | | | 0 | 1 |
| 10 | 13.251 | 75.354883 | 10 | 531.345932 | 10 |
| 5 | 2.311 | 50.3899174 | 11 | 184.444285 | 11 |
| | 0.116 | 6.71991444 | 1 | 109.13 | 1 |
| | 0.145 | 3.25001049 | 1 | 52.33 | 1 |
| 1 | 0.668 | 11.304749 | 1 | 33.67 | 1 |
| 1 | 0.438 | 5.7132535 | 3 | 22.89 | 3 |
| | 0.259 | 2.87085152 | 1 | 42.23 | 1 |
| | 0.212 | 3.18665385 | 1 | 32.81 | 1 |
| 1 | 0.179 | 0 | 1 | 0 | 1 |
| 1 | 0.304 | 16.9943097 | 3 | 125.593333 | 3 |
| 2 | 1.276 | 22.0571527 | 5 | 148.971913 | 5 |

| | | | | | |
|---|---|---|---|---|---|
| 11 | 23.245 | 79.1718603 | 13 | 387.117059 | 13 |
| 2 | 2.162 | 16.0495622 | 5 | 67.27 | 5 |
| | 0.292 | 2.6106267 | 1 | 0 | 1 |
| | 0.292 | 2.61410904 | 1 | 38.58 | 1 |
| | 0.389 | 0 | 1 | 0 | 1 |
| | 0.468 | 2.84894848 | 1 | 0 | 1 |
| 1 | 3.642 | 6.00894856 | 2 | 0 | 2 |
| | 0.122 | 2.27747059 | 1 | 0 | 1 |
| | 0.585 | 7.72578621 | 2 | 33.56 | 2 |
| 1 | 0.233 | 4.00517273 | 1 | 43.64 | 1 |
| | 1.154 | 6.84600592 | 2 | 40.58 | 2 |
| | 0.064 | 1.91498566 | 1 | 33.34 | 1 |
| 1 | 0.259 | 2.33512115 | 1 | 0 | 1 |
| 1 | 1.783 | 3.67449045 | 4 | 44.18 | 4 |
| 1 | 0.874 | 4.97390461 | 3 | 44.84 | 3 |
| 1 | 0.995 | 8.64385343 | 3 | 55.36 | 3 |
| | 1.154 | 2.71901822 | 1 | 49.94 | 1 |
| 2 | 0.311 | 5.08998823 | 2 | 0 | 2 |
| | 0.359 | 3.16476083 | 2 | 37.87 | 2 |
| 1 | 1.848 | 9.28924942 | 4 | 85.0419176 | 4 |
| 1 | 0.585 | 7.41125035 | 1 | 72.8198488 | 1 |
| 2 | 0.389 | 9.72164869 | 4 | 44.9070028 | 4 |
| 15 | 3.977 | 162.585695 | 21 | 588.575048 | 21 |
| 2 | 0.468 | 29.0514436 | 7 | 132.596667 | 7 |
| 2 | 643.947 | 415.173548 | 28 | 2179.87621 | 27 |
| 1 | 0.233 | 2.77372193 | 1 | 52.67 | 1 |
| 2 | 0.995 | 14.9718595 | 3 | 115.219374 | 3 |
| 7 | 4.08 | 62.2015095 | 11 | 352.038235 | 10 |
| | 0.066 | 3.16472673 | 1 | 0 | 1 |
| 3 | 6.197 | 38.3828602 | 8 | 157.310537 | 9 |
| 7 | 4.623 | 39.9072464 | 9 | 159.961405 | 10 |
| | 0.202 | 6.0714407 | 2 | 33.89 | 2 |
| 1 | 0.093 | 3.08602929 | 1 | 0 | 1 |
| 1 | 0.585 | 10.5693154 | 2 | 107.315984 | 2 |
| | 0.334 | 0 | 1 | 0 | 1 |
| 1 | 0.118 | 8.18072915 | 3 | 0 | 3 |
| | 0.668 | 4.39697933 | 2 | 32.17 | 2 |
| 1 | 0.077 | 3.58720183 | 1 | 0 | 1 |
| | 0.233 | 2.62356901 | 1 | 38.76 | 1 |
| 1 | 0.468 | 3.39721775 | 1 | 34.15 | 1 |
| | 0.116 | 9.08382082 | 1 | 46.84 | 1 |
| 2 | 0.52 | 6.03944111 | 2 | 61.67 | 2 |
| 1 | 0.468 | | | 0 | 1 |
| | 0.259 | 3.09671354 | 1 | 38.76 | 1 |
| 1 | 0.334 | 2.33684874 | 1 | 35.11 | 1 |
| 1 | 0.334 | 5.85664058 | 2 | 0 | 2 |
| 1 | 0.145 | 2.78646636 | 1 | 26.45 | 1 |
| 2 | 0.848 | 9.755687 | 4 | 41.8991236 | 4 |
| 1 | 0.175 | 14.9594288 | 4 | 182.352156 | 4 |
| 2 | 0.202 | 2.91274452 | 2 | 0 | 2 |
| 1 | 0.501 | 12.7908282 | 3 | 100.054625 | 3 |
| | 0.425 | 5.60547256 | 2 | 50.71 | 2 |
| 2 | 1.154 | 23.7750275 | 5 | 98.7716922 | 5 |
| | 0.468 | 2.79781556 | 1 | 48.29 | 1 |
| 4 | 2.728 | 24.5749226 | 8 | 148.614065 | 8 |
| | 0.233 | 5.91840935 | 2 | 37.74 | 1 |
| 1 | 0.179 | 4.5569787 | 1 | 53.7 | 1 |
| | 0.212 | 2.90940785 | 1 | 0 | 1 |
| 1 | 0.848 | 16.3652978 | 3 | 126.202163 | 3 |
| | 0.145 | 2.35289335 | 1 | 0 | 1 |
| 1 | 0.389 | 2.22309518 | 2 | 0 | 2 |
| 2 | 0.259 | 8.51152563 | 3 | 60.5003405 | 3 |

| | | | | | |
|---|---|---|---|---|---|
| 1 | 0.292 | 3.11250401 | 1 | 41.44 | 1 |
| 2 | 0.668 | 7.55470634 | 2 | 52.82 | 2 |
| | 0.212 | 9.31755471 | 2 | 115.606383 | 2 |
| 1 | 1.738 | 48.2086678 | 13 | 286.700925 | 13 |
| 1 | 1.154 | 1.94615793 | 2 | 0 | 2 |
| 3 | 2.162 | 20.5110652 | 6 | 88.6881691 | 6 |
| | 0.055 | 3.36864638 | 1 | 23.36 | 1 |
| 2 | 0.624 | 24.794378 | 4 | 212.78 | 4 |
| 2 | 0.638 | 18.0682449 | 3 | 64.4395368 | 3 |
| 1 | 0.823 | 16.8629289 | 5 | 118.217162 | 5 |
| | 0.413 | 2.92475939 | 2 | 0 | 2 |
| 2 | 2.162 | 18.7014577 | 4 | 121.104331 | 3 |
| | 0.585 | 0 | 1 | 0 | 1 |
| 1 | 0.116 | 5.43685579 | 1 | 42.47 | 1 |
| 2 | 0.778 | 16.6239278 | 5 | 110.179154 | 5 |
| | 0.389 | 3.53681564 | 1 | 42.64 | 1 |
| 3 | 2.981 | 17.7053213 | 6 | 116.692044 | 6 |
| 1 | 0.248 | 14.4683187 | 5 | 77.2716624 | 4 |
| 10 | 6.305 | 77.5260603 | 16 | 310.679506 | 16 |
| | 0.245 | 5.62384152 | 2 | 55.12 | 2 |
| 1 | 0.413 | 8.58658719 | 3 | 36.77 | 3 |
| 1 | 0.116 | 3.6226058 | 1 | 45.63 | 1 |
| 1 | 643.947 | 395.802438 | 28 | 1907.58456 | 28 |
| 1 | 0.585 | 8.88878632 | 3 | 69.23 | 3 |
| 1 | 2.162 | 15.7673392 | 5 | 85.7412323 | 5 |
| | 0.259 | 0 | 1 | 0 | 1 |
| 1 | 0.179 | 8.11043382 | 2 | 103.287791 | 2 |
| 1 | 0.425 | 6.87535 | 2 | 64.3475584 | 2 |
| | 0.334 | 15.820843 | 4 | 119.21 | 4 |
| | 0.241 | 7.81844258 | 3 | 27.33 | 3 |
| 1 | 0.413 | 7.78494596 | 3 | 39.9667571 | 2 |
| 3 | 0.778 | 7.03916693 | 4 | 71.5453688 | 4 |
| | 0.089 | 0 | 1 | | |
| 4 | 1.069 | 10.7722354 | 6 | 39.99 | 5 |
| 2 | 2.981 | 4.71390009 | 2 | 0 | 2 |
| | 0.292 | 4.20660925 | 1 | 38.45 | 1 |
| | 0.212 | 2.78055 | 1 | 39.72 | 1 |
| 1 | 0.52 | 7.36280036 | 2 | 59.11 | 1 |
| 8 | 7.913 | 118.25684 | 11 | 867.210151 | 12 |
| 1 | 0.389 | 3.99091411 | 1 | 49.3 | 2 |
| 1 | 0.501 | 7.44258642 | 3 | 36.22 | 3 |
| 6 | 4.08 | 41.8671124 | 11 | 266.193883 | 12 |
| 3 | 4.179 | 38.1359835 | 5 | 168.952309 | 6 |
| 3 | 1.31 | 29.3012664 | 3 | 260.789851 | 3 |
| 2 | 0.425 | 3.3350575 | 2 | 0 | 2 |
| | 2.162 | 3.41757512 | 1 | 47.28 | 1 |
| 1 | 0.778 | | | 60.32 | 1 |
| 9 | 1.929 | 89.0428603 | 20 | 376.953314 | 20 |
| 2 | 2.162 | 17.2473586 | 3 | 32.67 | 3 |
| 1 | 0.668 | 5.33365393 | 2 | 37.92 | 2 |
| 4 | 0.778 | 11.1943822 | 5 | 62.5033333 | 4 |
| | 1.346 | 54.089968 | 7 | 399.91643 | 7 |
| 7 | 4.248 | 56.9372535 | 14 | 267.568608 | 13 |
| 1 | 0.425 | 6.83213449 | 2 | 79.38 | 2 |
| | 0.11 | 3.65441656 | 1 | 0 | 1 |
| 2 | 0.668 | 10.1063802 | 4 | 49.8 | 4 |
| 1 | 0.199 | 12.6704326 | 3 | 140.98 | 3 |
| 1 | 0.292 | 4.28081465 | 1 | 46.09 | 1 |
| 1 | 0.233 | 6.62466693 | 2 | 42.5439039 | 2 |
| | 0.259 | 0 | 1 | 0 | 1 |
| 1 | 0.52 | 5.38867188 | 1 | 0 | 1 |
| | 0.189 | 8.03948951 | 3 | 0 | 3 |

| | | | | | |
|---|---|---|---|---|---|
| 1 | 0.304 | 8.91364932 | 3 | 39.75 | 3 |
| 2 | 0.259 | 10.6715944 | 2 | 63.6959898 | 2 |
| | 0.11 | 2.25161409 | 1 | 0 | 1 |
| 1 | 1.154 | 13.2626188 | 3 | 68.8816674 | 2 |
| 2 | 0.778 | 10.793427 | 3 | 58.43 | 3 |
| | 0.468 | 3.70721698 | 1 | 47.27 | 1 |
| | 0.194 | 2.42482924 | 1 | 0 | 1 |
| | 0.136 | 2.11939216 | 1 | | |
| 1 | 0.11 | 2.60521913 | 1 | 0 | 1 |
| | 0.468 | 2.48700285 | 1 | 40.41 | 1 |
| 2 | 0.52 | 17.1977522 | 6 | 86.8525686 | 6 |
| 2 | 2.162 | 11.7409644 | 3 | 55.3298746 | 3 |
| 2 | 0.245 | 7.60120249 | 2 | 62.2670035 | 2 |
| | 0.778 | 24.4931803 | 5 | 117.441159 | 5 |
| | 0.212 | 5.04481602 | 2 | 37.47 | 2 |
| | 0.369 | 10.6918142 | 2 | 109.649313 | 2 |
| | 0.292 | 2.94498229 | 1 | 0 | 1 |
| 10 | 4.456 | 60.2100277 | 11 | 385.949735 | 11 |
| 1 | 2.728 | 18.5569854 | 3 | 177.706647 | 3 |
| 1 | 0.585 | 13.3313448 | 3 | 0 | 2 |
| | 0.179 | 0 | 1 | 0 | 1 |
| 1 | 2.162 | 0 | 1 | 0 | 1 |
| 6 | 24.119 | 50.4351246 | 8 | 89.3352037 | 9 |
| | 0.425 | 5.93790317 | 2 | 50.74 | 2 |
| | 0.468 | 4.12171125 | 1 | 68.98 | 1 |
| 1 | 0.668 | 8.66248083 | 4 | 0 | 4 |
| 1 | 0.155 | 4.21570492 | 1 | 31.26 | 1 |
| 1 | 2.162 | 0 | 1 | 0 | 1 |
| | 5.813 | 15.5957198 | 4 | 57.6501925 | 4 |
| 1 | 0.194 | 2.35226655 | 2 | 35.81 | 2 |
| 1 | 0.292 | 10.1291602 | 3 | 71.5588595 | 3 |
| 1 | 0.212 | 8.06910443 | 2 | 34.14 | 2 |
| 1 | 0.778 | 1.9528625 | 1 | 0 | 1 |
| 1 | 0.327 | 25.9943998 | 6 | 175.777143 | 6 |
| 2 | 0.254 | 23.1512091 | 6 | 88.9535262 | 6 |
| | 0.11 | 4.09314394 | 1 | 33.49 | 1 |
| | 0.078 | 6.51160526 | 2 | 0 | 2 |
| | 0.166 | 3.96340156 | 1 | 49.24 | 1 |
| | 0.585 | 3.58532095 | 1 | 0 | 1 |
| 1 | 0.245 | 3.94992542 | 2 | 0 | 2 |
| 2 | 0.438 | 10.8997798 | 3 | 0 | 2 |
| 1 | 0.425 | 3.4456625 | 2 | 67.97 | 2 |
| | 0.048 | 2.45028329 | 1 | 49.02 | 1 |
| 1 | 0.292 | 4.82647848 | 2 | 61.58 | 2 |
| 2 | 0.292 | 9.95713735 | 3 | 111.999925 | 3 |
| 3 | 23.041 | 154.372475 | 16 | 633.46443 | 15 |
| 8 | 1.848 | 42.3375632 | 12 | 234.048033 | 12 |
| | 0.389 | 2.382195 | 1 | 33.73 | 1 |
| 1 | 0.212 | 2.5907836 | 1 | 33.92 | 1 |
| | 0.585 | 0 | 1 | 0 | 1 |
| 1 | 0.136 | 3.15668607 | 1 | 41.46 | 1 |
| | 0.334 | 2.34146118 | 1 | 0 | 1 |
| | 0.425 | 2.10364294 | 2 | 0 | 2 |
| | 0.52 | 7.44873905 | 2 | 0 | 2 |
| 2 | 0.668 | 7.81547785 | 2 | 88.56 | 2 |
| 2 | 0.778 | 6.7680223 | 2 | 42.18 | 3 |
| 1 | 0.259 | 3.94285727 | 1 | 54.04 | 1 |
| 1 | 0.468 | 11.3159723 | 2 | 82.9416773 | 2 |
| | 2.981 | 8.93290925 | 3 | 101.179501 | 2 |
| | 0.212 | 2.37705183 | 1 | 0 | 1 |
| 1 | 0.585 | 13.5360348 | 2 | 33.95 | 2 |
| 9 | 3.87 | 43.7883406 | 11 | 249.385705 | 10 |

| | | | | | |
|---|---|---|---|---|---|
| 1 | 2.162 | 2.83147597 | 1 | 37.17 | 1 |
| 5 | 2.384 | 63.8571775 | 17 | 359.149889 | 17 |
| 1 | 0.311 | 6.08251858 | 2 | 35.09 | 2 |
| 1 | 0.931 | 6.07800078 | 2 | 53.77 | 2 |
| | 0.179 | 4.66852617 | 1 | 68.25 | 1 |
| 1 | 1.683 | 9.75116444 | 3 | 81.78 | 3 |
| 1 | 2.162 | 4.03552675 | 1 | 107.002903 | 2 |
| | 0.931 | 2.97499013 | 2 | 34.21 | 2 |
| 1 | 0.292 | 0 | 1 | 0 | 1 |
| 6 | 32.246 | 183.023727 | 20 | 966.449332 | 20 |
| 2 | 0.995 | 10.4995027 | 3 | 96.871102 | 3 |
| 9 | 43.893 | 266.599284 | 23 | 1660.42383 | 23 |
| 1 | 1.512 | 4.65858507 | 1 | 0 | 1 |
| | 0.11 | 8.17287803 | 2 | 81.99 | 2 |
| | 0.067 | 5.20134234 | 2 | 59.23 | 2 |
| 1 | 2.981 | 10.3672752 | 2 | 47.3393539 | 2 |
| | 0.389 | 2.34130335 | 1 | 22.97 | 1 |
| 1 | 0.155 | 4.84774494 | 1 | 73.52 | 1 |
| 3 | 2.162 | 5.08979893 | 3 | 25.29 | 3 |
| 3 | 4.623 | 25.9458177 | 8 | 97.4039812 | 7 |
| | 0.042 | 3.38071561 | 1 | 0 | 1 |
| | 0.304 | 19.6575074 | 3 | 164.148555 | 3 |
| 4 | 0.668 | 32.5870287 | 10 | 260.113703 | 10 |
| 5 | 1.202 | 39.120523 | 10 | 259.182436 | 10 |
| 11 | 2.775 | 77.7231414 | 11 | 604.643567 | 12 |
| 3 | 1.254 | 21.7801371 | 6 | 92.5406284 | 6 |
| | 0.119 | 13.8342485 | 2 | 113.323333 | 2 |
| 1 | 0.03 | 0 | 1 | 0 | 1 |
| 2 | 0.931 | 11.8550892 | 4 | 73.01 | 4 |
| 1 | 0.086 | 2.66447663 | 1 | 36.79 | 1 |
| | 0.083 | 3.73661709 | 1 | 47.98 | 1 |
| | 0.848 | 18.7771277 | 3 | 74.3636234 | 3 |
| | 0.172 | 5.71420813 | 2 | 0 | 1 |
| 1 | 0.116 | 2.43707204 | 1 | 37.09 | 1 |
| | 0.077 | 0 | 1 | 0 | 1 |
| 12 | 2.511 | 82.4648323 | 16 | 336.810836 | 16 |
| 1 | 0.65 | 20.8560927 | 4 | 121.099812 | 4 |
| | 0.136 | 6.00108099 | 2 | 36.15 | 2 |
| 1 | 0.218 | 10.3621089 | 3 | 75.51 | 3 |
| 2 | 0.318 | 8.57493496 | 3 | 46.57 | 3 |
| 9 | 4.135 | 108.972057 | 24 | 539.432739 | 22 |
| 1 | 0.778 | 6.98428035 | 3 | 23.76 | 3 |
| | 0.259 | 3.05669856 | 1 | 38.79 | 1 |
| | 0.233 | 0 | 1 | 0 | 1 |
| 5 | 4.337 | 17.6946347 | 7 | 58.52 | 7 |
| 8 | 0.701 | 35.227452 | 9 | 212.299866 | 9 |
| | 0.274 | 5.72547817 | 1 | 92.85 | 1 |
| 1 | 0.194 | 8.17713046 | 2 | 55.05 | 2 |
| 8 | 4.766 | 151.447897 | 27 | 869.323362 | 26 |
| | 0.105 | 2.2591784 | 1 | 17.02 | 1 |
| 1 | 0.372 | 23.7504451 | 7 | 121.20187 | 7 |
| 2 | 0.433 | 24.6408162 | 5 | 119.311662 | 5 |
| 2 | 1.031 | 14.1007736 | 4 | 128.08116 | 4 |
| | 0.179 | 4.02050638 | 1 | 53.71 | 1 |
| 1 | 0.269 | 9.395015 | 3 | 22.86 | 3 |
| 3 | 0.509 | 15.4577627 | 5 | 81.4241211 | 5 |
| 1 | 0.109 | 11.2072706 | 3 | 51.6392199 | 3 |
| 1 | 0.11 | 2.70206833 | 1 | 33.83 | 2 |
| 7 | 9 | 45.4461691 | 10 | 224.523339 | 11 |
| 12 | 11.915 | 122.723128 | 10 | 498.182722 | 10 |
| | 0.292 | 3.15300393 | 1 | 29.76 | 1 |
| | 0.145 | 2.77420616 | 1 | 0 | 1 |

| | | | | | |
|---|---|---|---|---|---|
| 1 | 0.194 | 2.84208107 | 1 | 52.77 | 1 |
| | 0.292 | 3.95871902 | 1 | 50.66 | 1 |
| 10 | 5.449 | 80.3533947 | 13 | 530.435973 | 13 |
| | 0.334 | 3.03095579 | 1 | 47.03 | 1 |
| | 0.233 | 5.89411306 | 2 | 0 | 2 |
| | 0.155 | 2.5557344 | 1 | 35.19 | 1 |
| 1 | 0.274 | 5.30805373 | 2 | 35.07 | 2 |
| 1 | 0.585 | 8.76929903 | 2 | 128.27 | 2 |
| | 0.166 | 3.42646956 | 1 | 35.54 | 1 |
| 1 | 0.233 | 3.0307374 | 1 | 40.57 | 1 |
| 5 | 0.805 | 37.9450294 | 9 | 267.347738 | 9 |
| 1 | 0.129 | 2.944175 | 1 | 40.96 | 1 |
| 1 | 0.311 | 11.7490585 | 4 | 127.969678 | 4 |
| 2 | 0.212 | 13.8619843 | 3 | 119.015283 | 3 |
| 1 | 0.222 | 3.02822757 | 2 | 41.06 | 2 |
| 1 | 0.778 | 7.95976663 | 2 | 43.37 | 2 |
| 2 | 0.222 | 8.67619991 | 2 | 75.4281012 | 2 |
| | 0.089 | 3.4402492 | 1 | 42.32 | 1 |
| 1 | 0.101 | 5.31744432 | 1 | 44.03 | 1 |
| 3 | 0.271 | 22.7695284 | 5 | 120.342305 | 5 |
| | 0.047 | 3.2818644 | 1 | 0 | 1 |
| | 0.259 | 3.24704003 | 1 | 0 | 1 |
| 1 | 0.304 | 7.52421212 | 3 | 0 | 3 |
| | 0.08 | 5.35289025 | 1 | 38.8 | 1 |
| 12 | 8.12 | 85.2902987 | 19 | 490.229713 | 19 |
| | 0.116 | 5.23925257 | 2 | 0 | 2 |
| | 0.212 | 4.10805416 | 1 | 42.18 | 1 |
| 3 | 0.445 | 8.70372677 | 4 | 43.8408497 | 4 |
| 1 | 0.52 | 10.9847257 | 1 | 63.49 | 1 |
| 1 | 0.105 | 2.99745822 | 1 | 0 | 1 |
| 2 | 576.969 | 408.211788 | 28 | 2157.18068 | 27 |
| 5 | 463.159 | 379.77387 | 27 | 1779.17657 | 27 |
| 3 | 0.389 | 10.714335 | 4 | 82.6464373 | 4 |
| | 0.053 | 2.9323175 | 1 | 34.83 | 1 |
| | 0.08 | 3.66989231 | 1 | 58.53 | 1 |
| 2 | 0.304 | 19.3186259 | 6 | 60.7801923 | 6 |
| | 0.468 | 5.06759405 | 2 | 0 | 2 |
| 14 | 4.722 | 91.4805214 | 21 | 566.577181 | 20 |
| 1 | 0.122 | 2.47173548 | 1 | 33.39 | 1 |
| 2 | 0.417 | 15.8480756 | 5 | 98.4733333 | 5 |
| | 0.036 | 2.61119175 | 1 | 0 | 1 |
| 26 | 38.811 | 238.0499 | 25 | 1395.12237 | 25 |
| | 0.044 | 2.38536596 | 1 | 0 | 1 |
| | 0.202 | 6.09134722 | 2 | 41.56 | 2 |
| 1 | 0.668 | 17.228955 | 7 | 80.4497417 | 7 |
| 2 | 0.73 | 19.087826 | 4 | 102.295726 | 4 |
| | 0.03 | 5.83374 | 3 | 27.4742942 | 2 |
| 12 | 0.269 | 85.1704812 | 25 | 356.811354 | 27 |
| | 0.233 | 2.77151012 | 1 | 18.69 | 1 |
| 2 | 0.719 | 13.8043051 | 3 | 114.565395 | 3 |
| 2 | 0.585 | 8.30438495 | 2 | 88.76 | 2 |
| | 1.276 | 18.8505993 | 5 | 103.940633 | 5 |
| 3 | 0.346 | 10.1319041 | 4 | 85.7755229 | 4 |
| 1 | 0.283 | 11.6629643 | 4 | 103.821083 | 4 |
| 1 | 0.077 | 5.60954452 | 2 | 32.94 | 2 |
| | 0.233 | 2.91735387 | 1 | 41.98 | 1 |
| 5 | 0.643 | 43.2902539 | 11 | 252.731368 | 11 |
| 4 | 0.688 | 18.5500114 | 5 | 102.105344 | 5 |
| 3 | 0.322 | 6.06761289 | 3 | 43.17 | 4 |
| 2 | 0.413 | 13.3988264 | 3 | 130.771987 | 3 |
| 1 | 0.166 | 2.31633544 | 1 | 44.72 | 1 |
| 1 | 0.116 | 0 | 1 | 0 | 1 |

| | | | | | |
|---|---|---|---|---|---|
| | 0.116 | 2.36290693 | 1 | 39.11 | 1 |
| 1 | 0.083 | 3.48586631 | 1 | 0 | 1 |
| | 0.136 | 3.01820898 | 1 | 0 | 1 |
| 1 | 0.225 | 10.7299643 | 3 | 80.0836614 | 3 |
| | 0.859 | 22.5568531 | 7 | 108.252061 | 6 |
| | 0.096 | 3.11856771 | 1 | 36.46 | 1 |
| | 0.07 | 2.14582634 | 1 | 0 | 1 |
| | 0.179 | 2.08431125 | 1 | 0 | 1 |
| | 0.212 | 2.30735683 | 1 | 29.06 | 1 |
| 2 | 0.52 | 8.35164475 | 2 | 43.55 | 2 |
| 1 | 0.668 | 4.87775207 | 2 | 23.07 | 2 |
| 2 | 0.468 | 10.674274 | 2 | 133.315707 | 2 |
| 3 | 1.081 | 26.4916465 | 6 | 196.153508 | 6 |
| | 0.136 | 2.83032298 | 1 | 0 | 1 |
| 2 | 0.701 | 23.9170313 | 6 | 212.919753 | 6 |
| 4 | 0.374 | 29.0539346 | 8 | 141.045472 | 8 |
| 1 | 0.374 | 10.5030456 | 4 | 95.5963741 | 4 |
| 1 | 0.039 | 2.45995212 | 1 | 29.28 | 1 |
| 3 | 9 | 13.9026611 | 3 | 62.1308509 | 3 |
| 1 | 0.274 | 11.7085211 | 4 | 38.7798138 | 4 |
| 1 | 0.359 | 5.24368453 | 2 | 0 | 2 |
| 1 | 0.245 | 8.08206725 | 2 | 37.45 | 2 |
| | 0.468 | 10.6943254 | 2 | 95.7754523 | 2 |
| 31 | 21.067 | 243.980457 | 31 | 1465.77393 | 31 |
| | 0.194 | 2.42859721 | 1 | 31.38 | 1 |
| 1 | 0.468 | 4.03324604 | 1 | 45.48 | 1 |
| | 0.311 | 7.00980115 | 2 | 57.53 | 2 |
| 1 | 0.116 | 2.10775638 | 1 | 28.56 | 1 |
| | 0.389 | 0 | 1 | 0 | 1 |
| 7 | 154.052 | 323.145932 | 27 | 1636.08361 | 26 |
| | 0.778 | 3.31161141 | 1 | 49.27 | 1 |
| | 0.292 | 3.00719285 | 1 | 46.08 | 1 |
| | 0.11 | 3.4826293 | 1 | 0 | 1 |
| 1 | 0.129 | 5.21700096 | 1 | 44.18 | 1 |
| 2 | 0.292 | 11.3224118 | 3 | 70.4020329 | 3 |
| 1 | 0.145 | 3.34077382 | 1 | 30.54 | 1 |
| 2 | 1.818 | 37.6878641 | 8 | 125.889582 | 8 |
| 1 | 0.11 | 3.76057029 | 1 | 78.11 | 1 |
| 1 | 0.212 | 2.34215593 | 1 | 32.92 | 1 |
| 3 | 1.276 | 8.47966599 | 3 | 70.142663 | 4 |
| 1 | 0.52 | 8.83701634 | 2 | 54.9087368 | 2 |
| | 0.093 | 4.4852767 | 1 | 0 | 1 |
| | 0.212 | 6.80972552 | 1 | 69.7407664 | 1 |
| 2 | 0.389 | 21.8345037 | 5 | 130.620757 | 5 |
| 1 | 0.059 | 4.17307568 | 1 | 0 | 1 |
| 1 | 0.075 | 5.40569544 | 1 | 75.07 | 1 |
| | 0.668 | 18.7724864 | 4 | 134.0454 | 4 |
| 1 | 0.202 | 12.3419273 | 2 | 131.573814 | 2 |
| | 0.129 | 2.89132619 | 1 | 37.88 | 1 |
| 1 | 0.083 | 2.83026791 | 1 | 37.99 | 1 |
| 1 | 0.259 | 3.30350971 | 2 | 33.38 | 2 |
| 5 | 0.947 | 51.6541739 | 9 | 320.400885 | 9 |
| 3 | 0.655 | 21.2835171 | 7 | 147.111814 | 7 |
| | 0.155 | 3.13832569 | 1 | 32.16 | 1 |
| 3 | 1.848 | 10.1874959 | 4 | 52.91069 | 5 |
| 2 | 0.688 | 15.1137066 | 3 | 55.8882253 | 3 |
| 2 | 0.701 | 8.41673899 | 2 | 90.32 | 2 |
| | 0.585 | 5.04968214 | 1 | 51.99 | 1 |
| | 0.083 | 2.3712368 | 1 | 0 | 1 |
| | 0.194 | 3.50111771 | 1 | 60.6 | 1 |
| 1 | 0.136 | 4.54948807 | 1 | 36.71 | 1 |
| | 0.068 | 2.33716679 | 1 | 0 | 1 |

| | | | | | |
|---|---|---|---|---|---|
| | 0.17 | 8.96251893 | 3 | 40.51 | 3 |
| | 0.233 | 7.59831333 | 1 | 31.24 | 1 |
| 7 | 1.219 | 32.888284 | 8 | 140.811287 | 8 |
| | 0.145 | 5.79421592 | 2 | 22.27 | 2 |
| 1 | 0.334 | 5.56303191 | 1 | 24.26 | 1 |
| 1 | 0.061 | 2.8827188 | 1 | 50.49 | 1 |
| 1 | 0.194 | 2.31561589 | 1 | 29.77 | 1 |
| | 0.468 | 3.67112994 | 1 | 58.83 | 1 |
| 1 | 0.038 | 3.13807726 | 1 | 35.28 | 1 |
| 1 | 0.585 | 2.81249142 | 1 | 22.16 | 1 |
| 2 | 0.35 | 12.0938325 | 3 | 84.9946629 | 3 |
| | 0.202 | 5.13912773 | 2 | 49.76 | 2 |
| 2 | 0.318 | 15.4872262 | 2 | 156.473911 | 2 |
| 1 | 0.012 | 3.38156605 | 1 | 40.89 | 1 |
| 2 | 0.079 | 11.282443 | 3 | 55.6726881 | 3 |
| 1 | 0.155 | 3.36210847 | 1 | 41.51 | 1 |
| 1 | 0.389 | 5.67072535 | 2 | 32.73 | 2 |
| 6 | 2.162 | 61.4679284 | 11 | 203.613986 | 11 |
| 1 | 0.131 | 31.7564149 | 6 | 217.6718 | 6 |
| 1 | 1.031 | 14.4201381 | 4 | 110.79 | 4 |
| 1 | 0.468 | 3.5521605 | 2 | 60.11 | 2 |
| 1 | 0.389 | 0 | 1 | 0 | 1 |
| | 0.026 | 2.99115682 | 1 | 47.42 | 1 |
| 15 | 3.084 | 66.9985704 | 17 | 340.377478 | 18 |
| 1 | 0.585 | 4.96056128 | 3 | 56.46 | 3 |
| | 0.045 | 2.70056367 | 1 | 31.91 | 1 |
| 1 | 0.11 | 4.13846779 | 1 | 41.23 | 1 |
| | 0.179 | 4.04251766 | 1 | 40.05 | 1 |
| 27 | 1.596 | 200.669922 | 54 | 995.592429 | 54 |
| 5 | 3.87 | 24.988059 | 11 | 60.85 | 10 |
| 2 | 0.101 | 5.94501448 | 2 | 58.9340799 | 3 |
| 2 | 0.468 | 0 | 3 | 0 | 3 |
| 1 | 0.259 | 8.39853501 | 2 | 74.830227 | 2 |
| 1 | 0.425 | 7.21383739 | 2 | 40.5991025 | 2 |
| 1 | 0.417 | 15.452713 | 4 | 115.126912 | 4 |
| 1 | 0.233 | 3.21004295 | 1 | 44.54 | 1 |
| 1 | 0.105 | 6.35653901 | 2 | 74.26 | 2 |
| | 0.051 | 2.59847546 | 1 | 24.05 | 1 |
| | 0.194 | 3.83675694 | 1 | 57.79 | 1 |
| 1 | 0.054 | 4.6745429 | 1 | 63.64 | 1 |
| 1 | 0.059 | 5.48182726 | 1 | 101.84 | 1 |
| | 0.062 | 2.57863379 | 1 | 0 | 1 |
| 1 | 0.425 | 24.9901624 | 6 | 122.360079 | 6 |
| | 0.11 | 0 | 1 | 0 | 1 |
| | 0.292 | 6.63916802 | 2 | 42.22 | 2 |
| 1 | 0.212 | 2.73769283 | 1 | 0 | 1 |
| | 0.053 | 2.05385113 | 1 | 0 | 1 |
| 8 | 2.001 | 92.92135 | 19 | 469.742236 | 18 |
| 2 | 0.131 | 13.6308334 | 6 | 112.711876 | 6 |
| 1 | 1.069 | 14.4635277 | 5 | 119.25 | 5 |
| 1 | 0.369 | 15.1884346 | 6 | 91.3501017 | 5 |
| 1 | 0.072 | 4.20985985 | 1 | 58.18 | 1 |
| 3 | 2.548 | 62.104183 | 10 | 311.924693 | 10 |
| 1 | 0.389 | 16.7184262 | 5 | 120.203987 | 5 |
| | 0.136 | 7.77141333 | 1 | 39.8996309 | 1 |
| 1 | 0.233 | 4.16327 | 1 | 75.43 | 1 |
| | 0.334 | 0 | 1 | 0 | 1 |
| | 0.15 | 2.3740294 | 2 | 29.89 | 2 |
| 3 | 1.154 | 20.8912714 | 5 | 132.96229 | 5 |
| 1 | 0.468 | 6.3106029 | 3 | 60.57 | 3 |
| | 0.118 | 10.2377412 | 3 | 39.54 | 3 |
| | 0.089 | 2.617342 | 1 | 27.95 | 1 |

| | | | | | |
|---|---|---|---|---|---|
| 1 | 0.172 | 2.6052587 | 2 | 44.87 | 2 |
| | 0.025 | 3.4101069 | 1 | 27.5 | 1 |
| 10 | 1.721 | 57.6558335 | 16 | 255.302539 | 15 |
| 1 | 0.445 | 23.1280358 | 8 | 135.03 | 8 |
| 1 | 0.359 | 5.98707151 | 2 | 28.32 | 2 |
| 1 | 0.119 | 5.49281001 | 2 | 59.15 | 2 |
| 1 | 0.179 | 7.04133129 | 2 | 32.89 | 2 |
| | 0.089 | 2.77164221 | 1 | 30.24 | 1 |
| | 0.407 | 16.7011819 | 4 | 60.6043963 | 4 |
| 1 | 0.077 | 2.46098542 | 1 | 44.44 | 1 |
| | 0.122 | 2.65080523 | 1 | 23.58 | 1 |
| 1 | 0.359 | 6.5909586 | 2 | 95.1177554 | 2 |
| 4 | 5.31 | 31.4902601 | 6 | 31.0086036 | 6 |
| 5 | 1.894 | 33.8821018 | 4 | 271.041216 | 4 |
| 1 | 0.359 | 4.71030951 | 1 | 59.76 | 2 |
| 1 | 0.222 | 5.64221668 | 2 | 0 | 2 |
| | 0.274 | 5.59249091 | 2 | 26.18 | 2 |
| | 0.194 | 1.85060883 | 1 | 0 | 1 |
| | 0.162 | 10.4228253 | 3 | 97.9647214 | 3 |
| 1 | 0.198 | 8.77171087 | 4 | 68.7113742 | 4 |
| | 0.222 | 5.25001335 | 2 | 58.2 | 2 |
| | 0.145 | 3.87992454 | 1 | 54.46 | 1 |
| 3 | 2.415 | 43.30968 | 8 | 339.389841 | 8 |
| 1 | 0.179 | 5.57638359 | 1 | 78.18 | 1 |
| | 0.056 | 2.79131579 | 1 | 30.49 | 1 |
| | 0.129 | 3.29107165 | 1 | 71.91 | 1 |
| 2 | 0.269 | 4.6408534 | 3 | 0 | 3 |
| | 0.129 | 3.10237145 | 1 | 40.89 | 1 |
| 1 | 0.585 | 6.05892611 | 2 | 55.7457475 | 2 |
| 2 | 0.066 | 6.65847635 | 2 | 75.72 | 1 |
| 1 | 0.688 | 21.1041734 | 5 | 136.809621 | 5 |
| 1 | 0.292 | 8.12592769 | 2 | 67.8 | 2 |
| | 0.136 | 5.24398184 | 1 | 57.67 | 1 |
| | 0.11 | 2.21179652 | 1 | 0 | 1 |
| 2 | 3.217 | 17.4547775 | 4 | 111.625051 | 4 |
| 2 | 0.369 | 14.0535975 | 3 | 0 | 3 |
| 2 | 0.22 | 14.5343404 | 5 | 91.6701887 | 5 |
| 4 | 2.36 | 39.0157735 | 8 | 263.117366 | 8 |
| 5 | 0.201 | 36.3172944 | 7 | 253.016738 | 7 |
| | 0.099 | 5.13693929 | 2 | 33.94 | 2 |
| 1 | 1.894 | 25.5633488 | 5 | 127.716576 | 5 |
| 3 | 1.154 | 24.4922726 | 6 | 189.52192 | 6 |
| 1 | 0.359 | 19.5262136 | 6 | 134.428731 | 6 |
| 1 | 0.334 | 3.42966437 | 1 | 0 | 1 |
| 3 | 0.431 | 24.2915354 | 7 | 145.92355 | 7 |
| 1 | 0.222 | 5.71302557 | 2 | 29.2059409 | 2 |
| 1 | 0.145 | 6.1438663 | 2 | 92.0585635 | 2 |
| 5 | 4.817 | 47.763989 | 12 | 317.973869 | 12 |
| 2 | 0.369 | 19.0628011 | 3 | 99.195426 | 3 |
| | 0.08 | 5.22345328 | 1 | 45.41 | 1 |
| | 0.075 | 3.56892085 | 1 | 37.66 | 1 |
| 1 | 0.16 | 4.62291098 | 2 | 51.35 | 2 |
| 5 | 0.968 | 15.6817832 | 5 | 99.0570276 | 5 |
| 1 | 0.334 | 2.67021775 | 1 | 0 | 1 |
| | 0.049 | 2.09302258 | 1 | 0 | 1 |
| 17 | 4.541 | 127.45044 | 18 | 757.469049 | 17 |
| 1 | 0.995 | 6.31447864 | 3 | 53.72 | 3 |
| | 0.212 | 2.62583256 | 1 | 32.53 | 1 |
| | 0.212 | 3.14176774 | 1 | 0 | 1 |
| 7 | 111.202 | 179.716702 | 20 | 759.915597 | 20 |
| 5 | 0.468 | 26.1332076 | 7 | 205.818613 | 7 |
| | 0.068 | 3.9523375 | 1 | 43.44 | 1 |

| | | | | | |
|---|---|---|---|---|---|
| 3 | 0.45 | 21.3602464 | 4 | 85.5922956 | 4 |
| 2 | 0.233 | 10.4829111 | 4 | 55.6516652 | 4 |
| 5 | 0.632 | 31.598175 | 8 | 256.725696 | 8 |
| 8 | 3.062 | 46.5825076 | 12 | 185.130264 | 12 |
| 1 | 0.019 | 5.00689363 | 1 | 69.67 | 1 |
| | 0.145 | 0 | 1 | 0 | 1 |
| 1 | 0.334 | 4.15654278 | 1 | 57.3 | 1 |
| 16 | 8.522 | 220.868153 | 37 | 1257.53916 | 37 |
| 1 | 0.129 | 7.34402108 | 2 | 34.18 | 2 |
| 1 | 0.212 | 3.35086513 | 1 | 22.74 | 1 |
| | 0.047 | 2.22218299 | 1 | 0 | 1 |
| 4 | 0.433 | 13.3358195 | 5 | 91.09 | 5 |
| | 0.122 | 3.27202725 | 1 | 31.21 | 1 |
| 1 | 0.049 | 0 | 1 | 0 | 1 |
| 1 | 0.068 | | | 35.9 | 1 |
| 3 | 0.435 | 25.6570497 | 7 | 269.390384 | 7 |
| | 0.259 | 2.07544422 | 1 | 32.25 | 1 |
| 1 | 0.141 | 2.17520881 | 2 | 0 | 2 |
| 3 | 1.955 | 31.4840786 | 8 | 129.052909 | 8 |
| 1 | 0.778 | 9.974823 | 3 | 73.1288509 | 3 |
| | 0.083 | 5.14523888 | 1 | 49.71 | 1 |
| | 0.905 | 53.6210053 | 10 | 357.672537 | 10 |
| | 0.155 | 6.35501051 | 1 | 39.29 | 1 |
| | 0.116 | 2.3621583 | 1 | 27.44 | 1 |
| 1 | 0.028 | 0 | 1 | 0 | 1 |
| | 0.129 | 2.73609614 | 1 | 34.93 | 1 |
| 3 | 0.389 | 7.43373251 | 3 | 84.33 | 4 |
| | 0.292 | 5.33846307 | 2 | 35.69 | 2 |
| | 0.438 | 10.4531686 | 3 | 68.0921645 | 3 |
| 1 | 0.334 | 9.29528213 | 3 | 65.03 | 2 |
| 1 | 0.122 | 3.42764735 | 1 | 40.96 | 1 |
| 1 | 1.683 | 29.0222683 | 9 | 165.59762 | 9 |
| | 0.041 | 2.64734554 | 1 | 0 | 1 |
| 3 | 2.981 | 8.59130073 | 3 | 45 | 3 |
| 3 | 6.499 | 25.8850629 | 5 | 163.433581 | 5 |
| | 0.16 | 8.58566236 | 2 | 42.62 | 2 |
| 1 | 0.073 | 7.85203195 | 2 | 104.65 | 2 |
| 1 | 0.322 | 11.4420829 | 4 | 50.6731771 | 4 |
| | 0.008 | 2.21120453 | 1 | 0 | 1 |
| 10 | 15.379 | 86.8000257 | 14 | 336.147673 | 15 |
| | 4.36 | 114.913288 | 30 | 656.0817 | 29 |
| | 0.212 | 2.91898561 | 1 | 50.81 | 1 |
| | 0.122 | 3.83832216 | 1 | 0 | 1 |
| 1 | 0.194 | 10.3413889 | 3 | 104.579147 | 3 |
| 5 | 0.833 | 44.2282519 | 10 | 229.928386 | 10 |
| 6 | 0.668 | 43.1590528 | 11 | 283.089866 | 11 |
| | 0.068 | 4.32535172 | 1 | 49.01 | 1 |
| 1 | 0.179 | 3.0935812 | 1 | 33.93 | 1 |
| 4 | 0.284 | 47.0088453 | 13 | 216.150802 | 14 |
| 1 | 0.129 | 4.43891191 | 1 | 59.43 | 1 |
| | 0.778 | 0 | 1 | 0 | 1 |
| 2 | 0.341 | 22.7218304 | 7 | 118.076307 | 7 |
| 2 | 0.307 | 15.4619412 | 5 | 61.6634504 | 5 |
| 1 | 0.931 | 3.61715674 | 2 | 0 | 2 |
| 4 | 0.607 | 15.0038302 | 7 | 124.558622 | 7 |
| | 0.016 | 2.78707457 | 1 | 25.14 | 1 |
| | 0.186 | 5.73484302 | 2 | 47.96 | 2 |
| 2 | 1.371 | 16.6712701 | 6 | 109.633333 | 5 |
| | 0 | 0 | 1 | | |
| | 0.016 | 3.48830724 | 1 | 45.97 | 1 |
| | 0.222 | 12.0662122 | 2 | 90.6398343 | 2 |
| 3 | 9 | 23.5312965 | 4 | 155.015453 | 4 |

| | | | | | |
|---|---|---|---|---|---|
| 4 | 0.756 | 39.5647807 | 11 | 284.499613 | 11 |
| 8 | 3.806 | 59.3042092 | 13 | 411.38441 | 13 |
| 33 | 3.108 | 241.860111 | 46 | 1266.05559 | 46 |
| 4 | 0.359 | 18.8539302 | 6 | 98.12 | 6 |
| | 0.116 | 0 | 1 | 0 | 1 |
| 3 | 0.484 | 31.0051498 | 6 | 268.172566 | 6 |
| 4 | 1.783 | 11.9519477 | 4 | 72.41 | 4 |
| | 0.292 | 3.16925216 | 1 | 34.78 | 1 |
| | 0.389 | 5.24903154 | 1 | 82.14 | 1 |
| | 0.024 | 3.32145762 | 1 | 0 | 1 |
| | 0.033 | 2.64250636 | 1 | 31.45 | 1 |
| 4 | 0.931 | 18.3498647 | 5 | 101.893783 | 5 |
| 1 | 0.501 | 8.2989397 | 3 | 53.16 | 3 |
| | 0.093 | 2.30029058 | 1 | 31.66 | 1 |
| | 0.077 | 3.23607302 | 1 | 24.91 | 1 |
| 1 | 0.848 | 17.4857242 | 4 | 120.31 | 4 |
| 1 | 0.11 | 4.19216347 | 1 | 57.71 | 1 |
| 2 | 0.343 | 9.16006804 | 5 | 88.34 | 5 |
| 1 | 0.413 | 10.9660358 | 3 | 114.140644 | 3 |
| 5 | 1.637 | 22.3984194 | 7 | 84.5156241 | 6 |
| | 0.11 | 3.54422998 | 1 | 0 | 1 |
| | 0.166 | 1.7239331 | 1 | 0 | 1 |
| | 0.233 | 3.03165984 | 1 | 38.12 | 1 |
| 1 | 0.151 | 11.4182832 | 3 | 64.89 | 3 |
| 4 | 214.443 | 331.99707 | 27 | 1477.80735 | 27 |
| 10 | 9 | 137.732184 | 7 | 663.941439 | 6 |
| 1 | 0.035 | 3.01637363 | 1 | 34.07 | 1 |
| 1 | 0.172 | 6.25181222 | 2 | 27.66 | 2 |
| | 0.055 | 3.1816082 | 1 | 40.79 | 1 |
| | 0.136 | 3.59006786 | 1 | 0 | 1 |
| 1 | 0.585 | 3.00669527 | 1 | 40.63 | 1 |
| 1 | 0.105 | 5.24854946 | 1 | 64.11 | 1 |
| | 0.08 | 8.0681169 | 2 | 76.8085698 | 2 |
| | 0.212 | 3.38775873 | 1 | 44.86 | 1 |
| 1 | 0.061 | 0 | 1 | 0 | 1 |
| 1 | 4.623 | 24.2891643 | 6 | 160.433769 | 6 |
| 2 | 0.298 | 17.6312733 | 5 | 118.886667 | 5 |
| 6 | 1.61 | 47.3871284 | 9 | 172.594571 | 9 |
| | 0.126 | 13.4733295 | 3 | 46.79 | 3 |
| 1 | 0.105 | 7.07150745 | 2 | 70.0243589 | 2 |
| | 4.36 | 110.298822 | 30 | 665.583143 | 29 |
| 1 | 0.072 | 0 | 1 | 0 | 1 |
| 1 | 0.413 | 5.53597426 | 3 | 0 | 3 |
| 39 | 2.562 | 311.882816 | 67 | 1458.05674 | 67 |
| | 0.129 | 4.91454744 | 1 | 60.94 | 1 |
| 2 | 0.52 | 13.2453802 | 4 | 79.3494001 | 3 |
| | 0.083 | 3.16822672 | 1 | 36.69 | 1 |
| | 1.043 | 22.2919564 | 7 | 109.438326 | 7 |
| | 0.122 | 0 | 1 | | |
| 1 | 0.468 | 4.42844677 | 1 | 47.32 | 1 |
| 11 | 3.084 | 112.182191 | 17 | 622.726699 | 17 |
| | 0.122 | 4.4616127 | 1 | 66.54 | 1 |
| 1 | 0.624 | 9.99548578 | 4 | 0 | 2 |
| | 0.032 | 3.52402496 | 1 | 43.01 | 1 |
| | 0.108 | 4.50960803 | 1 | 48.08 | 2 |
| 1 | 0.334 | 3.60364079 | 1 | 40.33 | 1 |
| | 0.334 | 2.51871729 | 1 | 20.92 | 1 |
| | 0.096 | 2.25824881 | 1 | 0 | 1 |
| | 0.101 | 3.10868073 | 1 | 42.11 | 1 |
| 1 | 0.122 | 4.09151316 | 1 | 64.66 | 1 |
| 1 | 0.035 | 3.69962978 | 1 | 54.97 | 1 |
| | 0.259 | 2.41698313 | 1 | 0 | 1 |

| | | | | | |
|---|---|---|---|---|---|
| | 0.334 | 2.97632575 | 1 | 0 | 1 |
| 2 | 0.638 | 13.0494568 | 3 | 57.11 | 3 |
| 1 | 0.389 | 9.81984496 | 2 | 46.3321347 | 2 |
| 3 | 0.407 | 15.0842495 | 4 | 148.04 | 4 |
| | 0.062 | 3.60140944 | 1 | 36.48 | 1 |
| 5 | 0.182 | 19.6408682 | 5 | 130.400878 | 5 |
| | 0.058 | 3.59931421 | 1 | 26.38 | 1 |
| | 0.035 | 1.7629379 | 1 | 29.04 | 1 |
| 1 | 0.194 | 11.9019761 | 3 | 101.41 | 3 |
| | 0.086 | 2.56036448 | 1 | 51.13 | 1 |
| 1 | 0.068 | 0 | 1 | 0 | 1 |
| | 0.058 | 4.77434587 | 1 | 71.02 | 1 |
| 4 | 0.19 | 33.9019177 | 7 | 208.617056 | 7 |
| | 0.129 | | | 0 | 1 |
| | 0.053 | 0 | 1 | 0 | 1 |
| 1 | 0.026 | 4.84405422 | 1 | 70.83 | 1 |
| 1 | 0.139 | 18.9944637 | 4 | 45.150761 | 4 |
| 1 | 0.139 | 19.3300912 | 4 | 82.9796923 | 4 |
| | 0.037 | 5.84582782 | 1 | 55.5 | 1 |
| | 0.039 | 3.04776144 | 1 | 48.92 | 1 |
| | 0.266 | 7.06067872 | 4 | 53.7122135 | 4 |
| | 0.066 | 3.1475215 | 1 | 0 | 1 |
| | 0.122 | 2.46326256 | 1 | 46.37 | 1 |
| 1 | 0.116 | 2.35131311 | 1 | 33.31 | 1 |
| | 0.066 | 3.64079952 | 1 | 61.3 | 1 |
| 1 | 0.129 | 51.3787599 | 12 | 341.54314 | 12 |
| 2 | 4.337 | 30.3673756 | 6 | 217.564042 | 6 |
| | 0.096 | 3.89039826 | 1 | 58.24 | 1 |
| | 0.11 | 0 | 1 | 0 | 1 |
| | 0.585 | 2.73818707 | 1 | 46.35 | 1 |
| 2 | 0.269 | 12.5593805 | 3 | 107.60643 | 3 |
| 1 | 0.413 | 12.7459736 | 2 | 145.447753 | 2 |
| 1 | 0.083 | 4.43010902 | 1 | 47.8 | 1 |
| 5 | 0.859 | 7.03869283 | 6 | 46.2603334 | 7 |
| | 0.047 | 5.81175852 | 1 | 0 | 1 |
| 1 | 0.413 | 6.56689692 | 3 | 54.89 | 3 |
| 3 | 1.31 | 35.1529207 | 8 | 146.947273 | 8 |
| 1 | 0.198 | 14.1479902 | 4 | 127.276667 | 4 |
| | 0.07 | 3.06650376 | 1 | 51.42 | 1 |
| 1 | 1.297 | 41.4048295 | 11 | 250.601538 | 11 |
| 1 | 0.075 | 8.3038609 | 2 | 111.93 | 2 |
| | 0.179 | 4.57104683 | 1 | 62.03 | 1 |
| | 0.037 | 2.44085431 | 1 | 0 | 1 |
| 2 | 1.424 | 11.3949587 | 5 | 45.19 | 5 |
| 1 | 0.067 | 7.14817071 | 2 | 39.55 | 2 |
| 1 | 0.413 | 10.5749772 | 3 | 101.07 | 2 |
| | 0.044 | 3.01259279 | 1 | | |
| 2 | 0.245 | 4.00394678 | 2 | 34.82 | 2 |
| 1 | 0.425 | 9.9839642 | 2 | 54.4850254 | 2 |
| | 0.202 | 5.41027117 | 2 | 0 | 2 |
| | 0.122 | 2.89576817 | 1 | 0 | 1 |
| 2 | 1.069 | 39.0725913 | 5 | 255.822987 | 5 |
| 1 | 0.259 | 6.87001848 | 1 | 61.8 | 1 |
| | 0.072 | 3.00815678 | 1 | 0 | 1 |
| | 1.096 | 41.8617544 | 8 | 168.685943 | 8 |
| 2 | 0.148 | 20.7718282 | 3 | 255.236963 | 3 |
| | 0.172 | 8.80663919 | 2 | 31.572431 | 2 |
| 3 | 0.778 | 40.1378441 | 3 | 205.224663 | 3 |
| | 0.066 | 2.22451568 | 1 | 0 | 1 |
| | 0.124 | 5.78024507 | 3 | 41.3490033 | 3 |
| 25 | 29.599 | 184.589075 | 30 | 933.869712 | 30 |
| | 0.081 | 6.09651542 | 2 | 28.8 | 2 |

| | | | | | |
|---|---|---|---|---|---|
| 1 | 1.154 | 9.1558733 | 1 | 100.7 | 1 |
| 5 | 0.778 | 40.3086298 | 14 | 190.281256 | 13 |
| 1 | 0.585 | 5.1516633 | 3 | 0 | 3 |
| | 0.155 | 2.61832142 | 1 | 0 | 1 |
| | 0.136 | 5.55562592 | 1 | 79.91 | 1 |
| 1 | 0.155 | 4.17028713 | 1 | 49.38 | 1 |
| 5 | 0.532 | 37.4944663 | 10 | 263.270025 | 10 |
| 1 | 0.068 | 3.08606768 | 1 | 49.57 | 1 |
| 2 | 0.168 | 16.2255976 | 5 | 96.5716467 | 5 |
| 4 | 1.818 | 32.1620581 | 7 | 32.18 | 7 |
| | 0.056 | | | 0 | 1 |
| 2 | 0.139 | 21.3044567 | 6 | 136.614795 | 6 |
| 1 | 0.089 | 6.35172653 | 2 | 37.6237374 | 2 |
| 1 | 0.15 | 8.61871409 | 2 | 87.016192 | 2 |
| | 0.334 | 4.17816877 | 1 | 58.52 | 1 |
| | 0.334 | 4.71441531 | 1 | 20.53 | 1 |
| 1 | 0.73 | 17.1244293 | 5 | 82.6641506 | 5 |
| | 0.062 | 3.50643921 | 1 | 0 | 1 |
| | 0.044 | 2.11433935 | 2 | 0 | 2 |
| | 0.093 | 2.62750602 | 1 | 35.17 | 1 |
| | 0.068 | 4.29015923 | 1 | 37.98 | 1 |
| | 0.122 | 6.50384521 | 2 | 32.1690874 | 2 |
| | 0.059 | 4.84123087 | 1 | 88.59 | 1 |
| | 0.116 | 4.11641312 | 1 | 54.12 | 1 |
| 13 | 2.652 | 97.4855223 | 22 | 342.932177 | 22 |
| | 0.186 | 9.02609968 | 2 | 84.337022 | 2 |
| | 0.129 | 4.14313459 | 1 | 52.61 | 1 |
| | 0.039 | 4.91426897 | 1 | 61.18 | 1 |
| | 0.233 | 3.09380746 | 2 | 55.76 | 1 |
| | 0.018 | 2.45107245 | 1 | 34.38 | 1 |
| 3 | 0.995 | 7.27087092 | 3 | 40.2145735 | 2 |
| 3 | 0.141 | 11.7016759 | 4 | 49.35 | 4 |
| 5 | 2.793 | 34.7897592 | 10 | 109.457503 | 10 |
| 2 | 1.683 | 44.3762963 | 4 | 248.81841 | 4 |
| | 0.194 | 2.98549557 | 1 | 50.29 | 1 |
| | 0.034 | 3.24770975 | 1 | 0 | 1 |
| 1 | 0.055 | 2.62784266 | 1 | 0 | 1 |
| | 0.468 | 0 | 1 | | |
| 1 | 0.166 | 5.84966469 | 1 | 52.3247104 | 1 |
| | 0.116 | 2.4837873 | 1 | 44.91 | 1 |
| 1 | 0.077 | 9.13919806 | 3 | 62.44 | 3 |
| | 0.122 | 3.82007718 | 1 | 38.66 | 1 |
| 2 | 0.194 | 7.61557293 | 2 | 78.45 | 2 |
| | 0.136 | 2.21036386 | 1 | 0 | 1 |
| 2 | 0.186 | 6.31169677 | 2 | 45.91 | 2 |
| 4 | 0.374 | 28.7832153 | 8 | 200.650357 | 8 |
| 1 | 0.062 | 2.58021116 | 1 | 31.55 | 1 |
| 1 | 0.905 | 26.1526415 | 7 | 109.824167 | 7 |
| | 0.425 | 5.59275079 | 2 | 75.6303897 | 2 |
| | 0.233 | 3.60427594 | 1 | 49.9 | 1 |
| 2 | 2.162 | 22.2196481 | 5 | 30.2119164 | 5 |
| 1 | 1.154 | 26.4029219 | 7 | 142.966488 | 7 |
| 1 | 0.468 | 3.71821475 | 1 | 79.99 | 1 |
| 5 | 1.154 | 32.4153266 | 7 | 161.484905 | 5 |
| | 0.668 | 2.06957173 | 2 | 0 | 1 |
| 2 | 0.205 | 7.98886919 | 3 | 62.8665195 | 3 |
| 4 | 1.581 | 20.0059948 | 6 | 113.86 | 4 |

| Accession | Gene symbol | Protein description | Refseq protein accession |
|---|---|---|---|
| 4501857 | ACADL | long-chain specific acyl-CoA dehydrogenase, mi | NP_001599.1 |
| 4501867 | ACO2 | aconitate hydratase, mitochondrial precursor [H | NP_001089.1 |
| 4501881 | ACTA1 | actin, alpha skeletal muscle [Homo sapiens] | NP_001091.1 |
| 4501913 | ADAM23 | disintegrin and metalloproteinase domain-contair | NP_003803.1 |
| 4502049 | AKR1B1 | aldose reductase [Homo sapiens] | NP_001619.1 |
| 4502107 | ANXA5 | annexin A5 [Homo sapiens] | NP_001145.1 |
| 4502201 | ARF1 | ADP-ribosylation factor 1 [Homo sapiens] | NP_001649.1 |
| 4502209 | ARF5 | ADP-ribosylation factor 5 [Homo sapiens] | NP_001653.1 |
| 4502271 | ATP1A2 | sodium/potassium-transporting ATPase subunit | NP_000693.1 |
| 4502277 | ATP1B1 | sodium/potassium-transporting ATPase subunit | NP_001668.1 |
| 4502303 | ATP5O | ATP synthase subunit O, mitochondrial precurso | NP_001688.1 |
| 4502315 | ATP6V1C1 | V-type proton ATPase subunit C 1 [Homo sapier | NP_001686.1 |
| 4502317 | ATP6V1E1 | V-type proton ATPase subunit E 1 isoform a [Ho | NP_001687.1 |
| 4502643 | CCT6A | T-complex protein 1 subunit zeta isoform a [Hon | NP_001753.1 |
| 4502673 | CD47 | leukocyte surface antigen CD47 isoform 1 precu | NP_001768.1 |
| 4502993 | COX7C | cytochrome c oxidase subunit 7C, mitochondrial | NP_001858.1 |
| 4503009 | CPE | carboxypeptidase E preproprotein [Homo sapien | NP_001864.1 |
| 4503049 | CRIP2 | cysteine-rich protein 2 isoform 1 [Homo sapiens | NP_001303.1 |
| 4503065 | CRYM | ketimine reductase mu-crystallin [Homo sapiens | NP_001879.1 |
| 4503095 | CSNK2A1 | casein kinase II subunit alpha isoform a [Homo | NP_001886.1 |
| 4503377 | DPYSL2 | dihydropyrimidinase-related protein 2 isoform 2 [ | NP_001377.1 |
| 4503379 | DPYSL3 | dihydropyrimidinase-related protein 3 isoform 2 [ | NP_001378.1 |
| 4503471 | EEF1A1 | elongation factor 1-alpha 1 [Homo sapiens] | NP_001393.1 |
| 4503475 | EEF1A2 | elongation factor 1-alpha 2 [Homo sapiens] | NP_001949.1 |
| 4503477 | EEF1B2 | elongation factor 1-beta [Homo sapiens] | NP_001950.1 |
| 4503481 | EEF1G | elongation factor 1-gamma [Homo sapiens] | NP_001395.1 |
| 4503483 | EEF2 | elongation factor 2 [Homo sapiens] | NP_001952.1 |
| 4503529 | EIF4A1 | eukaryotic initiation factor 4A-I isoform 1 [Homo | NP_001407.1 |
| 4503571 | ENO1 | alpha-enolase isoform 1 [Homo sapiens] | NP_001419.1 |
| 4503607 | ETFA | electron transfer flavoprotein subunit alpha, mito | NP_000117.1 |
| 4503875 | GAD2 | glutamate decarboxylase 2 [Homo sapiens] | NP_000809.1 |
| 4503971 | GDI1 | rab GDP dissociation inhibitor alpha [Homo sapie | NP_001484.1 |
| 4504041 | GNAI2 | guanine nucleotide-binding protein G(i) subunit a | NP_002061.1 |
| 4504067 | GOT1 | aspartate aminotransferase, cytoplasmic [Homo | NP_002070.1 |
| 4504111 | GRB2 | growth factor receptor-bound protein 2 isoform 1 | NP_002077.1 |
| 4504251 | HIST2H2AA3 | histone H2A type 2-A [Homo sapiens] | NP_003507.1 |
| 4504255 | H2AFZ | histone H2A.Z [Homo sapiens] | NP_002097.1 |
| 4504317 | HIST1H4L | histone H4 [Homo sapiens] | NP_003537.1 |
| 4504347 | HBA1 | hemoglobin subunit alpha [Homo sapiens] | NP_000549.1 |
| 4504511 | DNAJA1 | dnaJ homolog subfamily A member 1 isoform 1 [ | NP_001530.1 |
| 4504523 | HSPE1 | 10 kDa heat shock protein, mitochondrial [Homo | NP_002148.1 |
| 4505185 | MIF | macrophage migration inhibitory factor [Homo sa | NP_002406.1 |
| 4505357 | NDUFA4 | cytochrome c oxidase subunit NDUFA4 [Homo s | NP_002480.1 |
| 4505369 | NDUFS4 | NADH dehydrogenase [ubiquinone] iron-sulfur pr | NP_002486.1 |
| 4505451 | NRAS | GTPase NRas [Homo sapiens] | NP_002515.1 |
| 4505531 | OSBP | oxysterol-binding protein 1 [Homo sapiens] | NP_002547.1 |
| 4505585 | PAFAH1B2 | platelet-activating factor acetylhydrolase IB sub | NP_002563.1 |
| 4505621 | PEBP1 | phosphatidylethanolamine-binding protein 1 [Hon | NP_002558.1 |
| 4505753 | PGAM1 | phosphoglycerate mutase 1 [Homo sapiens] | NP_002620.1 |
| 4505763 | PGK1 | phosphoglycerate kinase 1 [Homo sapiens] | NP_000282.1 |
| 4506013 | PPP1R7 | protein phosphatase 1 regulatory subunit 7 isofo | NP_002703.1 |
| 4506017 | PPP2CA | serine/threonine-protein phosphatase 2A catalyt | NP_002706.1 |
| 4506025 | PPP3R1 | calcineurin subunit B type 1 [Homo sapiens] | NP_000936.1 |
| 4506181 | PSMA2 | proteasome subunit alpha type-2 [Homo sapiens | NP_002778.1 |
| 4506183 | PSMA3 | proteasome subunit alpha type-3 isoform 1 [Hon | NP_002779.1 |
| 4506209 | PSMC2 | 26S protease regulatory subunit 7 isoform 1 [Ho | NP_002794.1 |
| 4506221 | PSMD12 | 26S proteasome non-ATPase regulatory subunit | NP_002807.1 |

| | | | |
|---|---|---|---|
| 4506365 | RAB2A | ras-related protein Rab-2A isoform a [Homo sapi | NP_002856.1 |
| 4506367 | RAB3A | ras-related protein Rab-3A [Homo sapiens] | NP_002857.1 |
| 4506371 | RAB5B | ras-related protein Rab-5B isoform 1 [Homo sapi | NP_002859.1 |
| 4506597 | RPL12 | 60S ribosomal protein L12 [Homo sapiens] | NP_000967.1 |
| 4506605 | RPL23 | 60S ribosomal protein L23 [Homo sapiens] | NP_000969.1 |
| 4506645 | RPL38 | 60S ribosomal protein L38 [Homo sapiens] | NP_000990.1 |
| 4506649 | RPL3 | 60S ribosomal protein L3 isoform a [Homo sapie | NP_000958.1 |
| 4506661 | RPL7A | 60S ribosomal protein L7a [Homo sapiens] | NP_000963.1 |
| 4506663 | RPL8 | 60S ribosomal protein L8 [Homo sapiens] | NP_000964.1 |
| 4506671 | RPLP2 | 60S acidic ribosomal protein P2 [Homo sapiens] | NP_000995.1 |
| 4506685 | RPS13 | 40S ribosomal protein S13 [Homo sapiens] | NP_001008.1 |
| 4506691 | RPS16 | 40S ribosomal protein S16 [Homo sapiens] | NP_001011.1 |
| 4506693 | RPS17 | 40S ribosomal protein S17 [Homo sapiens] | NP_001012.1 |
| 4506695 | RPS19 | 40S ribosomal protein S19 [Homo sapiens] | NP_001013.1 |
| 4506707 | RPS25 | 40S ribosomal protein S25 [Homo sapiens] | NP_001019.1 |
| 4506723 | RPS3A | 40S ribosomal protein S3a isoform 1 [Homo sapi | NP_000997.1 |
| 4506725 | RPS4X | 40S ribosomal protein S4, X isoform X isoform [I | NP_000998.1 |
| 4506741 | RPS7 | 40S ribosomal protein S7 [Homo sapiens] | NP_001002.1 |
| 4506743 | RPS8 | 40S ribosomal protein S8 [Homo sapiens] | NP_001003.1 |
| 4506929 | SH3GL1 | endophilin-A2 isoform 1 [Homo sapiens] | NP_003016.1 |
| 4507115 | FSCN1 | fascin [Homo sapiens] | NP_003079.1 |
| 4507137 | SNTA1 | alpha-1-syntrophin [Homo sapiens] | NP_003089.1 |
| 4507677 | HSP90B1 | endoplasmin precursor [Homo sapiens] | NP_003290.1 |
| 4507729 | TUBB2A | tubulin beta-2A chain isoform 1 [Homo sapiens] | NP_001060.1 |
| 4507793 | UBE2N | ubiquitin-conjugating enzyme E2 N [Homo sapie | NP_003339.1 |
| 4507879 | VDAC1 | voltage-dependent anion-selective channel prote | NP_003365.1 |
| 4507947 | YARS | tyrosine--tRNA ligase, cytoplasmic [Homo sapie | NP_003671.1 |
| 4507949 | YWHAB | 14-3-3 protein beta/alpha [Homo sapiens] | NP_003395.1 |
| 4507951 | YWHAH | 14-3-3 protein eta [Homo sapiens] | NP_003396.1 |
| 4557237 | ACAT1 | acetyl-CoA acetyltransferase, mitochondrial pre | NP_000010.1 |
| 4557367 | BLMH | bleomycin hydrolase [Homo sapiens] | NP_000377.1 |
| 4557395 | CA2 | carbonic anhydrase 2 isoform 1 [Homo sapiens] | NP_000058.1 |
| 4557707 | L1CAM | neural cell adhesion molecule L1 isoform 1 prec | NP_000416.1 |
| 4557735 | MAOA | amine oxidase [flavin-containing] A isoform 1 [H | NP_000231.1 |
| 4557817 | OXCT1 | succinyl-CoA:3-ketoacid coenzyme A transferas | NP_000427.1 |
| 4757774 | ARL3 | ADP-ribosylation factor-like protein 3 [Homo sap | NP_004302.1 |
| 4757900 | CALR | calreticulin precursor [Homo sapiens] | NP_004334.1 |
| 4758086 | CSRP1 | cysteine and glycine-rich protein 1 isoform 1 [Hc | NP_004069.1 |
| 4758152 | TIMM8A | mitochondrial import inner membrane translocase | NP_004076.1 |
| 4758256 | EIF2S1 | eukaryotic translation initiation factor 2 subunit | NP_004085.1 |
| 4758328 | FABP3 | fatty acid-binding protein, heart [Homo sapiens] | NP_004093.1 |
| 4758442 | GMFB | glia maturation factor beta [Homo sapiens] | NP_004115.1 |
| 4758582 | IDH3G | isocitrate dehydrogenase [NAD] subunit gamma, | NP_004126.1 |
| 4758638 | PRDX6 | peroxiredoxin-6 [Homo sapiens] | NP_004896.1 |
| 4758650 | KIF5C | kinesin heavy chain isoform 5C [Homo sapiens] | NP_004513.1 |
| 4758786 | NDUFS2 | NADH dehydrogenase [ubiquinone] iron-sulfur pr | NP_004541.1 |
| 4758788 | NDUFS3 | NADH dehydrogenase [ubiquinone] iron-sulfur pr | NP_004542.1 |
| 4758792 | NDUFS6 | NADH dehydrogenase [ubiquinone] iron-sulfur pr | NP_004544.1 |
| 4758950 | PPIB | peptidyl-prolyl cis-trans isomerase B precursor [ | NP_000933.1 |
| 4758988 | RAB1A | ras-related protein Rab-1A isoform 1 [Homo sapi | NP_004152.1 |
| 4759160 | SNRPD3 | small nuclear ribonucleoprotein Sm D3 [Homo sa | NP_004166.1 |
| 4759182 | STX1A | syntaxin-1A isoform 1 [Homo sapiens] | NP_004594.1 |
| 4759198 | HOMER1 | homer protein homolog 1 isoform 1 [Homo sapie | NP_004263.1 |
| 4759302 | VAPB | vesicle-associated membrane protein-associate | NP_004729.1 |
| 4759306 | LIN7A | protein lin-7 homolog A [Homo sapiens] | NP_004655.1 |
| 4826655 | CALB1 | calbindin [Homo sapiens] | NP_004920.1 |
| 4826675 | CDK5 | cyclin-dependent-like kinase 5 isoform 1 [Homo | NP_004926.1 |
| 4826734 | FUS | RNA-binding protein FUS isoform 1 [Homo sapie | NP_004951.1 |
| 4826816 | LGI1 | leucine-rich glioma-inactivated protein 1 isoform | NP_005088.1 |
| 4826898 | PFN1 | profilin-1 [Homo sapiens] | NP_005013.1 |
| 4826932 | PPID | peptidyl-prolyl cis-trans isomerase D [Homo sapi | NP_005029.1 |
| 4885281 | GLUD1 | glutamate dehydrogenase 1, mitochondrial prec | NP_005262.1 |

| | | | |
|---|---|---|---|
| 4885371 | H1F0 | histone H1.0 [Homo sapiens] | NP_005309.1 |
| 4885377 | HIST1H1D | histone H1.3 [Homo sapiens] | NP_005311.1 |
| 4885563 | PRKCE | protein kinase C epsilon type [Homo sapiens] | NP_005391.1 |
| 5031569 | ACTR1A | alpha-centractin [Homo sapiens] | NP_005727.1 |
| 5031571 | ACTR2 | actin-related protein 2 isoform b [Homo sapiens] | NP_005713.1 |
| 5031573 | ACTR3 | actin-related protein 3 isoform 1 [Homo sapiens] | NP_005712.1 |
| 5031599 | ARPC2 | actin-related protein 2/3 complex subunit 2 [Hom | NP_005722.1 |
| 5031635 | CFL1 | cofilin-1 [Homo sapiens] | NP_005498.1 |
| 5031741 | DNAJA2 | dnaJ homolog subfamily A member 2 [Homo sap | NP_005871.1 |
| 5031777 | IDH3A | isocitrate dehydrogenase [NAD] subunit alpha, r | NP_005521.1 |
| 5031985 | NUTF2 | nuclear transport factor 2 [Homo sapiens] | NP_005787.1 |
| 5032007 | PURA | transcriptional activator protein Pur-alpha [Homo | NP_005850.1 |
| 5032009 | PYGM | glycogen phosphorylase, muscle form isoform 1 | NP_005600.1 |
| 5032139 | SYT1 | synaptotagmin-1 isoform 1 [Homo sapiens] | NP_005630.1 |
| 5174391 | AKR1A1 | alcohol dehydrogenase [NADP(+)] [Homo sapien | NP_006057.1 |
| 5174447 | GNB2L1 | guanine nucleotide-binding protein subunit beta-2 | NP_006089.1 |
| 5174613 | NAP1L4 | nucleosome assembly protein 1-like 4 [Homo sa | NP_005960.1 |
| 5174735 | TUBB4B | tubulin beta-4B chain [Homo sapiens] | NP_006079.1 |
| 5453555 | RAN | GTP-binding nuclear protein Ran isoform 1 [Hom | NP_006316.1 |
| 5453559 | ATP5H | ATP synthase subunit d, mitochondrial isoform a | NP_006347.1 |
| 5453593 | CAP2 | adenylyl cyclase-associated protein 2 [Homo sa | NP_006357.1 |
| 5453599 | CAPZA2 | F-actin-capping protein subunit alpha-2 [Homo sa | NP_006127.1 |
| 5453603 | CCT2 | T-complex protein 1 subunit beta isoform 1 [Hom | NP_006422.1 |
| 5453607 | CCT7 | T-complex protein 1 subunit eta isoform a [Homo | NP_006420.1 |
| 5453629 | DCTN2 | dynactin subunit 2 isoform 1 [Homo sapiens] | NP_006391.1 |
| 5453710 | LASP1 | LIM and SH3 domain protein 1 isoform a [Homo s | NP_006139.1 |
| 5453908 | PITPNA | phosphatidylinositol transfer protein alpha isofor | NP_006215.1 |
| 5453958 | PPP5C | serine/threonine-protein phosphatase 5 isoform | NP_006238.1 |
| 5454034 | S100B | protein S100-B [Homo sapiens] | NP_006263.1 |
| 5579478 | MAP2K1 | dual specificity mitogen-activated protein kinase | NP_002746.1 |
| 5729781 | CPLX1 | complexin-1 [Homo sapiens] | NP_006642.1 |
| 5729875 | PGRMC1 | membrane-associated progesterone receptor co | NP_006658.1 |
| 5729991 | PSMC4 | 26S protease regulatory subunit 6B isoform 1 [H | NP_006494.1 |
| 5802966 | DSTN | destrin isoform a [Homo sapiens] | NP_006861.1 |
| 5803011 | ENO2 | gamma-enolase [Homo sapiens] | NP_001966.1 |
| 5803013 | ERP29 | endoplasmic reticulum resident protein 29 isofor | NP_006808.1 |
| 5803127 | PKIA | cAMP-dependent protein kinase inhibitor alpha [ | NP_006814.1 |
| 5803187 | TALDO1 | transaldolase [Homo sapiens] | NP_006746.1 |
| 5803225 | YWHAE | 14-3-3 protein epsilon [Homo sapiens] | NP_006752.1 |
| 5803227 | YWHAQ | 14-3-3 protein theta [Homo sapiens] | NP_006817.1 |
| 5902018 | TPPP | tubulin polymerization-promoting protein [Homo s | NP_008961.1 |
| 5902076 | SRSF1 | serine/arginine-rich splicing factor 1 isoform 1 [H | NP_008855.1 |
| 6005709 | PALM2-AKAP2 | PALM2-AKAP2 protein isoform 1 [Homo sapiens] | NP_009134.1 |
| 6005717 | ATP5I | ATP synthase subunit e, mitochondrial [Homo sa | NP_009031.1 |
| 6005942 | VCP | transitional endoplasmic reticulum ATPase [Hom | NP_009057.1 |
| 6005993 | CLTA | clathrin light chain A isoform b [Homo sapiens] | NP_009027.1 |
| 6005995 | CLTB | clathrin light chain B isoform b [Homo sapiens] | NP_009028.1 |
| 6031192 | SLC25A3 | phosphate carrier protein, mitochondrial isoform | NP_005879.1 |
| 6598323 | GDI2 | rab GDP dissociation inhibitor beta isoform 1 [Ho | NP_001485.2 |
| 6715568 | PPP3CA | serine/threonine-protein phosphatase 2B catalyt | NP_000935.1 |
| 6912238 | PRDX5 | peroxiredoxin-5, mitochondrial isoform a precurs | NP_036226.1 |
| 6912280 | AHSA1 | activator of 90 kDa heat shock protein ATPase | NP_036243.1 |
| 6912328 | DDAH1 | N(G),N(G)-dimethylarginine dimethylaminohydrola | NP_036269.1 |
| 6912482 | LETM1 | LETM1 and EF-hand domain-containing protein 1 | NP_036450.1 |
| 6912494 | MAPRE1 | microtubule-associated protein RP/EB family me | NP_036457.1 |
| 6912634 | RPL13A | 60S ribosomal protein L13a isoform 1 [Homo sap | NP_036555.1 |
| 6912646 | NPTN | neuroplastin isoform b precursor [Homo sapiens] | NP_036560.1 |
| 7019519 | PCSK1N | proSAAS preproprotein [Homo sapiens] | NP_037403.1 |
| 7549809 | PLS3 | plastin-3 isoform 1 [Homo sapiens] | NP_005023.2 |
| 7657015 | RTCB | tRNA-splicing ligase RtcB homolog [Homo sapier | NP_055121.1 |
| 7657033 | NT5C | 5'(3')-deoxyribonucleotidase, cytosolic type isof | NP_055410.1 |
| 7657056 | EHD3 | EH domain-containing protein 3 [Homo sapiens] | NP_055415.1 |

| | | | | |
|---|---|---|---|---|
| 7657381 | PRPF19 | | pre-mRNA-processing factor 19 [Homo sapiens] | NP_055317.1 |
| 7661678 | RAP1B | | ras-related protein Rap-1b isoform 1 precursor [I | NP_056461.1 |
| 7661862 | PPM1F | | protein phosphatase 1F [Homo sapiens] | NP_055449.1 |
| 7705419 | HPCAL4 | | hippocalcin-like protein 4 isoform 1 [Homo sapie | NP_057341.1 |
| 7705987 | GLTP | | glycolipid transfer protein [Homo sapiens] | NP_057517.1 |
| 7706497 | CMPK1 | | UMP-CMP kinase isoform a [Homo sapiens] | NP_057392.1 |
| 7706567 | GNG13 | | guanine nucleotide-binding protein G(I)/G(S)/G(C | NP_057625.1 |
| 7706757 | ATP6V1D | | V-type proton ATPase subunit D [Homo sapiens] | NP_057078.1 |
| 8922498 | PNPO | | pyridoxine-5'-phosphate oxidase [Homo sapiens] | NP_060599.1 |
| 8923911 | LANCL2 | | lanC-like protein 2 [Homo sapiens] | NP_061167.1 |
| 8923930 | CISD1 | | CDGSH iron-sulfur domain-containing protein 1 [I | NP_060934.1 |
| 9257257 | WDR1 | | WD repeat-containing protein 1 isoform 1 [Homo | NP_059830.1 |
| 9845509 | RAC1 | | ras-related C3 botulinum toxin substrate 1 isofor | NP_061485.1 |
| 9910382 | TOMM22 | | mitochondrial import receptor subunit TOM22 hor | NP_064628.1 |
| 9945322 | SLC17A7 | | vesicular glutamate transporter 1 [Homo sapiens | NP_064705.1 |
| 9945439 | | 5-Sep | septin-5 isoform 1 [Homo sapiens] | NP_002679.2 |
| 9951915 | AHCY | | adenosylhomocysteinase isoform 1 [Homo sapie | NP_000678.1 |
| 9966913 | ACTR3B | | actin-related protein 3B isoform 1 [Homo sapiens | NP_065178.1 |
| 10092657 | NDUFA12 | | NADH dehydrogenase [ubiquinone] 1 alpha subc | NP_061326.1 |
| 10092677 | PDXP | | pyridoxal phosphate phosphatase [Homo sapien | NP_064711.1 |
| 10518344 | RAP2A | | ras-related protein Rap-2a precursor [Homo sapi | NP_066361.1 |
| 10567816 | GNAO1 | | guanine nucleotide-binding protein G(o) subunit a | NP_066268.1 |
| 10800412 | MAPRE3 | | microtubule-associated protein RP/EB family me | NP_036458.2 |
| 10835063 | NPM1 | | nucleophosmin isoform 1 [Homo sapiens] | NP_002511.1 |
| 10863895 | TMSB10 | | thymosin beta-10 [Homo sapiens] | NP_066926.1 |
| 10863927 | PPIA | | peptidyl-prolyl cis-trans isomerase A isoform 1 [I | NP_066953.1 |
| 10863935 | RTN1 | | reticulon-1 isoform A [Homo sapiens] | NP_066959.1 |
| 11056044 | PPA1 | | inorganic pyrophosphatase [Homo sapiens] | NP_066952.1 |
| 11056046 | CADM3 | | cell adhesion molecule 3 isoform 1 precursor [Ho | NP_067012.1 |
| 11056061 | TMSB4X | | thymosin beta-4 [Homo sapiens] | NP_066932.1 |
| 11128019 | CYCS | | cytochrome c [Homo sapiens] | NP_061820.1 |
| 11225258 | MAG | | myelin-associated glycoprotein isoform a precurs | NP_002352.1 |
| 11321583 | SUCLA2 | | succinyl-CoA ligase [ADP-forming] subunit beta, | NP_003841.1 |
| 11545841 | OXCT2 | | succinyl-CoA:3-ketoacid coenzyme A transferas | NP_071403.1 |
| 11968182 | RPS18 | | 40S ribosomal protein S18 [Homo sapiens] | NP_072045.1 |
| 12025678 | ACTN4 | | alpha-actinin-4 [Homo sapiens] | NP_004915.2 |
| 12083581 | PLCB1 | | 1-phosphatidylinositol 4,5-bisphosphate phospho | NP_056007.1 |
| 12408675 | PFDN2 | | prefoldin subunit 2 [Homo sapiens] | NP_036526.2 |
| 12548785 | GABRB2 | | gamma-aminobutyric acid receptor subunit beta- | NP_068711.1 |
| 13259510 | DCTN1 | | dynactin subunit 1 isoform 1 [Homo sapiens] | NP_004073.2 |
| 13491174 | MARCKSL1 | | MARCKS-related protein [Homo sapiens] | NP_075385.1 |
| 13569885 | ITM2C | | integral membrane protein 2C isoform 1 [Homo s | NP_112188.1 |
| 13569962 | RAB1B | | ras-related protein Rab-1B [Homo sapiens] | NP_112243.1 |
| 13676857 | HSPA2 | | heat shock-related 70 kDa protein 2 [Homo sapi | NP_068814.2 |
| 13775198 | SH3BGRL3 | | SH3 domain-binding glutamic acid-rich-like protei | NP_112576.1 |
| 13775600 | SIRT2 | | NAD-dependent protein deacetylase sirtuin-2 isc | NP_036369.2 |
| 14043070 | HNRNPA1 | | heterogeneous nuclear ribonucleoprotein A1 isof | NP_112420.1 |
| 14043072 | HNRNPA2B1 | | heterogeneous nuclear ribonucleoproteins A2/B1 | NP_112533.1 |
| 14110420 | HNRNPD | | heterogeneous nuclear ribonucleoprotein D0 isof | NP_112738.1 |
| 14141152 | HNRNPM | | heterogeneous nuclear ribonucleoprotein M isofc | NP_005959.2 |
| 14141168 | PCBP2 | | poly(rC)-binding protein 2 isoform a [Homo sapie | NP_005007.2 |
| 14165437 | HNRNPK | | heterogeneous nuclear ribonucleoprotein K isofc | NP_112553.1 |
| 14210536 | TUBB6 | | tubulin beta-6 chain isoform 1 [Homo sapiens] | NP_115914.1 |
| 14249342 | INA | | alpha-internexin [Homo sapiens] | NP_116116.1 |
| 14277700 | RPS12 | | 40S ribosomal protein S12 [Homo sapiens] | NP_001007.2 |
| 14591909 | RPL5 | | 60S ribosomal protein L5 [Homo sapiens] | NP_000960.2 |
| 14719392 | CFL2 | | cofilin-2 isoform 1 [Homo sapiens] | NP_068733.1 |
| 15011936 | RPS26 | | 40S ribosomal protein S26 [Homo sapiens] | NP_001020.2 |
| 15055539 | RPS2 | | 40S ribosomal protein S2 [Homo sapiens] | NP_002943.2 |
| 15082258 | CBX3 | | chromobox protein homolog 3 [Homo sapiens] | NP_009207.2 |
| 15147219 | PURB | | transcriptional activator protein Pur-beta [Homo | NP_150093.1 |
| 15431288 | RPL10A | | 60S ribosomal protein L10a [Homo sapiens] | NP_009035.3 |

| | | | |
|---|---|---|---|
| 15431290 | RPL11 | 60S ribosomal protein L11 isoform 1 [Homo sapi | NP_000966.2 |
| 15431293 | RPL15 | 60S ribosomal protein L15 isoform 1 [Homo sapi | NP_002939.2 |
| 15431295 | RPL13 | 60S ribosomal protein L13 isoform 1 [Homo sapi | NP_150254.1 |
| 15431301 | RPL7 | 60S ribosomal protein L7 [Homo sapiens] | NP_000962.2 |
| 15812198 | FBXO2 | F-box only protein 2 [Homo sapiens] | NP_036300.2 |
| 16357472 | CDC42 | cell division control protein 42 homolog isoform 2 | NP_426359.1 |
| 16418379 | STX1B | syntaxin-1B [Homo sapiens] | NP_443106.1 |
| 16507237 | HSPA5 | 78 kDa glucose-regulated protein precursor [Hor | NP_005338.1 |
| 16579885 | RPL4 | 60S ribosomal protein L4 [Homo sapiens] | NP_000959.2 |
| 16753215 | PFN2 | profilin-2 isoform a [Homo sapiens] | NP_444252.1 |
| 16933546 | RPLP0 | 60S acidic ribosomal protein P0 [Homo sapiens] | NP_444505.1 |
| 17105394 | RPL23A | 60S ribosomal protein L23a [Homo sapiens] | NP_000975.2 |
| 17149836 | FKBP1A | peptidyl-prolyl cis-trans isomerase FKBP1A isof | NP_463460.1 |
| 17149842 | FKBP2 | peptidyl-prolyl cis-trans isomerase FKBP2 precu | NP_004461.2 |
| 17158044 | RPS6 | 40S ribosomal protein S6 [Homo sapiens] | NP_001001.2 |
| 17921989 | TUBA4A | tubulin alpha-4A chain isoform 1 [Homo sapiens] | NP_005991.1 |
| 17986258 | MYL6 | myosin light polypeptide 6 isoform 1 [Homo sapi | NP_066299.2 |
| 17986283 | TUBA1A | tubulin alpha-1A chain isoform 1 [Homo sapiens] | NP_006000.2 |
| 17999528 | COX6A1 | cytochrome c oxidase subunit 6A1, mitochondria | NP_004364.2 |
| 17999541 | VPS35 | vacuolar protein sorting-associated protein 35 [H | NP_060676.2 |
| 18079216 | CASKIN1 | caskin-1 [Homo sapiens] | NP_065815.1 |
| 18087855 | DYNLL2 | dynein light chain 2, cytoplasmic [Homo sapiens | NP_542408.1 |
| 18104948 | RPL21 | 60S ribosomal protein L21 [Homo sapiens] | NP_000973.2 |
| 18497300 | ATP6V1G2 | V-type proton ATPase subunit G 2 isoform a [Ho | NP_569730.1 |
| 18765733 | SNAP25 | synaptosomal-associated protein 25 isoform SN/ | NP_003072.2 |
| 19743875 | FH | fumarate hydratase, mitochondrial [Homo sapien | NP_000134.2 |
| 19913414 | AP2A1 | AP-2 complex subunit alpha-1 isoform 1 [Homo s | NP_055018.2 |
| 19913424 | ATP6V1A | V-type proton ATPase catalytic subunit A [Homo | NP_001681.2 |
| 19913428 | ATP6V1B2 | V-type proton ATPase subunit B, brain isoform [ | NP_001684.2 |
| 19913432 | ATP6V0D1 | V-type proton ATPase subunit d 1 [Homo sapien | NP_004682.2 |
| 19923142 | KPNB1 | importin subunit beta-1 isoform 1 [Homo sapiens | NP_002256.2 |
| 19923191 | MCM3AP | germinal-center associated nuclear protein [Hom | NP_003897.2 |
| 19923193 | ST13 | hsc70-interacting protein isoform 1 [Homo sapie | NP_003923.2 |
| 19923231 | RAB6A | ras-related protein Rab-6A isoform a [Homo sapi | NP_002860.2 |
| 19923233 | SCP2 | non-specific lipid-transfer protein isoform 1 prec | NP_002970.2 |
| 19923445 | ATL1 | atlastin-1 isoform a [Homo sapiens] | NP_056999.2 |
| 19923483 | RAB14 | ras-related protein Rab-14 [Homo sapiens] | NP_057406.2 |
| 19923748 | DLST | dihydrolipoyllysine-residue succinyltransferase c | NP_001924.2 |
| 19923750 | RAB3B | ras-related protein Rab-3B [Homo sapiens] | NP_002858.2 |
| 19923973 | KCTD12 | BTB/POZ domain-containing protein KCTD12 [Ho | NP_612453.1 |
| 19924099 | SYN1 | synapsin-1 isoform Ia [Homo sapiens] | NP_008881.2 |
| 19924103 | SYN2 | synapsin-2 isoform IIa [Homo sapiens] | NP_598328.1 |
| 20070125 | P4HB | protein disulfide-isomerase precursor [Homo sap | NP_000909.2 |
| 20127450 | PRKCB | protein kinase C beta type isoform 2 [Homo sapi | NP_002729.2 |
| 20149568 | NDUFV1 | NADH dehydrogenase [ubiquinone] flavoprotein | NP_009034.2 |
| 20149594 | HSP90AB1 | heat shock protein HSP 90-beta isoform a [Hom | NP_031381.2 |
| 20149675 | EFHD2 | EF-hand domain-containing protein D2 [Homo sa | NP_077305.2 |
| 20336761 | HEBP1 | heme-binding protein 1 [Homo sapiens] | NP_057071.2 |
| 20357529 | GNB2 | guanine nucleotide-binding protein G(I)/G(S)/G(T | NP_005264.2 |
| 21359867 | CYC1 | cytochrome c1, heme protein, mitochondrial prec | NP_001907.2 |
| 21361091 | UCHL1 | ubiquitin carboxyl-terminal hydrolase isozyme L1 | NP_004172.2 |
| 21361103 | SLC25A12 | calcium-binding mitochondrial carrier protein Aral | NP_003696.2 |
| 21361114 | SLC25A11 | mitochondrial 2-oxoglutarate/malate carrier prote | NP_003553.2 |
| 21361176 | ALDH1A1 | retinal dehydrogenase 1 [Homo sapiens] | NP_000680.2 |
| 21361181 | ATP1A1 | sodium/potassium-transporting ATPase subunit a | NP_000692.2 |
| 21361337 | EIF5 | eukaryotic translation initiation factor 5 [Homo s | NP_001960.2 |
| 21361370 | PYGB | glycogen phosphorylase, brain form [Homo sapie | NP_002853.2 |
| 21361399 | PPP2R1A | serine/threonine-protein phosphatase 2A 65 kDa | NP_055040.2 |
| 21361559 | VSNL1 | visinin-like protein 1 [Homo sapiens] | NP_003376.2 |
| 21361619 | TOLLIP | toll-interacting protein [Homo sapiens] | NP_061882.2 |
| 21361647 | AHCYL1 | adenosylhomocysteinase 2 isoform a [Homo sap | NP_006612.2 |
| 21361657 | PDIA3 | protein disulfide-isomerase A3 precursor [Homo | NP_005304.3 |

| | | | |
|---|---|---|---|
| 21361794 | CAND1 | cullin-associated NEDD8-dissociated protein 1 [I | NP_060918.2 |
| 21464101 | YWHAG | 14-3-3 protein gamma [Homo sapiens] | NP_036611.2 |
| 21536286 | CKB | creatine kinase B-type [Homo sapiens] | NP_001814.2 |
| 21626466 | MATR3 | matrin-3 isoform a [Homo sapiens] | NP_061322.2 |
| 21686977 | CADM4 | cell adhesion molecule 4 precursor [Homo sapie | NP_660339.1 |
| 21703367 | DIRAS2 | GTP-binding protein Di-Ras2 [Homo sapiens] | NP_060064.2 |
| 21735492 | PPP1R1B | protein phosphatase 1 regulatory subunit 1B iso | NP_115568.2 |
| 21735621 | MDH2 | malate dehydrogenase, mitochondrial isoform 1 | | NP_005909.2 |
| 22035696 | SYNGR1 | synaptogyrin-1 isoform 1a [Homo sapiens] | NP_004702.2 |
| 22538467 | PSMB4 | proteasome subunit beta type-4 [Homo sapiens] | NP_002787.2 |
| 22907052 | ARPC1A | actin-related protein 2/3 complex subunit 1A iso | NP_006400.2 |
| 23065544 | GSTM1 | glutathione S-transferase Mu 1 isoform 1 [Homo | NP_000552.2 |
| 23110925 | PSMB6 | proteasome subunit beta type-6 isoform 1 precu | NP_002789.1 |
| 23110935 | PSMA1 | proteasome subunit alpha type-1 isoform 1 [Hom | NP_683877.1 |
| 23110942 | PSMA5 | proteasome subunit alpha type-5 isoform 1 [Hom | NP_002781.2 |
| 23503295 | CSNK2B | casein kinase II subunit beta isoform 1 [Homo s | NP_001311.3 |
| 24234688 | HSPA9 | stress-70 protein, mitochondrial precursor [Hom | NP_004125.3 |
| 24307939 | CCT5 | T-complex protein 1 subunit epsilon isoform a [H | NP_036205.1 |
| 24307949 | NCDN | neurochondrin isoform 1 [Homo sapiens] | NP_055099.1 |
| 24307999 | GPD1L | glycerol-3-phosphate dehydrogenase 1-like prote | NP_055956.1 |
| 24308071 | CNRIP1 | CB1 cannabinoid receptor-interacting protein 1 is | NP_056278.1 |
| 24308257 | VAT1L | synaptic vesicle membrane protein VAT-1 homol | NP_065978.1 |
| 24497435 | PSMC5 | 26S protease regulatory subunit 8 isoform 1 [Ho | NP_002796.4 |
| 24638454 | ATP2A2 | sarcoplasmic/endoplasmic reticulum calcium ATI | NP_733765.1 |
| 25777713 | SKP1 | S-phase kinase-associated protein 1 isoform b [ | NP_733779.1 |
| 25777732 | ALDH2 | aldehyde dehydrogenase, mitochondrial isoform | NP_000681.2 |
| 25952114 | CAMK2A | calcium/calmodulin-dependent protein kinase typ | NP_057065.2 |
| 27436946 | LMNA | lamin isoform A [Homo sapiens] | NP_733821.1 |
| 27544939 | BRK1 | protein BRICK1 [Homo sapiens] | NP_060932.2 |
| 27764867 | SYP | synaptophysin [Homo sapiens] | NP_003170.1 |
| 28178832 | IDH2 | isocitrate dehydrogenase [NADP], mitochondrial | NP_002159.2 |
| 28302131 | HBG1 | hemoglobin subunit gamma-1 [Homo sapiens] | NP_000550.2 |
| 28872725 | PSMD11 | 26S proteasome non-ATPase regulatory subunit | NP_002806.2 |
| 29171702 | PPA2 | inorganic pyrophosphatase 2, mitochondrial isof | NP_789845.1 |
| 29788768 | TUBB2B | tubulin beta-2B chain [Homo sapiens] | NP_821080.1 |
| 29788785 | TUBB | tubulin beta chain isoform b [Homo sapiens] | NP_821133.1 |
| 29826321 | ADD1 | alpha-adducin isoform b [Homo sapiens] | NP_054908.2 |
| 30089916 | PACS1 | phosphofurin acidic cluster sorting protein 1 [Ho | NP_060496.2 |
| 30795231 | BASP1 | brain acid soluble protein 1 [Homo sapiens] | NP_006308.3 |
| 31541941 | HSPA4L | heat shock 70 kDa protein 4L [Homo sapiens] | NP_055093.2 |
| 31542528 | NECAP1 | adaptin ear-binding coat-associated protein 1 [H | NP_056324.2 |
| 31542947 | HSPD1 | 60 kDa heat shock protein, mitochondrial [Homo | NP_002147.2 |
| 31621303 | SFXN3 | sideroflexin-3 [Homo sapiens] | NP_112233.2 |
| 31711992 | DLAT | dihydrolipoyllysine-residue acetyltransferase coi | NP_001922.2 |
| 32189362 | PPFIA3 | liprin-alpha-3 [Homo sapiens] | NP_003651.1 |
| 32189394 | ATP5B | ATP synthase subunit beta, mitochondrial precu | NP_001677.2 |
| 32307148 | OGT | UDP-N-acetylglucosamine--peptide N-acetylgluc | NP_858058.1 |
| 32483416 | NEFH | neurofilament heavy polypeptide [Homo sapiens] | NP_066554.2 |
| 32528282 | ACOT7 | cytosolic acyl coenzyme A thioester hydrolase i | NP_863654.1 |
| 33350932 | DYNC1H1 | cytoplasmic dynein 1 heavy chain 1 [Homo sapi | NP_001367.2 |
| 33457311 | MDP1 | magnesium-dependent phosphatase 1 isoform 1 | NP_612485.2 |
| 33589861 | RAB31 | ras-related protein Rab-31 [Homo sapiens] | NP_006859.2 |
| 33620747 | MYEF2 | myelin expression factor 2 isoform a [Homo sapi | NP_057216.2 |
| 33946324 | GNAI1 | guanine nucleotide-binding protein G(i) subunit a | NP_002060.4 |
| 33946329 | RALA | ras-related protein Ral-A precursor [Homo sapier | NP_005393.2 |
| 34147513 | RAB7A | ras-related protein Rab-7a [Homo sapiens] | NP_004628.4 |
| 34147630 | TUFM | elongation factor Tu, mitochondrial precursor [H | NP_003312.3 |
| 34452715 | CADPS | calcium-dependent secretion activator 1 isoform | NP_003707.2 |
| 34482047 | AP3B2 | AP-3 complex subunit beta-2 isoform 2 [Homo s | NP_004635.2 |
| 34577061 | ADH1B | alcohol dehydrogenase 1B isoform 1 [Homo sapi | NP_000659.2 |
| 38327039 | HSPA4 | heat shock 70 kDa protein 4 [Homo sapiens] | NP_002145.3 |
| 38327625 | CS | citrate synthase, mitochondrial precursor [Homo | NP_004068.2 |

| | | | | |
|---|---|---|---|---|
| 38455427 | CCT4 | | T-complex protein 1 subunit delta isoform a [Hor | NP_006421.2 |
| 40254462 | GNAQ | | guanine nucleotide-binding protein G(q) subunit a | NP_002063.2 |
| 40254947 | TMX4 | | thioredoxin-related transmembrane protein 4 prec | NP_066979.2 |
| 40538799 | UBQLN4 | | ubiquilin-4 isoform 1 [Homo sapiens] | NP_064516.2 |
| 41349456 | PREP | | prolyl endopeptidase [Homo sapiens] | NP_002717.3 |
| 41393561 | LAP3 | | cytosol aminopeptidase [Homo sapiens] | NP_056991.2 |
| 42716277 | C2CD2L | | C2 domain-containing protein 2-like isoform 1 [H | NP_055622.3 |
| 42794769 | PAK1 | | serine/threonine-protein kinase PAK 1 isoform 2 | NP_002567.3 |
| 45439359 | TRIO | | triple functional domain protein [Homo sapiens] | NP_009049.2 |
| 45545411 | NCKAP1 | | nck-associated protein 1 isoform 2 [Homo sapie | NP_995314.1 |
| 45827776 | RTN1 | | reticulon-1 isoform C [Homo sapiens] | NP_996734.1 |
| 46249393 | RHOG | | rho-related GTP-binding protein RhoG precursor | NP_001656.2 |
| 46389550 | ENSA | | alpha-endosulfine isoform 1 [Homo sapiens] | NP_996925.1 |
| 46593007 | UQCRC1 | | cytochrome b-c1 complex subunit 1, mitochondr | NP_003356.2 |
| 47132585 | PRKAR2B | | cAMP-dependent protein kinase type II-beta reg | NP_002727.2 |
| 47717102 | ATP6V1H | | V-type proton ATPase subunit H isoform 1 [Hom | NP_998785.1 |
| 48255951 | ATP2B2 | | plasma membrane calcium-transporting ATPase | NP_001001331.1 |
| 48255966 | UGP2 | | UTP--glucose-1-phosphate uridylyltransferase is | NP_006750.3 |
| 48375173 | MAP6 | | microtubule-associated protein 6 isoform 1 [Hom | NP_149052.1 |
| 48762918 | PCP4 | | Purkinje cell protein 4 [Homo sapiens] | NP_006189.2 |
| 48762932 | CCT8 | | T-complex protein 1 subunit theta isoform 1 [Hor | NP_006576.2 |
| 49574491 | ATP1B2 | | sodium/potassium-transporting ATPase subunit I | NP_001669.3 |
| 49574532 | GSK3A | | glycogen synthase kinase-3 alpha [Homo sapier | NP_063937.2 |
| 50263048 | GPM6B | | neuronal membrane glycoprotein M6-b isoform 1 | NP_001001995.1 |
| 50345984 | ATP5A1 | | ATP synthase subunit alpha, mitochondrial isofo | NP_001001937.1 |
| 50345988 | ATP5C1 | | ATP synthase subunit gamma, mitochondrial iso | NP_001001973.1 |
| 50345991 | ATP5D | | ATP synthase subunit delta, mitochondrial precu | NP_001001975.1 |
| 50592988 | UQCRC2 | | cytochrome b-c1 complex subunit 2, mitochondr | NP_003357.2 |
| 50592996 | TUBB3 | | tubulin beta-3 chain isoform 1 [Homo sapiens] | NP_006077.2 |
| 51092270 | C1orf95 | | protein stum homolog [Homo sapiens] | NP_001003665.1 |
| 51317370 | NDUFA6 | | NADH dehydrogenase [ubiquinone] 1 alpha subc | NP_002481.2 |
| 52426787 | OMG | | oligodendrocyte-myelin glycoprotein precursor [H | NP_002535.3 |
| 52630440 | FKBP8 | | peptidyl-prolyl cis-trans isomerase FKBP8 isofor | NP_036313.3 |
| 52632383 | HNRNPL | | heterogeneous nuclear ribonucleoprotein L isofo | NP_001524.2 |
| 54112397 | CACNA2D3 | | voltage-dependent calcium channel subunit alph | NP_060868.2 |
| 54607031 | CADM2 | | cell adhesion molecule 2 isoform 3 precursor [H | NP_694854.2 |
| 54607064 | CACNB4 | | voltage-dependent L-type calcium channel subu | NP_000717.2 |
| 54607135 | TOMM70A | | mitochondrial import receptor subunit TOM70 [H | NP_055635.3 |
| 55749577 | SLC25A4 | | ADP/ATP translocase 1 [Homo sapiens] | NP_001142.2 |
| 55770834 | CENPF | | centromere protein F [Homo sapiens] | NP_057427.3 |
| 55770878 | NPTX1 | | neuronal pentraxin-1 precursor [Homo sapiens] | NP_002513.2 |
| 55956919 | HNRNPAB | | heterogeneous nuclear ribonucleoprotein A/B isc | NP_112556.2 |
| 56549123 | DNM2 | | dynamin-2 isoform 2 [Homo sapiens] | NP_001005361.1 |
| 56549135 | TAGLN3 | | transgelin-3 [Homo sapiens] | NP_037391.2 |
| 56676375 | TPPP3 | | tubulin polymerization-promoting protein family m | NP_057224.2 |
| 56788381 | MOG | | myelin-oligodendrocyte glycoprotein isoform alpl | NP_996532.2 |
| 57163987 | SLC8A2 | | sodium/calcium exchanger 2 precursor [Homo sa | NP_055878.1 |
| 57165410 | ACSL6 | | long-chain-fatty-acid--CoA ligase 6 isoform a [H | NP_056071.2 |
| 57863257 | TCP1 | | T-complex protein 1 subunit alpha isoform a [Ho | NP_110379.2 |
| 58331274 | | 5-Sep | septin-5 isoform 2 [Homo sapiens] | NP_001009939.1 |
| 58761500 | OLA1 | | obg-like ATPase 1 isoform 1 [Homo sapiens] | NP_037473.3 |
| 60685231 | TTC7B | | tetratricopeptide repeat protein 7B [Homo sapier | NP_001010854.1 |
| 61175239 | TCEAL5 | | transcription elongation factor A protein-like 5 [H | NP_001012997.1 |
| 61743942 | ABI1 | | abl interactor 1 isoform a [Homo sapiens] | NP_005461.2 |
| 62420877 | ETFB | | electron transfer flavoprotein subunit beta isofor | NP_001014763.1 |
| 62422571 | CRMP1 | | dihydropyrimidinase-related protein 1 isoform 1 [ | NP_001014809.1 |
| 63162572 | CCT3 | | T-complex protein 1 subunit gamma isoform a [H | NP_005989.3 |
| 66346679 | SERBP1 | | plasminogen activator inhibitor 1 RNA-binding pr | NP_001018077.1 |
| 66392205 | NME2 | | nucleoside diphosphate kinase B isoform a [Hon | NP_001018148.1 |
| 66932916 | MAPK1 | | mitogen-activated protein kinase 1 [Homo sapier | NP_002736.3 |
| 67782307 | SOD2 | | superoxide dismutase [Mn], mitochondrial isofor | NP_001019636.1 |
| 68163411 | ALCAM | | CD166 antigen isoform 1 precursor [Homo sapie | NP_001618.2 |

| | | | |
|---|---|---|---|
| 68303561 | PSMA8 | proteasome subunit alpha type-7-like isoform 1 [ | NP_653263.2 |
| 68509930 | MBP | myelin basic protein isoform 1 [Homo sapiens] | NP_001020252.1 |
| 71773329 | ANXA6 | annexin A6 isoform 1 [Homo sapiens] | NP_001146.2 |
| 72534660 | SRSF7 | serine/arginine-rich splicing factor 7 isoform 1 [H | NP_001026854.1 |
| 73486658 | GOT2 | aspartate aminotransferase, mitochondrial isofor | NP_002071.2 |
| 73760415 | STXBP1 | syntaxin-binding protein 1 isoform b [Homo sapie | NP_001027392.1 |
| 74136883 | HNRNPU | heterogeneous nuclear ribonucleoprotein U isofo | NP_114032.2 |
| 74271837 | GLUL | glutamine synthetase [Homo sapiens] | NP_001028216.1 |
| 76159293 | THEM4 | acyl-coenzyme A thioesterase THEM4 [Homo sa | NP_444283.2 |
| 78000181 | RPL14 | 60S ribosomal protein L14 [Homo sapiens] | NP_003964.3 |
| 83267868 | DYNLL1 | dynein light chain 1, cytoplasmic [Homo sapiens | NP_001032584.1 |
| 83776596 | CAMKV | caM kinase-like vesicle-associated protein [Hom | NP_076951.2 |
| 84875539 | PRPS2 | ribose-phosphate pyrophosphokinase 2 isoform | NP_001034180.1 |
| 87196351 | DDX3X | ATP-dependent RNA helicase DDX3X isoform 1 [ | NP_001347.3 |
| 91199540 | DLD | dihydrolipoyl dehydrogenase, mitochondrial isofo | NP_000099.2 |
| 91208420 | BSN | protein bassoon [Homo sapiens] | NP_003449.2 |
| 91208428 | PTPRZ1 | receptor-type tyrosine-protein phosphatase zeta | NP_002842.2 |
| 94721250 | VAPA | vesicle-associated membrane protein-associated | NP_003565.4 |
| 94721261 | CNP | 2',3'-cyclic-nucleotide 3'-phosphodiesterase [Ho | NP_149124.3 |
| 95147555 | MAP1A | microtubule-associated protein 1A [Homo sapien | NP_002364.5 |
| 96975097 | RAB6B | ras-related protein Rab-6B [Homo sapiens] | NP_057661.3 |
| 98986464 | TMED10 | transmembrane emp24 domain-containing protein | NP_006818.3 |
| 103472001 | NDUFA7 | NADH dehydrogenase [ubiquinone] 1 alpha subc | NP_004992.2 |
| 105990539 | NEFL | neurofilament light polypeptide [Homo sapiens] | NP_006149.2 |
| 108773797 | GDAP1 | ganglioside-induced differentiation-associated pr | NP_061845.2 |
| 109148508 | OTUB1 | ubiquitin thioesterase OTUB1 [Homo sapiens] | NP_060140.2 |
| 109240550 | PSPC1 | paraspeckle component 1 [Homo sapiens] | NP_001035879.1 |
| 109452591 | SUCLG1 | succinyl-CoA ligase [ADP/GDP-forming] subunit | NP_003840.2 |
| 112382250 | SPTBN1 | spectrin beta chain, non-erythrocytic 1 isoform 1 | NP_003119.2 |
| 113204622 | SMAP1 | stromal membrane-associated protein 1 isoform | NP_001037770.1 |
| 114155144 | TPM3 | tropomyosin alpha-3 chain isoform Tpm3.2cy [Ho | NP_001036816.1 |
| 115334682 | SRCIN1 | SRC kinase signaling inhibitor 1 [Homo sapiens] | NP_079524.2 |
| 115387094 | SDHB | succinate dehydrogenase [ubiquinone] iron-sulfu | NP_002991.2 |
| 115511049 | GNA11 | guanine nucleotide-binding protein subunit alpha | NP_002058.2 |
| 116256358 | SCAMP1 | secretory carrier-associated membrane protein 1 | NP_004857.4 |
| 117938759 | GNAS | protein GNAS isoform XLas [Homo sapiens] | NP_536350.2 |
| 118402586 | GLO1 | lactoylglutathione lyase [Homo sapiens] | NP_006699.2 |
| 140972063 | PPP1R9B | neurabin-2 [Homo sapiens] | NP_115984.3 |
| 148233338 | CTNNB1 | catenin beta-1 [Homo sapiens] | NP_001091679.1 |
| 148277037 | AAK1 | AP2-associated protein kinase 1 [Homo sapiens | NP_055726.3 |
| 148529011 | CORO2B | coronin-2B isoform 1 [Homo sapiens] | NP_006082.3 |
| 148529014 | DDB1 | DNA damage-binding protein 1 [Homo sapiens] | NP_001914.3 |
| 148612849 | KIF2A | kinesin-like protein KIF2A isoform 2 [Homo sapie | NP_001091981.1 |
| 148727331 | USP5 | ubiquitin carboxyl-terminal hydrolase 5 isoform 1 | NP_001092006.1 |
| 150170704 | PHYHIP | phytanoyl-CoA hydroxylase-interacting protein [I | NP_055574.3 |
| 153070260 | MARCKS | myristoylated alanine-rich C-kinase substrate [H | NP_002347.5 |
| 153792590 | HSP90AA1 | heat shock protein HSP 90-alpha isoform 1 [Hon | NP_001017963.2 |
| 153945728 | MAP1B | microtubule-associated protein 1B [Homo sapien | NP_005900.2 |
| 153946409 | CALB2 | calretinin isoform 1 [Homo sapiens] | NP_001731.2 |
| 154354964 | IMMT | MICOS complex subunit MIC60 isoform 1 [Homo | NP_006830.2 |
| 155030192 | RAP1GDS1 | rap1 GTPase-GDP dissociation stimulator 1 isofo | NP_001093896.1 |
| 156071459 | SLC25A5 | ADP/ATP translocase 2 [Homo sapiens] | NP_001143.2 |
| 156104878 | GLS | glutaminase kidney isoform, mitochondrial isofor | NP_055720.3 |
| 156104880 | GMPR | GMP reductase 1 [Homo sapiens] | NP_006868.3 |
| 156151366 | LGALSL | galectin-related protein [Homo sapiens] | NP_054900.2 |
| 156416003 | SDHA | succinate dehydrogenase [ubiquinone] flavoprot | NP_004159.2 |
| 156564403 | PDHB | pyruvate dehydrogenase E1 component subunit | NP_000916.2 |
| 156602655 | 3-Sep | neuronal-specific septin-3 isoform A [Homo sapi | NP_663786.2 |
| 157384973 | TNR | tenascin-R precursor [Homo sapiens] | NP_003276.3 |
| 157412328 | CORO1C | coronin-1C isoform a [Homo sapiens] | NP_001098707.1 |
| 157649073 | CAP1 | adenylyl cyclase-associated protein 1 [Homo sa | NP_001099000.1 |
| 157738649 | NEFM | neurofilament medium polypeptide isoform 1 [Ho | NP_005373.2 |

| | | | |
|---|---|---|---|
| 158937236 | NPEPPS | puromycin-sensitive aminopeptidase [Homo sapi | NP_006301.3 |
| 163644321 | UQCRFS1 | cytochrome b-c1 complex subunit Rieske, mitoc | NP_005994.2 |
| 163659856 | GRIA3 | glutamate receptor 3 isoform 1 precursor [Homo | NP_015564.4 |
| 164419734 | GRIA4 | glutamate receptor 4 isoform 1 precursor [Homo | NP_000820.3 |
| 166706879 | DMTN | dematin isoform 1 [Homo sapiens] | NP_001107608.1 |
| 166795297 | DYNC1LI1 | cytoplasmic dynein 1 light intermediate chain 1 [ | NP_057225.2 |
| 169404009 | SSR1 | translocon-associated protein subunit alpha isof | NP_003135.2 |
| 170932473 | ZNF786 | zinc finger protein 786 [Homo sapiens] | NP_689624.2 |
| 183076548 | HIST2H3D | histone H3.2 [Homo sapiens] | NP_001116847.1 |
| 187281616 | NDUFS7 | NADH dehydrogenase [ubiquinone] iron-sulfur pr | NP_077718.3 |
| 189083724 | GABRA1 | gamma-aminobutyric acid receptor subunit alpha | NP_001121115.1 |
| 190194363 | DPYSL4 | dihydropyrimidinase-related protein 4 [Homo sap | NP_006417.2 |
| 190358517 | RAB11B | ras-related protein Rab-11B [Homo sapiens] | NP_004209.2 |
| 190885499 | COX5A | cytochrome c oxidase subunit 5A, mitochondrial | NP_004246.2 |
| 192449447 | PLP1 | myelin proteolipid protein isoform 1 [Homo sapie | NP_001122306.1 |
| 194018537 | PRPSAP1 | phosphoribosyl pyrophosphate synthase-associ | NP_002757.2 |
| 194248068 | SYNGAP1 | ras/Rap GTPase-activating protein SynGAP [Ho | NP_006763.2 |
| 194272161 | ITPKB | inositol-trisphosphate 3-kinase B [Homo sapiens | NP_002212.3 |
| 194394161 | LRRC57 | leucine-rich repeat-containing protein 57 [Homo s | NP_694992.2 |
| 194473724 | AP1M1 | AP-1 complex subunit mu-1 isoform 1 [Homo sa | NP_001123996.1 |
| 194578909 | EIF4E | eukaryotic translation initiation factor 4E isoforn | NP_001124151.1 |
| 197927160 | SLC4A4 | electrogenic sodium bicarbonate cotransporter 1 | NP_001128214.1 |
| 198041678 | SLC12A5 | solute carrier family 12 member 5 isoform 1 [Hor | NP_001128243.1 |
| 208879465 | VDAC3 | voltage-dependent anion-selective channel prote | NP_001129166.1 |
| 208973246 | QDPR | dihydropteridine reductase isoform 1 [Homo sapi | NP_000311.2 |
| 214010226 | RPS24 | 40S ribosomal protein S24 isoform d [Homo sapi | NP_001135757.1 |
| 215422338 | PDK3 | pyruvate dehydrogenase kinase, isozyme 3 isof | NP_001135858.1 |
| 216548223 | SV2A | synaptic vesicle glycoprotein 2A isoform 1 [Hom | NP_055664.3 |
| 217330598 | GLOD4 | glyoxalase domain-containing protein 4 [Homo s | NP_057164.3 |
| 219555707 | EIF5A | eukaryotic translation initiation factor 5A-1 isofo | NP_001137232.1 |
| 219804203 | RPH3A | rabphilin-3A isoform 1 [Homo sapiens] | NP_001137326.1 |
| 221307584 | PHB2 | prohibitin-2 isoform 1 [Homo sapiens] | NP_001138303.1 |
| 221625487 | IMPA1 | inositol monophosphatase 1 isoform 2 [Homo sa | NP_001138350.1 |
| 222080062 | NDUFV2 | NADH dehydrogenase [ubiquinone] flavoprotein 2 | NP_066552.2 |
| 222352151 | PCBP1 | poly(rC)-binding protein 1 [Homo sapiens] | NP_006187.2 |
| 223556012 | ADGRF3 | adhesion G-protein coupled receptor F3 isoform | NP_001138640.1 |
| 224451142 | STMN1 | stathmin isoform b [Homo sapiens] | NP_001138926.1 |
| 224465190 | SCRN1 | secernin-1 isoform b [Homo sapiens] | NP_001138986.1 |
| 224831253 | OPA1 | dynamin-like 120 kDa protein, mitochondrial isof | NP_570850.2 |
| 225543166 | SAMM50 | sorting and assembly machinery component 50 | NP_056195.3 |
| 226246671 | RPS20 | 40S ribosomal protein S20 isoform 1 [Homo sapi | NP_001139699.1 |
| 226529917 | TPI1 | triosephosphate isomerase isoform 2 [Homo sap | NP_001152759.1 |
| 237757310 | SYNJ1 | synaptojanin-1 isoform a [Homo sapiens] | NP_003886.3 |
| 241982780 | PDCD6IP | programmed cell death 6-interacting protein isof | NP_001155901.1 |
| 256222019 | RAB10 | ras-related protein Rab-10 [Homo sapiens] | NP_057215.3 |
| 258613969 | PRKAR1B | cAMP-dependent protein kinase type I-beta regu | NP_001158230.1 |
| 260099723 | LDHA | L-lactate dehydrogenase A chain isoform 3 [Hon | NP_001158886.1 |
| 260436862 | AP1B1 | AP-1 complex subunit beta-1 isoform a [Homo s | NP_001118.3 |
| 271398239 | CNDP2 | cytosolic non-specific dipeptidase isoform 1 [Ho | NP_060705.2 |
| 283436220 | ATAD3A | ATPase family AAA domain-containing protein 3 | NP_060658.3 |
| 285002233 | GPD2 | glycerol-3-phosphate dehydrogenase, mitochonc | NP_000399.3 |
| 289666746 | FBXL16 | F-box/LRR-repeat protein 16 [Homo sapiens] | NP_699181.2 |
| 290463102 | PGM1 | phosphoglucomutase-1 isoform 2 [Homo sapiens | NP_001166289.1 |
| 291575128 | LDHB | L-lactate dehydrogenase B chain isoform LDHB | NP_001167568.1 |
| 294862256 | SUCLG2 | succinyl-CoA ligase [GDP-forming] subunit beta, | NP_001171070.1 |
| 296080746 | CTTN | src substrate cortactin isoform c [Homo sapiens | NP_001171669.1 |
| 296317337 | VDAC2 | voltage-dependent anion-selective channel prote | NP_001171712.1 |
| 308818195 | DPYSL2 | dihydropyrimidinase-related protein 2 isoform 1 [ | NP_001184222.1 |
| 311771647 | ARPC4 | actin-related protein 2/3 complex subunit 4 isofo | NP_001185709.1 |
| 313569822 | RPL17-C18orf32 | RPL17-C18orf32 protein isoform 1 [Homo sapien | NP_001186284.1 |
| 315259111 | NEDD8-MDP1 | NEDD8-MDP1 protein [Homo sapiens] | NP_001186752.1 |
| 315434249 | KCNAB2 | voltage-gated potassium channel subunit beta-2 | NP_001186791.1 |

| | | | |
|---|---|---|---|
| 316659409 | ACTG1 | actin, cytoplasmic 2 [Homo sapiens] | NP_001186883.1 |
| 316983160 | NDUFS1 | NADH-ubiquinone oxidoreductase 75 kDa subuni | NP_001186913.1 |
| 321117084 | RPS10-NUDT3 | RPS10-NUDT3 protein [Homo sapiens] | NP_001189399.1 |
| 332164775 | PKM | pyruvate kinase PKM isoform c [Homo sapiens] | NP_001193725.1 |
| 333944015 | CDH13 | cadherin-13 isoform 2 [Homo sapiens] | NP_001207417.1 |
| 334688844 | GFAP | glial fibrillary acidic protein isoform 3 [Homo sapi | NP_001229305.1 |
| 336285440 | NCAM1 | neural cell adhesion molecule 1 isoform 5 precur | NP_001229536.1 |
| 338827685 | AP2A2 | AP-2 complex subunit alpha-2 isoform 1 [Homo s | NP_001229766.1 |
| 342187211 | ALDOA | fructose-bisphosphate aldolase A isoform 2 [Hor | NP_001230106.1 |
| 354721184 | RAB5C | ras-related protein Rab-5C isoform b [Homo sapi | NP_001238968.1 |
| 354983493 | PCMT1 | protein-L-isoaspartate(D-aspartate) O-methyltrar | NP_001238978.1 |
| 363498929 | UQCRB | cytochrome b-c1 complex subunit 7 isoform 3 [H | NP_001241681.1 |
| 371940940 | ATP1A3 | sodium/potassium-transporting ATPase subunit a | NP_001243143.1 |
| 373432684 | UBE2L3 | ubiquitin-conjugating enzyme E2 L3 isoform 4 [H | NP_001243284.1 |
| 374081854 | RAB18 | ras-related protein Rab-18 isoform 2 [Homo sapi | NP_001243339.1 |
| 379030615 | DNAJC6 | putative tyrosine-protein phosphatase auxilin isc | NP_001243793.1 |
| 380837121 | SEC22B | vesicle-trafficking protein SEC22b precursor [Hc | NP_004883.3 |
| 384475521 | TKT | transketolase isoform 2 [Homo sapiens] | NP_001244957.1 |
| 386869503 | RPS3 | 40S ribosomal protein S3 isoform 2 [Homo sapie | NP_001247435.1 |
| 393195306 | ALDH1L1 | cytosolic 10-formyltetrahydrofolate dehydrogena | NP_001257293.1 |
| 409971401 | PPME1 | protein phosphatase methylesterase 1 isoform b | NP_001258522.1 |
| 430727947 | PPP2R2B | serine/threonine-protein phosphatase 2A 55 kDa | NP_858061.2 |
| 472235318 | FMN1 | formin-1 isoform a [Homo sapiens] | NP_001264242.1 |
| 510937025 | DNM1L | dynamin-1-like protein isoform 5 [Homo sapiens] | NP_001265393.1 |
| 514052670 | ARPC3 | actin-related protein 2/3 complex subunit 3 isofc | NP_001265485.1 |
| 520975477 | RANBP1 | ran-specific GTPase-activating protein isoform 1 | NP_001265568.1 |
| 525507379 | CPNE6 | copine-6 isoform 1 [Homo sapiens] | NP_001267487.1 |
| 527498279 | PHB | prohibitin isoform 1 [Homo sapiens] | NP_001268425.1 |
| 530360487 | PARK7 | protein DJ-1 isoform X1 [Homo sapiens] | XP_005263481.1 |
| 530363144 | SFPQ | splicing factor, proline- and glutamine-rich isofor | XP_005271170.1 |
| 530365929 | BPNT1 | 3'(2'),5'-bisphosphate nucleotidase 1 isoform X2 | XP_005273056.1 |
| 530366381 | RGS7 | regulator of G-protein signaling 7 isoform X1 [Hc | XP_005273275.1 |
| 530366978 | ROCK2 | rho-associated protein kinase 2 isoform X1 [Hon | XP_005246247.1 |
| 530369146 | BIN1 | myc box-dependent-interacting protein 1 isoform | XP_005263699.1 |
| 530369643 | ABI2 | abl interactor 2 isoform X2 [Homo sapiens] | XP_005246274.1 |
| 530369695 | LANCL1 | lanC-like protein 1 isoform X1 [Homo sapiens] | XP_005246300.1 |
| 530369990 | HNRNPA3 | heterogeneous nuclear ribonucleoprotein A3 isof | XP_005246437.1 |
| 530371611 | KIF1A | kinesin-like protein KIF1A isoform X2 [Homo sap | XP_005247079.1 |
| 530372834 | PRKAR2A | cAMP-dependent protein kinase type II-alpha re | XP_005265370.1 |
| 530374617 | LSAMP | limbic system-associated membrane protein isof | XP_005247511.1 |
| 530375053 | HEG1 | protein HEG homolog 1 isoform X1 [Homo sapier | XP_005247723.1 |
| 530378226 | CAMK2D | calcium/calmodulin-dependent protein kinase typ | XP_005263308.1 |
| 530378443 | RAPGEF2 | rap guanine nucleotide exchange factor 2 isofor | XP_005263415.1 |
| 530378702 | CTNND2 | catenin delta-2 isoform X1 [Homo sapiens] | XP_005248308.1 |
| 530380562 | PCDH1 | protocadherin-1 isoform X1 [Homo sapiens] | XP_005268509.1 |
| 530380819 | CPLX2 | complexin-2 isoform X1 [Homo sapiens] | XP_005265856.1 |
| 530381854 | HIST1H2BD | histone H2B type 1-D isoform X1 [Homo sapiens | XP_005249096.1 |
| 530383411 | EPB41L2 | band 4.1-like protein 2 isoform X3 [Homo sapien | XP_005266897.1 |
| 530387660 | PTK2B | protein-tyrosine kinase 2-beta isoform X1 [Homo | XP_005273504.1 |
| 530388568 | EFCAB1 | EF-hand calcium-binding domain-containing prote | XP_005251360.1 |
| 530388906 | PABPC1 | polyadenylate-binding protein 1 isoform X1 [Hom | XP_005250918.1 |
| 530389143 | PLEC | plectin isoform X3 [Homo sapiens] | XP_005251033.1 |
| 530389317 | YWHAZ | 14-3-3 protein zeta/delta isoform X1 [Homo sapi | XP_005251118.1 |
| 530390646 | DNM1 | dynamin-1 isoform X4 [Homo sapiens] | XP_005251820.1 |
| 530390694 | AK1 | adenylate kinase isoenzyme 1 isoform X2 [Homo | XP_005251843.1 |
| 530391273 | AUH | methylglutaconyl-CoA hydratase, mitochondrial i | XP_005252123.1 |
| 530391650 | ARPC5L | actin-related protein 2/3 complex subunit 5-like p | XP_005252307.1 |
| 530392189 | PFKP | ATP-dependent 6-phosphofructokinase, platelet | XP_005252522.1 |
| 530393366 | HSPA12A | heat shock 70 kDa protein 12A isoform X1 [Hom | XP_005269729.1 |
| 530393496 | HK1 | hexokinase-1 isoform X3 [Homo sapiens] | XP_005269792.1 |
| 530394328 | VCL | vinculin isoform X1 [Homo sapiens] | XP_005270199.1 |
| 530396203 | ME3 | NADP-dependent malic enzyme, mitochondrial is | XP_005273774.1 |

| | | | | |
|---|---|---|---|---|
| 530396818 | NDUFS8 | | NADH dehydrogenase [ubiquinone] iron-sulfur pr | XP_005274070.1 |
| 530398165 | OPCML | | opioid-binding protein/cell adhesion molecule iso | XP_005271631.1 |
| 530398179 | NTM | | neurotrimin isoform X2 [Homo sapiens] | XP_005271637.1 |
| 530399089 | GABARAPL1 | | gamma-aminobutyric acid receptor-associated pı | XP_005253401.1 |
| 530400348 | ATP2B1 | | plasma membrane calcium-transporting ATPase | XP_005268976.1 |
| 530400466 | PFKM | | ATP-dependent 6-phosphofructokinase, muscle | XP_005269033.1 |
| 530400534 | KIF21A | | kinesin-like protein KIF21A isoform X1 [Homo sa | XP_005269064.1 |
| 530402792 | DCLK1 | | serine/threonine-protein kinase DCLK1 isoform X | XP_005266649.1 |
| 530405580 | DMXL2 | | dmX-like protein 2 isoform X1 [Homo sapiens] | XP_005254312.1 |
| 530406386 | TPM1 | | tropomyosin alpha-1 chain isoform X15 [Homo sa | XP_005254694.1 |
| 530406406 | TPM1 | | tropomyosin alpha-1 chain isoform X8 [Homo sa | XP_005254704.1 |
| 530409932 | TOM1L2 | | TOM1-like protein 2 isoform X1 [Homo sapiens] | XP_005256518.1 |
| 530410491 | PRPSAP2 | | phosphoribosyl pyrophosphate synthase-associa | XP_005256782.1 |
| 530410596 | VAMP2 | | vesicle-associated membrane protein 2 isoform | XP_005256832.1 |
| 530410945 | AP2B1 | | AP-2 complex subunit beta isoform X1 [Homo sa | XP_005257994.1 |
| 530410969 | ALDOC | | fructose-bisphosphate aldolase C isoform X1 [H | XP_005258006.1 |
| 530411491 | CLTC | | clathrin heavy chain 1 isoform X1 [Homo sapien | XP_005257069.1 |
| 530412219 | MAPT | | microtubule-associated protein tau isoform X3 [H | XP_005257419.1 |
| 530412417 | ATP6V0A1 | | V-type proton ATPase 116 kDa subunit a isoforr | XP_005257516.1 |
| 530415146 | UBA52 | | ubiquitin-60S ribosomal protein L40 isoform X2 [ | XP_005260108.1 |
| 530417532 | PDCD5 | | programmed cell death protein 5 isoform X1 [Hor | XP_005259449.1 |
| 530418431 | DYNLRB1 | | dynein light chain roadblock-type 1 isoform X1 [ | XP_005260625.1 |
| 530418825 | GART | | trifunctional purine biosynthetic protein adenosir | XP_005260998.1 |
| 530423397 | TPP2 | | tripeptidyl-peptidase 2 isoform X1 [Homo sapien | XP_005254127.1 |
| 530425685 | ABHD12 | | monoacylglycerol lipase ABHD12 isoform X1 [Ho | XP_005260755.1 |
| 530425723 | IDH3B | | isocitrate dehydrogenase [NAD] subunit beta, m | XP_005260773.1 |
| 530427319 | CTBP1 | | C-terminal-binding protein 1 isoform X1 [Homo sa | XP_005272318.1 |
| 538917681 | IDH1 | | isocitrate dehydrogenase [NADP] cytoplasmic [H | NP_001269316.1 |
| 543871463 | OLFM1 | | noelin isoform 4 precursor [Homo sapiens] | NP_001269540.1 |
| 544063423 | STIP1 | | stress-induced-phosphoprotein 1 isoform a [Hon | NP_001269581.1 |
| 544186032 | PDIA6 | | protein disulfide-isomerase A6 isoform a [Homo s | NP_001269633.1 |
| 545688329 | NAPB | | beta-soluble NSF attachment protein isoform a [ | NP_001269947.1 |
| 546232203 | ADAP1 | | arf-GAP with dual PH domain-containing protein | NP_001271237.1 |
| 557357733 | HSPH1 | | heat shock protein 105 kDa isoform 3 [Homo sa | NP_001273433.1 |
| 574584816 | TUBB4A | | tubulin beta-4A chain isoform 3 [Homo sapiens] | NP_001276058.1 |
| 576583524 | GAPDH | | glyceraldehyde-3-phosphate dehydrogenase isof | NP_001276675.1 |
| 578799206 | SH3GLB1 | | endophilin-B1 isoform X1 [Homo sapiens] | XP_006710735.1 |
| 578799563 | SARS | | serine--tRNA ligase, cytoplasmic isoform X1 [Ho | XP_006710876.1 |
| 578799914 | SGIP1 | | SH3-containing GRB2-like protein 3-interacting p | XP_006711024.1 |
| 578803186 | FAM49A | | protein FAM49A isoform X1 [Homo sapiens] | XP_006712171.1 |
| 578804380 | | 2-Sep | septin-2 isoform X2 [Homo sapiens] | XP_006712610.1 |
| 578804626 | SLC4A10 | | sodium-driven chloride bicarbonate exchanger is | XP_006712706.1 |
| 578808960 | TRIM2 | | tripartite motif-containing protein 2 isoform X1 [H | XP_006714220.1 |
| 578810794 | SYNPO | | synaptopodin isoform X3 [Homo sapiens] | XP_006714818.1 |
| 578811231 | SNCB | | beta-synuclein isoform X2 [Homo sapiens] | XP_006714977.1 |
| 578813410 | AMPH | | amphiphysin isoform X1 [Homo sapiens] | XP_006715752.1 |
| 578813710 | | 7-Sep | septin-7 isoform X6 [Homo sapiens] | XP_006715869.1 |
| 578814598 | CACNA2D1 | | voltage-dependent calcium channel subunit alph | XP_006716181.1 |
| 578816062 | OXR1 | | oxidation resistance protein 1 isoform X1 [Homo | XP_006716657.1 |
| 578817145 | DNM1 | | dynamin-1 isoform X3 [Homo sapiens] | XP_006717055.1 |
| 578817664 | SH3GLB2 | | endophilin-B2 isoform X1 [Homo sapiens] | XP_006717251.1 |
| 578817797 | SPTAN1 | | spectrin alpha chain, non-erythrocytic 1 isoform | XP_006717308.1 |
| 578818230 | CELF2 | | CUGBP Elav-like family member 2 isoform X2 [H | XP_006717432.1 |
| 578818565 | VIM | | vimentin isoform X1 [Homo sapiens] | XP_006717563.1 |
| 578821687 | SPTBN2 | | spectrin beta chain, non-erythrocytic 2 isoform | XP_006718732.1 |
| 578822042 | HEPACAM | | hepatocyte cell adhesion molecule isoform X1 [H | XP_006718849.1 |
| 578825895 | GNG2 | | guanine nucleotide-binding protein G(I)/G(S)/G(C | XP_006720236.1 |
| 578826688 | SCAMP5 | | secretory carrier-associated membrane protein 5 | XP_006720483.1 |
| 578828264 | ROGDI | | protein rogdi homolog isoform X1 [Homo sapiens | XP_006721010.1 |
| 578832288 | MAPRE2 | | microtubule-associated protein RP/EB family me | XP_006722438.1 |
| 578832421 | WDR7 | | WD repeat-containing protein 7 isoform X1 [Hom | XP_006722494.1 |
| 578835125 | CYTH2 | | cytohesin-2 isoform X1 [Homo sapiens] | XP_006723535.1 |

| | | | |
|---|---|---|---|
| 578836565 | PFKL | ATP-dependent 6-phosphofructokinase, liver typ | XP_006724074.1 |
| 604723356 | NDUFA5 | NADH dehydrogenase [ubiquinone] 1 alpha subc | NP_001278233.1 |
| 662033922 | PEA15 | astrocytic phosphoprotein PEA-15 isoform a [Hc | NP_001284505.1 |
| 664805999 | EIF4B | eukaryotic translation initiation factor 4B isoforn | NP_001287750.1 |
| 729042247 | AIP | AH receptor-interacting protein isoform 1 [Homo | NP_003968.3 |
| 735997436 | ACAT2 | acetyl-CoA acetyltransferase, cytosolic isoform | NP_001290182.1 |
| 740086846 | ACLY | ATP-citrate synthase isoform 3 [Homo sapiens] | NP_001290203.1 |
| 740087210 | MYL12A | myosin regulatory light chain 12A isoform 2 [Hor | NP_001289978.1 |
| 747165389 | RPSA | 40S ribosomal protein SA isoform 2 [Homo sapie | NP_001291217.1 |
| 748585205 | PRKACA | cAMP-dependent protein kinase catalytic subuni | NP_001291278.1 |
| 751247014 | TIMM9 | mitochondrial import inner membrane translocas | NP_001291416.1 |
| 767903710 | PHGDH | D-3-phosphoglycerate dehydrogenase isoform X | XP_011539528.1 |
| 767906235 | CAPZB | F-actin-capping protein subunit beta isoform X1 | XP_011540530.1 |
| 767908737 | NFASC | neurofascin isoform X8 [Homo sapiens] | XP_011507620.1 |
| 767910140 | BCAN | brevican core protein isoform X1 [Homo sapiens | XP_011508168.1 |
| 767911065 | UBAP2L | ubiquitin-associated protein 2-like isoform X2 [H | XP_011508514.1 |
| 767913715 | ADD2 | beta-adducin isoform X1 [Homo sapiens] | XP_011530804.1 |
| 767913845 | CTNNA2 | catenin alpha-2 isoform X1 [Homo sapiens] | XP_011530858.1 |
| 767915105 | SLC8A1 | sodium/calcium exchanger 1 isoform X1 [Homo s | XP_011531358.1 |
| 767917469 | NGEF | ephexin-1 isoform X1 [Homo sapiens] | XP_011509225.1 |
| 767918167 | MAP2 | microtubule-associated protein 2 isoform X6 [Ho | XP_011509492.1 |
| 767922705 | CLASP2 | CLIP-associating protein 2 isoform X1 [Homo sa | XP_011531807.1 |
| 767922915 | ERC2 | ERC protein 2 isoform X1 [Homo sapiens] | XP_011531882.1 |
| 767924452 | CADPS | calcium-dependent secretion activator 1 isoform | XP_011532481.1 |
| 767924768 | IQSEC1 | IQ motif and SEC7 domain-containing protein 1 i | XP_011532606.1 |
| 767926887 | PEX5L | PEX5-related protein isoform X3 [Homo sapiens] | XP_011511184.1 |
| 767930167 | ATP8A1 | phospholipid-transporting ATPase IA isoform X1 | XP_011511917.1 |
| 767931770 | ANK2 | ankyrin-2 isoform X5 [Homo sapiens] | XP_011530193.1 |
| 767932537 | SNCA | alpha-synuclein isoform X2 [Homo sapiens] | XP_011530509.1 |
| 767933143 | G3BP2 | ras GTPase-activating protein-binding protein 2 i | XP_011530742.1 |
| 767935355 | HAPLN1 | hyaluronan and proteoglycan link protein 1 isofo | XP_011541470.1 |
| 767937841 | UBE2D2 | ubiquitin-conjugating enzyme E2 D2 isoform X1 | | XP_011535981.1 |
| 767938112 | DBN1 | drebrin isoform X1 [Homo sapiens] | XP_011532748.1 |
| 767938356 | HNRNPH1 | heterogeneous nuclear ribonucleoprotein H isofc | XP_011532847.1 |
| 767938662 | CANX | calnexin isoform X2 [Homo sapiens] | XP_011532966.1 |
| 767939866 | PACSIN1 | protein kinase C and casein kinase substrate in | XP_011512843.1 |
| 767943839 | WASF1 | wiskott-Aldrich syndrome protein family member | XP_011534535.1 |
| 767943934 | SNAP91 | clathrin coat assembly protein AP180 isoform X | XP_011534567.1 |
| 767945521 | OGDH | 2-oxoglutarate dehydrogenase, mitochondrial isc | XP_011513710.1 |
| 767945559 | MPP6 | MAGUK p55 subfamily member 6 isoform X1 [Hc | XP_011513727.1 |
| 767945681 | FIGNL1 | fidgetin-like protein 1 isoform X2 [Homo sapiens] | XP_011513772.1 |
| 767945839 | TTYH3 | protein tweety homolog 3 isoform X1 [Homo sapi | XP_011513837.1 |
| 767945863 | CAMK2B | calcium/calmodulin-dependent protein kinase typ | XP_011513849.1 |
| 767945997 | EIF3B | eukaryotic translation initiation factor 3 subunit | XP_011513901.1 |
| 767947915 | NRCAM | neuronal cell adhesion molecule isoform X2 [Hor | XP_011514557.1 |
| 767948171 | CHCHD3 | MICOS complex subunit MIC19 isoform X1 [Hom | XP_011514665.1 |
| 767948685 | EIF4H | eukaryotic translation initiation factor 4H isoforn | XP_011514858.1 |
| 767950720 | PSD3 | PH and SEC7 domain-containing protein 3 isofor | XP_011542764.1 |
| 767953162 | FAM49B | protein FAM49B isoform X2 [Homo sapiens] | XP_011515409.1 |
| 767953706 | NCALD | neurocalcin-delta isoform X1 [Homo sapiens] | XP_011515637.1 |
| 767954754 | KIAA1045 | protein KIAA1045 isoform X1 [Homo sapiens] | XP_011516129.1 |
| 767955207 | SH3GL2 | endophilin-A1 isoform X1 [Homo sapiens] | XP_011516307.1 |
| 767956550 | ABCA1 | ATP-binding cassette sub-family A member 1 isc | XP_011516642.1 |
| 767958030 | SET | protein SET isoform X1 [Homo sapiens] | XP_011517213.1 |
| 767958792 | GDA | guanine deaminase isoform X2 [Homo sapiens] | XP_011517524.1 |
| 767961213 | MICU1 | calcium uptake protein 1, mitochondrial isoform | XP_011537421.1 |
| 767963194 | PPP3CB | serine/threonine-protein phosphatase 2B catalyt | XP_011538223.1 |
| 767963256 | OGDHL | 2-oxoglutarate dehydrogenase-like, mitochondria | XP_011538249.1 |
| 767966201 | PSMC3 | 26S protease regulatory subunit 6A isoform X2 [ | XP_011518535.1 |
| 767966323 | SLC1A2 | excitatory amino acid transporter 2 isoform X1 [ | XP_011518586.1 |
| 767967537 | DLG2 | disks large homolog 2 isoform X1 [Homo sapiens | XP_011543080.1 |
| 767967976 | PGM2L1 | glucose 1,6-bisphosphate synthase isoform X1 | | XP_011543255.1 |

| | | | | |
|---|---|---|---|---|
| 767968186 | ARRB1 | | beta-arrestin-1 isoform X1 [Homo sapiens] | XP_011543336.1 |
| 767969491 | HYOU1 | | hypoxia up-regulated protein 1 isoform X1 [Homo | XP_011540859.1 |
| 767970091 | HSPA8 | | heat shock cognate 71 kDa protein isoform X1 [ | XP_011541100.1 |
| 767971402 | PTMS | | parathymosin isoform X1 [Homo sapiens] | XP_011519289.1 |
| 767973049 | OSBPL8 | | oxysterol-binding protein-related protein 8 isofor | XP_011536158.1 |
| 767973233 | CNTN1 | | contactin-1 isoform X1 [Homo sapiens] | XP_011536228.1 |
| 767974361 | NAP1L1 | | nucleosome assembly protein 1-like 1 isoform X | XP_011536692.1 |
| 767974682 | PPP1CC | | serine/threonine-protein phosphatase PP1-gamm | XP_011536806.1 |
| 767980422 | HNRNPC | | heterogeneous nuclear ribonucleoproteins C1/C2 | XP_011535010.1 |
| 767981118 | NDRG2 | | protein NDRG2 isoform X1 [Homo sapiens] | XP_011535300.1 |
| 767981762 | ACTN1 | | alpha-actinin-1 isoform X1 [Homo sapiens] | XP_011535567.1 |
| 767983063 | CYFIP1 | | cytoplasmic FMR1-interacting protein 1 isoform X | XP_011542175.1 |
| 767983219 | CKMT1B | | creatine kinase U-type, mitochondrial isoform X1 | XP_011519499.1 |
| 767984251 | MYO5A | | unconventional myosin-Va isoform X1 [Homo sa | XP_011519908.1 |
| 767985907 | SLC12A6 | | solute carrier family 12 member 6 isoform X1 [Ho | XP_011520569.1 |
| 767986980 | ABAT | | 4-aminobutyrate aminotransferase, mitochondria | XP_011520702.1 |
| 767987102 | CARHSP1 | | calcium-regulated heat stable protein 1 isoform X | XP_011520746.1 |
| 767987604 | UBE2I | | SUMO-conjugating enzyme UBC9 isoform X1 [Ho | XP_011520947.1 |
| 767988216 | CORO1A | | coronin-1A isoform X1 [Homo sapiens] | XP_011544016.1 |
| 767990939 | KIAA0513 | | uncharacterized protein KIAA0513 isoform X1 [H | XP_011521785.1 |
| 767991742 | DLG4 | | disks large homolog 4 isoform X1 [Homo sapiens | XP_011522000.1 |
| 767992184 | MYH10 | | myosin-10 isoform X2 [Homo sapiens] | XP_011522177.1 |
| 767993234 | BAIAP2 | | brain-specific angiogenesis inhibitor 1-associate | XP_011522496.1 |
| 767993263 | | 9-Sep | septin-9 isoform X3 [Homo sapiens] | XP_011522506.1 |
| 767994242 | TANC2 | | protein TANC2 isoform X2 [Homo sapiens] | XP_011522899.1 |
| 767994460 | GIT1 | | ARF GTPase-activating protein GIT1 isoform X1 | XP_011522986.1 |
| 767994920 | NSF | | vesicle-fusing ATPase isoform X2 [Homo sapien | XP_011523165.1 |
| 767995236 | PRKAR1A | | cAMP-dependent protein kinase type I-alpha reg | XP_011523286.1 |
| 767996099 | PIP4K2B | | phosphatidylinositol 5-phosphate 4-kinase type- | XP_011523628.1 |
| 767997328 | FASN | | fatty acid synthase isoform X1 [Homo sapiens] | XP_011521840.1 |
| 767997413 | ARHGDIA | | rho GDP-dissociation inhibitor 1 isoform X1 [Hon | XP_011521876.1 |
| 767997588 | EPB41L3 | | band 4.1-like protein 3 isoform X9 [Homo sapien | XP_011523918.1 |
| 767997921 | NAPG | | gamma-soluble NSF attachment protein isoform | XP_011524057.1 |
| 767999054 | KIAA1468 | | lisH domain and HEAT repeat-containing protein | XP_011524412.1 |
| 767999390 | HDHD2 | | haloacid dehalogenase-like hydrolase domain-co | XP_011524529.1 |
| 768001626 | PIP5K1C | | phosphatidylinositol 4-phosphate 5-kinase type- | XP_011526147.1 |
| 768003717 | ICAM5 | | intercellular adhesion molecule 5 isoform X1 [Ho | XP_011526531.1 |
| 768008086 | GPI | | glucose-6-phosphate isomerase isoform X2 [Hor | XP_011525056.1 |
| 768009376 | NOVA2 | | RNA-binding protein Nova-2 isoform X1 [Homo s | XP_011525296.1 |
| 768014078 | NSFL1C | | NSFL1 cofactor p47 isoform X1 [Homo sapiens] | XP_011527602.1 |
| 768016461 | EPB41L1 | | band 4.1-like protein 1 isoform X4 [Homo sapien | XP_011526969.1 |
| 768017580 | ARFGAP1 | | ADP-ribosylation factor GTPase-activating prote | XP_011527203.1 |
| 768018500 | MAP1LC3A | | microtubule-associated proteins 1A/1B light chai | XP_011527385.1 |
| 768020808 | NCAM2 | | neural cell adhesion molecule 2 isoform X1 [Hon | XP_011527878.1 |
| 768022887 | SLC25A18 | | mitochondrial glutamate carrier 2 isoform X2 [Ho | XP_011544451.1 |
| 768032495 | PDHA1 | | pyruvate dehydrogenase E1 component subunit | XP_011543833.1 |
| 768033352 | UBA1 | | ubiquitin-like modifier-activating enzyme 1 isofor | XP_011542255.1 |
| 768037765 | MSN | | moesin isoform X1 [Homo sapiens] | XP_011529261.1 |
| 768039038 | | 6-Sep | septin-6 isoform X2 [Homo sapiens] | XP_011529619.1 |
| 768039088 | HPRT1 | | hypoxanthine-guanine phosphoribosyltransferas | XP_011529630.1 |
| 779176942 | CALM2 | | calmodulin isoform 1 [Homo sapiens] | NP_001292553.1 |
| 807066321 | | 11-Sep | septin-11 isoform 1 [Homo sapiens] | NP_001293076.1 |
| 815891093 | RPS15 | | 40S ribosomal protein S15 isoform 1 [Homo sapi | NP_001295155.1 |
| 909618061 | AP2M1 | | AP-2 complex subunit mu isoform c [Homo sapie | NP_001298127.1 |
| 927669104 | RHOA | | transforming protein RhoA isoform 1 precursor [I | NP_001300870.1 |
| 937553895 | PVALB | | parvalbumin alpha [Homo sapiens] | NP_001302461.1 |
| 938403405 | PRKCG | | protein kinase C gamma type isoform 1 [Homo s | NP_001303258.1 |
| 939646432 | MDH1 | | malate dehydrogenase, cytoplasmic isoform MD | NP_001303303.1 |

| Exp. q-value | Sum PEP Score | Coverage | # Peptides | # PSMs | # Unique Peptides | # Protein Groups | # AAs | MW [kDa] |
|---|---|---|---|---|---|---|---|---|
| 0 | 4.16071054 | 3.02325581 | 1 | 2 | 1 | 1 | 430 | 47.625 |
| 0 | 57.1100315 | 18.4615385 | 13 | 39 | 13 | 1 | 780 | 85.372 |
| 0 | 93.0550913 | 34.7480106 | 19 | 86 | 3 | 1 | 377 | 42.024 |
| 0 | 3.54363854 | 3.36538462 | 2 | 4 | 2 | 1 | 832 | 91.867 |
| 0.00281162 | 1.83179725 | 9.49367089 | 1 | 2 | 1 | 1 | 316 | 35.83 |
| 0 | 2.90938929 | 4.6875 | 1 | 2 | 1 | 1 | 320 | 35.914 |
| 0 | 10.6193408 | 27.6243094 | 3 | 9 | 3 | 1 | 181 | 20.684 |
| 0 | 3.81163407 | 5.55555556 | 1 | 2 | 1 | 1 | 180 | 20.517 |
| 0 | 116.125673 | 26.7647059 | 23 | 72 | 12 | 1 | 1020 | 112.193 |
| 0 | 37.1405714 | 24.7524752 | 11 | 33 | 11 | 1 | 303 | 35.039 |
| 0 | 4.19185657 | 5.16431925 | 1 | 2 | 1 | 1 | 213 | 23.263 |
| 0 | 12.1337334 | 9.68586387 | 4 | 8 | 4 | 1 | 382 | 43.914 |
| 0 | 40.8916104 | 41.1504425 | 11 | 27 | 11 | 1 | 226 | 26.129 |
| 0.00281162 | 1.83416238 | 2.25988701 | 1 | 2 | 1 | 1 | 531 | 57.988 |
| 0 | 11.4905656 | 4.33436533 | 3 | 6 | 3 | 1 | 323 | 35.191 |
| 0.00339716 | 1.78595132 | 14.2857143 | 1 | 2 | 1 | 1 | 63 | 7.241 |
| 0 | 4.66978922 | 2.73109244 | 1 | 2 | 1 | 1 | 476 | 53.117 |
| 0.01680912 | 1.18876023 | 3.36538462 | 1 | 2 | 1 | 1 | 208 | 22.478 |
| 0 | 3.64092377 | 3.18471338 | 1 | 2 | 1 | 1 | 314 | 33.754 |
| 0 | 12.3634771 | 14.8337596 | 3 | 8 | 3 | 1 | 391 | 45.115 |
| 0 | 106.3973 | 50.6993007 | 21 | 68 | 1 | 1 | 572 | 62.255 |
| 0 | 29.4191576 | 20.7017544 | 9 | 27 | 4 | 1 | 570 | 61.924 |
| 0 | 38.6456081 | 21.6450216 | 10 | 26 | 3 | 1 | 462 | 50.109 |
| 0 | 33.3059209 | 26.349892 | 10 | 26 | 3 | 1 | 463 | 50.438 |
| 0.00711955 | 1.49281902 | 4 | 1 | 2 | 1 | 1 | 225 | 24.748 |
| 0 | 7.81805159 | 4.57665904 | 2 | 6 | 2 | 1 | 437 | 50.087 |
| 0 | 14.2124391 | 8.50815851 | 4 | 8 | 4 | 1 | 858 | 95.277 |
| 0 | 17.3341994 | 16.0098522 | 5 | 12 | 5 | 1 | 406 | 46.125 |
| 0 | 58.1407766 | 35.9447005 | 16 | 46 | 15 | 1 | 434 | 47.139 |
| 0 | 11.1436851 | 16.2162162 | 3 | 6 | 3 | 1 | 333 | 35.058 |
| 0.00161812 | 2.1177032 | 4.61538462 | 1 | 2 | 1 | 1 | 585 | 65.368 |
| 0 | 58.7030694 | 33.7807606 | 12 | 37 | 5 | 1 | 447 | 50.55 |
| 0 | 13.8519969 | 12.6760563 | 3 | 12 | 2 | 1 | 355 | 40.425 |
| 0 | 20.2366888 | 18.1598063 | 6 | 14 | 6 | 1 | 413 | 46.219 |
| 0 | 3.46042212 | 4.60829493 | 1 | 2 | 1 | 1 | 217 | 25.19 |
| 0 | 20.8018427 | 50.7692308 | 5 | 14 | 4 | 1 | 130 | 14.087 |
| 0 | 6.30338737 | 28.125 | 2 | 4 | 1 | 1 | 128 | 13.545 |
| 0 | 29.0695546 | 52.4271845 | 8 | 20 | 8 | 1 | 103 | 11.36 |
| 0 | 16.4467393 | 25.3521127 | 4 | 12 | 4 | 1 | 142 | 15.248 |
| 0 | 3.58566497 | 5.28967254 | 2 | 4 | 2 | 1 | 397 | 44.839 |
| 0 | 17.0375151 | 41.1764706 | 6 | 14 | 6 | 1 | 102 | 10.925 |
| 0 | 2.85573723 | 9.56521739 | 1 | 1 | 1 | 1 | 115 | 12.468 |
| 0 | 4.60049934 | 12.345679 | 1 | 2 | 1 | 1 | 81 | 9.364 |
| 0 | 4.40197593 | 5.71428571 | 1 | 2 | 1 | 1 | 175 | 20.095 |
| 0 | 14.3717266 | 41.2698413 | 5 | 12 | 1 | 1 | 189 | 21.216 |
| 0 | 3.34843126 | 1.11524164 | 1 | 2 | 1 | 1 | 807 | 89.365 |
| 0.00161812 | 2.14020145 | 3.930131 | 1 | 2 | 1 | 1 | 229 | 25.553 |
| 0 | 7.82887512 | 13.9037433 | 2 | 6 | 2 | 1 | 187 | 21.044 |
| 0 | 69.6760795 | 55.1181102 | 11 | 33 | 11 | 1 | 254 | 28.786 |
| 0 | 54.6677848 | 39.088729 | 14 | 49 | 14 | 1 | 417 | 44.586 |
| 0 | 9.24003014 | 5 | 1 | 2 | 1 | 1 | 360 | 41.539 |
| 0.00067889 | 2.67840157 | 2.26537217 | 1 | 2 | 1 | 1 | 309 | 35.571 |
| 0 | 10.0690248 | 44.7058824 | 3 | 7 | 3 | 1 | 170 | 19.288 |
| 0.00945347 | 1.36632959 | 8.97435897 | 1 | 1 | 1 | 1 | 234 | 25.882 |
| 0.00192431 | 1.98380265 | 5.49019608 | 1 | 2 | 1 | 1 | 255 | 28.415 |
| 0 | 14.3127241 | 11.3163972 | 3 | 6 | 3 | 1 | 433 | 48.603 |
| 0 | 7.07628981 | 2.85087719 | 1 | 2 | 1 | 1 | 456 | 52.871 |

| | | | | | | | | |
|---|---|---|---|---|---|---|---|---|
| 0 | 22.5114806 | 25.9433962 | 5 | 16 | 5 | 1 | 212 | 23.531 |
| 0 | 52.6644182 | 48.1818182 | 11 | 36 | 6 | 1 | 220 | 24.968 |
| 0 | 9.7973377 | 11.627907 | 2 | 4 | 1 | 1 | 215 | 23.692 |
| 0.00538761 | 1.56511188 | 5.45454545 | 1 | 2 | 1 | 1 | 165 | 17.808 |
| 0 | 3.00309449 | 5.71428571 | 1 | 2 | 1 | 1 | 140 | 14.856 |
| 0 | 4.86143011 | 34.2857143 | 2 | 4 | 2 | 1 | 70 | 8.213 |
| 0.00161812 | 2.09598812 | 2.72952854 | 1 | 2 | 1 | 1 | 403 | 46.08 |
| 0.00133245 | 2.30856485 | 3.38345865 | 1 | 2 | 1 | 1 | 266 | 29.977 |
| 0 | 3.79533749 | 6.22568093 | 1 | 2 | 1 | 1 | 257 | 28.007 |
| 0 | 6.56180713 | 14.7826087 | 2 | 4 | 2 | 1 | 115 | 11.658 |
| 0 | 3.23807216 | 7.94701987 | 1 | 2 | 1 | 1 | 151 | 17.212 |
| 0 | 6.96817134 | 13.6986301 | 3 | 6 | 3 | 1 | 146 | 16.435 |
| 0.00067889 | 2.58037464 | 7.40740741 | 1 | 2 | 1 | 1 | 135 | 15.54 |
| 0 | 9.71420387 | 12.4137931 | 4 | 7 | 4 | 1 | 145 | 16.051 |
| 0.01086637 | 1.32762502 | 6.4 | 1 | 1 | 1 | 1 | 125 | 13.734 |
| 0 | 5.40079826 | 8.33333333 | 2 | 4 | 2 | 1 | 264 | 29.926 |
| 0 | 3.64888368 | 7.2243346 | 2 | 3 | 2 | 1 | 263 | 29.579 |
| 0.00067889 | 2.55098468 | 4.12371134 | 1 | 2 | 1 | 1 | 194 | 22.113 |
| 0 | 10.3249539 | 11.5384615 | 2 | 6 | 2 | 1 | 208 | 24.19 |
| 0 | 9.01375853 | 10.0543478 | 4 | 8 | 1 | 1 | 368 | 41.464 |
| 0 | 15.0625657 | 15.4158215 | 6 | 12 | 6 | 1 | 493 | 54.496 |
| 0.01058201 | 1.34852815 | 2.77227723 | 1 | 2 | 1 | 1 | 505 | 53.862 |
| 0 | 22.3768256 | 9.09090909 | 7 | 17 | 5 | 1 | 803 | 92.411 |
| 0 | 211.757173 | 57.9775281 | 29 | 239 | 2 | 1 | 445 | 49.875 |
| 0 | 13.0278991 | 25.6578947 | 3 | 6 | 3 | 1 | 152 | 17.127 |
| 0 | 43.4060069 | 39.9293286 | 11 | 36 | 10 | 1 | 283 | 30.754 |
| 0 | 5.83534978 | 3.40909091 | 1 | 4 | 1 | 1 | 528 | 59.106 |
| 0 | 40.3670784 | 39.8373984 | 9 | 26 | 4 | 1 | 246 | 28.065 |
| 0 | 38.431683 | 38.2113821 | 10 | 26 | 7 | 1 | 246 | 28.201 |
| 0 | 5.20999648 | 3.7470726 | 1 | 2 | 1 | 1 | 427 | 45.171 |
| 0 | 3.68047755 | 2.85714286 | 1 | 2 | 1 | 1 | 455 | 52.528 |
| 0 | 12.2378051 | 11.1538462 | 2 | 5 | 2 | 1 | 260 | 29.228 |
| 0 | 4.32495526 | 1.03420843 | 1 | 2 | 1 | 1 | 1257 | 139.915 |
| 0.00161812 | 2.17855198 | 3.79506641 | 1 | 2 | 1 | 1 | 527 | 59.644 |
| 0 | 8.35176714 | 4.23076923 | 3 | 6 | 3 | 1 | 520 | 56.122 |
| 0.00067889 | 2.46407326 | 6.04395604 | 1 | 2 | 1 | 1 | 182 | 20.443 |
| 0 | 3.5407161 | 7.43405276 | 2 | 4 | 2 | 1 | 417 | 48.112 |
| 0.01746971 | 1.07930287 | 7.77202073 | 1 | 2 | 1 | 1 | 193 | 20.554 |
| 0 | 4.16027062 | 17.5257732 | 1 | 2 | 1 | 1 | 97 | 10.991 |
| 0 | 2.73802381 | 3.80952381 | 1 | 2 | 1 | 1 | 315 | 36.089 |
| 0 | 5.76736624 | 22.556391 | 3 | 6 | 3 | 1 | 133 | 14.849 |
| 0 | 4.53415395 | 12.6760563 | 2 | 4 | 2 | 1 | 142 | 16.702 |
| 0 | 4.1919944 | 4.32569975 | 2 | 4 | 2 | 1 | 393 | 42.767 |
| 0 | 4.90069789 | 15.1785714 | 2 | 3 | 2 | 1 | 224 | 25.019 |
| 0 | 8.81312624 | 4.17972832 | 2 | 4 | 2 | 1 | 957 | 109.427 |
| 0.01680912 | 1.20460667 | 2.15982721 | 1 | 2 | 1 | 1 | 463 | 52.512 |
| 0 | 12.9174568 | 10.9848485 | 3 | 8 | 3 | 1 | 264 | 30.223 |
| 0.00067889 | 2.56591036 | 8.06451613 | 1 | 2 | 1 | 1 | 124 | 13.703 |
| 0.00133245 | 2.37263414 | 6.01851852 | 1 | 2 | 1 | 1 | 216 | 23.728 |
| 0 | 18.3689446 | 31.2195122 | 4 | 10 | 2 | 1 | 205 | 22.663 |
| 0.00339716 | 1.75945075 | 7.14285714 | 1 | 2 | 1 | 1 | 126 | 13.907 |
| 0 | 30.7407669 | 21.5277778 | 8 | 16 | 8 | 1 | 288 | 33.003 |
| 0 | 4.96110044 | 6.21468927 | 2 | 3 | 2 | 1 | 354 | 40.252 |
| 0 | 8.38502993 | 11.5226337 | 3 | 6 | 2 | 1 | 243 | 27.211 |
| 0.00281162 | 1.83150252 | 3.00429185 | 1 | 2 | 1 | 1 | 233 | 25.981 |
| 0 | 13.1903225 | 18.3908046 | 3 | 6 | 2 | 1 | 261 | 30.006 |
| 0.00067889 | 2.49066304 | 2.73972603 | 1 | 2 | 1 | 1 | 292 | 33.283 |
| 0 | 5.00130232 | 7.79467681 | 2 | 4 | 2 | 1 | 526 | 53.394 |
| 0 | 9.83209222 | 6.10412926 | 3 | 6 | 3 | 1 | 557 | 63.777 |
| 0.00161812 | 2.03105032 | 5.71428571 | 1 | 4 | 1 | 1 | 140 | 15.045 |
| 0 | 7.06690345 | 7.83783784 | 2 | 4 | 2 | 1 | 370 | 40.738 |
| 0 | 42.8694475 | 24.5519713 | 12 | 26 | 1 | 1 | 558 | 61.359 |

| | | | | | | | | |
|---|---|---|---|---|---|---|---|---|
| 0 | 10.1691881 | 11.3402062 | 2 | 8 | 2 | 1 | 194 | 20.85 |
| 0 | 24.3907033 | 17.1945701 | 6 | 12 | 6 | 1 | 221 | 22.336 |
| 0 | 4.85620547 | 5.15603799 | 3 | 6 | 3 | 1 | 737 | 83.62 |
| 0 | 20.3263001 | 14.3617021 | 5 | 12 | 5 | 1 | 376 | 42.587 |
| 0 | 11.2692473 | 18.2741117 | 3 | 10 | 3 | 1 | 394 | 44.732 |
| 0 | 20.8352442 | 18.8995215 | 6 | 14 | 5 | 1 | 418 | 47.341 |
| 0 | 6.06775176 | 7.66666667 | 2 | 6 | 2 | 1 | 300 | 34.311 |
| 0 | 14.9972378 | 27.7108434 | 5 | 11 | 3 | 1 | 166 | 18.491 |
| 0.00538761 | 1.56399646 | 1.94174757 | 1 | 2 | 1 | 1 | 412 | 45.717 |
| 0 | 19.7737185 | 23.2240437 | 7 | 15 | 7 | 1 | 366 | 39.566 |
| 0.00827668 | 1.3864753 | 6.2992126 | 1 | 2 | 1 | 1 | 127 | 14.469 |
| 0 | 22.6491967 | 26.3975155 | 5 | 12 | 5 | 1 | 322 | 34.889 |
| 0 | 12.5167243 | 7.95724466 | 5 | 13 | 2 | 1 | 842 | 97.031 |
| 0 | 73.2125182 | 36.7298578 | 19 | 51 | 19 | 1 | 422 | 47.543 |
| 0.00067889 | 2.64955814 | 3.69230769 | 1 | 2 | 1 | 1 | 325 | 36.55 |
| 0 | 5.90492362 | 12.9337539 | 3 | 5 | 3 | 1 | 317 | 35.055 |
| 0 | 5.441162 | 5.86666667 | 2 | 4 | 1 | 1 | 375 | 42.797 |
| 0 | 207.117608 | 64.0449438 | 31 | 239 | 2 | 1 | 445 | 49.799 |
| 0 | 21.2563054 | 24.0740741 | 6 | 12 | 6 | 1 | 216 | 24.408 |
| 0.00827668 | 1.40616034 | 8.07453416 | 1 | 2 | 1 | 1 | 161 | 18.479 |
| 0 | 8.0086853 | 3.14465409 | 1 | 2 | 1 | 1 | 477 | 52.791 |
| 0 | 5.42520796 | 9.79020979 | 2 | 6 | 2 | 1 | 286 | 32.929 |
| 0 | 17.3688766 | 11.7757009 | 4 | 8 | 4 | 1 | 535 | 57.452 |
| 0 | 6.44181121 | 4.97237569 | 2 | 4 | 2 | 1 | 543 | 59.329 |
| 0 | 4.85189917 | 4.92610837 | 2 | 3 | 2 | 1 | 406 | 44.792 |
| 0 | 15.1071173 | 18.7739464 | 4 | 8 | 4 | 1 | 261 | 29.698 |
| 0 | 3.19307422 | 4.81481481 | 1 | 2 | 1 | 1 | 270 | 31.786 |
| 0 | 9.27401523 | 3.20641283 | 1 | 4 | 1 | 1 | 499 | 56.842 |
| 0.00161812 | 2.0791465 | 16.3043478 | 1 | 2 | 1 | 1 | 92 | 10.706 |
| 0 | 13.5716863 | 10.9414758 | 5 | 10 | 5 | 1 | 393 | 43.411 |
| 0.00192431 | 2.01019435 | 10.4477612 | 1 | 2 | 1 | 1 | 134 | 15.02 |
| 0.01004135 | 1.35428306 | 7.69230769 | 1 | 2 | 1 | 1 | 195 | 21.658 |
| 0.01730006 | 1.13715334 | 3.3492823 | 1 | 2 | 1 | 1 | 418 | 47.337 |
| 0 | 5.93824396 | 13.3333333 | 2 | 3 | 1 | 1 | 165 | 18.493 |
| 0 | 73.86818 | 37.7880184 | 13 | 60 | 12 | 1 | 434 | 47.239 |
| 0.00223642 | 1.92738252 | 4.59770115 | 1 | 2 | 1 | 1 | 261 | 28.975 |
| 0.01680912 | 1.22337145 | 23.6842105 | 1 | 1 | 1 | 1 | 76 | 7.984 |
| 0 | 3.41250073 | 5.9347181 | 2 | 4 | 2 | 1 | 337 | 37.516 |
| 0 | 26.0286865 | 36.8627451 | 9 | 24 | 7 | 1 | 255 | 29.155 |
| 0 | 37.1979268 | 24.0816327 | 6 | 22 | 3 | 1 | 245 | 27.747 |
| 0 | 24.8066368 | 13.6986301 | 3 | 15 | 3 | 1 | 219 | 23.679 |
| 0 | 6.27531347 | 13.3064516 | 2 | 7 | 2 | 1 | 248 | 27.728 |
| 0.01746971 | 1.06956041 | 0.63463282 | 1 | 1 | 1 | 1 | 1103 | 121.997 |
| 0.00133245 | 2.33077613 | 15.942029 | 1 | 1 | 1 | 1 | 69 | 7.928 |
| 0 | 57.5347838 | 27.5434243 | 16 | 33 | 16 | 1 | 806 | 89.266 |
| 0 | 12.0341083 | 10.0806452 | 3 | 8 | 3 | 1 | 248 | 27.06 |
| 0 | 7.39706564 | 12.2270742 | 4 | 8 | 4 | 1 | 229 | 25.175 |
| 0 | 3.8462051 | 5.24861878 | 2 | 4 | 2 | 1 | 362 | 40.069 |
| 0 | 40.7880508 | 22.247191 | 9 | 29 | 2 | 1 | 445 | 50.631 |
| 0 | 36.2666127 | 23.0326296 | 11 | 29 | 6 | 1 | 521 | 58.65 |
| 0 | 9.75457563 | 16.8224299 | 3 | 6 | 3 | 1 | 214 | 22.012 |
| 0 | 2.84496777 | 4.73372781 | 1 | 2 | 1 | 1 | 338 | 38.25 |
| 0.00538761 | 1.55767704 | 2.80701754 | 1 | 2 | 1 | 1 | 285 | 31.102 |
| 0 | 6.25442372 | 3.51826793 | 2 | 4 | 2 | 1 | 739 | 83.302 |
| 0 | 3.14880282 | 5.2238806 | 1 | 2 | 1 | 1 | 268 | 29.98 |
| 0 | 5.06610796 | 5.41871921 | 1 | 2 | 1 | 1 | 203 | 23.562 |
| 0 | 3.56911905 | 3.01507538 | 1 | 2 | 1 | 1 | 398 | 44.36 |
| 0 | 6.47071033 | 10.3846154 | 2 | 4 | 2 | 1 | 260 | 27.356 |
| 0.00133245 | 2.28701377 | 1.9047619 | 1 | 2 | 1 | 1 | 630 | 70.766 |
| 0 | 8.54337966 | 14.2574257 | 3 | 5 | 3 | 1 | 505 | 55.175 |
| 0.02239642 | 1.02571864 | 4.47761194 | 1 | 2 | 1 | 1 | 201 | 23.368 |
| 0 | 7.04909991 | 3.92523364 | 1 | 2 | 1 | 1 | 535 | 60.849 |

| | | | | | | | | |
|---|---|---|---|---|---|---|---|---|
| 0.00538761 | 1.56240797 | 2.77777778 | 1 | 2 | 1 | 1 | 504 | 55.146 |
| 0 | 5.65286022 | 14.1304348 | 2 | 4 | 2 | 1 | 184 | 20.812 |
| 0.00339716 | 1.76371472 | 4.40528634 | 1 | 2 | 1 | 1 | 454 | 49.8 |
| 0 | 8.71776564 | 23.0366492 | 2 | 5 | 2 | 1 | 191 | 22.188 |
| 0.01086637 | 1.31749391 | 4.3062201 | 1 | 2 | 1 | 1 | 209 | 23.834 |
| 0 | 15.1211693 | 14.9122807 | 4 | 8 | 4 | 1 | 228 | 25.838 |
| 0 | 4.88706002 | 26.8656716 | 1 | 2 | 1 | 1 | 67 | 7.944 |
| 0 | 10.1976283 | 10.5263158 | 2 | 4 | 2 | 1 | 247 | 28.245 |
| 0 | 3.6716204 | 8.04597701 | 1 | 2 | 1 | 1 | 261 | 29.969 |
| 0.00133245 | 2.41465209 | 4 | 1 | 2 | 1 | 1 | 450 | 50.821 |
| 0 | 4.22709752 | 14.8148148 | 2 | 4 | 2 | 1 | 108 | 12.191 |
| 0 | 12.9628316 | 12.0462046 | 6 | 10 | 6 | 1 | 606 | 66.152 |
| 0 | 11.9364145 | 16.1137441 | 3 | 7 | 3 | 1 | 211 | 23.452 |
| 0.00339716 | 1.7652297 | 7.74647887 | 1 | 2 | 1 | 1 | 142 | 15.512 |
| 0 | 4.38541913 | 1.96428571 | 1 | 2 | 1 | 1 | 560 | 61.573 |
| 0 | 20.8362384 | 18.9701897 | 5 | 14 | 1 | 1 | 369 | 42.75 |
| 0.00067889 | 2.63578625 | 4.62962963 | 2 | 3 | 1 | 1 | 432 | 47.685 |
| 0 | 10.6545327 | 7.89473684 | 2 | 6 | 1 | 1 | 418 | 47.577 |
| 0 | 3.9722428 | 11.7241379 | 1 | 2 | 1 | 1 | 145 | 17.104 |
| 0.00133245 | 2.37520242 | 5.74324324 | 1 | 2 | 1 | 1 | 296 | 31.678 |
| 0.01746971 | 1.08767164 | 6.01092896 | 1 | 2 | 1 | 1 | 183 | 20.602 |
| 0 | 39.5120941 | 26.2711864 | 8 | 30 | 7 | 1 | 354 | 40.025 |
| 0 | 9.4941047 | 11.7437722 | 4 | 7 | 3 | 1 | 281 | 31.962 |
| 0 | 7.70092874 | 7.14285714 | 1 | 2 | 1 | 1 | 294 | 32.555 |
| 0 | 3.07571658 | 43.1818182 | 2 | 3 | 2 | 1 | 44 | 5.023 |
| 0 | 43.5881095 | 39.3939394 | 8 | 43 | 8 | 1 | 165 | 18.001 |
| 0.00161812 | 2.10651565 | 1.28865979 | 1 | 2 | 1 | 1 | 776 | 83.566 |
| 0 | 6.40886928 | 8.30449827 | 3 | 6 | 2 | 1 | 289 | 32.639 |
| 0 | 3.68634365 | 4.16666667 | 1 | 2 | 1 | 1 | 432 | 46.991 |
| 0 | 3.71346764 | 54.5454545 | 2 | 3 | 2 | 1 | 44 | 5.05 |
| 0 | 3.22344409 | 7.61904762 | 1 | 4 | 1 | 1 | 105 | 11.741 |
| 0 | 3.93330145 | 3.83386581 | 1 | 4 | 1 | 1 | 626 | 69.025 |
| 0 | 9.27415563 | 7.99136069 | 4 | 8 | 4 | 1 | 463 | 50.285 |
| 0 | 6.33162556 | 3.48162476 | 1 | 4 | 1 | 1 | 517 | 56.104 |
| 0 | 2.91399629 | 7.23684211 | 1 | 2 | 1 | 1 | 152 | 17.708 |
| 0 | 21.0773041 | 8.01317234 | 7 | 16 | 2 | 1 | 911 | 104.788 |
| 0 | 26.1263181 | 7.97697368 | 7 | 14 | 7 | 1 | 1216 | 138.48 |
| 0 | 4.88605665 | 9.09090909 | 1 | 2 | 1 | 1 | 154 | 16.638 |
| 0 | 4.78251606 | 3.3203125 | 1 | 4 | 1 | 1 | 512 | 59.112 |
| 0 | 3.56431486 | 0.78247261 | 1 | 2 | 1 | 1 | 1278 | 141.607 |
| 0.02239642 | 1.03282658 | 3.58974359 | 1 | 1 | 1 | 1 | 195 | 19.517 |
| 0 | 4.42945706 | 8.23970037 | 1 | 2 | 1 | 1 | 267 | 30.204 |
| 0 | 14.4416349 | 19.4029851 | 3 | 8 | 1 | 1 | 201 | 22.157 |
| 0 | 48.3399659 | 17.370892 | 11 | 27 | 1 | 1 | 639 | 69.978 |
| 0 | 3.84588047 | 20.4301075 | 1 | 2 | 1 | 1 | 93 | 10.431 |
| 0.00067889 | 2.48798303 | 4.11311054 | 1 | 2 | 1 | 1 | 389 | 43.154 |
| 0 | 5.42666416 | 4.30107527 | 1 | 2 | 1 | 1 | 372 | 38.723 |
| 0 | 14.1857179 | 12.7478754 | 4 | 10 | 4 | 1 | 353 | 37.407 |
| 0 | 8.02188511 | 6.1971831 | 2 | 6 | 2 | 1 | 355 | 38.41 |
| 0 | 3.8471004 | 1.50684932 | 1 | 2 | 1 | 1 | 730 | 77.464 |
| 0 | 6.32979466 | 12.0218579 | 2 | 5 | 2 | 1 | 366 | 38.627 |
| 0 | 20.4570333 | 11.4224138 | 4 | 10 | 4 | 1 | 464 | 50.996 |
| 0 | 87.9170049 | 28.4753363 | 15 | 90 | 3 | 1 | 446 | 49.825 |
| 0 | 45.5628023 | 24.4488978 | 14 | 34 | 11 | 1 | 499 | 55.357 |
| 0.00067889 | 2.53298418 | 6.06060606 | 1 | 2 | 1 | 1 | 132 | 14.505 |
| 0 | 3.24252805 | 4.04040404 | 1 | 2 | 1 | 1 | 297 | 34.341 |
| 0 | 7.615645 | 18.0722892 | 3 | 5 | 1 | 1 | 166 | 18.725 |
| 0.00339716 | 1.75621808 | 13.0434783 | 1 | 2 | 1 | 1 | 115 | 13.007 |
| 0 | 3.66314018 | 3.75426621 | 1 | 2 | 1 | 1 | 293 | 31.305 |
| 0 | 3.7749493 | 7.10382514 | 1 | 2 | 1 | 1 | 183 | 20.798 |
| 0.00133245 | 2.38174266 | 3.52564103 | 1 | 2 | 1 | 1 | 312 | 33.22 |
| 0.00770142 | 1.46394684 | 5.99078341 | 1 | 2 | 1 | 1 | 217 | 24.816 |

| | | | | | | | | |
|---|---|---|---|---|---|---|---|---|
| 0 | 7.01322361 | 12.9213483 | 2 | 4 | 2 | 1 | 178 | 20.24 |
| 0 | 8.50447776 | 20.5882353 | 3 | 6 | 3 | 1 | 204 | 24.131 |
| 0 | 6.98743836 | 10.9004739 | 3 | 5 | 3 | 1 | 211 | 24.247 |
| 0 | 10.251725 | 13.7096774 | 3 | 6 | 3 | 1 | 248 | 29.207 |
| 0 | 3.87484417 | 4.72972973 | 1 | 2 | 1 | 1 | 296 | 33.307 |
| 0 | 6.89171228 | 16.7539267 | 2 | 12 | 2 | 1 | 191 | 21.297 |
| 0 | 47.4948027 | 26.3888889 | 9 | 22 | 9 | 1 | 288 | 33.224 |
| 0 | 69.0763105 | 26.2996942 | 16 | 44 | 14 | 1 | 654 | 72.288 |
| 0 | 8.03354709 | 6.79156909 | 2 | 6 | 2 | 1 | 427 | 47.667 |
| 0 | 8.67270654 | 17.1428571 | 3 | 6 | 3 | 1 | 140 | 15.036 |
| 0 | 5.56082526 | 3.78548896 | 1 | 4 | 1 | 1 | 317 | 34.252 |
| 0.00067889 | 2.53253989 | 8.33333333 | 1 | 2 | 1 | 1 | 156 | 17.684 |
| 0 | 8.57521632 | 25 | 2 | 3 | 2 | 1 | 108 | 11.943 |
| 0 | 5.62287596 | 14.084507 | 1 | 2 | 1 | 1 | 142 | 15.639 |
| 0 | 3.75547549 | 4.01606426 | 1 | 2 | 1 | 1 | 249 | 28.663 |
| 0 | 122.461818 | 45.0892857 | 22 | 115 | 9 | 1 | 448 | 49.892 |
| 0 | 12.6109814 | 29.1390728 | 3 | 6 | 3 | 1 | 151 | 16.919 |
| 0 | 132.670177 | 51.4412417 | 24 | 152 | 11 | 1 | 451 | 50.104 |
| 0 | 3.38526884 | 26.6055046 | 1 | 3 | 1 | 1 | 109 | 12.147 |
| 0.01746971 | 1.12142076 | 1.38190955 | 1 | 2 | 1 | 1 | 796 | 91.649 |
| 0 | 3.65344229 | 1.25786164 | 2 | 4 | 2 | 1 | 1431 | 149.722 |
| 0 | 7.5915768 | 13.4831461 | 2 | 6 | 2 | 1 | 89 | 10.343 |
| 0.00538761 | 1.6039751 | 4.375 | 1 | 2 | 1 | 1 | 160 | 18.553 |
| 0 | 13.9732357 | 30.5084746 | 3 | 6 | 3 | 1 | 118 | 13.596 |
| 0 | 29.0122714 | 42.2330097 | 6 | 16 | 6 | 1 | 206 | 23.321 |
| 0 | 20.6484062 | 10.3921569 | 3 | 8 | 3 | 1 | 510 | 54.602 |
| 0 | 37.215985 | 16.0696008 | 10 | 26 | 6 | 1 | 977 | 107.478 |
| 0 | 38.6927552 | 25.7698541 | 9 | 21 | 9 | 1 | 617 | 68.26 |
| 0 | 52.6010933 | 25.8317025 | 11 | 39 | 11 | 1 | 511 | 56.465 |
| 0 | 9.33411323 | 10.5413105 | 4 | 10 | 4 | 1 | 351 | 40.303 |
| 0 | 6.88836453 | 3.31050228 | 2 | 4 | 2 | 1 | 876 | 97.108 |
| 0.00281162 | 1.80354746 | 0.55555556 | 1 | 2 | 1 | 1 | 1980 | 218.267 |
| 0 | 10.2757866 | 8.1300813 | 3 | 6 | 3 | 1 | 369 | 41.305 |
| 0 | 7.98162774 | 15.3846154 | 3 | 6 | 1 | 1 | 208 | 23.534 |
| 0.01058201 | 1.35203054 | 1.46252285 | 1 | 2 | 1 | 1 | 547 | 58.956 |
| 0.00067889 | 2.65286522 | 1.7921147 | 1 | 2 | 1 | 1 | 558 | 63.503 |
| 0 | 16.5466497 | 13.0232558 | 3 | 8 | 1 | 1 | 215 | 23.882 |
| 0.00067889 | 2.48838398 | 1.98675497 | 1 | 2 | 1 | 1 | 453 | 48.724 |
| 0 | 13.9301027 | 21.4611872 | 5 | 12 | 1 | 1 | 219 | 24.742 |
| 0 | 3.3165927 | 3.69230769 | 1 | 2 | 1 | 1 | 325 | 35.679 |
| 0 | 94.864349 | 35.4609929 | 19 | 56 | 17 | 1 | 705 | 74.066 |
| 0 | 22.9124879 | 9.45017182 | 5 | 13 | 3 | 1 | 582 | 62.957 |
| 0 | 3.99524884 | 1.96850394 | 1 | 4 | 1 | 1 | 508 | 57.081 |
| 0 | 23.3874657 | 11.589896 | 6 | 14 | 6 | 1 | 673 | 76.962 |
| 0 | 6.49241464 | 5.60344828 | 2 | 4 | 2 | 1 | 464 | 50.785 |
| 0 | 83.377871 | 31.4917127 | 22 | 52 | 12 | 1 | 724 | 83.212 |
| 0.01058201 | 1.3483344 | 3.33333333 | 1 | 2 | 1 | 1 | 240 | 26.68 |
| 0.00133245 | 2.27886372 | 11.1111111 | 2 | 3 | 2 | 1 | 189 | 21.084 |
| 0 | 12.1701406 | 14.4117647 | 4 | 8 | 4 | 1 | 340 | 37.307 |
| 0.00770142 | 1.44551084 | 2.46153846 | 1 | 2 | 1 | 1 | 325 | 35.367 |
| 0 | 21.4026426 | 32.2869955 | 5 | 20 | 5 | 1 | 223 | 24.808 |
| 0 | 34.0659213 | 17.9941003 | 8 | 19 | 8 | 1 | 678 | 74.715 |
| 0 | 6.02863155 | 7.32484076 | 2 | 4 | 2 | 1 | 314 | 34.04 |
| 0 | 7.31641592 | 4.59081836 | 2 | 4 | 1 | 1 | 501 | 54.827 |
| 0 | 122.867364 | 26.7839687 | 25 | 82 | 11 | 1 | 1023 | 112.824 |
| 0.00067889 | 2.51342785 | 2.08816705 | 1 | 2 | 1 | 1 | 431 | 49.192 |
| 0 | 12.3476081 | 9.48991696 | 5 | 9 | 2 | 1 | 843 | 96.635 |
| 0 | 21.2973003 | 15.2801358 | 5 | 16 | 5 | 1 | 589 | 65.267 |
| 0 | 10.7979255 | 12.0418848 | 2 | 6 | 2 | 1 | 191 | 22.128 |
| 0 | 4.00594694 | 5.10948905 | 1 | 4 | 1 | 1 | 274 | 30.262 |
| 0 | 17.1649672 | 8.67924528 | 5 | 10 | 4 | 1 | 530 | 58.913 |
| 0 | 10.9480828 | 4.75247525 | 3 | 8 | 3 | 1 | 505 | 56.747 |

| | | | | | | | | |
|---|---|---|---|---|---|---|---|---|
| 0 | 16.9098695 | 7.88617886 | 4 | 8 | 4 | 1 | 1230 | 136.289 |
| 0 | 60.0957856 | 44.1295547 | 14 | 42 | 9 | 1 | 247 | 28.285 |
| 0 | 70.8601454 | 35.4330709 | 12 | 66 | 12 | 1 | 381 | 42.617 |
| 0.00133245 | 2.25649024 | 3.06965762 | 1 | 2 | 1 | 1 | 847 | 94.565 |
| 0.00067889 | 2.51655535 | 2.31958763 | 1 | 2 | 1 | 1 | 388 | 42.759 |
| 0 | 5.84193921 | 5.52763819 | 1 | 2 | 1 | 1 | 199 | 22.471 |
| 0 | 3.32367227 | 7.35294118 | 1 | 2 | 1 | 1 | 204 | 22.949 |
| 0 | 62.7149887 | 33.7278107 | 12 | 37 | 12 | 1 | 338 | 35.481 |
| 0.00133245 | 2.42759313 | 5.15021459 | 1 | 2 | 1 | 1 | 233 | 25.439 |
| 0 | 2.77572599 | 5.68181818 | 1 | 2 | 1 | 1 | 264 | 29.185 |
| 0 | 3.06048075 | 5.67567568 | 1 | 2 | 1 | 1 | 370 | 41.543 |
| 0.00133245 | 2.36251027 | 3.66972477 | 1 | 2 | 1 | 1 | 218 | 25.695 |
| 0.00067889 | 2.66534523 | 4.18410042 | 1 | 2 | 1 | 1 | 239 | 25.341 |
| 0 | 5.0897598 | 12.6394052 | 3 | 6 | 3 | 1 | 269 | 30.22 |
| 0 | 7.4251588 | 4.97925311 | 1 | 2 | 1 | 1 | 241 | 26.394 |
| 0.00192431 | 2.00585889 | 3.72093023 | 1 | 2 | 1 | 1 | 215 | 24.926 |
| 0 | 28.842563 | 15.1693667 | 8 | 16 | 8 | 1 | 679 | 73.635 |
| 0 | 7.60774764 | 8.50277264 | 2 | 4 | 2 | 1 | 541 | 59.633 |
| 0 | 16.5057618 | 10.5624143 | 4 | 8 | 4 | 1 | 729 | 78.814 |
| 0.00067889 | 2.55893359 | 2.27920228 | 1 | 2 | 1 | 1 | 351 | 38.394 |
| 0.00161812 | 2.16481693 | 6.70731707 | 1 | 2 | 1 | 1 | 164 | 18.637 |
| 0.00133245 | 2.35783237 | 2.86396181 | 1 | 2 | 1 | 1 | 419 | 45.87 |
| 0 | 5.1786175 | 5.17241379 | 1 | 2 | 1 | 1 | 406 | 45.597 |
| 0 | 23.0426186 | 9.50095969 | 6 | 15 | 6 | 1 | 1042 | 114.683 |
| 0.00457317 | 1.6430186 | 8.58895706 | 1 | 2 | 1 | 1 | 163 | 18.646 |
| 0 | 3.48103752 | 4.06189555 | 2 | 4 | 1 | 1 | 517 | 56.346 |
| 0 | 78.9268068 | 33.7423313 | 20 | 53 | 14 | 1 | 489 | 55.285 |
| 0 | 10.6768157 | 5.42168675 | 4 | 8 | 4 | 1 | 664 | 74.095 |
| 0 | 3.47560388 | 16 | 1 | 2 | 1 | 1 | 75 | 8.739 |
| 0 | 4.11391996 | 3.19488818 | 1 | 4 | 1 | 1 | 313 | 33.823 |
| 0 | 10.8666402 | 8.84955752 | 3 | 6 | 3 | 1 | 452 | 50.877 |
| 0 | 4.81061436 | 6.80272109 | 1 | 8 | 1 | 1 | 147 | 16.118 |
| 0 | 4.12854383 | 2.8436019 | 1 | 2 | 1 | 1 | 422 | 47.434 |
| 0 | 6.22453403 | 6.88622754 | 2 | 6 | 1 | 1 | 334 | 37.896 |
| 0 | 203.770744 | 57.9775281 | 28 | 235 | 1 | 1 | 445 | 49.921 |
| 0 | 195.71703 | 64.1891892 | 30 | 231 | 6 | 1 | 444 | 49.639 |
| 0 | 19.3362498 | 7.55208333 | 5 | 12 | 5 | 1 | 768 | 84.25 |
| 0 | 3.94058166 | 2.28452752 | 2 | 3 | 2 | 1 | 963 | 104.833 |
| 0 | 3.32890541 | 4.84581498 | 2 | 4 | 2 | 1 | 227 | 22.68 |
| 0 | 4.46566923 | 3.69487485 | 2 | 4 | 2 | 1 | 839 | 94.453 |
| 0.01680912 | 1.17069623 | 4.36363636 | 1 | 1 | 1 | 1 | 275 | 29.719 |
| 0 | 103.754775 | 37.521815 | 23 | 58 | 23 | 1 | 573 | 61.016 |
| 0.0039914 | 1.69164905 | 2.76923077 | 1 | 2 | 1 | 1 | 325 | 35.956 |
| 0 | 8.56729818 | 5.10046368 | 3 | 6 | 3 | 1 | 647 | 68.953 |
| 0 | 7.22355179 | 2.7638191 | 3 | 6 | 3 | 1 | 1194 | 133.414 |
| 0 | 186.977404 | 75.6143667 | 28 | 139 | 28 | 1 | 529 | 56.525 |
| 0.00067889 | 2.49214413 | 0.95602294 | 1 | 2 | 1 | 1 | 1046 | 116.85 |
| 0 | 17.9433482 | 8.23529412 | 7 | 14 | 5 | 1 | 1020 | 111.771 |
| 0 | 14.6698378 | 19.2105263 | 5 | 12 | 5 | 1 | 380 | 41.769 |
| 0 | 98.4104753 | 9.9009901 | 28 | 61 | 28 | 1 | 4646 | 532.072 |
| 0 | 4.33714777 | 14.2045455 | 1 | 3 | 1 | 1 | 176 | 20.096 |
| 0.01746971 | 1.11730474 | 5.64102564 | 1 | 2 | 1 | 1 | 195 | 21.686 |
| 0.01314636 | 1.28726614 | 1.5 | 1 | 1 | 1 | 1 | 600 | 64.139 |
| 0 | 8.13643124 | 7.34463277 | 2 | 8 | 1 | 1 | 354 | 40.335 |
| 0 | 9.83485828 | 12.6213592 | 2 | 4 | 2 | 1 | 206 | 23.552 |
| 0 | 15.4682593 | 28.9855072 | 5 | 10 | 5 | 1 | 207 | 23.475 |
| 0 | 17.5021527 | 9.67032967 | 5 | 10 | 5 | 1 | 455 | 49.843 |
| 0 | 21.1881338 | 8.72135994 | 7 | 16 | 1 | 1 | 1353 | 152.69 |
| 0.00067889 | 2.61726273 | 0.83179298 | 1 | 2 | 1 | 1 | 1082 | 118.985 |
| 0.0135603 | 1.26544018 | 3.46666667 | 1 | 1 | 1 | 1 | 375 | 39.81 |
| 0 | 45.7804322 | 16.0714286 | 10 | 23 | 10 | 1 | 840 | 94.271 |
| 0 | 12.1351396 | 9.65665236 | 4 | 8 | 4 | 1 | 466 | 51.68 |

| | | | | | | | | |
|---|---|---|---|---|---|---|---|---|
| 0 | 5.72509131 | 5.93692022 | 3 | 5 | 3 | 1 | 539 | 57.888 |
| 0 | 18.0102928 | 14.2061281 | 4 | 8 | 3 | 1 | 359 | 42.115 |
| 0.00067889 | 2.67612939 | 3.72492837 | 1 | 2 | 1 | 1 | 349 | 38.928 |
| 0.00339716 | 1.76270766 | 2.82861897 | 1 | 1 | 1 | 1 | 601 | 63.812 |
| 0.00457317 | 1.65619767 | 1.97183099 | 1 | 2 | 1 | 1 | 710 | 80.648 |
| 0 | 5.49882485 | 3.85356455 | 2 | 4 | 2 | 1 | 519 | 56.131 |
| 0.00133245 | 2.3990271 | 1.41442716 | 1 | 2 | 1 | 1 | 707 | 76.132 |
| 0 | 10.6865328 | 6.97247706 | 3 | 6 | 3 | 1 | 545 | 60.609 |
| 0 | 4.24169382 | 0.80723281 | 1 | 2 | 1 | 1 | 3097 | 346.683 |
| 0 | 6.9677843 | 1.32275132 | 1 | 2 | 1 | 1 | 1134 | 129.433 |
| 0 | 2.7070797 | 4.80769231 | 1 | 2 | 1 | 1 | 208 | 23.561 |
| 0 | 4.66015122 | 9.94764398 | 1 | 2 | 1 | 1 | 191 | 21.295 |
| 0 | 3.64955814 | 8.02919708 | 1 | 2 | 1 | 1 | 137 | 15.272 |
| 0 | 14.2650828 | 6.25 | 2 | 6 | 2 | 1 | 480 | 52.612 |
| 0 | 25.2901279 | 16.7464115 | 5 | 12 | 3 | 1 | 418 | 46.273 |
| 0 | 22.2838459 | 15.942029 | 5 | 14 | 5 | 1 | 483 | 55.847 |
| 0 | 26.4400109 | 9.33226066 | 9 | 16 | 5 | 1 | 1243 | 136.789 |
| 0 | 4.90757027 | 4.52755906 | 2 | 4 | 2 | 1 | 508 | 56.905 |
| 0 | 5.30741329 | 3.6900369 | 2 | 4 | 2 | 1 | 813 | 86.452 |
| 0 | 9.50768411 | 45.1612903 | 3 | 6 | 3 | 1 | 62 | 6.787 |
| 0 | 15.7205473 | 14.7810219 | 6 | 14 | 6 | 1 | 548 | 59.583 |
| 0 | 3.92093313 | 10 | 2 | 4 | 2 | 1 | 290 | 33.345 |
| 0 | 4.37860488 | 6.6252588 | 2 | 6 | 2 | 1 | 483 | 50.949 |
| 0 | 7.47394801 | 10.3658537 | 3 | 6 | 3 | 1 | 328 | 36.196 |
| 0 | 137.990062 | 48.1012658 | 30 | 113 | 30 | 1 | 553 | 59.714 |
| 0.00192431 | 2.02516604 | 3.3557047 | 1 | 2 | 1 | 1 | 298 | 32.975 |
| 0 | 3.18223637 | 5.35714286 | 1 | 2 | 1 | 1 | 168 | 17.479 |
| 0 | 3.25258819 | 2.20750552 | 1 | 2 | 1 | 1 | 453 | 48.413 |
| 0 | 161.760525 | 59.7777778 | 27 | 165 | 10 | 1 | 450 | 50.4 |
| 0.00161812 | 2.15131835 | 9.21985816 | 1 | 2 | 1 | 1 | 141 | 14.997 |
| 0 | 2.70421306 | 5.19480519 | 1 | 2 | 1 | 1 | 154 | 17.859 |
| 0.00067889 | 2.68110229 | 3.18181818 | 1 | 2 | 1 | 1 | 440 | 49.576 |
| 0 | 4.05036608 | 3.87409201 | 1 | 2 | 1 | 1 | 413 | 44.621 |
| 0 | 10.583602 | 5.77249576 | 2 | 4 | 2 | 1 | 589 | 64.092 |
| 0 | 4.47198366 | 1.64986251 | 1 | 2 | 1 | 1 | 1091 | 122.933 |
| 0 | 7.00395794 | 8.46681922 | 2 | 4 | 2 | 1 | 437 | 47.534 |
| 0 | 7.48254017 | 3.84615385 | 1 | 4 | 1 | 1 | 520 | 58.134 |
| 0 | 3.32230182 | 2.13815789 | 1 | 2 | 1 | 1 | 608 | 67.412 |
| 0 | 28.0442986 | 18.1208054 | 7 | 22 | 3 | 1 | 298 | 33.043 |
| 0.01680912 | 1.22061999 | 0.22479127 | 1 | 2 | 1 | 1 | 3114 | 357.306 |
| 0.01746971 | 1.08900228 | 2.5462963 | 1 | 1 | 1 | 1 | 432 | 47.093 |
| 0.00067889 | 2.66614985 | 2.40963855 | 1 | 3 | 1 | 1 | 332 | 35.945 |
| 0 | 12.0696637 | 6.55172414 | 6 | 13 | 1 | 1 | 870 | 97.905 |
| 0 | 14.0625546 | 30.1507538 | 4 | 8 | 4 | 1 | 199 | 22.458 |
| 0 | 5.94467015 | 15.9090909 | 2 | 4 | 2 | 1 | 176 | 18.974 |
| 0 | 3.15236566 | 5.26315789 | 1 | 2 | 1 | 1 | 247 | 28.175 |
| 0 | 14.1222028 | 7.27470141 | 3 | 7 | 3 | 1 | 921 | 100.304 |
| 0 | 4.96697856 | 2.49307479 | 1 | 2 | 1 | 1 | 722 | 80.477 |
| 0 | 10.4719913 | 5.75539568 | 2 | 4 | 2 | 1 | 556 | 60.306 |
| 0 | 15.7310714 | 16.4739884 | 5 | 12 | 1 | 1 | 346 | 39.307 |
| 0 | 12.2760317 | 6.56565657 | 2 | 6 | 2 | 1 | 396 | 44.715 |
| 0.00281162 | 1.86138157 | 1.66073547 | 1 | 2 | 1 | 1 | 843 | 94.12 |
| 0 | 3.88346557 | 6.7961165 | 2 | 3 | 2 | 1 | 206 | 23.293 |
| 0 | 5.89380563 | 4.92125984 | 2 | 4 | 1 | 1 | 508 | 55.047 |
| 0.00281162 | 1.82419837 | 3.46820809 | 1 | 2 | 1 | 1 | 346 | 37.411 |
| 0 | 35.7227418 | 18.9504373 | 8 | 20 | 5 | 1 | 686 | 74.216 |
| 0 | 27.4112646 | 17.2477064 | 6 | 15 | 6 | 1 | 545 | 60.495 |
| 0 | 5.68084614 | 6.61764706 | 2 | 4 | 2 | 1 | 408 | 44.938 |
| 0 | 13.9429518 | 30.9210526 | 4 | 10 | 4 | 1 | 152 | 17.287 |
| 0 | 6.93579001 | 10.2777778 | 3 | 6 | 3 | 1 | 360 | 41.363 |
| 0 | 7.29657105 | 10.3603604 | 2 | 7 | 2 | 1 | 222 | 24.735 |
| 0.01440507 | 1.2486439 | 2.40137221 | 1 | 2 | 1 | 1 | 583 | 65.061 |

| | | | | | | | | |
|---|---|---|---|---|---|---|---|---|
| 0 | 3.59808275 | 4.296875 | 1 | 2 | 1 | 1 | 256 | 28.512 |
| 0 | 30.537086 | 22.3350254 | 5 | 28 | 5 | 1 | 197 | 21.48 |
| 0 | 8.1522339 | 4.01188707 | 3 | 6 | 3 | 1 | 673 | 75.826 |
| 0 | 2.73660067 | 8.82352941 | 1 | 3 | 1 | 1 | 238 | 27.35 |
| 0 | 25.4539237 | 14.1860465 | 6 | 18 | 6 | 1 | 430 | 47.487 |
| 0 | 101.489612 | 49.1582492 | 26 | 103 | 26 | 1 | 594 | 67.526 |
| 0 | 4.32590777 | 2.78787879 | 2 | 3 | 2 | 1 | 825 | 90.528 |
| 0 | 3.37369686 | 5.09383378 | 2 | 4 | 2 | 1 | 373 | 42.037 |
| 0 | 3.35654732 | 3.33333333 | 1 | 2 | 1 | 1 | 240 | 27.112 |
| 0 | 4.49962629 | 5.58139535 | 1 | 2 | 1 | 1 | 215 | 23.417 |
| 0.00161812 | 2.09420412 | 13.4831461 | 1 | 2 | 1 | 1 | 89 | 10.359 |
| 0 | 10.9309702 | 5.38922156 | 2 | 4 | 2 | 1 | 501 | 54.32 |
| 0 | 5.15459186 | 4.98442368 | 1 | 2 | 1 | 1 | 321 | 35.032 |
| 0 | 2.68867005 | 2.11480363 | 1 | 2 | 1 | 1 | 662 | 73.198 |
| 0 | 4.54414297 | 5.89390963 | 2 | 4 | 2 | 1 | 509 | 54.143 |
| 0 | 7.38565959 | 0.89149261 | 3 | 6 | 3 | 1 | 3926 | 416.214 |
| 0 | 5.17398885 | 1.12311015 | 3 | 6 | 3 | 1 | 2315 | 254.429 |
| 0 | 10.3868447 | 8.50340136 | 3 | 8 | 2 | 1 | 294 | 32.593 |
| 0 | 45.8945966 | 15.2019002 | 10 | 28 | 10 | 1 | 421 | 47.549 |
| 0 | 32.1640779 | 3.56760614 | 7 | 16 | 6 | 1 | 2803 | 305.298 |
| 0 | 8.70048351 | 15.8653846 | 3 | 6 | 1 | 1 | 208 | 23.447 |
| 0 | 3.47846966 | 7.76255708 | 1 | 4 | 1 | 1 | 219 | 24.96 |
| 0 | 3.21218543 | 13.2743363 | 1 | 2 | 1 | 1 | 113 | 12.544 |
| 0 | 67.6287801 | 37.9373849 | 19 | 45 | 17 | 1 | 543 | 61.479 |
| 0 | 7.21235026 | 8.65921788 | 2 | 4 | 2 | 1 | 358 | 41.32 |
| 0 | 12.1091182 | 16.2361624 | 3 | 10 | 3 | 1 | 271 | 31.264 |
| 0 | 3.78542105 | 3.25047801 | 1 | 2 | 1 | 1 | 523 | 58.706 |
| 0 | 4.30909545 | 4.33526012 | 1 | 2 | 1 | 1 | 346 | 36.227 |
| 0 | 156.459681 | 19.4162437 | 42 | 102 | 40 | 1 | 2364 | 274.439 |
| 0 | 3.8706324 | 2.56959315 | 1 | 2 | 1 | 1 | 467 | 50.354 |
| 0 | 42.208523 | 41.9354839 | 12 | 30 | 6 | 1 | 248 | 28.853 |
| 0 | 11.8963852 | 4.31107354 | 4 | 7 | 4 | 1 | 1183 | 127.027 |
| 0.00067889 | 2.46029676 | 8.92857143 | 1 | 3 | 1 | 1 | 280 | 31.609 |
| 0 | 9.00944086 | 8.91364903 | 2 | 4 | 1 | 1 | 359 | 42.097 |
| 0 | 4.75294119 | 7.98816568 | 2 | 3 | 2 | 1 | 338 | 37.896 |
| 0 | 6.66282997 | 2.02507232 | 2 | 8 | 1 | 1 | 1037 | 110.956 |
| 0.00457317 | 1.63921731 | 10.8695652 | 1 | 2 | 1 | 1 | 184 | 20.764 |
| 0 | 2.96377046 | 1.59118727 | 1 | 2 | 1 | 1 | 817 | 89.28 |
| 0 | 5.14715418 | 1.53649168 | 1 | 2 | 1 | 1 | 781 | 85.442 |
| 0 | 17.2837505 | 4.578564 | 3 | 6 | 3 | 1 | 961 | 103.821 |
| 0.00596659 | 1.52870829 | 2.5 | 1 | 2 | 1 | 1 | 480 | 54.918 |
| 0 | 9.20480021 | 2.10526316 | 2 | 4 | 2 | 1 | 1140 | 126.887 |
| 0 | 4.70879697 | 3.76344086 | 2 | 4 | 2 | 1 | 744 | 84.037 |
| 0 | 30.1207539 | 11.5384615 | 7 | 14 | 7 | 1 | 858 | 95.725 |
| 0.00281162 | 1.86264589 | 3.63636364 | 1 | 2 | 1 | 1 | 330 | 37.548 |
| 0.00457317 | 1.65915945 | 2.10843373 | 1 | 2 | 1 | 1 | 332 | 31.536 |
| 0 | 62.7741362 | 20.2576112 | 17 | 41 | 9 | 1 | 854 | 98.099 |
| 0 | 15.4499163 | 1.66126418 | 4 | 10 | 3 | 1 | 2468 | 270.468 |
| 0 | 9.72435235 | 11.0701107 | 3 | 5 | 2 | 1 | 271 | 31.52 |
| 0 | 5.60432701 | 3.43007916 | 2 | 4 | 2 | 1 | 758 | 83.626 |
| 0 | 5.67695426 | 1.97368421 | 1 | 2 | 1 | 1 | 608 | 66.346 |
| 0 | 33.4297311 | 22.4832215 | 8 | 34 | 4 | 1 | 298 | 32.831 |
| 0 | 21.8512991 | 13.6023916 | 7 | 14 | 7 | 1 | 669 | 73.414 |
| 0 | 5.57495513 | 6.37681159 | 1 | 2 | 1 | 1 | 345 | 37.395 |
| 0 | 4.79236563 | 6.39534884 | 1 | 2 | 1 | 1 | 172 | 18.974 |
| 0.00067889 | 2.65099149 | 3.76506024 | 2 | 3 | 2 | 1 | 664 | 72.645 |
| 0 | 13.0866283 | 14.2061281 | 5 | 9 | 5 | 1 | 359 | 39.208 |
| 0 | 11.7649438 | 13.6871508 | 5 | 9 | 5 | 1 | 358 | 40.678 |
| 0 | 5.75055574 | 1.98821797 | 3 | 6 | 3 | 1 | 1358 | 149.467 |
| 0.00252525 | 1.89790947 | 5.5028463 | 1 | 2 | 1 | 1 | 527 | 58.91 |
| 0 | 2.76371472 | 2.73684211 | 1 | 2 | 1 | 1 | 475 | 51.641 |
| 0 | 60.1282575 | 20.7423581 | 20 | 49 | 17 | 1 | 916 | 102.411 |

| | | | | | | | | |
|---|---|---|---|---|---|---|---|---|
| 0 | 12.2818971 | 5.65832427 | 4 | 10 | 4 | 1 | 919 | 103.211 |
| 0.00624442 | 1.50584541 | 2.91970803 | 1 | 2 | 1 | 1 | 274 | 29.649 |
| 0 | 4.07857366 | 1.56599553 | 1 | 2 | 1 | 1 | 894 | 101.064 |
| 0 | 3.57593547 | 1.55210643 | 1 | 2 | 1 | 1 | 902 | 100.707 |
| 0 | 2.79128998 | 2.71604938 | 1 | 2 | 1 | 1 | 405 | 45.486 |
| 0 | 10.852696 | 7.07456979 | 3 | 6 | 3 | 1 | 523 | 56.544 |
| 0.01440507 | 1.24328784 | 2.7972028 | 1 | 2 | 1 | 1 | 286 | 32.215 |
| 0.01680912 | 1.16647042 | 2.30179028 | 1 | 1 | 1 | 1 | 782 | 89.757 |
| 0.00067889 | 2.46371152 | 10.2941176 | 2 | 4 | 2 | 1 | 136 | 15.379 |
| 0.00827668 | 1.39750593 | 4.69483568 | 1 | 2 | 1 | 1 | 213 | 23.548 |
| 0.0039914 | 1.72284939 | 2.19298246 | 1 | 2 | 1 | 1 | 456 | 51.769 |
| 0 | 5.5352039 | 4.02097902 | 2 | 4 | 2 | 1 | 572 | 61.838 |
| 0.00161812 | 2.02835312 | 5.04587156 | 1 | 2 | 1 | 1 | 218 | 24.473 |
| 0 | 20.5150483 | 44.6666667 | 5 | 22 | 5 | 1 | 150 | 16.752 |
| 0 | 25.9050507 | 16.967509 | 5 | 20 | 5 | 1 | 277 | 30.057 |
| 0 | 10.0525336 | 10.1298701 | 2 | 4 | 1 | 1 | 385 | 42.44 |
| 0.00711955 | 1.50058787 | 0.52122115 | 1 | 2 | 1 | 1 | 1343 | 148.191 |
| 0 | 2.70399333 | 0.95137421 | 1 | 2 | 1 | 1 | 946 | 102.312 |
| 0 | 4.41907502 | 5.43933054 | 1 | 4 | 1 | 1 | 239 | 26.737 |
| 0 | 3.93341261 | 7.12643678 | 2 | 4 | 2 | 1 | 435 | 49.809 |
| 0.00252525 | 1.88140463 | 2.82258065 | 1 | 2 | 1 | 1 | 248 | 28.76 |
| 0 | 7.96505406 | 2.37659963 | 2 | 4 | 2 | 1 | 1094 | 123.102 |
| 0 | 6.4688109 | 3.07287094 | 3 | 6 | 2 | 1 | 1139 | 126.102 |
| 0 | 31.9406887 | 27.8169014 | 7 | 20 | 6 | 1 | 284 | 30.77 |
| 0 | 5.97715939 | 6.55737705 | 1 | 2 | 1 | 1 | 244 | 25.773 |
| 0 | 2.81191563 | 4.15224913 | 1 | 4 | 1 | 1 | 289 | 32.41 |
| 0.00339716 | 1.7577071 | 3.61445783 | 1 | 2 | 1 | 1 | 415 | 48.013 |
| 0 | 8.33207144 | 5.79514825 | 3 | 6 | 3 | 1 | 742 | 82.642 |
| 0 | 3.4024145 | 3.3557047 | 1 | 2 | 1 | 1 | 298 | 33.212 |
| 0 | 10.5977635 | 11.4130435 | 3 | 8 | 3 | 1 | 184 | 20.157 |
| 0 | 6.38447078 | 2.16138329 | 1 | 2 | 1 | 1 | 694 | 76.824 |
| 0 | 13.6248122 | 18.3946488 | 4 | 8 | 4 | 1 | 299 | 33.276 |
| 0 | 6.87442229 | 8.03571429 | 2 | 4 | 2 | 1 | 336 | 36.671 |
| 0 | 3.75621808 | 4.01606426 | 1 | 2 | 1 | 1 | 249 | 27.374 |
| 0 | 7.19134672 | 12.3595506 | 2 | 4 | 2 | 1 | 356 | 37.474 |
| 0.02239642 | 1.02127182 | 1.11214087 | 1 | 1 | 1 | 1 | 1079 | 116.266 |
| 0 | 20.0997794 | 25.2873563 | 6 | 12 | 6 | 1 | 174 | 19.811 |
| 0 | 10.8526964 | 5.29953917 | 2 | 10 | 2 | 1 | 434 | 48.683 |
| 0 | 32.263527 | 17.4384236 | 12 | 24 | 12 | 1 | 1015 | 117.67 |
| 0.01746971 | 1.11571454 | 1.70575693 | 1 | 1 | 1 | 1 | 469 | 51.943 |
| 0.00223642 | 1.92154318 | 8.45070423 | 1 | 2 | 1 | 1 | 142 | 15.995 |
| 0 | 35.8786798 | 33.2167832 | 8 | 28 | 8 | 1 | 286 | 30.772 |
| 0 | 45.0016833 | 8.74689826 | 10 | 20 | 10 | 1 | 1612 | 177.438 |
| 0.00067889 | 2.55626776 | 1.60366552 | 1 | 1 | 1 | 1 | 873 | 96.711 |
| 0 | 20.5048648 | 23.5 | 5 | 14 | 2 | 1 | 200 | 22.527 |
| 0 | 3.68973163 | 4.72440945 | 1 | 2 | 1 | 1 | 381 | 43.046 |
| 0 | 27.2718166 | 15.5124654 | 4 | 10 | 4 | 1 | 361 | 39.812 |
| 0 | 41.7273586 | 9.5890411 | 8 | 16 | 1 | 1 | 949 | 104.54 |
| 0.00067889 | 2.54821356 | 2.31578947 | 1 | 2 | 1 | 1 | 475 | 52.845 |
| 0 | 3.88672531 | 2.20820189 | 1 | 2 | 1 | 1 | 634 | 71.325 |
| 0 | 31.2406071 | 13.6176066 | 7 | 18 | 7 | 1 | 727 | 80.802 |
| 0 | 5.07857366 | 4.59290188 | 1 | 4 | 1 | 1 | 479 | 51.624 |
| 0 | 12.8296075 | 8.79310345 | 4 | 8 | 4 | 1 | 580 | 63.75 |
| 0 | 28.6876335 | 19.1616766 | 8 | 16 | 8 | 1 | 334 | 36.615 |
| 0.00252525 | 1.90657831 | 1.59090909 | 1 | 4 | 1 | 1 | 440 | 47.702 |
| 0 | 7.1641649 | 4.7318612 | 2 | 4 | 2 | 1 | 634 | 70.915 |
| 0 | 23.0427929 | 22.3300971 | 7 | 14 | 7 | 1 | 309 | 33.351 |
| 0 | 109.839276 | 42.8360414 | 20 | 66 | 1 | 1 | 677 | 73.457 |
| 0 | 4.134104 | 10.1604278 | 2 | 4 | 2 | 1 | 187 | 21.574 |
| 0.00133245 | 2.34756025 | 4.8245614 | 1 | 4 | 1 | 1 | 228 | 26.356 |
| 0 | 7.76719319 | 9.84455959 | 2 | 4 | 2 | 1 | 193 | 22.014 |
| 0.00161812 | 2.19205928 | 3.13253012 | 1 | 2 | 1 | 1 | 415 | 46.497 |

| | | | | | | | | |
|---|---|---|---|---|---|---|---|---|
| 0 | 119.709258 | 59.2 | 24 | 105 | 8 | 1 | 375 | 41.766 |
| 0 | 25.9083205 | 11.3360324 | 6 | 14 | 6 | 1 | 741 | 80.945 |
| 0.00067889 | 2.54242085 | 3.09278351 | 1 | 2 | 1 | 1 | 291 | 33.12 |
| 0 | 80.2980218 | 29.2561983 | 16 | 55 | 16 | 1 | 605 | 65.889 |
| 0 | 7.40135239 | 4.86842105 | 2 | 4 | 2 | 1 | 760 | 83.345 |
| 0 | 13.2952398 | 7.99086758 | 4 | 8 | 4 | 1 | 438 | 50.258 |
| 0 | 2.81987412 | 1.80995475 | 1 | 2 | 1 | 1 | 884 | 97.284 |
| 0 | 38.0042796 | 15.4255319 | 9 | 25 | 5 | 1 | 940 | 104.024 |
| 0 | 42.5312374 | 26.5550239 | 11 | 26 | 11 | 1 | 418 | 45.232 |
| 0 | 8.83520189 | 12.8514056 | 3 | 6 | 2 | 1 | 249 | 27.019 |
| 0.00161812 | 2.05320397 | 5.94405594 | 1 | 2 | 1 | 1 | 286 | 30.339 |
| 0 | 3.90135627 | 7.14285714 | 1 | 2 | 1 | 1 | 140 | 16.256 |
| 0 | 178.781777 | 35.9649123 | 36 | 121 | 20 | 1 | 1026 | 112.987 |
| 0 | 3.66574736 | 7.0754717 | 1 | 2 | 1 | 1 | 212 | 23.989 |
| 0 | 3.36916548 | 4.68085106 | 1 | 2 | 1 | 1 | 235 | 26.393 |
| 0 | 5.48701493 | 2.88659794 | 2 | 4 | 2 | 1 | 970 | 105.608 |
| 0 | 2.99267905 | 9.30232558 | 1 | 2 | 1 | 1 | 215 | 24.725 |
| 0 | 21.7897134 | 9.50871632 | 6 | 12 | 6 | 1 | 631 | 68.77 |
| 0 | 6.20593962 | 20.4633205 | 3 | 6 | 3 | 1 | 259 | 28.468 |
| 0 | 4.9303319 | 1.64473684 | 1 | 2 | 1 | 1 | 912 | 99.69 |
| 0 | 5.40022606 | 2.75 | 1 | 2 | 1 | 1 | 400 | 43.842 |
| 0 | 12.9391652 | 6.19307832 | 2 | 4 | 2 | 1 | 549 | 62.465 |
| 0.01680912 | 1.20880075 | 1.69133192 | 1 | 1 | 1 | 1 | 1419 | 157.48 |
| 0 | 31.152254 | 15.0867824 | 7 | 14 | 7 | 1 | 749 | 83.347 |
| 0.00192431 | 1.99697053 | 6.17977528 | 1 | 2 | 1 | 1 | 178 | 20.533 |
| 0 | 3.63695241 | 3.95683453 | 1 | 2 | 1 | 1 | 278 | 31.884 |
| 0 | 9.05958258 | 6.04575163 | 3 | 8 | 3 | 1 | 612 | 67.598 |
| 0 | 21.4709221 | 25.3676471 | 6 | 12 | 6 | 1 | 272 | 29.786 |
| 0 | 10.7756929 | 12.1693122 | 3 | 6 | 3 | 1 | 189 | 19.878 |
| 0 | 9.02225134 | 3.96039604 | 2 | 4 | 2 | 1 | 707 | 76.102 |
| 0.00223642 | 1.92811799 | 2.47678019 | 1 | 2 | 1 | 1 | 323 | 35.246 |
| 0 | 9.98313553 | 4.64646465 | 2 | 4 | 2 | 1 | 495 | 57.632 |
| 0.00192431 | 1.96217525 | 0.98800282 | 1 | 2 | 1 | 1 | 1417 | 164.203 |
| 0 | 19.5121657 | 13.1578947 | 5 | 12 | 5 | 1 | 608 | 66.586 |
| 0 | 5.31743492 | 4.61254613 | 2 | 4 | 1 | 1 | 542 | 58.741 |
| 0 | 3.36231018 | 2.67639903 | 1 | 2 | 1 | 1 | 411 | 46.455 |
| 0 | 7.40909266 | 8.2010582 | 2 | 4 | 2 | 1 | 378 | 39.571 |
| 0.00516403 | 1.61403643 | 0.55555556 | 1 | 2 | 1 | 1 | 1800 | 203.003 |
| 0 | 16.3247263 | 12.1287129 | 4 | 8 | 2 | 1 | 404 | 45.49 |
| 0 | 4.57024772 | 3.04709141 | 1 | 2 | 1 | 1 | 361 | 39.92 |
| 0.00457317 | 1.6505282 | 2.09318028 | 1 | 1 | 1 | 1 | 1481 | 157.347 |
| 0 | 28.5746814 | 17.6360225 | 9 | 20 | 3 | 1 | 533 | 59.965 |
| 0 | 2.98088371 | 0.82304527 | 1 | 2 | 1 | 1 | 1701 | 190.599 |
| 0.00067889 | 2.51841406 | 0.8 | 1 | 2 | 1 | 1 | 1250 | 135.157 |
| 0.00067889 | 2.64569944 | 0.70257611 | 1 | 2 | 1 | 1 | 1281 | 138.748 |
| 0 | 11.7380762 | 19.4029851 | 3 | 10 | 3 | 1 | 134 | 15.385 |
| 0 | 21.1092023 | 24 | 5 | 18 | 5 | 1 | 150 | 16.347 |
| 0.00067889 | 2.56447415 | 1.52740341 | 1 | 2 | 1 | 1 | 1113 | 124.336 |
| 0 | 4.22773721 | 2.18037661 | 2 | 6 | 2 | 1 | 1009 | 115.8 |
| 0.01173365 | 1.30653687 | 7.58293839 | 1 | 1 | 1 | 1 | 211 | 24.472 |
| 0 | 5.85861039 | 5.66037736 | 3 | 6 | 3 | 1 | 636 | 70.626 |
| 0 | 4.16367588 | 0.40520367 | 1 | 2 | 1 | 1 | 4689 | 532.145 |
| 0 | 63.9969572 | 39.5918367 | 13 | 35 | 11 | 1 | 245 | 27.728 |
| 0 | 95.4422337 | 28.4562212 | 27 | 61 | 1 | 1 | 868 | 97.647 |
| 0.00067889 | 2.61672335 | 5.71428571 | 1 | 2 | 1 | 1 | 210 | 23.396 |
| 0 | 3.2432118 | 2.86532951 | 1 | 2 | 1 | 1 | 349 | 36.75 |
| 0 | 2.80299527 | 7.84313725 | 1 | 2 | 1 | 1 | 153 | 16.931 |
| 0 | 8.87981108 | 4.19161677 | 3 | 7 | 1 | 1 | 835 | 90.822 |
| 0 | 30.8635679 | 12.716763 | 7 | 18 | 7 | 1 | 692 | 76.97 |
| 0 | 32.436125 | 10.8333333 | 9 | 24 | 9 | 1 | 960 | 106.871 |
| 0 | 3.35733767 | 1.76211454 | 1 | 2 | 1 | 1 | 1135 | 123.85 |
| 0 | 3.76199878 | 5.46357616 | 2 | 5 | 2 | 1 | 604 | 67.026 |

| | | | | | | | | |
|---|---|---|---|---|---|---|---|---|
| 0.00067889 | 2.43097741 | 10.4761905 | 1 | 2 | 1 | 1 | 210 | 23.69 |
| 0 | 8.58047777 | 8.43023256 | 2 | 4 | 2 | 1 | 344 | 37.854 |
| 0 | 3.89756629 | 3.54223433 | 1 | 2 | 1 | 1 | 367 | 40.498 |
| 0.00067889 | 2.65876338 | 8.21917808 | 1 | 2 | 1 | 1 | 146 | 16.997 |
| 0 | 29.3416979 | 7.92634107 | 8 | 15 | 4 | 1 | 1249 | 137.717 |
| 0 | 12.7284989 | 5.32276331 | 5 | 11 | 3 | 1 | 883 | 96.846 |
| 0 | 4.08597435 | 1.55223881 | 1 | 2 | 1 | 1 | 1675 | 187.15 |
| 0 | 26.018856 | 15.1351351 | 7 | 13 | 7 | 1 | 740 | 82.173 |
| 0.00457317 | 1.66494348 | 0.58862001 | 1 | 1 | 1 | 1 | 3058 | 341.933 |
| 0 | 21.8087175 | 21.4723926 | 7 | 18 | 1 | 1 | 326 | 37.43 |
| 0 | 25.2897249 | 27.0916335 | 7 | 18 | 1 | 1 | 251 | 28.806 |
| 0 | 9.34144774 | 8.20895522 | 2 | 8 | 2 | 1 | 536 | 58.682 |
| 0 | 12.4063598 | 10.5691057 | 2 | 6 | 1 | 1 | 369 | 40.899 |
| 0 | 19.5346951 | 42.3728814 | 4 | 10 | 4 | 1 | 118 | 12.926 |
| 0 | 51.4627194 | 13.6698212 | 12 | 24 | 5 | 1 | 951 | 105.625 |
| 0 | 47.9780765 | 31.8681319 | 11 | 32 | 11 | 1 | 364 | 39.431 |
| 0 | 198.697903 | 31.510107 | 44 | 133 | 44 | 1 | 1682 | 192.276 |
| 0 | 10.1878048 | 5.33642691 | 5 | 9 | 5 | 1 | 862 | 89.739 |
| 0 | 17.2522669 | 6.39810427 | 4 | 10 | 4 | 1 | 844 | 97.194 |
| 0 | 8.40224209 | 16.5714286 | 2 | 5 | 2 | 1 | 175 | 19.446 |
| 0.00067889 | 2.53417118 | 8.52713178 | 1 | 2 | 1 | 1 | 129 | 14.988 |
| 0 | 7.3902989 | 13.2231405 | 1 | 4 | 1 | 1 | 121 | 13.354 |
| 0.01798258 | 1.05051241 | 2.77227723 | 1 | 1 | 1 | 1 | 1010 | 107.699 |
| 0.00192431 | 2.0125572 | 0.76687117 | 1 | 2 | 1 | 1 | 1304 | 144.36 |
| 0 | 10.7721154 | 9.38967136 | 2 | 4 | 2 | 1 | 426 | 46.948 |
| 0 | 20.9799869 | 15.245478 | 6 | 12 | 6 | 1 | 387 | 42.384 |
| 0.01058201 | 1.3357342 | 2.49433107 | 1 | 1 | 1 | 1 | 441 | 47.593 |
| 0.00252525 | 1.90274269 | 1.69082126 | 1 | 2 | 1 | 1 | 414 | 46.63 |
| 0 | 4.79102148 | 2.26804124 | 1 | 2 | 1 | 1 | 485 | 55.307 |
| 0 | 13.4147646 | 6.27118644 | 3 | 8 | 3 | 1 | 590 | 68.037 |
| 0.01314636 | 1.2866775 | 2.84552846 | 1 | 2 | 1 | 1 | 492 | 53.867 |
| 0 | 28.0421157 | 33.4437086 | 8 | 19 | 8 | 1 | 302 | 33.896 |
| 0.00161812 | 2.12222565 | 5.19480519 | 1 | 2 | 1 | 1 | 385 | 44.938 |
| 0 | 10.9651785 | 5.23255814 | 3 | 6 | 3 | 1 | 860 | 97.398 |
| 0 | 172.973432 | 64.1891892 | 28 | 198 | 7 | 1 | 444 | 49.554 |
| 0 | 71.4383397 | 17.9104478 | 6 | 43 | 6 | 1 | 335 | 36.03 |
| 0.003698 | 1.73002032 | 2.98507463 | 1 | 2 | 1 | 1 | 402 | 45.189 |
| 0 | 6.46521984 | 6.34328358 | 2 | 4 | 2 | 1 | 536 | 61.274 |
| 0 | 8.63590208 | 5.63549161 | 3 | 8 | 3 | 1 | 834 | 89.776 |
| 0 | 2.81844223 | 5.57275542 | 1 | 2 | 1 | 1 | 323 | 37.289 |
| 0 | 5.78015361 | 4.01891253 | 1 | 2 | 1 | 1 | 423 | 48.768 |
| 0 | 5.57446578 | 1.48083624 | 1 | 2 | 1 | 1 | 1148 | 129.473 |
| 0 | 3.45075144 | 2.06451613 | 1 | 2 | 1 | 1 | 775 | 84.878 |
| 0 | 7.31247104 | 1.48212729 | 1 | 2 | 1 | 1 | 1147 | 122.118 |
| 0 | 28.6590559 | 38.1578947 | 6 | 14 | 3 | 1 | 152 | 15.964 |
| 0 | 21.5963164 | 5.2244898 | 6 | 11 | 6 | 1 | 1225 | 134.798 |
| 0 | 42.2737162 | 30.1339286 | 10 | 28 | 9 | 1 | 448 | 52.011 |
| 0.00252525 | 1.89894065 | 1.89189189 | 1 | 2 | 1 | 1 | 1110 | 125.229 |
| 0 | 10.3128489 | 2.67857143 | 2 | 4 | 2 | 1 | 896 | 100.527 |
| 0 | 100.744688 | 29.2626728 | 29 | 67 | 2 | 1 | 868 | 97.502 |
| 0 | 3.75104638 | 3.27102804 | 1 | 2 | 1 | 1 | 428 | 47.239 |
| 0 | 314.251363 | 33.0278884 | 75 | 198 | 75 | 1 | 2510 | 288.945 |
| 0.00161812 | 2.10380459 | 4.57142857 | 1 | 2 | 1 | 1 | 525 | 56.095 |
| 0 | 23.135432 | 13.7339056 | 6 | 16 | 4 | 1 | 466 | 53.619 |
| 0 | 35.684475 | 5.2148519 | 10 | 18 | 8 | 1 | 2397 | 271.519 |
| 0.00252525 | 1.89245087 | 5.43735225 | 1 | 2 | 1 | 1 | 423 | 46.705 |
| 0.01086637 | 1.33385657 | 9.85915493 | 1 | 2 | 1 | 1 | 71 | 7.845 |
| 0 | 4.26153656 | 4.34782609 | 1 | 2 | 1 | 1 | 253 | 28.005 |
| 0 | 5.14417809 | 6.46258503 | 1 | 4 | 1 | 1 | 294 | 32.921 |
| 0 | 8.74617676 | 6.64652568 | 2 | 4 | 1 | 1 | 331 | 37.383 |
| 0 | 8.74612305 | 2.88590604 | 3 | 6 | 3 | 1 | 1490 | 163.705 |
| 0.00538761 | 1.58136731 | 2.6128266 | 1 | 2 | 1 | 1 | 421 | 48.754 |

| | | | | | | | | |
|---|---|---|---|---|---|---|---|---|
| 0 | 6.01783817 | 3.04806565 | 2 | 5 | 1 | 1 | 853 | 93.176 |
| 0 | 3.00467199 | 4.94505495 | 1 | 2 | 1 | 1 | 182 | 21.203 |
| 0 | 5.6183626 | 19.205298 | 2 | 5 | 2 | 1 | 151 | 17.296 |
| 0 | 9.91166245 | 7.14285714 | 2 | 6 | 2 | 1 | 616 | 69.657 |
| 0.00133245 | 2.43004118 | 4.24242424 | 1 | 2 | 1 | 1 | 330 | 37.64 |
| 0 | 3.45481663 | 2.58215962 | 1 | 2 | 1 | 1 | 426 | 44.614 |
| 0 | 14.6874291 | 3.72294372 | 3 | 6 | 3 | 1 | 1155 | 126.138 |
| 0 | 4.23829763 | 11.299435 | 1 | 2 | 1 | 1 | 177 | 20.444 |
| 0 | 6.38740431 | 10 | 2 | 4 | 2 | 1 | 300 | 33.293 |
| 0.00538761 | 1.59550838 | 2.10772834 | 1 | 2 | 1 | 1 | 427 | 49.103 |
| 0 | 5.49430749 | 16.8539326 | 1 | 2 | 1 | 1 | 89 | 10.371 |
| 0 | 11.5067798 | 5.60131796 | 3 | 6 | 3 | 1 | 607 | 64.653 |
| 0 | 7.16619448 | 9.80392157 | 2 | 4 | 2 | 1 | 306 | 34.481 |
| 0 | 17.2576524 | 4.65753425 | 5 | 10 | 5 | 1 | 1460 | 162.147 |
| 0.00339716 | 1.75178144 | 1.09769484 | 1 | 2 | 1 | 1 | 911 | 99.056 |
| 0.00339716 | 1.77624455 | 0.80717489 | 1 | 2 | 1 | 1 | 1115 | 117.616 |
| 0 | 7.35223992 | 3.58126722 | 2 | 4 | 2 | 1 | 726 | 80.803 |
| 0 | 4.07165536 | 1.25918153 | 1 | 2 | 1 | 1 | 953 | 105.247 |
| 0.00161812 | 2.15279764 | 1.13052415 | 1 | 2 | 1 | 1 | 973 | 108.478 |
| 0 | 5.4274769 | 3.09859155 | 1 | 2 | 1 | 1 | 710 | 82.445 |
| 0 | 28.8065464 | 5.44252289 | 8 | 19 | 8 | 1 | 1966 | 213.957 |
| 0 | 4.14377595 | 2.10057288 | 2 | 4 | 2 | 1 | 1571 | 171.721 |
| 0 | 6.67840157 | 2.40153698 | 1 | 2 | 1 | 1 | 1041 | 120.011 |
| 0 | 22.4612264 | 8.60366714 | 7 | 16 | 1 | 1 | 1418 | 159.903 |
| 0.00161812 | 2.19131649 | 1.39116203 | 1 | 2 | 1 | 1 | 1222 | 136.028 |
| 0 | 5.04220123 | 3.08370044 | 1 | 4 | 1 | 1 | 681 | 75.415 |
| 0 | 3.25777212 | 3.33048676 | 2 | 3 | 2 | 1 | 1171 | 132.179 |
| 0 | 35.4455359 | 3.29326923 | 10 | 21 | 10 | 1 | 4160 | 456.149 |
| 0 | 30.8732665 | 25.8823529 | 6 | 18 | 3 | 1 | 170 | 18.367 |
| 0 | 8.53913071 | 6.63900415 | 2 | 6 | 2 | 1 | 482 | 54.088 |
| 0.007414 | 1.48069715 | 2.25988701 | 1 | 2 | 1 | 1 | 354 | 40.14 |
| 0 | 4.40882405 | 6.3583815 | 1 | 2 | 1 | 1 | 173 | 19.618 |
| 0 | 11.9732006 | 8.60832138 | 4 | 8 | 4 | 1 | 697 | 76.422 |
| 0 | 7.93125993 | 7.20338983 | 2 | 4 | 2 | 1 | 472 | 51.197 |
| 0 | 9.15188112 | 3.09597523 | 2 | 4 | 2 | 1 | 646 | 73.364 |
| 0 | 10.9255105 | 12.1621622 | 6 | 9 | 6 | 1 | 444 | 50.934 |
| 0 | 12.6303225 | 10.7334526 | 5 | 10 | 5 | 1 | 559 | 61.614 |
| 0 | 23.7109708 | 10.4740904 | 7 | 14 | 7 | 1 | 907 | 92.444 |
| 0 | 20.3324454 | 9.1522158 | 5 | 10 | 4 | 1 | 1038 | 117.59 |
| 0 | 6.79317412 | 3.7037037 | 1 | 2 | 1 | 1 | 540 | 61.079 |
| 0.00161812 | 2.08365135 | 1.78041543 | 1 | 2 | 1 | 1 | 674 | 74.03 |
| 0 | 3.2142432 | 2.66159696 | 1 | 2 | 1 | 1 | 526 | 57.945 |
| 0 | 38.5591882 | 13.25811 | 11 | 26 | 7 | 1 | 709 | 77.208 |
| 0 | 3.94005811 | 3.56265356 | 1 | 2 | 1 | 1 | 814 | 92.424 |
| 0 | 7.46237064 | 2.21205187 | 2 | 6 | 2 | 1 | 1311 | 144.601 |
| 0 | 4.64801054 | 4.74137931 | 1 | 2 | 1 | 1 | 232 | 26.737 |
| 0 | 6.83597637 | 31.6 | 3 | 6 | 3 | 1 | 250 | 27.604 |
| 0 | 4.31966449 | 1.62664601 | 1 | 4 | 1 | 1 | 1291 | 142.185 |
| 0 | 6.86206324 | 9.30232558 | 2 | 4 | 2 | 1 | 344 | 38.866 |
| 0 | 4.57554791 | 17.0984456 | 2 | 6 | 2 | 1 | 193 | 22.231 |
| 0.00161812 | 2.03395218 | 2.95566502 | 1 | 2 | 1 | 1 | 406 | 45.81 |
| 0 | 32.5670817 | 22.0207254 | 8 | 24 | 5 | 1 | 386 | 44.295 |
| 0.01798258 | 1.06565307 | 1.26748252 | 1 | 1 | 1 | 1 | 2288 | 257.162 |
| 0 | 12.8458581 | 14.4827586 | 2 | 6 | 2 | 1 | 290 | 33.469 |
| 0 | 4.31623274 | 3.53634578 | 1 | 2 | 1 | 1 | 509 | 57.082 |
| 0 | 7.11611154 | 2.81954887 | 1 | 2 | 1 | 1 | 532 | 61.175 |
| 0 | 22.559753 | 18.5393258 | 8 | 19 | 3 | 1 | 534 | 60.187 |
| 0 | 13.5352745 | 4.85148515 | 3 | 6 | 2 | 1 | 1010 | 114.409 |
| 0 | 11.1775168 | 9.71922246 | 2 | 6 | 2 | 1 | 463 | 51.535 |
| 0 | 24.2904488 | 5.93220339 | 4 | 20 | 4 | 1 | 590 | 64.036 |
| 0.0135603 | 1.26841123 | 1.07317073 | 1 | 1 | 1 | 1 | 1025 | 115.134 |
| 0.01314636 | 1.2995559 | 1.39968896 | 1 | 2 | 1 | 1 | 643 | 72.849 |

| | | | | | | | | |
|---|---|---|---|---|---|---|---|---|
| 0 | 3.16033344 | 4.00890869 | 1 | 2 | 1 | 1 | 449 | 50.602 |
| 0 | 4.29636476 | 1.53256705 | 1 | 2 | 1 | 1 | 1044 | 116.288 |
| 0 | 148.526911 | 44.2724458 | 29 | 99 | 19 | 1 | 646 | 70.854 |
| 0 | 4.93554201 | 10.5769231 | 1 | 2 | 1 | 1 | 104 | 12.067 |
| 0.0039914 | 1.71783122 | 1.2195122 | 1 | 2 | 1 | 1 | 902 | 102.591 |
| 0 | 33.1984173 | 12.2789784 | 11 | 24 | 11 | 1 | 1018 | 113.249 |
| 0 | 7.85244169 | 8.18858561 | 3 | 6 | 2 | 1 | 403 | 46.803 |
| 0 | 2.73589084 | 3.86740331 | 1 | 2 | 1 | 1 | 362 | 41.406 |
| 0 | 5.74424521 | 3.92156863 | 1 | 2 | 1 | 1 | 306 | 33.65 |
| 0 | 3.70796556 | 2.69541779 | 1 | 2 | 1 | 1 | 371 | 40.772 |
| 0 | 47.4870237 | 19.8062433 | 14 | 32 | 9 | 1 | 929 | 106.714 |
| 0 | 8.40984962 | 2.81385281 | 3 | 6 | 3 | 1 | 1386 | 159.629 |
| 0 | 28.791076 | 21.3429257 | 7 | 22 | 7 | 1 | 417 | 47.007 |
| 0 | 9.01780503 | 1.64718385 | 3 | 6 | 3 | 1 | 1882 | 218.693 |
| 0 | 3.8802175 | 1.65217391 | 2 | 4 | 1 | 1 | 1150 | 127.534 |
| 0 | 15.156984 | 6 | 3 | 10 | 3 | 1 | 500 | 56.403 |
| 0.00516403 | 1.62488532 | 10.8843537 | 1 | 2 | 1 | 1 | 147 | 15.882 |
| 0 | 2.84133602 | 5.43478261 | 1 | 2 | 1 | 1 | 184 | 20.444 |
| 0 | 6.68884753 | 8.45986985 | 2 | 4 | 2 | 1 | 461 | 50.994 |
| 0 | 3.15403424 | 2.18978102 | 1 | 2 | 1 | 1 | 411 | 46.609 |
| 0.00067889 | 2.5902359 | 1.88205772 | 1 | 2 | 1 | 1 | 797 | 88.51 |
| 0 | 4.1686768 | 1.08001964 | 2 | 4 | 2 | 1 | 2037 | 235.705 |
| 0 | 7.53699621 | 7.3943662 | 2 | 4 | 2 | 1 | 568 | 62.982 |
| 0.00067889 | 2.59636481 | 1.62074554 | 1 | 2 | 1 | 1 | 617 | 68.635 |
| 0.01730006 | 1.13212022 | 0.38554217 | 1 | 2 | 1 | 1 | 2075 | 228.827 |
| 0 | 8.53043339 | 1.8018018 | 1 | 4 | 1 | 1 | 777 | 86.156 |
| 0 | 90.5555721 | 36.4109233 | 23 | 59 | 23 | 1 | 769 | 85.081 |
| 0 | 4.41555569 | 4.72440945 | 1 | 2 | 1 | 1 | 381 | 42.955 |
| 0.00281162 | 1.80882854 | 2.03160271 | 1 | 2 | 1 | 1 | 443 | 50.177 |
| 0.00161812 | 2.10441172 | 0.35842294 | 1 | 2 | 1 | 1 | 2511 | 273.254 |
| 0 | 8.97601135 | 12.4497992 | 3 | 6 | 3 | 1 | 249 | 27.501 |
| 0 | 9.17727043 | 4.81632653 | 4 | 8 | 4 | 1 | 1225 | 136.405 |
| 0 | 26.8575255 | 23.5474006 | 7 | 19 | 7 | 1 | 327 | 36.483 |
| 0.00538761 | 1.60607399 | 0.98039216 | 1 | 2 | 1 | 1 | 1224 | 135.485 |
| 0 | 4.24443002 | 5.01930502 | 1 | 2 | 1 | 1 | 259 | 28.518 |
| 0 | 2.91937351 | 1.64158687 | 1 | 2 | 1 | 1 | 731 | 80.057 |
| 0.00856974 | 1.36723911 | 1.2987013 | 1 | 2 | 1 | 1 | 924 | 97.056 |
| 0 | 3.16811264 | 1.3400335 | 1 | 4 | 1 | 1 | 597 | 67.243 |
| 0 | 4.22083658 | 2.69749518 | 1 | 2 | 1 | 1 | 519 | 51.992 |
| 0 | 3.03990993 | 3.2 | 1 | 2 | 1 | 1 | 375 | 41.205 |
| 0 | 7.78683508 | 1.89274448 | 2 | 6 | 2 | 1 | 1585 | 173.984 |
| 0.00133245 | 2.36845477 | 2.16346154 | 1 | 4 | 1 | 1 | 416 | 45.875 |
| 0.00133245 | 2.29576366 | 6.27802691 | 1 | 2 | 1 | 1 | 223 | 24.841 |
| 0.00161812 | 2.04556451 | 1.25858124 | 1 | 2 | 1 | 1 | 874 | 97.526 |
| 0 | 6.6383122 | 5.12195122 | 2 | 3 | 2 | 1 | 410 | 43.958 |
| 0 | 7.08738176 | 5.05747126 | 3 | 5 | 3 | 1 | 435 | 48.361 |
| 0 | 32.3914993 | 12.7071823 | 10 | 20 | 10 | 1 | 1086 | 120.85 |
| 0 | 4.70744056 | 3.1147541 | 2 | 4 | 2 | 1 | 610 | 71.32 |
| 0 | 18.1938432 | 13.878327 | 5 | 13 | 1 | 1 | 526 | 59.924 |
| 0 | 6.72420301 | 10.2678571 | 2 | 4 | 2 | 1 | 224 | 25.438 |
| 0 | 26.0916174 | 43.6548223 | 8 | 28 | 8 | 1 | 197 | 22.173 |
| 0 | 20.4928875 | 16.4009112 | 5 | 13 | 1 | 1 | 439 | 50.791 |
| 0.00457317 | 1.64897715 | 14.4736842 | 1 | 2 | 1 | 1 | 152 | 17.712 |
| 0 | 20.7744036 | 11.9565217 | 4 | 18 | 4 | 1 | 460 | 52.27 |
| 0 | 2.71669877 | 7.77202073 | 1 | 2 | 1 | 1 | 193 | 21.754 |
| 0 | 3.23498111 | 8.18181818 | 2 | 3 | 2 | 1 | 110 | 12.051 |
| 0 | 10.2636124 | 4.22535211 | 2 | 6 | 2 | 1 | 710 | 79.787 |
| 0 | 21.5297357 | 17.2804533 | 5 | 15 | 5 | 1 | 353 | 38.593 |

| calc. pI | Abundance Ratio: (F3, 127) / (F3, 126) | Abundance Ratio: (F3, 128) / (F3, 126) | Abundance Ratio: (F3, 129) / (F3, 126) |
|---|---|---|---|
| 7.8 | 0.846 | 0.867 | 0.864 |
| 7.61 | 0.825 | 0.873 | 0.778 |
| 5.39 | 0.639 | 0.746 | 0.838 |
| 7.64 | | | |
| 6.98 | 0.722 | 0.757 | 0.692 |
| 5.05 | | | |
| 6.8 | 0.756 | 0.769 | 0.807 |
| 6.79 | 0.796 | 0.723 | 0.743 |
| 5.66 | 0.797 | 0.802 | 0.683 |
| 8.53 | 0.806 | 0.781 | 0.71 |
| 9.96 | 0.986 | 0.94 | 0.767 |
| 7.46 | 0.723 | 0.847 | 0.802 |
| 8 | 0.72 | 0.797 | 0.765 |
| 6.68 | 1.228 | 0.97 | 0.956 |
| 7.21 | 0.712 | 0.801 | 0.743 |
| 10.27 | | | |
| 5.14 | 0.853 | 0.864 | 0.784 |
| 8.72 | | | |
| 5.14 | 0.763 | 0.838 | 0.723 |
| 7.74 | 0.839 | 0.756 | 0.847 |
| 6.38 | | | |
| 6.49 | 0.963 | 0.914 | 0.804 |
| 9.01 | 0.804 | 0.603 | 0.685 |
| 9.03 | 0.807 | 0.877 | 0.817 |
| 4.67 | | | |
| 6.67 | 0.973 | 0.822 | 0.691 |
| 6.83 | 0.89 | 0.858 | 0.77 |
| 5.48 | 0.765 | 0.851 | 0.926 |
| 7.39 | 0.704 | 0.738 | 0.704 |
| 8.38 | 0.922 | 0.802 | 0.61 |
| 6.9 | | | |
| 5.14 | 0.705 | 0.776 | 0.761 |
| 5.54 | 0.795 | 0.821 | 0.721 |
| 7.01 | 0.767 | 0.823 | 0.726 |
| 6.32 | 0.717 | 0.773 | 0.788 |
| 10.9 | 1.2 | 1.053 | 0.885 |
| 10.58 | 0.663 | 0.614 | 0.822 |
| 11.36 | 0.933 | 0.77 | 0.803 |
| 8.68 | 1.455 | 0.859 | 0.908 |
| 7.08 | 0.697 | 0.611 | 0.653 |
| 8.92 | 1.02 | 0.903 | 0.871 |
| 7.88 | | | |
| 9.38 | 0.773 | 0.794 | 0.742 |
| 10.3 | 0.81 | 0.774 | 0.783 |
| 5.17 | | | |
| 7.3 | | | |
| 5.92 | 0.667 | 0.718 | 0.685 |
| 7.53 | 0.763 | 0.698 | 0.727 |
| 7.18 | 0.965 | 0.862 | 0.764 |
| 8.1 | 0.845 | 0.843 | 0.778 |
| 4.91 | 0.871 | 0.78 | 0.722 |
| 5.54 | 0.754 | 0.771 | 0.747 |
| 4.81 | 0.673 | 0.711 | 0.767 |
| 7.43 | | | |
| 5.33 | | | |
| 5.95 | 1.027 | 0.827 | 0.764 |
| 7.65 | | | |

| | | | |
|---|---|---|---|
| 6.54 | 0.698 | 0.737 | 0.723 |
| 5.03 | 0.755 | 0.775 | 0.76 |
| 8.13 | | | |
| 9.42 | | | |
| 10.51 | | | |
| 10.1 | 0.799 | 0.703 | 0.618 |
| 10.18 | | | |
| 10.61 | | | |
| 11.03 | 1.17 | 0.98 | 0.841 |
| 4.54 | | | |
| 10.54 | 0.791 | 0.683 | 0.677 |
| 10.21 | | | |
| 9.85 | 0.848 | 0.891 | 0.819 |
| 10.32 | 1.168 | 0.912 | 0.747 |
| 10.11 | | | |
| 9.73 | 0.901 | 0.88 | 0.808 |
| 10.15 | 0.853 | 0.848 | 0.645 |
| 10.1 | 0.82 | 0.645 | 0.727 |
| 10.32 | 0.989 | 0.827 | 0.796 |
| 5.43 | | | |
| 7.24 | 0.934 | 0.821 | 0.824 |
| 6.8 | | | |
| 4.84 | 0.861 | 0.875 | 0.825 |
| 4.89 | 0.653 | 0.659 | 0.622 |
| 6.57 | 0.873 | 0.832 | 0.718 |
| 8.54 | 0.721 | 0.82 | 0.761 |
| 7.05 | 0.803 | 0.796 | 0.787 |
| 4.83 | 0.77 | 0.778 | 0.804 |
| 4.84 | 0.746 | 0.836 | 0.787 |
| 8.85 | 0.537 | 0.56 | 0.61 |
| 6.27 | | | |
| 7.4 | 1.634 | 1.665 | 1.062 |
| 6.24 | | | |
| 7.85 | 0.854 | 0.957 | 0.94 |
| 7.46 | 0.755 | 0.868 | 0.964 |
| 7.24 | | | |
| 4.44 | 1.176 | 0.997 | 0.851 |
| 8.57 | | | |
| 5.16 | 1.068 | 0.775 | 0.857 |
| 5.08 | 1.047 | 0.962 | 0.866 |
| 6.8 | | | |
| 5.29 | 0.749 | 0.775 | 0.802 |
| 8.5 | 0.64 | 0.698 | 0.673 |
| 6.38 | 0.795 | 0.545 | 0.568 |
| 6.19 | 1.04 | 1.178 | 0.604 |
| 7.55 | 1.004 | 0.92 | 0.853 |
| 7.5 | 0.928 | 0.949 | 0.859 |
| 8.28 | 1.16 | 0.932 | 0.826 |
| 9.41 | | | |
| 6.21 | 0.818 | 0.746 | 0.729 |
| 10.32 | | | |
| 5.24 | 0.675 | 0.758 | 0.778 |
| 5.44 | | | |
| 7.3 | 0.946 | 0.763 | 0.702 |
| 8.72 | | | |
| 4.83 | 1.034 | 1.034 | 1.247 |
| 7.66 | | | |
| 9.36 | 0.993 | 0.728 | 0.922 |
| 8.24 | | | |
| 8.27 | | | |
| 7.21 | 1.297 | 0.957 | 0.827 |
| 7.8 | 0.576 | 0.673 | 0.578 |

| | | | |
|---|---|---|---|
| 10.84 | | | |
| 11.02 | 1.032 | 0.703 | 0.787 |
| 7.12 | 0.764 | 0.688 | 0.608 |
| 6.64 | 0.874 | 0.809 | 0.762 |
| 6.74 | 0.849 | 0.83 | 0.805 |
| 5.88 | 0.862 | 0.885 | 0.858 |
| 7.36 | 0.742 | 0.709 | 0.776 |
| 8.09 | 0.681 | 0.595 | 0.676 |
| 6.48 | 1.049 | 1.025 | 0.926 |
| 6.92 | 0.747 | 0.842 | 0.82 |
| 5.38 | 0.704 | 0.733 | 0.67 |
| 6.44 | 0.863 | 0.789 | 0.713 |
| 7.03 | | | |
| 8.12 | 0.688 | 0.754 | 0.708 |
| 6.79 | 0.858 | 0.795 | 0.797 |
| 7.69 | 0.798 | 0.862 | 0.803 |
| 4.69 | | | |
| 4.89 | 1.091 | 0.818 | 0.707 |
| 7.49 | 0.726 | 0.747 | 0.679 |
| 5.3 | | | |
| 6.37 | 0.961 | 0.75 | 0.798 |
| 5.85 | 1.187 | 0.856 | 0.784 |
| 6.46 | 0.838 | 0.765 | 0.722 |
| 7.65 | 0.782 | 0.763 | 0.724 |
| 5.17 | 0.741 | 0.792 | 0.659 |
| 7.05 | 1.004 | 0.857 | 0.822 |
| 6.55 | | | |
| 6.28 | | | |
| 4.59 | | | |
| 6.62 | 0.792 | 0.777 | 0.722 |
| 4.97 | 0.695 | 0.652 | 0.671 |
| 4.7 | | | |
| 5.21 | | | |
| 7.85 | | | |
| 5.03 | 0.823 | 0.784 | 0.743 |
| 7.31 | | | |
| 4.54 | 7.25 | 1 | 1 |
| 6.81 | 0.754 | 0.924 | 0.717 |
| 4.74 | 0.726 | 0.8 | 0.751 |
| 4.78 | 0.852 | 0.82 | 0.765 |
| 9.44 | 0.921 | 0.831 | 0.823 |
| 10.36 | 0.913 | 0.863 | 0.693 |
| 5.06 | | | |
| 9.35 | 0.925 | 0.958 | 0.792 |
| 5.26 | 0.998 | 0.873 | 0.769 |
| 4.51 | 0.891 | 0.78 | 0.763 |
| 4.64 | 1.297 | 1.072 | 0.81 |
| 9.38 | 0.93 | 0.919 | 0.792 |
| 6.47 | | | |
| 5.86 | 0.753 | 0.727 | 0.761 |
| 8.57 | 0.644 | 0.687 | 0.627 |
| 5.53 | | | |
| 5.81 | | | |
| 6.7 | | | |
| 5.14 | | | |
| 10.93 | 0.797 | 0.706 | 0.681 |
| 7.99 | | | |
| 6.62 | 1.229 | 0.733 | 0.581 |
| 5.6 | | | |
| 7.23 | 0.788 | 0.671 | 0.684 |
| 6.64 | | | |
| 6.57 | 1.049 | 0.761 | 0.832 |

| | | | |
|---|---|---|---|
| 6.61 | 0.85 | 0.708 | 0.719 |
| 5.78 | 0.709 | 0.737 | 0.661 |
| 5.1 | | | |
| 4.89 | 0.682 | 0.674 | 0.669 |
| 7.39 | | | |
| 7.97 | 0.815 | 0.717 | 0.71 |
| 5.11 | | | |
| 9.36 | 0.845 | 0.938 | 0.967 |
| 7.06 | | | |
| 7.43 | 0.706 | 0.604 | 0.679 |
| 9.09 | | | |
| 6.65 | 0.759 | 0.79 | 0.702 |
| 8.63 | 0.789 | 0.808 | 0.707 |
| 4.34 | | | |
| 7.47 | 0.7 | 0.734 | 0.775 |
| 6.67 | 1.062 | 0.992 | 0.695 |
| 6.34 | | | |
| 5.88 | 0.634 | 0.737 | 0.664 |
| 9.63 | 0.978 | 1.014 | 1 |
| 6.55 | 0.924 | 0.752 | 0.804 |
| 4.82 | | | |
| 5.53 | 0.841 | 0.749 | 0.766 |
| 5.54 | 0.818 | 0.933 | 0.808 |
| 4.78 | 0.784 | 0.778 | 0.663 |
| 5.36 | 0.77 | 0.766 | 0.736 |
| 7.81 | 1.019 | 0.88 | 0.797 |
| 4.69 | | | |
| 5.86 | 1.179 | 1.069 | 0.94 |
| 5.38 | | | |
| 5.06 | 0.889 | 0.799 | 0.841 |
| 9.57 | 0.585 | 0.714 | 0.772 |
| 5.03 | | | |
| 7.42 | 0.828 | 0.646 | 0.678 |
| 7.14 | 0.832 | 0.848 | 0.811 |
| 10.99 | | | |
| 5.44 | 0.89 | 0.805 | 0.79 |
| 6.23 | 0.851 | 0.877 | 0.843 |
| 6.58 | | | |
| 9.36 | | | |
| 5.81 | | | |
| 4.67 | | | |
| 8 | | | |
| 5.73 | 0.987 | 0.929 | 0.716 |
| 5.74 | | | |
| 4.93 | 0.812 | 0.739 | 0.694 |
| 5.36 | | | |
| 9.13 | | | |
| 8.95 | 0.661 | 0.569 | 0.631 |
| 7.81 | 0.681 | 0.754 | 0.745 |
| 8.7 | | | |
| 6.79 | 0.721 | 0.805 | 0.622 |
| 5.33 | 0.855 | 0.908 | 0.707 |
| 4.88 | 0.855 | 0.816 | 0.792 |
| 5.4 | 0.795 | 0.816 | 0.696 |
| 7.21 | | | |
| 9.72 | | | |
| 7.88 | | | |
| 11 | | | |
| 10.24 | | | |
| 5.33 | 0.854 | 0.728 | 0.726 |
| 5.43 | | | |
| 9.94 | | | |

| | | | |
|---|---|---|---|
| 9.6 | 0.841 | 0.701 | 0.709 |
| 11.62 | 0.609 | 0.601 | 0.819 |
| 11.65 | 0.73 | 0.67 | 0.653 |
| 10.65 | 0.915 | 0.685 | 0.667 |
| 4.37 | 0.648 | 0.645 | 0.508 |
| 6.04 | 1.018 | 0.957 | 0.82 |
| 5.38 | 0.84 | 0.837 | 0.763 |
| 5.16 | 1.017 | 0.891 | 0.777 |
| 11.06 | 0.752 | 0.812 | 0.738 |
| 6.99 | 0.958 | 0.961 | 0.837 |
| 5.97 | | | |
| 10.45 | 0.934 | 0.883 | 0.953 |
| 8.16 | 0.855 | 0.861 | 0.792 |
| 9.13 | | | |
| 10.84 | 1.139 | 0.94 | 0.781 |
| 5.06 | 0.693 | 0.719 | 0.709 |
| 4.65 | 0.893 | 0.809 | 0.823 |
| 5.06 | 0.877 | 0.826 | 0.732 |
| 9.32 | 0.908 | 1 | 0.792 |
| 5.49 | | | |
| 9.14 | | | |
| 7.37 | 0.684 | 0.803 | 0.685 |
| 10.49 | | | |
| 10.26 | 0.948 | 0.952 | 0.8 |
| 4.86 | 0.827 | 0.788 | 0.772 |
| 8.76 | 0.705 | 0.813 | 0.757 |
| 7.03 | 0.715 | 0.763 | 0.752 |
| 5.52 | 0.861 | 0.976 | 0.82 |
| 5.81 | 0.881 | 0.893 | 0.802 |
| 5 | 0.837 | 0.847 | 0.774 |
| 4.78 | 0.999 | 0.727 | 0.71 |
| 6.39 | 0.626 | 0.739 | 0.654 |
| 5.27 | 0.741 | 0.895 | 0.828 |
| 5.54 | | | |
| 6.89 | 0.783 | 0.702 | 0.695 |
| 6.18 | | | |
| 6.21 | | | |
| 8.95 | | | |
| 5.02 | | | |
| 5.64 | | | |
| 9.83 | 0.779 | 0.775 | 0.76 |
| 8.41 | 0.752 | 0.835 | 0.733 |
| 4.87 | | | |
| 7.02 | 0.613 | 0.447 | 0.55 |
| 8.21 | 0.731 | 0.757 | 0.701 |
| 5.03 | 0.786 | 0.748 | 0.782 |
| 5.2 | 0.993 | 1.202 | 1.055 |
| 5.8 | 0.778 | 0.809 | 0.666 |
| 6 | 0.518 | 0.62 | 0.686 |
| 9 | | | |
| 5.48 | 0.801 | 0.786 | 0.713 |
| 8.38 | 0.846 | 0.817 | 0.794 |
| 9.91 | | | |
| 6.73 | 0.853 | 0.78 | 0.668 |
| 5.49 | 0.768 | 0.799 | 0.806 |
| 5.58 | | | |
| 6.86 | | | |
| 5.11 | 0.775 | 0.783 | 0.738 |
| 5.15 | 0.66 | 0.773 | 0.733 |
| 5.97 | 0.933 | 0.897 | 0.86 |
| 6.89 | 0.918 | 0.856 | 0.758 |
| 6.35 | 1.239 | 0.998 | 0.899 |

| | | | |
|---|---|---|---|
| 5.78 | 0.811 | 1.048 | 0.874 |
| 4.89 | 0.804 | 0.79 | 0.761 |
| 5.59 | 0.786 | 0.692 | 0.709 |
| 6.25 | | | |
| 6.3 | 0.775 | 0.887 | 0.775 |
| 8.76 | | | |
| 4.53 | 0.864 | 0.664 | 0.56 |
| 8.68 | 0.769 | 0.754 | 0.686 |
| 4.68 | | | |
| 5.97 | | | |
| 8.18 | | | |
| 6.7 | | | |
| 4.92 | | | |
| 6.99 | | | |
| 4.79 | | | |
| 5.55 | 0.787 | 0.862 | 0.729 |
| 6.16 | 0.847 | 0.742 | 0.71 |
| 5.66 | 0.625 | 1.023 | 0.898 |
| 5.48 | | | |
| 7.02 | | | |
| 7.94 | | | |
| 5.06 | | | |
| 7.55 | | | |
| 5.34 | 0.929 | 0.841 | 0.891 |
| 4.54 | | | |
| 7.05 | | | |
| 7.44 | 0.492 | 0.339 | 0.466 |
| 7.02 | | | |
| 5.45 | | | |
| 4.81 | 0.76 | 0.797 | 0.75 |
| 8.69 | 0.996 | 0.694 | 0.781 |
| 7.2 | 2.039 | 1.041 | 1.315 |
| 6.48 | | | |
| 7.39 | | | |
| 4.89 | 1.443 | 0.982 | 0.82 |
| 4.89 | 0.893 | 0.774 | 0.754 |
| 5.96 | 0.923 | 0.794 | 0.808 |
| 7.74 | | | |
| 4.63 | 0.696 | 0.75 | 0.803 |
| 5.88 | 1.374 | 1.033 | 0.802 |
| 6.8 | 0.998 | 0.647 | 0.67 |
| 5.87 | 0.96 | 0.865 | 0.734 |
| 9.09 | 0.829 | 0.714 | 0.566 |
| 7.84 | 0.799 | 0.87 | 0.723 |
| 5.68 | | | |
| 5.4 | 0.833 | 0.852 | 0.771 |
| 6.7 | | | |
| 6.11 | 0.587 | 0.814 | 0.56 |
| 8.54 | 0.965 | 0.859 | 0.819 |
| 6.4 | 0.911 | 0.789 | 0.729 |
| 6.39 | | | |
| 7.06 | | | |
| 8.87 | | | |
| 5.97 | 0.676 | 0.864 | 0.831 |
| 7.11 | | | |
| 6.7 | 0.728 | 0.808 | 0.838 |
| 7.61 | 0.792 | 0.577 | 0.618 |
| 5.71 | 0.801 | 0.844 | 0.692 |
| 5.59 | 0.804 | 0.814 | 0.767 |
| 8.19 | | | |
| 5.19 | 0.82 | 0.806 | 0.775 |
| 8.32 | 0.784 | 0.842 | 0.791 |

| | | | |
|---|---|---|---|
| 7.83 | | | |
| 5.68 | 0.702 | 0.686 | 0.682 |
| 4.37 | 0.747 | 0.772 | 0.83 |
| 5.22 | | | |
| 5.86 | | | |
| 7.93 | 0.829 | 0.933 | 0.707 |
| 7.46 | | | |
| 5.76 | | | |
| 6.37 | | | |
| 6.68 | | | |
| 8.91 | | | |
| 8.12 | 0.878 | 0.874 | 0.675 |
| 6.04 | 0.996 | 0.736 | 0.745 |
| 6.37 | 0.953 | 0.894 | 0.762 |
| 4.92 | 0.793 | 0.688 | 0.687 |
| 6.48 | 0.934 | 0.928 | 0.823 |
| 5.91 | 0.765 | 0.812 | 0.678 |
| 8.15 | 0.984 | 0.811 | 0.739 |
| 9.16 | 0.953 | 0.928 | 0.772 |
| 6.71 | 0.891 | 0.689 | 0.661 |
| 5.6 | 0.865 | 0.926 | 0.794 |
| 8.31 | 0.912 | 0.822 | 0.76 |
| 8.75 | | | |
| 6.09 | 0.853 | 0.766 | 0.793 |
| 9.13 | 0.806 | 0.828 | 0.746 |
| 9.22 | | | |
| 5.49 | 0.901 | 0.913 | 0.744 |
| 8.63 | | | |
| 4.93 | 0.822 | 0.83 | 0.758 |
| 7.21 | 0.937 | 1.242 | 1.005 |
| 10.14 | 0.846 | 0.861 | 0.711 |
| 7.94 | | | |
| 4.84 | 0.786 | 0.712 | 0.642 |
| 8.22 | 0.907 | 0.851 | 0.761 |
| 5.78 | | | |
| 5.33 | 0.905 | 1.155 | 0.887 |
| 9.32 | | | |
| 7.12 | | | |
| 9.76 | 0.614 | 0.646 | 0.78 |
| 5.1 | 0.914 | 0.72 | 0.64 |
| 6.55 | | | |
| 6.95 | | | |
| 7.46 | | | |
| 7.33 | 0.878 | 1.039 | 0.861 |
| 9.13 | 1.144 | 1.015 | 0.738 |
| 8.56 | | | |
| 5.15 | 0.83 | 0.638 | 0.544 |
| 7.18 | | | |
| 6.11 | 0.921 | 0.777 | 0.75 |
| 7.2 | | | |
| 7.81 | 0.934 | 0.948 | 0.89 |
| 6.89 | | | |
| 4.81 | 1.115 | 1.1 | 0.783 |
| 7.06 | | | |
| 7.17 | 0.997 | 0.763 | 0.712 |
| 6.86 | 0.861 | 0.901 | 0.824 |
| 6.49 | 0.857 | 0.98 | 0.881 |
| 8.65 | | | |
| 8.41 | 0.851 | 0.801 | 0.743 |
| 6.98 | 0.917 | 0.868 | 0.889 |
| 8.25 | | | |
| 6.25 | | | |

| | | | |
|---|---|---|---|
| 8.98 | | | |
| 11.34 | 0.943 | 0.898 | 0.65 |
| 5.6 | | | |
| 11.82 | | | |
| 9.01 | 0.741 | 0.805 | 0.716 |
| 6.96 | 0.866 | 0.861 | 0.779 |
| 6 | | | |
| 6.89 | 0.826 | 0.675 | 0.692 |
| 8.28 | | | |
| 10.93 | | | |
| 7.4 | 0.583 | 0.466 | 0.536 |
| 5.55 | 1.041 | 0.717 | 0.747 |
| 6.46 | | | |
| 7.18 | | | |
| 7.85 | 0.858 | 0.662 | 0.618 |
| 7.55 | 0.8 | 0.658 | 0.729 |
| 4.88 | 0.901 | 0.973 | 0.773 |
| 8.91 | 0.825 | 0.84 | 0.73 |
| 9.07 | 0.659 | 0.694 | 0.598 |
| 4.92 | 0.835 | 0.752 | 0.737 |
| 5.53 | 0.966 | 0.889 | 0.692 |
| 7.44 | | | |
| 10.18 | 1.154 | 0.925 | 1.032 |
| 4.65 | 0.781 | 0.817 | 0.606 |
| 8.34 | 0.785 | 0.741 | 0.669 |
| 4.94 | 0.885 | 0.825 | 0.782 |
| 6.67 | | | |
| 8.79 | | | |
| 5.57 | 0.872 | 0.915 | 0.836 |
| 8.75 | | | |
| 4.75 | 1.02 | 0.939 | 0.859 |
| 9.38 | 0.827 | 0.695 | 0.783 |
| 8.76 | | | |
| 5.69 | 1 | 0.679 | 0.702 |
| 7.42 | 0.798 | 0.916 | 0.777 |
| 5.03 | | | |
| 5.31 | | | |
| 4.97 | | | |
| 5.86 | 0.785 | 0.623 | 0.734 |
| 6.51 | 0.843 | 0.782 | 0.772 |
| 8.27 | | | |
| 5.26 | | | |
| 6.37 | | | |
| 5.03 | 0.713 | 0.468 | 0.507 |
| 7.01 | | | |
| 4.45 | | | |
| 5.16 | 0.872 | 0.752 | 0.725 |
| 4.81 | 0.923 | 0.832 | 0.717 |
| 5.15 | 0.835 | 0.938 | 0.577 |
| 6.48 | | | |
| 5.31 | 0.849 | 0.787 | 0.747 |
| 9.69 | 0.668 | 0.69 | 0.699 |
| 7.77 | 0.934 | 1.142 | 0.744 |
| 7.06 | 0.954 | 0.856 | 0.647 |
| 5.35 | 0.71 | 0.797 | 0.698 |
| 7.39 | 0.751 | 0.909 | 0.796 |
| 6.65 | 0.886 | 0.796 | 0.663 |
| 7.2 | 0.943 | 0.913 | 0.771 |
| 4.82 | 0.741 | 0.838 | 0.796 |
| 7.75 | | | |
| 8.02 | 0.806 | 0.713 | 0.709 |
| 4.91 | 0.838 | 0.814 | 0.648 |

| | | | |
|---|---|---|---|
| 5.72 | 0.79 | 0.863 | 0.809 |
| 8.32 | 0.802 | 0.754 | 0.672 |
| 8.54 | | | |
| 7.99 | | | |
| 8.88 | 0.85 | 0.757 | 0.851 |
| 6.42 | 0.855 | 0.883 | 0.83 |
| 4.49 | | | |
| 9.13 | | | |
| 11.27 | | | |
| 9.99 | | | |
| 9.22 | | | |
| 7.09 | 1.02 | 0.971 | 0.79 |
| 5.94 | | | |
| 6.79 | 0.935 | 0.873 | 0.807 |
| 8.35 | 0.797 | 0.696 | 0.505 |
| 8.47 | 0.892 | 1.008 | 0.811 |
| 8.98 | | | |
| 8.43 | 0.703 | 0.526 | 0.614 |
| 8.43 | 0.895 | 0.917 | 0.789 |
| 7.61 | 0.803 | 0.479 | 0.65 |
| 6.42 | | | |
| 6.87 | | | |
| 6.73 | 0.694 | 0.594 | 0.741 |
| 8.66 | 0.771 | 0.918 | 0.777 |
| 7.37 | 0.789 | 0.866 | 0.657 |
| 10.15 | | | |
| 8.69 | | | |
| 5.57 | 0.985 | 0.897 | 0.829 |
| 5.6 | | | |
| 7.01 | 0.477 | 0.736 | 0.706 |
| 8.34 | 0.826 | 0.748 | 0.745 |
| 9.83 | 0.963 | 0.955 | 0.843 |
| 7.91 | 1.141 | 0.972 | 0.896 |
| 8.06 | | | |
| 7.09 | | | |
| 7.46 | | | |
| 7.02 | 1.125 | 0.891 | 0.832 |
| 4.78 | 0.797 | 0.778 | 0.721 |
| 7.77 | 0.835 | 0.794 | 0.73 |
| 6.9 | | | |
| 9.32 | | | |
| 5.92 | 0.854 | 0.834 | 0.803 |
| 8.22 | 0.792 | 0.823 | 0.749 |
| 6.52 | | | |
| 8.38 | 1.081 | 0.593 | 0.711 |
| 5.71 | | | |
| 8.43 | 0.713 | 0.771 | 0.769 |
| 5.06 | | | |
| 5.97 | | | |
| 8.98 | | | |
| 7.69 | 0.754 | 0.721 | 0.733 |
| 6.52 | 0.768 | 0.645 | 0.694 |
| 5.83 | 0.742 | 0.743 | 0.637 |
| 6.05 | 0.842 | 0.853 | 0.719 |
| 6.43 | | | |
| 6.64 | 0.774 | 0.719 | 0.649 |
| 7.59 | 0.753 | 0.862 | 0.756 |
| 6.35 | 0.932 | 0.766 | 0.802 |
| 8.59 | 0.857 | 0.783 | 0.765 |
| 10.1 | 0.873 | 1.074 | 0.725 |
| 6.37 | | | |
| 8.72 | | | |

| | | | |
|---|---|---|---|
| 5.48 | 0.823 | 0.955 | 0.885 |
| 6.64 | 0.832 | 0.921 | 0.789 |
| 9.29 | | | |
| 7.87 | 0.75 | 0.818 | 0.724 |
| 5.12 | 0.841 | 0.832 | 0.655 |
| 5.67 | 0.799 | 0.811 | 0.735 |
| 4.93 | 1.236 | 0.953 | 0.834 |
| 6.86 | 0.893 | 0.816 | 0.779 |
| 8.25 | 1.04 | 0.972 | 0.833 |
| 8.66 | | | |
| 6.73 | | | |
| 8.97 | 0.999 | 0.912 | 0.825 |
| 5.38 | 0.924 | 0.902 | 0.759 |
| 8.75 | | | |
| 5.73 | | | |
| 7.47 | | | |
| 8.51 | | | |
| 7.52 | 0.895 | 0.821 | 0.735 |
| 9.73 | | | |
| 5.94 | 0.873 | 0.948 | 0.734 |
| 5.85 | | | |
| 6.79 | 0.857 | 0.777 | 0.801 |
| 8.44 | | | |
| 7.08 | 0.879 | 0.733 | 0.731 |
| 8.59 | | | |
| 8.75 | | | |
| 5.91 | | | |
| 5.76 | 0.901 | 0.843 | 0.818 |
| 6.79 | 0.955 | 0.821 | 0.687 |
| 9.44 | 0.838 | 0.757 | 0.769 |
| 6.05 | | | |
| 8.21 | 0.812 | 0.78 | 0.713 |
| 5.69 | | | |
| 5.45 | 1.009 | 0.963 | 0.799 |
| 6.02 | | | |
| 7.87 | | | |
| 9.01 | 0.656 | 0.589 | 0.712 |
| 5.53 | | | |
| 5.07 | 0.789 | 0.821 | 0.708 |
| 7.64 | 0.954 | 0.962 | 0.893 |
| 5.95 | | | |
| 7.27 | 0.674 | 0.598 | 0.548 |
| 6.71 | | | |
| 7.55 | 0.675 | 0.675 | 0.753 |
| 5.17 | | | |
| 5.08 | 0.886 | 0.844 | 0.78 |
| 10.32 | 0.877 | 0.755 | 0.774 |
| 5.4 | | | |
| 6.25 | | | |
| 5.06 | | | |
| 9.5 | 0.927 | 0.842 | 0.912 |
| 5.97 | | | |
| 4.79 | 0.799 | 0.789 | 0.762 |
| 7.34 | | | |
| 8.6 | | | |
| 9.38 | 0.785 | 0.683 | 0.69 |
| 6.6 | | | |
| 7.11 | 0.566 | 0.717 | 0.515 |
| 7.17 | 0.732 | 0.638 | 0.681 |
| 6.98 | 0.833 | 0.884 | 0.815 |
| 5.71 | 0.959 | 0.935 | 0.774 |
| 7.97 | | | |

| | | | |
|---|---|---|---|
| 6.34 | 0.829 | 1.21 | 0.906 |
| 7.24 | 0.838 | 0.831 | 0.709 |
| 7.64 | 0.846 | 0.83 | 0.928 |
| 9.5 | | | |
| 6 | 0.836 | 0.794 | 0.759 |
| 8.12 | 0.893 | 0.786 | 0.744 |
| 6.42 | | | |
| 8.66 | 0.81 | 0.751 | 0.751 |
| 6.34 | | | |
| 4.72 | | | |
| 4.83 | 1.176 | 1.111 | 0.937 |
| 4.79 | | | |
| 7.44 | 0.85 | 0.82 | 0.619 |
| 8.13 | 0.811 | 0.764 | 0.688 |
| 5.34 | 0.709 | 0.8 | 0.767 |
| 6.87 | 0.824 | 0.8 | 0.766 |
| 5.69 | 0.764 | 0.795 | 0.786 |
| 6.65 | 0.914 | 0.785 | 0.795 |
| 6.43 | 0.818 | 0.848 | 0.762 |
| 9.22 | 0.91 | 0.835 | 0.776 |
| 6.93 | | | |
| 10.08 | 0.836 | 0.871 | 0.818 |
| 6.7 | | | |
| 6.67 | | | |
| 8.31 | 0.919 | 0.9 | 0.811 |
| 8.46 | 0.819 | 0.791 | 0.694 |
| 6.77 | 0.629 | 0.658 | 0.722 |
| 7.01 | | | |
| 6.95 | 0.789 | 0.915 | 0.732 |
| 7.74 | 0.839 | 0.754 | 0.755 |
| 5.33 | | | |
| 5.47 | 0.745 | 0.771 | 0.692 |
| 8.95 | | | |
| 5.72 | | | |
| 4.88 | 0.697 | 0.712 | 0.701 |
| 8.46 | 0.85 | 0.809 | 0.74 |
| 6.21 | | | |
| 7.06 | 1.423 | 0.96 | 0.945 |
| 8.51 | 0.948 | 0.921 | 0.79 |
| 6.01 | | | |
| 7.17 | 0.919 | 0.878 | 0.728 |
| 6.8 | | | |
| 7.37 | | | |
| 9.36 | 0.621 | 0.66 | 0.813 |
| 4.7 | 0.834 | 0.834 | 0.711 |
| 4.55 | 0.636 | 0.77 | 0.822 |
| 8.73 | 0.924 | 0.876 | 0.777 |
| 5.36 | | | |
| 5.4 | 0.928 | 0.96 | 0.868 |
| 7.01 | 0.702 | 0.585 | 0.635 |
| 5.68 | 1.179 | 0.888 | 0.85 |
| 5.41 | 0.825 | 0.868 | 0.802 |
| 8.6 | | | |
| 5.12 | 0.896 | 0.919 | 0.694 |
| 6.23 | 0.754 | 0.748 | 0.813 |
| 9 | | | |
| 7.99 | | | |
| 8.81 | 0.601 | 0.689 | 0.718 |
| 7.88 | 0.994 | 0.837 | 0.87 |
| 5.91 | | | |
| 6.92 | 0.762 | 0.788 | 0.74 |
| 5.91 | | | |

| | | | |
|---|---|---|---|
| 8.16 | | | |
| 9.01 | 0.762 | 0.794 | 0.773 |
| 4.89 | 0.698 | 0.669 | 0.705 |
| 5.67 | 0.839 | 0.708 | 0.784 |
| 6.42 | | | |
| 8.46 | 0.571 | 0.81 | 0.612 |
| 8.18 | 0.835 | 0.756 | 0.672 |
| 4.75 | | | |
| 4.87 | 0.741 | 0.843 | 0.716 |
| 8.46 | | | |
| 7.21 | 0.839 | 0.882 | 0.965 |
| 6.87 | 0.875 | 0.946 | 0.789 |
| 5.96 | 0.876 | 1.108 | 1.034 |
| 7.15 | 0.804 | 0.816 | 0.741 |
| 4.64 | | | |
| 7.12 | | | |
| 5.92 | 0.982 | 0.88 | 0.832 |
| 5.71 | | | |
| 5 | 0.676 | 0.815 | 0.795 |
| 5.57 | | | |
| 4.92 | 0.879 | 0.789 | 0.704 |
| 8.47 | | | |
| 7.11 | | | |
| 5.9 | 0.753 | 0.721 | 0.67 |
| 8.47 | | | |
| 5.36 | | | |
| 6.74 | | | |
| 5.12 | 0.771 | 0.82 | 0.677 |
| 10.01 | 1.076 | 1.038 | 0.899 |
| 5.55 | 0.841 | 0.751 | 0.707 |
| 7.42 | | | |
| 5.52 | 0.701 | 0.648 | 0.663 |
| 4.58 | 0.775 | 0.852 | 0.862 |
| 6.8 | 0.793 | 0.628 | 0.682 |
| 4.65 | 0.787 | 0.963 | 0.807 |
| 5.24 | 1.127 | 0.993 | 0.83 |
| 6.46 | 0.924 | 0.937 | 0.782 |
| 4.86 | 0.947 | 0.753 | 0.8 |
| 6.92 | 0.875 | 0.93 | 0.781 |
| 6.18 | | | |
| 7.85 | | | |
| 5.31 | | | |
| 7.2 | 0.57 | 0.437 | 0.446 |
| 5 | | | |
| 5.71 | 1.017 | 0.942 | 0.864 |
| 7.75 | 0.706 | 0.789 | 0.678 |
| 9.07 | 1.165 | 0.823 | 0.823 |
| 6.67 | 0.888 | 0.655 | 0.73 |
| 6.38 | 0.829 | 0.864 | 0.692 |
| 5.35 | 0.695 | 0.713 | 0.675 |
| 5.67 | | | |
| 5.48 | 0.774 | 0.859 | 0.841 |
| 7.18 | | | |
| 4.32 | 0.653 | 0.4 | 0.8 |
| 6.25 | 0.78 | 0.763 | 0.73 |
| 9.38 | | | |
| 6.1 | 0.831 | 0.817 | 0.71 |
| 6.65 | 0.948 | 0.843 | 0.813 |
| 5.43 | 1.008 | 0.922 | 0.916 |
| 6.87 | 0.996 | 0.869 | 0.774 |
| 6.33 | | | |
| 7.33 | | | |

| | | | |
|---|---|---|---|
| 6.67 | 0.729 | 0.799 | 0.708 |
| 5.38 | | | |
| 5.52 | 0.859 | 0.807 | 0.731 |
| 11 | | | |
| 6.71 | | | |
| 5.9 | 0.806 | 0.832 | 0.73 |
| 4.46 | | | |
| 6.07 | 0.734 | 0.691 | 0.66 |
| 5.08 | 0.806 | 0.72 | 0.743 |
| 5.21 | 0.71 | 0.729 | 0.645 |
| 6.05 | 0.983 | 0.916 | 0.921 |
| 6.98 | | | |
| 8.34 | 0.938 | 0.966 | 0.733 |
| 8.47 | 0.776 | 0.736 | 0.761 |
| 7.08 | | | |
| 7.96 | 1.364 | 1 | 1.205 |
| 8.21 | | | |
| 8.46 | | | |
| 6.68 | 1.338 | 1.156 | 1.078 |
| 5.06 | | | |
| 6.1 | | | |
| 5.6 | | | |
| 9.13 | | | |
| 8.97 | | | |
| 8.31 | 0.863 | 0.878 | 0.721 |
| 7.03 | 0.705 | 0.509 | 0.58 |
| 7.62 | 0.775 | 0.641 | 0.631 |
| 5.35 | | | |
| 7.81 | | | |
| 6.44 | | | |
| 5.01 | 0.98 | 0.91 | 0.787 |
| 5.3 | 0.914 | 0.8 | 0.775 |
| 5.67 | 0.874 | 0.791 | 0.749 |
| 5.5 | | | |
| 6.24 | | | |
| 5.54 | | | |
| 5.95 | | | |
| 8.97 | 0.725 | 0.755 | 0.653 |
| 7.72 | | | |
| 5.21 | | | |
| 5.53 | 0.96 | 0.788 | 0.816 |
| 5.78 | | | |
| 8.98 | 0.776 | 0.751 | 0.669 |
| 5.55 | 0.983 | 0.876 | 0.783 |
| 9.22 | 0.809 | 0.884 | 0.754 |
| 8.54 | 0.779 | 0.736 | 0.688 |
| 5.71 | 0.86 | 0.706 | 0.715 |
| 6.06 | | | |
| 7.37 | 0.91 | 0.878 | 0.824 |
| 7.83 | 0.749 | 0.918 | 0.829 |
| 4.63 | 0.785 | 0.838 | 0.866 |
| 7.01 | 0.828 | 0.916 | 0.645 |
| 10.39 | 1.149 | 1.426 | 1.277 |
| 9.5 | 0.774 | 0.764 | 0.709 |
| 6.1 | | | |
| 5.19 | 1.183 | 1.01 | 0.911 |
| 7.55 | 0.736 | 0.583 | 0.538 |
| 7.3 | 1.037 | 0.911 | 0.767 |

| Abundance: F3: 126, Control | Abundance: F3: 127, Sample | Abundance: F3: 128, Sample | Abundance: F3: 129, Sample | Abundances Count: F3: 126, Control | Abundances Count: F3: 127, Sample | Abundances Count: F3: 128, Sample | Abundances Count: F3: 129, Sample | emPAI |
|---|---|---|---|---|---|---|---|---|
| 77.3 | 65.4 | 67 | 66.8 | 1 | 1 | 1 | 1 | 0.129 |
| 659.4 | 544.3 | 575.4 | 512.9 | 7 | 7 | 7 | 7 | 1.482 |
| 172.1 | 109.9 | 128.4 | 144.3 | 3 | 3 | 3 | 3 | 55.234 |
| | | | | | | | | 0.119 |
| 26.3 | 19 | 19.9 | 18.2 | 1 | 1 | 1 | 1 | 0.136 |
| | | | | | | | | 0.11 |
| 74.5 | 56.3 | 57.3 | 60.1 | 2 | 2 | 2 | 2 | 2.162 |
| 149.6 | 119.1 | 108.2 | 111.1 | 1 | 1 | 1 | 1 | 0.233 |
| 1020.2 | 812.6 | 818.7 | 696.9 | 9 | 9 | 9 | 9 | 3.095 |
| 1101.6 | 887.9 | 860.8 | 781.9 | 9 | 9 | 9 | 9 | 6.499 |
| 115.2 | 113.6 | 108.3 | 88.4 | 1 | 1 | 1 | 1 | 0.212 |
| 131.9 | 95.3 | 111.7 | 105.8 | 1 | 1 | 1 | 1 | 0.389 |
| 582.3 | 419.3 | 463.9 | 445.6 | 7 | 7 | 7 | 7 | 9 |
| 50.1 | 61.5 | 48.6 | 47.9 | 1 | 1 | 1 | 1 | 0.086 |
| 284.6 | 202.6 | 228 | 211.4 | 2 | 2 | 2 | 2 | 0.778 |
| | | | | | | | | 0.585 |
| 118.2 | 100.8 | 102.1 | 92.7 | 1 | 1 | 1 | 1 | 0.089 |
| | | | | | | | | 0.389 |
| 151 | 115.2 | 126.6 | 109.1 | 1 | 1 | 1 | 1 | 0.166 |
| 96.9 | 81.3 | 73.3 | 82.1 | 2 | 2 | 2 | 2 | 0.468 |
| | | | | | | | | 9.771 |
| 664.7 | 640.4 | 607.4 | 534.3 | 4 | 4 | 4 | 4 | 1.102 |
| 179.6 | 144.4 | 108.3 | 123.1 | 2 | 2 | 2 | 2 | 2.728 |
| 82.4 | 66.5 | 72.3 | 67.3 | 3 | 3 | 3 | 3 | 2.728 |
| | | | | | | | | 0.194 |
| 142.3 | 138.5 | 116.9 | 98.3 | 2 | 2 | 2 | 2 | 0.304 |
| 62.6 | 55.7 | 53.7 | 48.2 | 1 | 1 | 1 | 1 | 0.216 |
| 84.1 | 64.3 | 71.6 | 77.9 | 1 | 1 | 1 | 1 | 0.65 |
| 731.4 | 514.6 | 539.5 | 514.8 | 8 | 8 | 8 | 8 | 5.579 |
| 33.3 | 30.7 | 26.7 | 20.3 | 1 | 1 | 1 | 1 | 0.413 |
| | | | | | | | | 0.101 |
| 350 | 246.8 | 271.5 | 266.4 | 4 | 4 | 4 | 4 | 3.365 |
| 124.9 | 99.3 | 102.5 | 90 | 2 | 2 | 2 | 2 | 0.668 |
| 454.1 | 348.4 | 373.8 | 329.7 | 5 | 5 | 5 | 5 | 0.823 |
| 145.1 | 104 | 112.2 | 114.4 | 1 | 1 | 1 | 1 | 0.179 |
| 93.6 | 112.3 | 98.6 | 82.8 | 3 | 3 | 3 | 3 | 14.849 |
| 10.1 | 6.7 | 6.2 | 8.3 | 1 | 1 | 1 | 1 | 1.512 |
| 136.3 | 127.1 | 105 | 109.5 | 3 | 3 | 3 | 3 | 30.623 |
| 372.7 | 542.1 | 320.3 | 338.4 | 3 | 3 | 3 | 3 | 4.623 |
| 54.7 | 38.1 | 33.4 | 35.7 | 1 | 1 | 1 | 1 | 0.274 |
| 459.4 | 468.4 | 415 | 400 | 5 | 5 | 5 | 5 | 6.499 |
| | | | | | | | | 0.778 |
| 253.3 | 195.8 | 201.2 | 188 | 1 | 1 | 1 | 1 | 0.468 |
| 126 | 102 | 97.5 | 98.7 | 1 | 1 | 1 | 1 | 0.292 |
| | | | | | | | | 2.162 |
| | | | | | | | | 0.056 |
| 48.6 | 32.4 | 34.9 | 33.3 | 1 | 1 | 1 | 1 | 0.259 |
| 330.1 | 251.8 | 230.4 | 239.9 | 2 | 2 | 2 | 2 | 0.874 |
| 535.4 | 516.6 | 461.4 | 408.8 | 10 | 10 | 10 | 10 | 13.251 |
| 766.7 | 647.7 | 646.7 | 596.7 | 8 | 8 | 8 | 8 | 3.786 |
| 89.1 | 77.6 | 69.5 | 64.3 | 1 | 1 | 1 | 1 | 0.116 |
| 183.2 | 138.1 | 141.3 | 136.8 | 1 | 1 | 1 | 1 | 0.145 |
| 130.7 | 87.9 | 92.9 | 100.2 | 3 | 3 | 3 | 3 | 1.783 |
| | | | | | | | | 0.259 |
| | | | | | | | | 0.212 |
| 76.4 | 78.5 | 63.2 | 58.4 | 1 | 1 | 1 | 1 | 0.304 |
| | | | | | | | | 0.096 |

| | | | | | | | | |
|---|---|---|---|---|---|---|---|---|
| 330.1 | 230.5 | 243.3 | 238.7 | 3 | 3 | 3 | 3 | 1.276 |
| 622 | 469.8 | 482.1 | 472.9 | 5 | 5 | 5 | 5 | 13.251 |
| | | | | | | | | 0.52 |
| | | | | | | | | 0.292 |
| | | | | | | | | 0.292 |
| 54.2 | 43.3 | 38.1 | 33.5 | 1 | 1 | 1 | 1 | 3.642 |
| | | | | | | | | 0.122 |
| | | | | | | | | 0.259 |
| 59.9 | 70.1 | 58.7 | 50.4 | 1 | 1 | 1 | 1 | 0.233 |
| | | | | | | | | 1.154 |
| 104.4 | 82.6 | 71.3 | 70.7 | 1 | 1 | 1 | 1 | 0.292 |
| | | | | | | | | 0.874 |
| 13.8 | 11.7 | 12.3 | 11.3 | 1 | 1 | 1 | 1 | 0.585 |
| 48.7 | 56.9 | 44.4 | 36.4 | 1 | 1 | 1 | 1 | 1.512 |
| | | | | | | | | 0.585 |
| 115.9 | 104.4 | 102 | 93.6 | 1 | 1 | 1 | 1 | 0.311 |
| 40.9 | 34.9 | 34.7 | 26.4 | 1 | 1 | 1 | 1 | 0.359 |
| 124 | 101.7 | 80 | 90.2 | 1 | 1 | 1 | 1 | 0.233 |
| 122.1 | 120.7 | 101 | 97.2 | 2 | 2 | 2 | 2 | 0.995 |
| | | | | | | | | 0.585 |
| 215.7 | 201.5 | 177 | 177.8 | 3 | 3 | 3 | 3 | 0.638 |
| | | | | | | | | 0.089 |
| 306.1 | 263.4 | 267.7 | 252.4 | 3 | 3 | 3 | 3 | 0.468 |
| 340.3 | 222.3 | 224.1 | 211.6 | 2 | 2 | 2 | 2 | 576.969 |
| 126.5 | 110.4 | 105.3 | 90.8 | 2 | 2 | 2 | 2 | 0.995 |
| 775.1 | 558.9 | 635.6 | 589.7 | 7 | 7 | 7 | 7 | 3.437 |
| 107.4 | 86.2 | 85.5 | 84.5 | 1 | 1 | 1 | 1 | 0.066 |
| 208.6 | 160.7 | 162.3 | 167.8 | 4 | 4 | 4 | 4 | 6.197 |
| 552.7 | 412.2 | 461.9 | 434.9 | 7 | 7 | 7 | 7 | 3.87 |
| 125.4 | 67.4 | 70.2 | 76.5 | 1 | 1 | 1 | 1 | 0.096 |
| | | | | | | | | 0.093 |
| 16.1 | 26.3 | 26.8 | 17.1 | 2 | 2 | 2 | 2 | 0.585 |
| | | | | | | | | 0.038 |
| 23.3 | 19.9 | 22.3 | 21.9 | 1 | 1 | 1 | 1 | 0.089 |
| 47.7 | 36 | 41.4 | 46 | 1 | 1 | 1 | 1 | 0.28 |
| | | | | | | | | 0.233 |
| 37.5 | 44.1 | 37.4 | 31.9 | 2 | 2 | 2 | 2 | 0.245 |
| | | | | | | | | 0.233 |
| 88.6 | 94.6 | 68.7 | 75.9 | 1 | 1 | 1 | 1 | 0.468 |
| 66.6 | 69.7 | 64.1 | 57.7 | 1 | 1 | 1 | 1 | 0.11 |
| | | | | | | | | 0.995 |
| 78.8 | 59 | 61.1 | 63.2 | 1 | 1 | 1 | 1 | 0.778 |
| 36.1 | 23.1 | 25.2 | 24.3 | 1 | 1 | 1 | 1 | 0.311 |
| 4.4 | 3.5 | 2.4 | 2.5 | 1 | 1 | 1 | 1 | 0.359 |
| 10.1 | 10.5 | 11.9 | 6.1 | 1 | 1 | 1 | 1 | 0.084 |
| 92.4 | 92.8 | 85 | 78.8 | 1 | 1 | 1 | 1 | 0.096 |
| 218.4 | 202.6 | 207.2 | 187.5 | 1 | 1 | 1 | 1 | 0.501 |
| 105.8 | 122.7 | 98.6 | 87.4 | 1 | 1 | 1 | 1 | 0.334 |
| | | | | | | | | 0.194 |
| 116.8 | 95.5 | 87.1 | 85.1 | 2 | 2 | 2 | 2 | 0.848 |
| | | | | | | | | 0.468 |
| 371.9 | 251 | 281.8 | 289.3 | 5 | 5 | 5 | 5 | 2.728 |
| | | | | | | | | 0.233 |
| 29.9 | 28.3 | 22.8 | 21 | 1 | 1 | 1 | 1 | 0.638 |
| | | | | | | | | 0.212 |
| 8.9 | 9.2 | 9.2 | 11.1 | 1 | 1 | 1 | 1 | 0.585 |
| | | | | | | | | 0.145 |
| 28.3 | 28.1 | 20.6 | 26.1 | 1 | 1 | 1 | 1 | 0.389 |
| | | | | | | | | 0.259 |
| | | | | | | | | 0.292 |
| 76.8 | 99.6 | 73.5 | 63.5 | 1 | 1 | 1 | 1 | 0.212 |
| 45.3 | 26.1 | 30.5 | 26.2 | 1 | 1 | 1 | 1 | 1.548 |

| | | | | | | | | |
|---|---|---|---|---|---|---|---|---|
| | | | | | | | | 1.154 |
| 476 | 491.2 | 334.6 | 374.4 | 5 | 5 | 5 | 5 | 2.162 |
| 66.9 | 51.1 | 46 | 40.7 | 1 | 1 | 1 | 1 | 0.174 |
| 259.4 | 226.8 | 209.8 | 197.6 | 3 | 3 | 3 | 3 | 0.833 |
| 233.7 | 198.4 | 194 | 188.1 | 2 | 2 | 2 | 2 | 0.638 |
| 115.1 | 99.2 | 101.9 | 98.7 | 2 | 2 | 2 | 2 | 1.015 |
| 93.8 | 69.6 | 66.5 | 72.8 | 1 | 1 | 1 | 1 | 0.259 |
| 188.7 | 128.5 | 112.3 | 127.6 | 1 | 1 | 1 | 1 | 2.162 |
| 47.3 | 49.6 | 48.5 | 43.8 | 1 | 1 | 1 | 1 | 0.116 |
| 183.8 | 137.3 | 154.7 | 150.8 | 3 | 3 | 3 | 3 | 1.239 |
| 52.4 | 36.9 | 38.4 | 35.1 | 1 | 1 | 1 | 1 | 0.389 |
| 199.3 | 172 | 157.2 | 142.2 | 3 | 3 | 3 | 3 | 2.162 |
| | | | | | | | | 0.248 |
| 757.5 | 521.2 | 571.3 | 536.1 | 10 | 10 | 10 | 10 | 10.103 |
| 74.8 | 64.2 | 59.5 | 59.6 | 1 | 1 | 1 | 1 | 0.116 |
| 115.5 | 92.2 | 99.6 | 92.7 | 2 | 2 | 2 | 2 | 0.413 |
| | | | | | | | | 0.311 |
| 217.4 | 237.1 | 177.8 | 153.8 | 2 | 2 | 2 | 2 | 643.947 |
| 558.2 | 405 | 417.2 | 379 | 4 | 4 | 4 | 4 | 2.981 |
| | | | | | | | | 0.259 |
| 89.7 | 86.2 | 67.3 | 71.6 | 1 | 1 | 1 | 1 | 0.086 |
| 13.9 | 16.5 | 11.9 | 10.9 | 1 | 1 | 1 | 1 | 0.425 |
| 70.2 | 58.8 | 53.7 | 50.7 | 1 | 1 | 1 | 1 | 0.334 |
| 131.4 | 102.7 | 100.3 | 95.1 | 1 | 1 | 1 | 1 | 0.155 |
| 95.4 | 70.7 | 75.6 | 62.9 | 1 | 1 | 1 | 1 | 0.259 |
| 252.2 | 253.1 | 216.1 | 207.3 | 3 | 3 | 3 | 3 | 0.778 |
| | | | | | | | | 0.136 |
| | | | | | | | | 0.186 |
| | | | | | | | | 0.778 |
| 253.9 | 201.2 | 197.2 | 183.3 | 5 | 5 | 5 | 5 | 0.833 |
| 169.4 | 117.7 | 110.4 | 113.6 | 1 | 1 | 1 | 1 | 0.585 |
| | | | | | | | | 0.292 |
| | | | | | | | | 0.101 |
| | | | | | | | | 0.52 |
| 1184.3 | 975.1 | 928.2 | 879.4 | 9 | 9 | 9 | 9 | 6.943 |
| | | | | | | | | 0.179 |
| 1.2 | 8.7 | 1.2 | 1.2 | 1 | 1 | 1 | 1 | 0.585 |
| 72.8 | 54.9 | 67.3 | 52.2 | 1 | 1 | 1 | 1 | 0.311 |
| 359.6 | 261 | 287.7 | 269.9 | 5 | 5 | 5 | 5 | 2.875 |
| 316.7 | 269.9 | 259.7 | 242.2 | 4 | 4 | 4 | 4 | 4.179 |
| 507.9 | 468 | 422.2 | 418.1 | 3 | 3 | 3 | 3 | 1.848 |
| 87.6 | 80 | 75.6 | 60.7 | 2 | 2 | 2 | 2 | 0.425 |
| | | | | | | | | 0.046 |
| 57.1 | 52.8 | 54.7 | 45.2 | 1 | 1 | 1 | 1 | 0.778 |
| 307.9 | 307.3 | 268.8 | 236.9 | 8 | 8 | 8 | 8 | 1.268 |
| 377.1 | 336 | 294.2 | 287.8 | 3 | 3 | 3 | 3 | 2.162 |
| 56.9 | 73.8 | 61 | 46.1 | 1 | 1 | 1 | 1 | 1.783 |
| 66.8 | 62.1 | 61.4 | 52.9 | 1 | 1 | 1 | 1 | 0.259 |
| | | | | | | | | 2.03 |
| 784.8 | 591 | 570.5 | 597.3 | 5 | 5 | 5 | 5 | 2.02 |
| 139.2 | 89.6 | 95.7 | 87.3 | 1 | 1 | 1 | 1 | 0.701 |
| | | | | | | | | 0.11 |
| | | | | | | | | 0.136 |
| | | | | | | | | 0.129 |
| | | | | | | | | 0.166 |
| 161.3 | 128.6 | 113.8 | 109.9 | 1 | 1 | 1 | 1 | 0.292 |
| | | | | | | | | 0.11 |
| 21 | 25.8 | 15.4 | 12.2 | 2 | 2 | 2 | 2 | 0.52 |
| | | | | | | | | 0.059 |
| 23.1 | 18.2 | 15.5 | 15.8 | 1 | 1 | 1 | 1 | 0.304 |
| | | | | | | | | 0.233 |
| 22.6 | 23.7 | 17.2 | 18.8 | 1 | 1 | 1 | 1 | 0.08 |

| | | | | | | | | |
|---|---|---|---|---|---|---|---|---|
| 52 | 44.2 | 36.8 | 37.4 | 1 | 1 | 1 | 1 | 0.11 |
| 77.9 | 55.2 | 57.4 | 51.5 | 1 | 1 | 1 | 1 | 0.585 |
| | | | | | | | | 0.096 |
| 135.4 | 92.4 | 91.3 | 90.6 | 2 | 2 | 2 | 2 | 0.468 |
| | | | | | | | | 0.179 |
| 173.4 | 141.3 | 124.3 | 123.1 | 3 | 3 | 3 | 3 | 1.154 |
| | | | | | | | | 0.468 |
| 62.7 | 53 | 58.8 | 60.6 | 1 | 1 | 1 | 1 | 0.425 |
| | | | | | | | | 0.194 |
| 77 | 54.4 | 46.5 | 52.3 | 1 | 1 | 1 | 1 | 0.11 |
| | | | | | | | | 1.154 |
| 345.2 | 262 | 272.7 | 242.5 | 2 | 2 | 2 | 2 | 0.52 |
| 412 | 325 | 333 | 291.3 | 4 | 4 | 4 | 4 | 2.162 |
| | | | | | | | | 0.389 |
| 173.2 | 121.3 | 127.1 | 134.2 | 1 | 1 | 1 | 1 | 0.116 |
| 35.4 | 37.6 | 35.1 | 24.6 | 2 | 2 | 2 | 2 | 0.995 |
| | | | | | | | | 0.212 |
| 81.6 | 51.7 | 60.1 | 54.2 | 1 | 1 | 1 | 1 | 0.369 |
| 41.4 | 40.5 | 42 | 41.4 | 1 | 1 | 1 | 1 | 0.292 |
| 67.4 | 62.3 | 50.7 | 54.2 | 1 | 1 | 1 | 1 | 0.136 |
| | | | | | | | | 0.212 |
| 475.3 | 399.7 | 356.2 | 364.3 | 5 | 5 | 5 | 5 | 2.793 |
| 93.2 | 76.2 | 87 | 75.3 | 2 | 2 | 2 | 2 | 0.848 |
| 83.7 | 65.6 | 65.1 | 55.5 | 1 | 1 | 1 | 1 | 0.179 |
| 135.1 | 104 | 103.5 | 99.4 | 1 | 1 | 1 | 1 | 9 |
| 1105.5 | 1127 | 972.4 | 881.6 | 7 | 7 | 7 | 7 | 24.119 |
| | | | | | | | | 0.07 |
| 21.8 | 25.7 | 23.3 | 20.5 | 1 | 1 | 1 | 1 | 0.468 |
| | | | | | | | | 0.155 |
| 125.9 | 111.9 | 100.6 | 105.9 | 1 | 1 | 1 | 1 | 9 |
| 70.9 | 41.5 | 50.6 | 54.7 | 1 | 1 | 1 | 1 | 0.468 |
| | | | | | | | | 0.093 |
| 119.7 | 99.1 | 77.3 | 81.2 | 1 | 1 | 1 | 1 | 0.407 |
| 61.9 | 51.5 | 52.5 | 50.2 | 1 | 1 | 1 | 1 | 0.212 |
| | | | | | | | | 0.292 |
| 96.7 | 86.1 | 77.8 | 76.4 | 1 | 1 | 1 | 1 | 0.381 |
| 171.2 | 145.7 | 150.1 | 144.4 | 3 | 3 | 3 | 3 | 0.302 |
| | | | | | | | | 0.259 |
| | | | | | | | | 0.11 |
| | | | | | | | | 0.038 |
| | | | | | | | | 0.292 |
| | | | | | | | | 0.166 |
| 75.6 | 74.6 | 70.2 | 54.1 | 1 | 1 | 1 | 1 | 0.638 |
| | | | | | | | | 1.297 |
| 46.4 | 37.7 | 34.3 | 32.2 | 1 | 1 | 1 | 1 | 0.585 |
| | | | | | | | | 0.116 |
| | | | | | | | | 0.129 |
| 328.3 | 216.9 | 186.9 | 207.2 | 4 | 4 | 4 | 4 | 0.624 |
| 300.9 | 205 | 226.9 | 224.2 | 2 | 2 | 2 | 2 | 0.425 |
| | | | | | | | | 0.048 |
| 118.4 | 85.4 | 95.3 | 73.6 | 1 | 1 | 1 | 1 | 0.292 |
| 139.8 | 119.5 | 126.9 | 98.8 | 3 | 3 | 3 | 3 | 0.407 |
| 259 | 221.5 | 211.3 | 205.2 | 4 | 4 | 4 | 4 | 14.505 |
| 1272.6 | 1011.9 | 1037.9 | 885.4 | 9 | 9 | 9 | 9 | 2.054 |
| | | | | | | | | 0.389 |
| | | | | | | | | 0.212 |
| | | | | | | | | 0.874 |
| | | | | | | | | 0.585 |
| | | | | | | | | 0.136 |
| 78.7 | 67.2 | 57.3 | 57.1 | 1 | 1 | 1 | 1 | 0.334 |
| | | | | | | | | 0.194 |
| | | | | | | | | 0.233 |

| | | | | | | | | |
|---|---|---|---|---|---|---|---|---|
| 174 | 146.3 | 122 | 123.4 | 2 | 2 | 2 | 2 | 0.668 |
| 13.8 | 8.4 | 8.3 | 11.3 | 1 | 1 | 1 | 1 | 0.778 |
| 70.3 | 51.3 | 47.1 | 45.9 | 1 | 1 | 1 | 1 | 0.995 |
| 21.3 | 19.5 | 14.6 | 14.2 | 2 | 2 | 2 | 2 | 0.778 |
| 32.1 | 20.8 | 20.7 | 16.3 | 1 | 1 | 1 | 1 | 0.212 |
| 50.6 | 51.5 | 48.4 | 41.5 | 1 | 1 | 1 | 1 | 0.585 |
| 590.6 | 496.1 | 494.6 | 450.9 | 7 | 7 | 7 | 7 | 3.217 |
| 471.2 | 479.3 | 419.8 | 365.9 | 6 | 6 | 6 | 6 | 2.384 |
| 14.9 | 11.2 | 12.1 | 11 | 1 | 1 | 1 | 1 | 0.501 |
| 145.6 | 139.5 | 139.9 | 121.9 | 2 | 2 | 2 | 2 | 1.683 |
| | | | | | | | | 0.179 |
| 136.8 | 127.8 | 120.8 | 130.4 | 1 | 1 | 1 | 1 | 0.389 |
| 199.7 | 170.7 | 171.9 | 158.1 | 2 | 2 | 2 | 2 | 2.162 |
| | | | | | | | | 0.389 |
| 129.9 | 147.9 | 122.1 | 101.5 | 1 | 1 | 1 | 1 | 0.292 |
| 552.6 | 383.1 | 397.2 | 391.8 | 6 | 6 | 6 | 6 | 53.844 |
| 268.9 | 240.2 | 217.6 | 221.4 | 2 | 2 | 2 | 2 | 0.995 |
| 1071.6 | 939.4 | 885 | 784.7 | 9 | 9 | 9 | 9 | 59.619 |
| 12 | 10.9 | 12 | 9.5 | 1 | 1 | 1 | 1 | 1.512 |
| | | | | | | | | 0.054 |
| | | | | | | | | 0.067 |
| 138.1 | 94.5 | 110.9 | 94.6 | 1 | 1 | 1 | 1 | 2.981 |
| | | | | | | | | 0.389 |
| 151.4 | 143.6 | 144.2 | 121.1 | 3 | 3 | 3 | 3 | 2.162 |
| 219.3 | 181.4 | 172.8 | 169.2 | 5 | 5 | 5 | 5 | 3.642 |
| 92 | 64.9 | 74.8 | 69.6 | 1 | 1 | 1 | 1 | 0.304 |
| 461.7 | 330.3 | 352.5 | 347.1 | 5 | 5 | 5 | 5 | 0.598 |
| 223.6 | 192.5 | 218.3 | 183.4 | 5 | 5 | 5 | 5 | 0.931 |
| 1036.8 | 913.9 | 926.1 | 831 | 9 | 9 | 9 | 9 | 2.162 |
| 392.5 | 328.7 | 332.3 | 303.6 | 2 | 2 | 2 | 2 | 0.719 |
| 72.1 | 72 | 52.4 | 51.2 | 1 | 1 | 1 | 1 | 0.119 |
| 48.6 | 30.4 | 35.9 | 31.8 | 1 | 1 | 1 | 1 | 0.03 |
| 103.2 | 76.5 | 92.4 | 85.4 | 1 | 1 | 1 | 1 | 0.638 |
| | | | | | | | | 0.701 |
| 59.1 | 46.3 | 41.5 | 41.1 | 1 | 1 | 1 | 1 | 0.086 |
| | | | | | | | | 0.083 |
| | | | | | | | | 0.848 |
| | | | | | | | | 0.116 |
| | | | | | | | | 1.424 |
| | | | | | | | | 0.136 |
| 1226.7 | 956.1 | 950.7 | 931.7 | 14 | 14 | 14 | 14 | 3.977 |
| 147.7 | 111.1 | 123.3 | 108.2 | 3 | 3 | 3 | 3 | 0.823 |
| | | | | | | | | 0.066 |
| 146.6 | 89.9 | 65.6 | 80.7 | 2 | 2 | 2 | 2 | 0.585 |
| 233.8 | 170.8 | 177.1 | 164 | 2 | 2 | 2 | 2 | 0.202 |
| 531.8 | 418.1 | 397.9 | 415.8 | 8 | 8 | 8 | 8 | 3.281 |
| 74.9 | 74.4 | 90 | 79 | 1 | 1 | 1 | 1 | 0.212 |
| 63.4 | 49.3 | 51.3 | 42.2 | 1 | 1 | 1 | 1 | 0.52 |
| 396.8 | 205.4 | 246.1 | 272.3 | 2 | 2 | 2 | 2 | 0.778 |
| | | | | | | | | 0.136 |
| 678.1 | 543.2 | 533.3 | 483.6 | 5 | 5 | 5 | 5 | 3.329 |
| 307 | 259.8 | 250.7 | 243.7 | 4 | 4 | 4 | 4 | 0.701 |
| | | | | | | | | 0.274 |
| 38.2 | 32.6 | 29.8 | 25.5 | 1 | 1 | 1 | 1 | 0.194 |
| 598.4 | 459.5 | 478.4 | 482.3 | 7 | 7 | 7 | 7 | 4.217 |
| | | | | | | | | 0.105 |
| | | | | | | | | 0.253 |
| 370.4 | 286.9 | 290.1 | 273.4 | 4 | 4 | 4 | 4 | 0.433 |
| 167.7 | 110.6 | 129.6 | 123 | 1 | 1 | 1 | 1 | 0.425 |
| 79.3 | 74 | 71.1 | 68.2 | 1 | 1 | 1 | 1 | 0.179 |
| 88.6 | 81.3 | 75.8 | 67.2 | 1 | 1 | 1 | 1 | 0.487 |
| 102.6 | 127.1 | 102.4 | 92.2 | 1 | 1 | 1 | 1 | 0.389 |

| | | | | | | | | |
|---|---|---|---|---|---|---|---|---|
| 57.8 | 46.9 | 60.6 | 50.5 | 2 | 2 | 2 | 2 | 0.147 |
| 943.6 | 759.1 | 745.2 | 718.3 | 9 | 9 | 9 | 9 | 19.535 |
| 1296.6 | 1018.8 | 897.2 | 919.9 | 13 | 13 | 13 | 13 | 10.365 |
| | | | | | | | | 0.059 |
| 60 | 46.5 | 53.2 | 46.5 | 1 | 1 | 1 | 1 | 0.145 |
| | | | | | | | | 0.194 |
| 84.3 | 72.8 | 56 | 47.2 | 1 | 1 | 1 | 1 | 0.292 |
| 1183.3 | 909.8 | 892.2 | 811.2 | 9 | 9 | 9 | 9 | 3.642 |
| | | | | | | | | 0.334 |
| | | | | | | | | 0.233 |
| | | | | | | | | 0.11 |
| | | | | | | | | 0.155 |
| | | | | | | | | 0.194 |
| | | | | | | | | 0.438 |
| | | | | | | | | 0.259 |
| 42.8 | 33.7 | 36.9 | 31.2 | 1 | 1 | 1 | 1 | 0.233 |
| 525.4 | 444.9 | 389.6 | 373.1 | 6 | 6 | 6 | 6 | 0.604 |
| 8.8 | 5.5 | 9 | 7.9 | 1 | 1 | 1 | 1 | 0.145 |
| | | | | | | | | 0.292 |
| | | | | | | | | 0.105 |
| | | | | | | | | 0.334 |
| | | | | | | | | 0.105 |
| | | | | | | | | 0.101 |
| 137.5 | 127.7 | 115.6 | 122.5 | 2 | 2 | 2 | 2 | 0.334 |
| | | | | | | | | 0.259 |
| | | | | | | | | 0.194 |
| 1328.4 | 653.2 | 450.6 | 618.8 | 7 | 7 | 7 | 7 | 8.12 |
| | | | | | | | | 0.245 |
| | | | | | | | | 0.468 |
| 371.1 | 282 | 295.8 | 278.2 | 2 | 2 | 2 | 2 | 0.468 |
| 27.8 | 27.7 | 19.3 | 21.7 | 1 | 1 | 1 | 1 | 0.318 |
| 41.3 | 84.2 | 43 | 54.3 | 1 | 1 | 1 | 1 | 0.52 |
| | | | | | | | | 0.089 |
| | | | | | | | | 0.222 |
| 67.2 | 97 | 66 | 55.1 | 1 | 1 | 1 | 1 | 463.159 |
| 730.1 | 652.1 | 565.1 | 550.8 | 5 | 5 | 5 | 5 | 463.159 |
| 249.1 | 229.8 | 197.7 | 201.3 | 3 | 3 | 3 | 3 | 0.509 |
| | | | | | | | | 0.108 |
| 111.2 | 77.4 | 83.4 | 89.3 | 1 | 1 | 1 | 1 | 0.52 |
| 32.9 | 45.2 | 34 | 26.4 | 1 | 1 | 1 | 1 | 0.093 |
| 43.3 | 43.2 | 28 | 29 | 1 | 1 | 1 | 1 | 0.212 |
| 1285.6 | 1234.7 | 1111.9 | 943.9 | 16 | 16 | 16 | 16 | 5.136 |
| 66.8 | 55.4 | 47.7 | 37.8 | 1 | 1 | 1 | 1 | 0.122 |
| 170.9 | 136.6 | 148.6 | 123.6 | 2 | 2 | 2 | 2 | 0.233 |
| | | | | | | | | 0.112 |
| 1911.8 | 1593.1 | 1628.2 | 1473.1 | 27 | 27 | 27 | 27 | 35.869 |
| | | | | | | | | 0.044 |
| 76.3 | 44.8 | 62.1 | 42.7 | 1 | 1 | 1 | 1 | 0.565 |
| 155.5 | 150.1 | 133.6 | 127.4 | 3 | 3 | 3 | 3 | 0.73 |
| 220.8 | 201.2 | 174.3 | 160.9 | 7 | 7 | 7 | 7 | 0.28 |
| | | | | | | | | 0.233 |
| | | | | | | | | 0.194 |
| | | | | | | | | 0.08 |
| 78.9 | 53.3 | 68.2 | 65.6 | 1 | 1 | 1 | 1 | 0.501 |
| | | | | | | | | 0.585 |
| 158.1 | 115.1 | 127.8 | 132.5 | 2 | 2 | 2 | 2 | 1.276 |
| 290.5 | 230.1 | 167.5 | 179.4 | 3 | 3 | 3 | 3 | 0.45 |
| 40.3 | 32.3 | 34 | 27.9 | 1 | 1 | 1 | 1 | 0.272 |
| 70 | 56.3 | 57 | 53.7 | 1 | 1 | 1 | 1 | 0.038 |
| | | | | | | | | 0.122 |
| 392.2 | 321.5 | 316.3 | 303.9 | 6 | 6 | 6 | 6 | 0.571 |
| 102.9 | 80.7 | 86.6 | 81.4 | 2 | 2 | 2 | 2 | 0.52 |

| | | | | | | | | |
|---|---|---|---|---|---|---|---|---|
| | | | | | | | | 0.233 |
| 130 | 91.3 | 89.2 | 88.7 | 1 | 1 | 1 | 1 | 0.585 |
| 28.9 | 21.6 | 22.3 | 24 | 1 | 1 | 1 | 1 | 0.166 |
| | | | | | | | | 0.116 |
| | | | | | | | | 0.051 |
| 50.8 | 42.1 | 47.4 | 35.9 | 1 | 1 | 1 | 1 | 0.145 |
| | | | | | | | | 0.096 |
| | | | | | | | | 0.304 |
| | | | | | | | | 0.014 |
| | | | | | | | | 0.037 |
| | | | | | | | | 0.212 |
| 41.9 | 36.8 | 36.6 | 28.3 | 1 | 1 | 1 | 1 | 0.233 |
| 196.3 | 195.5 | 144.4 | 146.2 | 1 | 1 | 1 | 1 | 0.292 |
| 47 | 44.8 | 42 | 35.8 | 1 | 1 | 1 | 1 | 0.334 |
| 121 | 96 | 83.3 | 83.1 | 2 | 2 | 2 | 2 | 0.688 |
| 153.3 | 143.2 | 142.2 | 126.2 | 2 | 2 | 2 | 2 | 0.557 |
| 155.1 | 118.6 | 125.9 | 105.2 | 3 | 3 | 3 | 3 | 0.429 |
| 61.3 | 60.3 | 49.7 | 45.3 | 1 | 1 | 1 | 1 | 0.172 |
| 68.5 | 65.3 | 63.6 | 52.9 | 1 | 1 | 1 | 1 | 0.08 |
| 108.4 | 96.6 | 74.7 | 71.7 | 1 | 1 | 1 | 1 | 4.623 |
| 88.3 | 76.4 | 81.8 | 70.1 | 1 | 1 | 1 | 1 | 0.438 |
| 104 | 94.8 | 85.5 | 79 | 2 | 2 | 2 | 2 | 0.359 |
| | | | | | | | | 0.245 |
| 58 | 49.5 | 44.4 | 46 | 1 | 1 | 1 | 1 | 0.778 |
| 2791.9 | 2250.4 | 2313 | 2081.6 | 27 | 27 | 27 | 27 | 18.11 |
| | | | | | | | | 0.194 |
| 132.7 | 119.6 | 121.1 | 98.7 | 1 | 1 | 1 | 1 | 0.468 |
| | | | | | | | | 0.116 |
| 430 | 353.5 | 356.7 | 325.9 | 9 | 9 | 9 | 9 | 154.052 |
| 74.7 | 70 | 92.8 | 75.1 | 1 | 1 | 1 | 1 | 0.778 |
| 86.5 | 73.2 | 74.5 | 61.5 | 1 | 1 | 1 | 1 | 0.292 |
| | | | | | | | | 0.11 |
| 81 | 63.7 | 57.7 | 52 | 1 | 1 | 1 | 1 | 0.129 |
| 58.2 | 52.8 | 49.5 | 44.3 | 1 | 1 | 1 | 1 | 0.186 |
| | | | | | | | | 0.046 |
| 44.4 | 40.2 | 51.3 | 39.4 | 2 | 2 | 2 | 2 | 0.311 |
| | | | | | | | | 0.086 |
| | | | | | | | | 0.07 |
| 191.5 | 117.5 | 123.8 | 149.3 | 1 | 1 | 1 | 1 | 1.512 |
| 62.8 | 57.4 | 45.2 | 40.2 | 1 | 1 | 1 | 1 | 0.012 |
| | | | | | | | | 0.11 |
| | | | | | | | | 0.212 |
| | | | | | | | | 0.342 |
| 116.6 | 102.4 | 121.1 | 100.4 | 2 | 2 | 2 | 2 | 0.931 |
| 114.3 | 130.8 | 116 | 84.4 | 2 | 2 | 2 | 2 | 0.52 |
| | | | | | | | | 0.212 |
| 39.5 | 32.8 | 25.2 | 21.5 | 1 | 1 | 1 | 1 | 0.218 |
| | | | | | | | | 0.059 |
| 76.1 | 70.1 | 59.1 | 57.1 | 1 | 1 | 1 | 1 | 0.155 |
| | | | | | | | | 0.896 |
| 62 | 57.9 | 58.8 | 55.2 | 1 | 1 | 1 | 1 | 0.318 |
| | | | | | | | | 0.056 |
| 74.7 | 83.3 | 82.2 | 58.5 | 2 | 2 | 2 | 2 | 0.425 |
| | | | | | | | | 0.274 |
| 29.5 | 29.4 | 22.5 | 21 | 1 | 1 | 1 | 1 | 0.122 |
| 196.8 | 169.4 | 177.3 | 162.1 | 3 | 3 | 3 | 3 | 0.725 |
| 61.5 | 52.7 | 60.3 | 54.2 | 2 | 2 | 2 | 2 | 0.655 |
| | | | | | | | | 0.259 |
| 171 | 145.5 | 137 | 127 | 2 | 2 | 2 | 2 | 1.848 |
| 114.9 | 105.4 | 99.7 | 102.2 | 1 | 1 | 1 | 1 | 0.369 |
| | | | | | | | | 0.425 |
| | | | | | | | | 0.083 |

| | | | | | | | | |
|---|---|---|---|---|---|---|---|---|
| | | | | | | | | 0.194 |
| 479.2 | 452.1 | 430.3 | 311.3 | 3 | 3 | 3 | 3 | 5.813 |
| | | | | | | | | 0.17 |
| | | | | | | | | 0.233 |
| 813.4 | 603 | 655.1 | 582.4 | 6 | 6 | 6 | 6 | 1.031 |
| 1635.6 | 1416.6 | 1408.8 | 1273.4 | 20 | 20 | 20 | 20 | 8.326 |
| | | | | | | | | 0.145 |
| 61.6 | 50.9 | 41.6 | 42.6 | 2 | 2 | 2 | 2 | 0.334 |
| | | | | | | | | 0.194 |
| | | | | | | | | 0.468 |
| 44.4 | 25.9 | 20.7 | 23.8 | 1 | 1 | 1 | 1 | 0.585 |
| 88.6 | 92.2 | 63.5 | 66.2 | 2 | 2 | 2 | 2 | 0.194 |
| | | | | | | | | 0.155 |
| | | | | | | | | 0.058 |
| 20.4 | 17.5 | 13.5 | 12.6 | 1 | 1 | 1 | 1 | 0.202 |
| 187.2 | 149.7 | 123.2 | 136.5 | 1 | 1 | 1 | 1 | 0.037 |
| 52.4 | 47.2 | 51 | 40.5 | 2 | 2 | 2 | 2 | 0.079 |
| 51.9 | 42.8 | 43.6 | 37.9 | 1 | 1 | 1 | 1 | 0.638 |
| 782.2 | 515.8 | 542.8 | 467.6 | 6 | 6 | 6 | 6 | 1.471 |
| 254.5 | 212.4 | 191.4 | 187.6 | 4 | 4 | 4 | 4 | 0.179 |
| 88 | 85 | 78.2 | 60.9 | 1 | 1 | 1 | 1 | 0.701 |
| | | | | | | | | 0.212 |
| 28 | 32.3 | 25.9 | 28.9 | 1 | 1 | 1 | 1 | 0.389 |
| 937.8 | 732.8 | 766.3 | 568.5 | 13 | 13 | 13 | 13 | 2.371 |
| 86.4 | 67.8 | 64 | 57.8 | 1 | 1 | 1 | 1 | 0.359 |
| 115.1 | 101.9 | 95 | 90 | 3 | 3 | 3 | 3 | 0.848 |
| | | | | | | | | 0.11 |
| | | | | | | | | 0.179 |
| 1458.5 | 1271.2 | 1333.9 | 1218.9 | 20 | 20 | 20 | 20 | 1.131 |
| | | | | | | | | 0.259 |
| 283.8 | 289.4 | 266.6 | 243.9 | 4 | 4 | 4 | 4 | 6.499 |
| 68.8 | 56.9 | 47.8 | 53.9 | 1 | 1 | 1 | 1 | 0.136 |
| | | | | | | | | 0.136 |
| 13.1 | 13.1 | 8.9 | 9.2 | 1 | 1 | 1 | 1 | 0.259 |
| 61.8 | 49.3 | 56.6 | 48 | 1 | 1 | 1 | 1 | 0.425 |
| | | | | | | | | 0.233 |
| | | | | | | | | 0.233 |
| | | | | | | | | 0.077 |
| 126.7 | 99.4 | 78.9 | 93 | 1 | 1 | 1 | 1 | 0.054 |
| 255.1 | 215 | 199.5 | 196.9 | 3 | 3 | 3 | 3 | 0.189 |
| | | | | | | | | 0.096 |
| | | | | | | | | 0.083 |
| | | | | | | | | 0.129 |
| 109.1 | 77.8 | 51.1 | 55.3 | 1 | 1 | 1 | 1 | 0.512 |
| | | | | | | | | 0.136 |
| | | | | | | | | 0.212 |
| 223.8 | 195.2 | 168.2 | 162.2 | 6 | 6 | 6 | 6 | 1.703 |
| 194.4 | 179.5 | 161.8 | 139.3 | 2 | 2 | 2 | 2 | 0.108 |
| 69.3 | 57.9 | 65 | 40 | 1 | 1 | 1 | 1 | 0.438 |
| | | | | | | | | 0.11 |
| 102.5 | 87 | 80.7 | 76.6 | 1 | 1 | 1 | 1 | 0.072 |
| 584 | 390.2 | 403.1 | 408.3 | 2 | 2 | 2 | 2 | 1.512 |
| 52.8 | 49.3 | 60.3 | 39.3 | 3 | 3 | 3 | 3 | 0.585 |
| 41.1 | 39.2 | 35.2 | 26.6 | 1 | 1 | 1 | 1 | 0.136 |
| 103.2 | 73.3 | 82.3 | 72 | 1 | 1 | 1 | 1 | 0.233 |
| 44.1 | 33.1 | 40.1 | 35.1 | 1 | 1 | 1 | 1 | 0.15 |
| 442.2 | 391.8 | 352.1 | 293 | 4 | 4 | 4 | 4 | 0.896 |
| 133 | 125.4 | 121.4 | 102.5 | 2 | 2 | 2 | 2 | 0.896 |
| 50.1 | 37.1 | 42 | 39.9 | 1 | 1 | 1 | 1 | 0.118 |
| | | | | | | | | 0.089 |
| 25.8 | 20.8 | 18.4 | 18.3 | 1 | 1 | 1 | 1 | 0.083 |
| 592.6 | 496.8 | 482.1 | 384.3 | 11 | 11 | 11 | 11 | 2.162 |

| | | | | | | | | |
|---|---|---|---|---|---|---|---|---|
| 64.4 | 50.9 | 55.6 | 52.1 | 1 | 1 | 1 | 1 | 0.202 |
| 102.3 | 82 | 77.1 | 68.7 | 1 | 1 | 1 | 1 | 0.166 |
| | | | | | | | | 0.058 |
| | | | | | | | | 0.05 |
| 101.8 | 86.5 | 77.1 | 86.6 | 1 | 1 | 1 | 1 | 0.11 |
| 68.4 | 58.5 | 60.4 | 56.8 | 1 | 1 | 1 | 1 | 0.28 |
| | | | | | | | | 0.233 |
| | | | | | | | | 0.07 |
| | | | | | | | | 1.512 |
| | | | | | | | | 0.233 |
| | | | | | | | | 0.089 |
| 174.7 | 178.2 | 169.6 | 138.1 | 2 | 2 | 2 | 2 | 0.16 |
| | | | | | | | | 0.166 |
| 534.2 | 499.7 | 466.1 | 431 | 5 | 5 | 5 | 5 | 4.012 |
| 1005.6 | 801 | 699.7 | 507.8 | 6 | 6 | 6 | 6 | 2.455 |
| 24.9 | 22.2 | 25.1 | 20.2 | 1 | 1 | 1 | 1 | 0.222 |
| | | | | | | | | 0.04 |
| 62.7 | 44.1 | 33 | 38.5 | 1 | 1 | 1 | 1 | 0.044 |
| 107.8 | 96.5 | 98.9 | 85.1 | 1 | 1 | 1 | 1 | 0.166 |
| 11.7 | 9.4 | 5.6 | 7.6 | 1 | 1 | 1 | 1 | 0.194 |
| | | | | | | | | 0.194 |
| | | | | | | | | 0.105 |
| 17 | 11.8 | 10.1 | 12.6 | 2 | 2 | 2 | 2 | 0.145 |
| 415.5 | 320.2 | 381.6 | 322.8 | 5 | 5 | 5 | 5 | 1.929 |
| 125.9 | 99.3 | 109 | 82.7 | 1 | 1 | 1 | 1 | 0.179 |
| | | | | | | | | 0.212 |
| | | | | | | | | 0.11 |
| 117 | 115.3 | 104.9 | 97 | 2 | 2 | 2 | 2 | 0.269 |
| | | | | | | | | 0.129 |
| 23.5 | 11.2 | 17.3 | 16.6 | 2 | 2 | 2 | 2 | 1.512 |
| 139.4 | 115.1 | 104.3 | 103.9 | 1 | 1 | 1 | 1 | 0.066 |
| 68.3 | 65.8 | 65.2 | 57.6 | 1 | 1 | 1 | 1 | 0.52 |
| 151.2 | 172.5 | 146.9 | 135.4 | 2 | 2 | 2 | 2 | 0.292 |
| | | | | | | | | 0.145 |
| | | | | | | | | 0.292 |
| | | | | | | | | 0.077 |
| 279.1 | 314.1 | 248.6 | 232.2 | 4 | 4 | 4 | 4 | 4.623 |
| 111.7 | 89 | 86.9 | 80.5 | 1 | 1 | 1 | 1 | 0.369 |
| 475.8 | 397.5 | 378 | 347.4 | 7 | 7 | 7 | 7 | 0.61 |
| | | | | | | | | 0.086 |
| | | | | | | | | 0.468 |
| 776 | 662.8 | 647 | 623.5 | 8 | 8 | 8 | 8 | 2.793 |
| 390 | 309 | 320.9 | 292.1 | 6 | 6 | 6 | 6 | 0.299 |
| | | | | | | | | 0.048 |
| 13.5 | 14.6 | 8 | 9.6 | 1 | 1 | 1 | 1 | 1.894 |
| | | | | | | | | 0.145 |
| 257.8 | 183.7 | 198.8 | 198.3 | 3 | 3 | 3 | 3 | 0.73 |
| | | | | | | | | 0.506 |
| | | | | | | | | 0.093 |
| | | | | | | | | 0.064 |
| 167.9 | 126.6 | 121.1 | 123 | 3 | 3 | 3 | 3 | 0.431 |
| 112.7 | 86.6 | 72.7 | 78.2 | 1 | 1 | 1 | 1 | 0.105 |
| 128.2 | 95.1 | 95.3 | 81.7 | 1 | 1 | 1 | 1 | 0.311 |
| 623.1 | 524.5 | 531.7 | 448.3 | 6 | 6 | 6 | 6 | 1.955 |
| | | | | | | | | 0.08 |
| 208.8 | 161.6 | 150.2 | 135.6 | 1 | 1 | 1 | 1 | 0.16 |
| 559.7 | 421.3 | 482.3 | 422.9 | 5 | 5 | 5 | 5 | 1.581 |
| 115.9 | 108 | 88.8 | 92.9 | 1 | 1 | 1 | 1 | 5.236 |
| 104.7 | 89.7 | 82 | 80.1 | 1 | 1 | 1 | 1 | 0.585 |
| 20.4 | 17.8 | 21.9 | 14.8 | 1 | 1 | 1 | 1 | 0.212 |
| | | | | | | | | 0.468 |
| | | | | | | | | 0.11 |

| | | | | | | | | |
|---|---|---|---|---|---|---|---|---|
| 375.5 | 309.2 | 358.6 | 332.4 | 8 | 8 | 8 | 8 | 111.202 |
| 450.7 | 375.2 | 415.3 | 355.7 | 5 | 5 | 5 | 5 | 0.389 |
| | | | | | | | | 0.155 |
| 983.3 | 737.1 | 804.7 | 712.3 | 10 | 10 | 10 | 10 | 3.184 |
| 100.5 | 84.5 | 83.6 | 65.8 | 1 | 1 | 1 | 1 | 0.141 |
| 105.1 | 84 | 85.2 | 77.3 | 2 | 2 | 2 | 2 | 0.346 |
| 74.2 | 91.7 | 70.7 | 61.9 | 1 | 1 | 1 | 1 | 0.054 |
| 131.5 | 117.4 | 107.3 | 102.5 | 3 | 3 | 3 | 3 | 0.8 |
| 532.7 | 553.9 | 517.8 | 443.6 | 8 | 8 | 8 | 8 | 3.062 |
| | | | | | | | | 0.778 |
| | | | | | | | | 0.145 |
| 222.3 | 222.1 | 202.7 | 183.5 | 1 | 1 | 1 | 1 | 0.334 |
| 1449 | 1338.2 | 1306.3 | 1099.8 | 18 | 18 | 18 | 18 | 10.029 |
| | | | | | | | | 0.212 |
| | | | | | | | | 0.166 |
| | | | | | | | | 0.129 |
| | | | | | | | | 0.212 |
| 256.5 | 229.5 | 210.7 | 188.6 | 4 | 4 | 4 | 4 | 0.54 |
| | | | | | | | | 0.413 |
| 132.7 | 115.9 | 125.8 | 97.4 | 1 | 1 | 1 | 1 | 0.043 |
| | | | | | | | | 0.136 |
| 131.9 | 113 | 102.5 | 105.7 | 2 | 2 | 2 | 2 | 0.212 |
| | | | | | | | | 0.031 |
| 336.8 | 295.9 | 247 | 246.3 | 4 | 4 | 4 | 4 | 0.372 |
| | | | | | | | | 0.259 |
| | | | | | | | | 0.194 |
| | | | | | | | | 0.218 |
| 105.9 | 95.4 | 89.3 | 86.6 | 1 | 1 | 1 | 1 | 1.254 |
| 154.1 | 147.1 | 126.5 | 105.8 | 2 | 2 | 2 | 2 | 0.778 |
| 195.5 | 163.8 | 148 | 150.3 | 2 | 2 | 2 | 2 | 0.172 |
| | | | | | | | | 0.116 |
| 102.4 | 83.2 | 79.9 | 73 | 1 | 1 | 1 | 1 | 0.212 |
| | | | | | | | | 0.028 |
| 195 | 196.8 | 187.7 | 155.8 | 4 | 4 | 4 | 4 | 0.638 |
| | | | | | | | | 0.292 |
| | | | | | | | | 0.116 |
| 16.3 | 10.7 | 9.6 | 11.6 | 1 | 1 | 1 | 1 | 0.274 |
| | | | | | | | | 0.024 |
| 91.4 | 72.1 | 75 | 64.7 | 1 | 1 | 1 | 1 | 0.468 |
| 71.2 | 67.9 | 68.5 | 63.6 | 1 | 1 | 1 | 1 | 0.122 |
| | | | | | | | | 0.038 |
| 464.4 | 312.8 | 277.5 | 254.7 | 2 | 2 | 2 | 2 | 1.471 |
| | | | | | | | | 0.03 |
| 44.6 | 30.1 | 30.1 | 33.6 | 1 | 1 | 1 | 1 | 0.038 |
| | | | | | | | | 0.041 |
| 309.3 | 274.1 | 261.2 | 241.3 | 3 | 3 | 3 | 3 | 5.31 |
| 329.3 | 288.7 | 248.7 | 255 | 3 | 3 | 3 | 3 | 4.623 |
| | | | | | | | | 0.036 |
| | | | | | | | | 0.073 |
| | | | | | | | | 0.166 |
| 53.3 | 49.4 | 44.9 | 48.6 | 1 | 1 | 1 | 1 | 0.233 |
| | | | | | | | | 0.008 |
| 975.9 | 779.4 | 769.7 | 744.1 | 9 | 9 | 9 | 9 | 10.788 |
| | | | | | | | | 2.831 |
| | | | | | | | | 0.212 |
| 56.4 | 44.3 | 38.5 | 38.9 | 1 | 1 | 1 | 1 | 0.122 |
| | | | | | | | | 0.292 |
| 9.9 | 5.6 | 7.1 | 5.1 | 1 | 1 | 1 | 1 | 0.194 |
| 189.8 | 138.9 | 121.1 | 129.2 | 2 | 2 | 2 | 2 | 0.528 |
| 605.5 | 504.6 | 535 | 493.3 | 5 | 5 | 5 | 5 | 0.468 |
| 56.6 | 54.3 | 52.9 | 43.8 | 1 | 1 | 1 | 1 | 0.032 |
| | | | | | | | | 0.141 |

| | | | | | | | | |
|---|---|---|---|---|---|---|---|---|
| 36.2 | 30 | 43.8 | 32.8 | 1 | 1 | 1 | 1 | 0.179 |
| 138.7 | 116.2 | 115.2 | 98.4 | 1 | 1 | 1 | 1 | 0.274 |
| 125.9 | 106.5 | 104.5 | 116.8 | 1 | 1 | 1 | 1 | 0.129 |
| | | | | | | | | 0.778 |
| 183.8 | 153.6 | 145.9 | 139.5 | 2 | 2 | 2 | 2 | 0.398 |
| 149.8 | 133.8 | 117.7 | 111.5 | 2 | 2 | 2 | 2 | 0.307 |
| | | | | | | | | 0.032 |
| 355.1 | 287.6 | 266.6 | 266.7 | 4 | 4 | 4 | 4 | 0.607 |
| | | | | | | | | 0.016 |
| | | | | | | | | 1.637 |
| 46.1 | 54.2 | 51.2 | 43.2 | 1 | 1 | 1 | 1 | 2.162 |
| | | | | | | | | 0.222 |
| 51.2 | 43.5 | 42 | 31.7 | 1 | 1 | 1 | 1 | 0.35 |
| 252.1 | 204.4 | 192.7 | 173.5 | 3 | 3 | 3 | 3 | 5.31 |
| 314.1 | 222.8 | 251.4 | 241 | 3 | 3 | 3 | 3 | 0.848 |
| 972.1 | 801.3 | 777.9 | 745.1 | 10 | 10 | 10 | 10 | 2.899 |
| 1289.2 | 985.1 | 1024.5 | 1013.4 | 22 | 22 | 22 | 22 | 2.798 |
| 432.7 | 395.3 | 339.8 | 343.9 | 4 | 4 | 4 | 4 | 0.292 |
| 197.8 | 161.8 | 167.8 | 150.8 | 3 | 3 | 3 | 3 | 0.301 |
| 212.1 | 193 | 177 | 164.5 | 2 | 2 | 2 | 2 | 0.668 |
| | | | | | | | | 0.292 |
| 76.2 | 63.7 | 66.4 | 62.3 | 1 | 1 | 1 | 1 | 0.389 |
| | | | | | | | | 0.045 |
| | | | | | | | | 0.033 |
| 66.8 | 61.4 | 60.1 | 54.2 | 1 | 1 | 1 | 1 | 0.222 |
| 277.4 | 227.2 | 219.5 | 192.5 | 2 | 2 | 2 | 2 | 0.931 |
| 175.4 | 110.3 | 115.4 | 126.6 | 1 | 1 | 1 | 1 | 0.116 |
| | | | | | | | | 0.093 |
| 116.6 | 92 | 106.7 | 85.3 | 1 | 1 | 1 | 1 | 0.11 |
| 239.9 | 201.2 | 180.9 | 181.1 | 3 | 3 | 3 | 3 | 0.194 |
| | | | | | | | | 0.122 |
| 485.9 | 362.1 | 374.4 | 336.3 | 5 | 5 | 5 | 5 | 1.976 |
| | | | | | | | | 0.11 |
| | | | | | | | | 0.151 |
| 551 | 383.9 | 392.2 | 386.2 | 6 | 6 | 6 | 6 | 192.07 |
| 418.2 | 355.3 | 338.2 | 309.3 | 8 | 8 | 8 | 8 | 6.743 |
| | | | | | | | | 0.136 |
| 20.1 | 28.6 | 19.3 | 19 | 1 | 1 | 1 | 1 | 0.172 |
| 207.9 | 197 | 191.4 | 164.2 | 2 | 2 | 2 | 2 | 0.174 |
| | | | | | | | | 0.136 |
| 67 | 61.6 | 58.8 | 48.8 | 1 | 1 | 1 | 1 | 0.105 |
| | | | | | | | | 0.039 |
| | | | | | | | | 0.061 |
| 149.8 | 93.1 | 98.9 | 121.8 | 1 | 1 | 1 | 1 | 0.039 |
| 30.1 | 25.1 | 25.1 | 21.4 | 1 | 1 | 1 | 1 | 6.499 |
| 146.6 | 93.2 | 112.9 | 120.5 | 2 | 2 | 2 | 2 | 0.298 |
| 360.7 | 333.3 | 315.8 | 280.4 | 6 | 6 | 6 | 6 | 2.831 |
| | | | | | | | | 0.04 |
| 92.5 | 85.8 | 88.8 | 80.3 | 1 | 1 | 1 | 1 | 0.105 |
| 92.6 | 65 | 54.2 | 58.8 | 1 | 1 | 1 | 1 | 3.217 |
| 58 | 68.4 | 51.5 | 49.3 | 1 | 1 | 1 | 1 | 0.122 |
| 3301.9 | 2723.6 | 2864.5 | 2647.8 | 46 | 46 | 46 | 46 | 2.918 |
| | | | | | | | | 0.129 |
| 137.8 | 123.5 | 126.7 | 95.6 | 2 | 2 | 2 | 2 | 0.748 |
| 108.6 | 81.9 | 81.2 | 88.3 | 2 | 2 | 2 | 2 | 0.195 |
| | | | | | | | | 0.122 |
| | | | | | | | | 0.778 |
| 165.3 | 99.4 | 113.9 | 118.7 | 1 | 1 | 1 | 1 | 0.468 |
| 121.2 | 120.5 | 101.4 | 105.5 | 1 | 1 | 1 | 1 | 0.122 |
| | | | | | | | | 0.274 |
| 146.9 | 112 | 115.7 | 108.7 | 2 | 2 | 2 | 2 | 0.098 |
| | | | | | | | | 0.116 |

| | | | | | | | | |
|---|---|---|---|---|---|---|---|---|
| | | | | | | | | 0.108 |
| 100.1 | 76.3 | 79.5 | 77.4 | 1 | 1 | 1 | 1 | 0.334 |
| 13.9 | 9.7 | 9.3 | 9.8 | 1 | 1 | 1 | 1 | 0.778 |
| 68.5 | 57.5 | 48.5 | 53.7 | 1 | 1 | 1 | 1 | 0.202 |
| | | | | | | | | 0.145 |
| 14.7 | 8.4 | 11.9 | 9 | 1 | 1 | 1 | 1 | 0.122 |
| 86.1 | 71.9 | 65.1 | 57.9 | 1 | 1 | 1 | 1 | 0.109 |
| | | | | | | | | 0.259 |
| 61.3 | 45.4 | 51.7 | 43.9 | 1 | 1 | 1 | 1 | 0.389 |
| | | | | | | | | 0.11 |
| 54.1 | 45.4 | 47.7 | 52.2 | 1 | 1 | 1 | 1 | 0.468 |
| 127.6 | 111.6 | 120.7 | 100.7 | 1 | 1 | 1 | 1 | 0.292 |
| 49.9 | 43.7 | 55.3 | 51.6 | 1 | 1 | 1 | 1 | 0.292 |
| 385.2 | 309.7 | 314.4 | 285.5 | 4 | 4 | 4 | 4 | 0.182 |
| | | | | | | | | 0.058 |
| | | | | | | | | 0.07 |
| 94.3 | 92.6 | 83 | 78.5 | 1 | 1 | 1 | 1 | 0.125 |
| | | | | | | | | 0.046 |
| 103 | 69.6 | 83.9 | 81.9 | 1 | 1 | 1 | 1 | 0.068 |
| | | | | | | | | 0.058 |
| 421.3 | 370.3 | 332.6 | 296.4 | 4 | 4 | 4 | 4 | 0.216 |
| | | | | | | | | 0.053 |
| | | | | | | | | 0.046 |
| 58.7 | 44.2 | 42.3 | 39.3 | 1 | 1 | 1 | 1 | 0.259 |
| | | | | | | | | 0.039 |
| | | | | | | | | 0.066 |
| | | | | | | | | 0.081 |
| 214.7 | 165.5 | 176.1 | 145.4 | 3 | 3 | 3 | 3 | 0.106 |
| 101.2 | 108.9 | 105 | 91 | 2 | 2 | 2 | 2 | 4.337 |
| 143.9 | 121 | 108 | 101.7 | 1 | 1 | 1 | 1 | 0.202 |
| | | | | | | | | 0.11 |
| 114.4 | 80.2 | 74.1 | 75.9 | 1 | 1 | 1 | 1 | 0.585 |
| 146.9 | 113.9 | 125.1 | 126.7 | 2 | 2 | 2 | 2 | 0.374 |
| 148.5 | 117.7 | 93.3 | 101.3 | 2 | 2 | 2 | 2 | 0.259 |
| 71 | 55.9 | 68.4 | 57.3 | 1 | 1 | 1 | 1 | 0.172 |
| 174.3 | 196.4 | 173 | 144.6 | 4 | 4 | 4 | 4 | 0.859 |
| 38 | 35.1 | 35.6 | 29.7 | 1 | 1 | 1 | 1 | 0.778 |
| 119.6 | 113.3 | 90 | 95.7 | 2 | 2 | 2 | 2 | 1.081 |
| 45.6 | 39.9 | 42.4 | 35.6 | 1 | 1 | 1 | 1 | 0.253 |
| | | | | | | | | 0.07 |
| | | | | | | | | 0.061 |
| | | | | | | | | 0.122 |
| 256.3 | 146 | 112 | 114.4 | 2 | 2 | 2 | 2 | 1.154 |
| | | | | | | | | 0.062 |
| 220.3 | 224 | 207.5 | 190.3 | 2 | 2 | 2 | 2 | 0.075 |
| 188.9 | 133.3 | 149 | 128.1 | 1 | 1 | 1 | 1 | 0.179 |
| 67.8 | 79 | 55.8 | 55.8 | 3 | 3 | 3 | 3 | 0.701 |
| 71.4 | 63.4 | 46.8 | 52.1 | 1 | 1 | 1 | 1 | 0.033 |
| 105.5 | 87.5 | 91.1 | 73 | 2 | 2 | 2 | 2 | 0.245 |
| 72.9 | 50.7 | 52 | 49.2 | 1 | 1 | 1 | 1 | 0.425 |
| | | | | | | | | 0.122 |
| 443.4 | 343.4 | 381.1 | 373.1 | 4 | 4 | 4 | 4 | 2.36 |
| | | | | | | | | 0.02 |
| 9.5 | 6.2 | 3.8 | 7.6 | 1 | 1 | 1 | 1 | 0.585 |
| 64.9 | 50.6 | 49.5 | 47.4 | 1 | 1 | 1 | 1 | 0.072 |
| | | | | | | | | 0.08 |
| 203.9 | 169.5 | 166.5 | 144.7 | 3 | 3 | 3 | 3 | 0.931 |
| 96 | 91 | 80.9 | 78 | 1 | 1 | 1 | 1 | 0.148 |
| 47.7 | 48.1 | 44 | 43.7 | 1 | 1 | 1 | 1 | 0.172 |
| 324.1 | 322.8 | 281.6 | 250.7 | 4 | 4 | 4 | 4 | 0.778 |
| | | | | | | | | 0.042 |
| | | | | | | | | 0.066 |

| | | | | | | | | |
|---|---|---|---|---|---|---|---|---|
| 42.8 | 31.2 | 34.2 | 30.3 | 1 | 1 | 1 | 1 | 0.11 |
| | | | | | | | | 0.04 |
| 2263.6 | 1943.7 | 1825.6 | 1654.1 | 20 | 20 | 20 | 20 | 18.307 |
| | | | | | | | | 1.154 |
| | | | | | | | | 0.045 |
| 261.1 | 210.4 | 217.3 | 190.7 | 4 | 4 | 4 | 4 | 0.572 |
| | | | | | | | | 0.585 |
| 18.8 | 13.8 | 13 | 12.4 | 1 | 1 | 1 | 1 | 0.116 |
| 92.5 | 74.6 | 66.6 | 68.7 | 1 | 1 | 1 | 1 | 0.136 |
| 161 | 114.3 | 117.3 | 103.8 | 1 | 1 | 1 | 1 | 0.155 |
| 179.9 | 176.8 | 164.7 | 165.7 | 5 | 5 | 5 | 5 | 0.896 |
| | | | | | | | | 0.098 |
| 177.4 | 166.4 | 171.3 | 130.1 | 4 | 4 | 4 | 4 | 1.818 |
| 134.5 | 104.4 | 99 | 102.4 | 1 | 1 | 1 | 1 | 0.067 |
| | | | | | | | | 0.089 |
| 4.4 | 6 | 4.4 | 5.3 | 1 | 1 | 1 | 1 | 0.322 |
| | | | | | | | | 0.334 |
| | | | | | | | | 0.334 |
| 7.7 | 10.3 | 8.9 | 8.3 | 1 | 1 | 1 | 1 | 0.245 |
| | | | | | | | | 0.122 |
| | | | | | | | | 0.062 |
| | | | | | | | | 0.044 |
| | | | | | | | | 0.141 |
| | | | | | | | | 0.059 |
| 98.6 | 85.1 | 86.6 | 71.1 | 1 | 1 | 1 | 1 | 0.024 |
| 80.9 | 57 | 41.2 | 46.9 | 1 | 1 | 1 | 1 | 0.122 |
| 1256.9 | 974.4 | 805.6 | 792.6 | 14 | 14 | 14 | 14 | 2.652 |
| | | | | | | | | 0.129 |
| | | | | | | | | 0.11 |
| | | | | | | | | 0.018 |
| 163.7 | 160.5 | 148.9 | 128.9 | 2 | 2 | 2 | 2 | 0.995 |
| 111.9 | 102.3 | 89.5 | 86.7 | 3 | 3 | 3 | 3 | 0.141 |
| 270.5 | 236.4 | 214 | 202.6 | 4 | 4 | 4 | 4 | 1.637 |
| | | | | | | | | 0.038 |
| | | | | | | | | 0.194 |
| | | | | | | | | 0.083 |
| | | | | | | | | 0.055 |
| 124.9 | 90.5 | 94.3 | 81.6 | 1 | 1 | 1 | 1 | 0.166 |
| | | | | | | | | 0.116 |
| | | | | | | | | 0.116 |
| 119.3 | 114.5 | 94 | 97.4 | 1 | 1 | 1 | 1 | 0.051 |
| | | | | | | | | 0.122 |
| 24.5 | 19 | 18.4 | 16.4 | 1 | 1 | 1 | 1 | 0.233 |
| 58.1 | 57.1 | 50.9 | 45.5 | 1 | 1 | 1 | 1 | 0.047 |
| 121.1 | 98 | 107.1 | 91.3 | 1 | 1 | 1 | 1 | 0.194 |
| 67.3 | 52.4 | 49.5 | 46.3 | 1 | 1 | 1 | 1 | 0.292 |
| 427.2 | 367.6 | 301.8 | 305.4 | 5 | 5 | 5 | 5 | 0.487 |
| | | | | | | | | 0.129 |
| 27.9 | 25.4 | 24.5 | 23 | 1 | 1 | 1 | 1 | 0.585 |
| 128.3 | 96.1 | 117.8 | 106.3 | 1 | 1 | 1 | 1 | 0.425 |
| 413.5 | 324.5 | 346.5 | 358 | 6 | 6 | 6 | 6 | 5.813 |
| 27.3 | 22.6 | 25 | 17.6 | 1 | 1 | 1 | 1 | 0.73 |
| 4.7 | 5.4 | 6.7 | 6 | 1 | 1 | 1 | 1 | 0.468 |
| 481.3 | 372.6 | 367.8 | 341.2 | 6 | 6 | 6 | 6 | 0.817 |
| | | | | | | | | 0.292 |
| 41.5 | 49.1 | 41.9 | 37.8 | 1 | 1 | 1 | 1 | 0.778 |
| 124.8 | 91.9 | 72.8 | 67.1 | 1 | 1 | 1 | 1 | 0.133 |
| 603.4 | 625.6 | 549.8 | 462.9 | 5 | 5 | 5 | 5 | 1.581 |

| Score Sequest HT | # Peptides Sequest HT | Score Mascot | # Peptides Mascot |
|---|---|---|---|
| 3.53699303 | 1 | 60.96 | 1 |
| 73.1875775 | 12 | 511.658601 | 13 |
| 121.489293 | 19 | 560.144068 | 18 |
| 2.57188654 | 2 | 17.26 | 2 |
| 2.75617385 | 1 | 0 | 1 |
| 0 | 1 | 0 | 1 |
| 7.90604734 | 3 | 36.49 | 3 |
| 3.88024259 | 1 | 48.69 | 1 |
| 134.152901 | 23 | 960.116518 | 23 |
| 49.1202271 | 11 | 147.094639 | 10 |
| 4.77476645 | 1 | 61.41 | 1 |
| 11.0043051 | 4 | 87.3340023 | 4 |
| 43.7470155 | 11 | 320.316662 | 9 |
| 0 | 1 | 0 | 1 |
| 10.4321461 | 3 | 60.2933198 | 3 |
| 2.80789876 | 1 | 0 | 1 |
| 5.09457684 | 1 | 65.75 | 1 |
| 2.40065765 | 1 | 19.33 | 1 |
| 3.85539913 | 1 | 55.94 | 1 |
| 13.9861975 | 3 | 54.2252772 | 3 |
| 114.488229 | 21 | 487.924376 | 21 |
| 34.6972246 | 9 | 159.34539 | 8 |
| 48.0457356 | 10 | 245.353278 | 9 |
| 44.5355365 | 10 | 156.417001 | 9 |
| 2.42937613 | 1 | 37.97 | 1 |
| 8.08201957 | 2 | 17.85 | 2 |
| 11.9094455 | 4 | 85.6454776 | 4 |
| 17.9704585 | 5 | 145.03 | 5 |
| 73.6678152 | 16 | 360.417774 | 16 |
| 12.5964608 | 3 | 69.8246168 | 3 |
| 3.68484473 | 1 | 0 | 1 |
| 73.3107007 | 12 | 346.767394 | 12 |
| 22.2566237 | 3 | 153.285049 | 3 |
| 26.5054328 | 6 | 173.784287 | 6 |
| 3.27926588 | 1 | 45.25 | 1 |
| 26.2382903 | 5 | 245.489923 | 5 |
| 6.98862553 | 2 | 74.3278071 | 2 |
| 28.1419604 | 8 | 120.385714 | 8 |
| 19.1648884 | 4 | 143.429014 | 3 |
| 2.9705565 | 2 | 0 | 2 |
| 20.6890037 | 6 | 53.8 | 6 |
| | | 40.55 | 1 |
| 3.83657241 | 1 | 48.98 | 1 |
| 3.73275948 | 1 | 44.77 | 1 |
| 18.2505227 | 5 | 62.16 | 5 |
| 2.6342411 | 1 | 55.37 | 1 |
| 2.42821741 | 1 | 50.16 | 1 |
| 10.6748865 | 2 | 36.3 | 2 |
| 65.6430125 | 11 | 566.628776 | 11 |
| 75.7967153 | 14 | 342.704673 | 13 |
| 6.58941126 | 1 | 100.98 | 1 |
| 2.47812939 | 1 | 0 | 1 |
| 12.2073727 | 3 | 56.09 | 3 |
| 0 | 1 | | |
| 0 | 1 | 14.55 | 1 |
| 11.5985718 | 3 | 107.47 | 3 |
| 4.71577215 | 1 | 83.32 | 1 |

| | | | |
|---|---|---|---|
| 30.282975 | 5 | 173.369962 | 5 |
| 55.6010778 | 11 | 270.249503 | 11 |
| 7.69213033 | 2 | 108.37 | 2 |
| 2.55750299 | 1 | 0 | 1 |
| 3.15458608 | 1 | 38.88 | 1 |
| 4.61023521 | 2 | 0 | 2 |
| 2.21366811 | 1 | 0 | 1 |
| 2.43969464 | 1 | 24.58 | 1 |
| 4.27128458 | 1 | 35.86 | 1 |
| 5.64175868 | 2 | 38.89 | 2 |
| 2.2365036 | 1 | 0 | 1 |
| 8.35992956 | 3 | 52.4365428 | 3 |
| 2.51163507 | 1 | 37.68 | 1 |
| 8.74416661 | 4 | 73.7866667 | 3 |
| 0 | 1 | | |
| 5.45207143 | 2 | 25.52 | 2 |
| 2.82322001 | 2 | 27.03 | 1 |
| 2.81909108 | 1 | 46.89 | 1 |
| 10.9127631 | 2 | 95.816957 | 2 |
| 8.89739203 | 4 | 41.52 | 4 |
| 12.4936953 | 6 | 63.1228098 | 6 |
| 1.90903628 | 1 | 18.63 | 1 |
| 30.821682 | 7 | 107.384726 | 6 |
| 400.1854 | 29 | 1980.95513 | 28 |
| 13.8233612 | 3 | 102.818848 | 3 |
| 55.4710665 | 11 | 325.131637 | 10 |
| 8.86595392 | 1 | 49.678372 | 1 |
| 45.9360381 | 9 | 221.624485 | 9 |
| 49.7720183 | 10 | 196.842897 | 10 |
| 3.48971009 | 1 | 0 | 1 |
| 3.15886879 | 1 | 31.89 | 1 |
| 6.08948278 | 2 | 106.8 | 2 |
| 3.29111886 | 1 | 54.44 | 1 |
| 4.02664804 | 1 | 33.56 | 1 |
| 6.35259104 | 3 | 71.86 | 3 |
| 3.02072334 | 1 | 43.09 | 1 |
| 3.48158169 | 2 | 0 | 1 |
| 0 | 1 | 0 | 1 |
| 2.34801769 | 1 | 22.03 | 1 |
| 3.9575119 | 1 | 48.16 | 1 |
| 7.48052621 | 3 | 41.74 | 3 |
| 5.21594977 | 2 | 43.96 | 2 |
| 6.09808755 | 2 | 22.22 | 2 |
| 3.95455575 | 2 | 47.68 | 1 |
| 8.44456244 | 2 | 40.73 | 2 |
| 0 | 1 | 0 | 1 |
| 15.1414423 | 3 | 163.07 | 3 |
| 2.08960366 | 1 | 0 | 1 |
| 1.87123108 | 1 | 30.45 | 1 |
| 19.042382 | 4 | 115.791393 | 4 |
| 2.39418459 | 1 | 31.95 | 1 |
| 27.6315136 | 8 | 142.042521 | 8 |
| 2.893188 | 2 | 46.88 | 1 |
| 9.04071903 | 3 | 63.1852257 | 3 |
| 2.70948029 | 1 | 0 | 1 |
| 12.2711847 | 3 | 97.0034536 | 3 |
| 2.56001401 | 1 | 40.42 | 1 |
| 5.44182158 | 2 | 37.86 | 2 |
| 8.80810809 | 3 | 114.806667 | 3 |
| 5.33711362 | 1 | 51.91 | 1 |
| 6.56534815 | 2 | 61.33 | 2 |
| 48.0973594 | 12 | 272.457287 | 12 |

| | | | |
|---|---|---|---|
| 18.3460836 | 2 | 194.992608 | 2 |
| 20.1779065 | 6 | 121.496344 | 6 |
| 2.20638323 | 3 | 0 | 3 |
| 23.4274688 | 5 | 180.316188 | 5 |
| 17.2013736 | 3 | 83.31 | 3 |
| 18.9238079 | 6 | 121.944732 | 6 |
| 9.42574263 | 2 | 25.98 | 2 |
| 20.4263525 | 5 | 166.369149 | 2 |
| 2.06905127 | 1 | 0 | 1 |
| 23.5608251 | 7 | 133.285318 | 6 |
| 2.40320992 | 1 | 28.77 | 1 |
| 25.481503 | 5 | 153.892324 | 5 |
| 19.0050496 | 5 | 59.3474283 | 4 |
| 79.7683327 | 19 | 327.435516 | 18 |
| 2.4869349 | 1 | 0 | 1 |
| 5.00884891 | 3 | 0 | 2 |
| 6.40645194 | 2 | 39.68 | 2 |
| 403.214079 | 31 | 1917.75964 | 30 |
| 20.5434897 | 6 | 112.984222 | 6 |
| 0 | 1 | 0 | 1 |
| 4.70782042 | 1 | 90.83 | 1 |
| 9.83839297 | 2 | 73.3420987 | 2 |
| 16.9474192 | 4 | 102.67245 | 4 |
| 7.34152961 | 2 | 68.21 | 2 |
| 3.02577066 | 2 | 48.2 | 1 |
| 13.9782922 | 4 | 94.9807821 | 4 |
| 3.96428108 | 1 | 0 | 1 |
| 8.89656115 | 1 | 38.66 | 1 |
| 2.00055218 | 1 | 29.98 | 1 |
| 10.6240854 | 5 | 40.3634203 | 5 |
| 2.18311214 | 1 | 0 | 1 |
| 0 | 1 | 0 | 1 |
| 0 | 1 | 0 | 1 |
| 6.05424905 | 2 | 72.27 | 1 |
| 120.117494 | 13 | 877.589049 | 13 |
| 2.16682243 | 1 | 35.01 | 1 |
| 0 | 1 | | |
| 2.58264518 | 2 | 35.14 | 2 |
| 32.6000198 | 9 | 165.366241 | 8 |
| 44.5376035 | 6 | 252.397154 | 6 |
| 32.5490234 | 3 | 243.546224 | 3 |
| 11.1842189 | 2 | 64.6914358 | 2 |
| 2.43647385 | 1 | | |
| | | 54.63 | 1 |
| 60.6714964 | 16 | 281.530551 | 15 |
| 12.2231092 | 3 | 41.71 | 3 |
| 9.19934011 | 4 | 33.5 | 4 |
| 6.68093824 | 2 | 49.59 | 2 |
| 49.0750935 | 9 | 205.107339 | 9 |
| 40.1246562 | 11 | 267.157916 | 11 |
| 9.79277754 | 3 | 81.7040737 | 3 |
| 2.67095399 | 1 | 0 | 1 |
| 2.22463799 | 1 | 0 | 1 |
| 7.44691658 | 2 | 45.8311961 | 2 |
| 2.70063496 | 1 | 0 | 1 |
| 3.52487755 | 1 | 69.28 | 1 |
| 3.77079868 | 1 | 31.54 | 1 |
| 5.65457129 | 2 | 35.04 | 2 |
| 0 | 1 | 0 | 1 |
| 6.82848978 | 3 | 60.92 | 2 |
| 1.83999085 | 1 | 17.17 | 1 |
| 6.43935633 | 1 | 76.45 | 1 |

| | | | |
|---|---|---|---|
| 2.0960865 | 1 | 0 | 1 |
| 5.92907763 | 2 | 69.87 | 2 |
| 2.21618652 | 1 | 0 | 1 |
| 10.4148436 | 2 | 60.4175634 | 1 |
| 3.11774015 | 1 | 25.02 | 1 |
| 13.0916588 | 4 | 102.198493 | 4 |
| 4.35331774 | 1 | 40.49 | 1 |
| 8.33723092 | 2 | 91.548906 | 2 |
| 4.38167286 | 1 | 0 | 1 |
| 2.10198665 | 1 | 0 | 1 |
| 5.08957362 | 2 | 0 | 2 |
| 13.2638168 | 6 | 62.3159212 | 4 |
| 10.6653686 | 3 | 117.998025 | 2 |
| 2.44128871 | 1 | 0 | 1 |
| 3.98574209 | 1 | 63.15 | 1 |
| 20.3339314 | 5 | 100.017107 | 5 |
| 1.68455076 | 2 | 0 | 1 |
| 9.63171387 | 2 | 72.2398707 | 2 |
| 4.24776125 | 1 | 52.98 | 1 |
| 3.65481901 | 1 | 55.25 | 1 |
| 2.1899209 | 1 | 16.54 | 1 |
| 53.787323 | 8 | 321.04602 | 8 |
| 6.62213635 | 4 | 37.14 | 3 |
| 6.49764919 | 1 | 81.05 | 1 |
| 2.912817 | 2 | 0 | 1 |
| 62.1640429 | 8 | 141.350165 | 8 |
| 2.94978809 | 1 | 35.23 | 1 |
| 8.22054195 | 3 | 38.39 | 3 |
| 3.55210209 | 1 | 31.58 | 1 |
| 3.14465308 | 2 | 0 | 1 |
| 5.26351428 | 1 | 58.01 | 1 |
| 6.82039452 | 1 | 0 | 1 |
| 11.1642065 | 4 | 86.9934205 | 4 |
| 5.17461061 | 1 | 33.95 | 1 |
| 2.1581459 | 1 | 0 | 1 |
| 23.6152947 | 7 | 172.305714 | 7 |
| 28.7734344 | 7 | 113.545807 | 7 |
| 4.33255577 | 1 | 57.58 | 1 |
| 7.60634089 | 1 | 0 | 1 |
| 3.33388686 | 1 | 0 | 1 |
| 1.81383026 | 1 | | |
| 3.35680914 | 1 | 47.35 | 1 |
| 13.1940463 | 3 | 138.284619 | 3 |
| 45.4434786 | 11 | 362.262069 | 11 |
| 5.16639566 | 1 | 0 | 1 |
| 3.30178666 | 1 | 27.85 | 1 |
| 3.96171737 | 1 | 0 | 1 |
| 17.7904205 | 4 | 52.1473111 | 4 |
| 11.68134 | 2 | 94.5685147 | 2 |
| 3.54384732 | 1 | 56.2 | 1 |
| 5.40814924 | 2 | 67.33 | 2 |
| 16.4103982 | 4 | 184.633333 | 4 |
| 150.749125 | 14 | 588.769186 | 15 |
| 45.0212142 | 14 | 255.170477 | 13 |
| 2.88107514 | 1 | 34.33 | 1 |
| 3.33595061 | 1 | 0 | 1 |
| 10.1544516 | 3 | 50.9322916 | 2 |
| 0 | 1 | 0 | 1 |
| 3.76692295 | 1 | 47.57 | 1 |
| 3.37002444 | 1 | 21.9 | 1 |
| 0 | 1 | 0 | 1 |
| 2.67303348 | 1 | 0 | 1 |

| | | | |
|---|---|---|---|
| 6.58616149 | 2 | 70.67 | 2 |
| 10.1355257 | 3 | 41.58 | 3 |
| 6.17988777 | 3 | 48.52 | 2 |
| 11.2405624 | 3 | 52.15 | 3 |
| 3.80641079 | 1 | 43.03 | 1 |
| 25.0498605 | 2 | 0 | 2 |
| 50.1066079 | 9 | 321.619477 | 9 |
| 77.2207172 | 16 | 392.165938 | 16 |
| 3.70499468 | 2 | 44.11 | 2 |
| 8.81126738 | 3 | 60.9439027 | 3 |
| 8.17932963 | 1 | 91.2112742 | 1 |
| 2.77713895 | 1 | 48.89 | 1 |
| 4.09869432 | 1 | 86.9464223 | 2 |
| 5.68294668 | 1 | 57.99 | 1 |
| 3.5312736 | 1 | 0 | 1 |
| 180.032695 | 22 | 1045.98534 | 21 |
| 12.2570152 | 3 | 74.6734805 | 3 |
| 258.005616 | 24 | 1692.12543 | 23 |
| 4.66800022 | 1 | 0 | 1 |
| 1.98327625 | 1 | 33.28 | 1 |
| 4.33539963 | 2 | 39.6299213 | 2 |
| 7.68204069 | 2 | 52.3352051 | 2 |
| 2.01601982 | 1 | 0 | 1 |
| 11.8591511 | 3 | 58.9807418 | 3 |
| 26.4445925 | 6 | 162.801246 | 6 |
| 20.1803336 | 3 | 171.853257 | 3 |
| 40.8349886 | 10 | 389.014669 | 10 |
| 32.0200691 | 9 | 261.945391 | 8 |
| 70.5917311 | 11 | 522.852067 | 11 |
| 13.9432416 | 4 | 45.5508048 | 4 |
| 7.06713176 | 2 | 46.18 | 2 |
| 0 | 1 | 0 | 1 |
| 10.9037955 | 3 | 110.085194 | 3 |
| 8.28865242 | 3 | 62.74 | 3 |
| 1.99611962 | 1 | 31.54 | 1 |
| 3.51020145 | 1 | 0 | 1 |
| 14.67729 | 3 | 87.490208 | 3 |
| 2.2962954 | 1 | 32.53 | 1 |
| 16.0401757 | 5 | 133.97891 | 5 |
| 3.76519728 | 1 | 39.35 | 1 |
| 115.242702 | 19 | 455.672989 | 19 |
| 24.012569 | 5 | 178.371243 | 4 |
| 7.46752882 | 1 | 76.03 | 1 |
| 18.5656508 | 6 | 83.2443059 | 6 |
| 6.60924935 | 2 | 47.7616819 | 2 |
| 94.4088023 | 22 | 517.000709 | 21 |
| 0 | 1 | 0 | 1 |
| 0 | 2 | 0 | 1 |
| 12.4998643 | 4 | 113.016667 | 4 |
| 0 | 1 | 0 | 1 |
| 30.1134799 | 5 | 30.88 | 5 |
| 34.688549 | 8 | 301.318533 | 8 |
| 6.90378916 | 2 | 49.57 | 2 |
| 6.13728595 | 2 | 50.42 | 2 |
| 157.293605 | 25 | 1145.10619 | 24 |
| 2.33609772 | 1 | 42.09 | 1 |
| 13.7174686 | 5 | 59.3474283 | 4 |
| 31.2691813 | 5 | 157.577346 | 5 |
| 13.0820513 | 2 | 161.673333 | 2 |
| 8.96527195 | 1 | 57.35 | 1 |
| 14.3613491 | 5 | 69.9949824 | 5 |
| 6.77031684 | 3 | 59.44 | 3 |

| | | | |
|---|---|---|---|
| 15.2632688 | 4 | 92.7761653 | 4 |
| 65.5098556 | 14 | 366.797705 | 14 |
| 118.409423 | 12 | 394.526472 | 11 |
| 2.73965025 | 1 | 22.49 | 1 |
| 2.72559953 | 1 | 38.74 | 1 |
| 4.25047112 | 1 | 77.8 | 1 |
| 4.14062119 | 1 | 47.73 | 1 |
| 68.9425607 | 12 | 514.339215 | 11 |
| 3.22901511 | 1 | 55.45 | 1 |
| 2.9632473 | 1 | 0 | 1 |
| 2.55591965 | 1 | 0 | 1 |
| 2.88426828 | 1 | 33.75 | 1 |
| 2.95528579 | 1 | 38.85 | 1 |
| 5.21354318 | 3 | 34.04 | 3 |
| 3.88885665 | 1 | 87.79 | 1 |
| 2.69599485 | 1 | 35.5 | 1 |
| 25.6167288 | 8 | 260.247731 | 8 |
| 7.02529907 | 2 | 75.8 | 2 |
| 16.0559826 | 4 | 105.07859 | 4 |
| 2.54517388 | 1 | 38.17 | 1 |
| 2.17866278 | 1 | 0 | 1 |
| 2.55758309 | 1 | 41.9 | 1 |
| 4.98047066 | 1 | 55.73 | 1 |
| 25.3867829 | 6 | 91.0412403 | 6 |
| 0 | 1 | 20.99 | 1 |
| 4.66186213 | 2 | 0 | 2 |
| 81.3504071 | 20 | 462.659878 | 19 |
| 11.8779571 | 4 | 33.9595781 | 4 |
| 2.74486613 | 1 | 0 | 1 |
| 5.28455234 | 1 | 65.6 | 1 |
| 9.85870504 | 3 | 64.8481508 | 3 |
| 9.14233017 | 1 | 71.17 | 1 |
| 3.06466365 | 1 | 0 | 1 |
| 8.82704091 | 2 | 35.32 | 2 |
| 391.239004 | 28 | 1909.70599 | 27 |
| 386.957585 | 30 | 1738.7131 | 29 |
| 17.7802444 | 5 | 105.378581 | 5 |
| 3.06870246 | 2 | 45.78 | 1 |
| 2.75147343 | 2 | 0 | 2 |
| 6.07850075 | 2 | 37.43 | 2 |
| 0 | 1 | | |
| 101.32837 | 23 | 607.365752 | 21 |
| 1.93069148 | 1 | 0 | 1 |
| 8.26288867 | 3 | 77.7500047 | 3 |
| 4.28892303 | 3 | 0 | 3 |
| 254.538221 | 28 | 1683.14942 | 28 |
| 2.66873217 | 1 | 36.17 | 1 |
| 16.8619118 | 7 | 95.8168969 | 6 |
| 17.2252977 | 5 | 120.099985 | 5 |
| 95.9528803 | 28 | 428.723364 | 25 |
| 4.06984615 | 1 | 28.98 | 1 |
| 2.62873769 | 1 | 0 | 1 |
| 0 | 1 | | |
| 12.2668555 | 2 | 120.368598 | 2 |
| 7.19589877 | 2 | 79.6 | 2 |
| 16.1877601 | 5 | 105.46408 | 5 |
| 16.642905 | 5 | 127.812343 | 5 |
| 24.3342371 | 7 | 62.8229764 | 7 |
| 2.79855585 | 1 | 41.31 | 1 |
| 0 | 1 | | |
| 53.3797781 | 10 | 318.318791 | 9 |
| 13.299984 | 4 | 82.3 | 4 |

| | | | |
|---|---|---|---|
| 6.51298714 | 3 | 0 | 2 |
| 16.9440911 | 4 | 146.155261 | 4 |
| 2.30675316 | 1 | 45.17 | 1 |
| | | 0 | 1 |
| 0 | 1 | 23.59 | 1 |
| 5.4759872 | 2 | 45.7211974 | 2 |
| 2.81260109 | 1 | 35.07 | 1 |
| 8.13844585 | 3 | 61.9102832 | 3 |
| 3.16543031 | 1 | 43.18 | 1 |
| 5.5927825 | 1 | 73.9 | 1 |
| 2.74866962 | 1 | 38.2 | 1 |
| 4.01180077 | 1 | 47.13 | 1 |
| 3.50552368 | 1 | 51.1 | 1 |
| 14.6695168 | 2 | 131.147586 | 2 |
| 23.1784697 | 5 | 166.12 | 5 |
| 26.0428278 | 5 | 226.262554 | 5 |
| 23.7689557 | 9 | 98.6234982 | 7 |
| 6.37647986 | 2 | 33.12 | 2 |
| 7.5170033 | 2 | 33.02 | 2 |
| 10.2769256 | 3 | 78.9475207 | 3 |
| 16.0467775 | 6 | 87.754825 | 6 |
| 4.88989735 | 2 | 25.95 | 2 |
| 2.10507536 | 2 | 0 | 2 |
| 7.92532539 | 3 | 65.01 | 3 |
| 195.466247 | 30 | 1203.33995 | 30 |
| 3.62199926 | 1 | 0 | 1 |
| 3.7827034 | 1 | 45.04 | 1 |
| 3.11726499 | 1 | 45.42 | 1 |
| 286.926837 | 27 | 1479.49614 | 26 |
| 3.14810038 | 1 | 42.13 | 1 |
| 3.2278223 | 1 | 44.11 | 1 |
| 3.9304297 | 1 | 0 | 1 |
| 4.90943861 | 1 | 46.95 | 1 |
| 7.17974663 | 2 | 83.4652051 | 2 |
| 3.0818224 | 1 | 38.77 | 1 |
| 7.70312309 | 2 | 90.9161177 | 2 |
| 9.56422043 | 1 | 113.680709 | 1 |
| 3.15602612 | 1 | 26.46 | 1 |
| 38.1797786 | 7 | 153.7029 | 7 |
| 0 | 1 | 0 | 1 |
| 0 | 1 | | |
| 2.65673757 | 1 | 43.16 | 1 |
| 13.084717 | 6 | 60.83 | 5 |
| 14.3299561 | 4 | 115.096007 | 4 |
| 4.73780441 | 2 | 55.5 | 2 |
| 3.1365068 | 1 | 42.7 | 1 |
| 15.7445478 | 3 | 98.1461414 | 3 |
| 4.89552879 | 1 | 34.24 | 1 |
| 8.92480206 | 2 | 85.58 | 2 |
| 15.4470041 | 5 | 102.519962 | 5 |
| 7.65757775 | 2 | 114.384121 | 2 |
| 0 | 1 | 0 | 1 |
| 3.14976072 | 2 | 0 | 1 |
| 6.43650913 | 2 | 0 | 2 |
| 2.48033905 | 1 | 32.06 | 1 |
| 39.5411272 | 8 | 196.408308 | 8 |
| 20.1495757 | 6 | 175.09059 | 5 |
| 5.92363167 | 2 | 55.26 | 2 |
| 13.0687082 | 4 | 23.74 | 4 |
| 8.61962318 | 3 | 35.4 | 3 |
| 12.7294376 | 2 | 146.199837 | 1 |
| 0 | 1 | 0 | 1 |

| | | | |
|---|---|---|---|
| 3.24687076 | 1 | 51.93 | 1 |
| 44.5771799 | 5 | 217.74544 | 5 |
| 9.76336002 | 3 | 47.05 | 3 |
| 3.81160784 | 1 | 0 | 1 |
| 33.8561161 | 6 | 158.58133 | 6 |
| 180.635407 | 26 | 897.711414 | 24 |
| 5.21416306 | 2 | 0 | 1 |
| 5.21751261 | 2 | 30.51 | 2 |
| 2.86990261 | 1 | 39.7 | 1 |
| 3.29890156 | 1 | 62.82 | 1 |
| 2.5280571 | 1 | 23.1 | 1 |
| 12.6345983 | 2 | 66.6961513 | 2 |
| 4.83693457 | 1 | 48.56 | 1 |
| 2.91385221 | 1 | 0 | 1 |
| 5.52059293 | 2 | 31.35 | 2 |
| 4.25199604 | 3 | 55.01 | 3 |
| 5.35641766 | 3 | 0 | 3 |
| 11.7899497 | 3 | 52.97 | 3 |
| 55.4069073 | 10 | 203.873876 | 10 |
| 28.3479314 | 7 | 215.29308 | 7 |
| 8.91337967 | 3 | 63.35 | 3 |
| 6.93721724 | 1 | 39.8367005 | 1 |
| 4.8140645 | 1 | 0 | 1 |
| 70.4738983 | 19 | 358.836156 | 19 |
| 3.27560997 | 2 | 0 | 2 |
| 12.0486937 | 3 | 33.06 | 3 |
| 3.66186118 | 1 | 0 | 1 |
| 4.98103476 | 1 | 45.29 | 1 |
| 154.586076 | 42 | 983.6406 | 41 |
| 4.32767916 | 1 | 38.28 | 1 |
| 42.3525763 | 12 | 122.377091 | 12 |
| 6.65193796 | 4 | 63.57 | 3 |
| 2.98320413 | 1 | 0 | 1 |
| 8.55214953 | 2 | 91.0926829 | 2 |
| 3.9082613 | 2 | 34.98 | 1 |
| 11.3276947 | 2 | 105.197357 | 2 |
| 0 | 1 | 0 | 1 |
| 2.73518777 | 1 | 0 | 1 |
| 5.30597782 | 1 | 66.1 | 1 |
| 12.3631206 | 3 | 148.560738 | 3 |
| 2.11016607 | 1 | 36.18 | 1 |
| 8.64366317 | 2 | 50.87 | 2 |
| 5.35109735 | 2 | 0 | 2 |
| 29.0511947 | 7 | 161.486679 | 7 |
| 2.46114874 | 1 | 0 | 1 |
| 2.63470483 | 1 | 35.21 | 1 |
| 70.1990242 | 17 | 425.867305 | 14 |
| 13.1723533 | 4 | 119.503333 | 4 |
| 8.74871159 | 3 | 101.81 | 2 |
| 6.59646583 | 2 | 39.53 | 2 |
| 4.27436209 | 1 | 67.15 | 1 |
| 65.3569643 | 8 | 301.16929 | 8 |
| 16.4312432 | 7 | 98.7640045 | 6 |
| 3.64736652 | 1 | 35.35 | 1 |
| 3.79927754 | 1 | 68.13 | 1 |
| 0 | 2 | 0 | 1 |
| 15.6718664 | 5 | 95.4733333 | 4 |
| 12.4571352 | 5 | 97.2292437 | 4 |
| 5.524647 | 3 | 30.03 | 3 |
| 0 | 1 | 0 | 1 |
| 2.93954539 | 1 | 0 | 1 |
| 55.4133651 | 20 | 179.205046 | 19 |

| | | | |
|---|---|---|---|
| 13.2092571 | 4 | 42.15 | 4 |
| 2.78359389 | 1 | 25.23 | 1 |
| 4.21073294 | 1 | 71.86 | 1 |
| 3.25289321 | 1 | 53.34 | 1 |
| 4.01944733 | 1 | 46.15 | 1 |
| 10.0883245 | 3 | 60.1499495 | 3 |
| 1.62325203 | 1 | 0 | 1 |
| 0 | 1 | | |
| 4.6170156 | 2 | 22.72 | 2 |
| 2.44351768 | 1 | 0 | 1 |
| 2.12607694 | 1 | 26.47 | 1 |
| 3.9303062 | 2 | 53.89 | 2 |
| 2.57186508 | 1 | 46.61 | 1 |
| 37.3366494 | 5 | 32.88 | 4 |
| 33.7039814 | 5 | 235.312552 | 5 |
| 9.21758366 | 2 | 94.05 | 2 |
| 1.70668709 | 1 | 27.59 | 1 |
| 2.81936622 | 1 | 37.6 | 1 |
| 8.78532314 | 1 | 86.95 | 1 |
| 2.33940101 | 2 | 30.04 | 2 |
| 2.35008264 | 1 | 0 | 1 |
| 6.17052245 | 2 | 79.6046978 | 2 |
| 5.70126271 | 3 | 52.18 | 3 |
| 38.8504651 | 7 | 287.779505 | 7 |
| 3.74127626 | 1 | 72.72 | 1 |
| 5.04333448 | 1 | 40.6 | 1 |
| 2.77479649 | 1 | 38.17 | 1 |
| 5.18166208 | 3 | 0 | 3 |
| 3.33947086 | 1 | 50.52 | 1 |
| 8.41821194 | 3 | 78.9847483 | 3 |
| 5.87243128 | 1 | 62.01 | 1 |
| 13.9656146 | 4 | 65.7026325 | 4 |
| 7.83421707 | 2 | 63.52 | 2 |
| 3.22927356 | 1 | 43.73 | 1 |
| 8.96978092 | 2 | 65.94 | 2 |
| 0 | 1 | | |
| 17.3518193 | 6 | 127.230822 | 6 |
| 15.724148 | 2 | 61.78 | 2 |
| 27.5409651 | 12 | 144.417944 | 12 |
| 2.54920363 | 1 | | |
| 0 | 1 | 29.94 | 1 |
| 45.667326 | 8 | 281.879491 | 8 |
| 37.1916733 | 10 | 254.156404 | 10 |
| 0 | 1 | | |
| 22.1160214 | 5 | 146.681896 | 5 |
| 3.62134838 | 1 | 0 | 1 |
| 24.4694283 | 4 | 219.512966 | 4 |
| 34.3737018 | 8 | 280.357439 | 8 |
| 3.01759171 | 1 | 31.53 | 1 |
| 3.33019733 | 1 | 58.51 | 1 |
| 36.3638778 | 7 | 234.711195 | 7 |
| 7.12911344 | 1 | 33.38 | 1 |
| 14.6619811 | 4 | 152.366173 | 4 |
| 25.9268243 | 8 | 195.832736 | 7 |
| 5.03329277 | 1 | 34 | 1 |
| 7.68251014 | 2 | 61.973103 | 2 |
| 25.0008481 | 7 | 126.138843 | 7 |
| 119.15906 | 20 | 545.779333 | 20 |
| 6.41195297 | 2 | 53.86 | 2 |
| 0 | 1 | 38.33 | 1 |
| 7.26214266 | 2 | 0 | 2 |
| 0 | 1 | 0 | 1 |

| | | | |
|---|---|---|---|
| 181.780799 | 24 | 790.683716 | 23 |
| 23.4814885 | 6 | 157.642392 | 6 |
| 3.05912638 | 1 | 43.09 | 1 |
| 90.1396413 | 16 | 541.404982 | 16 |
| 8.31810641 | 2 | 51.96 | 2 |
| 15.0569627 | 4 | 56.9481915 | 4 |
| 3.02548695 | 1 | 0 | 1 |
| 38.5388985 | 9 | 275.076414 | 9 |
| 42.5789607 | 11 | 171.544245 | 11 |
| 8.03782678 | 3 | 69.0412032 | 3 |
| 2.4093225 | 1 | 0 | 1 |
| 3.59664631 | 1 | 59.1 | 1 |
| 240.655669 | 36 | 1571.13096 | 33 |
| 4.04241371 | 1 | 36.57 | 1 |
| 3.13350749 | 1 | 45.53 | 1 |
| 7.3320868 | 2 | 34.09 | 2 |
| 2.98140454 | 1 | 23.49 | 1 |
| 17.3958364 | 6 | 173.033437 | 6 |
| 9.26894855 | 3 | 0 | 3 |
| 4.24954796 | 1 | 53.9 | 1 |
| 3.9692328 | 1 | 70.56 | 1 |
| 9.42395115 | 2 | 111.322429 | 2 |
| 0 | 1 | | |
| 27.968697 | 7 | 216.163688 | 7 |
| 2.08201003 | 1 | 35.27 | 1 |
| 3.94634676 | 1 | 40.81 | 1 |
| 5.13470197 | 3 | 0 | 3 |
| 19.8648367 | 6 | 148.659489 | 6 |
| 10.1273727 | 3 | 89.9675654 | 3 |
| 7.73286724 | 2 | 55.7557263 | 2 |
| 2.63435006 | 1 | 27.45 | 1 |
| 6.98855162 | 2 | 83.029946 | 2 |
| 2.52597284 | 1 | 0 | 1 |
| 18.1215377 | 5 | 83.6913763 | 5 |
| 6.05140209 | 2 | 0 | 2 |
| 3.21792912 | 1 | 66.82 | 1 |
| 6.96437979 | 2 | 48.02 | 2 |
| 2.56508827 | 1 | 30.91 | 1 |
| 14.5403841 | 4 | 94.9969927 | 4 |
| 3.87956309 | 1 | 58.81 | 1 |
| 0 | 1 | | |
| 29.2235222 | 9 | 104.032509 | 7 |
| 2.97381544 | 1 | 43.45 | 1 |
| 2.80292583 | 1 | 46.17 | 1 |
| 2.57120681 | 1 | 47.2 | 1 |
| 9.1952548 | 3 | 49.5775876 | 3 |
| 30.1888666 | 5 | 200.155486 | 5 |
| 3.1988523 | 1 | 0 | 1 |
| 2.48845315 | 2 | 41.58 | 2 |
| 0 | 1 | | |
| 3.59715366 | 3 | 45.88 | 3 |
| 3.64494467 | 1 | 35.97 | 1 |
| 69.4377855 | 13 | 343.832639 | 13 |
| 95.9842639 | 26 | 560.734075 | 26 |
| 2.78907037 | 1 | 42.46 | 1 |
| 2.21552777 | 1 | 21.53 | 1 |
| 2.82946873 | 1 | 59.03 | 1 |
| 7.95116496 | 3 | 77.25 | 3 |
| 32.1505203 | 7 | 209.016583 | 7 |
| 42.2325566 | 9 | 253.066474 | 9 |
| 4.01398659 | 1 | 0 | 1 |
| 2.78882694 | 2 | 0 | 2 |

| | | | |
|---|---|---|---|
| 3.02456617 | 1 | 0 | 1 |
| 8.87967634 | 2 | 51.1370028 | 2 |
| 4.71933508 | 1 | 50.24 | 1 |
| 3.13607645 | 1 | 0 | 1 |
| 26.7330523 | 8 | 170.107738 | 7 |
| 13.3015277 | 5 | 0 | 5 |
| 2.97344661 | 1 | 18.26 | 1 |
| 25.089051 | 7 | 188.0717 | 6 |
| | | 34.3 | 1 |
| 19.6181853 | 7 | 83.1533333 | 7 |
| 23.7590873 | 7 | 133.08 | 7 |
| 22.3359654 | 2 | 99.6730453 | 2 |
| 14.1221404 | 2 | 74.97 | 2 |
| 17.6862199 | 4 | 161.195229 | 4 |
| 48.3390965 | 12 | 368.047577 | 12 |
| 61.7547398 | 11 | 424.055236 | 11 |
| 230.955074 | 44 | 1193.45629 | 44 |
| 10.2705557 | 5 | 82.5133333 | 4 |
| 18.7868218 | 4 | 113.431854 | 4 |
| 8.41237688 | 2 | 80.39 | 2 |
| 2.16999507 | 1 | 0 | 1 |
| 11.0727873 | 1 | 132.97 | 1 |
| 3.10116053 | 1 | | |
| 2.07303333 | 1 | 0 | 1 |
| 8.82476354 | 2 | 94.7 | 2 |
| 17.0715616 | 6 | 93.8988699 | 6 |
| | | 34.44 | 1 |
| 2.55351996 | 1 | 31.18 | 1 |
| 3.99052215 | 1 | 65.31 | 1 |
| 9.85463381 | 3 | 85.54 | 3 |
| 2.31712723 | 1 | 27.93 | 1 |
| 32.8304305 | 8 | 124.51285 | 7 |
| 2.52061439 | 1 | 0 | 1 |
| 10.7252395 | 3 | 83.97 | 3 |
| 334.33264 | 28 | 1562.3789 | 28 |
| 88.799479 | 6 | 468.969864 | 6 |
| 0 | 1 | 0 | 1 |
| 5.32252026 | 2 | 39.09 | 2 |
| 12.6605504 | 3 | 89.5995893 | 3 |
| 3.71154523 | 1 | 35.28 | 1 |
| 4.98372316 | 1 | 73.44 | 1 |
| 4.93628073 | 1 | 67.33 | 1 |
| 3.72380304 | 1 | 0 | 1 |
| 6.09024811 | 1 | 108.95 | 1 |
| 24.2117987 | 6 | 167.852336 | 6 |
| 18.7836308 | 6 | 123.859438 | 5 |
| 44.1531575 | 10 | 157.147507 | 10 |
| 3.4548595 | 1 | 0 | 1 |
| 6.59070349 | 2 | 117.064337 | 2 |
| 104.276455 | 28 | 587.901179 | 28 |
| 3.11515236 | 1 | 0 | 1 |
| 363.186615 | 75 | 1608.59571 | 73 |
| 3.19486856 | 1 | 0 | 1 |
| 15.3660512 | 6 | 79.362511 | 6 |
| 31.428144 | 10 | 221.68 | 8 |
| 0 | 1 | 0 | 1 |
| 2.42846751 | 1 | 0 | 1 |
| 4.71209764 | 1 | 41.52 | 1 |
| 8.36041117 | 1 | 107.06 | 1 |
| 7.1265564 | 2 | 67.39 | 2 |
| 10.6601584 | 3 | 56.8255371 | 3 |
| 2.31624055 | 1 | 0 | 1 |

| | | | |
|---|---|---|---|
| 8.00094485 | 2 | 33.62 | 2 |
| 3.56324959 | 1 | 34.69 | 1 |
| 5.22342062 | 2 | 44.98 | 1 |
| 14.67472 | 2 | 59.65 | 2 |
| 2.95902586 | 1 | 36.83 | 1 |
| 3.61493731 | 1 | 50.49 | 1 |
| 8.49690723 | 3 | 126.37 | 3 |
| 4.3924222 | 1 | 43.53 | 1 |
| 7.67352247 | 2 | 93.12 | 2 |
| 2.08248925 | 1 | 0 | 1 |
| 4.4231534 | 1 | 52.93 | 1 |
| 10.7606924 | 3 | 127.186667 | 3 |
| 7.63867307 | 2 | 40.91 | 2 |
| 19.1596539 | 5 | 108.229909 | 5 |
| 2.31651092 | 1 | 31.81 | 1 |
| 2.36918068 | 1 | 28.11 | 1 |
| 5.92949653 | 2 | 99.9453997 | 2 |
| 2.96856093 | 1 | 43.77 | 1 |
| 2.31153321 | 1 | 0 | 1 |
| 4.97229528 | 1 | 66.52 | 1 |
| 33.2434864 | 8 | 232.633216 | 7 |
| 3.12415838 | 2 | 34.07 | 2 |
| 5.12468672 | 1 | 49.74 | 1 |
| 25.4691138 | 7 | 59.5293589 | 7 |
| 2.10187006 | 1 | 36.71 | 1 |
| 3.59733129 | 1 | 0 | 1 |
| 2.3338573 | 2 | 31.43 | 1 |
| 34.9019823 | 10 | 243.405242 | 10 |
| 32.675169 | 6 | 168.841147 | 6 |
| 14.4327338 | 2 | 94.583239 | 2 |
| 2.57320309 | 1 | 0 | 1 |
| 3.41293073 | 1 | 64.88 | 1 |
| 11.9345872 | 4 | 62.92 | 4 |
| 6.57766819 | 2 | 92.7689206 | 2 |
| 9.92774534 | 2 | 70.1317267 | 2 |
| 6.75794649 | 5 | 40.84 | 3 |
| 13.9066426 | 5 | 0 | 5 |
| 22.5112257 | 7 | 143.910871 | 6 |
| 16.8224738 | 5 | 95.952291 | 5 |
| 5.56821871 | 1 | 81.01 | 1 |
| 2.29209733 | 1 | 0 | 1 |
| 3.81516075 | 1 | 36.97 | 1 |
| 40.1416051 | 11 | 142.2176 | 10 |
| 3.6000967 | 1 | 39.53 | 1 |
| 10.7694652 | 2 | 113.74 | 2 |
| 4.48811483 | 1 | 62.02 | 1 |
| 12.393609 | 3 | 30.4 | 3 |
| 8.3524332 | 1 | 45.62 | 1 |
| 6.94943953 | 2 | 36.3 | 2 |
| 4.56524611 | 2 | 39.52 | 2 |
| 0 | 1 | 0 | 1 |
| 42.9141272 | 8 | 249.650979 | 7 |
| 0 | 1 | | |
| 18.56213 | 2 | 121.719691 | 2 |
| 4.26195383 | 1 | 36.95 | 1 |
| 4.74158096 | 1 | 88.22 | 1 |
| 25.2307799 | 8 | 149.181046 | 8 |
| 13.1413641 | 3 | 101.894727 | 3 |
| 13.4776564 | 2 | 58.6314467 | 2 |
| 36.2498631 | 4 | 221.50495 | 4 |
| 0 | 1 | | |
| 1.97836375 | 1 | 0 | 1 |

| | | | |
|---|---|---|---|
| 2.36967206 | 1 | 0 | 1 |
| 3.55437779 | 1 | 0 | 1 |
| 159.239096 | 29 | 832.657551 | 28 |
| 4.68597174 | 1 | 68.49 | 1 |
| 2.20098734 | 1 | 0 | 1 |
| 34.8952532 | 11 | 180.454444 | 11 |
| 9.12618709 | 3 | 47.8316723 | 3 |
| 2.77935624 | 1 | 0 | 1 |
| 4.24389029 | 1 | 68.13 | 1 |
| 3.72551441 | 1 | 48.87 | 1 |
| 49.1514459 | 14 | 233.012221 | 14 |
| 9.48723483 | 3 | 98.65 | 3 |
| 33.8078101 | 7 | 161.4022 | 7 |
| 10.1651897 | 3 | 66.0874891 | 3 |
| 5.3379283 | 2 | 32.94 | 2 |
| 13.001266 | 3 | 126.096064 | 3 |
| 0 | 1 | 0 | 1 |
| 2.99953485 | 1 | 0 | 1 |
| 7.33844638 | 2 | 49.91 | 2 |
| 2.64483261 | 1 | 27.85 | 1 |
| 3.88559866 | 1 | 0 | 1 |
| 3.02004409 | 2 | 25.03 | 2 |
| 7.84711504 | 2 | 0 | 2 |
| 3.33195829 | 1 | 44.77 | 1 |
| 3.40870905 | 1 | | |
| 8.15792108 | 1 | 102.4 | 1 |
| 96.4463909 | 23 | 476.784 | 23 |
| 4.66211653 | 1 | 47.07 | 1 |
| 1.63742185 | 1 | 33.9 | 1 |
| 1.71067905 | 1 | 45.83 | 1 |
| 8.97868395 | 3 | 33.22 | 3 |
| 10.4608049 | 4 | 0 | 4 |
| 33.9170287 | 7 | 90.1257108 | 6 |
| 1.70085466 | 1 | 0 | 1 |
| 2.14238477 | 1 | 60.79 | 1 |
| 0 | 1 | 0 | 1 |
| 2.44928765 | 1 | 15.38 | 1 |
| 5.51784492 | 1 | 53.2666072 | 1 |
| 3.95546484 | 1 | 54.63 | 1 |
| 3.88615513 | 1 | 0 | 1 |
| 9.96094513 | 2 | 98.62 | 2 |
| 4.88471913 | 1 | 0 | 1 |
| 2.59867525 | 1 | 0 | 1 |
| 2.21523571 | 1 | 0 | 1 |
| 6.47029185 | 2 | 77.91 | 1 |
| 7.4495132 | 3 | 39.24 | 2 |
| 30.3354036 | 10 | 174.523774 | 10 |
| 5.61901307 | 2 | 28.28 | 2 |
| 21.3001924 | 5 | 133.991237 | 5 |
| 6.27767968 | 2 | 75.85 | 2 |
| 21.6429446 | 8 | 81.9757042 | 7 |
| 20.7086973 | 5 | 182.117124 | 5 |
| 2.34712219 | 1 | 0 | 1 |
| 28.4584491 | 4 | 195.036249 | 4 |
| 0 | 1 | 0 | 1 |
| 2.56522799 | 2 | 27.21 | 1 |
| 14.3900259 | 2 | 68.3531946 | 2 |
| 23.2039244 | 5 | 123.72 | 4 |
