## Supplemental table 2 for "A TMT-based quantitative proteomics approach toward α-syn PFF associated Lewy Body Dementia (LBD) using α-syn PFF-injected mouse brain tissues"

| Gene_Symbol | Set.1.Ratio.127/126 | Set.1.Ratio.128/126 | Set.1.Ratio.129/126 |
|---|---|---|---|
| HBG1 | 1.949 | 0.947 | 1.137 |
| HBA1 | 1.495 | 0.827 | 0.855 |
| PCSK1N | 1.286 | 0.891 | 0.798 |
| TPPP3 | 1.367 | 1.2 | 0.763 |
| RAB10 | 1.409 | 1.705 | 1.318 |
| TUBB2B | 1.025 | 0.838 | 0.738 |
| NDUFA7 | 1.288 | 0.712 | 0.808 |
| TPM1 | 1.24 | 1.031 | 0.906 |
| PYGM | 1.363 | 1.308 | 0.835 |
| HSPA4L | 0.976 | 0.868 | 0.753 |
| SARS | 0.921 | 0.913 | 0.739 |
| CLASP2 | 1.333 | 0.908 | 0.867 |
| PPID | 1 | 1 | 1 |
| CA2 | 0.641 | 0.875 | 0.703 |
| SDHB | 1.27 | 1.066 | 0.869 |
| DNAJA2 | 1.195 | 0.87 | 1.221 |
| RPS19 | 1.074 | 0.823 | 0.863 |
| TPM1 | 1.24 | 1.031 | 0.906 |
| ETFA | 1.286 | 1.274 | 0.619 |
| GLO1 | 1.208 | 1.16 | 0.752 |
| STMN1 | 1.081 | 0.874 | 0.785 |
| ALDOA | 1.165 | 0.994 | 0.805 |
| HIST2H2AA3 | 1 | 1 | 1 |
| RPS6 | 1.058 | 0.907 | 0.92 |
| PPA1 | 1.015 | 0.888 | 0.882 |
| CLTB | 0.893 | 0.788 | 0.768 |
| CALR | 1 | 1 | 1 |
| TIMM8A | 1.102 | 0.778 | 0.842 |
| HIST1H1D | 1.114 | 0.704 | 0.806 |
| SNCA | 1.052 | 0.908 | 0.933 |
| IMPA1 | 0.983 | 0.844 | 0.776 |
| SH3GLB2 | 0.945 | 0.989 | 0.663 |
| PACSIN1 | 0.992 | 0.943 | 0.81 |
| PDIA3 | 0.878 | 0.763 | 0.789 |
| ABAT | 0.747 | 0.768 | 0.676 |
| RPL8 | 0.94 | 0.949 | 0.843 |
| NCAM1 | 0.866 | 0.795 | 0.764 |
| PFKP | 1.532 | 1.277 | 0.66 |
| HSPA5 | 1.077 | 0.993 | 0.851 |
| NDUFS3 | 1.166 | 0.947 | 0.795 |
| CORO1A | 0.747 | 0.575 | 0.759 |
| PFN2 | 1.123 | 1.015 | 0.877 |

| | | | |
|---|---|---|---|
| LSAMP | 1.125 | 1.037 | 1.015 |
| CAPZA2 | 0.888 | 0.837 | 0.98 |
| CCT4 | 1.073 | 0.97 | 0.816 |
| 4-Sep | 1 | 1 | 1 |
| 4-Sep | 1 | 1 | 1 |
| UGP2 | 1.066 | 1.102 | 0.889 |
| ATP6V1G2 | 1.089 | 0.843 | 0.881 |
| ERP29 | 1.032 | 0.774 | 0.874 |
| PTMS | 1.029 | 0.901 | 0.836 |
| HSPH1 | 1.026 | 1.035 | 0.894 |
| NPTX1 | 1.026 | 1.035 | 0.907 |
| H1F0 | 1.022 | 0.93 | 1.054 |
| PSMC2 | 0.987 | 0.826 | 0.708 |
| PSMC3 | 1 | 1 | 1 |
| 4-Sep | 1 | 1 | 1 |
| 4-Sep | 1 | 1 | 1 |
| 8-Sep | 1 | 1 | 1 |
| ABI1 | 1 | 1 | 1 |
| ABI2 | 1 | 1 | 1 |
| ADAM23 | 1 | 1 | 1 |
| ADAP1 | 1 | 1 | 1 |
| AHCY | 1 | 1 | 1 |
| AHSA1 | 1 | 1 | 1 |
| AK1 | 1 | 1 | 1 |
| ALCAM | 1 | 1 | 1 |
| ANXA6 | 1 | 1 | 1 |
| ARFGAP1 | 1 | 1 | 1 |
| ARL3 | 1 | 1 | 1 |
| ARPC1A | 1 | 1 | 1 |
| ARPC3 | 1 | 1 | 1 |
| ATL1 | 1 | 1 | 1 |
| ATP5C1 | 1 | 1 | 1 |
| ATP5H | 1 | 1 | 1 |
| BAIAP2 | 1 | 1 | 1 |
| BCAN | 1 | 1 | 1 |
| BPNT1 | 1 | 1 | 1 |
| CACNA2D1 | 1 | 1 | 1 |
| CARHSP1 | 1 | 1 | 1 |
| CASKIN1 | 1 | 1 | 1 |
| CDK5 | 1 | 1 | 1 |
| CELF2 | 1 | 1 | 1 |
| CISD1 | 1 | 1 | 1 |
| CORO1C | 1 | 1 | 1 |

| | | | |
|---|---|---|---|
| COX7C | 1 | 1 | 1 |
| DCTN1 | 1 | 1 | 1 |
| DLG4 | 1 | 1 | 1 |
| DMXL2 | 1 | 1 | 1 |
| DNM1 | 1 | 1 | 1 |
| DNM1 | 1 | 1 | 1 |
| EIF4E | 1 | 1 | 1 |
| EIF5 | 1 | 1 | 1 |
| FABP3 | 1 | 1 | 1 |
| FAM49A | 1 | 1 | 1 |
| FASN | 1 | 1 | 1 |
| FKBP2 | 1 | 1 | 1 |
| GABARAPL1 | 1 | 1 | 1 |
| GABRA1 | 1 | 1 | 1 |
| GABRB2 | 1 | 1 | 1 |
| GDI2 | 1 | 1 | 1 |
| GLOD4 | 1 | 1 | 1 |
| GNG13 | 1 | 1 | 1 |
| GSTM1 | 1 | 1 | 1 |
| HAPLN1 | 1 | 1 | 1 |
| HDHD2 | 1 | 1 | 1 |
| HEPACAM | 1 | 1 | 1 |
| HNRNPM | 1 | 1 | 1 |
| HNRNPU | 1 | 1 | 1 |
| HOMER1 | 1 | 1 | 1 |
| HYOU1 | 1 | 1 | 1 |
| IDH1 | 1 | 1 | 1 |
| IQSEC1 | 1 | 1 | 1 |
| ITM2C | 1 | 1 | 1 |
| KIAA1045 | 1 | 1 | 1 |
| KIF2A | 1 | 1 | 1 |
| LIN7A | 1 | 1 | 1 |
| LMNA | 1 | 1 | 1 |
| MDP1 | 1 | 1 | 1 |
| ME3 | 1 | 1 | 1 |
| MIF | 1 | 1 | 1 |
| MOG | 1 | 1 | 1 |
| MPP6 | 1 | 1 | 1 |
| MYH10 | 1 | 1 | 1 |
| MYL12A | 1 | 1 | 1 |
| NEDD8-MDP1 | 1 | 1 | 1 |
| NGEF | 1 | 1 | 1 |
| NRAS | 1 | 1 | 1 |

| | | | |
|---|---|---|---|
| NSFL1C | 1 | 1 | 1 |
| OGT | 1 | 1 | 1 |
| OMG | 1 | 1 | 1 |
| OSBP | 1 | 1 | 1 |
| P4HB | 1 | 1 | 1 |
| PACS1 | 1 | 1 | 1 |
| PAK1 | 1 | 1 | 1 |
| PCBP1 | 1 | 1 | 1 |
| PCDH1 | 1 | 1 | 1 |
| PCMT1 | 1 | 1 | 1 |
| PDCD5 | 1 | 1 | 1 |
| PDCD6IP | 1 | 1 | 1 |
| PEX5L | 1 | 1 | 1 |
| PFKL | 1 | 1 | 1 |
| PGM2L1 | 1 | 1 | 1 |
| PGRMC1 | 1 | 1 | 1 |
| PHYHIP | 1 | 1 | 1 |
| PIP4K2B | 1 | 1 | 1 |
| PLEC | 1 | 1 | 1 |
| PLS3 | 1 | 1 | 1 |
| PPFIA3 | 1 | 1 | 1 |
| PPIB | 1 | 1 | 1 |
| PPP5C | 1 | 1 | 1 |
| PRKAR1A | 1 | 1 | 1 |
| PSMA2 | 1 | 1 | 1 |
| PSMA3 | 1 | 1 | 1 |
| PSMA8 | 1 | 1 | 1 |
| PURB | 1 | 1 | 1 |
| RAB14 | 1 | 1 | 1 |
| RAB18 | 1 | 1 | 1 |
| RHOA | 1 | 1 | 1 |
| RPL10A | 1 | 1 | 1 |
| RPL12 | 1 | 1 | 1 |
| RPL14 | 1 | 1 | 1 |
| RPL21 | 1 | 1 | 1 |
| RPL23 | 1 | 1 | 1 |
| RPL3 | 1 | 1 | 1 |
| RPL7A | 1 | 1 | 1 |
| RPLP0 | 1 | 1 | 1 |
| RPLP2 | 1 | 1 | 1 |
| RPS12 | 1 | 1 | 1 |
| RPS26 | 1 | 1 | 1 |
| RPS3 | 1 | 1 | 1 |

| | | | |
|---|---|---|---|
| RTN1 | 1 | 1 | 1 |
| RTN1 | 1 | 1 | 1 |
| SKP1 | 1 | 1 | 1 |
| SLC25A11 | 1 | 1 | 1 |
| SLC4A10 | 1 | 1 | 1 |
| SLC4A4 | 1 | 1 | 1 |
| SNRPD3 | 1 | 1 | 1 |
| SRSF7 | 1 | 1 | 1 |
| SUCLG1 | 1 | 1 | 1 |
| SUCLG2 | 1 | 1 | 1 |
| SYNGR1 | 1 | 1 | 1 |
| TOM1L2 | 1 | 1 | 1 |
| TPP2 | 1 | 1 | 1 |
| UBE2I | 1 | 1 | 1 |
| VPS35 | 1 | 1 | 1 |
| KPNB1 | 1 | 1 | 1 |
| NECAP1 | 1 | 1 | 1 |
| LASP1 | 0.99 | 0.876 | 0.783 |
| ROGDI | 1 | 1 | 1 |
| PSPC1 | 0.993 | 0.901 | 0.737 |
| EHD3 | 0.943 | 0.815 | 0.649 |
| SCRN1 | 1.188 | 1.014 | 0.919 |
| DNAJC6 | 0.978 | 0.936 | 0.76 |
| NDUFA12 | 1 | 1 | 1 |
| PHB2 | 1.015 | 0.966 | 0.844 |
| GNA11 | 0.976 | 0.794 | 0.683 |
| EPB41L1 | 1.009 | 0.82 | 0.709 |
| GSK3A | 0.957 | 0.856 | 0.759 |
| GMPR | 1 | 1 | 1 |
| SRSF1 | 1.04 | 0.685 | 0.75 |
| ETFB | 0.955 | 0.988 | 0.653 |
| PPIA | 0.931 | 0.853 | 0.784 |
| THEM4 | 0.95 | 0.763 | 0.636 |
| SGIP1 | 1 | 1 | 1 |
| L1CAM | 0.943 | 0.859 | 0.872 |
| VAT1L | 0.941 | 0.88 | 0.675 |
| EPB41L3 | 1.025 | 0.891 | 0.766 |
| ATP1B2 | 1.026 | 0.895 | 0.658 |
| C1orf95 | 1 | 1 | 1 |
| SLC1A2 | 0.938 | 0.911 | 0.785 |
| TOLLIP | 1 | 1 | 1 |
| TPPP | 1.011 | 0.944 | 0.841 |
| EEF1G | 0.957 | 0.853 | 0.776 |

| | | | |
|---|---|---|---|
| PABPC1 | 1.003 | 0.898 | 0.866 |
| NDUFS2 | 0.925 | 0.936 | 0.744 |
| CDC42 | 0.908 | 0.887 | 0.9 |
| MAP6 | 0.972 | 0.937 | 0.868 |
| CMPK1 | 1.109 | 0.916 | 0.831 |
| ICAM5 | 0.923 | 0.769 | 0.753 |
| VAPB | 0.974 | 0.848 | 0.819 |
| ADD2 | 0.935 | 0.865 | 0.838 |
| MBP | 0.972 | 0.899 | 0.66 |
| AHCYL1 | 0.995 | 0.949 | 0.842 |
| HSP90AA1 | 1.041 | 0.888 | 0.773 |
| CAP2 | 0.951 | 0.781 | 0.906 |
| RPS15 | 0.756 | 0.733 | 0.674 |
| UBQLN4 | 0.904 | 0.872 | 0.78 |
| ROCK2 | 0.903 | 0.852 | 0.932 |
| KIF5C | 0.86 | 0.754 | 0.835 |
| ARHGDIA | 0.919 | 0.885 | 0.807 |
| ENSA | 0.902 | 0.727 | 0.807 |
| GNAS | 0.898 | 0.817 | 0.663 |
| OXR1 | 0.969 | 0.952 | 0.872 |
| PSD3 | 1.009 | 0.852 | 0.889 |
| TPM3 | 0.876 | 0.884 | 0.821 |
| CALB1 | 0.861 | 0.84 | 0.671 |
| COX6A1 | 0.987 | 0.888 | 0.978 |
| RPS2 | 0.895 | 0.826 | 0.853 |
| TMED10 | 0.895 | 0.807 | 0.76 |
| RAB11B | 0.894 | 0.828 | 0.754 |
| COX5A | 0.955 | 0.926 | 0.79 |
| SV2A | 0.903 | 0.876 | 0.733 |
| UQCRB | 0.889 | 0.825 | 0.71 |
| CKMT1B | 0.949 | 1.008 | 0.818 |
| HSPE1 | 0.867 | 0.837 | 0.787 |
| MDH1 | 0.848 | 0.907 | 0.789 |
| MAPK1 | 0.967 | 0.892 | 0.749 |
| PYGB | 0.883 | 0.784 | 0.784 |
| DPYSL4 | 0.858 | 0.897 | 0.866 |
| NRCAM | 0.861 | 0.909 | 0.883 |
| PSMA5 | 0.877 | 1.095 | 0.801 |
| EFHD2 | 0.883 | 1.056 | 0.973 |
| RPL17-C18orf32 | 1 | 1 | 1 |
| GLS | 0.937 | 1.008 | 0.929 |
| PPP1R7 | 1 | 1 | 1 |
| BIN1 | 0.861 | 0.883 | 0.711 |

| | | | |
|---|---|---|---|
| PSMA1 | 0.87 | 0.881 | 0.828 |
| PHB | 0.965 | 0.907 | 0.771 |
| IDH2 | 0.869 | 0.798 | 0.764 |
| PPP1R1B | 1 | 1 | 1 |
| MAPT | 0.946 | 0.823 | 0.766 |
| CADM3 | 0.859 | 0.871 | 0.704 |
| AKR1A1 | 1 | 1 | 1 |
| EIF4H | 0.692 | 0.842 | 0.692 |
| SNAP91 | 0.91 | 0.831 | 0.752 |
| TUBA1A | 0.98 | 0.824 | 0.737 |
| HNRNPAB | 0.855 | 0.797 | 0.862 |
| PDIA6 | 0.855 | 0.845 | 0.831 |
| CBX3 | 1 | 1 | 1 |
| GPM6B | 1 | 1 | 1 |
| LGI1 | 0.853 | 0.814 | 0.75 |
| RPS4X | 1 | 1 | 1 |
| CNRIP1 | 0.851 | 0.72 | 0.829 |
| DPYSL3 | 0.888 | 0.858 | 0.763 |
| ATP5O | 0.864 | 0.79 | 0.758 |
| PRPF19 | 1 | 1 | 1 |
| PSMC5 | 0.849 | 0.885 | 0.656 |
| TUBB6 | 0.992 | 0.763 | 0.792 |
| NDUFA6 | 1 | 1 | 1 |
| ACTN4 | 0.955 | 0.921 | 0.85 |
| ATP6V1D | 1 | 1 | 1 |
| DPYSL2 | 0.844 | 0.855 | 0.755 |
| HSPD1 | 0.884 | 0.82 | 0.745 |
| CDH13 | 1 | 1 | 1 |
| G3BP2 | 1 | 1 | 1 |
| GRIA3 | 0.841 | 0.849 | 0.803 |
| PCP4 | 0.95 | 0.736 | 0.702 |
| FUS | 0.847 | 0.831 | 0.949 |
| CAPZB | 0.963 | 0.94 | 0.969 |
| EIF4B | 1 | 1 | 1 |
| CCT2 | 1 | 1 | 1 |
| SFPQ | 1 | 1 | 1 |
| DYNLRB1 | 1 | 1 | 1 |
| PARK7 | 0.879 | 0.673 | 0.781 |
| PPP3CB | 1 | 1 | 1 |
| OLA1 | 0.896 | 0.884 | 0.764 |
| NAP1L1 | 0.829 | 0.641 | 0.765 |
| ALDH1A1 | 0.974 | 0.853 | 0.714 |
| DIRAS2 | 0.824 | 0.814 | 0.712 |

| | | | |
|---|---|---|---|
| DSTN | 0.82 | 0.788 | 0.727 |
| MAG | 0.82 | 0.755 | 0.592 |
| RPL5 | 0.818 | 0.8 | 0.716 |
| PPA2 | 0.817 | 0.881 | 0.701 |
| CPLX2 | 0.93 | 0.809 | 0.734 |
| MAP1B | 0.893 | 0.772 | 0.678 |
| PGAM1 | 0.849 | 0.812 | 0.728 |
| TUBB | 0.921 | 0.799 | 0.723 |
| BLMH | 0.812 | 0.739 | 0.7 |
| LDHB | 0.97 | 0.864 | 0.748 |
| SH3BGRL3 | 1 | 1 | 1 |
| ATP2A2 | 0.878 | 0.879 | 0.842 |
| HNRNPC | 1 | 1 | 1 |
| MARCKS | 0.804 | 0.749 | 0.68 |
| ACLY | 0.968 | 1.078 | 0.747 |
| YARS | 1 | 1 | 1 |
| HIST1H2BD | 0.925 | 0.753 | 0.859 |
| ATP6V1B2 | 0.92 | 0.866 | 0.784 |
| VCP | 0.803 | 0.803 | 0.739 |
| VIM | 0.905 | 0.804 | 0.681 |
| CSRP1 | 0.8 | 0.912 | 0.791 |
| PRPSAP1 | 0.907 | 0.813 | 0.813 |
| GPD1L | 0.798 | 0.887 | 0.791 |
| TAGLN3 | 0.919 | 0.962 | 0.844 |
| MAP2 | 0.917 | 0.738 | 0.7 |
| ANXA5 | 0.794 | 0.723 | 0.608 |
| FSCN1 | 0.858 | 0.808 | 0.774 |
| NCDN | 0.792 | 0.615 | 0.594 |
| ANK2 | 1.015 | 0.888 | 0.81 |
| HIST1H4L | 0.853 | 0.785 | 0.813 |
| AUH | 1 | 1 | 1 |
| TUBB4B | 0.694 | 0.67 | 0.637 |
| CRMP1 | 0.923 | 0.899 | 0.838 |
| NPM1 | 1 | 1 | 1 |
| CCT7 | 1 | 1 | 1 |
| ACSL6 | 0.781 | 0.684 | 0.63 |
| RPS16 | 0.781 | 0.852 | 0.735 |
| GDA | 1 | 1 | 1 |
| LETM1 | 0.78 | 0.855 | 0.745 |
| HEBP1 | 1 | 1 | 1 |
| DPYSL2 | 0.844 | 0.855 | 0.755 |
| CADM4 | 1 | 1 | 1 |
| PTK2B | 0.774 | 0.769 | 0.689 |

| | | | |
|---|---|---|---|
| UQCRC2 | 0.774 | 0.776 | 0.779 |
| DYNC1LI1 | 0.916 | 0.948 | 0.705 |
| 1-Sep | 0.849 | 0.792 | 0.707 |
| ADD1 | 0.845 | 0.758 | 0.737 |
| HSPA8 | 0.907 | 0.847 | 0.759 |
| SNAP25 | 0.939 | 0.798 | 0.649 |
| PRKCE | 1 | 1 | 1 |
| TKT | 0.869 | 0.839 | 0.723 |
| ACOT7 | 0.798 | 0.858 | 0.72 |
| CRYM | 1 | 1 | 1 |
| RPS3A | 0.862 | 0.821 | 0.76 |
| SPTBN1 | 0.891 | 0.911 | 0.832 |
| ATP6V1A | 0.901 | 0.899 | 0.803 |
| WDR7 | 1 | 1 | 1 |
| ATP1A3 | 0.837 | 0.833 | 0.749 |
| GAPDH | 0.91 | 0.869 | 0.789 |
| SYP | 1 | 1 | 1 |
| RAP1GDS1 | 0.91 | 0.932 | 0.86 |
| ACTN1 | 0.772 | 0.848 | 0.827 |
| CYFIP1 | 0.755 | 0.768 | 0.756 |
| PPP2CA | 1 | 1 | 1 |
| RPS8 | 0.764 | 0.739 | 0.552 |
| SDHA | 1 | 1 | 1 |
| SPTAN1 | 0.925 | 0.91 | 0.837 |
| DDAH1 | 0.749 | 0.853 | 0.804 |
| HPRT1 | 1 | 1 | 1 |
| RALA | 0.749 | 0.795 | 0.685 |
| SLC12A6 | 0.749 | 0.648 | 0.704 |
| RPL23A | 0.813 | 0.777 | 0.747 |
| CAP1 | 0.94 | 1.015 | 0.814 |
| ATP6V1H | 0.811 | 0.758 | 0.747 |
| ST13 | 1.003 | 0.982 | 0.765 |
| TMSB4X | 0.854 | 0.637 | 0.657 |
| 2-Sep | 0.799 | 0.724 | 0.617 |
| ARPC2 | 1 | 1 | 1 |
| TNR | 1 | 1 | 1 |
| EEF2 | 0.85 | 0.803 | 0.718 |
| DYNC1H1 | 0.828 | 0.791 | 0.736 |
| PLCB1 | 0.888 | 1.003 | 0.952 |
| DLD | 0.88 | 0.848 | 0.63 |
| EEF1A2 | 0.931 | 0.902 | 0.825 |
| NAPG | 0.864 | 0.879 | 0.759 |
| HK1 | 0.904 | 1.016 | 0.891 |

| | | | |
|---|---|---|---|
| PFKM | 0.842 | 0.902 | 0.8 |
| TPI1 | 0.879 | 0.878 | 0.815 |
| GLUL | 0.906 | 1 | 0.906 |
| SEC22B | 0.732 | 0.989 | 0.684 |
| DNM1L | 0.849 | 0.839 | 0.688 |
| RAB7A | 1 | 1 | 1 |
| ATP5D | 0.825 | 0.754 | 0.65 |
| OGDHL | 0.778 | 0.829 | 0.781 |
| SFXN3 | 0.897 | 0.882 | 0.805 |
| ATP2B1 | 0.888 | 0.792 | 0.742 |
| VAPA | 0.898 | 0.97 | 0.869 |
| CPNE6 | 0.721 | 0.597 | 0.693 |
| RAB1A | 0.902 | 0.8 | 0.685 |
| NPTN | 0.719 | 0.756 | 0.755 |
| ACTR1A | 0.844 | 0.768 | 0.739 |
| OTUB1 | 0.832 | 0.882 | 0.844 |
| CANX | 0.929 | 0.971 | 0.794 |
| RHOG | 0.838 | 0.793 | 0.689 |
| UQCRC1 | 0.762 | 0.713 | 0.732 |
| HNRNPK | 0.859 | 0.832 | 0.828 |
| CADPS | 0.909 | 0.84 | 0.746 |
| ALDH2 | 0.709 | 0.756 | 0.77 |
| RPL7 | 0.794 | 0.861 | 0.782 |
| TCP1 | 0.788 | 0.825 | 0.703 |
| CHCHD3 | 1 | 1 | 1 |
| CTNNB1 | 0.921 | 0.751 | 0.767 |
| PTPRZ1 | 0.805 | 0.885 | 0.783 |
| FH | 1 | 1 | 1 |
| GIT1 | 1 | 1 | 1 |
| NUTF2 | 1 | 1 | 1 |
| DNM1 | 1 | 1 | 1 |
| DNM1 | 1 | 1 | 1 |
| UBE2D2 | 1 | 1 | 1 |
| OGDH | 0.825 | 0.839 | 0.814 |
| HSPA9 | 0.852 | 0.718 | 0.673 |
| NPEPPS | 0.908 | 0.969 | 0.835 |
| PEA15 | 1 | 1 | 1 |
| UBA1 | 0.838 | 0.686 | 0.715 |
| DNAJA1 | 1 | 1 | 1 |
| WDR1 | 0.936 | 0.853 | 0.8 |
| RPH3A | 0.868 | 0.807 | 0.744 |
| CCT3 | 0.836 | 0.872 | 0.794 |
| PRKACA | 0.693 | 0.628 | 0.766 |

| | | | |
|---|---|---|---|
| SNCB | 0.859 | 0.882 | 0.756 |
| LAP3 | 0.863 | 0.782 | 0.715 |
| NME2 | 0.841 | 0.857 | 0.786 |
| 5-Sep | 0.781 | 0.696 | 0.646 |
| NEFM | 0.853 | 0.806 | 0.662 |
| IMMT | 0.689 | 0.841 | 0.753 |
| 6-Sep | 0.764 | 0.813 | 0.758 |
| ATP5B | 0.855 | 0.845 | 0.737 |
| GNB2L1 | 0.89 | 0.88 | 0.782 |
| MSN | 0.687 | 0.706 | 0.554 |
| CPE | 0.832 | 0.807 | 0.699 |
| NTM | 0.838 | 0.765 | 0.803 |
| STXBP1 | 0.814 | 0.822 | 0.77 |
| DLST | 0.679 | 0.778 | 0.64 |
| FKBP1A | 0.824 | 0.737 | 0.796 |
| VAMP2 | 0.868 | 0.869 | 0.725 |
| AP2A2 | 0.785 | 0.808 | 0.751 |
| HNRNPL | 0.77 | 0.732 | 0.694 |
| ACO2 | 0.85 | 0.876 | 0.758 |
| OPCML | 0.837 | 0.863 | 0.826 |
| YWHAQ | 0.823 | 0.778 | 0.733 |
| CCT8 | 0.809 | 0.592 | 0.796 |
| SLC25A12 | 0.826 | 0.778 | 0.752 |
| ATP5A1 | 0.863 | 0.88 | 0.751 |
| NDUFS1 | 0.837 | 0.847 | 0.764 |
| OPA1 | 0.833 | 0.798 | 0.72 |
| ACTR2 | 0.818 | 0.825 | 0.776 |
| ATP5I | 0.742 | 0.695 | 0.604 |
| CADM2 | 0.761 | 0.857 | 0.839 |
| MAP1A | 0.831 | 0.795 | 0.872 |
| IDH3A | 0.918 | 0.883 | 0.814 |
| RPL38 | 0.866 | 0.709 | 0.687 |
| STIP1 | 0.826 | 0.801 | 0.755 |
| ATP1A2 | 0.866 | 0.861 | 0.712 |
| ATP6V0D1 | 0.826 | 0.844 | 0.789 |
| CADPS | 0.909 | 0.84 | 0.746 |
| SOD2 | 0.661 | 0.742 | 0.679 |
| SYN1 | 0.882 | 0.861 | 0.775 |
| SCP2 | 0.877 | 0.718 | 0.737 |
| MAPRE2 | 0.659 | 0.596 | 0.567 |
| GFAP | 0.857 | 0.865 | 0.745 |
| HNRNPA3 | 1 | 1 | 1 |
| WASF1 | 0.731 | 0.805 | 0.826 |

| | | | |
|---|---|---|---|
| RAB6B | 0.688 | 0.697 | 0.667 |
| DBN1 | 0.877 | 0.768 | 0.848 |
| UBA52 | 0.742 | 0.705 | 0.772 |
| CSNK2A1 | 0.812 | 0.803 | 0.727 |
| MAP2K1 | 0.859 | 0.848 | 0.729 |
| NDUFS8 | 0.821 | 0.917 | 0.924 |
| PDHB | 0.764 | 0.775 | 0.728 |
| FBXO2 | 1 | 1 | 1 |
| RPL11 | 0.805 | 0.79 | 0.669 |
| UBE2N | 0.772 | 0.825 | 0.729 |
| ENO2 | 0.821 | 0.838 | 0.715 |
| AP1B1 | 0.643 | 0.692 | 0.696 |
| ACTR3 | 0.779 | 0.792 | 0.87 |
| PFN1 | 0.64 | 0.656 | 0.64 |
| CNTN1 | 0.833 | 0.898 | 0.79 |
| GNAO1 | 0.798 | 0.709 | 0.772 |
| RTCB | 0.851 | 0.651 | 0.687 |
| GPD2 | 0.883 | 0.883 | 0.866 |
| MYL6 | 0.743 | 0.812 | 0.818 |
| CS | 0.851 | 0.902 | 0.786 |
| VDAC2 | 0.882 | 0.93 | 0.778 |
| ACTR3B | 1 | 1 | 1 |
| ATP6V0A1 | 0.815 | 0.861 | 0.743 |
| STX1B | 0.792 | 0.859 | 0.771 |
| SCAMP1 | 0.833 | 0.872 | 0.803 |
| BSN | 0.83 | 0.697 | 0.702 |
| CLTA | 0.739 | 0.784 | 0.753 |
| SLC25A3 | 0.7 | 0.816 | 0.735 |
| RPSA | 0.888 | 0.81 | 0.731 |
| GNAI2 | 0.832 | 0.678 | 0.675 |
| DCTN2 | 0.884 | 1.092 | 0.98 |
| CALB2 | 0.788 | 0.844 | 0.542 |
| CADPS | 0.82 | 0.896 | 0.671 |
| SIRT2 | 0.62 | 0.786 | 0.722 |
| PURA | 0.756 | 0.713 | 0.73 |
| AAK1 | 0.774 | 0.634 | 0.575 |
| ACTG1 | 0.793 | 0.95 | 0.94 |
| ALDOC | 0.787 | 0.801 | 0.699 |
| ARF5 | 0.813 | 0.699 | 0.819 |
| RPS13 | 0.818 | 0.81 | 0.683 |
| CAND1 | 0.797 | 0.883 | 0.74 |
| UQCRFS1 | 0.806 | 0.813 | 0.743 |
| QDPR | 0.815 | 0.819 | 0.622 |

| | | | |
|---|---|---|---|
| ATP1B1 | 0.797 | 0.798 | 0.694 |
| YWHAG | 0.799 | 0.824 | 0.783 |
| EEF1A1 | 0.796 | 0.618 | 0.624 |
| PCBP2 | 0.875 | 0.648 | 0.795 |
| OXCT2 | 0.763 | 0.811 | 0.745 |
| HSPA4 | 0.77 | 0.773 | 0.748 |
| IDH3B | 0.77 | 0.7 | 0.642 |
| TUBB3 | 0.766 | 0.769 | 0.73 |
| SLC25A18 | 0.777 | 0.827 | 0.795 |
| CYCS | 1 | 1 | 1 |
| DLAT | 0.786 | 0.834 | 0.834 |
| HSP90B1 | 0.723 | 0.805 | 0.778 |
| SRCIN1 | 0.757 | 0.794 | 0.8 |
| FAM49B | 0.751 | 0.851 | 0.714 |
| PHGDH | 0.704 | 0.763 | 0.722 |
| HSP90AB1 | 0.792 | 0.72 | 0.747 |
| CSNK2B | 0.79 | 0.783 | 0.673 |
| TRIM2 | 0.576 | 0.819 | 0.799 |
| AKR1B1 | 0.853 | 0.821 | 0.79 |
| FKBP8 | 0.789 | 0.683 | 0.604 |
| CADPS | 0.82 | 0.896 | 0.671 |
| MDH2 | 0.804 | 0.783 | 0.711 |
| NDUFS4 | 0.763 | 0.88 | 0.805 |
| 10-Sep | 0.744 | 0.654 | 0.82 |
| CD47 | 0.86 | 0.82 | 0.834 |
| PRKCG | 0.834 | 0.561 | 0.55 |
| HNRNPH1 | 0.773 | 0.609 | 0.584 |
| ATP2B2 | 0.8 | 0.838 | 0.845 |
| SYNJ1 | 0.773 | 0.77 | 0.742 |
| AP2M1 | 0.789 | 0.783 | 0.742 |
| PGK1 | 0.716 | 0.727 | 0.695 |
| TMX4 | 0.812 | 0.798 | 0.671 |
| RPL13A | 0.76 | 0.831 | 0.632 |
| PRKAR2B | 0.763 | 0.672 | 0.61 |
| SPTBN2 | 0.802 | 0.814 | 0.806 |
| YWHAH | 0.81 | 0.773 | 0.756 |
| PDHA1 | 0.775 | 0.689 | 0.685 |
| PPP2R1A | 0.779 | 0.732 | 0.789 |
| DCLK1 | 0.743 | 0.728 | 0.744 |
| NFASC | 0.749 | 0.783 | 0.664 |
| PEBP1 | 0.788 | 0.534 | 0.579 |
| AP3B2 | 0.746 | 0.672 | 0.762 |
| NAPB | 0.802 | 0.858 | 0.739 |

| | | | |
|---|---|---|---|
| HSPA12A | 0.814 | 0.726 | 0.706 |
| CTTN | 0.77 | 0.641 | 0.722 |
| OLFM1 | 0.753 | 0.861 | 0.789 |
| RAC1 | 0.752 | 0.811 | 0.726 |
| ACAT1 | 1 | 1 | 1 |
| RPL4 | 0.785 | 0.824 | 0.729 |
| TMSB10 | 0.767 | 0.794 | 0.644 |
| PGM1 | 0.794 | 0.887 | 0.83 |
| YWHAZ | 0.734 | 0.743 | 0.723 |
| GDI1 | 0.826 | 0.867 | 0.766 |
| VDAC3 | 0.76 | 0.858 | 0.747 |
| MAPRE3 | 0.712 | 0.695 | 0.753 |
| INA | 0.732 | 0.802 | 0.656 |
| SUCLA2 | 0.698 | 0.59 | 0.673 |
| NEFL | 0.742 | 0.778 | 0.61 |
| CPLX1 | 0.824 | 0.687 | 0.678 |
| CALM2 | 0.733 | 0.822 | 0.751 |
| IDH3G | 0.877 | 1.071 | 0.785 |
| NDUFA5 | 0.754 | 0.779 | 0.74 |
| SLC8A2 | 0.685 | 0.655 | 0.696 |
| VDAC1 | 0.794 | 0.875 | 0.769 |
| FBXL16 | 0.746 | 0.74 | 0.74 |
| TUBA4A | 0.818 | 0.741 | 0.725 |
| AP2A1 | 0.795 | 0.767 | 0.741 |
| SH3GL2 | 0.736 | 0.816 | 0.763 |
| NEFH | 0.922 | 0.878 | 0.322 |
| PPP3CA | 0.756 | 0.756 | 0.748 |
| SLC17A7 | 0.809 | 0.781 | 0.837 |
| CKB | 0.722 | 0.635 | 0.648 |
| GDAP1 | 0.719 | 0.699 | 0.613 |
| ATP6V1C1 | 0.78 | 0.805 | 0.794 |
| NSF | 0.728 | 0.637 | 0.645 |
| PRKAR2A | 0.713 | 0.896 | 0.782 |
| MYO5A | 0.725 | 0.713 | 0.765 |
| CNP | 0.84 | 0.768 | 0.675 |
| PKM | 0.749 | 0.817 | 0.756 |
| ARPC4 | 0.641 | 0.821 | 0.833 |
| TALDO1 | 0.744 | 0.731 | 0.669 |
| NDUFA4 | 0.724 | 0.787 | 0.754 |
| TUFM | 0.703 | 0.52 | 0.644 |
| ATP1A1 | 0.726 | 0.818 | 0.786 |
| UCHL1 | 0.693 | 0.749 | 0.627 |
| PRDX6 | 0.697 | 0.663 | 0.702 |

| | | | |
|---|---|---|---|
| GMFB | 0.739 | 0.678 | 0.697 |
| LDHA | 0.774 | 0.737 | 0.801 |
| CLTC | 0.721 | 0.769 | 0.769 |
| PAFAH1B2 | 0.815 | 0.923 | 0.743 |
| GOT1 | 0.714 | 0.742 | 0.684 |
| NDUFV1 | 0.75 | 0.752 | 0.683 |
| ATP6V1E1 | 0.754 | 0.836 | 0.723 |
| TUBB4A | 0.772 | 0.715 | 0.698 |
| PRKCB | 0.855 | 0.77 | 0.702 |
| PLP1 | 0.663 | 0.663 | 0.51 |
| GNAI1 | 0.782 | 0.874 | 0.767 |
| YWHAB | 0.686 | 0.684 | 0.738 |
| SYN2 | 0.702 | 0.807 | 0.68 |
| LGALSL | 0.741 | 0.743 | 0.785 |
| VSNL1 | 0.783 | 0.828 | 0.683 |
| CFL1 | 0.76 | 0.714 | 0.712 |
| GOT2 | 0.696 | 0.758 | 0.669 |
| GPI | 0.699 | 0.693 | 0.656 |
| RAB3A | 0.668 | 0.76 | 0.757 |
| NDRG2 | 0.712 | 0.742 | 0.627 |
| USP5 | 0.707 | 0.472 | 0.543 |
| ENO1 | 0.714 | 0.725 | 0.71 |
| RPL13 | 0.679 | 0.595 | 0.612 |
| PPP3R1 | 0.734 | 0.694 | 0.655 |
| RAB2A | 0.709 | 0.708 | 0.72 |
| SYT1 | 0.714 | 0.781 | 0.767 |
| PRDX5 | 0.757 | 0.821 | 0.812 |
| AP2B1 | 0.674 | 0.723 | 0.758 |
| YWHAE | 0.654 | 0.75 | 0.706 |
| SLC8A1 | 0.701 | 0.845 | 0.79 |
| GNAQ | 0.673 | 0.737 | 0.74 |
| ARF1 | 0.612 | 0.678 | 0.725 |
| CAMK2D | 0.684 | 0.523 | 0.551 |
| HNRNPD | 0.676 | 0.686 | 0.63 |
| STX1A | 0.681 | 0.765 | 0.784 |
| LANCL2 | 0.64 | 0.711 | 0.639 |
| RAN | 0.619 | 0.702 | 0.713 |
| NCALD | 0.644 | 0.627 | 0.653 |
| HNRNPA2B1 | 0.672 | 0.564 | 0.684 |
| SLC25A5 | 0.661 | 0.703 | 0.671 |
| SCAMP5 | 0.722 | 0.792 | 0.745 |
| ACTA1 | 0.671 | 0.711 | 0.768 |
| EIF5A | 0.83 | 0.749 | 0.705 |

| | | | |
|---|---|---|---|
| SLC12A5 | 0.609 | 0.359 | 0.511 |
| TUBB2A | 0.643 | 0.626 | 0.635 |
| HPCAL4 | 0.608 | 0.72 | 0.61 |
| GLUD1 | 0.688 | 0.632 | 0.644 |
| SLC25A4 | 0.634 | 0.767 | 0.738 |
| RPL15 | 0.637 | 0.704 | 0.644 |
| DYNLL2 | 0.55 | 0.659 | 0.672 |
| ACAT2 | 0.65 | 0.848 | 0.924 |
| AMPH | 0.585 | 0.685 | 0.693 |
| SET | 0.548 | 1.151 | 0.753 |
| CCT5 | 0.527 | 0.571 | 0.637 |
| RPS7 | 0.306 | 0.531 | 0.347 |
| MCM3AP | 0.491 | 0.431 | 0.69 |
| DYNLL1 | 0.517 | 0.591 | 0.739 |
| CAMK2B | 0.47 | 0.363 | 0.435 |
| CAMK2A | 0.532 | 0.378 | 0.483 |

| Set.2.Ratio.127/126 | Set.2.Ratio.128/126 | Set.2.Ratio.129/126 | Average_FC_127/126 |
|---|---|---|---|
| 2.039 | 1.041 | 1.315 | 1.994 |
| 1.455 | 0.859 | 0.908 | 1.475 |
| 1.229 | 0.733 | 0.581 | 1.2575 |
| 1.144 | 1.015 | 0.738 | 1.2555 |
| 1.081 | 0.593 | 0.711 | 1.245 |
| 1.443 | 0.982 | 0.82 | 1.234 |
| 1.154 | 0.925 | 1.032 | 1.221 |
| 1.176 | 1.111 | 0.937 | 1.208 |
| 1 | 1 | 1 | 1.1815 |
| 1.374 | 1.033 | 0.802 | 1.175 |
| 1.423 | 0.96 | 0.945 | 1.172 |
| 1 | 1 | 1 | 1.1665 |
| 1.297 | 0.957 | 0.827 | 1.1485 |
| 1.634 | 1.665 | 1.062 | 1.1375 |
| 1 | 1 | 1 | 1.135 |
| 1.049 | 1.025 | 0.926 | 1.122 |
| 1.168 | 0.912 | 0.747 | 1.121 |
| 1 | 1 | 1 | 1.12 |
| 0.922 | 0.802 | 0.61 | 1.104 |
| 1 | 1 | 1 | 1.104 |
| 1.125 | 0.891 | 0.832 | 1.103 |
| 1.04 | 0.972 | 0.833 | 1.1025 |
| 1.2 | 1.053 | 0.885 | 1.1 |
| 1.139 | 0.94 | 0.781 | 1.0985 |
| 1.179 | 1.069 | 0.94 | 1.097 |
| 1.297 | 1.072 | 0.81 | 1.095 |
| 1.176 | 0.997 | 0.851 | 1.088 |
| 1.068 | 0.775 | 0.857 | 1.085 |
| 1.032 | 0.703 | 0.787 | 1.073 |
| 1.076 | 1.038 | 0.899 | 1.064 |
| 1.141 | 0.972 | 0.896 | 1.062 |
| 1.179 | 0.888 | 0.85 | 1.062 |
| 1.127 | 0.993 | 0.83 | 1.0595 |
| 1.239 | 0.998 | 0.899 | 1.0585 |
| 1.364 | 1 | 1.205 | 1.0555 |
| 1.17 | 0.98 | 0.841 | 1.055 |
| 1.236 | 0.953 | 0.834 | 1.051 |
| 0.566 | 0.717 | 0.515 | 1.049 |
| 1.017 | 0.891 | 0.777 | 1.047 |
| 0.928 | 0.949 | 0.859 | 1.047 |
| 1.338 | 1.156 | 1.078 | 1.0425 |
| 0.958 | 0.961 | 0.837 | 1.0405 |

| | | | |
|---|---|---|---|
| 0.954 | 0.962 | 0.893 | 1.0395 |
| 1.187 | 0.856 | 0.784 | 1.0375 |
| 1 | 1 | 1 | 1.0365 |
| 1.062 | 0.992 | 0.695 | 1.031 |
| 1.062 | 0.992 | 0.695 | 1.031 |
| 0.984 | 0.811 | 0.739 | 1.025 |
| 0.948 | 0.952 | 0.8 | 1.0185 |
| 1 | 1 | 1 | 1.016 |
| 1 | 1 | 1 | 1.0145 |
| 1 | 1 | 1 | 1.013 |
| 1 | 1 | 1 | 1.013 |
| 1 | 1 | 1 | 1.011 |
| 1.027 | 0.827 | 0.764 | 1.007 |
| 1.008 | 0.922 | 0.916 | 1.004 |
| 1 | 1 | 1 | 1 |
| 1 | 1 | 1 | 1 |
| 1 | 1 | 1 | 1 |
| 1 | 1 | 1 | 1 |
| 1 | 1 | 1 | 1 |
| 1 | 1 | 1 | 1 |
| 1 | 1 | 1 | 1 |
| 1 | 1 | 1 | 1 |
| 1 | 1 | 1 | 1 |
| 1 | 1 | 1 | 1 |
| 1 | 1 | 1 | 1 |
| 1 | 1 | 1 | 1 |
| 1 | 1 | 1 | 1 |
| 1 | 1 | 1 | 1 |
| 1 | 1 | 1 | 1 |
| 1 | 1 | 1 | 1 |
| 1 | 1 | 1 | 1 |
| 1 | 1 | 1 | 1 |
| 1 | 1 | 1 | 1 |
| 1 | 1 | 1 | 1 |
| 1 | 1 | 1 | 1 |
| 1 | 1 | 1 | 1 |
| 1 | 1 | 1 | 1 |
| 1 | 1 | 1 | 1 |
| 1 | 1 | 1 | 1 |
| 1 | 1 | 1 | 1 |
| 1 | 1 | 1 | 1 |
| 1 | 1 | 1 | 1 |
| 1 | 1 | 1 | 1 |

| | | | |
|---|---|---|---|
| 1 | 1 | 1 | 1 |
| 1 | 1 | 1 | 1 |
| 1 | 1 | 1 | 1 |
| 1 | 1 | 1 | 1 |
| 1 | 1 | 1 | 1 |
| 1 | 1 | 1 | 1 |
| 1 | 1 | 1 | 1 |
| 1 | 1 | 1 | 1 |
| 1 | 1 | 1 | 1 |
| 1 | 1 | 1 | 1 |
| 1 | 1 | 1 | 1 |
| 1 | 1 | 1 | 1 |
| 1 | 1 | 1 | 1 |
| 1 | 1 | 1 | 1 |
| 1 | 1 | 1 | 1 |
| 1 | 1 | 1 | 1 |
| 1 | 1 | 1 | 1 |
| 1 | 1 | 1 | 1 |
| 1 | 1 | 1 | 1 |
| 1 | 1 | 1 | 1 |
| 1 | 1 | 1 | 1 |
| 1 | 1 | 1 | 1 |
| 1 | 1 | 1 | 1 |
| 1 | 1 | 1 | 1 |
| 1 | 1 | 1 | 1 |
| 1 | 1 | 1 | 1 |
| 1 | 1 | 1 | 1 |
| 1 | 1 | 1 | 1 |
| 1 | 1 | 1 | 1 |
| 1 | 1 | 1 | 1 |
| 1 | 1 | 1 | 1 |
| 1 | 1 | 1 | 1 |
| 1 | 1 | 1 | 1 |
| 1 | 1 | 1 | 1 |
| 1 | 1 | 1 | 1 |
| 1 | 1 | 1 | 1 |
| 1 | 1 | 1 | 1 |
| 1 | 1 | 1 | 1 |
| 1 | 1 | 1 | 1 |
| 1 | 1 | 1 | 1 |
| 1 | 1 | 1 | 1 |
| 1 | 1 | 1 | 1 |
| 1 | 1 | 1 | 1 |

| | | | |
|---|---|---|---|
| 1 | 1 | 1 | 1 |
| 1 | 1 | 1 | 1 |
| 1 | 1 | 1 | 1 |
| 1 | 1 | 1 | 1 |
| 1 | 1 | 1 | 1 |
| 1 | 1 | 1 | 1 |
| 1 | 1 | 1 | 1 |
| 1 | 1 | 1 | 1 |
| 1 | 1 | 1 | 1 |
| 1 | 1 | 1 | 1 |
| 1 | 1 | 1 | 1 |
| 1 | 1 | 1 | 1 |
| 1 | 1 | 1 | 1 |
| 1 | 1 | 1 | 1 |
| 1 | 1 | 1 | 1 |
| 1 | 1 | 1 | 1 |
| 1 | 1 | 1 | 1 |
| 1 | 1 | 1 | 1 |
| 1 | 1 | 1 | 1 |
| 1 | 1 | 1 | 1 |
| 1 | 1 | 1 | 1 |
| 1 | 1 | 1 | 1 |
| 1 | 1 | 1 | 1 |
| 1 | 1 | 1 | 1 |
| 1 | 1 | 1 | 1 |
| 1 | 1 | 1 | 1 |
| 1 | 1 | 1 | 1 |
| 1 | 1 | 1 | 1 |
| 1 | 1 | 1 | 1 |
| 1 | 1 | 1 | 1 |
| 1 | 1 | 1 | 1 |
| 1 | 1 | 1 | 1 |
| 1 | 1 | 1 | 1 |
| 1 | 1 | 1 | 1 |
| 1 | 1 | 1 | 1 |
| 1 | 1 | 1 | 1 |
| 1 | 1 | 1 | 1 |
| 1 | 1 | 1 | 1 |
| 1 | 1 | 1 | 1 |
| 1 | 1 | 1 | 1 |
| 1 | 1 | 1 | 1 |
| 1 | 1 | 1 | 1 |
| 1 | 1 | 1 | 1 |

| | | | |
|---|---|---|---|
| 1 | 1 | 1 | 1 |
| 1 | 1 | 1 | 1 |
| 1 | 1 | 1 | 1 |
| 1 | 1 | 1 | 1 |
| 1 | 1 | 1 | 1 |
| 1 | 1 | 1 | 1 |
| 1 | 1 | 1 | 1 |
| 1 | 1 | 1 | 1 |
| 1 | 1 | 1 | 1 |
| 1 | 1 | 1 | 1 |
| 1 | 1 | 1 | 1 |
| 1 | 1 | 1 | 1 |
| 1 | 1 | 1 | 1 |
| 1 | 1 | 1 | 1 |
| 1 | 1 | 1 | 1 |
| 0.999 | 0.727 | 0.71 | 0.9995 |
| 0.998 | 0.647 | 0.67 | 0.999 |
| 1.004 | 0.857 | 0.822 | 0.997 |
| 0.994 | 0.837 | 0.87 | 0.997 |
| 1 | 1 | 1 | 0.9965 |
| 1.049 | 0.761 | 0.832 | 0.996 |
| 0.797 | 0.778 | 0.721 | 0.9925 |
| 1 | 1 | 1 | 0.989 |
| 0.978 | 1.014 | 1 | 0.989 |
| 0.963 | 0.955 | 0.843 | 0.989 |
| 1 | 0.679 | 0.702 | 0.988 |
| 0.96 | 0.788 | 0.816 | 0.9845 |
| 1 | 1 | 1 | 0.9785 |
| 0.954 | 0.856 | 0.647 | 0.977 |
| 0.913 | 0.863 | 0.693 | 0.9765 |
| 0.997 | 0.763 | 0.712 | 0.976 |
| 1.019 | 0.88 | 0.797 | 0.975 |
| 1 | 1 | 1 | 0.975 |
| 0.948 | 0.921 | 0.79 | 0.974 |
| 1 | 1 | 1 | 0.9715 |
| 1 | 1 | 1 | 0.9705 |
| 0.914 | 0.8 | 0.775 | 0.9695 |
| 0.912 | 0.822 | 0.76 | 0.969 |
| 0.937 | 1.242 | 1.005 | 0.9685 |
| 0.996 | 0.869 | 0.774 | 0.967 |
| 0.933 | 0.897 | 0.86 | 0.9665 |
| 0.921 | 0.831 | 0.823 | 0.966 |
| 0.973 | 0.822 | 0.691 | 0.965 |

| | | | |
|---|---|---|---|
| 0.927 | 0.842 | 0.912 | 0.965 |
| 1.004 | 0.92 | 0.853 | 0.9645 |
| 1.018 | 0.957 | 0.82 | 0.963 |
| 0.953 | 0.928 | 0.772 | 0.9625 |
| 0.815 | 0.717 | 0.71 | 0.962 |
| 1 | 1 | 1 | 0.9615 |
| 0.946 | 0.763 | 0.702 | 0.96 |
| 0.982 | 0.88 | 0.832 | 0.9585 |
| 0.943 | 0.898 | 0.65 | 0.9575 |
| 0.918 | 0.856 | 0.758 | 0.9565 |
| 0.872 | 0.752 | 0.725 | 0.9565 |
| 0.961 | 0.75 | 0.798 | 0.956 |
| 1.149 | 1.426 | 1.277 | 0.9525 |
| 1 | 1 | 1 | 0.952 |
| 1 | 1 | 1 | 0.9515 |
| 1.04 | 1.178 | 0.604 | 0.95 |
| 0.98 | 0.91 | 0.787 | 0.9495 |
| 0.996 | 0.736 | 0.745 | 0.949 |
| 1 | 1 | 1 | 0.949 |
| 0.928 | 0.96 | 0.868 | 0.9485 |
| 0.888 | 0.655 | 0.73 | 0.9485 |
| 1.02 | 0.939 | 0.859 | 0.948 |
| 1.034 | 1.034 | 1.247 | 0.9475 |
| 0.908 | 1 | 0.792 | 0.9475 |
| 1 | 1 | 1 | 0.9475 |
| 1 | 1 | 1 | 0.9475 |
| 1 | 1 | 1 | 0.947 |
| 0.935 | 0.873 | 0.807 | 0.945 |
| 0.985 | 0.897 | 0.829 | 0.944 |
| 0.999 | 0.912 | 0.825 | 0.944 |
| 0.938 | 0.966 | 0.733 | 0.9435 |
| 1.02 | 0.903 | 0.871 | 0.9435 |
| 1.037 | 0.911 | 0.767 | 0.9425 |
| 0.917 | 0.868 | 0.889 | 0.942 |
| 1 | 1 | 1 | 0.9415 |
| 1.02 | 0.971 | 0.79 | 0.939 |
| 1.017 | 0.942 | 0.864 | 0.939 |
| 1 | 1 | 1 | 0.9385 |
| 0.993 | 1.202 | 1.055 | 0.938 |
| 0.873 | 1.074 | 0.725 | 0.9365 |
| 0.934 | 1.142 | 0.744 | 0.9355 |
| 0.871 | 0.78 | 0.722 | 0.9355 |
| 1.009 | 0.963 | 0.799 | 0.935 |

| | | | |
|---|---|---|---|
| 1 | 1 | 1 | 0.935 |
| 0.901 | 0.843 | 0.818 | 0.933 |
| 0.996 | 0.694 | 0.781 | 0.9325 |
| 0.864 | 0.664 | 0.56 | 0.932 |
| 0.914 | 0.785 | 0.795 | 0.93 |
| 1 | 1 | 1 | 0.9295 |
| 0.858 | 0.795 | 0.797 | 0.929 |
| 1.165 | 0.823 | 0.823 | 0.9285 |
| 0.947 | 0.753 | 0.8 | 0.9285 |
| 0.877 | 0.826 | 0.732 | 0.9285 |
| 1 | 1 | 1 | 0.9275 |
| 1 | 1 | 1 | 0.9275 |
| 0.854 | 0.728 | 0.726 | 0.927 |
| 0.853 | 0.766 | 0.793 | 0.9265 |
| 1 | 1 | 1 | 0.9265 |
| 0.853 | 0.848 | 0.645 | 0.9265 |
| 1 | 1 | 1 | 0.9255 |
| 0.963 | 0.914 | 0.804 | 0.9255 |
| 0.986 | 0.94 | 0.767 | 0.925 |
| 0.85 | 0.708 | 0.719 | 0.925 |
| 1 | 1 | 1 | 0.9245 |
| 0.855 | 0.816 | 0.792 | 0.9235 |
| 0.846 | 0.861 | 0.711 | 0.923 |
| 0.89 | 0.805 | 0.79 | 0.9225 |
| 0.845 | 0.938 | 0.967 | 0.9225 |
| 1 | 1 | 1 | 0.922 |
| 0.96 | 0.865 | 0.734 | 0.922 |
| 0.841 | 0.832 | 0.655 | 0.9205 |
| 0.841 | 0.751 | 0.707 | 0.9205 |
| 1 | 1 | 1 | 0.9205 |
| 0.891 | 0.689 | 0.661 | 0.9205 |
| 0.993 | 0.728 | 0.922 | 0.92 |
| 0.876 | 1.108 | 1.034 | 0.9195 |
| 0.839 | 0.708 | 0.784 | 0.9195 |
| 0.838 | 0.765 | 0.722 | 0.919 |
| 0.838 | 0.757 | 0.769 | 0.919 |
| 0.836 | 0.871 | 0.818 | 0.918 |
| 0.955 | 0.821 | 0.687 | 0.917 |
| 0.831 | 0.817 | 0.71 | 0.9155 |
| 0.934 | 0.948 | 0.89 | 0.915 |
| 1 | 1 | 1 | 0.9145 |
| 0.853 | 0.78 | 0.668 | 0.9135 |
| 1 | 1 | 1 | 0.912 |

| | | | |
|---|---|---|---|
| 1 | 1 | 1 | 0.91 |
| 1 | 1 | 1 | 0.91 |
| 1 | 1 | 1 | 0.909 |
| 1 | 1 | 1 | 0.9085 |
| 0.886 | 0.844 | 0.78 | 0.908 |
| 0.923 | 0.832 | 0.717 | 0.908 |
| 0.965 | 0.862 | 0.764 | 0.907 |
| 0.893 | 0.774 | 0.754 | 0.907 |
| 1 | 1 | 1 | 0.906 |
| 0.842 | 0.853 | 0.719 | 0.906 |
| 0.812 | 0.739 | 0.694 | 0.906 |
| 0.929 | 0.841 | 0.891 | 0.9035 |
| 0.806 | 0.72 | 0.743 | 0.903 |
| 1 | 1 | 1 | 0.902 |
| 0.835 | 0.756 | 0.672 | 0.9015 |
| 0.803 | 0.796 | 0.787 | 0.9015 |
| 0.877 | 0.755 | 0.774 | 0.901 |
| 0.881 | 0.893 | 0.802 | 0.9005 |
| 0.998 | 0.873 | 0.769 | 0.9005 |
| 0.896 | 0.919 | 0.694 | 0.9005 |
| 1 | 1 | 1 | 0.9 |
| 0.892 | 1.008 | 0.811 | 0.8995 |
| 1 | 1 | 1 | 0.899 |
| 0.878 | 1.039 | 0.861 | 0.8985 |
| 0.879 | 0.789 | 0.704 | 0.898 |
| 1 | 1 | 1 | 0.897 |
| 0.934 | 0.821 | 0.824 | 0.896 |
| 1 | 1 | 1 | 0.896 |
| 0.771 | 0.82 | 0.677 | 0.893 |
| 0.933 | 0.77 | 0.803 | 0.893 |
| 0.785 | 0.683 | 0.69 | 0.8925 |
| 1.091 | 0.818 | 0.707 | 0.8925 |
| 0.861 | 0.901 | 0.824 | 0.892 |
| 0.784 | 0.778 | 0.663 | 0.892 |
| 0.782 | 0.763 | 0.724 | 0.891 |
| 1 | 1 | 1 | 0.8905 |
| 1 | 1 | 1 | 0.8905 |
| 0.78 | 0.763 | 0.73 | 0.89 |
| 1 | 1 | 1 | 0.89 |
| 0.778 | 0.809 | 0.666 | 0.889 |
| 0.932 | 0.766 | 0.802 | 0.888 |
| 0.775 | 0.887 | 0.775 | 0.8875 |
| 1 | 1 | 1 | 0.887 |

| | | | |
|---|---|---|---|
| 1 | 1 | 1 | 0.887 |
| 0.855 | 0.883 | 0.83 | 0.8855 |
| 0.919 | 0.878 | 0.728 | 0.884 |
| 0.923 | 0.794 | 0.808 | 0.884 |
| 0.859 | 0.807 | 0.731 | 0.883 |
| 0.827 | 0.788 | 0.772 | 0.883 |
| 0.764 | 0.688 | 0.608 | 0.882 |
| 0.895 | 0.821 | 0.735 | 0.882 |
| 0.965 | 0.859 | 0.819 | 0.8815 |
| 0.763 | 0.838 | 0.723 | 0.8815 |
| 0.901 | 0.88 | 0.808 | 0.8815 |
| 0.872 | 0.915 | 0.836 | 0.8815 |
| 0.861 | 0.976 | 0.82 | 0.881 |
| 0.762 | 0.788 | 0.74 | 0.881 |
| 0.924 | 0.902 | 0.759 | 0.8805 |
| 0.85 | 0.809 | 0.74 | 0.88 |
| 0.76 | 0.797 | 0.75 | 0.88 |
| 0.849 | 0.787 | 0.747 | 0.8795 |
| 0.983 | 0.916 | 0.921 | 0.8775 |
| 1 | 1 | 1 | 0.8775 |
| 0.754 | 0.771 | 0.747 | 0.877 |
| 0.989 | 0.827 | 0.796 | 0.8765 |
| 0.751 | 0.909 | 0.796 | 0.8755 |
| 0.825 | 0.868 | 0.802 | 0.875 |
| 1 | 1 | 1 | 0.8745 |
| 0.749 | 0.918 | 0.829 | 0.8745 |
| 1 | 1 | 1 | 0.8745 |
| 1 | 1 | 1 | 0.8745 |
| 0.934 | 0.883 | 0.953 | 0.8735 |
| 0.806 | 0.713 | 0.709 | 0.873 |
| 0.934 | 0.928 | 0.823 | 0.8725 |
| 0.741 | 0.895 | 0.828 | 0.872 |
| 0.889 | 0.799 | 0.841 | 0.8715 |
| 0.943 | 0.913 | 0.771 | 0.871 |
| 0.742 | 0.709 | 0.776 | 0.871 |
| 0.741 | 0.838 | 0.796 | 0.8705 |
| 0.89 | 0.858 | 0.77 | 0.87 |
| 0.911 | 0.789 | 0.729 | 0.8695 |
| 0.851 | 0.877 | 0.843 | 0.8695 |
| 0.858 | 0.662 | 0.618 | 0.869 |
| 0.807 | 0.877 | 0.817 | 0.869 |
| 0.874 | 0.791 | 0.749 | 0.869 |
| 0.833 | 0.884 | 0.815 | 0.8685 |

| | | | |
|---|---|---|---|
| 0.893 | 0.786 | 0.744 | 0.8675 |
| 0.854 | 0.834 | 0.803 | 0.8665 |
| 0.826 | 0.675 | 0.692 | 0.866 |
| 1 | 1 | 1 | 0.866 |
| 0.879 | 0.733 | 0.731 | 0.864 |
| 0.728 | 0.808 | 0.838 | 0.864 |
| 0.901 | 0.913 | 0.744 | 0.863 |
| 0.948 | 0.843 | 0.813 | 0.863 |
| 0.829 | 0.714 | 0.566 | 0.863 |
| 0.836 | 0.794 | 0.759 | 0.862 |
| 0.825 | 0.84 | 0.73 | 0.8615 |
| 1 | 1 | 1 | 0.8605 |
| 0.818 | 0.746 | 0.729 | 0.86 |
| 1 | 1 | 1 | 0.8595 |
| 0.874 | 0.809 | 0.762 | 0.859 |
| 0.885 | 0.825 | 0.782 | 0.8585 |
| 0.787 | 0.963 | 0.807 | 0.858 |
| 0.878 | 0.874 | 0.675 | 0.858 |
| 0.953 | 0.894 | 0.762 | 0.8575 |
| 0.855 | 0.908 | 0.707 | 0.857 |
| 0.801 | 0.844 | 0.692 | 0.855 |
| 1 | 1 | 1 | 0.8545 |
| 0.915 | 0.685 | 0.667 | 0.8545 |
| 0.921 | 0.777 | 0.75 | 0.8545 |
| 0.706 | 0.789 | 0.678 | 0.853 |
| 0.785 | 0.623 | 0.734 | 0.853 |
| 0.901 | 0.973 | 0.773 | 0.853 |
| 0.705 | 0.813 | 0.757 | 0.8525 |
| 0.705 | 0.509 | 0.58 | 0.8525 |
| 0.704 | 0.733 | 0.67 | 0.852 |
| 0.702 | 0.585 | 0.635 | 0.851 |
| 0.702 | 0.585 | 0.635 | 0.851 |
| 0.701 | 0.648 | 0.663 | 0.8505 |
| 0.875 | 0.93 | 0.781 | 0.85 |
| 0.847 | 0.742 | 0.71 | 0.8495 |
| 0.79 | 0.863 | 0.809 | 0.849 |
| 0.698 | 0.669 | 0.705 | 0.849 |
| 0.86 | 0.706 | 0.715 | 0.849 |
| 0.697 | 0.611 | 0.653 | 0.8485 |
| 0.759 | 0.79 | 0.702 | 0.8475 |
| 0.826 | 0.748 | 0.745 | 0.847 |
| 0.857 | 0.98 | 0.881 | 0.8465 |
| 1 | 1 | 1 | 0.8465 |

| | | | |
|---|---|---|---|
| 0.834 | 0.834 | 0.711 | 0.8465 |
| 0.829 | 0.933 | 0.707 | 0.846 |
| 0.851 | 0.801 | 0.743 | 0.846 |
| 0.91 | 0.878 | 0.824 | 0.8455 |
| 0.838 | 0.814 | 0.648 | 0.8455 |
| 1 | 1 | 1 | 0.8445 |
| 0.924 | 0.876 | 0.777 | 0.844 |
| 0.833 | 0.852 | 0.771 | 0.844 |
| 0.798 | 0.862 | 0.803 | 0.844 |
| 1 | 1 | 1 | 0.8435 |
| 0.853 | 0.864 | 0.784 | 0.8425 |
| 0.846 | 0.83 | 0.928 | 0.842 |
| 0.866 | 0.861 | 0.779 | 0.84 |
| 1 | 1 | 1 | 0.8395 |
| 0.855 | 0.861 | 0.792 | 0.8395 |
| 0.811 | 0.764 | 0.688 | 0.8395 |
| 0.893 | 0.816 | 0.779 | 0.839 |
| 0.907 | 0.851 | 0.761 | 0.8385 |
| 0.825 | 0.873 | 0.778 | 0.8375 |
| 0.838 | 0.831 | 0.709 | 0.8375 |
| 0.852 | 0.82 | 0.765 | 0.8375 |
| 0.865 | 0.926 | 0.794 | 0.837 |
| 0.846 | 0.817 | 0.794 | 0.836 |
| 0.806 | 0.828 | 0.746 | 0.8345 |
| 0.832 | 0.921 | 0.789 | 0.8345 |
| 0.835 | 0.794 | 0.73 | 0.834 |
| 0.849 | 0.83 | 0.805 | 0.8335 |
| 0.925 | 0.958 | 0.792 | 0.8335 |
| 0.905 | 1.155 | 0.887 | 0.833 |
| 0.835 | 0.752 | 0.737 | 0.833 |
| 0.747 | 0.842 | 0.82 | 0.8325 |
| 0.799 | 0.703 | 0.618 | 0.8325 |
| 0.839 | 0.754 | 0.755 | 0.8325 |
| 0.797 | 0.802 | 0.683 | 0.8315 |
| 0.837 | 0.847 | 0.774 | 0.8315 |
| 0.753 | 0.721 | 0.67 | 0.831 |
| 1 | 1 | 1 | 0.8305 |
| 0.779 | 0.775 | 0.76 | 0.8305 |
| 0.783 | 0.702 | 0.695 | 0.83 |
| 1 | 1 | 1 | 0.8295 |
| 0.799 | 0.811 | 0.735 | 0.828 |
| 0.656 | 0.589 | 0.712 | 0.828 |
| 0.924 | 0.937 | 0.782 | 0.8275 |

| | | | |
|---|---|---|---|
| 0.966 | 0.889 | 0.692 | 0.827 |
| 0.775 | 0.852 | 0.862 | 0.826 |
| 0.91 | 0.835 | 0.776 | 0.826 |
| 0.839 | 0.756 | 0.847 | 0.8255 |
| 0.792 | 0.777 | 0.722 | 0.8255 |
| 0.829 | 1.21 | 0.906 | 0.825 |
| 0.886 | 0.796 | 0.663 | 0.825 |
| 0.648 | 0.645 | 0.508 | 0.824 |
| 0.841 | 0.701 | 0.709 | 0.823 |
| 0.873 | 0.832 | 0.718 | 0.8225 |
| 0.823 | 0.784 | 0.743 | 0.822 |
| 1 | 1 | 1 | 0.8215 |
| 0.862 | 0.885 | 0.858 | 0.8205 |
| 1 | 1 | 1 | 0.82 |
| 0.806 | 0.832 | 0.73 | 0.8195 |
| 0.841 | 0.749 | 0.766 | 0.8195 |
| 0.788 | 0.671 | 0.684 | 0.8195 |
| 0.754 | 0.721 | 0.733 | 0.8185 |
| 0.893 | 0.809 | 0.823 | 0.818 |
| 0.784 | 0.842 | 0.791 | 0.8175 |
| 0.753 | 0.862 | 0.756 | 0.8175 |
| 0.634 | 0.737 | 0.664 | 0.817 |
| 0.818 | 0.848 | 0.762 | 0.8165 |
| 0.84 | 0.837 | 0.763 | 0.816 |
| 0.798 | 0.916 | 0.777 | 0.8155 |
| 0.8 | 0.658 | 0.729 | 0.815 |
| 0.891 | 0.78 | 0.763 | 0.815 |
| 0.93 | 0.919 | 0.792 | 0.815 |
| 0.741 | 0.843 | 0.716 | 0.8145 |
| 0.795 | 0.821 | 0.721 | 0.8135 |
| 0.741 | 0.792 | 0.659 | 0.8125 |
| 0.835 | 0.938 | 0.577 | 0.8115 |
| 0.801 | 0.844 | 0.692 | 0.8105 |
| 1 | 1 | 1 | 0.81 |
| 0.863 | 0.789 | 0.713 | 0.8095 |
| 0.843 | 0.782 | 0.772 | 0.8085 |
| 0.823 | 0.955 | 0.885 | 0.808 |
| 0.824 | 0.8 | 0.766 | 0.8055 |
| 0.796 | 0.723 | 0.743 | 0.8045 |
| 0.791 | 0.683 | 0.677 | 0.8045 |
| 0.811 | 1.048 | 0.874 | 0.804 |
| 0.802 | 0.754 | 0.672 | 0.804 |
| 0.789 | 0.866 | 0.657 | 0.802 |

| | | | |
|---|---|---|---|
| 0.806 | 0.781 | 0.71 | 0.8015 |
| 0.804 | 0.79 | 0.761 | 0.8015 |
| 0.804 | 0.603 | 0.685 | 0.8 |
| 0.721 | 0.805 | 0.622 | 0.798 |
| 0.832 | 0.848 | 0.811 | 0.7975 |
| 0.82 | 0.806 | 0.775 | 0.795 |
| 0.819 | 0.791 | 0.694 | 0.7945 |
| 0.822 | 0.83 | 0.758 | 0.794 |
| 0.809 | 0.884 | 0.754 | 0.793 |
| 0.585 | 0.714 | 0.772 | 0.7925 |
| 0.799 | 0.87 | 0.723 | 0.7925 |
| 0.861 | 0.875 | 0.825 | 0.792 |
| 0.827 | 0.695 | 0.783 | 0.792 |
| 0.829 | 0.864 | 0.692 | 0.79 |
| 0.875 | 0.946 | 0.789 | 0.7895 |
| 0.786 | 0.748 | 0.782 | 0.789 |
| 0.787 | 0.862 | 0.729 | 0.7885 |
| 1 | 1 | 1 | 0.788 |
| 0.722 | 0.757 | 0.692 | 0.7875 |
| 0.786 | 0.712 | 0.642 | 0.7875 |
| 0.753 | 0.721 | 0.67 | 0.7865 |
| 0.769 | 0.754 | 0.686 | 0.7865 |
| 0.81 | 0.774 | 0.783 | 0.7865 |
| 0.828 | 0.916 | 0.645 | 0.786 |
| 0.712 | 0.801 | 0.743 | 0.786 |
| 0.736 | 0.583 | 0.538 | 0.785 |
| 0.793 | 0.628 | 0.682 | 0.783 |
| 0.765 | 0.812 | 0.678 | 0.7825 |
| 0.792 | 0.823 | 0.749 | 0.7825 |
| 0.774 | 0.764 | 0.709 | 0.7815 |
| 0.845 | 0.843 | 0.778 | 0.7805 |
| 0.747 | 0.772 | 0.83 | 0.7795 |
| 0.797 | 0.706 | 0.681 | 0.7785 |
| 0.793 | 0.688 | 0.687 | 0.778 |
| 0.754 | 0.748 | 0.813 | 0.778 |
| 0.746 | 0.836 | 0.787 | 0.778 |
| 0.779 | 0.736 | 0.688 | 0.777 |
| 0.775 | 0.783 | 0.738 | 0.777 |
| 0.81 | 0.751 | 0.751 | 0.7765 |
| 0.804 | 0.816 | 0.741 | 0.7765 |
| 0.763 | 0.698 | 0.727 | 0.7755 |
| 0.804 | 0.814 | 0.767 | 0.775 |
| 0.745 | 0.771 | 0.692 | 0.7735 |

| | | | |
|---|---|---|---|
| 0.732 | 0.638 | 0.681 | 0.773 |
| 0.774 | 0.719 | 0.649 | 0.772 |
| 0.789 | 0.915 | 0.732 | 0.771 |
| 0.789 | 0.808 | 0.707 | 0.7705 |
| 0.537 | 0.56 | 0.61 | 0.7685 |
| 0.752 | 0.812 | 0.738 | 0.7685 |
| 0.77 | 0.766 | 0.736 | 0.7685 |
| 0.742 | 0.743 | 0.637 | 0.768 |
| 0.799 | 0.789 | 0.762 | 0.7665 |
| 0.705 | 0.776 | 0.761 | 0.7655 |
| 0.771 | 0.918 | 0.777 | 0.7655 |
| 0.818 | 0.933 | 0.808 | 0.765 |
| 0.795 | 0.816 | 0.696 | 0.7635 |
| 0.828 | 0.646 | 0.678 | 0.763 |
| 0.781 | 0.817 | 0.606 | 0.7615 |
| 0.695 | 0.652 | 0.671 | 0.7595 |
| 0.785 | 0.838 | 0.866 | 0.759 |
| 0.64 | 0.698 | 0.673 | 0.7585 |
| 0.762 | 0.794 | 0.773 | 0.758 |
| 0.83 | 0.638 | 0.544 | 0.7575 |
| 0.721 | 0.82 | 0.761 | 0.7575 |
| 0.768 | 0.645 | 0.694 | 0.757 |
| 0.693 | 0.719 | 0.709 | 0.7555 |
| 0.715 | 0.763 | 0.752 | 0.755 |
| 0.774 | 0.859 | 0.841 | 0.755 |
| 0.587 | 0.814 | 0.56 | 0.7545 |
| 0.753 | 0.727 | 0.761 | 0.7545 |
| 0.7 | 0.734 | 0.775 | 0.7545 |
| 0.786 | 0.692 | 0.709 | 0.754 |
| 0.785 | 0.741 | 0.669 | 0.752 |
| 0.723 | 0.847 | 0.802 | 0.7515 |
| 0.775 | 0.641 | 0.631 | 0.7515 |
| 0.789 | 0.821 | 0.708 | 0.751 |
| 0.776 | 0.736 | 0.761 | 0.7505 |
| 0.659 | 0.694 | 0.598 | 0.7495 |
| 0.75 | 0.818 | 0.724 | 0.7495 |
| 0.857 | 0.783 | 0.765 | 0.749 |
| 0.754 | 0.924 | 0.717 | 0.749 |
| 0.773 | 0.794 | 0.742 | 0.7485 |
| 0.792 | 0.577 | 0.618 | 0.7475 |
| 0.768 | 0.799 | 0.806 | 0.747 |
| 0.801 | 0.786 | 0.713 | 0.747 |
| 0.795 | 0.545 | 0.568 | 0.746 |

| | | | |
|---|---|---|---|
| 0.749 | 0.775 | 0.802 | 0.744 |
| 0.713 | 0.771 | 0.769 | 0.7435 |
| 0.764 | 0.795 | 0.786 | 0.7425 |
| 0.667 | 0.718 | 0.685 | 0.741 |
| 0.767 | 0.823 | 0.726 | 0.7405 |
| 0.731 | 0.757 | 0.701 | 0.7405 |
| 0.72 | 0.797 | 0.765 | 0.737 |
| 0.697 | 0.712 | 0.701 | 0.7345 |
| 0.613 | 0.447 | 0.55 | 0.734 |
| 0.797 | 0.696 | 0.505 | 0.73 |
| 0.676 | 0.864 | 0.831 | 0.729 |
| 0.77 | 0.778 | 0.804 | 0.728 |
| 0.752 | 0.835 | 0.733 | 0.727 |
| 0.71 | 0.797 | 0.698 | 0.7255 |
| 0.66 | 0.773 | 0.733 | 0.7215 |
| 0.681 | 0.595 | 0.676 | 0.7205 |
| 0.741 | 0.805 | 0.716 | 0.7185 |
| 0.725 | 0.755 | 0.653 | 0.712 |
| 0.755 | 0.775 | 0.76 | 0.7115 |
| 0.71 | 0.729 | 0.645 | 0.711 |
| 0.713 | 0.468 | 0.507 | 0.71 |
| 0.704 | 0.738 | 0.704 | 0.709 |
| 0.73 | 0.67 | 0.653 | 0.7045 |
| 0.673 | 0.711 | 0.767 | 0.7035 |
| 0.698 | 0.737 | 0.723 | 0.7035 |
| 0.688 | 0.754 | 0.708 | 0.701 |
| 0.644 | 0.687 | 0.627 | 0.7005 |
| 0.709 | 0.8 | 0.767 | 0.6915 |
| 0.726 | 0.8 | 0.751 | 0.69 |
| 0.676 | 0.815 | 0.795 | 0.6885 |
| 0.702 | 0.686 | 0.682 | 0.6875 |
| 0.756 | 0.769 | 0.807 | 0.684 |
| 0.674 | 0.598 | 0.548 | 0.679 |
| 0.681 | 0.754 | 0.745 | 0.6785 |
| 0.675 | 0.758 | 0.778 | 0.678 |
| 0.706 | 0.604 | 0.679 | 0.673 |
| 0.726 | 0.747 | 0.679 | 0.6725 |
| 0.695 | 0.713 | 0.675 | 0.6695 |
| 0.661 | 0.569 | 0.631 | 0.6665 |
| 0.668 | 0.69 | 0.699 | 0.6645 |
| 0.601 | 0.689 | 0.718 | 0.6615 |
| 0.639 | 0.746 | 0.838 | 0.655 |
| 0.477 | 0.736 | 0.706 | 0.6535 |

| | | | |
|---|---|---|---|
| 0.694 | 0.594 | 0.741 | 0.6515 |
| 0.653 | 0.659 | 0.622 | 0.648 |
| 0.682 | 0.674 | 0.669 | 0.645 |
| 0.576 | 0.673 | 0.578 | 0.632 |
| 0.614 | 0.646 | 0.78 | 0.624 |
| 0.609 | 0.601 | 0.819 | 0.623 |
| 0.684 | 0.803 | 0.685 | 0.617 |
| 0.571 | 0.81 | 0.612 | 0.6105 |
| 0.636 | 0.77 | 0.822 | 0.6105 |
| 0.653 | 0.4 | 0.8 | 0.6005 |
| 0.625 | 1.023 | 0.898 | 0.576 |
| 0.82 | 0.645 | 0.727 | 0.563 |
| 0.626 | 0.739 | 0.654 | 0.5585 |
| 0.583 | 0.466 | 0.536 | 0.55 |
| 0.57 | 0.437 | 0.446 | 0.52 |
| 0.492 | 0.339 | 0.466 | 0.512 |
