## Supplemental table 3 for "A TMT-based quantitative proteomics approach toward α-syn PFF associated Lewy Body Dementia (LBD) using α-syn PFF-injected mouse brain tissues"

| GOID |
| --- |
| GO:0021762 |
| KEGG:04540 |
| KEGG:04721 |
| KEGG:00020 |
| KEGG:05012 |
| KEGG:05016 |
| KEGG:00010 |
| KEGG:00620 |
| KEGG:04066 |
| GO:0099003 |
| GO:0099504 |
| GO:0036465 |
| GO:0016079 |
| KEGG:00010 |
| KEGG:04020 |
| KEGG:04066 |
| KEGG:04114 |
| KEGG:04261 |
| KEGG:04720 |
| KEGG:04911 |
| KEGG:04922 |
| KEGG:04925 |
| KEGG:04961 |
| KEGG:04971 |
| KEGG:05031 |

| GOTerm |
| --- |
| substantia nigra development |
| Gap junction |
| Synaptic vesicle cycle |
| Citrate cycle (TCA cycle) |
| Parkinson disease |
| Huntington disease |
| Glycolysis / Gluconeogenesis |
| Pyruvate metabolism |
| HIF-1 signaling pathway |
| vesicle-mediated transport in synapse |
| synaptic vesicle cycle |
| synaptic vesicle recycling |
| synaptic vesicle exocytosis |
| Glycolysis / Gluconeogenesis |
| Calcium signaling pathway |
| HIF-1 signaling pathway |
| Oocyte meiosis |
| Adrenergic signaling in cardiomyocytes |
| Long-term potentiation |
| Insulin secretion |
| Glucagon signaling pathway |
| Aldosterone synthesis and secretion |
| Endocrine and other factor-regulated calcium reabsorption |
| Gastric acid secretion |
| Amphetamine addiction |

| Ontology Source | Term PValue |
|---|---|
| GO_BiologicalProcess-EBI-UniProt-GOA-ACAP-ARAP_30.07.2022_00h00 | 0.00 |
| KEGG_08.05.2020 | 0.00 |
| KEGG_08.05.2020 | 0.00 |
| KEGG_08.05.2020 | 0.00 |
| KEGG_08.05.2020 | 0.00 |
| KEGG_08.05.2020 | 0.00 |
| KEGG_08.05.2020 | 0.00 |
| KEGG_08.05.2020 | 0.00 |
| KEGG_08.05.2020 | 0.00 |
| GO_BiologicalProcess-EBI-UniProt-GOA-ACAP-ARAP_30.07.2022_00h00 | 0.00 |
| GO_BiologicalProcess-EBI-UniProt-GOA-ACAP-ARAP_30.07.2022_00h00 | 0.00 |
| GO_BiologicalProcess-EBI-UniProt-GOA-ACAP-ARAP_30.07.2022_00h00 | 0.00 |
| GO_BiologicalProcess-EBI-UniProt-GOA-ACAP-ARAP_30.07.2022_00h00 | 0.00 |
| KEGG_08.05.2020 | 0.00 |
| KEGG_08.05.2020 | 0.00 |
| KEGG_08.05.2020 | 0.00 |
| KEGG_08.05.2020 | 0.00 |
| KEGG_08.05.2020 | 0.00 |
| KEGG_08.05.2020 | 0.00 |
| KEGG_08.05.2020 | 0.00 |
| KEGG_08.05.2020 | 0.00 |
| KEGG_08.05.2020 | 0.00 |
| KEGG_08.05.2020 | 0.00 |
| KEGG_08.05.2020 | 0.00 |
| KEGG_08.05.2020 | 0.00 |

| Term PValue Corrected with Bonferroni step dow | Group PValu |
|---|---|
| 0.00 | 0.00 |
| 0.00 | 0.00 |
| 0.00 | 0.00 |
| 0.00 | 0.00 |
| 0.00 | 0.00 |
| 0.00 | 0.00 |
| 0.00 | 0.00 |
| 0.00 | 0.00 |
| 0.00 | 0.00 |
| 0.00 | 0.00 |
| 0.00 | 0.00 |
| 0.00 | 0.00 |
| 0.00 | 0.00 |
| 0.00 | 0.00 |
| 0.00 | 0.00 |
| 0.00 | 0.00 |
| 0.00 | 0.00 |
| 0.00 | 0.00 |
| 0.00 | 0.00 |
| 0.00 | 0.00 |
| 0.00 | 0.00 |
| 0.00 | 0.00 |
| 0.00 | 0.00 |
| 0.00 | 0.00 |
| 0.00 | 0.00 |

| Group PValue Corrected with Bonferroni step dow | GOLevels | GOGrou |
|---|---|---|
| 0.00 | [4, 6, 7, 8, 9] | Group0 |
| 0.00 | [-1] | Group1 |
| 0.00 | [-1] | Group2 |
| 0.00 | [-1] | Group3 |
| 0.00 | [-1] | Group4 |
| 0.00 | [-1] | Group4 |
| 0.00 | [-1] | Group5 |
| 0.00 | [-1] | Group5 |
| 0.00 | [-1] | Group5 |
| 0.00 | [5] | Group6 |
| 0.00 | [4, 6] | Group6 |
| 0.00 | [4, 5, 7] | Group6 |
| 0.00 | [4, 5, 6, 7, 8, 9] | Group6 |
| 0.00 | [-1] | Group7 |
| 0.00 | [-1] | Group7 |
| 0.00 | [-1] | Group7 |
| 0.00 | [-1] | Group7 |
| 0.00 | [-1] | Group7 |
| 0.00 | [-1] | Group7 |
| 0.00 | [-1] | Group7 |
| 0.00 | [-1] | Group7 |
| 0.00 | [-1] | Group7 |
| 0.00 | [-1] | Group7 |
| 0.00 | [-1] | Group7 |
| 0.00 | [-1] | Group7 |

| % Associated Gen | Nr. Genes |
|---|---|
| 18.18 | 10.00 |
| 10.23 | 9.00 |
| 16.67 | 13.00 |
| 23.33 | 7.00 |
| 9.24 | 23.00 |
| 8.49 | 23.00 |
| 13.24 | 9.00 |
| 20.51 | 8.00 |
| 10.09 | 11.00 |
| 8.62 | 20.00 |
| 8.10 | 17.00 |
| 13.58 | 11.00 |
| 8.00 | 10.00 |
| 13.24 | 9.00 |
| 8.29 | 16.00 |
| 10.09 | 11.00 |
| 8.59 | 11.00 |
| 8.05 | 12.00 |
| 13.43 | 9.00 |
| 10.47 | 9.00 |
| 10.38 | 11.00 |
| 9.18 | 9.00 |
| 22.64 | 12.00 |
| 13.16 | 10.00 |
| 13.04 | 9.00 |

| Associated Genes Found |
|---|
| [CALM2, CKB, CNP, DYNLL1, GLUD1, INA, NDRG2, PLP1, YWHAE, YWHAH] |
| [GNAI1, GNAQ, PRKCB, PRKCG, TUBA4A, TUBB2A, TUBB2B, TUBB3, TUBB4A] |
| [AP2A1, AP2B1, AP2M1, ATP6V1C1, ATP6V1E1, CLTB, CLTC, CPLX1, NSF, RAB3A, SLC17A7, STX1A, SYT1] |
| [DLAT, IDH3B, IDH3G, MDH2, PDHA1, SDHB, SUCLA2] |
| [CALM2, CAMK2A, CAMK2B, CAMK2D, CYCS, GNAI1, NDUFA4, NDUFA5, NDUFA7, NDUFS4, NDUFV1, SDHB, SL[illegible] |
| [AP2A1, AP2B1, AP2M1, CLTB, CLTC, CYCS, GNAQ, NDUFA4, NDUFA5, NDUFA7, NDUFS4, NDUFV1, SDHB, SLC[illegible] |
| [ALDOA, DLAT, ENO1, GPI, LDHA, PDHA1, PGK1, PGM1, PKM] |
| [ACAT1, ACAT2, DLAT, GLO1, LDHA, MDH2, PDHA1, PKM] |
| [ALDOA, CAMK2A, CAMK2B, CAMK2D, ENO1, LDHA, PDHA1, PGK1, PRKCB, PRKCG, RPS6] |
| [AMPH, AP2A1, AP2B1, AP2M1, AP3B2, CADPS, CPLX1, NAPB, PACSIN1, PPP3R1, PRKCB, PRKCG, RAB3A, S[illegible] |
| [AMPH, AP2M1, AP3B2, CADPS, CPLX1, NAPB, PACSIN1, PRKCB, PRKCG, RAB3A, SH3GL2, SLC17A7, SNCA, [illegible] |
| [AMPH, AP2M1, AP3B2, PACSIN1, RAB3A, SH3GL2, SLC17A7, SNCA, STX1A, SYNJ1, SYT1] |
| [CADPS, CPLX1, NAPB, PRKCB, PRKCG, RAB3A, SNCA, STX1A, SYNJ1, SYT1] |
| [ALDOA, DLAT, ENO1, GPI, LDHA, PDHA1, PGK1, PGM1, PKM] |
| [ATP2B2, CALM2, CAMK2A, CAMK2B, CAMK2D, GNAQ, PPP3CA, PPP3R1, PRKCB, PRKCG, SLC25A4, SLC25A[illegible] |
| [ALDOA, CAMK2A, CAMK2B, CAMK2D, ENO1, LDHA, PDHA1, PGK1, PRKCB, PRKCG, RPS6] |
| [CALM2, CAMK2A, CAMK2B, CAMK2D, PPP2R1A, PPP3CA, PPP3R1, YWHAB, YWHAE, YWHAH, YWHAZ] |
| [ATP1A1, ATP2B2, CALM2, CAMK2A, CAMK2B, CAMK2D, GNAI1, GNAQ, PPP2R1A, SLC8A1, SLC8A2, TPM1] |
| [CALM2, CAMK2A, CAMK2B, CAMK2D, GNAQ, PPP3CA, PPP3R1, PRKCB, PRKCG] |
| [ATP1A1, CAMK2A, CAMK2B, CAMK2D, GNAQ, PRKCB, PRKCG, RAB3A, STX1A] |
| [CALM2, CAMK2A, CAMK2B, CAMK2D, GNAQ, LDHA, PDHA1, PKM, PPP3CA, PPP3R1, PYGM] |
| [ATP1A1, ATP2B2, CALM2, CAMK2A, CAMK2B, CAMK2D, GNAQ, PRKCB, PRKCG] |
| [AP2A1, AP2B1, AP2M1, ATP1A1, ATP2B2, CLTB, CLTC, GNAQ, PRKCB, PRKCG, SLC8A1, SLC8A2] |
| [ATP1A1, CA2, CALM2, CAMK2A, CAMK2B, CAMK2D, GNAI1, GNAQ, PRKCB, PRKCG] |
| [CALM2, CAMK2A, CAMK2B, CAMK2D, PPP3CA, PPP3R1, PRKCB, PRKCG, STX1A] |
