## Supplemental table 4 for "A TMT-based quantitative proteomics approach toward α-syn PFF associated Lewy Body Dementia (LBD) using α-syn PFF-injected mouse brain tissues"

| 1-month after a |
|---|
| **GO ID** |
| 5515 |
| 166 |
| 32553 |
| 32555 |
| 17076 |
| 3924 |
| 5525 |
| 19001 |
| 32561 |
| 5488 |
| 48037 |

| 2-months after a |
|---|
| **GO ID** |
| 3924 |
| 5525 |
| 19001 |
| 32561 |
| 5515 |
| 166 |
| 32553 |
| 32555 |
| 17076 |
| 17111 |
| 16462 |
| 16818 |
| 16817 |
| 5198 |
| 5488 |
| 5516 |
| 3735 |

| 3-months after a |
|---|
| **GO ID** |

166
32553
32555
17111
17076
5515
16462
16818
16817
3924
5525
19001
32561
15077
42625
22890
5488
5198
19829
15078
3824
42626
43492
16820
51287
32559
30554
15662
15399
15405
5524
46961
16787
50662
16887
22804
5391
48037
16491
42623
8092
5215
8556
149
22892

8324
15075
5516
22891
46933
19905
15079
19904

**lpha-synuclein PFF injection**

### GO description

protein binding
nucleotide binding
ribonucleotide binding
purine ribonucleotide binding
purine nucleotide binding
GTPase activity
GTP binding
guanyl nucleotide binding
guanyl ribonucleotide binding
binding
cofactor binding

**alpha-synuclein PFF injection**

### GO description

GTPase activity
GTP binding
guanyl nucleotide binding
guanyl ribonucleotide binding
protein binding
nucleotide binding
ribonucleotide binding
purine ribonucleotide binding
purine nucleotide binding
nucleoside-triphosphatase activity
pyrophosphatase activity
hydrolase activity, acting on acid anhydrides, in phosphorus-containing anhydrides
hydrolase activity, acting on acid anhydrides
structural molecule activity
binding
calmodulin binding
structural constituent of ribosome

**alpha-synuclein PFF injection**

### GO description

nucleotide binding
ribonucleotide binding
purine ribonucleotide binding
nucleoside-triphosphatase activity
purine nucleotide binding
protein binding
pyrophosphatase activity
hydrolase activity, acting on acid anhydrides, in phosphorus-containing anhydrides
hydrolase activity, acting on acid anhydrides
GTPase activity
GTP binding
guanyl nucleotide binding
guanyl ribonucleotide binding
monovalent inorganic cation transmembrane transporter activity
ATPase activity, coupled to transmembrane movement of ions
inorganic cation transmembrane transporter activity
binding
structural molecule activity
cation-transporting ATPase activity
hydrogen ion transmembrane transporter activity
catalytic activity
ATPase activity, coupled to transmembrane movement of substances
ATPase activity, coupled to movement of substances
hydrolase activity, acting on acid anhydrides, catalyzing transmembrane movement of subs
NAD or NADH binding
adenyl ribonucleotide binding
adenyl nucleotide binding
ATPase activity, coupled to transmembrane movement of ions, phosphorylative mechanism
primary active transmembrane transporter activity
P-P-bond-hydrolysis-driven transmembrane transporter activity
ATP binding
proton-transporting ATPase activity, rotational mechanism
hydrolase activity
coenzyme binding
ATPase activity
active transmembrane transporter activity
sodium:potassium-exchanging ATPase activity
cofactor binding
oxidoreductase activity
ATPase activity, coupled
cytoskeletal protein binding
transporter activity
potassium-transporting ATPase activity
SNARE binding
substrate-specific transporter activity

cation transmembrane transporter activity
ion transmembrane transporter activity
calmodulin binding
substrate-specific transmembrane transporter activity
hydrogen ion transporting ATP synthase activity, rotational mechanism
syntaxin binding
potassium ion transmembrane transporter activity
protein domain specific binding

| p-val | Corrected p-val | Cluster frequency |
|---|---|---|
| 6.96E-12 | 3.31E-09 | 133/171 77.7% |
| 4.77E-10 | 1.13E-07 | 57/171 33.3% |
| 2.56E-09 | 3.05E-07 | 49/171 28.6% |
| 2.56E-09 | 3.05E-07 | 49/171 28.6% |
| 3.73E-09 | 3.54E-07 | 50/171 29.2% |
| 8.47E-08 | 6.70E-06 | 14/171 8.1% |
| 1.75E-07 | 1.19E-05 | 18/171 10.5% |
| 2.79E-07 | 1.47E-05 | 18/171 10.5% |
| 2.79E-07 | 1.47E-05 | 18/171 10.5% |
| 5.83E-07 | 2.77E-05 | 160/171 93.5% |
| 8.72E-07 | 3.77E-05 | 14/171 8.1% |

| p-val | Corrected p-val | Cluster frequency |
|---|---|---|
| 1.07E-16 | 4.95E-14 | 23/172 13.3% |
| 8.29E-15 | 1.92E-12 | 27/172 15.6% |
| 1.80E-14 | 2.09E-12 | 27/172 15.6% |
| 1.80E-14 | 2.09E-12 | 27/172 15.6% |
| 3.23E-14 | 2.99E-12 | 138/172 80.2% |
| 4.16E-14 | 3.21E-12 | 65/172 37.7% |
| 2.47E-12 | 1.44E-10 | 55/172 31.9% |
| 2.47E-12 | 1.44E-10 | 55/172 31.9% |
| 4.06E-12 | 2.09E-10 | 56/172 32.5% |
| 3.42E-11 | 1.59E-09 | 31/172 18.0% |
| 8.88E-11 | 3.74E-09 | 31/172 18.0% |
| 9.84E-11 | 3.80E-09 | 31/172 18.0% |
| 1.13E-10 | 4.03E-09 | 31/172 18.0% |
| 3.29E-09 | 1.09E-07 | 26/172 15.1% |
| 1.32E-07 | 4.07E-06 | 162/172 94.1% |
| 5.05E-07 | 1.46E-05 | 11/172 6.3% |
| 1.47E-06 | 4.02E-05 | 11/172 6.3% |

| p-val | Corrected p-val | Cluster frequency |
|---|---|---|

| | | |
|---|---|---|
| 1.94E-23 | 1.21E-20 | 113/300 37.6% |
| 4.99E-20 | 8.01E-18 | 95/300 31.6% |
| 4.99E-20 | 8.01E-18 | 95/300 31.6% |
| 5.14E-20 | 8.01E-18 | 56/300 18.6% |
| 8.55E-20 | 1.07E-17 | 97/300 32.3% |
| 1.22E-19 | 1.27E-17 | 233/300 77.6% |
| 3.05E-19 | 2.72E-17 | 56/300 18.6% |
| 3.69E-19 | 2.88E-17 | 56/300 18.6% |
| 4.77E-19 | 3.30E-17 | 56/300 18.6% |
| 6.83E-17 | 4.26E-15 | 29/300 9.6% |
| 9.60E-15 | 5.44E-13 | 35/300 11.6% |
| 2.50E-14 | 1.20E-12 | 35/300 11.6% |
| 2.50E-14 | 1.20E-12 | 35/300 11.6% |
| 1.78E-12 | 7.94E-11 | 22/300 7.3% |
| 2.13E-12 | 8.85E-11 | 15/300 5.0% |
| 3.67E-12 | 1.43E-10 | 24/300 8.0% |
| 5.31E-12 | 1.95E-10 | 282/300 94.0% |
| 1.11E-11 | 3.85E-10 | 40/300 13.3% |
| 1.43E-10 | 4.68E-09 | 10/300 3.3% |
| 2.22E-10 | 6.92E-09 | 15/300 5.0% |
| 2.74E-10 | 8.13E-09 | 151/300 50.3% |
| 1.01E-09 | 2.87E-08 | 15/300 5.0% |
| 1.16E-09 | 3.16E-08 | 15/300 5.0% |
| 1.34E-09 | 3.48E-08 | 15/300 5.0% |
| 1.50E-09 | 3.74E-08 | 11/300 3.6% |
| 2.38E-09 | 5.71E-08 | 63/300 21.0% |
| 2.88E-09 | 6.43E-08 | 65/300 21.6% |
| 3.00E-09 | 6.46E-08 | 11/300 3.6% |
| 4.40E-09 | 8.58E-08 | 15/300 5.0% |
| 4.40E-09 | 8.58E-08 | 15/300 5.0% |
| 8.58E-09 | 1.62E-07 | 61/300 20.3% |
| 4.06E-08 | 7.45E-07 | 7/300 2.3% |
| 4.65E-08 | 8.29E-07 | 79/300 26.3% |
| 8.46E-08 | 1.47E-06 | 17/300 5.6% |
| 1.31E-07 | 2.21E-06 | 22/300 7.3% |
| 2.27E-07 | 3.73E-06 | 23/300 7.6% |
| 3.19E-07 | 5.10E-06 | 5/300 1.6% |
| 5.15E-07 | 7.99E-06 | 19/300 6.3% |
| 5.25E-07 | 7.99E-06 | 34/300 11.3% |
| 5.81E-07 | 8.63E-06 | 19/300 6.3% |
| 7.13E-07 | 1.03E-05 | 28/300 9.3% |
| 1.12E-06 | 1.57E-05 | 47/300 15.6% |
| 1.13E-06 | 1.57E-05 | 5/300 1.6% |
| 1.36E-06 | 1.85E-05 | 7/300 2.3% |
| 1.44E-06 | 1.91E-05 | 41/300 13.6% |

| | | |
|---|---|---|
| 2.37E-06 | 3.08E-05 | 28/300 9.3% |
| 3.18E-06 | 4.04E-05 | 33/300 11.0% |
| 3.48E-06 | 4.34E-05 | 13/300 4.3% |
| 3.74E-06 | 4.57E-05 | 36/300 12.0% |
| 4.67E-06 | 5.61E-05 | 5/300 1.6% |
| 5.17E-06 | 6.08E-05 | 6/300 2.0% |
| 6.90E-06 | 7.97E-05 | 5/300 1.6% |
| 7.63E-06 | 8.66E-05 | 22/300 7.3% |

**Total frequency**

8120/15432 52.6%
2251/15432 14.5%
1842/15432 11.9%
1842/15432 11.9%
1925/15432 12.4%
209/15432 1.3%
373/15432 2.4%
385/15432 2.4%
385/15432 2.4%
12359/15432 80.0%
253/15432 1.6%

**Total frequency**

208/15428 1.3%
372/15428 2.4%
384/15428 2.4%
384/15428 2.4%
8117/15428 52.6%
2251/15428 14.5%
1843/15428 11.9%
1843/15428 11.9%
1925/15428 12.4%
697/15428 4.5%
724/15428 4.6%
727/15428 4.7%
731/15428 4.7%
605/15428 3.9%
12357/15428 80.0%
141/15428 0.9%
157/15428 1.0%

**Total frequency**

2249/15420 14.5%
1841/15420 11.9%
1841/15420 11.9%
695/15420 4.5%
1923/15420 12.4%
8116/15420 52.6%
722/15420 4.6%
725/15420 4.7%
729/15420 4.7%
208/15420 1.3%
372/15420 2.4%
384/15420 2.4%
384/15420 2.4%
169/15420 1.0%
67/15420 0.4%
212/15420 1.3%
12351/15420 80.0%
605/15420 3.9%
30/15420 0.1%
91/15420 0.5%
5086/15420 32.9%
101/15420 0.6%
102/15420 0.6%
103/15420 0.6%
48/15420 0.3%
1493/15420 9.6%
1573/15420 10.2%
51/15420 0.3%
112/15420 0.7%
112/15420 0.7%
1470/15420 9.5%
19/15420 0.1%
2236/15420 14.5%
180/15420 1.1%
304/15420 1.9%
340/15420 2.2%
9/15420 0.0%
252/15420 1.6%
685/15420 4.4%
254/15420 1.6%
507/15420 3.2%
1158/15420 7.5%
11/15420 0.0%
30/15420 0.1%
954/15420 6.1%

539/15420 3.4%
709/15420 4.5%
140/15420 0.9%
816/15420 5.2%
14/15420 0.0%
24/15420 0.1%
15/15420 0.0%
388/15420 2.5%

### Genes

RPL4 GMFB RAB3A PEBP1 ABAT ENO1 SLC8A1 SLC8A2 SCAMP5 RPS19 PPP2R1A CFL1 STMN1 ATP6V1E1 TMSB
ARF1 RAB3A HSP90AB1 PEBP1 ETFA PYGM TUBB3 IDH3B PHGDH RAC1 NSF PRKCG RAB2A PRKCB SARS SYN2
ARF1 RAB3A HSP90AB1 PEBP1 PYGM TUBB3 RAC1 NSF PRKCG RAB2A PRKCB SARS SYN2 TUBA4A TUFM EEF1
ARF1 RAB3A HSP90AB1 PEBP1 PYGM TUBB3 RAC1 NSF PRKCG RAB2A PRKCB SARS SYN2 TUBA4A TUFM EEF1
ARF1 RAB3A HSP90AB1 PEBP1 ETFA PYGM TUBB3 RAC1 NSF PRKCG RAB2A PRKCB SARS SYN2 TUBA4A TUFM
RAB2A NAPB ARF1 RAB3A TUBA4A GNAI1 TUFM EEF1A1 TUBB2B TUBB2A TUBB3 GNAQ RAC1 RAN
RAB2A NAPB SEPT10 ARF1 RAB3A TUBA4A GNAI1 TUFM EEF1A1 RAB10 GLUD1 TUBB2B TUBB2A TUBB3 GNA
RAB2A NAPB SEPT10 ARF1 RAB3A TUBA4A GNAI1 TUFM EEF1A1 RAB10 GLUD1 TUBB2B TUBB2A TUBB3 GNA
RAB2A NAPB SEPT10 ARF1 RAB3A TUBA4A GNAI1 TUFM EEF1A1 RAB10 GLUD1 TUBB2B TUBB2A TUBB3 GNA
RPL4 GMFB RAB3A PEBP1 ABAT ENO1 RPL8 SLC8A1 SLC8A2 SCAMP5 RPS19 PPP2R1A TUBB3 CFL1 STMN1 ATP
GOT1 IDH3G GOT2 ABAT ETFA PYGM SDHB ACAT2 ACAT1 GLUD1 IDH3B PHGDH DLAT NDUFV1

### Genes

RAB2A RAB1A ARF1 RAB3A TUBB SEPT5 TUBA4A DNM1 TUFM GNAI2 EEF1A1 GNAO1 TUBB6 EHD3 TUBB2A
RAB1A ARF1 RAB3A GNAI2 TUBB6 OPA1 TUBB3 GNA11 DNM1L RAB6B RAB2A SEPT10 TUBB SEPT5 TUBA4A
RAB1A ARF1 RAB3A GNAI2 TUBB6 OPA1 TUBB3 GNA11 DNM1L RAB6B RAB2A SEPT10 TUBB SEPT5 TUBA4A
RAB1A ARF1 RAB3A GNAI2 TUBB6 OPA1 TUBB3 GNA11 DNM1L RAB6B RAB2A SEPT10 TUBB SEPT5 TUBA4A
GMFB RAB3A PEBP1 ENO1 PARK7 GLS RPL7 SLC8A2 RPH3A RPS15 SCAMP5 SCP2 PPP2R1A CFL1 RPL38 TMSB1
ARF1 RAB3A HSP90AB1 PEBP1 PYGM TUBB6 ACTR1A OPA1 TUBB3 IDH3B DNM1L NSF PRKCG RAB2A CSNK2A
ARF1 RAB3A HSP90AB1 PEBP1 PYGM TUBB6 ACTR1A OPA1 TUBB3 DNM1L NSF PRKCG RAB2A CSNK2A1 PRKC
ARF1 RAB3A HSP90AB1 PEBP1 PYGM TUBB6 ACTR1A OPA1 TUBB3 DNM1L NSF PRKCG RAB2A CSNK2A1 PRKC
ARF1 RAB3A HSP90AB1 PEBP1 PYGM TUBB6 ACTR1A OPA1 TUBB3 DNM1L NSF PRKCG RAB2A CSNK2A1 PRKC
RAB1A ARF1 RAB3A GNAI2 TUBB6 OPA1 TUBB3 GNA11 CCT8 DNM1L RAB6B NSF RAB2A DYNC1H1 TUBB MY
RAB1A ARF1 RAB3A GNAI2 TUBB6 OPA1 TUBB3 GNA11 CCT8 DNM1L RAB6B NSF RAB2A DYNC1H1 TUBB MY
RAB1A ARF1 RAB3A GNAI2 TUBB6 OPA1 TUBB3 GNA11 CCT8 DNM1L RAB6B NSF RAB2A DYNC1H1 TUBB MY
RAB1A ARF1 RAB3A GNAI2 TUBB6 OPA1 TUBB3 GNA11 CCT8 DNM1L RAB6B NSF RAB2A DYNC1H1 TUBB MY
CLTC RPL11 CLTA RPL7 RPS15 TUBB6 TUBB3 MAP2 NEFL RPL13 RPL38 RPL15 RPS13 SPTBN2 RPS7 RPS8 TUBB
GMFB RAB3A CPNE6 PEBP1 ENO1 PARK7 GLS RPL7 SLC8A2 RPH3A RPS15 SCAMP5 TUBB6 SCP2 PPP2R1A TUB
CAMK2B PPP3CA PPP3R1 CAMK2D SYT1 MAP2 CAMK2A MYO5A ATP2B1 ADD1 SLC8A2
RPS15 RPS7 RPS8 RPL11 RPL13A RPL13 RPL38 RPL15 UBA52 RPS13 RPL7

### Genes

RAB3A PEBP1 TUBB6 KIF5C TUBB3 CKMT1B GLUL PRKCG NSF RAB2A MAP2K1 CSNK2A1 PRKCB TUBA4A TUF
ARF1 RAB3A HSP90AB1 PEBP1 ACTR1A TUBB6 SEPT6 TUBA1A OPA1 KIF5C TUBB3 CKMT1B RAC1 DNM1L GLU
ARF1 RAB3A HSP90AB1 PEBP1 ACTR1A TUBB6 SEPT6 TUBA1A OPA1 KIF5C TUBB3 CKMT1B RAC1 DNM1L GLU
ARF1 RAB3A TUBB6 TUBA1A OPA1 KIF5C TUBB3 RAC1 ATP6V1E1 DNM1L NSF RAB2A TUBB RHOG ATP1B2 DY
RAB3A PEBP1 TUBB6 KIF5C TUBB3 CKMT1B GLUL PRKCG NSF RAB2A MAP2K1 CSNK2A1 PRKCB TUBA4A TUF
RPL4 GMFB ENO1 ENO2 RPL7 SCAMP5 SCP2 PPP2R1A DPYSL2 RPL38 ATP6V1E1 GLUL RPS13 CSNK2A1 PRKCB
ARF1 RAB3A TUBB6 TUBA1A OPA1 KIF5C TUBB3 RAC1 ATP6V1E1 DNM1L NSF RAB2A TUBB RHOG ATP1B2 DY
ARF1 RAB3A TUBB6 TUBA1A OPA1 KIF5C TUBB3 RAC1 ATP6V1E1 DNM1L NSF RAB2A TUBB RHOG ATP1B2 DY
ARF1 RAB3A TUBB6 TUBA1A OPA1 KIF5C TUBB3 RAC1 ATP6V1E1 DNM1L NSF RAB2A TUBB RHOG ATP1B2 DY
NAPB RAB1A ARF1 RAB3A GNAI1 GNAI2 TUBB6 TUBA1A OPA1 TUBB3 GNA11 RAC1 DNM1L RAB6B RAB2A T
NAPB RAB1A ARF1 RAB3A GNAI1 GNAI2 TUBB6 SEPT6 TUBA1A OPA1 TUBB3 GNA11 RAC1 DNM1L RAB6B RA
NAPB RAB1A ARF1 RAB3A GNAI1 GNAI2 TUBB6 SEPT6 TUBA1A OPA1 TUBB3 GNA11 RAC1 DNM1L RAB6B RA
NAPB RAB1A ARF1 RAB3A GNAI1 GNAI2 TUBB6 SEPT6 TUBA1A OPA1 TUBB3 GNA11 RAC1 DNM1L RAB6B RA
UQCRB ATP5A1 SLC1A2 ATP1A3 ATP5I ATP1A2 ATP1B2 ATP1A1 ATP5O ATP1B1 COX5A SLC8A1 ATP5B ATP5D U
ATP5A1 ATP1A3 ATP1A2 ATP2B2 ATP1B2 ATP1A1 ATP2B1 ATP1B1 ATP5B ATP5D ATP6V1B2 ATP6V1H ATP6V1
UQCRB ATP5A1 SLC1A2 ATP1A3 ATP5I ATP1A2 ATP2B2 ATP1B2 ATP1A1 ATP2B1 ATP5O ATP1B1 COX5A SLC8A
RPL4 AHCYL1 GMFB ENO1 ENO2 RPL7 SCAMP5 SCP2 PPP2R1A DPYSL2 CKMT1B RPL38 ATP6V1E1 BSN GLUL RP
RPL4 CLTC RPL11 CLTB CLTA RPL7 TUBB6 TUBA1A EPB41L1 MAP2 TUBB3 EPB41L3 NEFL RPL13 RPL38 NEFM
ATP5B ATP5D ATP5A1 ATP6V1B2 ATP6V1H ATP1A2 ATP2B2 ATP2B1 ATP6V1E1 ATP6V1C1
UQCRB ATP5A1 ATP5I ATP5O COX5A ATP5B ATP5D UQCRC1 ATP6V1B2 ATP6V1H UQCRFS1 ATP6V1E1 ATP6V0
AHCYL1 RAB3A OGDHL ENO1 PARK7 ENO2 TUBB6 SCP2 KIF5C TUBB3 DPYSL2 DPYSL3 CKMT1B ATP6V1E1 GLU
ATP5A1 ATP1A3 ATP1A2 ATP2B2 ATP1B2 ATP1A1 ATP2B1 ATP1B1 ATP5B ATP5D ATP6V1B2 ATP6V1H ATP6V1
ATP5A1 ATP1A3 ATP1A2 ATP2B2 ATP1B2 ATP1A1 ATP2B1 ATP1B1 ATP5B ATP5D ATP6V1B2 ATP6V1H ATP6V1
ATP5A1 ATP1A3 ATP1A2 ATP2B2 ATP1B2 ATP1A1 ATP2B1 ATP1B1 ATP5B ATP5D ATP6V1B2 ATP6V1H ATP6V1
LDHB GLUD1 MDH1 IDH3G IDH2 IDH3B NDUFS2 PHGDH DLD NDUFV1 GAPDH
HSP90AB1 PEBP1 ACTR1A KIF5C CKMT1B GLUL PRKCG NSF ACTR2 MAP2K1 HSP90AA1 CSNK2A1 PRKCB NME2
HSP90AB1 PEBP1 ETFA ACTR1A KIF5C CKMT1B GLUL PRKCG NSF ACTR2 MAP2K1 HSP90AA1 CSNK2A1 PRKCB
ATP5B ATP1A3 ATP1A2 ATP2B2 ATP1B2 ATP1A1 ATP2B1 ATP6V1E1 ATP6V0D1 ATP1B1 ATP6V1C1
ATP5A1 ATP1A3 ATP1A2 ATP2B2 ATP1B2 ATP1A1 ATP2B1 ATP1B1 ATP5B ATP5D ATP6V1B2 ATP6V1H ATP6V1
ATP5A1 ATP1A3 ATP1A2 ATP2B2 ATP1B2 ATP1A1 ATP2B1 ATP1B1 ATP5B ATP5D ATP6V1B2 ATP6V1H ATP6V1
HSP90AB1 PEBP1 ACTR1A KIF5C CKMT1B GLUL PRKCG NSF ACTR2 MAP2K1 HSP90AA1 CSNK2A1 PRKCB NME2
ATP5B ATP5D ATP5A1 ATP6V1B2 ATP6V1H ATP6V1E1 ATP6V1C1
GPI ARF1 AHCYL1 RAB3A ENO1 PPP3CA UCHL1 TUBB6 TUBA1A SCP2 OPA1 KIF5C TUBB3 DPYSL2 DPYSL3 RAC1
ACOT7 MDH1 IDH3G IDH2 OGDHL ETFA ACAT2 LDHB GLUD1 OGDH IDH3B NDUFS2 PHGDH DLAT DLD NDUFV1
NSF DYNC1H1 HSPA8 VCP ATP5A1 ATP1A3 ATP1A2 ATP2B2 ATP1B2 ATP1A1 ATP2B1 ATP1B1 HSPD1 ATP5B AT
SLC25A3 SLC12A5 ATP5A1 SLC1A2 ATP1A3 ATP1A2 ATP2B2 ATP1B2 ATP1A1 ATP2B1 ATP1B1 SLC8A1 SLC8A2 A
ATP1A3 ATP1A2 ATP1B2 ATP1A1 ATP1B1
ACOT7 MDH1 GOT1 IDH3G GOT2 IDH2 OGDHL ETFA ACAT2 LDHB GLUD1 OGDH IDH3B NDUFS2 PHGDH DLAT D
UQCRB AKR1B1 OGDHL ETFA PARK7 PDHB COX5A LDHB LDHA PRDX5 SCP2 IDH3B UQCRFS1 PHGDH NDUFV1 P
NSF DYNC1H1 HSPA8 ATP5A1 ATP1A3 ATP1A2 ATP2B2 ATP1B2 ATP1A1 ATP2B1 ATP1B1 ATP5B ATP5D ATP6V1
SNAP25 GMFB WDR1 ADD1 NCALD RPH3A EPB41L1 TMSB4X EPB41L3 CFL1 TMSB10 YWHAH CAP1 ACTR2 MYC
SNAP25 SLC25A3 UQCRB ATP5A1 SLC1A2 ATP1A3 ATP5I AP2A1 ATP1A2 ATP1A1 ATP5O AP2A2 COX5A SLC8A1
ATP1A3 ATP1A2 ATP1B2 ATP1A1 ATP1B1
NSF SNAP25 SYT1 STXBP1 CPLX2 CPLX1 STX1A
SNAP25 SLC25A3 UQCRB ATP5A1 SLC1A2 ATP1A3 ATP5I AP2A1 ATP1A2 ATP1A1 ATP5O AP2A2 COX5A SLC8A1

SNAP25 UQCRB ATP5A1 SLC1A2 ATP1A3 ATP5I ATP1A2 ATP1A1 ATP5O COX5A SLC8A1 SLC8A2 ATP5B ATP5D A
SNAP25 SLC25A3 UQCRB ATP5A1 SLC1A2 ATP1A3 ATP5I ATP1A2 ATP1A1 ATP5O COX5A SLC8A1 SLC8A2 ATP5B
CAMK2B CAMK2D SYT1 CAMK2A MYO5A ATP2B2 ATP2B1 SLC8A1 ADD1 SLC8A2 PPP3CA PPP3R1 MAP2
SNAP25 SLC25A3 UQCRB ATP5A1 SLC1A2 ATP1A3 ATP5I ATP1A2 ATP1A1 ATP5O COX5A SLC8A1 SLC8A2 ATP5B
ATP5B ATP5D ATP5A1 ATP6V1B2 ATP5O
NSF SNAP25 SYT1 STXBP1 CPLX2 CPLX1
ATP1A3 ATP1A2 ATP1B2 ATP1A1 ATP1B1
YWHAE VCP HSP90AA1 HSP90AB1 YWHAB ATP2B2 DYNLL1 HPCAL4 YWHAZ STX1B TUBA1A SYNJ1 OPA1 YWHA

X1 ACAT2 ACAT1 HSP90B1 LDHA PCSK1N PPP3R1 PRKAR2B PRKAR2A PCBP2 ATP6V1C1 PACS
10 HBG1 HSPA4 IDH3G MYO5A ATP2B2 HSPA12A DCLK1 RAB10 GLUD1 SUCLA2 GNAQ DNAJ
A DCLK1 RAB10 GLUD1 SUCLA2 GNAQ DNAJA2 RAN
A DCLK1 RAB10 GLUD1 SUCLA2 GNAQ DNAJA2 RAN
SPA12A DCLK1 RAB10 GLUD1 SUCLA2 GNAQ DNAJA2 RAN

VDAC1 PPID STX1A SET CNP GLO1 HSPA4L CPLX1 ACAT2 ACAT1 HSP90B1 LDHA PCSK1N PPP

P2 RAB6B SPTBN2 HSPA9 PDHA1 SYT1 FUS HIST1H4L GOT2 MYO5A SNAP91 GNAO1 HNRNP
CCT5 HSPA9 SEPT10 DYNC1H1 HNRNPA3 HSPA4 IDH2 MYO5A ATP2B1 HSPA12A DCLK1 GNA
MYO5A ATP2B1 HSPA12A DCLK1 GNAO1 RAB10 GLUD1 EHD3 SUCLA2 GNAQ AAK1 UBA1 RA
MYO5A ATP2B1 HSPA12A DCLK1 GNAO1 RAB10 GLUD1 EHD3 SUCLA2 GNAQ AAK1 UBA1 RA
PA4 MYO5A ATP2B1 HSPA12A DCLK1 GNAO1 RAB10 GLUD1 EHD3 SUCLA2 GNAQ AAK1 UBA

T CNP GLO1 CPLX1 ACAT1 STIP1 LDHA PPP3R1 MAP2 PRKAR2B PCBP2 PGK1 NDUFV1 RAB6B

LUD1 EHD3 GNAQ HNRNPD AAK1 UBE2N UBA1 ARF1 HSP90AB1 ETFA ACTR1A SEPT6 TUBA
2 LANCL2 TKT ARF5 CAMK2B NAPB RAB1A VCP CAMK2D CNP HSPA4L ATP5A1 CAMK2A ATP1
2 LANCL2 TKT ARF5 CAMK2B NAPB RAB1A VCP CAMK2D CNP HSPA4L ATP5A1 CAMK2A ATP1
D ATP6V1H CCT8 RAB6B ATP6V1C1 DYNC1H1 HSPA8 MYO5A ATP2B2 ATP2B1 EEF2 GNAO1 E
HSP90AB1 ETFA ACTR1A SEPT6 TUBA1A OPA1 RAC1 DNM1L ACTR2 HSP90AA1 TUBB NME2
2 PRKAR2B PRKAR2A ATP5D EPB41L3 PCBP2 ATP6V1C1 ATP6V0A1 ST13 MYO5A SNAP91 CS
D ATP6V1H CCT8 RAB6B ATP6V1C1 DYNC1H1 HSPA8 MYO5A ATP2B2 ATP2B1 EEF2 GNAO1 E
D ATP6V1H CCT8 RAB6B ATP6V1C1 DYNC1H1 HSPA8 MYO5A ATP2B2 ATP2B1 EEF2 GNAO1 E
D ATP6V1H CCT8 RAB6B ATP6V1C1 DYNC1H1 HSPA8 MYO5A ATP2B2 ATP2B1 EEF2 GNAO1 E

O ATP5B LDHB LDHA PPP3R1 EPB41L1 MAP2 PRKAR2B PRKAR2A ATP5D EPB41L3 PCBP2 ATP

CAT2 ATP5B LDHB LDHA PPP3R1 PRDX5 MAP2 ATP5D PGK1 ATP6V1H NDUFV1 RAB6B ATP6V

HSPA9 DYNC1H1 HSPA8 HBG1 HSPA4 IDH3G MYO5A ATP2B2 ATP2B1 HSPA12A DCLK1 GLUD
CKB CCT5 HSPA9 DYNC1H1 HSPA8 HBG1 HSPA4 IDH3G MYO5A ATP2B2 ATP2B1 HSPA12A D

SPA8 HBG1 HSPA4 IDH3G MYO5A ATP2B2 ATP2B1 HSPA12A DCLK1 GLUD1 EHD3 SUCLA2 PS

6V0D1 ARF5 PAFAH1B2 NAPB RAB1A VCP CNP ATP5A1 ATP1A3 ATP1A2 ATP1A1 GNAI1 GNA

3 SFXN3 ATP6V1B2 VDAC2 VDAC1 ATP6V0D1 SLC25A12 SLC25A5 SLC25A4

D1 SLC25A12 SLC25A5 SLC25A4

IN1 SPTBN2 PDHA1 SYT1 HBG1 GOT2 MYO5A ARPC4 GLUD1 SYNJ1 GNAQ HNRNPD CYCS CAL
\2 HNRNPA2B1 HNRNPD RAN

R1 PRKAR2B PRKAR2A PCBP2 PGK1 NDUFV1 ATP6V1C1 PACSIN1 SPTBN2 PDHA1 GOT1 SYT1

L GLUD1 EHD3 SYNJ1 GNAQ HNRNPD UBA1 YWHAE GPI SNAP25 ARF1 HSP90AB1 YWHAB CL
)1 HNRNPL RAB10 GLUD1 EHD3 SUCLA2 GNAQ HNRNPA2B1 AAK1 HNRNPD UBA1 RAN
N
N
L RAN

SPTBN2 HSPA9 PDHA1 GOT1 SYT1 FUS HIST1H4L HSPA4 GOT2 IDH2 MYO5A HSPA12A DCLK1

1A OPA1 IDH3B PHGDH RAC1 DNM1L ACTR2 HSP90AA1 TUBB NME2 RHOG SEPT1 SYN2 SEPT
A3 ATP1A2 ATP1A1 GNAI1 GNAI2 HSPD1 ATP5B PRKAR2B PRKAR2A GNA11 PGK1 CCT8 CKB R
A3 ATP1A2 ATP1A1 GNAI1 GNAI2 HSPD1 ATP5B PRKAR2B PRKAR2A GNA11 PGK1 CCT8 CKB R
HD3 GNAQ PSMC2 RAN
RHOG SEPT1 SYN2 SEPT5 SYN1 SEPT2 TUBB2B TUBB2A PKM TCP1 ATP6V1B2 LANCL2 TKT D
INRNPL GLUD1 EHD3 HNRNPK HNRNPD GPI ARF1 GDI1 CLTC CLTB AKR1B1 CLTA AP2A1 AP2A
HD3 GNAQ PSMC2 RAN
HD3 GNAQ PSMC2 RAN
HD3 GNAQ PSMC2 RAN

6V1C1 ATP6V0A1 ST13 MYO5A HSPA12A DCLK1 SNAP91 CS HNRNPL GLUD1 EHD3 HNRNPK H

1C1 HSPA8 PDHA1 GOT1 HBG1 GOT2 IDH2 MYO5A EEF2 PRDX6 DCLK1 CS GNAO1 QDPR GLUD

1 EHD3 SUCLA2 PSMC2 DNAJA2 AAK1 UBE2N UBA1 PFKM PFKP
CLK1 GLUD1 EHD3 SUCLA2 PSMC2 DNAJA2 AAK1 UBE2N UBA1 PFKM PFKP

MC2 DNAJA2 AAK1 UBE2N UBA1 PFKM PFKP

2 HSPD1 ATP5B PPP3R1 PTPRZ1 MAP2 GNA11 ATP5D ATP6V1H CCT8 RAB6B ATP6V1C1 DYNC

.R CALM2 YWHAE GPI ARF1 HSP90AB1 GDI1 YWHAB CLTC CLTB AKR1B1 AP2A1 PYGM PPP3C/

HBG1 HSPA4 GOT2 MYO5A ARPC4 HSPA12A DCLK1 GLUD1 SYNJ1 GNAQ HNRNPD CYCS CALF

TC AKR1B1 CLTA AP2A1 PYGM PPP3CA ACTR1A UCHL1 OPA1 YWHAQ CA2 TMSB4X ENSA EFH

SNAP91 GNAO1 HNRNPL GLUD1 EHD3 SYNJ1 GNAQ AAK1 HNRNPD UBA1 YWHAE GPI SNAP:

5 SYN1 SEPT2 TUBB2B TUBB2A PKM HNRNPH1 TCP1 ATP6V1B2 LANCL2 NDUFS2 TKT DLD GA
AB6B CCT5 HSPA9 DYNC1H1 HSPA8 SEPT10 HBG1 HSPA4 IDH3G MYO5A ATP2B2 ATP2B1 HSP
AB6B CCT5 HSPA9 DYNC1H1 HSPA8 SEPT10 HBG1 HSPA4 IDH3G MYO5A ATP2B2 ATP2B1 HSP

LD ARF5 CAMK2B NAPB RAB1A CAMK2D ATP5A1 CAMK2A ATP1A3 ATP1A2 ATP1A1 GNAI1 G
2 PPP3CA ACTR1A UCHL1 TUBA1A TMSB4X ENSA NEFL NEFM DLAT RAC1 NEFH GIT1 CAP1 P

NRNPD OGDH GPI ARF1 GDI1 CLTC CLTB AKR1B1 CLTA AP2A1 AP2A2 PPP3CA ACTR1A UCHL1

O1 EHD3 SYNJ1 GNAQ OGDH AAK1 UBE2N CPE UBA1 ACO2 GPI ARF1 AKR1B1 ETFA PPP3CA U

C1H1 HSPA8 MYO5A ATP2B2 ATP2B1 EEF2 PRDX6 GNAO1 EHD3 SYNJ1 GNAQ PSMC2 CPE RA

UCHL1 IMPA1 CA2 TRIM2 HIST1H1D NEFL DLAT RAC1 NEFH YWHAH EIF5A DYNLL1 AP3B2

CALM2 HIST2H2AA3 YWHAE GPI ARF1 HSP90AB1 GDI1 YWHAB CLTC CLTB AKR1B1 AP2A1

D2 HIST1H1D NEFL UQCRFS1 DNM1L GIT1 EIF5A TUBB DYNLL1 SEPT5 AP3B2 DYNLL2

25 ARF1 HSP90AB1 YWHAB CLTC AKR1B1 CLTA AP2A1 PYGM PPP3CA ACTR1A UCHL1 OPA1

APDH ARF5 CAMK2B NAPB RAB1A CAMK2D ATP5A1 CAMK2A SRSF1 ATP1A3 ATP1A2 ATP1A1
PA12A EEF2 DCLK1 GNAO1 GLUD1 EHD3 SUCLA2 GNAQ PSMC2 DNAJA2 AAK1 UBE2N UBA1 P
PA12A EEF2 DCLK1 GNAO1 GLUD1 EHD3 SUCLA2 GNAQ PSMC2 DNAJA2 AAK1 UBE2N UBA1 P

NAI2 HSPD1 GNA11 CCT8 CKB CCT5 DYNC1H1 SEPT10 IDH3G ATP2B2 ATP2B1 SUCLA2 PSMC2
GAM1 TUBB NME2 RHOG DYNLL1 AP3B2 DYNLL2 SYN1 TUBB2A CTTN HNRNPH1 TCP1 ALDOC

TUBA1A TMSB4X ENSA NEFL NEFM PHGDH DLAT RAC1 NEFH GIT1 CAP1 PGAM1 TUBB NME

UCHL1 TUBA1A OPA1 IDH3B UQCRFS1 PHGDH DLAT RAC1 DNM1L PGM1 CAP1 PGAM1 TUBB

N

YNLL2 YWHAZ ACTA1 PKM TUBB2A CTTN HNRNPH1 MAPRE3 CD47 ALDOA SLC25A5 SLC25A4

TFA PYGM PPP3CA UCHL1 IMPA1 CA2 TRIM2 HIST1H1D IDH3B NEFL PHGDH DLAT RAC1 NEFH

DNM1 ACTA1 TUBB2A CTTN HNRNPH1 SLC25A12 SLC25A5 DLD MAPRE2 SLC25A4 ARF5 CA[illegible]

WHAQ CA2 TMSB4X ENSA EFHD2 HIST1H1D IDH3B NEFL UQCRFS1 DNM1L GIT1 EIF5A TUBB

GNAI1 GNAI2 HSPD1 GNA11 EIF4H CCT8 CKB CCT5 DYNC1H1 SEPT10 MDH1 IDH3G ATP2B2 A
FKM PFKP RAN
FKM PFKP RAN

DNAJA2 PFKM PFKP RAN
C MAPRE3 CD47 MAPT TKT DLD MAPRE2 ARF5 PAFAH1B2 VAMP2 NAPB ATP5A1 RPL11 GNAI1

2 RHOG DYNLL1 AP3B2 DYNLL2 SYN2 SYN1 TUBB2B TUBB2A CTTN VSNL1 HNRNPH1 TCP1 AL

NME2 RHOG DYNLL1 DYNLL2 SYN2 SEPT5 SYN1 TUBB2B TUBB2A PKM ALDH1A1 NDUFS4 ATI

PAFAH1B2 CAMK2B NAPB CAMK2D CAMK2A ATP1A1 GNAI1 NCALD RPL13 CKB CCT5 CLASP2

PGM1 YWHAH EIF5A DYNLL1 AP3B2 DYNLL2 SYN2 YWHAZ ACTA1 SLC25A18 TUBB2B PKM TU

MK2B CAMK2D CAMK2A RPL11 SRSF1 ADD1 NCALD GNAI2 PURA GNA11 RPL13 CCT8 CKB CCT

DYNLL1 SEPT5 AP3B2 DYNLL2 YWHAZ DNM1 ACTA1 NDUFS8 TUBB2A CTTN HNRNPH1 LANCL

TP2B1 SUCLA2 HNRNPA2B1 PSMC2 DNAJA2 PFKM PFKP RAN

NAPG GNAI2 PURA RPL13 CCT8 CKB CCT5 DYNC1H1 MDH2 AP2B1 FKBP1A TPPP3 HNRNPA2[illegible]

DOC MAPRE3 CD47 MAPT TKT DLD MAPRE2 ARF5 PAFAH1B2 VAMP2 NAPB ATP5A1 RPL11 C[illegible]

6V1B2 LANCL2 NDUFS2 ALDOC NDUFS1 TKT DLD GAPDH ARF5 PAFAH1B2 CAMK2B NAPB RA[illegible]

SNCA SEPT10 TIMM8A SLC12A5 MDH2 ATP2B2 AP2B1 RAB10 NFASC TPPP3 SUCLA2 DNAJA2

UBB2A CTTN VSNL1 HNRNPH1 LANCL2 MAPRE3 CD47 ALDOA SLC25A5 SLC25A4 PAFAH1B2 CA

5 CAP2 SEPT10 DYNC1H1 TIMM8A HNRNPA3 SLC12A5 MDH2 AP2B1 ATP2B1 FKBP1A RAB10 N

.2 SLC25A12 SLC25A5 DLD MAPRE2 SLC25A4 ARF5 CAMK2B RAB1A CAMK2D CAMK2A RPL11 S

B1 PSMC2 PLP1 CTNNB1 RAB3A PEBP1 PARK7 SLC8A1 SLC8A2 RPH3A KIF5C ARHGDIA CFL1 T

OX5A GNAI1 NAPG GNAI2 PURA RPL13 CCT8 CKB RPL15 CCT5 DYNC1H1 MDH1 MDH2 AP2B1

AB1A CAMK2D ATP5A1 CAMK2A ATP1A3 ATP1A2 ATP1A1 PDHB COX5A GNAI1 GNAI2 HSPD1

HNRNPA2B1 PLP1 RAN

AMK2B NAPB CAMK2D CAMK2A ATP1A1 GNAI1 NCALD RPL13 CKB RPL15 CCT5 CLASP2 SNCA

NFASC TPPP3 SUCLA2 HNRNPA2B1 PLP1 CTNNB1 RAN

SRSF1 ADD1 NCALD GNAI2 PURA GNA11 RPL13 CCT8 CKB RPL15 CCT5 CAP2 SEPT10 DYNC1H1

MSB10 AP2M1 NSF SH3GLB2 MAP2K1 RPS7 USP5 RPS8 ATP1B2 ATP1B1 SRCIN1 TUBA4A TUI

FKBP1A TPPP3 HNRNPA2B1 PSMC2 PLP1 CTNNB1 RAB3A PEBP1 OGDHL PARK7 SLC8A1 SLC8A

PTPRZ1 GNA11 OXCT2 CCT8 CKB DYNC1H1 MDH1 NDUFA5 MDH2 IDH3G NDUFA4 ATP2B2 ATI

SEPT10 TIMM8A SLC12A5 IDH3G MDH2 CADPS ATP2B2 AP2B1 RAB10 MCM3AP NFASC TPPP3

. TIMM8A HNRNPA3 SLC12A5 MDH2 CADPS AP2B1 ATP2B1 FKBP1A RAB10 MCM3AP NFASC T

FM EEF1A1 ACLY OLFM1 FSCN1 AMPH PPIA VCP CNP PHB CPLX2 CPLX1 ACAT2 STIP1 PCSK1N

2 RPH3A TUBB6 KIF5C TUBB3 ARHGDIA CFL1 TMSB10 AP2M1 NSF RAB2A SH3GLB2 MAP2K1

2B1 FKBP1A SUCLA2 PSMC2 PFKM PFKP RAN

[illegible] SUCLA2 DNAJA2 HNRNPA2B1 PLP1 RAN

TPPP3 SUCLA2 HNRNPA2B1 PLP1 CTNNB1 RAN

ATP6V1H MBP RAB6B HSPA9 HSPA8 PDHA1 SYT1 HBG1 GOT2 MSN ARPC4 EEF2 GNB2

[illegible] RPS7 USP5 RPS8 RPSA ATP1B2 ATP1B1 SRCIN1 TUBA4A TUFM EEF1A1 ACLY OLFM1 MAG
